# Colora: A Snakemake Workflow for Complete Chromosome-scale *De Novo* Genome Assembly

**DOI:** 10.1101/2024.09.10.612003

**Authors:** Lia Obinu, Timothy Booth, Heleen De Weerd, Urmi Trivedi, Andrea Porceddu

## Abstract

**Background:** *De novo* assembly creates reference genomes that underpin many modern biodiversity and conservation studies. Large numbers of new genomes are being assembled by labs around the world. To avoid duplication of efforts and variable data quality, we desire a best-practice assembly process, implemented as an automated portable workflow.

**Results:** Here we present Colora, a Snakemake workflow that produces chromosome-scale *de novo* primary or phased genome assemblies complete with organelles using PacBio HiFi, Hi-C, and optionally ONT reads as input. The source code of Colora is available on GitHub: https://github.com/LiaOb21/colora. Colora is also available at the Snakemake Workflow Catalog (https://snakemake.github.io/snakemake-workflow-catalog/?usage=LiaOb21%2Fcolora).

**Conclusion:** Colora is a user-friendly, versatile, and reproducible pipeline that is ready to use by researchers looking for an automated way to obtain high-quality *de novo* genome assemblies.

## Background

Third-generation sequencing technologies, such as the platforms from Pacific Biosciences (PacBio) and Oxford Nanopore Technologies (ONT), can be used to obtain high-quality *de novo* genome assemblies for model and non-model species [1, 2]. However, long-read assemblies alone are unlikely to create chromosome-scale assemblies, and, therefore, Hi-C reads are widely used to perform assembly scaffolding [3].

Currently, *de novo* genome assemblies are the basis of cutting-edge biodiversity and conservation studies. However, the bioinformatics procedures used to obtain such results are often challenging in terms of time, study, and human and computational resources. This is particularly true for small labs, which do not have access to the same resources that are available to big institutions. Therefore, automating workflows by combining up-to-date techniques and tools in an easily implementable, modifiable, and portable way is fundamental.

Snakemake is a popular Python-based workflow engine that allows the automation of workflows for single machines, such as personal laptops, and High-Performance Computer (HPC) clusters [4]. It allows the creation of separate environments for each bioinformatics tool through Conda (https://anaconda.org/), Docker [5] or Singularity [6], making the installation of the whole pipeline automatic and thus easy for the user and ensuring reproducibility.

Here we present Colora, a Snakemake workflow that generates complete chromosome-scale assemblies, including organelles. Colora requires PacBio HiFi and Hi-C reads as mandatory inputs, and ONT reads can be optionally integrated into the process. With Colora, it is possible to obtain a scaffolded primary assembly or a phased assembly with separate haplotypes. Colora was primarily developed for plant genome assembly, as it includes plant-specific tools such as Oatk (https://github.com/c-zhou/oatk), but it can also be used for other organisms. Colora is the first automated workflow for *de novo* genome assembly implemented in Snakemake that integrates PacBio Hifi, ONT, and Hi-C reads to produce complete chromosome-scale assemblies.

## Implementation

Colora was constructed using the Snakemake workflow manager (version ≥ 8.0.0. The whole workflow is installed and runs using virtual environments created by Conda [7]. We provide frozen environments containing the tools with the version already tested to avoid breakage due to dependency updates.

Figure 1 shows a schematic representation of Colora.

**Figure 1:**
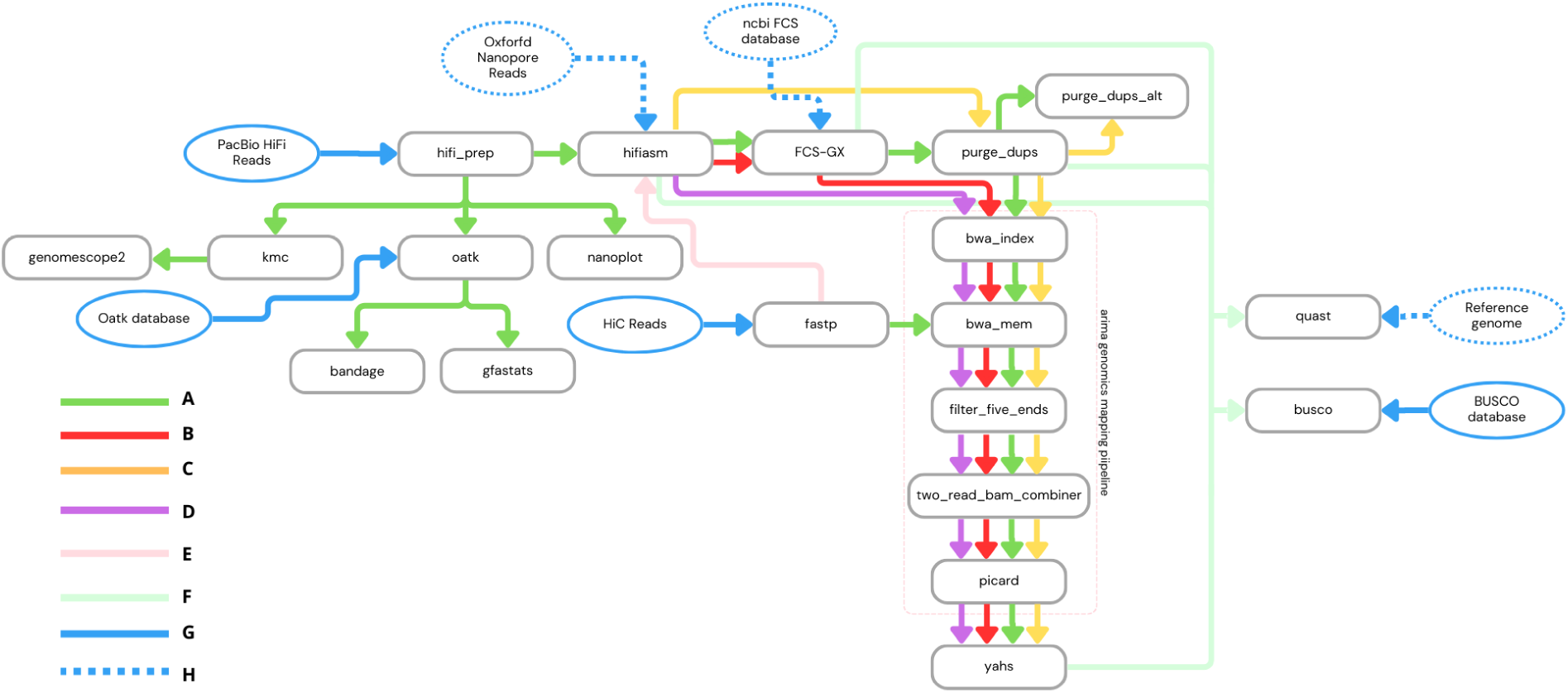
Flowchart illustrating the Colora Snakemake workflow. A: full workflow in case of primary assembly; B: skip Purge_dups step, full workflow in case of phased assembly; C: skip FCS-GX step; D: skip Purge_dups and FCS-GX steps; E: executed only in case of phased assembly; F: assembly quality inspection (checkpoints are dependent on the workflow configuration); G: mandatory input; H: optional input.

The choice of tools integrated into the workflow is based on those chosen by world-leading biodiversity projects, such as the Earth BioGenome Project (EBP) (https://www.earthbiogenome.org/), the Darwin Tree of Life (DToL) (https://www.darwintreeoflife.org/), and the Vertebrate Genomes Project (VGP) (https://vertebrategenomesproject.org/), and on our previous benchmarking [8].

### Reads pre-processing

The workflow executes the quality assessment of the reads used to perform the assembly process. After being joined in a single file with the rule hifi prep if needed, PacBio HiFi reads are checked for quality with NanoPlot [9]. As this kind of read typically does not need to be filtered before the assembly process, we did not implement an automatic filtering step. Hi-C reads are automatically checked and filtered for quality, and adapters are removed with Fastp [10]. ONT reads (if available) must be previously quality inspected and filtered if necessary.

### K-mers analysis, organelles and genome assembly

The k-mers are counted with KMC [11, 12, 13] from the PacBio HiFi reads, and the k-mer spectrum is plotted with GenomeScope2 [14].

The organelles are assembled from the PacBio HiFi reads using Oatk (https://github.com/c-zhou/oatk).

The assembly process starts with Hifiasm. Hifiasm can be run with PacBio HiFi reads only, in which case the result will be a primary assembly, or in “Hi-C mode”, i.e., integrating Hi-C reads to separate the haplotypes in heterozygous species, resulting in two separate assemblies representing the two haplotypes (hap1 and hap2). In both cases, ONT reads can be optionally integrated if available.

Table 1 outlines the differences between workflow paths that can be chosen by the user after the contig-level assembly with Hifiasm. It shows the possibility of performing the steps of contaminants removal with FCS-GX [15], removal of duplications and overlaps with Purge_dups [16], Hi-C reads mapping to primary contigs with the Arima Genomics pipeline (https://github.com/ArimaGenomics/mapping_pipeline), and assembly scaf-folding with YaHS [17] depending on the chosen configuration.

**Table 1:** Comparison of possible workflow paths following the primary contigs assembly with Hifiasm. Symbols: ✓ for “yes”, × for “no”.

| Workflow path | Contaminants removal from primary contigs | Overlap and duplications removal | Hi-C reads mapping to primary contigs | Assembly scaffolding | Applicable for primary assembly | Applicable for phased assembly | Arrow colour in Figure 1 |
| --- | --- | --- | --- | --- | --- | --- | --- |
| Skip contaminants, overlaps, and duplications removal | × | × | ✓ | ✓ | ✓ | ✓ | Purple |
| Skip overlaps and duplications removal | ✓ | × | ✓ | ✓ | ✓ | ✓ | Red |
| Skip contaminants removal | × | ✓ | ✓ | ✓ | ✓ | × | Yellow |
| Full workflow | ✓ | ✓ | ✓ | ✓ | ✓ | × | Green |

The Arima Genomics Mapping Pipeline (https://github.com/ArimaGenomics/mapping_pipeline) has been adapted to Snakemake from its original bash form. In addition, the flag -M has been added to the bwa mem commands [18].

### Results quality assessment

Assemblies produced at different steps along the workflow, depending on the configuration, are submitted to quality assessment. The genome assembly quality assessment is performed using BUSCO [19, 20] and QUAST [21].

The quality assessment of organelle assemblies is performed with Gfastats [22] and Bandage [23].

Hi-C contact maps of the scaffolded assemblies were produced after the completion of the workflow to gain further evidence about the success of the assembly process. They were obtained using the code shown in Additional Files S0 and visualised with PretextView (https://github.com/sanger-tol/PretextView).

### User specifications

The user can specify the desired parameters to use along the workflow and the route to follow by editing the config.yaml file. This is essential for a successful run of Colora.

Before starting the workflow, the user must be sure to have all the mandatory inputs, which are PacBio Hifi and Hi-C reads, the Oatk database (https://github.com/c-zhou/OatkDB), and the BUSCO database for the species studied (light blue continuous lines in Figure 1). Depending on the chosen configuration, other optional inputs that must be available before starting are ONT reads, the NCBI FCS-GX database, and the reference genome, optionally with the related annotation GFF file, for the species studied (light blue dotted lines in Figure 1). These inputs can be specified through the config.yaml file, as explained in detail in config/README.md and in the examples on our GitHub repository wiki (https://github.com/LiaOb21/colora).

The config.yaml allows the users to change several parameters for customisation of the tools according to their needs for the studied species. In the config/README.md, we have highlighted the key parameters that a user should check in the config.yaml and have provided links to the upstream documentation where users may find further details for each tool.

### Testing of the workflow

To test the pipeline, we used three publicly available datasets:

1. *Rhizophagus irregularis*, strain G1, was obtained from the BioProject PRJNA922099; the dataset included PacBio HiFi and Hi-C reads. This organism is a heterokaryotic arbuscular mycorrhizal fungus, in which cells two different kinds of haploid nuclei coexist [24]. The size of the reference genome is 146.8 Mbp, and it is organised into 32 chromosomes [25].
2. For *Arabidopsis thaliana*, the BioProject PRJCA005809 dataset included PacBio HiFi, ONT, and Hi-C reads [26]. This organism is the model plant for genomics, and its genome size is 135 Mbp organised in 5 chromosomes. The size of the reference genome TAIR10.1 is 119.1 Mbp [27]. Since this species is autogamous, the two haplotypes are identical by descendent (IBD), and its genome is therefore represented by a single haplotype.
3. *Malus domestica*, cultivar Fuji, was obtained from BioProject PRJNA814760; the dataset included PacBio HiFi and Hi-C reads [28]. This is an agronomically important species, and the size of the reference genomes is 703 Mbp organised in 17 chromosomes. This species is allogamous and heterozygous, and its genome is therefore expected to be diploid.

These datasets were used to produce *de novo* genome assemblies in previous studies. The assemblies produced in the original studies and the reference genome for each species were quality inspected using QUAST and BUSCO to compare the results with the quality of the assemblies produced with Colora.

For the *R. irregularis* and *M. domestica* datasets, the reads were downloaded from NCBI (https://www.ncbi.nlm.nih.gov/) using SRA tools (https://github.com/ncbi/sra-tools/wiki/01.-Downloading-SRA-Toolkit). *A. thaliana* data were downloaded from NGDC (https://ngdc.cncb.ac.cn/).

The codes used to test the workflow are available in the wiki on our GitHub repository. In addition, we provide a test dataset for users to test the workflow within a limited amount of time. This dataset is a subset of reads of *Saccharomyces cerevisiae* obtained from the BioProject PRJNA1075684 for PacBio and ONT reads and from the BioProject PRJNA1013711 for Hi-C reads [29]. This dataset is for testing purposes only.

Before running Colora using the *A. thaliana* dataset, we inspected ONT reads with NanoPlot [9] to assess read length and quality distributions. To improve data quality, the reads were filtered using NanoFilt [9] with the parameter -l 500 to exclude reads shorter than 500 bp. Given the limited mean read length of the dataset, we applied a shallow filter to avoid over-exclusion of data.

All the other parameters used to produce the assemblies are shown in the relative config.yaml files in Additional files (S1, S2, and S3).

Colora was tested on different systems, including HPCs and personal computers with CPUs Intel-i7 and Ubuntu-based OS.

While testing, we monitored the memory consumption along the workflow to assess performance and provide users with suggestions for memory requirements. These data are reported in the config/README.md file and in the config.yaml files in Additional files (S1, S2, and S3).

After the successful workflow, we produced a Snakemake report using the command snakemake --report to monitor the runtime of each rule.

## Results and Discussion

### Workflow performance

Figure 2 shows the runtime graphs obtained from the SnakeMake report for the three species. The Hifiasm rule was the most demanding rule in terms of time for the three datasets. Hifiasm required the highest amount of time for the *M. domestica* dataset.

**Figure 2:**
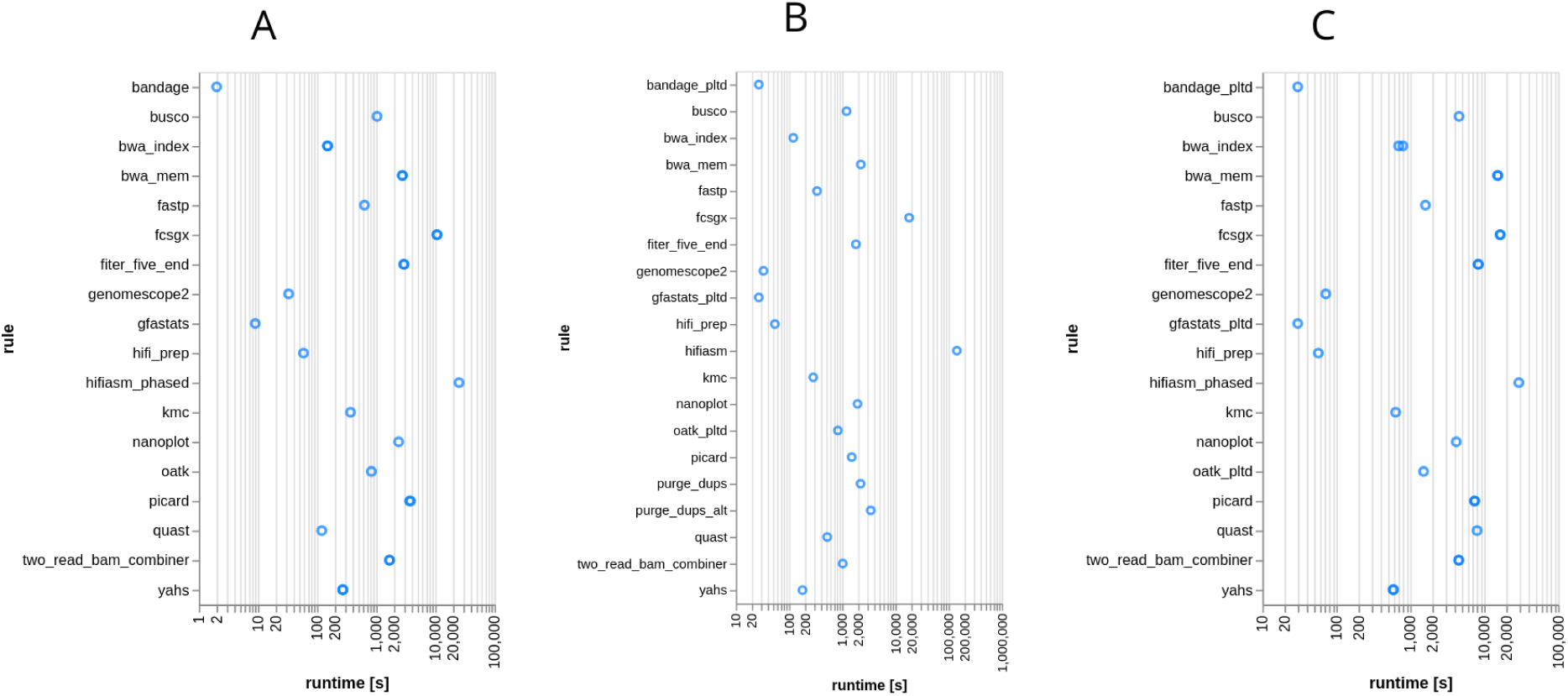
Runtime (seconds) required by each rule for *R. irregularis* (A), *A. thaliana* (B), and *M. domestica* (C).

Figure 3 shows the maximum resident size (kbytes) for each rule measured for the three datasets, i.e., the highest amount of RAM each rule required while it was running. Hifiasm was the most demanding rule in terms of memory for the three datasets, and it required the highest amount of RAM for the *M. domestica* dataset.

**Figure 3:**
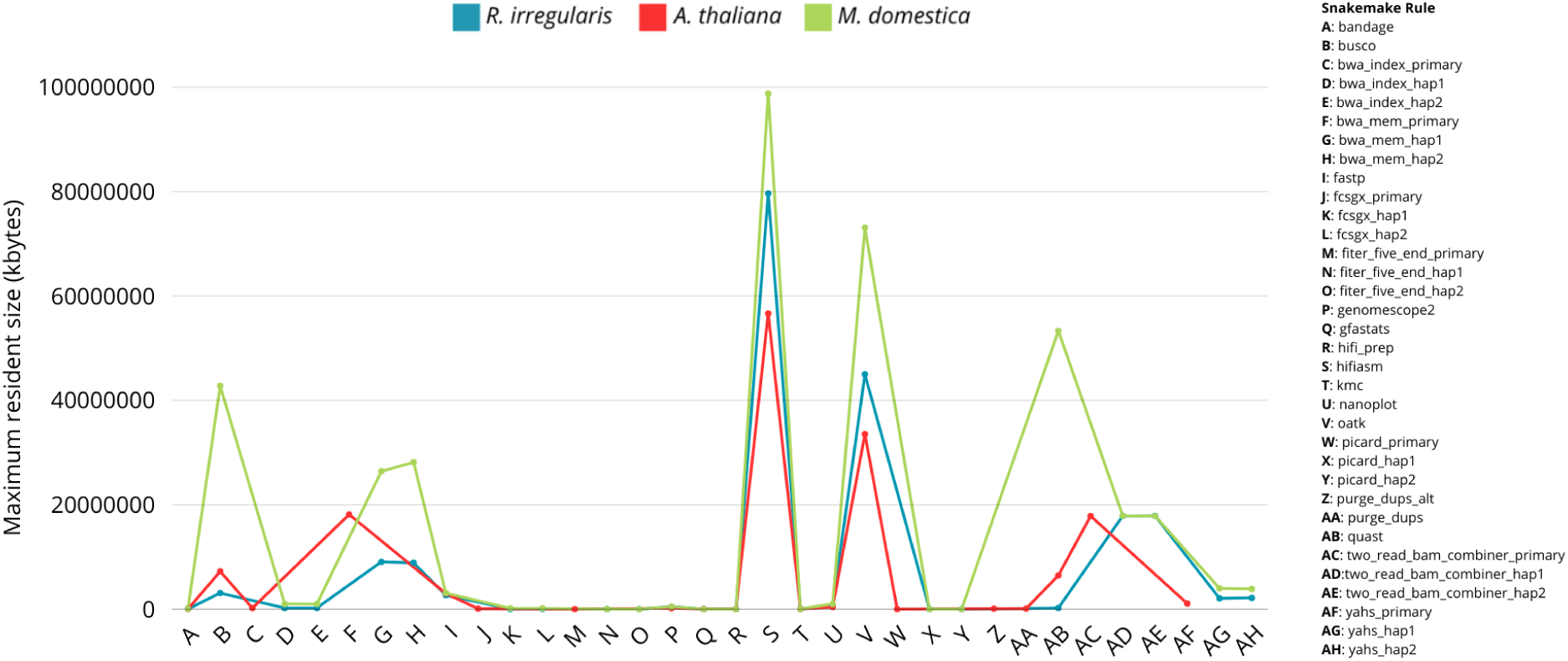
Maximum resident size (kbytes) required by each rule for the three datasets.

This is in line with the expectations, as the *M. domestica* genome was the largest among those analysed. Overall, the runtime and RAM requirements for each rule are dependent on the size of the dataset. Therefore, the workflow performance is strictly dependent on the dataset analysed.

### Reads quality assessment

The complete NanoPlot reports for PacBio HiFi reads (S4, S5, and S6), Fastp reports for Hi-C reads (S7, S8, and S9) for the three datasets, and the NanoPlot report for ONT *A. thaliana* dataset (S10) are shown in Additional files.

Table 2 summarises the read characteristics of the *R. irregularis*, *A. thaliana*, and *M. domestica* datasets.

**Table 2:** Overview of the read characteristics of the *R. irregularis* dataset (BioProject PRJNA922099), *A. thaliana* (BioProject PRJCA005809), and *M. domestica* (BioProject PRJNA814760)

| Species | Dataset | Mean read length | Mean read quality | Number of reads | Total bases | Coverage |
| --- | --- | --- | --- | --- | --- | --- |
| *R. irregularis* | PacBio HiFi | 12,812.1 | 21.0 | 2,341,873 | 30,004,306,961 | 204.11 |
|  | Hi-C | 150 | >30 | 236,467,312 | 35,470,097,000 | 241.29 |
| *A. thaliana* | PacBio HiFi | 15,094.4 | 31.8 | 1,517,433 | 22,904,700,074 | 169.66 |
|  | Hi-C | 150 | >30 | 140,957,500 | 21,143,625,000 | 156.62 |
|  | ONT | 18,541.3 | 11.1 | 3,064,191 | 56,814,196,989 | 420.84 |
| *M. domestica* | PacBio HiFi | 11,749.8 | 27.2 | 4,523,532 | 53,150,548,964 | 75.9 |
|  | Hi-C | 148 | >30 | 676,416,866 | 100,522,162,000 | 143.60 |

### GenomeScope2 profiles

Figure 4 shows the k-mer spectra of the three datasets under study.

**Figure 4:**
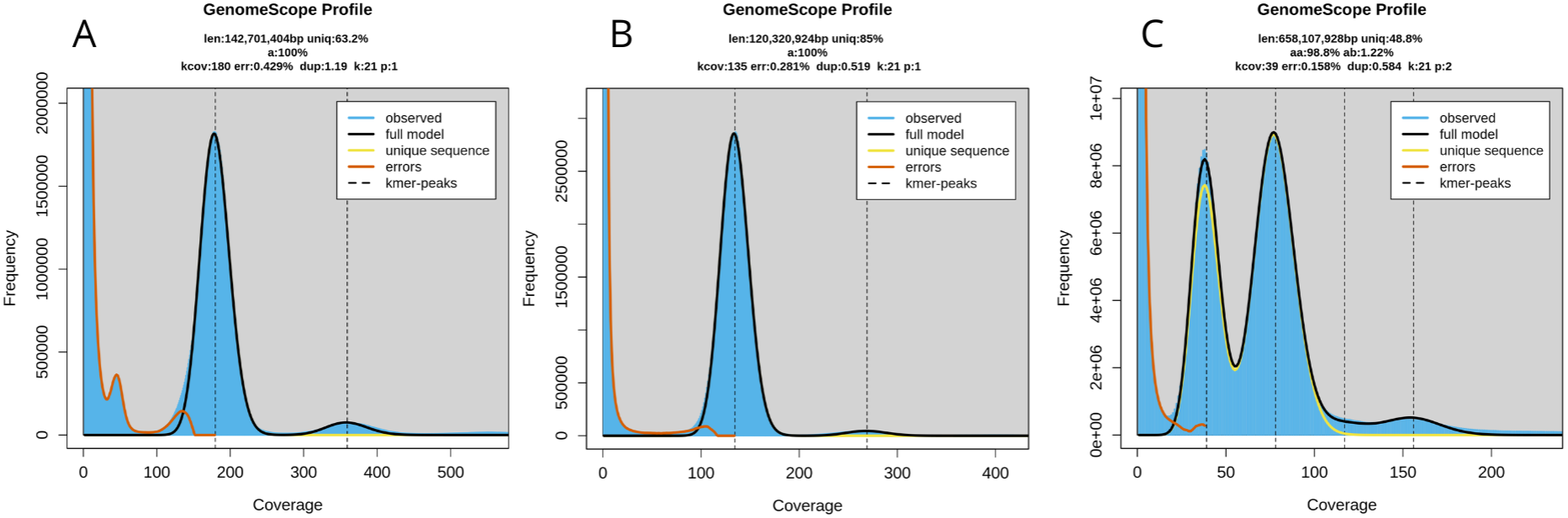
GenomeScope2 k-mer profiles for *R. irregularis* (A), *A. thaliana* (B), and *M. domestica* (C).

For *R. irregularis* (Figure 4A), GenomeScope2 calculated an estimated genome size of 142,701,404 bp, which is comparable to the size of the reference genome ASM2621079v1, and the genome size obtained in the original paper (Additional file S11). It was possible to obtain a similar estimate from the k-mer analysis only using the parameter -p 1 for GenomeScope2, which is used for homozygous genomes. This species is known to be heterokaryotic, i.e., it carries two different types of haploid nuclei in its cells [24]. It is likely that the small peak observed at coverage ∼50, erroneously placed by the model among the errors, represents the second haploid nucleus.

GenomeScope2 estimated a genome size of 120,320,924 bp for *A. thaliana* (Figure 4B), which is comparable to the reference genome of the species TAIR10.1, while the genome size of the assembly obtained in the original paper was 133.7 Mbp. As expected, the k-mer spectrum fits the haploid genome model, showing only one peak.

For *M. domestica*, GenomeScope2 estimated genome size of 658,107,928 bp (Figure 4C), close in size to the haploid assemblies from the original paper but lower compared to the reference genome ASM211411v1 and to the consensus haploid assembly from the original paper (Additional file S11). The k-mer spectrum was consistent with what is expected for a heterozygous genome, showing two peaks.

### Organelle assembly quality assessment

Figure 5 shows the statistics obtained with Gfastats [22] and the plot obtained with Bandage [23] for the mitochondrial assembly of the three species. For all the datasets, the mitochondrial assemblies were composed of a single, gapless contig and segment. Compared to the reference mitochondrion, the assemblies displayed close concordance in length.

**Figure 5:**
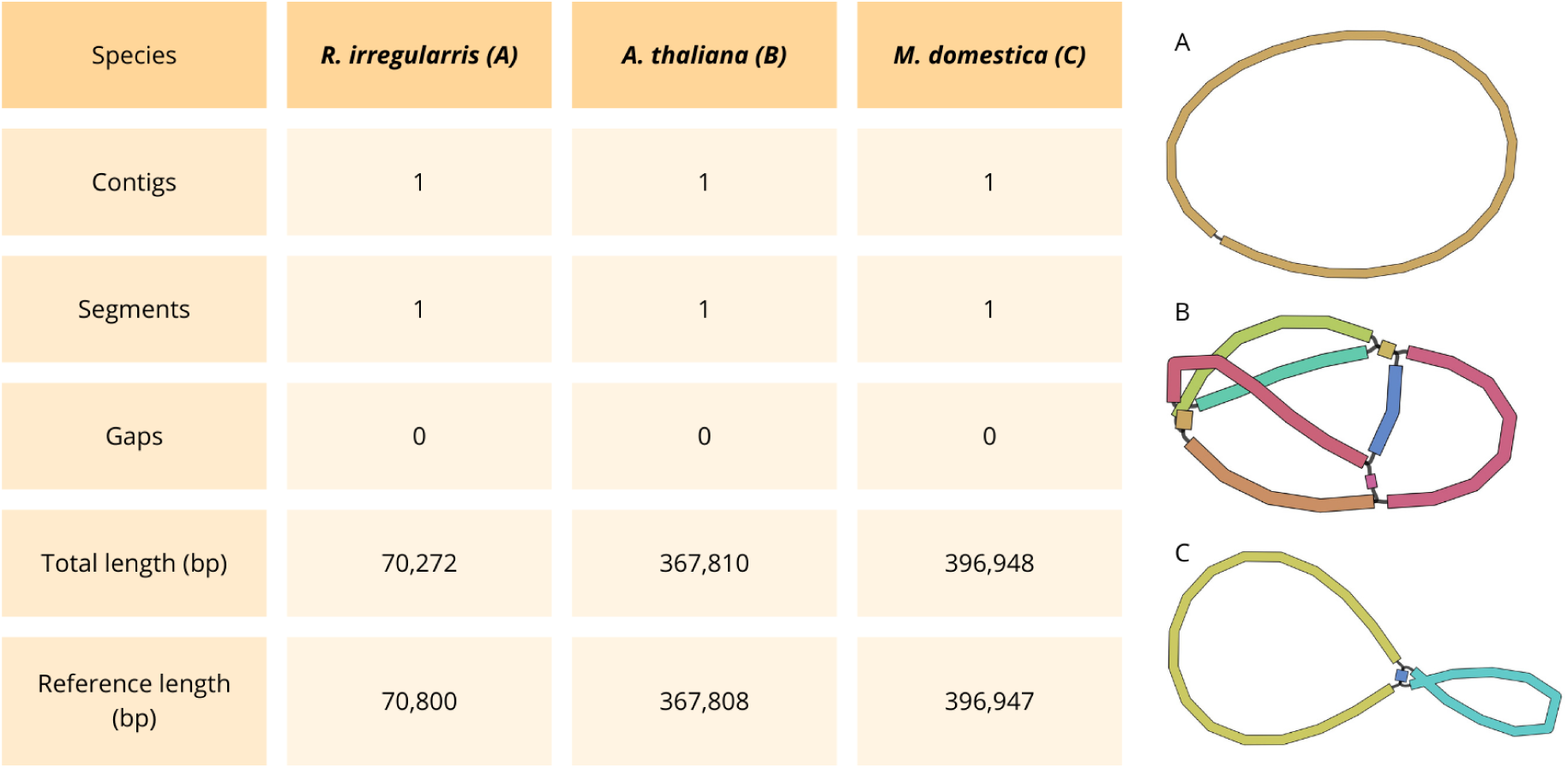
Gfastats statistics and Bandage visualisation for *R. irregularis* (A), *A. thaliana* (B), and *M. domestica* (C) mitochondria. NCBI reference sequences (RefSeq) are NC_014489.1, NC_037304.1, and NC_018554.1 respectively.

Figure 6 displays the statistics generated using Gfastats and the visualisation created with Bandage for the chloroplast assemblies of *A. thaliana* and *M. domestica*. In both datasets, the chloroplast assemblies consisted of a single, gapless contig and segment. The assembly lengths were compared to the reference chloroplast length, revealing comparable results.

**Figure 6:**
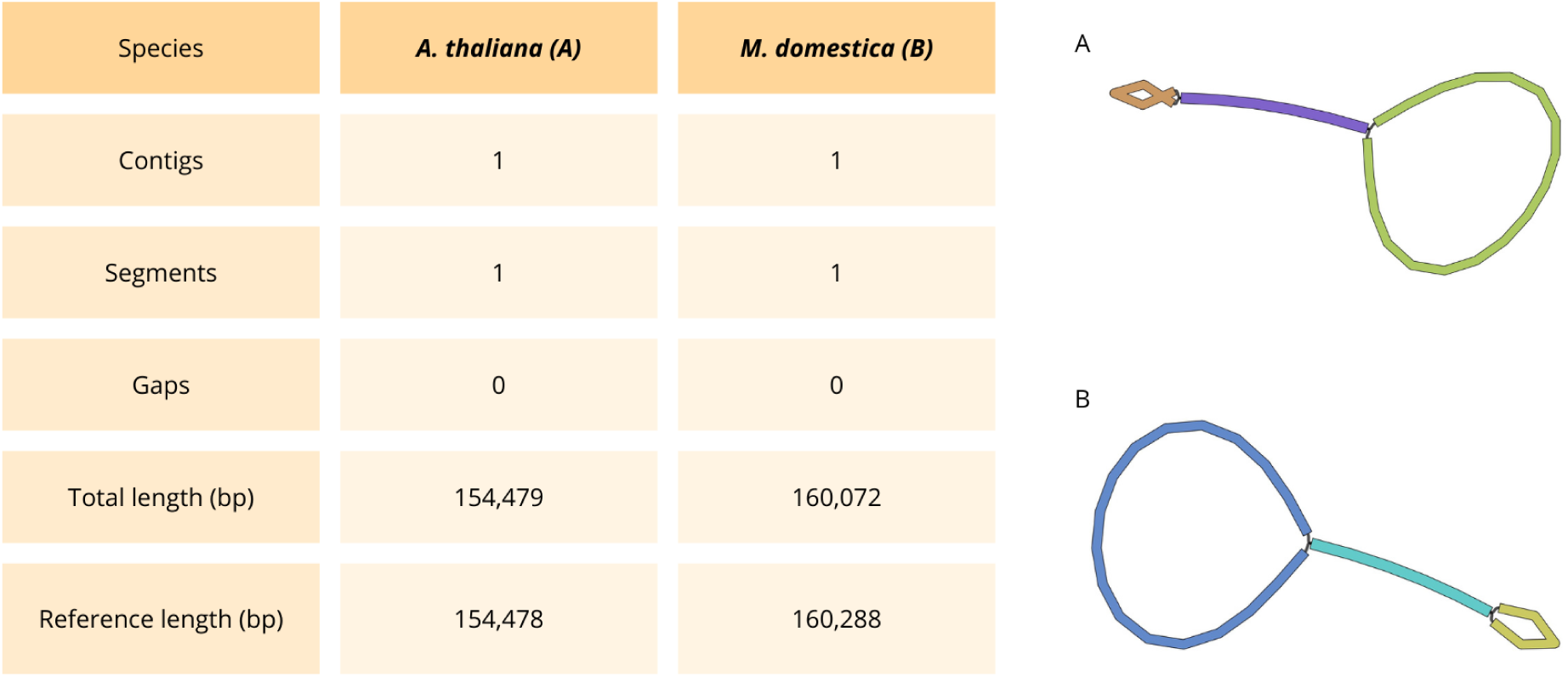
Chloroplast gfastats statistics and Bandage visualisation for *A. thaliana* (A), and *M. domestica* (B). NCBI reference sequences (RefSeq) are NC_000932.1 and NC_061549.1 respectively.

Complete Gfastats results for mitochondria (S12, S13, and S14) and chloroplast (S15, and S16) are shown in Additional files.

### Genome assembly quality assessment

Figure 7 shows the QUAST metrics for *R. irregularis*.

**Figure 7:**
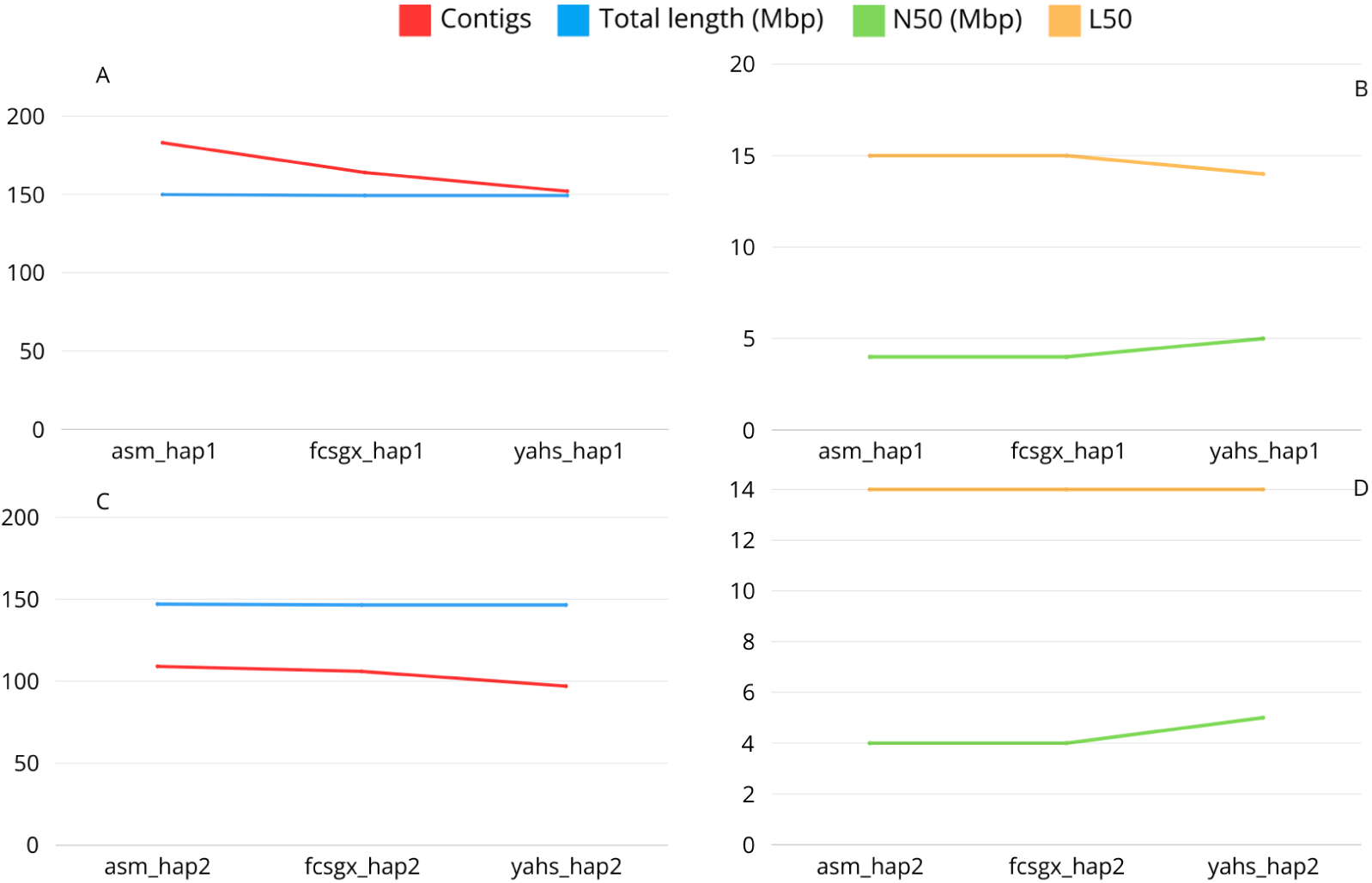
Quast metrics for *R.irregularis* for haplotype 1 (A,B) and haplotype 2 (C,D).

The number of contigs for the two haplotypes decreased along the workflow from 183 to 152 for hap1 and from 109 to 97 for hap2 due to the process of decontamination and scaffolding.

The number of contigs in the reference genome and the assemblies obtained in the original paper is lower compared to our assemblies (Additional File S11). However, these assemblies were manually curated [24, 25]. For our assemblies, further improvements may be achieved with manual curation, as shown by the Hi-C contact maps (Additional file S17 A, B), which displayed the presence of small contigs not included in the chromosome-length scaffolds for both assemblies. The assemblies could be improved further by removing contaminations from mitochondrial sequences, which are not automatically removed by FCS-GX.

The total length of the final assembly was 149.3 Mbp for hap1 and 146.5 Mbp for hap2, which is close to the genome size predicted by GenomeScope2 and comparable to the total length obtained for the assemblies of the original paper and the reference genome (Additional files S11).

The N50 of the final assembly was 4.9 Mbp for both haplotypes, which is comparable to the results obtained for the assemblies of the original paper and the reference genome. The L50 was 14 for both haplotypes in the final scaffolded assembly.

The complete QUAST report for *R. irregularis* is shown in Additional files (S11).

The QUAST metrics for *A. thaliana* are shown in Figure 8.

**Figure 8:**
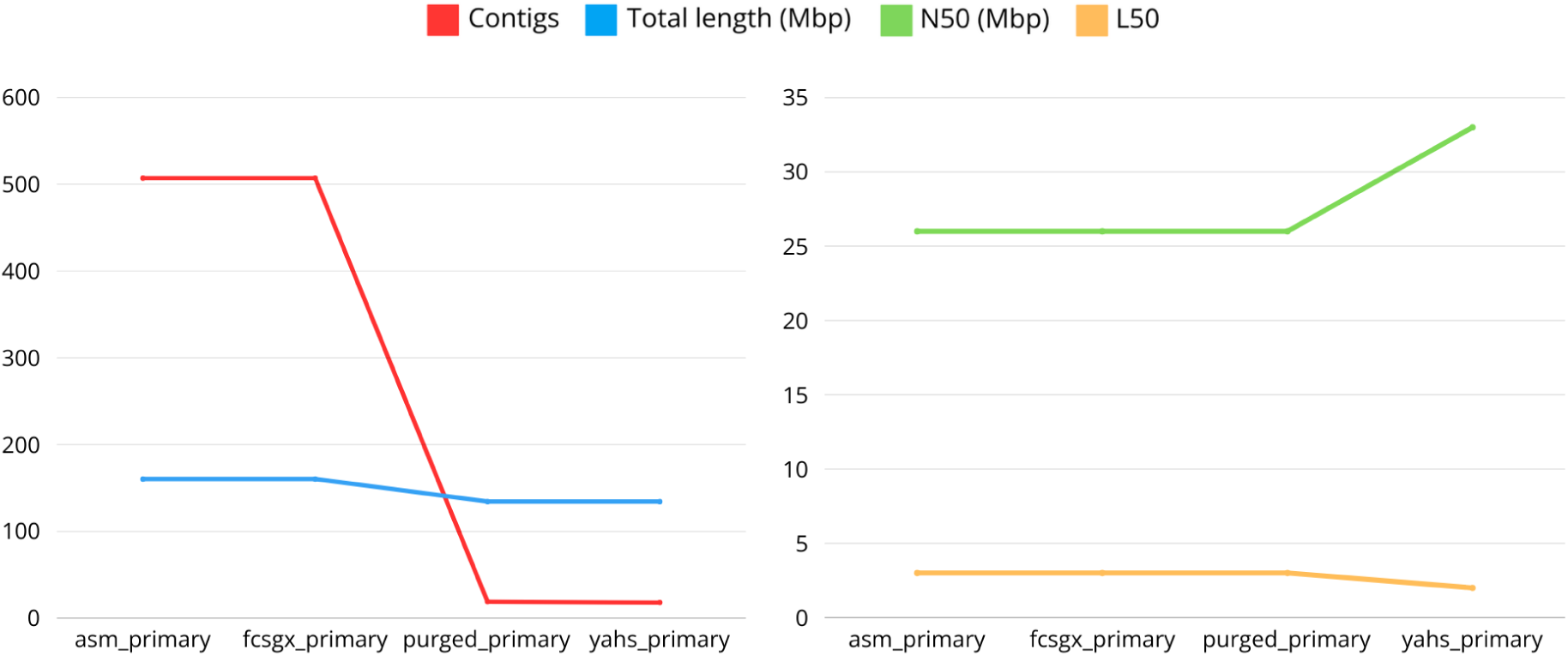
Quast metrics for *A. thaliana*

The number of contigs was drastically reduced during the workflow, from the initial 507 to the final 18, especially due to the removal of haplotigs and overlaps by Purge_dups. The total length of the final assembly was 134.38 Mbp. This result is distant from the estimated genome size obtained with GenomeScope2, but it is in line with the results obtained in the original paper, where the authors obtained a genome assembly of 133.7

Mbp. The reference genome TAIR10.1 has a genome size of 119.1 Mbp.

The L50 was equal to 2, indicating the possible introduction of a false join during the scaffolding process since we would expect this value to be equal to 3. The false join between the first two chromosomes was clearly shown by the Hi-C contact map (Additional file S17 C).

The N50 for the final assembly was 32.6 Mbp, which is higher compared to the results obtained for the reference genome and the original paper due to the presence of the false join between two scaffolds.

For this species, we expect 5 chromosomes, and this result can be achieved with manual curation of the assembly and by removing contaminations from mitochondrial and chloroplast sequences.

The complete QUAST report for *A. thaliana* is shown in Additional Files (S11).

Figure 9 shows the QUAST metrics for *M. domestica*.

**Figure 9:**
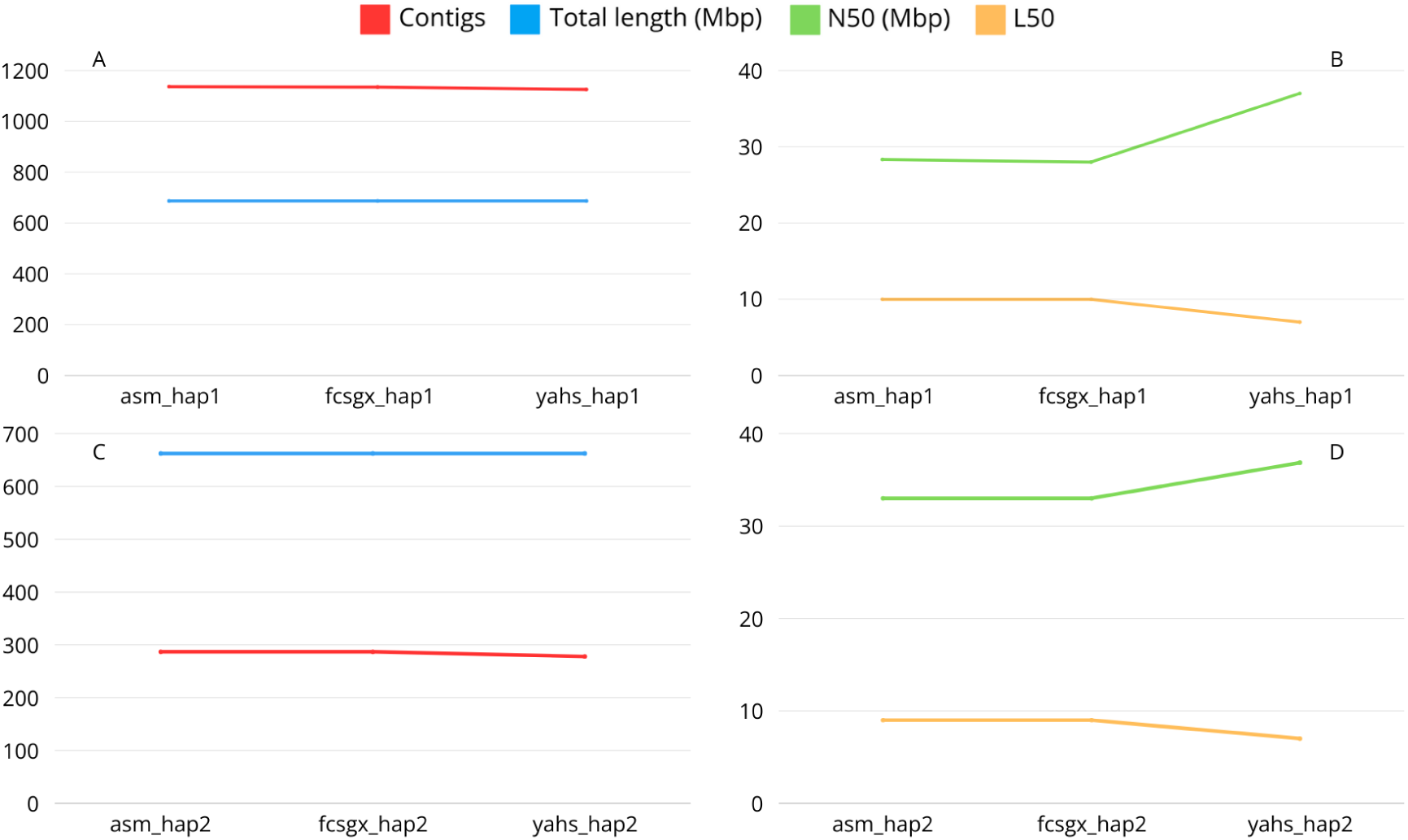
Quast metrics for *M. domestica* haplotype 1 (A,B) and haplotype 2 (C,D).

The number of contigs decreased along the workflow for both haplotypes. However, they did not reach a number of contigs matching the expected number of chromosomes for this species, which is 17. The same is true for the number of contigs observed in the reference genome and the assemblies produced in the original paper (Additional file S11). However, the assemblies can be improved with further manual curation and by removing contaminations from mitochondrial sequences.

The total length of the final assembly was 686.8 Mbp and 662.6 Mbp for hap1 and hap2, respectively. These results are comparable with the expected genome size shown by Genomescope2 and with the size obtained in the original paper for the two haplotypes. The reference genome ASM211411v1 for this species is 703 Mbp, while the consensus assembly obtained in the original paper showed a length equal to 736.9 Mbp (Additional Files S11).

The N50 was 37.2 Mbp and 36.9 Mbp for hap1 and hap2, respectively, showing values comparable to the assemblies from the original paper and to the reference genome (Additional Files S11).

The L50 was equal to 7 for both the haplotypes. For this species, we would expect an L50=9; therefore, this is an indication of the presence of false joins introduced during the process of scaffolding, which was confirmed by the Hi-C contact maps (Additional files S17 D, E).

Figure 10 shows the BUSCO assessment results for the three species under study. *R. irregularis* assemblies were compared with the BUSCO lineage mucoromycota odb10 (Figure 10A). The BUSCO scores for the final hap1 assembly were 98.2% complete BUSCOs (96.3% single copy, 1.9% duplicated), 0.3% fragmented BUSCOs, and 1.5% missing BUSCOs. For hap2, the results showed 97.8% complete BUSCOs, (97.0% single copy and 0.8% duplicated), 0.4% fragmented BUSCOs, and 1.8% missing BUSCOs. Overall, the results show that BUSCO scores remained unaltered throughout the workflow.

**Figure 10:**
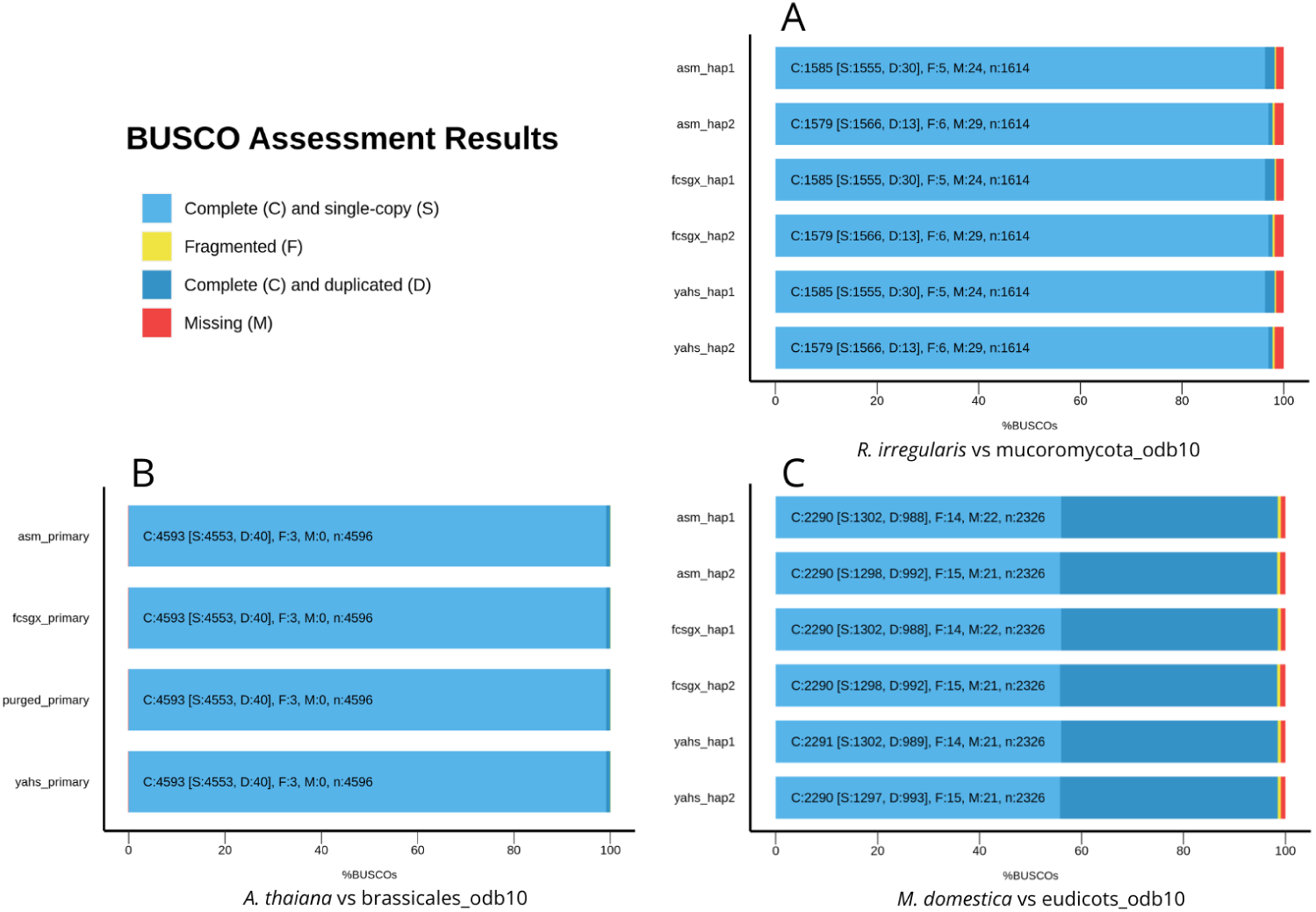
BUSCO Assessment results for *R. irregularis* (A), *A. thaliana* (B), and *M. domestica* (C).

*A. thaliana* assemblies were compared with the BUSCO lineage brassicales odb10 (Figure 10B). The BUSCO scores for the final assembly are 100.0% of complete BUSCOs (99.1% single copy and 0.9% duplicated), 0.1% fragmented BUSCOs, and no missing BUSCOs. Overall, the results show that BUSCO scores remained unaltered along the workflow.

*M. domestica* assemblies were compared with the BUSCO lineage eudicots odb10 (Figure 10C). The final BUSCO scores for hap1 were 98.5% complete BUSCOs (56.0% single copy and 42.5% duplicated), 0.6% fragmented BUSCOs, and 0.9% missing BUSCOs. The final BUSCO scores for hap2 were 98.5% complete BUSCOs (55.8% single copy and 42.7% duplicated), 0.6% fragmented BUSCOs, and 0.9% missing BUSCOs. For hap1, complete and duplicated BUSCOs slightly increased along the workflow, and missing BUSCOs slightly decreased. For hap2, complete and single copy BUSCOs slightly decreased along the workflow, while complete and duplicated BUSCOs slightly increased. The high degree of duplication in M. domestica genome is known and is the result of small-scale and large-scale whole-genome duplication events during its evolutionary history [30, 31].

Overall, the BUSCO metrics for the assemblies obtained with Colora were comparable to those observed for the reference genomes and the assemblies published in the original papers (Additional file S18).

Colora is the first Snakemake workflow for *de novo* genome assembly that combines PacBio Hifi, ONT, and Hi-C reads to obtain complete chromosome-scale assemblies. A previously published Snakemake workflow for *de novo* genome assembly, SnakeCube, utilises MinION ONT and Illumina reads as input [32]. However, there are Nextflow and Galaxy pipelines that share part of the assembly strategy and tools with Colora, and which inspired Colora. These pipelines are the Earth Biogenome assembly pipeline (https://github.com/NBISweden/Earth-Biogenome-Project-pilot) and the DToL assembly pipeline (https://pipelines.tol.sanger.ac.uk/genomeassembly), which are implemented in Nextflow, and the VGP pipeline, which is implemented in Galaxy [33]. Colora has many aspects in common with these pipelines but also its peculiarities. For example, the decontamination process is performed using FCS-GX and applied to the contig-level assembly. FCS-GX has only recently been implemented in the Earth BioGenome assembly pipeline. In Colora, this step is optional: this allows the user to use the pipeline in small systems, as the FCS-GX database needs a substantial amount of disk space, and the process requires a large amount of RAM to run. The removal of haplotigs with Purge_dups is also optional (in the case of primary assembly), as we observed that in large and highly repetitive genomes this can lead to a substantial reduction in complete BUSCOs (unpublished work). The tool used to assemble the organelles is Oatk, which was only recently implemented in the DToL pipeline. We chose this tool because it is optimised for the assembly of complex plant organelles, but it can also be used for simple organelle genomes. Colora also has the peculiarity of evaluating organelle assemblies using Gfastats and Bandage.

Overall, Colora automatically produces high-quality *de novo* assemblies, which are comparable to the assemblies obtained in the original papers with a step-by-step workflow. Subsequent manual curation, as suggested by Howe et al. (2021) [34], is necessary to further improve the quality of the assemblies.

## Limitations and future work

Colora is set up to run on Linux systems. The version described in this paper is v 1.0.0. In the future, we aim to incorporate Merqury [35] for genome quality assessment as soon as an updated Conda recipe is available. We plan to add a method that automatically removes contigs identified as organelles. We aim to keep all the tools included in the pipeline up to date to their latest versions, and to implement the containerisation with Docker and Singularity.

## Conclusions

Colora is a reliable, state-of-the-art, and portable pipeline for automated *de novo* genome assembly, specifically designed with plant genomes in mind. The primary goal of the workflow is to be user-friendly and adaptable to various systems and datasets. With detailed documentation to support users, we believe it will be valuable for researchers worldwide seeking a customisable, automated, and easy-to-implement solution for obtaining highquality genome assemblies.

## Availability and requirements

**Project name:** Colora

**Project home page**: https://github.com/LiaOb21/colora

**Operating system(s)**: Linux

**Programming language**: Python, Bash, Perl, Awk

**Other requirements**: Miniconda, Snakemake

**License**: MIT

Colora is also available at the Snakemake Workflow Catalog (https://snakemake.github.io/snakemake-workflow-catalog/?usage=LiaOb21%2Fcolora).

The public raw data used to test the workflow are available from the NCBI SRA database for *R. irregularis* (https://www.ncbi.nlm.nih.gov/sra?LinkName=biosample_sra&from_uid=32643414) and *M. domestica* (https://www.ncbi.nlm.nih.gov/sra?LinkName=biosample_sra&from_uid=26566345) and from the NGDC for *A. thaliana* (https://ngdc.cncb.ac.cn/bioproject/browse/PRJCA005809).

## List of abbreviations

PacBio: Pacific Biosciences
(PacBio) HiFi: High Fidelity
ONT: Oxford Nanopore Technlogies
HPC: High-Performance Computer
hap1: Haplotype 1
hap2: Haplotype 2
NCBI: National Center for Biotechnology Information
NGDC: National Genomics Data Center

## Acknowledgments

The authors would like to thank the organisers of Biodiversity Genomics Academy 2023 (https://bga23.org/), which was essential for acquiring the knowledge that determined the success of this project.

## Authors’ contributions

LO designed, developed and tested the workflow. TB provided substantial technical support with Snakemake and suggestions about the design of the workflow. HDW provided suggestions about the design of the workflow and helped with the testing. UT and AP provided guidance and support. LO, TB, HDW, UT and AP wrote the manuscript.

