## Additional files for "Colora: A Snakemake Workflow for Complete Chromosome-scale *De Novo* Genome Assembly": S0_Hi_C_maps_code.pdf

### Code used to visualise the Hi-C contact maps

The example code is referring to a phased assembly. In case of primary assembly substitute `hap1` with `primary` in directory names.

Required files:

- `~/colora/results/bwa_index_hap1/asm.fa`
- `~/colora/results/yahs_hap1/asm_yahs_scaffolds_final.agp`
- `~/colora/results/yahs_hap1/asm_yahs.bin`
- `~/colora/results/yahs_hap1/asm_yahs_scaffolds_final.fa`

Required scripts and tools:

- `juicer pre` from YaHS (https://github.com/c-zhou/yahs/blob/main/juicer.c)
- Juicer Tools (https://github.com/aidenlab/JuicerTools)
- PretextMap (https://github.com/sanger-tol/PretextMap)

```
samtools faidx asm.fa

(juicer pre asm_yahs.bin asm_yahs_scaffolds_final.agp asm.fa.fai | sort -k2,2d -k6,6d -T ./ --parallel=8 -S64G | awk 'NF' > alignments_sorted.txt.part) && (mv alignments_sorted.txt.part alignments_sorted.txt)

samtools faidx asm_yahs_scaffolds_final.fa

awk '{print $1, $2}' asm_yahs_scaffolds_final.fa.fai > scaffolds_final.chrom.sizes

( awk 'BEGIN{print "## pairs format v1.0"} {print "#chromsize:\t"$1"\t"$2} END{print "#columns:\treadID\tchr1\tpos1\tchr2\tpos2\tstrand1\tstrand2"}' scaffolds_final.chrom.sizes ; awk '{print ".\t"$2"\t"$3"\t"$6"\t"$7"\t.\t."}' alignments_sorted.txt ) > temp_output.txt

PretextMap -o out_scaffolds_final.pretext --mapq 0 --highres < temp_output.txt
```

The file `out_scaffolds_final.pretext` can be subsequently visualised in PretextView (https://github.com/sanger-tol/PretextView).
