## Additional files for "Colora: A Snakemake Workflow for Complete Chromosome-scale *De Novo* Genome Assembly": S2_A.thaliana_config.yaml

```
# config.yaml for A. thaliana

# Set memory and threads for high demanding rules
high:
  mem_mb: 153600 # memory in MB
  t: 50 # number of threads

# Set memory and threads for medium demanding rules
medium:
  mem_mb: 20480 # memory in MB
  t: 20 # number of threads

# Set memory and threads for low demanding rules
low:
  mem_mb: 10240 # memory in MB
  t: 8 # number of threads

# Path to hifi reads
hifi_path: "resources/raw_hifi/"

# Path to hic reads
hic_path: "resources/raw_hic/"

# Customisable parameters for kmc
kmc:
  k: 21 # kmer size, it will be the same used for genomescope2
  ci: 1 # exclude k-mers occurring less than <value> times (default: 2)
  cs: 10000 #maximal value of a counter (default: 255)

# Customisable parameters for kmc_tools transform
kmc_tools:
  cx: 10000 # exclude k-mers occurring more of than <value> times

# Customisable parameters for genomescope2
genomescope2:
  optional_params:
    "-p": "1"
    "-l": ""

# Customisable parameters for oatk
oatk:
  k: 1001 # kmer size [1001]
  c: 150 #  minimum kmer coverage [3]
  m: "resources/oatkDB/embryophyta_mito.fam" # mitochondria gene annotation HMM profile database [NULL]
  optional_params: 
    "-p": "resources/oatkDB/embryophyta_pltd.fam" # to use for species that have a plastid db

# Customisable parameters for fastp
fastp:
  optional_params:
    "--cut_front": False # to use only with Arima Hi-C library prep kit generated data
    "--cut_front_window_size": "" # to use only with Arima Hi-C library prep kit generated data

# Customisable parameters for hifiasm
hifiasm:
  phased_assembly: False # set to true if you want to obtain a phased assembly
  optional_params: 
    "-f": "" # used for small datasets
    "-l": "" # purge level. 0: no purging; 1: light; 2/3: aggressive [0 for trio; 3 for unzip]
    "--ul": "/home/edg01/edg01/lia/colora_paper/ara_thal/colora/resources/raw_ont/CRR302667_trimmed.fastq.gz" # use this if you have also ont data you want to integrate in your assembly

#Set this to False if you want to skip the fcsgx step:
include_fcsgx: True #inlcude this rule only if you have previously downloaded the database (recommended to run fcsgx only on a HPC. It requires around 500 GB of space on your disk and a large RAM)

# Customisable parameters for fcsgx
fcsgx:
  ncbi_tax_id: 3702
  path_to_gx_db: "resources/gxdb"

# Set this to False if you want to skip purge_dups steps:
include_purge_dups: True

# Customisable parameters for arima mapping pipeline:
arima:
  MAPQ_FILTER: 10

# Customisable parameters for yahs
yahs:
  optional_params: 
    "-e": "" # you can specify the restriction enzyme(s) used by the Hi-C experiment

# Customisable parameters for quast
quast:
  optional_params: 
    "--fragmented": ""
    "--large": ""
    "-r": "resources/reference/GCA_000001735.2_TAIR10.1_genomic.fna"
    "-g": "resources/reference/GCA_000001735.2_TAIR10.1_genomic.gff"

# Customisable parameters for busco
busco:
  lineage: "resources/busco_db/brassicales_odb10" # lineage to be used for busco analysis
```
