## Additional files for "Colora: A Snakemake Workflow for Complete Chromosome-scale *De Novo* Genome Assembly": S4_NanoPlot_R.irregularis.html

NanoPlot Report


### NanoPlot statistics report

#### Menu

- Summary Statistics
- Plots
  - Weighted histogram of read lengths
  - Weighted histogram of read lengths after log transformation
  - Non weighted histogram of read lengths
  - Non weighted histogram of read lengths after log transformation
  - Yield by length
  - Read lengths vs Average read quality plot using dots
  - Read lengths vs Average read quality plot using dots after log transformation of read lengths
- Report issue on Github


#### NanoPlot reports

##### Summary statistics

|  |  |
| --- | --- |
| General summary |  |
| Mean read length | 12,812.1 |
| Mean read quality | 21.0 |
| Median read length | 13,219.0 |
| Median read quality | 30.6 |
| Number of reads | 2,341,873.0 |
| Read length N50 | 13,983.0 |
| STDEV read length | 4,115.0 |
| Total bases | 30,004,306,961.0 |
| Number, percentage and megabases of reads above quality cutoffs |  |
| >Q5 | 2341873 (100.0%) 30004.3Mb |
| >Q7 | 2340830 (100.0%) 29973.2Mb |
| >Q10 | 2327686 (99.4%) 29711.5Mb |
| >Q12 | 2303673 (98.4%) 29357.2Mb |
| >Q15 | 2206465 (94.2%) 28008.5Mb |
| Top 5 highest mean basecall quality scores and their read lengths |  |
| 1 | 93.0 (5864) |
| 2 | 93.0 (1928) |
| 3 | 93.0 (3567) |
| 4 | 93.0 (2927) |
| 5 | 93.0 (1736) |
| Top 5 longest reads and their mean basecall quality score |  |
| 1 | 132789 (7.1) |
| 2 | 128512 (6.1) |
| 3 | 123726 (9.0) |
| 4 | 119378 (6.3) |
| 5 | 116180 (6.7) |

##### Plots

Weighted histogram of read lengths

###### Weighted histogram of read lengths

Weighted histogram of read lengths after log transformation

###### Weighted histogram of read lengths after log transformation

Non weighted histogram of read lengths

###### Non weighted histogram of read lengths

Non weighted histogram of read lengths after log transformation

###### Non weighted histogram of read lengths after log transformation

Yield by length

###### Yield by length

Read lengths vs Average read quality plot using dots

###### Read lengths vs Average read quality plot using dots

Read lengths vs Average read quality plot using dots after log transformation of read lengths

###### Read lengths vs Average read quality plot using dots after log transformation of read lengths
