## Additional files for "Colora: A Snakemake Workflow for Complete Chromosome-scale *De Novo* Genome Assembly": S5_NanoPlot_A.thaliana.html

NanoPlot Report


### NanoPlot statistics report

#### Menu

- Summary Statistics
- Plots
  - Weighted histogram of read lengths
  - Weighted histogram of read lengths after log transformation
  - Non weighted histogram of read lengths
  - Non weighted histogram of read lengths after log transformation
  - Yield by length
  - Read lengths vs Average read quality plot using dots
  - Read lengths vs Average read quality plot using dots after log transformation of read lengths
- Report issue on Github


#### NanoPlot reports

##### Summary statistics

|  |  |
| --- | --- |
| General summary |  |
| Mean read length | 15,094.4 |
| Mean read quality | 27.1 |
| Median read length | 14,936.0 |
| Median read quality | 31.4 |
| Number of reads | 1,517,433.0 |
| Read length N50 | 15,424.0 |
| STDEV read length | 2,661.7 |
| Total bases | 22,904,700,074.0 |
| Number, percentage and megabases of reads above quality cutoffs |  |
| >Q5 | 1517433 (100.0%) 22904.7Mb |
| >Q7 | 1517433 (100.0%) 22904.7Mb |
| >Q10 | 1517433 (100.0%) 22904.7Mb |
| >Q12 | 1517433 (100.0%) 22904.7Mb |
| >Q15 | 1517433 (100.0%) 22904.7Mb |
| Top 5 highest mean basecall quality scores and their read lengths |  |
| 1 | 93.0 (89) |
| 2 | 93.0 (1825) |
| 3 | 93.0 (3415) |
| 4 | 93.0 (1581) |
| 5 | 93.0 (52) |
| Top 5 longest reads and their mean basecall quality score |  |
| 1 | 45035 (22.6) |
| 2 | 43917 (21.8) |
| 3 | 43745 (20.5) |
| 4 | 43682 (23.6) |
| 5 | 43626 (22.0) |

##### Plots

Weighted histogram of read lengths

###### Weighted histogram of read lengths

Weighted histogram of read lengths after log transformation

###### Weighted histogram of read lengths after log transformation

Non weighted histogram of read lengths

###### Non weighted histogram of read lengths

Non weighted histogram of read lengths after log transformation

###### Non weighted histogram of read lengths after log transformation

Yield by length

###### Yield by length

Read lengths vs Average read quality plot using dots

###### Read lengths vs Average read quality plot using dots

Read lengths vs Average read quality plot using dots after log transformation of read lengths

###### Read lengths vs Average read quality plot using dots after log transformation of read lengths
