## Additional files for "Colora: A Snakemake Workflow for Complete Chromosome-scale *De Novo* Genome Assembly": S6_NanoPlot_M.domestica.html

NanoPlot Report


### NanoPlot statistics report

#### Menu

- Summary Statistics
- Plots
  - Weighted histogram of read lengths
  - Weighted histogram of read lengths after log transformation
  - Non weighted histogram of read lengths
  - Non weighted histogram of read lengths after log transformation
  - Yield by length
  - Read lengths vs Average read quality plot using dots
  - Read lengths vs Average read quality plot using dots after log transformation of read lengths
- Report issue on Github


#### NanoPlot reports

##### Summary statistics

|  |  |
| --- | --- |
| General summary |  |
| Mean read length | 11,749.8 |
| Mean read quality | 27.2 |
| Median read length | 11,619.0 |
| Median read quality | 32.3 |
| Number of reads | 4,523,532.0 |
| Read length N50 | 12,074.0 |
| STDEV read length | 1,871.6 |
| Total bases | 53,150,548,964.0 |
| Number, percentage and megabases of reads above quality cutoffs |  |
| >Q5 | 4523532 (100.0%) 53150.5Mb |
| >Q7 | 4523532 (100.0%) 53150.5Mb |
| >Q10 | 4523532 (100.0%) 53150.5Mb |
| >Q12 | 4523532 (100.0%) 53150.5Mb |
| >Q15 | 4523532 (100.0%) 53150.5Mb |
| Top 5 highest mean basecall quality scores and their read lengths |  |
| 1 | 93.0 (90) |
| 2 | 93.0 (2699) |
| 3 | 93.0 (2069) |
| 4 | 93.0 (1748) |
| 5 | 93.0 (48) |
| Top 5 longest reads and their mean basecall quality score |  |
| 1 | 39353 (21.3) |
| 2 | 36707 (21.0) |
| 3 | 35561 (26.7) |
| 4 | 35051 (23.6) |
| 5 | 34635 (21.7) |

##### Plots

Weighted histogram of read lengths

###### Weighted histogram of read lengths

Weighted histogram of read lengths after log transformation

###### Weighted histogram of read lengths after log transformation

Non weighted histogram of read lengths

###### Non weighted histogram of read lengths

Non weighted histogram of read lengths after log transformation

###### Non weighted histogram of read lengths after log transformation

Yield by length

###### Yield by length

Read lengths vs Average read quality plot using dots

###### Read lengths vs Average read quality plot using dots

Read lengths vs Average read quality plot using dots after log transformation of read lengths

###### Read lengths vs Average read quality plot using dots after log transformation of read lengths
