## Additional files for "Colora: A Snakemake Workflow for Complete Chromosome-scale *De Novo* Genome Assembly": S7_fastp_R.irregularis.html

fastp report at 2024-04-26 16:54:08 

### fastp report Summary General | | | | --- | --- | | fastp version: | 0.23.4 (https://github.com/OpenGene/fastp) | | sequencing: | paired end (150 cycles + 150 cycles) | | mean length before filtering: | 150bp, 150bp | | mean length after filtering: | 148bp, 148bp | | duplication rate: | 5.692436% | | Insert size peak: | 269 | Before filtering | | | | --- | --- | | total reads: | 236.467312 M | | total bases: | 35.470097 G | | Q20 bases: | 33.735107 G (95.108585%) | | Q30 bases: | 31.768578 G (89.564396%) | | GC content: | 29.808992% | After filtering | | | | --- | --- | | total reads: | 227.063358 M | | total bases: | 33.791405 G | | Q20 bases: | 32.502895 G (96.186870%) | | Q30 bases: | 30.812385 G (91.184089%) | | GC content: | 29.568264% | Filtering result | | | | --- | --- | | reads passed filters: | 227.063358 M (96.023148%) | | reads with low quality: | 9.394622 M (3.972905%) | | reads with too many N: | 9.332000 K (0.003946%) | | reads too short: | 0 (0.000000%) | Adapters Adapter or bad ligation of read1 The input has little adapter percentage (~0.756375%), probably it's trimmed before. | | | | --- | --- | | Sequence | Occurrences | | C | 140030 | | CT | 134938 | | CTG | 131484 | | CTGT | 127786 | | CTGTC | 124636 | | CTGTCT | 121232 | | CTGTCTC | 117853 | | CTGTCTCT | 114124 | | CTGTCTCTT | 110525 | | CTGTCTCTTA | 106777 | | CTGTCTCTTAT | 103290 | | CTGTCTCTTATA | 100165 | | CTGTCTCTTATAC | 97081 | | CTGTCTCTTATACA | 94809 | | CTGTCTCTTATACAC | 91620 | | CTGTCTCTTATACACA | 90413 | | CTGTCTCTTATACACAT | 87622 | | CTGTCTCTTATACACATC | 84469 | | CTGTCTCTTATACACATCT | 81544 | | CTGTCTCTTATACACATCTC | 78966 | | CTGTCTCTTATACACATCTCC | 75950 | | CTGTCTCTTATACACATCTCCG | 73568 | | CTGTCTCTTATACACATCTCCGA | 71045 | | CTGTCTCTTATACACATCTCCGAG | 68240 | | CTGTCTCTTATACACATCTCCGAGC | 66646 | | CTGTCTCTTATACACATCTCCGAGCC | 64842 | | CTGTCTCTTATACACATCTCCGAGCCC | 63553 | | CTGTCTCTTATACACATCTCCGAGCCCA | 61565 | | CTGTCTCTTATACACATCTCCGAGCCCAC | 59059 | | CTGTCTCTTATACACATCTCCGAGCCCACG | 56659 | | CTGTCTCTTATACACATCTCCGAGCCCACGA | 54065 | | CTGTCTCTTATACACATCTCCGAGCCCACGAG | 51882 | | CTGTCTCTTATACACATCTCCGAGCCCACGAGA | 50235 | | CTGTCTCTTATACACATCTCCGAGCCCACGAGAC | 48712 | | CTGTCTCTTATACACATCTCCGAGCCCACGAGACG | 47512 | | other adapter sequences | 1633000 | Adapter or bad ligation of read2 The input has little adapter percentage (~0.756375%), probably it's trimmed before. | | | | --- | --- | | Sequence | Occurrences | | C | 140045 | | CT | 135069 | | CTG | 131333 | | CTGT | 128079 | | CTGTC | 124875 | | CTGTCT | 121303 | | CTGTCTC | 117970 | | CTGTCTCT | 114219 | | CTGTCTCTT | 110700 | | CTGTCTCTTA | 107103 | | CTGTCTCTTAT | 103136 | | CTGTCTCTTATA | 99958 | | CTGTCTCTTATAC | 96855 | | CTGTCTCTTATACA | 94740 | | CTGTCTCTTATACAC | 91985 | | CTGTCTCTTATACACA | 90461 | | CTGTCTCTTATACACAT | 87547 | | CTGTCTCTTATACACATC | 84553 | | CTGTCTCTTATACACATCT | 81691 | | CTGTCTCTTATACACATCTG | 78954 | | CTGTCTCTTATACACATCTGA | 75966 | | CTGTCTCTTATACACATCTGAC | 73693 | | CTGTCTCTTATACACATCTGACG | 71373 | | CTGTCTCTTATACACATCTGACGC | 68595 | | CTGTCTCTTATACACATCTGACGCT | 67082 | | CTGTCTCTTATACACATCTGACGCTG | 65545 | | CTGTCTCTTATACACATCTGACGCTGC | 64479 | | CTGTCTCTTATACACATCTGACGCTGCC | 62541 | | CTGTCTCTTATACACATCTGACGCTGCCG | 60192 | | CTGTCTCTTATACACATCTGACGCTGCCGA | 57851 | | CTGTCTCTTATACACATCTGACGCTGCCGAC | 55505 | | CTGTCTCTTATACACATCTGACGCTGCCGACG | 53316 | | CTGTCTCTTATACACATCTGACGCTGCCGACGA | 51599 | | CTGTCTCTTATACACATCTGACGCTGCCGACGAC | 49445 | | CTGTCTCTTATACACATCTGACGCTGCCGACGACT | 48120 | | other adapter sequences | 1620019 | Insert size estimation This estimation is based on paired-end overlap analysis, and there are 60.315413% reads found not overlapped. The nonoverlapped read pairs may have insert size <30 or >270, or contain too much sequencing errors to be detected as overlapped. Before filtering Before filtering: read1: quality Value of each position will be shown on mouse over. Before filtering: read1: base contents Value of each position will be shown on mouse over. Before filtering: read1: KMER counting Darker background means larger counts. The count will be shown on mouse over. | | | | | | | | | | | | | | | | | | | --- | --- | --- | --- | --- | --- | --- | --- | --- | --- | --- | --- | --- | --- | --- | --- | --- | | | AA | AT | AC | AG | TA | TT | TC | TG | CA | CT | CC | CG | GA | GT | GC | GG | | AAA | AAAAA | AAAAT | AAAAC | AAAAG | AAATA | AAATT | AAATC | AAATG | AAACA | AAACT | AAACC | AAACG | AAAGA | AAAGT | AAAGC | AAAGG | | AAT | AATAA | AATAT | AATAC | AATAG | AATTA | AATTT | AATTC | AATTG | AATCA | AATCT | AATCC | AATCG | AATGA | AATGT | AATGC | AATGG | | AAC | AACAA | AACAT | AACAC | AACAG | AACTA | AACTT | AACTC | AACTG | AACCA | AACCT | AACCC | AACCG | AACGA | AACGT | AACGC | AACGG | | AAG | AAGAA | AAGAT | AAGAC | AAGAG | AAGTA | AAGTT | AAGTC | AAGTG | AAGCA | AAGCT | AAGCC | AAGCG | AAGGA | AAGGT | AAGGC | AAGGG | | ATA | ATAAA | ATAAT | ATAAC | ATAAG | ATATA | ATATT | ATATC | ATATG | ATACA | ATACT | ATACC | ATACG | ATAGA | ATAGT | ATAGC | ATAGG | | ATT | ATTAA | ATTAT | ATTAC | ATTAG | ATTTA | ATTTT | ATTTC | ATTTG | ATTCA | ATTCT | ATTCC | ATTCG | ATTGA | ATTGT | ATTGC | ATTGG | | ATC | ATCAA | ATCAT | ATCAC | ATCAG | ATCTA | ATCTT | ATCTC | ATCTG | ATCCA | ATCCT | ATCCC | ATCCG | ATCGA | ATCGT | ATCGC | ATCGG | | ATG | ATGAA | ATGAT | ATGAC | ATGAG | ATGTA | ATGTT | ATGTC | ATGTG | ATGCA | ATGCT | ATGCC | ATGCG | ATGGA | ATGGT | ATGGC | ATGGG | | ACA | ACAAA | ACAAT | ACAAC | ACAAG | ACATA | ACATT | ACATC | ACATG | ACACA | ACACT | ACACC | ACACG | ACAGA | ACAGT | ACAGC | ACAGG | | ACT | ACTAA | ACTAT | ACTAC | ACTAG | ACTTA | ACTTT | ACTTC | ACTTG | ACTCA | ACTCT | ACTCC | ACTCG | ACTGA | ACTGT | ACTGC | ACTGG | | ACC | ACCAA | ACCAT | ACCAC | ACCAG | ACCTA | ACCTT | ACCTC | ACCTG | ACCCA | ACCCT | ACCCC | ACCCG | ACCGA | ACCGT | ACCGC | ACCGG | | ACG | ACGAA | ACGAT | ACGAC | ACGAG | ACGTA | ACGTT | ACGTC | ACGTG | ACGCA | ACGCT | ACGCC | ACGCG | ACGGA | ACGGT | ACGGC | ACGGG | | AGA | AGAAA | AGAAT | AGAAC | AGAAG | AGATA | AGATT | AGATC | AGATG | AGACA | AGACT | AGACC | AGACG | AGAGA | AGAGT | AGAGC | AGAGG | | AGT | AGTAA | AGTAT | AGTAC | AGTAG | AGTTA | AGTTT | AGTTC | AGTTG | AGTCA | AGTCT | AGTCC | AGTCG | AGTGA | AGTGT | AGTGC | AGTGG | | AGC | AGCAA | AGCAT | AGCAC | AGCAG | AGCTA | AGCTT | AGCTC | AGCTG | AGCCA | AGCCT | AGCCC | AGCCG | AGCGA | AGCGT | AGCGC | AGCGG | | AGG | AGGAA | AGGAT | AGGAC | AGGAG | AGGTA | AGGTT | AGGTC | AGGTG | AGGCA | AGGCT | AGGCC | AGGCG | AGGGA | AGGGT | AGGGC | AGGGG | | TAA | TAAAA | TAAAT | TAAAC | TAAAG | TAATA | TAATT | TAATC | TAATG | TAACA | TAACT | TAACC | TAACG | TAAGA | TAAGT | TAAGC | TAAGG | | TAT | TATAA | TATAT | TATAC | TATAG | TATTA | TATTT | TATTC | TATTG | TATCA | TATCT | TATCC | TATCG | TATGA | TATGT | TATGC | TATGG | | TAC | TACAA | TACAT | TACAC | TACAG | TACTA | TACTT | TACTC | TACTG | TACCA | TACCT | TACCC | TACCG | TACGA | TACGT | TACGC | TACGG | | TAG | TAGAA | TAGAT | TAGAC | TAGAG | TAGTA | TAGTT | TAGTC | TAGTG | TAGCA | TAGCT | TAGCC | TAGCG | TAGGA | TAGGT | TAGGC | TAGGG | | TTA | TTAAA | TTAAT | TTAAC | TTAAG | TTATA | TTATT | TTATC | TTATG | TTACA | TTACT | TTACC | TTACG | TTAGA | TTAGT | TTAGC | TTAGG | | TTT | TTTAA | TTTAT | TTTAC | TTTAG | TTTTA | TTTTT | TTTTC | TTTTG | TTTCA | TTTCT | TTTCC | TTTCG | TTTGA | TTTGT | TTTGC | TTTGG | | TTC | TTCAA | TTCAT | TTCAC | TTCAG | TTCTA | TTCTT | TTCTC | TTCTG | TTCCA | TTCCT | TTCCC | TTCCG | TTCGA | TTCGT | TTCGC | TTCGG | | TTG | TTGAA | TTGAT | TTGAC | TTGAG | TTGTA | TTGTT | TTGTC | TTGTG | TTGCA | TTGCT | TTGCC | TTGCG | TTGGA | TTGGT | TTGGC | TTGGG | | TCA | TCAAA | TCAAT | TCAAC | TCAAG | TCATA | TCATT | TCATC | TCATG | TCACA | TCACT | TCACC | TCACG | TCAGA | TCAGT | TCAGC | TCAGG | | TCT | TCTAA | TCTAT | TCTAC | TCTAG | TCTTA | TCTTT | TCTTC | TCTTG | TCTCA | TCTCT | TCTCC | TCTCG | TCTGA | TCTGT | TCTGC | TCTGG | | TCC | TCCAA | TCCAT | TCCAC | TCCAG | TCCTA | TCCTT | TCCTC | TCCTG | TCCCA | TCCCT | TCCCC | TCCCG | TCCGA | TCCGT | TCCGC | TCCGG | | TCG | TCGAA | TCGAT | TCGAC | TCGAG | TCGTA | TCGTT | TCGTC | TCGTG | TCGCA | TCGCT | TCGCC | TCGCG | TCGGA | TCGGT | TCGGC | TCGGG | | TGA | TGAAA | TGAAT | TGAAC | TGAAG | TGATA | TGATT | TGATC | TGATG | TGACA | TGACT | TGACC | TGACG | TGAGA | TGAGT | TGAGC | TGAGG | | TGT | TGTAA | TGTAT | TGTAC | TGTAG | TGTTA | TGTTT | TGTTC | TGTTG | TGTCA | TGTCT | TGTCC | TGTCG | TGTGA | TGTGT | TGTGC | TGTGG | | TGC | TGCAA | TGCAT | TGCAC | TGCAG | TGCTA | TGCTT | TGCTC | TGCTG | TGCCA | TGCCT | TGCCC | TGCCG | TGCGA | TGCGT | TGCGC | TGCGG | | TGG | TGGAA | TGGAT | TGGAC | TGGAG | TGGTA | TGGTT | TGGTC | TGGTG | TGGCA | TGGCT | TGGCC | TGGCG | TGGGA | TGGGT | TGGGC | TGGGG | | CAA | CAAAA | CAAAT | CAAAC | CAAAG | CAATA | CAATT | CAATC | CAATG | CAACA | CAACT | CAACC | CAACG | CAAGA | CAAGT | CAAGC | CAAGG | | CAT | CATAA | CATAT | CATAC | CATAG | CATTA | CATTT | CATTC | CATTG | CATCA | CATCT | CATCC | CATCG | CATGA | CATGT | CATGC | CATGG | | CAC | CACAA | CACAT | CACAC | CACAG | CACTA | CACTT | CACTC | CACTG | CACCA | CACCT | CACCC | CACCG | CACGA | CACGT | CACGC | CACGG | | CAG | CAGAA | CAGAT | CAGAC | CAGAG | CAGTA | CAGTT | CAGTC | CAGTG | CAGCA | CAGCT | CAGCC | CAGCG | CAGGA | CAGGT | CAGGC | CAGGG | | CTA | CTAAA | CTAAT | CTAAC | CTAAG | CTATA | CTATT | CTATC | CTATG | CTACA | CTACT | CTACC | CTACG | CTAGA | CTAGT | CTAGC | CTAGG | | CTT | CTTAA | CTTAT | CTTAC | CTTAG | CTTTA | CTTTT | CTTTC | CTTTG | CTTCA | CTTCT | CTTCC | CTTCG | CTTGA | CTTGT | CTTGC | CTTGG | | CTC | CTCAA | CTCAT | CTCAC | CTCAG | CTCTA | CTCTT | CTCTC | CTCTG | CTCCA | CTCCT | CTCCC | CTCCG | CTCGA | CTCGT | CTCGC | CTCGG | | CTG | CTGAA | CTGAT | CTGAC | CTGAG | CTGTA | CTGTT | CTGTC | CTGTG | CTGCA | CTGCT | CTGCC | CTGCG | CTGGA | CTGGT | CTGGC | CTGGG | | CCA | CCAAA | CCAAT | CCAAC | CCAAG | CCATA | CCATT | CCATC | CCATG | CCACA | CCACT | CCACC | CCACG | CCAGA | CCAGT | CCAGC | CCAGG | | CCT | CCTAA | CCTAT | CCTAC | CCTAG | CCTTA | CCTTT | CCTTC | CCTTG | CCTCA | CCTCT | CCTCC | CCTCG | CCTGA | CCTGT | CCTGC | CCTGG | | CCC | CCCAA | CCCAT | CCCAC | CCCAG | CCCTA | CCCTT | CCCTC | CCCTG | CCCCA | CCCCT | CCCCC | CCCCG | CCCGA | CCCGT | CCCGC | CCCGG | | CCG | CCGAA | CCGAT | CCGAC | CCGAG | CCGTA | CCGTT | CCGTC | CCGTG | CCGCA | CCGCT | CCGCC | CCGCG | CCGGA | CCGGT | CCGGC | CCGGG | | CGA | CGAAA | CGAAT | CGAAC | CGAAG | CGATA | CGATT | CGATC | CGATG | CGACA | CGACT | CGACC | CGACG | CGAGA | CGAGT | CGAGC | CGAGG | | CGT | CGTAA | CGTAT | CGTAC | CGTAG | CGTTA | CGTTT | CGTTC | CGTTG | CGTCA | CGTCT | CGTCC | CGTCG | CGTGA | CGTGT | CGTGC | CGTGG | | CGC | CGCAA | CGCAT | CGCAC | CGCAG | CGCTA | CGCTT | CGCTC | CGCTG | CGCCA | CGCCT | CGCCC | CGCCG | CGCGA | CGCGT | CGCGC | CGCGG | | CGG | CGGAA | CGGAT | CGGAC | CGGAG | CGGTA | CGGTT | CGGTC | CGGTG | CGGCA | CGGCT | CGGCC | CGGCG | CGGGA | CGGGT | CGGGC | CGGGG | | GAA | GAAAA | GAAAT | GAAAC | GAAAG | GAATA | GAATT | GAATC | GAATG | GAACA | GAACT | GAACC | GAACG | GAAGA | GAAGT | GAAGC | GAAGG | | GAT | GATAA | GATAT | GATAC | GATAG | GATTA | GATTT | GATTC | GATTG | GATCA | GATCT | GATCC | GATCG | GATGA | GATGT | GATGC | GATGG | | GAC | GACAA | GACAT | GACAC | GACAG | GACTA | GACTT | GACTC | GACTG | GACCA | GACCT | GACCC | GACCG | GACGA | GACGT | GACGC | GACGG | | GAG | GAGAA | GAGAT | GAGAC | GAGAG | GAGTA | GAGTT | GAGTC | GAGTG | GAGCA | GAGCT | GAGCC | GAGCG | GAGGA | GAGGT | GAGGC | GAGGG | | GTA | GTAAA | GTAAT | GTAAC | GTAAG | GTATA | GTATT | GTATC | GTATG | GTACA | GTACT | GTACC | GTACG | GTAGA | GTAGT | GTAGC | GTAGG | | GTT | GTTAA | GTTAT | GTTAC | GTTAG | GTTTA | GTTTT | GTTTC | GTTTG | GTTCA | GTTCT | GTTCC | GTTCG | GTTGA | GTTGT | GTTGC | GTTGG | | GTC | GTCAA | GTCAT | GTCAC | GTCAG | GTCTA | GTCTT | GTCTC | GTCTG | GTCCA | GTCCT | GTCCC | GTCCG | GTCGA | GTCGT | GTCGC | GTCGG | | GTG | GTGAA | GTGAT | GTGAC | GTGAG | GTGTA | GTGTT | GTGTC | GTGTG | GTGCA | GTGCT | GTGCC | GTGCG | GTGGA | GTGGT | GTGGC | GTGGG | | GCA | GCAAA | GCAAT | GCAAC | GCAAG | GCATA | GCATT | GCATC | GCATG | GCACA | GCACT | GCACC | GCACG | GCAGA | GCAGT | GCAGC | GCAGG | | GCT | GCTAA | GCTAT | GCTAC | GCTAG | GCTTA | GCTTT | GCTTC | GCTTG | GCTCA | GCTCT | GCTCC | GCTCG | GCTGA | GCTGT | GCTGC | GCTGG | | GCC | GCCAA | GCCAT | GCCAC | GCCAG | GCCTA | GCCTT | GCCTC | GCCTG | GCCCA | GCCCT | GCCCC | GCCCG | GCCGA | GCCGT | GCCGC | GCCGG | | GCG | GCGAA | GCGAT | GCGAC | GCGAG | GCGTA | GCGTT | GCGTC | GCGTG | GCGCA | GCGCT | GCGCC | GCGCG | GCGGA | GCGGT | GCGGC | GCGGG | | GGA | GGAAA | GGAAT | GGAAC | GGAAG | GGATA | GGATT | GGATC | GGATG | GGACA | GGACT | GGACC | GGACG | GGAGA | GGAGT | GGAGC | GGAGG | | GGT | GGTAA | GGTAT | GGTAC | GGTAG | GGTTA | GGTTT | GGTTC | GGTTG | GGTCA | GGTCT | GGTCC | GGTCG | GGTGA | GGTGT | GGTGC | GGTGG | | GGC | GGCAA | GGCAT | GGCAC | GGCAG | GGCTA | GGCTT | GGCTC | GGCTG | GGCCA | GGCCT | GGCCC | GGCCG | GGCGA | GGCGT | GGCGC | GGCGG | | GGG | GGGAA | GGGAT | GGGAC | GGGAG | GGGTA | GGGTT | GGGTC | GGGTG | GGGCA | GGGCT | GGGCC | GGGCG | GGGGA | GGGGT | GGGGC | GGGGG | Before filtering: read1: overrepresented sequences Sampling rate: 1 / 20 | | | | | --- | --- | --- | | overrepresented sequence | count (% of bases) | distribution: cycle 1 ~ cycle 150 | | AAAAAAAAAA | 238423 (0.268872%) | | | ACACATCTCCGAGCCCACGAGACGCTACTCTATCTCGTAT | 16 (0.000072%) | | | ACATCTCCGAGCCCACGAGACGCTACTCTATCTCGTATGC | 16 (0.000072%) | | | ACGAGACGCTACTCTATCTCGTATGCCGTCTTCTGCTTGA | 17 (0.000077%) | | | AGCCCACGAGACGCTACTCTATCTCGTATGCCGTCTTCTG | 28 (0.000126%) | | | ATACACATCTCCGAGCCCACGAGACGCTACTCTATCTCGT | 37 (0.000167%) | | | ATCTCCGAGCCCACGAGACGCTACTCTATCTCGTATGCCG | 17 (0.000077%) | | | ATGATCGATC | 389998 (0.439805%) | | | CACATCTCCGAGCCCACGAGACGCTACTCTATCTCGTATG | 20 (0.000090%) | | | CACGAGACGCTACTCTATCTCGTATGCCGTCTTCTGCTTG | 105 (0.000474%) | | | CATCTCCGAGCCCACGAGACGCTACTCTATCTCGTATGCC | 29 (0.000131%) | | | CCACGAGACGCTACTCTATCTCGTATGCCGTCTTCTGCTT | 25 (0.000113%) | | | CCCACGAGACGCTACTCTATCTCGTATGCCGTCTTCTGCT | 35 (0.000158%) | | | CCGAGCCCACGAGACGCTACTCTATCTCGTATGCCGTCTT | 75 (0.000338%) | | | CCTGTCTCTTATACACATCTCCGAGCCCACGAGACGCTAC | 21328 (0.096207%) | | | CGAGCCCACGAGACGCTACTCTATCTCGTATGCCGTCTTC | 54 (0.000244%) | | | CTCCGAGCCCACGAGACGCTACTCTATCTCGTATGCCGTC | 19 (0.000086%) | | | CTCTTATACACATCTCCGAGCCCACGAGACGCTACTCTAT | 192 (0.000866%) | | | CTGTCTCTTATACACATCTCCGAGCCCACGAGACGCTACT | 7189 (0.032428%) | | | CTTATACACATCTCCGAGCCCACGAGACGCTACTCTATCT | 76 (0.000343%) | | | GAGCCCACGAGACGCTACTCTATCTCGTATGCCGTCTTCT | 32 (0.000144%) | | | GATCGATCAT | 203059 (0.228992%) | | | GCCCACGAGACGCTACTCTATCTCGTATGCCGTCTTCTGC | 33 (0.000149%) | | | GCTGTCTCTTATACACATCTCCGAGCCCACGAGACGCTAC | 11006 (0.049646%) | | | GTCTCTTATACACATCTCCGAGCCCACGAGACGCTACTCT | 102 (0.000460%) | | | TACACATCTCCGAGCCCACGAGACGCTACTCTATCTCGTA | 21 (0.000095%) | | | TATACACATCTCCGAGCCCACGAGACGCTACTCTATCTCG | 33 (0.000149%) | | | TCCGAGCCCACGAGACGCTACTCTATCTCGTATGCCGTCT | 45 (0.000203%) | | | TCTCCGAGCCCACGAGACGCTACTCTATCTCGTATGCCGT | 26 (0.000117%) | | | TCTCTTATACACATCTCCGAGCCCACGAGACGCTACTCTA | 26 (0.000117%) | | | TCTGTCTCTTATACACATCTCCGAGCCCACGAGACGCTAC | 17110 (0.077181%) | | | TCTTATACACATCTCCGAGCCCACGAGACGCTACTCTATC | 33 (0.000149%) | | | TGATCGATCA | 228596 (0.257790%) | | | TGTCTCTTATACACATCTCCGAGCCCACGAGACGCTACTC | 102 (0.000460%) | | | TTATACACATCTCCGAGCCCACGAGACGCTACTCTATCTC | 29 (0.000131%) | | | TTTTTTTTTT | 243874 (0.275019%) | | Before filtering: read2: quality Value of each position will be shown on mouse over. Before filtering: read2: base contents Value of each position will be shown on mouse over. Before filtering: read2: KMER counting Darker background means larger counts. The count will be shown on mouse over. | | | | | | | | | | | | | | | | | | | --- | --- | --- | --- | --- | --- | --- | --- | --- | --- | --- | --- | --- | --- | --- | --- | --- | | | AA | AT | AC | AG | TA | TT | TC | TG | CA | CT | CC | CG | GA | GT | GC | GG | | AAA | AAAAA | AAAAT | AAAAC | AAAAG | AAATA | AAATT | AAATC | AAATG | AAACA | AAACT | AAACC | AAACG | AAAGA | AAAGT | AAAGC | AAAGG | | AAT | AATAA | AATAT | AATAC | AATAG | AATTA | AATTT | AATTC | AATTG | AATCA | AATCT | AATCC | AATCG | AATGA | AATGT | AATGC | AATGG | | AAC | AACAA | AACAT | AACAC | AACAG | AACTA | AACTT | AACTC | AACTG | AACCA | AACCT | AACCC | AACCG | AACGA | AACGT | AACGC | AACGG | | AAG | AAGAA | AAGAT | AAGAC | AAGAG | AAGTA | AAGTT | AAGTC | AAGTG | AAGCA | AAGCT | AAGCC | AAGCG | AAGGA | AAGGT | AAGGC | AAGGG | | ATA | ATAAA | ATAAT | ATAAC | ATAAG | ATATA | ATATT | ATATC | ATATG | ATACA | ATACT | ATACC | ATACG | ATAGA | ATAGT | ATAGC | ATAGG | | ATT | ATTAA | ATTAT | ATTAC | ATTAG | ATTTA | ATTTT | ATTTC | ATTTG | ATTCA | ATTCT | ATTCC | ATTCG | ATTGA | ATTGT | ATTGC | ATTGG | | ATC | ATCAA | ATCAT | ATCAC | ATCAG | ATCTA | ATCTT | ATCTC | ATCTG | ATCCA | ATCCT | ATCCC | ATCCG | ATCGA | ATCGT | ATCGC | ATCGG | | ATG | ATGAA | ATGAT | ATGAC | ATGAG | ATGTA | ATGTT | ATGTC | ATGTG | ATGCA | ATGCT | ATGCC | ATGCG | ATGGA | ATGGT | ATGGC | ATGGG | | ACA | ACAAA | ACAAT | ACAAC | ACAAG | ACATA | ACATT | ACATC | ACATG | ACACA | ACACT | ACACC | ACACG | ACAGA | ACAGT | ACAGC | ACAGG | | ACT | ACTAA | ACTAT | ACTAC | ACTAG | ACTTA | ACTTT | ACTTC | ACTTG | ACTCA | ACTCT | ACTCC | ACTCG | ACTGA | ACTGT | ACTGC | ACTGG | | ACC | ACCAA | ACCAT | ACCAC | ACCAG | ACCTA | ACCTT | ACCTC | ACCTG | ACCCA | ACCCT | ACCCC | ACCCG | ACCGA | ACCGT | ACCGC | ACCGG | | ACG | ACGAA | ACGAT | ACGAC | ACGAG | ACGTA | ACGTT | ACGTC | ACGTG | ACGCA | ACGCT | ACGCC | ACGCG | ACGGA | ACGGT | ACGGC | ACGGG | | AGA | AGAAA | AGAAT | AGAAC | AGAAG | AGATA | AGATT | AGATC | AGATG | AGACA | AGACT | AGACC | AGACG | AGAGA | AGAGT | AGAGC | AGAGG | | AGT | AGTAA | AGTAT | AGTAC | AGTAG | AGTTA | AGTTT | AGTTC | AGTTG | AGTCA | AGTCT | AGTCC | AGTCG | AGTGA | AGTGT | AGTGC | AGTGG | | AGC | AGCAA | AGCAT | AGCAC | AGCAG | AGCTA | AGCTT | AGCTC | AGCTG | AGCCA | AGCCT | AGCCC | AGCCG | AGCGA | AGCGT | AGCGC | AGCGG | | AGG | AGGAA | AGGAT | AGGAC | AGGAG | AGGTA | AGGTT | AGGTC | AGGTG | AGGCA | AGGCT | AGGCC | AGGCG | AGGGA | AGGGT | AGGGC | AGGGG | | TAA | TAAAA | TAAAT | TAAAC | TAAAG | TAATA | TAATT | TAATC | TAATG | TAACA | TAACT | TAACC | TAACG | TAAGA | TAAGT | TAAGC | TAAGG | | TAT | TATAA | TATAT | TATAC | TATAG | TATTA | TATTT | TATTC | TATTG | TATCA | TATCT | TATCC | TATCG | TATGA | TATGT | TATGC | TATGG | | TAC | TACAA | TACAT | TACAC | TACAG | TACTA | TACTT | TACTC | TACTG | TACCA | TACCT | TACCC | TACCG | TACGA | TACGT | TACGC | TACGG | | TAG | TAGAA | TAGAT | TAGAC | TAGAG | TAGTA | TAGTT | TAGTC | TAGTG | TAGCA | TAGCT | TAGCC | TAGCG | TAGGA | TAGGT | TAGGC | TAGGG | | TTA | TTAAA | TTAAT | TTAAC | TTAAG | TTATA | TTATT | TTATC | TTATG | TTACA | TTACT | TTACC | TTACG | TTAGA | TTAGT | TTAGC | TTAGG | | TTT | TTTAA | TTTAT | TTTAC | TTTAG | TTTTA | TTTTT | TTTTC | TTTTG | TTTCA | TTTCT | TTTCC | TTTCG | TTTGA | TTTGT | TTTGC | TTTGG | | TTC | TTCAA | TTCAT | TTCAC | TTCAG | TTCTA | TTCTT | TTCTC | TTCTG | TTCCA | TTCCT | TTCCC | TTCCG | TTCGA | TTCGT | TTCGC | TTCGG | | TTG | TTGAA | TTGAT | TTGAC | TTGAG | TTGTA | TTGTT | TTGTC | TTGTG | TTGCA | TTGCT | TTGCC | TTGCG | TTGGA | TTGGT | TTGGC | TTGGG | | TCA | TCAAA | TCAAT | TCAAC | TCAAG | TCATA | TCATT | TCATC | TCATG | TCACA | TCACT | TCACC | TCACG | TCAGA | TCAGT | TCAGC | TCAGG | | TCT | TCTAA | TCTAT | TCTAC | TCTAG | TCTTA | TCTTT | TCTTC | TCTTG | TCTCA | TCTCT | TCTCC | TCTCG | TCTGA | TCTGT | TCTGC | TCTGG | | TCC | TCCAA | TCCAT | TCCAC | TCCAG | TCCTA | TCCTT | TCCTC | TCCTG | TCCCA | TCCCT | TCCCC | TCCCG | TCCGA | TCCGT | TCCGC | TCCGG | | TCG | TCGAA | TCGAT | TCGAC | TCGAG | TCGTA | TCGTT | TCGTC | TCGTG | TCGCA | TCGCT | TCGCC | TCGCG | TCGGA | TCGGT | TCGGC | TCGGG | | TGA | TGAAA | TGAAT | TGAAC | TGAAG | TGATA | TGATT | TGATC | TGATG | TGACA | TGACT | TGACC | TGACG | TGAGA | TGAGT | TGAGC | TGAGG | | TGT | TGTAA | TGTAT | TGTAC | TGTAG | TGTTA | TGTTT | TGTTC | TGTTG | TGTCA | TGTCT | TGTCC | TGTCG | TGTGA | TGTGT | TGTGC | TGTGG | | TGC | TGCAA | TGCAT | TGCAC | TGCAG | TGCTA | TGCTT | TGCTC | TGCTG | TGCCA | TGCCT | TGCCC | TGCCG | TGCGA | TGCGT | TGCGC | TGCGG | | TGG | TGGAA | TGGAT | TGGAC | TGGAG | TGGTA | TGGTT | TGGTC | TGGTG | TGGCA | TGGCT | TGGCC | TGGCG | TGGGA | TGGGT | TGGGC | TGGGG | | CAA | CAAAA | CAAAT | CAAAC | CAAAG | CAATA | CAATT | CAATC | CAATG | CAACA | CAACT | CAACC | CAACG | CAAGA | CAAGT | CAAGC | CAAGG | | CAT | CATAA | CATAT | CATAC | CATAG | CATTA | CATTT | CATTC | CATTG | CATCA | CATCT | CATCC | CATCG | CATGA | CATGT | CATGC | CATGG | | CAC | CACAA | CACAT | CACAC | CACAG | CACTA | CACTT | CACTC | CACTG | CACCA | CACCT | CACCC | CACCG | CACGA | CACGT | CACGC | CACGG | | CAG | CAGAA | CAGAT | CAGAC | CAGAG | CAGTA | CAGTT | CAGTC | CAGTG | CAGCA | CAGCT | CAGCC | CAGCG | CAGGA | CAGGT | CAGGC | CAGGG | | CTA | CTAAA | CTAAT | CTAAC | CTAAG | CTATA | CTATT | CTATC | CTATG | CTACA | CTACT | CTACC | CTACG | CTAGA | CTAGT | CTAGC | CTAGG | | CTT | CTTAA | CTTAT | CTTAC | CTTAG | CTTTA | CTTTT | CTTTC | CTTTG | CTTCA | CTTCT | CTTCC | CTTCG | CTTGA | CTTGT | CTTGC | CTTGG | | CTC | CTCAA | CTCAT | CTCAC | CTCAG | CTCTA | CTCTT | CTCTC | CTCTG | CTCCA | CTCCT | CTCCC | CTCCG | CTCGA | CTCGT | CTCGC | CTCGG | | CTG | CTGAA | CTGAT | CTGAC | CTGAG | CTGTA | CTGTT | CTGTC | CTGTG | CTGCA | CTGCT | CTGCC | CTGCG | CTGGA | CTGGT | CTGGC | CTGGG | | CCA | CCAAA | CCAAT | CCAAC | CCAAG | CCATA | CCATT | CCATC | CCATG | CCACA | CCACT | CCACC | CCACG | CCAGA | CCAGT | CCAGC | CCAGG | | CCT | CCTAA | CCTAT | CCTAC | CCTAG | CCTTA | CCTTT | CCTTC | CCTTG | CCTCA | CCTCT | CCTCC | CCTCG | CCTGA | CCTGT | CCTGC | CCTGG | | CCC | CCCAA | CCCAT | CCCAC | CCCAG | CCCTA | CCCTT | CCCTC | CCCTG | CCCCA | CCCCT | CCCCC | CCCCG | CCCGA | CCCGT | CCCGC | CCCGG | | CCG | CCGAA | CCGAT | CCGAC | CCGAG | CCGTA | CCGTT | CCGTC | CCGTG | CCGCA | CCGCT | CCGCC | CCGCG | CCGGA | CCGGT | CCGGC | CCGGG | | CGA | CGAAA | CGAAT | CGAAC | CGAAG | CGATA | CGATT | CGATC | CGATG | CGACA | CGACT | CGACC | CGACG | CGAGA | CGAGT | CGAGC | CGAGG | | CGT | CGTAA | CGTAT | CGTAC | CGTAG | CGTTA | CGTTT | CGTTC | CGTTG | CGTCA | CGTCT | CGTCC | CGTCG | CGTGA | CGTGT | CGTGC | CGTGG | | CGC | CGCAA | CGCAT | CGCAC | CGCAG | CGCTA | CGCTT | CGCTC | CGCTG | CGCCA | CGCCT | CGCCC | CGCCG | CGCGA | CGCGT | CGCGC | CGCGG | | CGG | CGGAA | CGGAT | CGGAC | CGGAG | CGGTA | CGGTT | CGGTC | CGGTG | CGGCA | CGGCT | CGGCC | CGGCG | CGGGA | CGGGT | CGGGC | CGGGG | | GAA | GAAAA | GAAAT | GAAAC | GAAAG | GAATA | GAATT | GAATC | GAATG | GAACA | GAACT | GAACC | GAACG | GAAGA | GAAGT | GAAGC | GAAGG | | GAT | GATAA | GATAT | GATAC | GATAG | GATTA | GATTT | GATTC | GATTG | GATCA | GATCT | GATCC | GATCG | GATGA | GATGT | GATGC | GATGG | | GAC | GACAA | GACAT | GACAC | GACAG | GACTA | GACTT | GACTC | GACTG | GACCA | GACCT | GACCC | GACCG | GACGA | GACGT | GACGC | GACGG | | GAG | GAGAA | GAGAT | GAGAC | GAGAG | GAGTA | GAGTT | GAGTC | GAGTG | GAGCA | GAGCT | GAGCC | GAGCG | GAGGA | GAGGT | GAGGC | GAGGG | | GTA | GTAAA | GTAAT | GTAAC | GTAAG | GTATA | GTATT | GTATC | GTATG | GTACA | GTACT | GTACC | GTACG | GTAGA | GTAGT | GTAGC | GTAGG | | GTT | GTTAA | GTTAT | GTTAC | GTTAG | GTTTA | GTTTT | GTTTC | GTTTG | GTTCA | GTTCT | GTTCC | GTTCG | GTTGA | GTTGT | GTTGC | GTTGG | | GTC | GTCAA | GTCAT | GTCAC | GTCAG | GTCTA | GTCTT | GTCTC | GTCTG | GTCCA | GTCCT | GTCCC | GTCCG | GTCGA | GTCGT | GTCGC | GTCGG | | GTG | GTGAA | GTGAT | GTGAC | GTGAG | GTGTA | GTGTT | GTGTC | GTGTG | GTGCA | GTGCT | GTGCC | GTGCG | GTGGA | GTGGT | GTGGC | GTGGG | | GCA | GCAAA | GCAAT | GCAAC | GCAAG | GCATA | GCATT | GCATC | GCATG | GCACA | GCACT | GCACC | GCACG | GCAGA | GCAGT | GCAGC | GCAGG | | GCT | GCTAA | GCTAT | GCTAC | GCTAG | GCTTA | GCTTT | GCTTC | GCTTG | GCTCA | GCTCT | GCTCC | GCTCG | GCTGA | GCTGT | GCTGC | GCTGG | | GCC | GCCAA | GCCAT | GCCAC | GCCAG | GCCTA | GCCTT | GCCTC | GCCTG | GCCCA | GCCCT | GCCCC | GCCCG | GCCGA | GCCGT | GCCGC | GCCGG | | GCG | GCGAA | GCGAT | GCGAC | GCGAG | GCGTA | GCGTT | GCGTC | GCGTG | GCGCA | GCGCT | GCGCC | GCGCG | GCGGA | GCGGT | GCGGC | GCGGG | | GGA | GGAAA | GGAAT | GGAAC | GGAAG | GGATA | GGATT | GGATC | GGATG | GGACA | GGACT | GGACC | GGACG | GGAGA | GGAGT | GGAGC | GGAGG | | GGT | GGTAA | GGTAT | GGTAC | GGTAG | GGTTA | GGTTT | GGTTC | GGTTG | GGTCA | GGTCT | GGTCC | GGTCG | GGTGA | GGTGT | GGTGC | GGTGG | | GGC | GGCAA | GGCAT | GGCAC | GGCAG | GGCTA | GGCTT | GGCTC | GGCTG | GGCCA | GGCCT | GGCCC | GGCCG | GGCGA | GGCGT | GGCGC | GGCGG | | GGG | GGGAA | GGGAT | GGGAC | GGGAG | GGGTA | GGGTT | GGGTC | GGGTG | GGGCA | GGGCT | GGGCC | GGGCG | GGGGA | GGGGT | GGGGC | GGGGG | Before filtering: read2: overrepresented sequences Sampling rate: 1 / 20 | | | | | --- | --- | --- | | overrepresented sequence | count (% of bases) | distribution: cycle 1 ~ cycle 150 | | AAAAAAAAAAAAAAAAAAAA | 8202 (0.018499%) | | | ACACATCTGACGCTGCCGACGACTCGAACAGTGTAGATCT | 28 (0.000126%) | | | ACATCTGACGCTGCCGACGACTCGAACAGTGTAGATCTCG | 19 (0.000086%) | | | ACCTGTCTCTTATACACATCTGACGCTGCCGACGACTCGA | 9977 (0.045005%) | | | ACGACTCGAACAGTGTAGATCTCGGTGGTCGCCGTATCAT | 7 (0.000032%) | | | ACGCTGCCGACGACTCGAACAGTGTAGATCTCGGTGGTCG | 61 (0.000275%) | | | ATACACATCTGACGCTGCCGACGACTCGAACAGTGTAGAT | 21 (0.000095%) | | | ATCTGACGCTGCCGACGACTCGAACAGTGTAGATCTCGGT | 19 (0.000086%) | | | ATCTGTCTCTTATACACATCTGACGCTGCCGACGACTCGA | 10065 (0.045402%) | | | ATGATCGATC | 377615 (0.425840%) | | | CACATCTGACGCTGCCGACGACTCGAACAGTGTAGATCTC | 27 (0.000122%) | | | CATCTGACGCTGCCGACGACTCGAACAGTGTAGATCTCGG | 17 (0.000077%) | | | CCGACGACTCGAACAGTGTAGATCTCGGTGGTCGCCGTAT | 21 (0.000095%) | | | CCTGTCTCTTATACACATCTGACGCTGCCGACGACTCGAA | 11553 (0.052114%) | | | CGACGACTCGAACAGTGTAGATCTCGGTGGTCGCCGTATC | 20 (0.000090%) | | | CGACTCGAACAGTGTAGATCTCGGTGGTCGCCGTATCATT | 20 (0.000090%) | | | CGCTGCCGACGACTCGAACAGTGTAGATCTCGGTGGTCGC | 36 (0.000162%) | | | CTCTTATACACATCTGACGCTGCCGACGACTCGAACAGTG | 81 (0.000365%) | | | CTGACGCTGCCGACGACTCGAACAGTGTAGATCTCGGTGG | 38 (0.000171%) | | | CTGCCGACGACTCGAACAGTGTAGATCTCGGTGGTCGCCG | 37 (0.000167%) | | | CTGTCTCTTATACACATCTGACGCTGCCGACGACTCGAAC | 7262 (0.032758%) | | | CTTATACACATCTGACGCTGCCGACGACTCGAACAGTGTA | 79 (0.000356%) | | | GACGACTCGAACAGTGTAGATCTCGGTGGTCGCCGTATCA | 16 (0.000072%) | | | GACGCTGCCGACGACTCGAACAGTGTAGATCTCGGTGGTC | 52 (0.000235%) | | | GATCGATCAT | 197782 (0.223041%) | | | GCCGACGACTCGAACAGTGTAGATCTCGGTGGTCGCCGTA | 11 (0.000050%) | | | GCTGCCGACGACTCGAACAGTGTAGATCTCGGTGGTCGCC | 43 (0.000194%) | | | GCTGTCTCTTATACACATCTGACGCTGCCGACGACTCGAA | 10903 (0.049182%) | | | GTCTCTTATACACATCTGACGCTGCCGACGACTCGAACAG | 77 (0.000347%) | | | TACACATCTGACGCTGCCGACGACTCGAACAGTGTAGATC | 34 (0.000153%) | | | TATACACATCTGACGCTGCCGACGACTCGAACAGTGTAGA | 26 (0.000117%) | | | TCTCTTATACACATCTGACGCTGCCGACGACTCGAACAGT | 54 (0.000244%) | | | TCTGACGCTGCCGACGACTCGAACAGTGTAGATCTCGGTG | 20 (0.000090%) | | | TCTGTCTCTTATACACATCTGACGCTGCCGACGACTCGAA | 7681 (0.034648%) | | | TCTTATACACATCTGACGCTGCCGACGACTCGAACAGTGT | 48 (0.000217%) | | | TGACGCTGCCGACGACTCGAACAGTGTAGATCTCGGTGGT | 49 (0.000221%) | | | TGATCGATCA | 221717 (0.250033%) | | | TGCCGACGACTCGAACAGTGTAGATCTCGGTGGTCGCCGT | 23 (0.000104%) | | | TGTCTCTTATACACATCTGACGCTGCCGACGACTCGAACA | 88 (0.000397%) | | | TTATACACATCTGACGCTGCCGACGACTCGAACAGTGTAG | 40 (0.000180%) | | | TTTTTTTTTT | 226123 (0.255001%) | | After filtering After filtering: read1: quality Value of each position will be shown on mouse over. After filtering: read1: base contents Value of each position will be shown on mouse over. After filtering: read1: KMER counting Darker background means larger counts. The count will be shown on mouse over. | | | | | | | | | | | | | | | | | | | --- | --- | --- | --- | --- | --- | --- | --- | --- | --- | --- | --- | --- | --- | --- | --- | --- | | | AA | AT | AC | AG | TA | TT | TC | TG | CA | CT | CC | CG | GA | GT | GC | GG | | AAA | AAAAA | AAAAT | AAAAC | AAAAG | AAATA | AAATT | AAATC | AAATG | AAACA | AAACT | AAACC | AAACG | AAAGA | AAAGT | AAAGC | AAAGG | | AAT | AATAA | AATAT | AATAC | AATAG | AATTA | AATTT | AATTC | AATTG | AATCA | AATCT | AATCC | AATCG | AATGA | AATGT | AATGC | AATGG | | AAC | AACAA | AACAT | AACAC | AACAG | AACTA | AACTT | AACTC | AACTG | AACCA | AACCT | AACCC | AACCG | AACGA | AACGT | AACGC | AACGG | | AAG | AAGAA | AAGAT | AAGAC | AAGAG | AAGTA | AAGTT | AAGTC | AAGTG | AAGCA | AAGCT | AAGCC | AAGCG | AAGGA | AAGGT | AAGGC | AAGGG | | ATA | ATAAA | ATAAT | ATAAC | ATAAG | ATATA | ATATT | ATATC | ATATG | ATACA | ATACT | ATACC | ATACG | ATAGA | ATAGT | ATAGC | ATAGG | | ATT | ATTAA | ATTAT | ATTAC | ATTAG | ATTTA | ATTTT | ATTTC | ATTTG | ATTCA | ATTCT | ATTCC | ATTCG | ATTGA | ATTGT | ATTGC | ATTGG | | ATC | ATCAA | ATCAT | ATCAC | ATCAG | ATCTA | ATCTT | ATCTC | ATCTG | ATCCA | ATCCT | ATCCC | ATCCG | ATCGA | ATCGT | ATCGC | ATCGG | | ATG | ATGAA | ATGAT | ATGAC | ATGAG | ATGTA | ATGTT | ATGTC | ATGTG | ATGCA | ATGCT | ATGCC | ATGCG | ATGGA | ATGGT | ATGGC | ATGGG | | ACA | ACAAA | ACAAT | ACAAC | ACAAG | ACATA | ACATT | ACATC | ACATG | ACACA | ACACT | ACACC | ACACG | ACAGA | ACAGT | ACAGC | ACAGG | | ACT | ACTAA | ACTAT | ACTAC | ACTAG | ACTTA | ACTTT | ACTTC | ACTTG | ACTCA | ACTCT | ACTCC | ACTCG | ACTGA | ACTGT | ACTGC | ACTGG | | ACC | ACCAA | ACCAT | ACCAC | ACCAG | ACCTA | ACCTT | ACCTC | ACCTG | ACCCA | ACCCT | ACCCC | ACCCG | ACCGA | ACCGT | ACCGC | ACCGG | | ACG | ACGAA | ACGAT | ACGAC | ACGAG | ACGTA | ACGTT | ACGTC | ACGTG | ACGCA | ACGCT | ACGCC | ACGCG | ACGGA | ACGGT | ACGGC | ACGGG | | AGA | AGAAA | AGAAT | AGAAC | AGAAG | AGATA | AGATT | AGATC | AGATG | AGACA | AGACT | AGACC | AGACG | AGAGA | AGAGT | AGAGC | AGAGG | | AGT | AGTAA | AGTAT | AGTAC | AGTAG | AGTTA | AGTTT | AGTTC | AGTTG | AGTCA | AGTCT | AGTCC | AGTCG | AGTGA | AGTGT | AGTGC | AGTGG | | AGC | AGCAA | AGCAT | AGCAC | AGCAG | AGCTA | AGCTT | AGCTC | AGCTG | AGCCA | AGCCT | AGCCC | AGCCG | AGCGA | AGCGT | AGCGC | AGCGG | | AGG | AGGAA | AGGAT | AGGAC | AGGAG | AGGTA | AGGTT | AGGTC | AGGTG | AGGCA | AGGCT | AGGCC | AGGCG | AGGGA | AGGGT | AGGGC | AGGGG | | TAA | TAAAA | TAAAT | TAAAC | TAAAG | TAATA | TAATT | TAATC | TAATG | TAACA | TAACT | TAACC | TAACG | TAAGA | TAAGT | TAAGC | TAAGG | | TAT | TATAA | TATAT | TATAC | TATAG | TATTA | TATTT | TATTC | TATTG | TATCA | TATCT | TATCC | TATCG | TATGA | TATGT | TATGC | TATGG | | TAC | TACAA | TACAT | TACAC | TACAG | TACTA | TACTT | TACTC | TACTG | TACCA | TACCT | TACCC | TACCG | TACGA | TACGT | TACGC | TACGG | | TAG | TAGAA | TAGAT | TAGAC | TAGAG | TAGTA | TAGTT | TAGTC | TAGTG | TAGCA | TAGCT | TAGCC | TAGCG | TAGGA | TAGGT | TAGGC | TAGGG | | TTA | TTAAA | TTAAT | TTAAC | TTAAG | TTATA | TTATT | TTATC | TTATG | TTACA | TTACT | TTACC | TTACG | TTAGA | TTAGT | TTAGC | TTAGG | | TTT | TTTAA | TTTAT | TTTAC | TTTAG | TTTTA | TTTTT | TTTTC | TTTTG | TTTCA | TTTCT | TTTCC | TTTCG | TTTGA | TTTGT | TTTGC | TTTGG | | TTC | TTCAA | TTCAT | TTCAC | TTCAG | TTCTA | TTCTT | TTCTC | TTCTG | TTCCA | TTCCT | TTCCC | TTCCG | TTCGA | TTCGT | TTCGC | TTCGG | | TTG | TTGAA | TTGAT | TTGAC | TTGAG | TTGTA | TTGTT | TTGTC | TTGTG | TTGCA | TTGCT | TTGCC | TTGCG | TTGGA | TTGGT | TTGGC | TTGGG | | TCA | TCAAA | TCAAT | TCAAC | TCAAG | TCATA | TCATT | TCATC | TCATG | TCACA | TCACT | TCACC | TCACG | TCAGA | TCAGT | TCAGC | TCAGG | | TCT | TCTAA | TCTAT | TCTAC | TCTAG | TCTTA | TCTTT | TCTTC | TCTTG | TCTCA | TCTCT | TCTCC | TCTCG | TCTGA | TCTGT | TCTGC | TCTGG | | TCC | TCCAA | TCCAT | TCCAC | TCCAG | TCCTA | TCCTT | TCCTC | TCCTG | TCCCA | TCCCT | TCCCC | TCCCG | TCCGA | TCCGT | TCCGC | TCCGG | | TCG | TCGAA | TCGAT | TCGAC | TCGAG | TCGTA | TCGTT | TCGTC | TCGTG | TCGCA | TCGCT | TCGCC | TCGCG | TCGGA | TCGGT | TCGGC | TCGGG | | TGA | TGAAA | TGAAT | TGAAC | TGAAG | TGATA | TGATT | TGATC | TGATG | TGACA | TGACT | TGACC | TGACG | TGAGA | TGAGT | TGAGC | TGAGG | | TGT | TGTAA | TGTAT | TGTAC | TGTAG | TGTTA | TGTTT | TGTTC | TGTTG | TGTCA | TGTCT | TGTCC | TGTCG | TGTGA | TGTGT | TGTGC | TGTGG | | TGC | TGCAA | TGCAT | TGCAC | TGCAG | TGCTA | TGCTT | TGCTC | TGCTG | TGCCA | TGCCT | TGCCC | TGCCG | TGCGA | TGCGT | TGCGC | TGCGG | | TGG | TGGAA | TGGAT | TGGAC | TGGAG | TGGTA | TGGTT | TGGTC | TGGTG | TGGCA | TGGCT | TGGCC | TGGCG | TGGGA | TGGGT | TGGGC | TGGGG | | CAA | CAAAA | CAAAT | CAAAC | CAAAG | CAATA | CAATT | CAATC | CAATG | CAACA | CAACT | CAACC | CAACG | CAAGA | CAAGT | CAAGC | CAAGG | | CAT | CATAA | CATAT | CATAC | CATAG | CATTA | CATTT | CATTC | CATTG | CATCA | CATCT | CATCC | CATCG | CATGA | CATGT | CATGC | CATGG | | CAC | CACAA | CACAT | CACAC | CACAG | CACTA | CACTT | CACTC | CACTG | CACCA | CACCT | CACCC | CACCG | CACGA | CACGT | CACGC | CACGG | | CAG | CAGAA | CAGAT | CAGAC | CAGAG | CAGTA | CAGTT | CAGTC | CAGTG | CAGCA | CAGCT | CAGCC | CAGCG | CAGGA | CAGGT | CAGGC | CAGGG | | CTA | CTAAA | CTAAT | CTAAC | CTAAG | CTATA | CTATT | CTATC | CTATG | CTACA | CTACT | CTACC | CTACG | CTAGA | CTAGT | CTAGC | CTAGG | | CTT | CTTAA | CTTAT | CTTAC | CTTAG | CTTTA | CTTTT | CTTTC | CTTTG | CTTCA | CTTCT | CTTCC | CTTCG | CTTGA | CTTGT | CTTGC | CTTGG | | CTC | CTCAA | CTCAT | CTCAC | CTCAG | CTCTA | CTCTT | CTCTC | CTCTG | CTCCA | CTCCT | CTCCC | CTCCG | CTCGA | CTCGT | CTCGC | CTCGG | | CTG | CTGAA | CTGAT | CTGAC | CTGAG | CTGTA | CTGTT | CTGTC | CTGTG | CTGCA | CTGCT | CTGCC | CTGCG | CTGGA | CTGGT | CTGGC | CTGGG | | CCA | CCAAA | CCAAT | CCAAC | CCAAG | CCATA | CCATT | CCATC | CCATG | CCACA | CCACT | CCACC | CCACG | CCAGA | CCAGT | CCAGC | CCAGG | | CCT | CCTAA | CCTAT | CCTAC | CCTAG | CCTTA | CCTTT | CCTTC | CCTTG | CCTCA | CCTCT | CCTCC | CCTCG | CCTGA | CCTGT | CCTGC | CCTGG | | CCC | CCCAA | CCCAT | CCCAC | CCCAG | CCCTA | CCCTT | CCCTC | CCCTG | CCCCA | CCCCT | CCCCC | CCCCG | CCCGA | CCCGT | CCCGC | CCCGG | | CCG | CCGAA | CCGAT | CCGAC | CCGAG | CCGTA | CCGTT | CCGTC | CCGTG | CCGCA | CCGCT | CCGCC | CCGCG | CCGGA | CCGGT | CCGGC | CCGGG | | CGA | CGAAA | CGAAT | CGAAC | CGAAG | CGATA | CGATT | CGATC | CGATG | CGACA | CGACT | CGACC | CGACG | CGAGA | CGAGT | CGAGC | CGAGG | | CGT | CGTAA | CGTAT | CGTAC | CGTAG | CGTTA | CGTTT | CGTTC | CGTTG | CGTCA | CGTCT | CGTCC | CGTCG | CGTGA | CGTGT | CGTGC | CGTGG | | CGC | CGCAA | CGCAT | CGCAC | CGCAG | CGCTA | CGCTT | CGCTC | CGCTG | CGCCA | CGCCT | CGCCC | CGCCG | CGCGA | CGCGT | CGCGC | CGCGG | | CGG | CGGAA | CGGAT | CGGAC | CGGAG | CGGTA | CGGTT | CGGTC | CGGTG | CGGCA | CGGCT | CGGCC | CGGCG | CGGGA | CGGGT | CGGGC | CGGGG | | GAA | GAAAA | GAAAT | GAAAC | GAAAG | GAATA | GAATT | GAATC | GAATG | GAACA | GAACT | GAACC | GAACG | GAAGA | GAAGT | GAAGC | GAAGG | | GAT | GATAA | GATAT | GATAC | GATAG | GATTA | GATTT | GATTC | GATTG | GATCA | GATCT | GATCC | GATCG | GATGA | GATGT | GATGC | GATGG | | GAC | GACAA | GACAT | GACAC | GACAG | GACTA | GACTT | GACTC | GACTG | GACCA | GACCT | GACCC | GACCG | GACGA | GACGT | GACGC | GACGG | | GAG | GAGAA | GAGAT | GAGAC | GAGAG | GAGTA | GAGTT | GAGTC | GAGTG | GAGCA | GAGCT | GAGCC | GAGCG | GAGGA | GAGGT | GAGGC | GAGGG | | GTA | GTAAA | GTAAT | GTAAC | GTAAG | GTATA | GTATT | GTATC | GTATG | GTACA | GTACT | GTACC | GTACG | GTAGA | GTAGT | GTAGC | GTAGG | | GTT | GTTAA | GTTAT | GTTAC | GTTAG | GTTTA | GTTTT | GTTTC | GTTTG | GTTCA | GTTCT | GTTCC | GTTCG | GTTGA | GTTGT | GTTGC | GTTGG | | GTC | GTCAA | GTCAT | GTCAC | GTCAG | GTCTA | GTCTT | GTCTC | GTCTG | GTCCA | GTCCT | GTCCC | GTCCG | GTCGA | GTCGT | GTCGC | GTCGG | | GTG | GTGAA | GTGAT | GTGAC | GTGAG | GTGTA | GTGTT | GTGTC | GTGTG | GTGCA | GTGCT | GTGCC | GTGCG | GTGGA | GTGGT | GTGGC | GTGGG | | GCA | GCAAA | GCAAT | GCAAC | GCAAG | GCATA | GCATT | GCATC | GCATG | GCACA | GCACT | GCACC | GCACG | GCAGA | GCAGT | GCAGC | GCAGG | | GCT | GCTAA | GCTAT | GCTAC | GCTAG | GCTTA | GCTTT | GCTTC | GCTTG | GCTCA | GCTCT | GCTCC | GCTCG | GCTGA | GCTGT | GCTGC | GCTGG | | GCC | GCCAA | GCCAT | GCCAC | GCCAG | GCCTA | GCCTT | GCCTC | GCCTG | GCCCA | GCCCT | GCCCC | GCCCG | GCCGA | GCCGT | GCCGC | GCCGG | | GCG | GCGAA | GCGAT | GCGAC | GCGAG | GCGTA | GCGTT | GCGTC | GCGTG | GCGCA | GCGCT | GCGCC | GCGCG | GCGGA | GCGGT | GCGGC | GCGGG | | GGA | GGAAA | GGAAT | GGAAC | GGAAG | GGATA | GGATT | GGATC | GGATG | GGACA | GGACT | GGACC | GGACG | GGAGA | GGAGT | GGAGC | GGAGG | | GGT | GGTAA | GGTAT | GGTAC | GGTAG | GGTTA | GGTTT | GGTTC | GGTTG | GGTCA | GGTCT | GGTCC | GGTCG | GGTGA | GGTGT | GGTGC | GGTGG | | GGC | GGCAA | GGCAT | GGCAC | GGCAG | GGCTA | GGCTT | GGCTC | GGCTG | GGCCA | GGCCT | GGCCC | GGCCG | GGCGA | GGCGT | GGCGC | GGCGG | | GGG | GGGAA | GGGAT | GGGAC | GGGAG | GGGTA | GGGTT | GGGTC | GGGTG | GGGCA | GGGCT | GGGCC | GGGCG | GGGGA | GGGGT | GGGGC | GGGGG | After filtering: read1: overrepresented sequences Sampling rate: 1 / 20 | | | | | --- | --- | --- | | overrepresented sequence | count (% of bases) | distribution: cycle 1 ~ cycle 150 | | AAAAAAAAAA | 227817 (0.269674%) | | | ATGATCGATC | 381082 (0.451099%) | | | CCTGTCTCTTATACACATCTCCGAGCCCACGAGACGCTAC | 72 (0.000341%) | | | CTGTCTCTTATACACATCTCCGAGCCCACGAGACGCTACT | 35 (0.000166%) | | | GATCGATCAT | 198329 (0.234769%) | | | GCTGTCTCTTATACACATCTCCGAGCCCACGAGACGCTAC | 32 (0.000152%) | | | TCTGTCTCTTATACACATCTCCGAGCCCACGAGACGCTAC | 52 (0.000246%) | | | TGATCGATCA | 224034 (0.265196%) | | | TTTTTTTTTT | 233118 (0.275949%) | | After filtering: read2: quality Value of each position will be shown on mouse over. After filtering: read2: base contents Value of each position will be shown on mouse over. After filtering: read2: KMER counting Darker background means larger counts. The count will be shown on mouse over. | | | | | | | | | | | | | | | | | | | --- | --- | --- | --- | --- | --- | --- | --- | --- | --- | --- | --- | --- | --- | --- | --- | --- | | | AA | AT | AC | AG | TA | TT | TC | TG | CA | CT | CC | CG | GA | GT | GC | GG | | AAA | AAAAA | AAAAT | AAAAC | AAAAG | AAATA | AAATT | AAATC | AAATG | AAACA | AAACT | AAACC | AAACG | AAAGA | AAAGT | AAAGC | AAAGG | | AAT | AATAA | AATAT | AATAC | AATAG | AATTA | AATTT | AATTC | AATTG | AATCA | AATCT | AATCC | AATCG | AATGA | AATGT | AATGC | AATGG | | AAC | AACAA | AACAT | AACAC | AACAG | AACTA | AACTT | AACTC | AACTG | AACCA | AACCT | AACCC | AACCG | AACGA | AACGT | AACGC | AACGG | | AAG | AAGAA | AAGAT | AAGAC | AAGAG | AAGTA | AAGTT | AAGTC | AAGTG | AAGCA | AAGCT | AAGCC | AAGCG | AAGGA | AAGGT | AAGGC | AAGGG | | ATA | ATAAA | ATAAT | ATAAC | ATAAG | ATATA | ATATT | ATATC | ATATG | ATACA | ATACT | ATACC | ATACG | ATAGA | ATAGT | ATAGC | ATAGG | | ATT | ATTAA | ATTAT | ATTAC | ATTAG | ATTTA | ATTTT | ATTTC | ATTTG | ATTCA | ATTCT | ATTCC | ATTCG | ATTGA | ATTGT | ATTGC | ATTGG | | ATC | ATCAA | ATCAT | ATCAC | ATCAG | ATCTA | ATCTT | ATCTC | ATCTG | ATCCA | ATCCT | ATCCC | ATCCG | ATCGA | ATCGT | ATCGC | ATCGG | | ATG | ATGAA | ATGAT | ATGAC | ATGAG | ATGTA | ATGTT | ATGTC | ATGTG | ATGCA | ATGCT | ATGCC | ATGCG | ATGGA | ATGGT | ATGGC | ATGGG | | ACA | ACAAA | ACAAT | ACAAC | ACAAG | ACATA | ACATT | ACATC | ACATG | ACACA | ACACT | ACACC | ACACG | ACAGA | ACAGT | ACAGC | ACAGG | | ACT | ACTAA | ACTAT | ACTAC | ACTAG | ACTTA | ACTTT | ACTTC | ACTTG | ACTCA | ACTCT | ACTCC | ACTCG | ACTGA | ACTGT | ACTGC | ACTGG | | ACC | ACCAA | ACCAT | ACCAC | ACCAG | ACCTA | ACCTT | ACCTC | ACCTG | ACCCA | ACCCT | ACCCC | ACCCG | ACCGA | ACCGT | ACCGC | ACCGG | | ACG | ACGAA | ACGAT | ACGAC | ACGAG | ACGTA | ACGTT | ACGTC | ACGTG | ACGCA | ACGCT | ACGCC | ACGCG | ACGGA | ACGGT | ACGGC | ACGGG | | AGA | AGAAA | AGAAT | AGAAC | AGAAG | AGATA | AGATT | AGATC | AGATG | AGACA | AGACT | AGACC | AGACG | AGAGA | AGAGT | AGAGC | AGAGG | | AGT | AGTAA | AGTAT | AGTAC | AGTAG | AGTTA | AGTTT | AGTTC | AGTTG | AGTCA | AGTCT | AGTCC | AGTCG | AGTGA | AGTGT | AGTGC | AGTGG | | AGC | AGCAA | AGCAT | AGCAC | AGCAG | AGCTA | AGCTT | AGCTC | AGCTG | AGCCA | AGCCT | AGCCC | AGCCG | AGCGA | AGCGT | AGCGC | AGCGG | | AGG | AGGAA | AGGAT | AGGAC | AGGAG | AGGTA | AGGTT | AGGTC | AGGTG | AGGCA | AGGCT | AGGCC | AGGCG | AGGGA | AGGGT | AGGGC | AGGGG | | TAA | TAAAA | TAAAT | TAAAC | TAAAG | TAATA | TAATT | TAATC | TAATG | TAACA | TAACT | TAACC | TAACG | TAAGA | TAAGT | TAAGC | TAAGG | | TAT | TATAA | TATAT | TATAC | TATAG | TATTA | TATTT | TATTC | TATTG | TATCA | TATCT | TATCC | TATCG | TATGA | TATGT | TATGC | TATGG | | TAC | TACAA | TACAT | TACAC | TACAG | TACTA | TACTT | TACTC | TACTG | TACCA | TACCT | TACCC | TACCG | TACGA | TACGT | TACGC | TACGG | | TAG | TAGAA | TAGAT | TAGAC | TAGAG | TAGTA | TAGTT | TAGTC | TAGTG | TAGCA | TAGCT | TAGCC | TAGCG | TAGGA | TAGGT | TAGGC | TAGGG | | TTA | TTAAA | TTAAT | TTAAC | TTAAG | TTATA | TTATT | TTATC | TTATG | TTACA | TTACT | TTACC | TTACG | TTAGA | TTAGT | TTAGC | TTAGG | | TTT | TTTAA | TTTAT | TTTAC | TTTAG | TTTTA | TTTTT | TTTTC | TTTTG | TTTCA | TTTCT | TTTCC | TTTCG | TTTGA | TTTGT | TTTGC | TTTGG | | TTC | TTCAA | TTCAT | TTCAC | TTCAG | TTCTA | TTCTT | TTCTC | TTCTG | TTCCA | TTCCT | TTCCC | TTCCG | TTCGA | TTCGT | TTCGC | TTCGG | | TTG | TTGAA | TTGAT | TTGAC | TTGAG | TTGTA | TTGTT | TTGTC | TTGTG | TTGCA | TTGCT | TTGCC | TTGCG | TTGGA | TTGGT | TTGGC | TTGGG | | TCA | TCAAA | TCAAT | TCAAC | TCAAG | TCATA | TCATT | TCATC | TCATG | TCACA | TCACT | TCACC | TCACG | TCAGA | TCAGT | TCAGC | TCAGG | | TCT | TCTAA | TCTAT | TCTAC | TCTAG | TCTTA | TCTTT | TCTTC | TCTTG | TCTCA | TCTCT | TCTCC | TCTCG | TCTGA | TCTGT | TCTGC | TCTGG | | TCC | TCCAA | TCCAT | TCCAC | TCCAG | TCCTA | TCCTT | TCCTC | TCCTG | TCCCA | TCCCT | TCCCC | TCCCG | TCCGA | TCCGT | TCCGC | TCCGG | | TCG | TCGAA | TCGAT | TCGAC | TCGAG | TCGTA | TCGTT | TCGTC | TCGTG | TCGCA | TCGCT | TCGCC | TCGCG | TCGGA | TCGGT | TCGGC | TCGGG | | TGA | TGAAA | TGAAT | TGAAC | TGAAG | TGATA | TGATT | TGATC | TGATG | TGACA | TGACT | TGACC | TGACG | TGAGA | TGAGT | TGAGC | TGAGG | | TGT | TGTAA | TGTAT | TGTAC | TGTAG | TGTTA | TGTTT | TGTTC | TGTTG | TGTCA | TGTCT | TGTCC | TGTCG | TGTGA | TGTGT | TGTGC | TGTGG | | TGC | TGCAA | TGCAT | TGCAC | TGCAG | TGCTA | TGCTT | TGCTC | TGCTG | TGCCA | TGCCT | TGCCC | TGCCG | TGCGA | TGCGT | TGCGC | TGCGG | | TGG | TGGAA | TGGAT | TGGAC | TGGAG | TGGTA | TGGTT | TGGTC | TGGTG | TGGCA | TGGCT | TGGCC | TGGCG | TGGGA | TGGGT | TGGGC | TGGGG | | CAA | CAAAA | CAAAT | CAAAC | CAAAG | CAATA | CAATT | CAATC | CAATG | CAACA | CAACT | CAACC | CAACG | CAAGA | CAAGT | CAAGC | CAAGG | | CAT | CATAA | CATAT | CATAC | CATAG | CATTA | CATTT | CATTC | CATTG | CATCA | CATCT | CATCC | CATCG | CATGA | CATGT | CATGC | CATGG | | CAC | CACAA | CACAT | CACAC | CACAG | CACTA | CACTT | CACTC | CACTG | CACCA | CACCT | CACCC | CACCG | CACGA | CACGT | CACGC | CACGG | | CAG | CAGAA | CAGAT | CAGAC | CAGAG | CAGTA | CAGTT | CAGTC | CAGTG | CAGCA | CAGCT | CAGCC | CAGCG | CAGGA | CAGGT | CAGGC | CAGGG | | CTA | CTAAA | CTAAT | CTAAC | CTAAG | CTATA | CTATT | CTATC | CTATG | CTACA | CTACT | CTACC | CTACG | CTAGA | CTAGT | CTAGC | CTAGG | | CTT | CTTAA | CTTAT | CTTAC | CTTAG | CTTTA | CTTTT | CTTTC | CTTTG | CTTCA | CTTCT | CTTCC | CTTCG | CTTGA | CTTGT | CTTGC | CTTGG | | CTC | CTCAA | CTCAT | CTCAC | CTCAG | CTCTA | CTCTT | CTCTC | CTCTG | CTCCA | CTCCT | CTCCC | CTCCG | CTCGA | CTCGT | CTCGC | CTCGG | | CTG | CTGAA | CTGAT | CTGAC | CTGAG | CTGTA | CTGTT | CTGTC | CTGTG | CTGCA | CTGCT | CTGCC | CTGCG | CTGGA | CTGGT | CTGGC | CTGGG | | CCA | CCAAA | CCAAT | CCAAC | CCAAG | CCATA | CCATT | CCATC | CCATG | CCACA | CCACT | CCACC | CCACG | CCAGA | CCAGT | CCAGC | CCAGG | | CCT | CCTAA | CCTAT | CCTAC | CCTAG | CCTTA | CCTTT | CCTTC | CCTTG | CCTCA | CCTCT | CCTCC | CCTCG | CCTGA | CCTGT | CCTGC | CCTGG | | CCC | CCCAA | CCCAT | CCCAC | CCCAG | CCCTA | CCCTT | CCCTC | CCCTG | CCCCA | CCCCT | CCCCC | CCCCG | CCCGA | CCCGT | CCCGC | CCCGG | | CCG | CCGAA | CCGAT | CCGAC | CCGAG | CCGTA | CCGTT | CCGTC | CCGTG | CCGCA | CCGCT | CCGCC | CCGCG | CCGGA | CCGGT | CCGGC | CCGGG | | CGA | CGAAA | CGAAT | CGAAC | CGAAG | CGATA | CGATT | CGATC | CGATG | CGACA | CGACT | CGACC | CGACG | CGAGA | CGAGT | CGAGC | CGAGG | | CGT | CGTAA | CGTAT | CGTAC | CGTAG | CGTTA | CGTTT | CGTTC | CGTTG | CGTCA | CGTCT | CGTCC | CGTCG | CGTGA | CGTGT | CGTGC | CGTGG | | CGC | CGCAA | CGCAT | CGCAC | CGCAG | CGCTA | CGCTT | CGCTC | CGCTG | CGCCA | CGCCT | CGCCC | CGCCG | CGCGA | CGCGT | CGCGC | CGCGG | | CGG | CGGAA | CGGAT | CGGAC | CGGAG | CGGTA | CGGTT | CGGTC | CGGTG | CGGCA | CGGCT | CGGCC | CGGCG | CGGGA | CGGGT | CGGGC | CGGGG | | GAA | GAAAA | GAAAT | GAAAC | GAAAG | GAATA | GAATT | GAATC | GAATG | GAACA | GAACT | GAACC | GAACG | GAAGA | GAAGT | GAAGC | GAAGG | | GAT | GATAA | GATAT | GATAC | GATAG | GATTA | GATTT | GATTC | GATTG | GATCA | GATCT | GATCC | GATCG | GATGA | GATGT | GATGC | GATGG | | GAC | GACAA | GACAT | GACAC | GACAG | GACTA | GACTT | GACTC | GACTG | GACCA | GACCT | GACCC | GACCG | GACGA | GACGT | GACGC | GACGG | | GAG | GAGAA | GAGAT | GAGAC | GAGAG | GAGTA | GAGTT | GAGTC | GAGTG | GAGCA | GAGCT | GAGCC | GAGCG | GAGGA | GAGGT | GAGGC | GAGGG | | GTA | GTAAA | GTAAT | GTAAC | GTAAG | GTATA | GTATT | GTATC | GTATG | GTACA | GTACT | GTACC | GTACG | GTAGA | GTAGT | GTAGC | GTAGG | | GTT | GTTAA | GTTAT | GTTAC | GTTAG | GTTTA | GTTTT | GTTTC | GTTTG | GTTCA | GTTCT | GTTCC | GTTCG | GTTGA | GTTGT | GTTGC | GTTGG | | GTC | GTCAA | GTCAT | GTCAC | GTCAG | GTCTA | GTCTT | GTCTC | GTCTG | GTCCA | GTCCT | GTCCC | GTCCG | GTCGA | GTCGT | GTCGC | GTCGG | | GTG | GTGAA | GTGAT | GTGAC | GTGAG | GTGTA | GTGTT | GTGTC | GTGTG | GTGCA | GTGCT | GTGCC | GTGCG | GTGGA | GTGGT | GTGGC | GTGGG | | GCA | GCAAA | GCAAT | GCAAC | GCAAG | GCATA | GCATT | GCATC | GCATG | GCACA | GCACT | GCACC | GCACG | GCAGA | GCAGT | GCAGC | GCAGG | | GCT | GCTAA | GCTAT | GCTAC | GCTAG | GCTTA | GCTTT | GCTTC | GCTTG | GCTCA | GCTCT | GCTCC | GCTCG | GCTGA | GCTGT | GCTGC | GCTGG | | GCC | GCCAA | GCCAT | GCCAC | GCCAG | GCCTA | GCCTT | GCCTC | GCCTG | GCCCA | GCCCT | GCCCC | GCCCG | GCCGA | GCCGT | GCCGC | GCCGG | | GCG | GCGAA | GCGAT | GCGAC | GCGAG | GCGTA | GCGTT | GCGTC | GCGTG | GCGCA | GCGCT | GCGCC | GCGCG | GCGGA | GCGGT | GCGGC | GCGGG | | GGA | GGAAA | GGAAT | GGAAC | GGAAG | GGATA | GGATT | GGATC | GGATG | GGACA | GGACT | GGACC | GGACG | GGAGA | GGAGT | GGAGC | GGAGG | | GGT | GGTAA | GGTAT | GGTAC | GGTAG | GGTTA | GGTTT | GGTTC | GGTTG | GGTCA | GGTCT | GGTCC | GGTCG | GGTGA | GGTGT | GGTGC | GGTGG | | GGC | GGCAA | GGCAT | GGCAC | GGCAG | GGCTA | GGCTT | GGCTC | GGCTG | GGCCA | GGCCT | GGCCC | GGCCG | GGCGA | GGCGT | GGCGC | GGCGG | | GGG | GGGAA | GGGAT | GGGAC | GGGAG | GGGTA | GGGTT | GGGTC | GGGTG | GGGCA | GGGCT | GGGCC | GGGCG | GGGGA | GGGGT | GGGGC | GGGGG | After filtering: read2: overrepresented sequences Sampling rate: 1 / 20 | | | | | --- | --- | --- | | overrepresented sequence | count (% of bases) | distribution: cycle 1 ~ cycle 150 | | AAAAAAAAAAAAAAAAAAAA | 7671 (0.018161%) | | | ACCTGTCTCTTATACACATCTGACGCTGCCGACGACTCGA | 27 (0.000128%) | | | ACGCTGCCGACGACTCGAACAGTGTAGATCTCGGTGGTCG | 6 (0.000028%) | | | ATCTGTCTCTTATACACATCTGACGCTGCCGACGACTCGA | 26 (0.000123%) | | | ATGATCGATC | 374903 (0.443785%) | | | CCTGTCTCTTATACACATCTGACGCTGCCGACGACTCGAA | 42 (0.000199%) | | | CTGTCTCTTATACACATCTGACGCTGCCGACGACTCGAAC | 18 (0.000085%) | | | GACGCTGCCGACGACTCGAACAGTGTAGATCTCGGTGGTC | 11 (0.000052%) | | | GATCGATCAT | 195357 (0.231251%) | | | GCTGTCTCTTATACACATCTGACGCTGCCGACGACTCGAA | 20 (0.000095%) | | | TCTGTCTCTTATACACATCTGACGCTGCCGACGACTCGAA | 26 (0.000123%) | | | TGACGCTGCCGACGACTCGAACAGTGTAGATCTCGGTGGT | 9 (0.000043%) | | | TGATCGATCA | 221193 (0.261833%) | | | TTTTTTTTTT | 212492 (0.251534%) | |

fastp -p -i resources/raw\_hic/SRR23080414\_1.fastq.gz -I resources/raw\_hic/SRR23080414\_2.fastq.gz -o results/fastp/hic\_trim\_1.fastq.gz -O results/fastp/hic\_trim\_2.fastq.gz --detect\_adapter\_for\_pe --json results/fastp/hic\_report\_fastp.HiC.json --html results/fastp/hic\_report\_fastp.HiC.html --thread 20

fastp 0.23.4, at 2024-04-26 16:54:08
