## Additional files for "Colora: A Snakemake Workflow for Complete Chromosome-scale *De Novo* Genome Assembly": S8_fastp_A.thaliana.html

fastp report at 2024-04-29 23:10:53 

### fastp report Summary General | | | | --- | --- | | fastp version: | 0.23.4 (https://github.com/OpenGene/fastp) | | sequencing: | paired end (150 cycles + 150 cycles) | | mean length before filtering: | 150bp, 150bp | | mean length after filtering: | 148bp, 148bp | | duplication rate: | 9.464361% | | Insert size peak: | 269 | Before filtering | | | | --- | --- | | total reads: | 140.957500 M | | total bases: | 21.143625 G | | Q20 bases: | 20.447749 G (96.708815%) | | Q30 bases: | 19.262000 G (91.100747%) | | GC content: | 39.431368% | After filtering | | | | --- | --- | | total reads: | 140.120386 M | | total bases: | 20.871882 G | | Q20 bases: | 20.234323 G (96.945371%) | | Q30 bases: | 19.079098 G (91.410531%) | | GC content: | 39.356446% | Filtering result | | | | --- | --- | | reads passed filters: | 140.120386 M (99.406123%) | | reads with low quality: | 822.994000 K (0.583860%) | | reads with too many N: | 360 (0.000255%) | | reads too short: | 13.760000 K (0.009762%) | Adapters Adapter or bad ligation of read1 The input has little adapter percentage (~0.663794%), probably it's trimmed before. | | | | --- | --- | | Sequence | Occurrences | | A | 44602 | | AG | 39964 | | AGA | 38018 | | AGAT | 37142 | | AGATC | 34863 | | AGATCG | 34099 | | AGATCGG | 33010 | | AGATCGGA | 32658 | | AGATCGGAA | 31158 | | AGATCGGAAG | 31217 | | AGATCGGAAGA | 30159 | | AGATCGGAAGAG | 29458 | | AGATCGGAAGAGC | 28073 | | AGATCGGAAGAGCA | 27423 | | AGATCGGAAGAGCAC | 26519 | | AGATCGGAAGAGCACA | 25496 | | AGATCGGAAGAGCACAC | 24565 | | AGATCGGAAGAGCACACG | 24403 | | AGATCGGAAGAGCACACGT | 23364 | | AGATCGGAAGAGCACACGTC | 23233 | | AGATCGGAAGAGCACACGTCT | 23077 | | AGATCGGAAGAGCACACGTCTG | 22236 | | AGATCGGAAGAGCACACGTCTGA | 21548 | | AGATCGGAAGAGCACACGTCTGAACTCCAGTCACGAGATTCCATCTCGTATGCCGTCTTCTGCTTGAAAA | 29027 | | other adapter sequences | 1419410 | Adapter or bad ligation of read2 The input has little adapter percentage (~0.674501%), probably it's trimmed before. | | | | --- | --- | | Sequence | Occurrences | | A | 45050 | | AG | 40601 | | AGA | 38484 | | AGAT | 37579 | | AGATC | 35307 | | AGATCG | 34507 | | AGATCGG | 33528 | | AGATCGGA | 33053 | | AGATCGGAA | 31476 | | AGATCGGAAG | 31588 | | AGATCGGAAGA | 30418 | | AGATCGGAAGAG | 29773 | | AGATCGGAAGAGC | 28051 | | AGATCGGAAGAGCG | 27669 | | AGATCGGAAGAGCGT | 26690 | | AGATCGGAAGAGCGTC | 25571 | | AGATCGGAAGAGCGTCG | 24541 | | AGATCGGAAGAGCGTCGT | 24065 | | AGATCGGAAGAGCGTCGTG | 23189 | | AGATCGGAAGAGCGTCGTGT | 22876 | | AGATCGGAAGAGCGTCGTGTA | 22524 | | AGATCGGAAGAGCGTCGTGTAG | 21782 | | AGATCGGAAGAGCGTCGTGTAGGGAAAGAGTGTTCAGAGCCGTGTAGATCTCGGTGGTCGCCGTATCATTAAAAA | 25870 | | other adapter sequences | 1441969 | Insert size estimation This estimation is based on paired-end overlap analysis, and there are 68.372577% reads found not overlapped. The nonoverlapped read pairs may have insert size <30 or >270, or contain too much sequencing errors to be detected as overlapped. Before filtering Before filtering: read1: quality Value of each position will be shown on mouse over. Before filtering: read1: base contents Value of each position will be shown on mouse over. Before filtering: read1: KMER counting Darker background means larger counts. The count will be shown on mouse over. | | | | | | | | | | | | | | | | | | | --- | --- | --- | --- | --- | --- | --- | --- | --- | --- | --- | --- | --- | --- | --- | --- | --- | | | AA | AT | AC | AG | TA | TT | TC | TG | CA | CT | CC | CG | GA | GT | GC | GG | | AAA | AAAAA | AAAAT | AAAAC | AAAAG | AAATA | AAATT | AAATC | AAATG | AAACA | AAACT | AAACC | AAACG | AAAGA | AAAGT | AAAGC | AAAGG | | AAT | AATAA | AATAT | AATAC | AATAG | AATTA | AATTT | AATTC | AATTG | AATCA | AATCT | AATCC | AATCG | AATGA | AATGT | AATGC | AATGG | | AAC | AACAA | AACAT | AACAC | AACAG | AACTA | AACTT | AACTC | AACTG | AACCA | AACCT | AACCC | AACCG | AACGA | AACGT | AACGC | AACGG | | AAG | AAGAA | AAGAT | AAGAC | AAGAG | AAGTA | AAGTT | AAGTC | AAGTG | AAGCA | AAGCT | AAGCC | AAGCG | AAGGA | AAGGT | AAGGC | AAGGG | | ATA | ATAAA | ATAAT | ATAAC | ATAAG | ATATA | ATATT | ATATC | ATATG | ATACA | ATACT | ATACC | ATACG | ATAGA | ATAGT | ATAGC | ATAGG | | ATT | ATTAA | ATTAT | ATTAC | ATTAG | ATTTA | ATTTT | ATTTC | ATTTG | ATTCA | ATTCT | ATTCC | ATTCG | ATTGA | ATTGT | ATTGC | ATTGG | | ATC | ATCAA | ATCAT | ATCAC | ATCAG | ATCTA | ATCTT | ATCTC | ATCTG | ATCCA | ATCCT | ATCCC | ATCCG | ATCGA | ATCGT | ATCGC | ATCGG | | ATG | ATGAA | ATGAT | ATGAC | ATGAG | ATGTA | ATGTT | ATGTC | ATGTG | ATGCA | ATGCT | ATGCC | ATGCG | ATGGA | ATGGT | ATGGC | ATGGG | | ACA | ACAAA | ACAAT | ACAAC | ACAAG | ACATA | ACATT | ACATC | ACATG | ACACA | ACACT | ACACC | ACACG | ACAGA | ACAGT | ACAGC | ACAGG | | ACT | ACTAA | ACTAT | ACTAC | ACTAG | ACTTA | ACTTT | ACTTC | ACTTG | ACTCA | ACTCT | ACTCC | ACTCG | ACTGA | ACTGT | ACTGC | ACTGG | | ACC | ACCAA | ACCAT | ACCAC | ACCAG | ACCTA | ACCTT | ACCTC | ACCTG | ACCCA | ACCCT | ACCCC | ACCCG | ACCGA | ACCGT | ACCGC | ACCGG | | ACG | ACGAA | ACGAT | ACGAC | ACGAG | ACGTA | ACGTT | ACGTC | ACGTG | ACGCA | ACGCT | ACGCC | ACGCG | ACGGA | ACGGT | ACGGC | ACGGG | | AGA | AGAAA | AGAAT | AGAAC | AGAAG | AGATA | AGATT | AGATC | AGATG | AGACA | AGACT | AGACC | AGACG | AGAGA | AGAGT | AGAGC | AGAGG | | AGT | AGTAA | AGTAT | AGTAC | AGTAG | AGTTA | AGTTT | AGTTC | AGTTG | AGTCA | AGTCT | AGTCC | AGTCG | AGTGA | AGTGT | AGTGC | AGTGG | | AGC | AGCAA | AGCAT | AGCAC | AGCAG | AGCTA | AGCTT | AGCTC | AGCTG | AGCCA | AGCCT | AGCCC | AGCCG | AGCGA | AGCGT | AGCGC | AGCGG | | AGG | AGGAA | AGGAT | AGGAC | AGGAG | AGGTA | AGGTT | AGGTC | AGGTG | AGGCA | AGGCT | AGGCC | AGGCG | AGGGA | AGGGT | AGGGC | AGGGG | | TAA | TAAAA | TAAAT | TAAAC | TAAAG | TAATA | TAATT | TAATC | TAATG | TAACA | TAACT | TAACC | TAACG | TAAGA | TAAGT | TAAGC | TAAGG | | TAT | TATAA | TATAT | TATAC | TATAG | TATTA | TATTT | TATTC | TATTG | TATCA | TATCT | TATCC | TATCG | TATGA | TATGT | TATGC | TATGG | | TAC | TACAA | TACAT | TACAC | TACAG | TACTA | TACTT | TACTC | TACTG | TACCA | TACCT | TACCC | TACCG | TACGA | TACGT | TACGC | TACGG | | TAG | TAGAA | TAGAT | TAGAC | TAGAG | TAGTA | TAGTT | TAGTC | TAGTG | TAGCA | TAGCT | TAGCC | TAGCG | TAGGA | TAGGT | TAGGC | TAGGG | | TTA | TTAAA | TTAAT | TTAAC | TTAAG | TTATA | TTATT | TTATC | TTATG | TTACA | TTACT | TTACC | TTACG | TTAGA | TTAGT | TTAGC | TTAGG | | TTT | TTTAA | TTTAT | TTTAC | TTTAG | TTTTA | TTTTT | TTTTC | TTTTG | TTTCA | TTTCT | TTTCC | TTTCG | TTTGA | TTTGT | TTTGC | TTTGG | | TTC | TTCAA | TTCAT | TTCAC | TTCAG | TTCTA | TTCTT | TTCTC | TTCTG | TTCCA | TTCCT | TTCCC | TTCCG | TTCGA | TTCGT | TTCGC | TTCGG | | TTG | TTGAA | TTGAT | TTGAC | TTGAG | TTGTA | TTGTT | TTGTC | TTGTG | TTGCA | TTGCT | TTGCC | TTGCG | TTGGA | TTGGT | TTGGC | TTGGG | | TCA | TCAAA | TCAAT | TCAAC | TCAAG | TCATA | TCATT | TCATC | TCATG | TCACA | TCACT | TCACC | TCACG | TCAGA | TCAGT | TCAGC | TCAGG | | TCT | TCTAA | TCTAT | TCTAC | TCTAG | TCTTA | TCTTT | TCTTC | TCTTG | TCTCA | TCTCT | TCTCC | TCTCG | TCTGA | TCTGT | TCTGC | TCTGG | | TCC | TCCAA | TCCAT | TCCAC | TCCAG | TCCTA | TCCTT | TCCTC | TCCTG | TCCCA | TCCCT | TCCCC | TCCCG | TCCGA | TCCGT | TCCGC | TCCGG | | TCG | TCGAA | TCGAT | TCGAC | TCGAG | TCGTA | TCGTT | TCGTC | TCGTG | TCGCA | TCGCT | TCGCC | TCGCG | TCGGA | TCGGT | TCGGC | TCGGG | | TGA | TGAAA | TGAAT | TGAAC | TGAAG | TGATA | TGATT | TGATC | TGATG | TGACA | TGACT | TGACC | TGACG | TGAGA | TGAGT | TGAGC | TGAGG | | TGT | TGTAA | TGTAT | TGTAC | TGTAG | TGTTA | TGTTT | TGTTC | TGTTG | TGTCA | TGTCT | TGTCC | TGTCG | TGTGA | TGTGT | TGTGC | TGTGG | | TGC | TGCAA | TGCAT | TGCAC | TGCAG | TGCTA | TGCTT | TGCTC | TGCTG | TGCCA | TGCCT | TGCCC | TGCCG | TGCGA | TGCGT | TGCGC | TGCGG | | TGG | TGGAA | TGGAT | TGGAC | TGGAG | TGGTA | TGGTT | TGGTC | TGGTG | TGGCA | TGGCT | TGGCC | TGGCG | TGGGA | TGGGT | TGGGC | TGGGG | | CAA | CAAAA | CAAAT | CAAAC | CAAAG | CAATA | CAATT | CAATC | CAATG | CAACA | CAACT | CAACC | CAACG | CAAGA | CAAGT | CAAGC | CAAGG | | CAT | CATAA | CATAT | CATAC | CATAG | CATTA | CATTT | CATTC | CATTG | CATCA | CATCT | CATCC | CATCG | CATGA | CATGT | CATGC | CATGG | | CAC | CACAA | CACAT | CACAC | CACAG | CACTA | CACTT | CACTC | CACTG | CACCA | CACCT | CACCC | CACCG | CACGA | CACGT | CACGC | CACGG | | CAG | CAGAA | CAGAT | CAGAC | CAGAG | CAGTA | CAGTT | CAGTC | CAGTG | CAGCA | CAGCT | CAGCC | CAGCG | CAGGA | CAGGT | CAGGC | CAGGG | | CTA | CTAAA | CTAAT | CTAAC | CTAAG | CTATA | CTATT | CTATC | CTATG | CTACA | CTACT | CTACC | CTACG | CTAGA | CTAGT | CTAGC | CTAGG | | CTT | CTTAA | CTTAT | CTTAC | CTTAG | CTTTA | CTTTT | CTTTC | CTTTG | CTTCA | CTTCT | CTTCC | CTTCG | CTTGA | CTTGT | CTTGC | CTTGG | | CTC | CTCAA | CTCAT | CTCAC | CTCAG | CTCTA | CTCTT | CTCTC | CTCTG | CTCCA | CTCCT | CTCCC | CTCCG | CTCGA | CTCGT | CTCGC | CTCGG | | CTG | CTGAA | CTGAT | CTGAC | CTGAG | CTGTA | CTGTT | CTGTC | CTGTG | CTGCA | CTGCT | CTGCC | CTGCG | CTGGA | CTGGT | CTGGC | CTGGG | | CCA | CCAAA | CCAAT | CCAAC | CCAAG | CCATA | CCATT | CCATC | CCATG | CCACA | CCACT | CCACC | CCACG | CCAGA | CCAGT | CCAGC | CCAGG | | CCT | CCTAA | CCTAT | CCTAC | CCTAG | CCTTA | CCTTT | CCTTC | CCTTG | CCTCA | CCTCT | CCTCC | CCTCG | CCTGA | CCTGT | CCTGC | CCTGG | | CCC | CCCAA | CCCAT | CCCAC | CCCAG | CCCTA | CCCTT | CCCTC | CCCTG | CCCCA | CCCCT | CCCCC | CCCCG | CCCGA | CCCGT | CCCGC | CCCGG | | CCG | CCGAA | CCGAT | CCGAC | CCGAG | CCGTA | CCGTT | CCGTC | CCGTG | CCGCA | CCGCT | CCGCC | CCGCG | CCGGA | CCGGT | CCGGC | CCGGG | | CGA | CGAAA | CGAAT | CGAAC | CGAAG | CGATA | CGATT | CGATC | CGATG | CGACA | CGACT | CGACC | CGACG | CGAGA | CGAGT | CGAGC | CGAGG | | CGT | CGTAA | CGTAT | CGTAC | CGTAG | CGTTA | CGTTT | CGTTC | CGTTG | CGTCA | CGTCT | CGTCC | CGTCG | CGTGA | CGTGT | CGTGC | CGTGG | | CGC | CGCAA | CGCAT | CGCAC | CGCAG | CGCTA | CGCTT | CGCTC | CGCTG | CGCCA | CGCCT | CGCCC | CGCCG | CGCGA | CGCGT | CGCGC | CGCGG | | CGG | CGGAA | CGGAT | CGGAC | CGGAG | CGGTA | CGGTT | CGGTC | CGGTG | CGGCA | CGGCT | CGGCC | CGGCG | CGGGA | CGGGT | CGGGC | CGGGG | | GAA | GAAAA | GAAAT | GAAAC | GAAAG | GAATA | GAATT | GAATC | GAATG | GAACA | GAACT | GAACC | GAACG | GAAGA | GAAGT | GAAGC | GAAGG | | GAT | GATAA | GATAT | GATAC | GATAG | GATTA | GATTT | GATTC | GATTG | GATCA | GATCT | GATCC | GATCG | GATGA | GATGT | GATGC | GATGG | | GAC | GACAA | GACAT | GACAC | GACAG | GACTA | GACTT | GACTC | GACTG | GACCA | GACCT | GACCC | GACCG | GACGA | GACGT | GACGC | GACGG | | GAG | GAGAA | GAGAT | GAGAC | GAGAG | GAGTA | GAGTT | GAGTC | GAGTG | GAGCA | GAGCT | GAGCC | GAGCG | GAGGA | GAGGT | GAGGC | GAGGG | | GTA | GTAAA | GTAAT | GTAAC | GTAAG | GTATA | GTATT | GTATC | GTATG | GTACA | GTACT | GTACC | GTACG | GTAGA | GTAGT | GTAGC | GTAGG | | GTT | GTTAA | GTTAT | GTTAC | GTTAG | GTTTA | GTTTT | GTTTC | GTTTG | GTTCA | GTTCT | GTTCC | GTTCG | GTTGA | GTTGT | GTTGC | GTTGG | | GTC | GTCAA | GTCAT | GTCAC | GTCAG | GTCTA | GTCTT | GTCTC | GTCTG | GTCCA | GTCCT | GTCCC | GTCCG | GTCGA | GTCGT | GTCGC | GTCGG | | GTG | GTGAA | GTGAT | GTGAC | GTGAG | GTGTA | GTGTT | GTGTC | GTGTG | GTGCA | GTGCT | GTGCC | GTGCG | GTGGA | GTGGT | GTGGC | GTGGG | | GCA | GCAAA | GCAAT | GCAAC | GCAAG | GCATA | GCATT | GCATC | GCATG | GCACA | GCACT | GCACC | GCACG | GCAGA | GCAGT | GCAGC | GCAGG | | GCT | GCTAA | GCTAT | GCTAC | GCTAG | GCTTA | GCTTT | GCTTC | GCTTG | GCTCA | GCTCT | GCTCC | GCTCG | GCTGA | GCTGT | GCTGC | GCTGG | | GCC | GCCAA | GCCAT | GCCAC | GCCAG | GCCTA | GCCTT | GCCTC | GCCTG | GCCCA | GCCCT | GCCCC | GCCCG | GCCGA | GCCGT | GCCGC | GCCGG | | GCG | GCGAA | GCGAT | GCGAC | GCGAG | GCGTA | GCGTT | GCGTC | GCGTG | GCGCA | GCGCT | GCGCC | GCGCG | GCGGA | GCGGT | GCGGC | GCGGG | | GGA | GGAAA | GGAAT | GGAAC | GGAAG | GGATA | GGATT | GGATC | GGATG | GGACA | GGACT | GGACC | GGACG | GGAGA | GGAGT | GGAGC | GGAGG | | GGT | GGTAA | GGTAT | GGTAC | GGTAG | GGTTA | GGTTT | GGTTC | GGTTG | GGTCA | GGTCT | GGTCC | GGTCG | GGTGA | GGTGT | GGTGC | GGTGG | | GGC | GGCAA | GGCAT | GGCAC | GGCAG | GGCTA | GGCTT | GGCTC | GGCTG | GGCCA | GGCCT | GGCCC | GGCCG | GGCGA | GGCGT | GGCGC | GGCGG | | GGG | GGGAA | GGGAT | GGGAC | GGGAG | GGGTA | GGGTT | GGGTC | GGGTG | GGGCA | GGGCT | GGGCC | GGGCG | GGGGA | GGGGT | GGGGC | GGGGG | Before filtering: read1: overrepresented sequences Sampling rate: 1 / 20 | | | | | --- | --- | --- | | overrepresented sequence | count (% of bases) | distribution: cycle 1 ~ cycle 150 | | AAAAAAAAAA | 59956 (0.113426%) | | | AAAAAGCAACGTCGCTATGAACGCTTGGCTGCCACAAGCCAGTTATCCCTGTGGTAACTTTTCTGACACCTCTAGCTTCAAATTCCGAAGGTCTAAAGGA | 339 (0.006413%) | | | AAAAGATAACGCAGGTGTCCTAAGATGAGCTCAACGAGAACAGAAATCTCGTGTGGAACAAAAGGGTAAAAGCTCGTTTGATTCTGATTTTCAGTACGAA | 12 (0.000227%) | | | AAAAGCAACGTCGCTATGAACGCTTGGCTGCCACAAGCCAGTTATCCCTGTGGTAACTTTTCTGACACCTCTAGCTTCAAATTCCGAAGGTCTAAAGGAT | 33 (0.000624%) | | | AAAAGCCTAAGTAGTGTTTCCTTGTTAGAAGACACAAAGCCAAAGACTCATATGGACTTTGGCTACACCATGAAAGCTTTGAGAAGCAAGAAGAAGGTTG | 25 (0.000473%) | | | AAAATCAGAATCAAACGAGCTTTTACCCTTTTGTTCCACACGAGATTTCTGTTCTCGTTGAGCTCATCTTAGGACACCTGCGTTATCTTTTAACAGATGT | 3 (0.000057%) | | | AAACCGCAACCGGATCTTAAAGGCGTAAGAATTGTATCCTTGTTAGAAGACACAAAGCCAAAGACTCATATGGACTTTGGCTACACCATGAAAGCTTTGA | 74 (0.001400%) | | | AAACGAGCTTTTACCCTTTTGTTCCACACGAGATTTCTGTTCTCGTTGAGCTCATCTTAGGACACCTGCGTTATCTTTTAACAGATGTGCCGCCCCAGCC | 4 (0.000076%) | | | AAACTGCGAATGGCTCATTAAATCAGTTATAGTTTGTTTGATGGTAACTACTACTCGGATAACCGTAGTAATTCTAGAGCTAATACGTGCAACAAACCCC | 4 (0.000076%) | | | AAAGAAGACCCTGTTGAGCTTGACTCTAGTCCGACTTTGTGAAATGACTTGAGAGGTGTAGGATAAGTGGGAGCTTCGGCGCAAGTGAAATACCACTACT | 15 (0.000284%) | | | AAAGACTCATATGGACTTTGGCTACACCATGAAAGCTTTGAGAAGCAAGAAGAAGGTTGGTTAGTGTTTTGGAGTCGAATATGACTTGATGTCATGTGTA | 90 (0.001703%) | | | AAAGAGCCCGGTATTGTTATTTATTGTCACTACCTCCCCGTGTCAGGATTGGGTAATTTGCGCGCCTGCTGCCTTCCTTGGATGTGGTAGCCGTTTCTCA | 6 (0.000114%) | | | AAAGATTAAGCCATGCATGTGTAAGTATGAACGAATTCAGACTGTGAAACTGCGAATGGCTCATTAAATCAGTTATAGTTTGTTTGATGGTAACTACTAC | 18 (0.000341%) | | | AAAGCAACGTCGCTATGAACGCTTGGCTGCCACAAGCCAGTTATCCCTGTGGTAACTTTTCTGACACCTCTAGCTTCAAATTCCGAAGGTCTAAAGGATC | 269 (0.005089%) | | | AAAGCCAAAGACTCATATGGACTTTGGCTACACCATGAAAGCTTTGAGAAGCAAGAAGAAGGTTGGTTAGTGTTTTGGAGTCGAATATGACTTGATGTCA | 314 (0.005940%) | | | AAAGCCTAAGTAGTGTTTCCTTGTTAGAAGACACAAAGCCAAAGACTCATATGGACTTTGGCTACACCATGAAAGCTTTGAGAAGCAAGAAGAAGGTTGG | 8 (0.000151%) | | | AAAGCTTTCATGGTGTAGCCAAAGTCCATATGAGTCTTTGGCTTTGTGTCTTCTAACAAGGAAACACTACTTAGGCTTATAAGATGCGGTTGCGGTTTAA | 8 (0.000151%) | | | AAAGGCGTAAGAATTGTATCCTTGTTAAAAGACACAAAGCCAAAGACTCATATGGACTTTGGCTACACCATGAAAGCTTTGAGAAGCAAGAAGAAGGTTG | 13 (0.000246%) | | | AAAGGCGTAAGAATTGTATCCTTGTTAGAAGACACAAAGCCAAAGACTCATATGGACTTTGGCTACACCATGAAAGCTTTGAGAAGCAAGAAGAAGGTTG | 27 (0.000511%) | | | AAATCAGTTATAGTTTGTTTGATGGTAACTACTACTCGGATAACCGTAGTAATTCTAGAGCTAATACGTGCAACAAACCCCGACTTATGGAAGGGACGCA | 3 (0.000057%) | | | AAATCCGCTAAGGAGTGTGTAACAACTCACCTGCCGAATCAACTAGCCCCGAAAATGGATGGCGCTTAAGCGCGCGACCTATACCCGGCCGTCGGGGCAA | 9 (0.000170%) | | | AAATCCTATGATGTTATCCCATGCTAATGTATCCAGAGCGTAGGCTTGCTTTGAGCACTCTAATTTCTTCAAAGTAACAGCGCCGGAGGCACGACCCGGC | 32 (0.000605%) | | | AAATCCTATTTTCTGGTAAATTTTCATAATTTTTTGACACCTCTAGCTAGGTCATTTGACCTGATACAACATCGGATTTTCATGGTCTAGTTGGGGCTCC | 5 (0.000095%) | | | AAATTTTCATAATTTTTTGACACCTCTAGCTAGGTCATTTGACCTGATACAACATCGGATTTTCATGGTCTAGTTGGGGCTCCGTGGGCATATTTGATGC | 8 (0.000151%) | | | AACAGGGACAGTCGGGGGCATTCGTATTTCATAGTCAGAGGTGAAATTCTTGGATTTATGAAAGACGAACAACTGCGAAAGCATTTGCCAAGGATGTTTT | 19 (0.000359%) | | | AACAGGTCTGTGATGCCCTTAGATGTTCTGGGCCGCACGCGCGCTACACTGATGTATTCAACGAGTTCACACCTTGGCCGACAGGCCCGGGTAATCTTTG | 9 (0.000170%) | | | AACCGCAACCGGATCTTAAAGGCGTAAGAATTGTATCCTTGTTAGAAGACACAAAGCCAAAGACTCATATGGACTTTGGCTACACCATGAAAGCTTTGAG | 3 (0.000057%) | | | AACCTAGGCGAGACAAGGGTTCACATTTCGTTCATCACCCTTGGCCGGCTATCGAACAGCCGGACTCCCATCAAAAGATGGTTGCCAAGAACATCTTCGT | 3 (0.000057%) | | | AACGAAGCACGCCCATCCAACCTAGGCGAGACAAGGGTTCACATTTCGTTCATCACCCTTGGCCGGCTATCGAACAGCCGGACTCCCATCAAAAGATGGT | 19 (0.000359%) | | | AACGAAGCACGCCCATCCAACCTAGGCGAGACAAGGGTTCACATTTCGTTCATCACCCTTGGCCGGCTTTCGAACAGCCGGACTCCCATCAAAAGATGGT | 21 (0.000397%) | | | AACGCCTCGAAGAACTAATGGCAGCCCACGCAAGGCAAGCCCATTCTCCTCGACGATTCAGCAGTTTTTGTCCGAGAACTGCTGAGAAAACTCGGAAAAA | 11 (0.000208%) | | | AACGGCGTGCCTCGGCATCAGCGTGCTCCGGGCGTCGGCCTGTGGGCTCCCCATTCGACCCGTCTTGAAACACGGACCAAGGAGTCTGACATGTGTGCGA | 3 (0.000057%) | | | AACGGGCAGAGCCCGCGTCGACCTTTTATCTAATAAATGCGTCCCTTCCATAAGTCGGGGTTTGTTGCACGTATTAGCTCTAGAATTACTACGGTTATCC | 13 (0.000246%) | | | AAGAAGACCCTGTTGAGCTTGACTCTAGTCCGACTTTGTGAAATGACTTGAGAGGTGTAGGATAAGTGGGAGCTTCGGCGCAAGTGAAATACCACTACTT | 7 (0.000132%) | | | AAGAATTGTATCCTTGTTAGAAGACACAAAGCCAAAGACTCATATGGACTTTGGCTACACCATGAAAGCTTTGAGAAGCAAGAAGAAGGTTGGTTAGTGT | 17 (0.000322%) | | | AAGACACAAAGCCAAAGACTCATATGGACTTTGGCTACACCATGAAAGCTTTGAGAAGCAAGAAGAAGGTTGGTTAGTGTTTTGGAGTCGAATATGACTT | 658 (0.012448%) | | | AAGACCCTGTTGAGCTTGACTCTAGTCCGACTTTGTGAAATGACTTGAGAGGTGTAGGATAAGTGGGAGCTTCGGCGCAAGTGAAATACCACTACTTTTA | 3 (0.000057%) | | | AAGACTCATATGGACTTTGGCTACACCATGAAAGCTTTGAGAAGCAAGAAGAAGGTTGGTTAGTGTTTTGGAGTCGAATATGACTTGATGTCATGTGTAT | 23 (0.000435%) | | | AAGAGCACACGTCTGAACTCCAGTCACGAGATTCCATCTC | 8 (0.000061%) | | | AAGAGCCCGGTATTGTTATTTATTGTCACTACCTCCCCGTGTCAGGATTGGGTAATTTGCGCGCCTGCTGCCTTCCTTGGATGTGGTAGCCGTTTCTCAG | 7 (0.000132%) | | | AAGATCGATC | 202437 (0.382975%) | | | AAGCAACGTCGCTATGAACGCTTGGCTGCCACAAGCCAGTTATCCCTGTGGTAACTTTTCTGACACCTCTAGCTTCAAATTCCGAAGGTCTAAAGGATCG | 138 (0.002611%) | | | AAGCACGCCCATCCAACCTAGGCGAGACAAGGGTTCACATTTCGTTCATCACCCTTGGCCGGCTTTCGAACAGCCGGACTCCCATCAAAAGATGGTTGCC | 3 (0.000057%) | | | AAGCCAAAGACTCATATGGACTTTGGCTACACCATGAAAGCTTTGAGAAGCAAGAAGAAGGTTGGTTAGTGTTTTGGAGTCGAATATGACTTGATGTCAT | 11 (0.000208%) | | | AAGCCATGCATGTGTAAGTATGAACGAATTCAGACTGTGAAACTGCGAATGGCTCATTAAATCAGTTATAGTTTGTTTGATGGTAACTACTACTCGGATA | 3 (0.000057%) | | | AAGCCTAAGTAGTGTTTCCTTGTTAGAAGACACAAAGCCAAAGACTCATATGGACTTTGGCTACACCATGAAAGCTTTGAGAAGCAAGAAGAAGGTTGGT | 1202 (0.022740%) | | | AAGCGCGCGACCTATACCCGGCCGTCGGGGCAAGAGCCAGGCCTCGATGAGTAGGAGGGCGCGGCGGTCGCTGCAAAACCTAGGGCGCGAGCCCGGGCGG | 12 (0.000227%) | | | AAGCTTTCATGGTGTAGCCAAAGTCCATATGAGTCTTTGGCTTTGTGTCTTCTAACAAGGAAACACTACTTAGGCTTATAAGATGCGGTTGCGGTTTAAG | 4 (0.000076%) | | | AAGGATACTAAATCCTATTTTCTGGTAAATTTTCATAATTTTTTGACACCTCTAGCTAGGTCATTTGACCTGATACAACATCGGATTTTCATGGTCTAGT | 9 (0.000170%) | | | AAGGCGTAAGAATTGTATCCTTGTTAAAAGACACAAAGCCAAAGACTCATATGGACTTTGGCTACACCATGAAAGCTTTGAGAAGCAAGAAGAAGGTTGG | 7 (0.000132%) | | | AAGGCGTAAGAATTGTATCCTTGTTAGAAGACACAAAGCCAAAGACTCATATGGACTTTGGCTACACCATGAAAGCTTTGAGAAGCAAGAAGAAGGTTGG | 10 (0.000189%) | | | AAGGGATTTAGATTGTACTCATTCCAATTACCAGACTCGAAAGAGCCCGGTATTGTTATTTATTGTCACTACCTCCCCGTGTCAGGATTGGGTAATTTGC | 57 (0.001078%) | | | AAGTAGTGTTTCCTTGTTAGAAGACACAAAGCCAAAGACTCATATGGACTTTGGCTACACCATGAAAGCTTTGAGAAGCAAGAAGAAGGTTGGTTAGTGT | 18 (0.000341%) | | | AAGTCATATTCGACTCCAAAACACTAACCAACCTTCTTCTTGCTTCTCAAAGCTTTCATGGTGTAGCCAAAGTCCATATGAGTCTTTGGCTTTGTGTCTT | 136 (0.002573%) | | | AAGTCGAAATCCGCTAAGGAGTGTGTAACAACTCACCTGCCGAATCAACTAGCCCCGAAAATGGATGGCGCTTAAGCGCGCGACCTATACCCGGCCGTCG | 7 (0.000132%) | | | AAGTCTGGTGCCAGCAGCCGCGGTAATTCCAGCTCCAATAGCGTATATTTAAGTTGTTGCAGTTAAAAAGCTCGTAGTTGAACCTTGGGATGGGTCGGCC | 3 (0.000057%) | | | AAGTTCACCACCGCATGTCGGTACGCTCCAGGCGTCCTTGGCTCGGATTTAGGCCAACCGCGTGCGGTAACACACGGGAGACCAGCTTCCGTCCCGCATC | 23 (0.000435%) | | | AAGTTCTTATACTCAATCATACACATGACATCAAGTCATATTCGACTCCAAAACACTAACCAACCTTCTTCTTGCTTCTCAAAGCTTTCATGGTGTAGCC | 35 (0.000662%) | | | AAGTTGTTATACTCAATCATACACATGACATCAAGTCATA | 116 (0.000878%) | | | AATAACAGGTCTGTGATGCCCTTAGATGTTCTGGGCCGCACGCGCGCTACACTGATGTATTCAACGAGTTCACACCTTGGCCGACAGGCCCGGGTAATCT | 20 (0.000378%) | | | AATATGACTTGATGTCATGTGTATGATTGAGTATAAGAAC | 1390 (0.010519%) | | | AATCAAACGAGCTTTTACCCTTTTGTTCCACACGAGATTTCTGTTCTCGTTGAGCTCATCTTAGGACACCTGCGTTATCTTTTAACAGATGTGCCGCCCC | 32 (0.000605%) | | | AATCAGAATCAAACGAGCTTTTACCCTTTTGTTCCACACGAGATTTCTGTTCTCGTTGAGCTCATCTTAGGACACCTGCGTTATCTTTTAACAGATGTGC | 4 (0.000076%) | | | AATCAGTTATAGTTTGTTTGATGGTAACTACTACTCGGATAACCGTAGTAATTCTAGAGCTAATACGTGCAACAAACCCCGACTTATGGAAGGGACGCAT | 8 (0.000151%) | | | AATCATACACATGACAACAAGTCATATTCGACTCCAAAACACTAACCAACCTTCTTCTTGCTTCTCAAAGCTTTCATGGTGTAGCCAAAGTCCATATGAG | 21 (0.000397%) | | | AATCATACACATGACATCAAGTCATATTCGACTCCAAAACACTAACCAACCTTCTTCTTGCTTCTCAAAGCTTTCATGGTGTAGCCAAAGTCCATATGAG | 63 (0.001192%) | | | AATCATCAGAGCAACGGGCAGAGCCCGCGTCGACCTTTTATCTAATAAATGCGTCCCTTCCATAAGTCGGGGTTTGTTGCACGTATTAGCTCTAGAATTA | 51 (0.000965%) | | | AATCCGCTAAGGAGTGTGTAACAACTCACCTGCCGAATCAACTAGCCCCGAAAATGGATGGCGCTTAAGCGCGCGACCTATACCCGGCCGTCGGGGCAAG | 14 (0.000265%) | | | AATCCTATGATGTTATCCCATGCTAATGTATCCAGAGCGTAGGCTTGCTTTGAGCACTCTAATTTCTTCAAAGTAACAGCGCCGGAGGCACGACCCGGCC | 42 (0.000795%) | | | AATCCTTGTTAGTTTCTTTTCCTCCGCTTATTGATATGCTTAAACTCAGCGGGTAATCCCGCCTGACCTGGGGTCGCTATATGGACTTTGGGTCATCTAC | 8 (0.000151%) | | | AATCGGTAGGAGCGACGGGCGGTGTGTACAAAGGGCAGGGACGTAGTCAACGCGAGCTGATGACTCGCGCTTACTAGGAATTCCTCGTTGAAGACCAACA | 71 (0.001343%) | | | AATGATTAACAGGGACAGTCGGGGGCATTCGTATTTCATAGTCAGAGGTGAAATTCTTGGATTTATGAAAGACGAACAACTGCGAAAGCATTTGCCAAGG | 93 (0.001759%) | | | AATGGAAAGAGAGAAAGGACGAGGTCTTGACCGTCATCTTTTGCCCGAAGGACGGATGAGCTTTGGCGGGACTGAATCACTTCGAGTCACCGTCGACAAC | 4 (0.000076%) | | | AATGTATCCAGAGCGTAGGCTTGCTTTGAGCACTCTAATTTCTTCAAAGTAACAGCGCCGGAGGCACGACCCGGCCAATTAAGACCAGGAGCGTATCGCC | 7 (0.000132%) | | | AATTACCAGACTCGAAAGAGCCCGGTATTGTTATTTATTGTCACTACCTCCCCGTGTCAGGATTGGGTAATTTGCGCGCCTGCTGCCTTCCTTGGATGTG | 23 (0.000435%) | | | AATTAGGGTTCGATTCCGGAGAGGGAGCCTGAGAAACGGCTACCACATCCAAGGAAGGCAGCAGGCGCGCAAATTACCCAATCCTGACACGGGGAGGTAG | 27 (0.000511%) | | | AATTCCAGCTCCAATAGCGTATATTTAAGTTGTTGCAGTTAAAAAGCTCGTAGTTGAACCTTGGGATGGGTCGGCCGGTCCGCCTTTGGTGTGCATTGGT | 12 (0.000227%) | | | AATTCCCCGCCACATCCTCTCAAACGCAATGGAAAGAGAGAAAGGACGAGGTCTTGACCGTCATCTTTTGCCCGAAGGACGGATGAGCTTTGGCGGGACT | 11 (0.000208%) | | | AATTGTATCCTTGTTAGAAGACACAAAGCCAAAGACTCATATGGACTTTGGCTACACCATGAAAGCTTTGAGAAGCAAGAAGAAGGTTGGTTAGTGTTTT | 4 (0.000076%) | | | AATTGTTGGTCTTCAACGAGGAATTCCTAGTAAGCGCGAGTCATCAGCTCGCGTTGACTACGTCCCTGCCCTTTGTACACACCGCCCGTCGCTCCTACCG | 1353 (0.025596%) | | | AATTTTCATAATTTTTTGACACCTCTAGCTAGGTCATTTGACCTGATACAACATCGGATTTTCATGGTCTAGTTGGGGCTCCGTGGGCATATTTGATGCA | 8 (0.000151%) | | | ACAAAGCCAAAGACTCATATGGACTTTGGCTACACCATGAAAGCTTTGAGAAGCAAGAAGAAGGTTGGTTAGTGTTTTGGAGTCGAATATGACTTGATGT | 167 (0.003159%) | | | ACACAAAGCCAAAGACTCATATGGACTTTGGCTACACCATGAAAGCTTTGAGAAGCAAGAAGAAGGTTGGTTAGTGTTTTGGAGTCGAATATGACTTGAT | 185 (0.003500%) | | | ACACATGACATCAAGTCATATTCGACTCCAAAACACTAACCAACCTTCTTCTTGCTTCTCAAAGCTTTCATGGTGTAGCCAAAGTCCATATGAGTCTTTG | 19 (0.000359%) | | | ACACCATGAAAGCTTTGAGAAGCAAGAAGAAGGTTGGTTAGTGTTTTGGAGTCGAATATGACTTGATGTCATGTGTATGATTGAGTATAACAACTTAAAC | 7 (0.000132%) | | | ACACGTCTGAACTCCAGTCACGAGATTCCATCTCGTATGC | 19 (0.000144%) | | | ACATCAAGTCATATTCGACTCCAAAACACTAACCAACCTTCTTCTTGCTTCTCAAAGCTTTCATGGTGTAGCCAAAGTCCATATGAGTCTTTGGCTTTGT | 58 (0.001097%) | | | ACATCTGTTAAAAGATAACGCAGGTGTCCTAAGATGAGCTCAACGAGAACAGAAATCTCGTGTGGAACAAAAGGGTAAAAGCTCGTTTGATTCTGATTTT | 4 (0.000076%) | | | ACATGACATCAAGTCATATTCGACTCCAAAACACTAACCAACCTTCTTCTTGCTTCTCAAAGCTTTCATGGTGTAGCCAAAGTCCATATGAGTCTTTGGC | 70 (0.001324%) | | | ACATTGTCAGGTGGGGAGTTTGGCTGGGGCGGCACATCTGTTAAAAGATAACGCAGGTGTCCTAAGATGAGCTCAACGAGAACAGAAATCTCGTGTGGAA | 8 (0.000151%) | | | ACCACCGCATGTCGGTACGCTCCAGGCGTCCTTGGCTCGGATTTAGGCCAACCGCGTGCGGTAACACACGGGAGACCAGCTTCCGTCCCGCATCAGCAAA | 12 (0.000227%) | | | ACCAGACTCGAAAGAGCCCGGTATTGTTATTTATTGTCACTACCTCCCCGTGTCAGGATTGGGTAATTTGCGCGCCTGCTGCCTTCCTTGGATGTGGTAG | 6 (0.000114%) | | | ACCAGAGGCTGTTCACCTTGGAGACCTGATGCGGTTATGAGTACGACCGGGCGTGAGCGGCACTCGGTCCTCCGGATTTTCAAGGGCCGCCGGGGGCGCA | 10 (0.000189%) | | | ACCATCCTTTGCTGATGCGGGACGGAAGCTGGTCTCCCGTGTGTTACCGCACGCGGTTGGCCTAAATCCGAGCCAAGGACGCCTGGAGCGTACCGACATG | 6 (0.000114%) | | | ACCCGGCCGTCGGGGCAAGAGCCAGGCCTCGATGAGTAGGAGGGCGCGGCGGTCGCTGCAAAACCTAGGGCGCGAGCCCGGGCGGAGCGGCCGTCGGTGC | 3 (0.000057%) | | | ACCCGTTGACTCGCACACATGTCAGACTCCTTGGTCCGTGTTTCAAGACGGGTCGAATGGGGAGCCCACAGGCCGACGCCCGGAGCACGCTGATGCCGAG | 4 (0.000076%) | | | ACCCTGTTGAGCTTGACTCTAGTCCGACTTTGTGAAATGACTTGAGAGGTGTAGGATAAGTGGGAGCTTCGGCGCAAGTGAAATACCACTACTTTTAACG | 5 (0.000095%) | | | ACCGCATGTCGGTACGCTCCAGGCGTCCTTGGCTCGGATTTAGGCCAACCGCGTGCGGTAACACACGGGAGACCAGCTTCCGTCCCGCATCAGCAAAGGA | 6 (0.000114%) | | | ACCGGATTGCTCCGTTCCGCATCCGACCAGGACGCATCGCCGGCCCCCATCCGCTTCCCTCCCGACAATTTCAAGCACTCTTTGACTCTCTTTTCAAAGT | 15 (0.000284%) | | | ACCTAGGCGAGACAAGGGTTCACATTTCGTTCATCACCCTTGGCCGGCTATCGAACAGCCGGACTCCCATCAAAAGATGGTTGCCAAGAACATCTTCGTT | 5 (0.000095%) | | | ACCTAGGCGAGACAAGGGTTCACATTTCGTTCATCACCCTTGGCCGGCTTTCGAACAGCCGGACTCCCATCAAAAGATGGTTGCCAAGAACATCTTCGTT | 3 (0.000057%) | | | ACGAAGCACGCCCATCCAACCTAGGCGAGACAAGGGTTCACATTTCGTTCATCACCCTTGGCCGGCTATCGAACAGCCGGACTCCCATCAAAAGATGGTT | 5 (0.000095%) | | | ACGAAGCACGCCCATCCAACCTAGGCGAGACAAGGGTTCACATTTCGTTCATCACCCTTGGCCGGCTTTCGAACAGCCGGACTCCCATCAAAAGATGGTT | 8 (0.000151%) | | | ACGAGGAATTCCTAGTAAGCGCGAGTCATCAGCTCGCGTTGACTACGTCCCTGCCCTTTGTACACACCGCCCGTCGCTCCTACCGATTGAATGATCGATC | 4 (0.000076%) | | | ACGCGCCTAACGGCGTGCCTCGGCATCAGCGTGCTCCGGGCGTCGGCCTGTGGGCTCCCCATTCGACCCGTCTTGAAACACGGACCAAGGAGTCTGACAT | 14 (0.000265%) | | | ACGCTTTCACGGTTCGTATTCGTACTGAAAATCAGAATCAAACGAGCTTTTACCCTTTTGTTCCACACGAGATTTCTGTTCTCGTTGAGCTCATCTTAGG | 7 (0.000132%) | | | ACGGCAATTCCCCGCCACATCCTCTCAAACGCAATGGAAAGAGAGAAAGGACGAGGTCTTGACCGTCATCTTTTGCCCGAAGGACGGATGAGCTTTGGCG | 7 (0.000132%) | | | ACGGCGTGCCTCGGCATCAGCGTGCTCCGGGCGTCGGCCTGTGGGCTCCCCATTCGACCCGTCTTGAAACACGGACCAAGGAGTCTGACATGTGTGCGAG | 4 (0.000076%) | | | ACGGGCAGAGCCCGCGTCGACCTTTTATCTAATAAATGCGTCCCTTCCATAAGTCGGGGTTTGTTGCACGTATTAGCTCTAGAATTACTACGGTTATCCG | 3 (0.000057%) | | | ACGGGCGGTGTGTACAAAGGGCAGGGACGTAGTCAACGCGAGCTGATGACTCGCGCTTACTAGGAATTCCTCGTTGAAGACCAACAATTGCAATGATCGA | 3 (0.000057%) | | | ACGTCTGAACTCCAGTCACGAGATTCCATCTCGTATGCCG | 18 (0.000136%) | | | ACTAAATCCTATTTTCTGGTAAATTTTCATAATTTTTTGACACCTCTAGCTAGGTCATTTGACCTGATACAACATCGGATTTTCATGGTCTAGTTGGGGC | 24 (0.000454%) | | | ACTCAATCATACACATGACAACAAGTCATATTCGACTCCAAAACACTAACCAACCTTCTTCTTGCTTCTCAAAGCTTTCATGGTGTAGCCAAAGTCCATA | 8 (0.000151%) | | | ACTCAATCATACACATGACATCAAGTCATATTCGACTCCAAAACACTAACCAACCTTCTTCTTGCTTCTCAAAGCTTTCATGGTGTAGCCAAAGTCCATA | 26 (0.000492%) | | | ACTCATATGGACTTTGGCTACACCATGAAAGCTTTGAGAAGCAAGAAGAAGGTTGGTTAGTGTTTTGGAGTCGAATATGACTTGATGTCATGTGTATGAT | 88 (0.001665%) | | | ACTCATTCCAATTACCAGACTCGAAAGAGCCCGGTATTGTTATTTATTGTCACTACCTCCCCGTGTCAGGATTGGGTAATTTGCGCGCCTGCTGCCTTCC | 30 (0.000568%) | | | ACTCCAAAACACTAACCAACCTTCTTCTTGCTTCTCAAAGCTTTCATGGTGTAGCCAAAGTCCATATGAGTCTTTGGCTTTGTGTCTTCTAACAAGGAAA | 203 (0.003840%) | | | ACTCGAAAGAGCCCGGTATTGTTATTTATTGTCACTACCTCCCCGTGTCAGGATTGGGTAATTTGCGCGCCTGCTGCCTTCCTTGGATGTGGTAGCCGTT | 4 (0.000076%) | | | ACTGAAAATCAGAATCAAACGAGCTTTTACCCTTTTGTTCCACACGAGATTTCTGTTCTCGTTGAGCTCATCTTAGGACACCTGCGTTATCTTTTAACAG | 5 (0.000095%) | | | ACTTCCCTTGCCTACATTGTTCCATCGACCAGAGGCTGTTCACCTTGGAGACCTGATGCGGTTATGAGTACGACCGGGCGTGAGCGGCACTCGGTCCTCC | 13 (0.000246%) | | | ACTTGATGTCATGTGTATGATTGAGTATAAGAACTTAAAC | 358 (0.002709%) | | | ACTTTGTGAAATGACTTGAGAGGTGTAGGATAAGTGGGAGCTTCGGCGCAAGTGAAATACCACTACTTTTAACGTTATTTTACTTACTCCGTGAATCGGA | 5 (0.000095%) | | | AGAAGACACAAAGCCAAAGACTCATATGGACTTTGGCTACACCATGAAAGCTTTGAGAAGCAAGAAGAAGGTTGGTTAGTGTTTTGGAGTCGAATATGAC | 81 (0.001532%) | | | AGAAGACCCTGTTGAGCTTGACTCTAGTCCGACTTTGTGAAATGACTTGAGAGGTGTAGGATAAGTGGGAGCTTCGGCGCAAGTGAAATACCACTACTTT | 51 (0.000965%) | | | AGAATTGTATCCTTGTTAGAAGACACAAAGCCAAAGACTCATATGGACTTTGGCTACACCATGAAAGCTTTGAGAAGCAAGAAGAAGGTTGGTTAGTGTT | 6 (0.000114%) | | | AGACAAGGGTTCACATTTCGTTCATCACCCTTGGCCGGCTATCGAACAGCCGGACTCCCATCAAAAGATGGTTGCCAAGAACATCTTCGTTACGGTTTGC | 3 (0.000057%) | | | AGACACAAAGCCAAAGACTCATATGGACTTTGGCTACACCATGAAAGCTTTGAGAAGCAAGAAGAAGGTTGGTTAGTGTTTTGGAGTCGAATATGACTTG | 39 (0.000738%) | | | AGACCCTGTTGAGCTTGACTCTAGTCCGACTTTGTGAAATGACTTGAGAGGTGTAGGATAAGTGGGAGCTTCGGCGCAAGTGAAATACCACTACTTTTAA | 6 (0.000114%) | | | AGACTCATATGGACTTTGGCTACACCATGAAAGCTTTGAGAAGCAAGAAGAAGGTTGGTTAGTGTTTTGGAGTCGAATATGACTTGATGTCATGTGTATG | 15 (0.000284%) | | | AGACTCGAAAGAGCCCGGTATTGTTATTTATTGTCACTACCTCCCCGTGTCAGGATTGGGTAATTTGCGCGCCTGCTGCCTTCCTTGGATGTGGTAGCCG | 5 (0.000095%) | | | AGAGCAACGGGCAGAGCCCGCGTCGACCTTTTATCTAATAAATGCGTCCCTTCCATAAGTCGGGGTTTGTTGCACGTATTAGCTCTAGAATTACTACGGT | 33 (0.000624%) | | | AGAGCACACGTCTGAACTCCAGTCACGAGATTCCATCTCG | 15 (0.000114%) | | | AGAGCCCGCGTCGACCTTTTATCTAATAAATGCGTCCCTTCCATAAGTCGGGGTTTGTTGCACGTATTAGCTCTAGAATTACTACGGTTATCCGAGTAGT | 7 (0.000132%) | | | AGAGCCCGGTATTGTTATTTATTGTCACTACCTCCCCGTGTCAGGATTGGGTAATTTGCGCGCCTGCTGCCTTCCTTGGATGTGGTAGCCGTTTCTCAGG | 3 (0.000057%) | | | AGAGCGTAGGCTTGCTTTGAGCACTCTAATTTCTTCAAAGTAACAGCGCCGGAGGCACGACCCGGCCAATTAAGACCAGGAGCGTATCGCCGACCGAAGG | 14 (0.000265%) | | | AGATCGGAAGAGCACACGTCTGAACTCCAGTCACGAGATT | 20073 (0.151898%) | | | AGATGTTCTGGGCCGCACGCGCGCTACACTGATGTATTCAACGAGTTCACACCTTGGCCGACAGGCCCGGGTAATCTTTGAAATTTCATCGTGATGGGGA | 8 (0.000151%) | | | AGATTAAGCCATGCATGTGTAAGTATGAACGAATTCAGACTGTGAAACTGCGAATGGCTCATTAAATCAGTTATAGTTTGTTTGATGGTAACTACTACTC | 4 (0.000076%) | | | AGATTGTACTCATTCCAATTACCAGACTCGAAAGAGCCCGGTATTGTTATTTATTGTCACTACCTCCCCGTGTCAGGATTGGGTAATTTGCGCGCCTGCT | 12 (0.000227%) | | | AGCAACGGGCAGAGCCCGCGTCGACCTTTTATCTAATAAATGCGTCCCTTCCATAAGTCGGGGTTTGTTGCACGTATTAGCTCTAGAATTACTACGGTTA | 4 (0.000076%) | | | AGCAACGTCGCTATGAACGCTTGGCTGCCACAAGCCAGTTATCCCTGTGGTAACTTTTCTGACACCTCTAGCTTCAAATTCCGAAGGTCTAAAGGATCGA | 32 (0.000605%) | | | AGCACACGTCTGAACTCCAGTCACGAGATTCCATCTCGTA | 8 (0.000061%) | | | AGCACGCCCATCCAACCTAGGCGAGACAAGGGTTCACATTTCGTTCATCACCCTTGGCCGGCTATCGAACAGCCGGACTCCCATCAAAAGATGGTTGCCA | 12 (0.000227%) | | | AGCACGCCCATCCAACCTAGGCGAGACAAGGGTTCACATTTCGTTCATCACCCTTGGCCGGCTTTCGAACAGCCGGACTCCCATCAAAAGATGGTTGCCA | 16 (0.000303%) | | | AGCACGCGCCTAACGGCGTGCCTCGGCATCAGCGTGCTCCGGGCGTCGGCCTGTGGGCTCCCCATTCGACCCGTCTTGAAACACGGACCAAGGAGTCTGA | 31 (0.000586%) | | | AGCAGCCGCGGTAATTCCAGCTCCAATAGCGTATATTTAAGTTGTTGCAGTTAAAAAGCTCGTAGTTGAACCTTGGGATGGGTCGGCCGGTCCGCCTTTG | 7 (0.000132%) | | | AGCCAAAGACTCATATGGACTTTGGCTACACCATGAAAGCTTTGAGAAGCAAGAAGAAGGTTGGTTAGTGTTTTGGAGTCGAATATGACTTGATGTCATG | 10 (0.000189%) | | | AGCCCGCGTCGACCTTTTATCTAATAAATGCGTCCCTTCCATAAGTCGGGGTTTGTTGCACGTATTAGCTCTAGAATTACTACGGTTATCCGAGTAGTAG | 3 (0.000057%) | | | AGCCGCGGTAATTCCAGCTCCAATAGCGTATATTTAAGTTGTTGCAGTTAAAAAGCTCGTAGTTGAACCTTGGGATGGGTCGGCCGGTCCGCCTTTGGTG | 9 (0.000170%) | | | AGCCTAAGTAGTGTTTCCTTGTTAGAAGACACAAAGCCAAAGACTCATATGGACTTTGGCTACACCATGAAAGCTTTGAGAAGCAAGAAGAAGGTTGGTT | 34 (0.000643%) | | | AGCGACGGGCGGTGTGTACAAAGGGCAGGGACGTAGTCAACGCGAGCTGATGACTCGCGCTTACTAGGAATTCCTCGTTGAAGACCAACAATTGCAATGA | 5 (0.000095%) | | | AGCGAGGTGTGAGTGTCGCCCATGGGCATCGACACCTTGCGGCTAGGAACTGGAACGAGACGGGTGGCAAAGATTTCGAGTAGCACTTCATACTACCGTG | 6 (0.000114%) | | | AGCGTAGGCTTGCTTTGAGCACTCTAATTTCTTCAAAGTAACAGCGCCGGAGGCACGACCCGGCCAATTAAGACCAGGAGCGTATCGCCGACCGAAGGGA | 6 (0.000114%) | | | AGGACATTGTCAGGTGGGGAGTTTGGCTGGGGCGGCACATCTGTTAAAAGATAACGCAGGTGTCCTAAGATGAGCTCAACGAGAACAGAAATCTCGTGTG | 188 (0.003557%) | | | AGGAGCGACGGGCGGTGTGTACAAAGGGCAGGGACGTAGTCAACGCGAGCTGATGACTCGCGCTTACTAGGAATTCCTCGTTGAAGACCAACAATTGCAA | 17 (0.000322%) | | | AGGATACTAAATCCTATTTTCTGGTAAATTTTCATAATTTTTTGACACCTCTAGCTAGGTCATTTGACCTGATACAACATCGGATTTTCATGGTCTAGTT | 4 (0.000076%) | | | AGGCCACGCTTTCACGGTTCGTATTCGTACTGAAAATCAGAATCAAACGAGCTTTTACCCTTTTGTTCCACACGAGATTTCTGTTCTCGTTGAGCTCATC | 15 (0.000284%) | | | AGGCGAGACAAGGGTTCACATTTCGTTCATCACCCTTGGCCGGCTTTCGAACAGCCGGACTCCCATCAAAAGATGGTTGCCAAGAACATCTTCGTTACGG | 7 (0.000132%) | | | AGGCGTAAGAATTGTATCCTTGTTAAAAGACACAAAGCCAAAGACTCATATGGACTTTGGCTACACCATGAAAGCTTTGAGAAGCAAGAAGAAGGTTGGT | 6 (0.000114%) | | | AGGCGTAAGAATTGTATCCTTGTTAGAAGACACAAAGCCAAAGACTCATATGGACTTTGGCTACACCATGAAAGCTTTGAGAAGCAAGAAGAAGGTTGGT | 3 (0.000057%) | | | AGGCTGTCCCGAGTGTGAGCGAGGTGTGAGTGTCGCCCATGGGCATCGACACCTTGCGGCTAGGAACTGGAACGAGACGGGTAGCAAAGATTTCGAGTAG | 16 (0.000303%) | | | AGGCTGTCCCGAGTGTGAGCGAGGTGTGAGTGTCGCCCATGGGCATCGACACCTTGCGGCTAGGAACTGGAACGAGACGGGTGGCAAAGATTTCGAGTAG | 11 (0.000208%) | | | AGGGAATCCTTGTTAGTTTCTTTTCCTCCGCTTATTGATATGCTTAAACTCAGCGGGTAATCCCGCCTGACCTGGGGTCGCTATATGGACTTTGGGTCAT | 227 (0.004294%) | | | AGGGACAGTCGGGGGCATTCGTATTTCATAGTCAGAGGTGAAATTCTTGGATTTATGAAAGACGAACAACTGCGAAAGCATTTGCCAAGGATGTTTTCAT | 5 (0.000095%) | | | AGGGATTTAGATTGTACTCATTCCAATTACCAGACTCGAAAGAGCCCGGTATTGTTATTTATTGTCACTACCTCCCCGTGTCAGGATTGGGTAATTTGCG | 10 (0.000189%) | | | AGGGCAAGTCTGGTGCCAGCAGCCGCGGTAATTCCAGCTCCAATAGCGTATATTTAAGTTGTTGCAGTTAAAAAGCTCGTAGTTGAACCTTGGGATGGGT | 25 (0.000473%) | | | AGGTCTGTGATGCCCTTAGATGTTCTGGGCCGCACGCGCGCTACACTGATGTATTCAACGAGTTCACACCTTGGCCGACAGGCCCGGGTAATCTTTGAAA | 6 (0.000114%) | | | AGTAGTCATATGCTTGTCTCAAAGATTAAGCCATGCATGTGTAAGTATGAACGAATTCAGACTGTGAAACTGCGAATGGCTCATTAAATCAGTTATAGTT | 73 (0.001381%) | | | AGTAGTGTTTCCTTGTTAGAAGACACAAAGCCAAAGACTCATATGGACTTTGGCTACACCATGAAAGCTTTGAGAAGCAAGAAGAAGGTTGGTTAGTGTT | 27 (0.000511%) | | | AGTCAGAGGTGAAATTCTTGGATTTATGAAAGACGAACAACTGCGAAAGCATTTGCCAAGGATGTTTTCATTAATCAAGAACGAAAGTTGGGGGCTCGAA | 9 (0.000170%) | | | AGTCATATGCTTGTCTCAAAGATTAAGCCATGCATGTGTAAGTATGAACGAATTCAGACTGTGAAACTGCGAATGGCTCATTAAATCAGTTATAGTTTGT | 11 (0.000208%) | | | AGTCATATTCGACTCCAAAACACTAACCAACCTTCTTCTTGCTTCTCAAAGCTTTCATGGTGTAGCCAAAGTCCATATGAGTCTTTGGCTTTGTGTCTTC | 6 (0.000114%) | | | AGTCGAAATCCGCTAAGGAGTGTGTAACAACTCACCTGCCGAATCAACTAGCCCCGAAAATGGATGGCGCTTAAGCGCGCGACCTATACCCGGCCGTCGG | 9 (0.000170%) | | | AGTCGGGGGCATTCGTATTTCATAGTCAGAGGTGAAATTCTTGGATTTATGAAAGACGAACAACTGCGAAAGCATTTGCCAAGGATGTTTTCATTAATCA | 8 (0.000151%) | | | AGTGTATCCTTGTTAGAAGACACAAAGCCAAAGACTCATATGGACTTTGGCTACACCATGAAAGCTTTGAGAAGCAAGAAGAAGGTTGGTTAGTGTTTTG | 768 (0.014529%) | | | AGTGTGAGCGAGGTGTGAGTGTCGCCCATGGGCATCGACACCTTGCGGCTAGGAACTGGAACGAGACGGGTGGCAAAGATTTCGAGTAGCACTTCATACT | 4 (0.000076%) | | | AGTGTTTCCTTGTTAGAAGACACAAAGCCAAAGACTCATATGGACTTTGGCTACACCATGAAAGCTTTGAGAAGCAAGAAGAAGGTTGGTTAGTGTTTTG | 12 (0.000227%) | | | AGTTATCATGAATCATCAGAGCAACGGGCAGAGCCCGCGTCGACCTTTTATCTAATAAATGCGTCCCTTCCATAAGTCGGGGTTTGTTGCACGTATTAGC | 12 (0.000227%) | | | AGTTCACCACCGCATGTCGGTACGCTCCAGGCGTCCTTGGCTCGGATTTAGGCCAACCGCGTGCGGTAACACACGGGAGACCAGCTTCCGTCCCGCATCA | 6 (0.000114%) | | | AGTTCTTATACTCAATCATACACATGACATCAAGTCATATTCGACTCCAAAACACTAACCAACCTTCTTCTTGCTTCTCAAAGCTTTCATGGTGTAGCCA | 30 (0.000568%) | | | AGTTGTTATACTCAATCATACACATGACATCAAGTCATAT | 98 (0.000742%) | | | AGTTTCTTTTCCTCCGCTTATTGATATGCTTAAACTCAGCGGGTAATCCCGCCTGACCTGGGGTCGCTATATGGACTTTGGGTCATCTACAGCTTCCGGA | 4 (0.000076%) | | | ATAACAGGTCTGTGATGCCCTTAGATGTTCTGGGCCGCACGCGCGCTACACTGATGTATTCAACGAGTTCACACCTTGGCCGACAGGCCCGGGTAATCTT | 27 (0.000511%) | | | ATAAGGATACTAAATCCTATTTTCTGGTAAATTTTCATAATTTTTTGACACCTCTAGCTAGGTCATTTGACCTGATACAACATCGGATTTTCATGGTCTA | 13 (0.000246%) | | | ATAATTTTTTGACACCTCTAGCTAGGTCATTTGACCTGATACAACATCGGATTTTCATGGTCTAGTTGGGGCTCCGTGGGCATATTTGATGCAAACTTGA | 10 (0.000189%) | | | ATACACATGACATCAAGTCATATTCGACTCCAAAACACTAACCAACCTTCTTCTTGCTTCTCAAAGCTTTCATGGTGTAGCCAAAGTCCATATGAGTCTT | 291 (0.005505%) | | | ATACCCGGCCGTCGGGGCAAGAGCCAGGCCTCGATGAGTAGGAGGGCGCGGCGGTCGCTGCAAAACCTAGGGCGCGAGCCCGGGCGGAGCGGCCGTCGGT | 11 (0.000208%) | | | ATACTCAATCATACACATGACAACAAGTCATATTCGACTCCAAAACACTAACCAACCTTCTTCTTGCTTCTCAAAGCTTTCATGGTGTAGCCAAAGTCCA | 14 (0.000265%) | | | ATACTCAATCATACACATGACATCAAGTCATATTCGACTCCAAAACACTAACCAACCTTCTTCTTGCTTCTCAAAGCTTTCATGGTGTAGCCAAAGTCCA | 30 (0.000568%) | | | ATAGGCCACGCTTTCACGGTTCGTATTCGTACTGAAAATCAGAATCAAACGAGCTTTTACCCTTTTGTTCCACACGAGATTTCTGTTCTCGTTGAGCTCA | 88 (0.001665%) | | | ATAGTCAGAGGTGAAATTCTTGGATTTATGAAAGACGAACAACTGCGAAAGCATTTGCCAAGGATGTTTTCATTAATCAAGAACGAAAGTTGGGGGCTCG | 7 (0.000132%) | | | ATAGTGGCCTACCATGGTGGTAACGGGTGACGGAGAATTAGGGTTCGATTCCGGAGAGGGAGCCTGAGAAACGGCTACCACATCCAAGGAAGGCAGCAGG | 18 (0.000341%) | | | ATAGTTTGTTTGATGGTAACTACTACTCGGATAACCGTAGTAATTCTAGAGCTAATACGTGCAACAAACCCCGACTTATGGAAGGGACGCATTTATTAGA | 6 (0.000114%) | | | ATATGACTTGATGTCATGTGTATGATTGAGTATAAGAACT | 154 (0.001165%) | | | ATATGCTTGTCTCAAAGATTAAGCCATGCATGTGTAAGTATGAACGAATTCAGACTGTGAAACTGCGAATGGCTCATTAAATCAGTTATAGTTTGTTTGA | 57 (0.001078%) | | | ATATGGACTTTGGCTACACCATGAAAGCTTTGAGAAGCAAGAAGAAGGTTGGTTAGTGTTTTGGAGTCGAATATGACTTGATGTCATGTGTATGATTGAG | 26 (0.000492%) | | | ATATTCGACTCCAAAACACTAACCAACCTTCTTCTTGCTTCTCAAAGCTTTCATGGTGTAGCCAAAGTCCATATGAGTCTTTGGCTTTGTGTCTTCTAAC | 164 (0.003103%) | | | ATCAAACGAGCTTTTACCCTTTTGTTCCACACGAGATTTCTGTTCTCGTTGAGCTCATCTTAGGACACCTGCGTTATCTTTTAACAGATGTGCCGCCCCA | 7 (0.000132%) | | | ATCAAGTCATATTCGACTCCAAAACACTAACCAACCTTCTTCTTGCTTCTCAAAGCTTTCATGGTGTAGCCAAAGTCCATATGAGTCTTTGGCTTTGTGT | 1462 (0.027658%) | | | ATCAAGTTCACCACCGCATGTCGGTACGCTCCAGGCGTCCTTGGCTCGGATTTAGGCCAACCGCGTGCGGTAACACACGGGAGACCAGCTTCCGTCCCGC | 106 (0.002005%) | | | ATCACGGCAATTCCCCGCCACATCCTCTCAAACGCAATGGAAAGAGAGAAAGGACGAGGTCTTGACCGTCATCTTTTGCCCGAAGGACGGATGAGCTTTG | 8 (0.000151%) | | | ATCAGAATCAAACGAGCTTTTACCCTTTTGTTCCACACGAGATTTCTGTTCTCGTTGAGCTCATCTTAGGACACCTGCGTTATCTTTTAACAGATGTGCC | 12 (0.000227%) | | | ATCAGAGCAACGGGCAGAGCCCGCGTCGACCTTTTATCTAATAAATGCGTCCCTTCCATAAGTCGGGGTTTGTTGCACGTATTAGCTCTAGAATTACTAC | 51 (0.000965%) | | | ATCAGCGTGCTCCGGGCGTCGGCCTGTGGGCTCCCCATTCGACCCGTCTTGAAACACGGACCAAGGAGTCTGACATGTGTGCGAGTCAACGGGTGAGTAA | 7 (0.000132%) | | | ATCAGTTATAGTTTGTTTGATGGTAACTACTACTCGGATAACCGTAGTAATTCTAGAGCTAATACGTGCAACAAACCCCGACTTATGGAAGGGACGCATT | 6 (0.000114%) | | | ATCATAAGGATACTAAATCCTATTTTCTGGTAAATTTTCATAATTTTTTGACACCTCTAGCTAGGTCATTTGACCTGATACAACATCGGATTTTCATGGT | 3 (0.000057%) | | | ATCATACACATGACAACAAGTCATATTCGACTCCAAAACACTAACCAACCTTCTTCTTGCTTCTCAAAGCTTTCATGGTGTAGCCAAAGTCCATATGAGT | 4 (0.000076%) | | | ATCATACACATGACATCAAGTCATATTCGACTCCAAAACACTAACCAACCTTCTTCTTGCTTCTCAAAGCTTTCATGGTGTAGCCAAAGTCCATATGAGT | 30 (0.000568%) | | | ATCATCAGAGCAACGGGCAGAGCCCGCGTCGACCTTTTATCTAATAAATGCGTCCCTTCCATAAGTCGGGGTTTGTTGCACGTATTAGCTCTAGAATTAC | 8 (0.000151%) | | | ATCATGAATCATCAGAGCAACGGGCAGAGCCCGCGTCGACCTTTTATCTAATAAATGCGTCCCTTCCATAAGTCGGGGTTTGTTGCACGTATTAGCTCTA | 75 (0.001419%) | | | ATCATTCAATCGGTAGGAGCGACGGGCGGTGTGTACAAAGGGCAGGGACGTAGTCAACGCGAGCTGATGACTCGCGCTTACTAGGAATTCCTCGTTGAAG | 20 (0.000378%) | | | ATCCAACCTAGGCGAGACAAGGGTTCACATTTCGTTCATCACCCTTGGCCGGCTATCGAACAGCCGGACTCCCATCAAAAGATGGTTGCCAAGAACATCT | 23 (0.000435%) | | | ATCCAACCTAGGCGAGACAAGGGTTCACATTTCGTTCATCACCCTTGGCCGGCTTTCGAACAGCCGGACTCCCATCAAAAGATGGTTGCCAAGAACATCT | 24 (0.000454%) | | | ATCCAGAGCGTAGGCTTGCTTTGAGCACTCTAATTTCTTCAAAGTAACAGCGCCGGAGGCACGACCCGGCCAATTAAGACCAGGAGCGTATCGCCGACCG | 13 (0.000246%) | | | ATCCATGCTTTCCAACGAAGCACGCCCATCCAACCTAGGCGAGACAAGGGTTCACATTTCGTTCATCACCCTTGGCCGGCTATCGAACAGCCGGACTCCC | 7 (0.000132%) | | | ATCCATTGGAGGGCAAGTCTGGTGCCAGCAGCCGCGGTAATTCCAGCTCCAATAGCGTATATTTAAGTTGTTGCAGTTAAAAAGCTCGTAGTTGAACCTT | 29 (0.000549%) | | | ATCCCATGCTAATGTATCCAGAGCGTAGGCTTGCTTTGAGCACTCTAATTTCTTCAAAGTAACAGCGCCGGAGGCACGACCCGGCCAATTAAGACCAGGA | 13 (0.000246%) | | | ATCCGCCTAGGCTGTCCCGAGTGTGAGCGAGGTGTGAGTGTCGCCCATGGGCATCGACACCTTGCGGCTAGGAACTGGAACGAGACGGGTAGCAAAGATT | 60 (0.001135%) | | | ATCCGCCTAGGCTGTCCCGAGTGTGAGCGAGGTGTGAGTGTCGCCCATGGGCATCGACACCTTGCGGCTAGGAACTGGAACGAGACGGGTGGCAAAGATT | 114 (0.002157%) | | | ATCCGCTAAGGAGTGTGTAACAACTCACCTGCCGAATCAACTAGCCCCGAAAATGGATGGCGCTTAAGCGCGCGACCTATACCCGGCCGTCGGGGCAAGA | 8 (0.000151%) | | | ATCCGGTTAAAATTCCGGAACCGGGACGTGGCGGTTGACGGCAACGTTAGGGAGTCCGGAGACGTCGGCGGGGGCCTCGGGAAGAGTTATCTTTTCTGTT | 19 (0.000359%) | | | ATCCGTAACTTCGGGAAAAGGATTGGCTCTGAGGGCTGGGCTCGGGGGTCCCAGTTCCGAACCCGTCGGCTGTCAGCGGACTGCTCGAGCTGCTTCCGCG | 4 (0.000076%) | | | ATCCGTCGAGTTATCATGAATCATCAGAGCAACGGGCAGAGCCCGCGTCGACCTTTTATCTAATAAATGCGTCCCTTCCATAAGTCGGGGTTTGTTGCAC | 7 (0.000132%) | | | ATCCTATGATGTTATCCCATGCTAATGTATCCAGAGCGTAGGCTTGCTTTGAGCACTCTAATTTCTTCAAAGTAACAGCGCCGGAGGCACGACCCGGCCA | 62 (0.001173%) | | | ATCCTCGTTAAGGGATTTAGATTGTACTCATTCCAATTACCAGACTCGAAAGAGCCCGGTATTGTTATTTATTGTCACTACCTCCCCGTGTCAGGATTGG | 12 (0.000227%) | | | ATCCTCTCAAACGCAATGGAAAGAGAGAAAGGACGAGGTCTTGACCGTCATCTTTTGCCCGAAGGACGGATGAGCTTTGGCGGGACTGAATCACTTCGAG | 3 (0.000057%) | | | ATCCTTGTTAGAAGACACAAAGCCAAAGACTCATATGGACTTTGGCTACACCATGAAAGCTTTGAGAAGCAAGAAGAAGGTTGGTTAGTGTTTTGGAGTC | 6 (0.000114%) | | | ATCCTTGTTAGTTTCTTTTCCTCCGCTTATTGATATGCTTAAACTCAGCGGGTAATCCCGCCTGACCTGGGGTCGCTATATGGACTTTGGGTCATCTACA | 7 (0.000132%) | | | ATCCTTTGCTGATGCGGGACGGAAGCTGGTCTCCCGTGTGTTACCGCACGCGGTTGGCCTAAATCCGAGCCAAGGACGCCTGGAGCGTACCGACATGCGG | 6 (0.000114%) | | | ATCGAAATCCTATGATGTTATCCCATGCTAATGTATCCAGAGCGTAGGCTTGCTTTGAGCACTCTAATTTCTTCAAAGTAACAGCGCCGGAGGCACGACC | 8 (0.000151%) | | | ATCGACACCTTGCGGCTAGGAACTGGAACGAGACGGGTGGCAAAGATTTCGAGTAGCACTTCATACTACCGTGGGTTTTTTAAACCTTCCGAGTTTTGTT | 13 (0.000246%) | | | ATCGACCAGAGGCTGTTCACCTTGGAGACCTGATGCGGTTATGAGTACGACCGGGCGTGAGCGGCACTCGGTCCTCCGGATTTTCAAGGGCCGCCGGGGG | 34 (0.000643%) | | | ATCGATCAAGTTCACCACCGCATGTCGGTACGCTCCAGGCGTCCTTGGCTCGGATTTAGGCCAACCGCGTGCGGTAACACACGGGAGACCAGCTTCCGTC | 5 (0.000095%) | | | ATCGATCACGGCAATTCCCCGCCACATCCTCTCAAACGCAATGGAAAGAGAGAAAGGACGAGGTCTTGACCGTCATCTTTTGCCCGAAGGACGGATGAGC | 131 (0.002478%) | | | ATCGATCCATGCTTTCCAACGAAGCACGCCCATCCAACCTAGGCGAGACAAGGGTTCACATTTCGTTCATCACCCTTGGCCGGCTATCGAACAGCCGGAC | 38 (0.000719%) | | | ATCGATCCGCCTAGGCTGTCCCGAGTGTGAGCGAGGTGTGAGTGTCGCCCATGGGCATCGACACCTTGCGGCTAGGAACTGGAACGAGACGGGTAGCAAA | 12 (0.000227%) | | | ATCGATCCGCCTAGGCTGTCCCGAGTGTGAGCGAGGTGTGAGTGTCGCCCATGGGCATCGACACCTTGCGGCTAGGAACTGGAACGAGACGGGTGGCAAA | 24 (0.000454%) | | | ATCGATCCGGTTAAAATTCCGGAACCGGGACGTGGCGGTTGACGGCAACGTTAGGGAGTCCGGAGACGTCGGCGGGGGCCTCGGGAAGAGTTATCTTTTC | 17 (0.000322%) | | | ATCGATCCGTAACTTCGGGAAAAGGATTGGCTCTGAGGGCTGGGCTCGGGGGTCCCAGTTCCGAACCCGTCGGCTGTCAGCGGACTGCTCGAGCTGCTTC | 36 (0.000681%) | | | ATCGATCCGTCGAGTTATCATGAATCATCAGAGCAACGGGCAGAGCCCGCGTCGACCTTTTATCTAATAAATGCGTCCCTTCCATAAGTCGGGGTTTGTT | 17 (0.000322%) | | | ATCGATCCTCGTTAAGGGATTTAGATTGTACTCATTCCAATTACCAGACTCGAAAGAGCCCGGTATTGTTATTTATTGTCACTACCTCCCCGTGTCAGGA | 45 (0.000851%) | | | ATCGATCGAAATCCTATGATGTTATCCCATGCTAATGTATCCAGAGCGTAGGCTTGCTTTGAGCACTCTAATTTCTTCAAAGTAACAGCGCCGGAGGCAC | 11 (0.000208%) | | | ATCGATCGGCGGACCGGATTGCTCCGTTCCGCATCCGACCAGGACGCATCGCCGGCCCCCATCCGCTTCCCTCCCGACAATTTCAAGCACTCTTTGACTC | 18 (0.000341%) | | | ATCGATCGGGTTGCGGTTTAAGTTGTTATACTCAATCATA | 23 (0.000174%) | | | ATCGGAAGAGCACACGTCTGAACTCCAGTCACGAGATTCC | 43 (0.000325%) | | | ATCGGCGGACCGGATTGCTCCGTTCCGCATCCGACCAGGACGCATCGCCGGCCCCCATCCGCTTCCCTCCCGACAATTTCAAGCACTCTTTGACTCTCTT | 5 (0.000095%) | | | ATCGGGTTGCGGTTTAAGTTGTTATACTCAATCATACACA | 132 (0.000999%) | | | ATCGGTAGGAGCGACGGGCGGTGTGTACAAAGGGCAGGGACGTAGTCAACGCGAGCTGATGACTCGCGCTTACTAGGAATTCCTCGTTGAAGACCAACAA | 74 (0.001400%) | | | ATCTAATAAATGCGTCCCTTCCATAAGTCGGGGTTTGTTGCACGTATTAGCTCTAGAATTACTACGGTTATCCGAGTAGTAGTTACCATCAAACAAACTA | 5 (0.000095%) | | | ATCTGTTAAAAGATAACGCAGGTGTCCTAAGATGAGCTCAACGAGAACAGAAATCTCGTGTGGAACAAAAGGGTAAAAGCTCGTTTGATTCTGATTTTCA | 6 (0.000114%) | | | ATCTTAAAAGCCTAAGTAGTGTTTCCTTGTTAGAAGACACAAAGCCAAAGACTCATATGGACTTTGGCTACACCATGAAAGCTTTGAGAAGCAAGAAGAA | 593 (0.011219%) | | | ATCTTAAAGGCGTAAGAATTGTATCCTTGTTAAAAGACACAAAGCCAAAGACTCATATGGACTTTGGCTACACCATGAAAGCTTTGAGAAGCAAGAAGAA | 43 (0.000813%) | | | ATCTTAAAGGCGTAAGAATTGTATCCTTGTTAGAAGACACAAAGCCAAAGACTCATATGGACTTTGGCTACACCATGAAAGCTTTGAGAAGCAAGAAGAA | 28 (0.000530%) | | | ATGAATCATCAGAGCAACGGGCAGAGCCCGCGTCGACCTTTTATCTAATAAATGCGTCCCTTCCATAAGTCGGGGTTTGTTGCACGTATTAGCTCTAGAA | 82 (0.001551%) | | | ATGACATCAAGTCATATTCGACTCCAAAACACTAACCAACCTTCTTCTTGCTTCTCAAAGCTTTCATGGTGTAGCCAAAGTCCATATGAGTCTTTGGCTT | 117 (0.002213%) | | | ATGACTTGATGTCATGTGTATGATTGAGTATAAGAACTTA | 285 (0.002157%) | | | ATGATGTTATCCCATGCTAATGTATCCAGAGCGTAGGCTTGCTTTGAGCACTCTAATTTCTTCAAAGTAACAGCGCCGGAGGCACGACCCGGCCAATTAA | 66 (0.001249%) | | | ATGATTAACAGGGACAGTCGGGGGCATTCGTATTTCATAGTCAGAGGTGAAATTCTTGGATTTATGAAAGACGAACAACTGCGAAAGCATTTGCCAAGGA | 100 (0.001892%) | | | ATGCATCATAAGGATACTAAATCCTATTTTCTGGTAAATTTTCATAATTTTTTGACACCTCTAGCTAGGTCATTTGACCTGATACAACATCGGATTTTCA | 7 (0.000132%) | | | ATGCCAGCCGTTCGTTTGCATGTTCCTTGACACTTTTCGTGCCGGGGTTTTGTGATATCCGGAAGCAACGCGCACGACAAGACCGAGATAAAAGCTCCCG | 11 (0.000208%) | | | ATGCCCTTAGATGTTCTGGGCCGCACGCGCGCTACACTGATGTATTCAACGAGTTCACACCTTGGCCGACAGGCCCGGGTAATCTTTGAAATTTCATCGT | 6 (0.000114%) | | | ATGCTAATGTATCCAGAGCGTAGGCTTGCTTTGAGCACTCTAATTTCTTCAAAGTAACAGCGCCGGAGGCACGACCCGGCCAATTAAGACCAGGAGCGTA | 14 (0.000265%) | | | ATGCTTGTCTCAAAGATTAAGCCATGCATGTGTAAGTATGAACGAATTCAGACTGTGAAACTGCGAATGGCTCATTAAATCAGTTATAGTTTGTTTGATG | 16 (0.000303%) | | | ATGCTTTCCAACGAAGCACGCCCATCCAACCTAGGCGAGACAAGGGTTCACATTTCGTTCATCACCCTTGGCCGGCTATCGAACAGCCGGACTCCCATCA | 297 (0.005619%) | | | ATGGACTTTGGCTACACCATGAAAGCTTTGAGAAGCAAGAAGAAGGTTGGTTAGTGTTTTGGAGTCGAATATGACTTGATGTCATGTGTATGATTGAGTA | 81 (0.001532%) | | | ATGGCTCATTAAATCAGTTATAGTTTGTTTGATGGTAACTACTACTCGGATAACCGTAGTAATTCTAGAGCTAATACGTGCAACAAACCCCGACTTATGG | 8 (0.000151%) | | | ATGGGCATCGACACCTTGCGGCTAGGAACTGGAACGAGACGGGTGGCAAAGATTTCGAGTAGCACTTCATACTACCGTGGGTTTTTTAAACCTTCCGAGT | 10 (0.000189%) | | | ATGTATCCAGAGCGTAGGCTTGCTTTGAGCACTCTAATTTCTTCAAAGTAACAGCGCCGGAGGCACGACCCGGCCAATTAAGACCAGGAGCGTATCGCCG | 3 (0.000057%) | | | ATGTCATGTGTATGATTGAGTATAAGAACTTAAACCGCAA | 125 (0.000946%) | | | ATGTCGGTACGCTCCAGGCGTCCTTGGCTCGGATTTAGGCCAACCGCGTGCGGTAACACACGGGAGACCAGCTTCCGTCCCGCATCAGCAAAGGATGGTG | 4 (0.000076%) | | | ATGTGTAAGTATGAACGAATTCAGACTGTGAAACTGCGAATGGCTCATTAAATCAGTTATAGTTTGTTTGATGGTAACTACTACTCGGATAACCGTAGTA | 10 (0.000189%) | | | ATGTTATCCCATGCTAATGTATCCAGAGCGTAGGCTTGCTTTGAGCACTCTAATTTCTTCAAAGTAACAGCGCCGGAGGCACGACCCGGCCAATTAAGAC | 17 (0.000322%) | | | ATGTTCTGGGCCGCACGCGCGCTACACTGATGTATTCAACGAGTTCACACCTTGGCCGACAGGCCCGGGTAATCTTTGAAATTTCATCGTGATGGGGATA | 5 (0.000095%) | | | ATTAAATCAGTTATAGTTTGTTTGATGGTAACTACTACTCGGATAACCGTAGTAATTCTAGAGCTAATACGTGCAACAAACCCCGACTTATGGAAGGGAC | 38 (0.000719%) | | | ATTAACAGGGACAGTCGGGGGCATTCGTATTTCATAGTCAGAGGTGAAATTCTTGGATTTATGAAAGACGAACAACTGCGAAAGCATTTGCCAAGGATGT | 102 (0.001930%) | | | ATTAAGCCATGCATGTGTAAGTATGAACGAATTCAGACTGTGAAACTGCGAATGGCTCATTAAATCAGTTATAGTTTGTTTGATGGTAACTACTACTCGG | 19 (0.000359%) | | | ATTACCAGACTCGAAAGAGCCCGGTATTGTTATTTATTGTCACTACCTCCCCGTGTCAGGATTGGGTAATTTGCGCGCCTGCTGCCTTCCTTGGATGTGG | 5 (0.000095%) | | | ATTAGGGTTCGATTCCGGAGAGGGAGCCTGAGAAACGGCTACCACATCCAAGGAAGGCAGCAGGCGCGCAAATTACCCAATCCTGACACGGGGAGGTAGT | 6 (0.000114%) | | | ATTCAATCGGTAGGAGCGACGGGCGGTGTGTACAAAGGGCAGGGACGTAGTCAACGCGAGCTGATGACTCGCGCTTACTAGGAATTCCTCGTTGAAGACC | 339 (0.006413%) | | | ATTCCAATTACCAGACTCGAAAGAGCCCGGTATTGTTATTTATTGTCACTACCTCCCCGTGTCAGGATTGGGTAATTTGCGCGCCTGCTGCCTTCCTTGG | 23 (0.000435%) | | | ATTCCAGCTCCAATAGCGTATATTTAAGTTGTTGCAGTTAAAAAGCTCGTAGTTGAACCTTGGGATGGGTCGGCCGGTCCGCCTTTGGTGTGCATTGGTC | 5 (0.000095%) | | | ATTCCCCGCCACATCCTCTCAAACGCAATGGAAAGAGAGAAAGGACGAGGTCTTGACCGTCATCTTTTGCCCGAAGGACGGATGAGCTTTGGCGGGACTG | 10 (0.000189%) | | | ATTCGACTCCAAAACACTAACCAACCTTCTTCTTGCTTCTCAAAGCTTTCATGGTGTAGCCAAAGTCCATATGAGTCTTTGGCTTTGTGTCTTCTAACAA | 112 (0.002119%) | | | ATTCGTACTGAAAATCAGAATCAAACGAGCTTTTACCCTTTTGTTCCACACGAGATTTCTGTTCTCGTTGAGCTCATCTTAGGACACCTGCGTTATCTTT | 10 (0.000189%) | | | ATTCGTATTTCATAGTCAGAGGTGAAATTCTTGGATTTATGAAAGACGAACAACTGCGAAAGCATTTGCCAAGGATGTTTTCATTAATCAAGAACGAAAG | 8 (0.000151%) | | | ATTGGAGGGCAAGTCTGGTGCCAGCAGCCGCGGTAATTCCAGCTCCAATAGCGTATATTTAAGTTGTTGCAGTTAAAAAGCTCGTAGTTGAACCTTGGGA | 205 (0.003878%) | | | ATTGTACTCATTCCAATTACCAGACTCGAAAGAGCCCGGTATTGTTATTTATTGTCACTACCTCCCCGTGTCAGGATTGGGTAATTTGCGCGCCTGCTGC | 8 (0.000151%) | | | ATTGTATCCTTGTTAGAAGACACAAAGCCAAAGACTCATATGGACTTTGGCTACACCATGAAAGCTTTGAGAAGCAAGAAGAAGGTTGGTTAGTGTTTTG | 38 (0.000719%) | | | ATTGTCAGGTGGGGAGTTTGGCTGGGGCGGCACATCTGTTAAAAGATAACGCAGGTGTCCTAAGATGAGCTCAACGAGAACAGAAATCTCGTGTGGAACA | 38 (0.000719%) | | | ATTGTTCCATCGACCAGAGGCTGTTCACCTTGGAGACCTGATGCGGTTATGAGTACGACCGGGCGTGAGCGGCACTCGGTCCTCCGGATTTTCAAGGGCC | 13 (0.000246%) | | | ATTGTTGGTCTTCAACGAGGAATTCCTAGTAAGCGCGAGTCATCAGCTCGCGTTGACTACGTCCCTGCCCTTTGTACACACCGCCCGTCGCTCCTACCGA | 40 (0.000757%) | | | ATTTAGATTGTACTCATTCCAATTACCAGACTCGAAAGAGCCCGGTATTGTTATTTATTGTCACTACCTCCCCGTGTCAGGATTGGGTAATTTGCGCGCC | 18 (0.000341%) | | | ATTTCATAGTCAGAGGTGAAATTCTTGGATTTATGAAAGACGAACAACTGCGAAAGCATTTGCCAAGGATGTTTTCATTAATCAAGAACGAAAGTTGGGG | 10 (0.000189%) | | | ATTTCGTTCATCACCCTTGGCCGGCTATCGAACAGCCGGACTCCCATCAAAAGATGGTTGCCAAGAACATCTTCGTTACGGTTTGCTAATTCTCGGAATA | 24 (0.000454%) | | | ATTTTCATAATTTTTTGACACCTCTAGCTAGGTCATTTGACCTGATACAACATCGGATTTTCATGGTCTAGTTGGGGCTCCGTGGGCATATTTGATGCAA | 4 (0.000076%) | | | ATTTTCTGGTAAATTTTCATAATTTTTTGACACCTCTAGCTAGGTCATTTGACCTGATACAACATCGGATTTTCATGGTCTAGTTGGGGCTCCGTGGGCA | 8 (0.000151%) | | | CAAAAAGCAACGTCGCTATGAACGCTTGGCTGCCACAAGCCAGTTATCCCTGTGGTAACTTTTCTGACACCTCTAGCTTCAAATTCCGAAGGTCTAAAGG | 1357 (0.025672%) | | | CAAACGCAATGGAAAGAGAGAAAGGACGAGGTCTTGACCGTCATCTTTTGCCCGAAGGACGGATGAGCTTTGGCGGGACTGAATCACTTCGAGTCACCGT | 6 (0.000114%) | | | CAAAGACTCATATGGACTTTGGCTACACCATGAAAGCTTTGAGAAGCAAGAAGAAGGTTGGTTAGTGTTTTGGAGTCGAATATGACTTGATGTCATGTGT | 119 (0.002251%) | | | CAAAGATTAAGCCATGCATGTGTAAGTATGAACGAATTCAGACTGTGAAACTGCGAATGGCTCATTAAATCAGTTATAGTTTGTTTGATGGTAACTACTA | 64 (0.001211%) | | | CAAAGCCAAAGACTCATATGGACTTTGGCTACACCATGAAAGCTTTGAGAAGCAAGAAGAAGGTTGGTTAGTGTTTTGGAGTCGAATATGACTTGATGTC | 108 (0.002043%) | | | CAACCTAGGCGAGACAAGGGTTCACATTTCGTTCATCACCCTTGGCCGGCTATCGAACAGCCGGACTCCCATCAAAAGATGGTTGCCAAGAACATCTTCG | 9 (0.000170%) | | | CAACCTAGGCGAGACAAGGGTTCACATTTCGTTCATCACCCTTGGCCGGCTTTCGAACAGCCGGACTCCCATCAAAAGATGGTTGCCAAGAACATCTTCG | 5 (0.000095%) | | | CAACGAAGCACGCCCATCCAACCTAGGCGAGACAAGGGTTCACATTTCGTTCATCACCCTTGGCCGGCTATCGAACAGCCGGACTCCCATCAAAAGATGG | 67 (0.001268%) | | | CAACGAAGCACGCCCATCCAACCTAGGCGAGACAAGGGTTCACATTTCGTTCATCACCCTTGGCCGGCTTTCGAACAGCCGGACTCCCATCAAAAGATGG | 86 (0.001627%) | | | CAACGAGGAATTCCTAGTAAGCGCGAGTCATCAGCTCGCGTTGACTACGTCCCTGCCCTTTGTACACACCGCCCGTCGCTCCTACCGATTGAATGATCGA | 6 (0.000114%) | | | CAACGGGCAGAGCCCGCGTCGACCTTTTATCTAATAAATGCGTCCCTTCCATAAGTCGGGGTTTGTTGCACGTATTAGCTCTAGAATTACTACGGTTATC | 19 (0.000359%) | | | CAACGTCGCTATGAACGCTTGGCTGCCACAAGCCAGTTATCCCTGTGGTAACTTTTCTGACACCTCTAGCTTCAAATTCCGAAGGTCTAAAGGATCGATC | 22 (0.000416%) | | | CAAGGGTTCACATTTCGTTCATCACCCTTGGCCGGCTATCGAACAGCCGGACTCCCATCAAAAGATGGTTGCCAAGAACATCTTCGTTACGGTTTGCTAA | 22 (0.000416%) | | | CAAGGGTTCACATTTCGTTCATCACCCTTGGCCGGCTTTCGAACAGCCGGACTCCCATCAAAAGATGGTTGCCAAGAACATCTTCGTTACGGTTTGCTAA | 35 (0.000662%) | | | CAAGTCATATTCGACTCCAAAACACTAACCAACCTTCTTCTTGCTTCTCAAAGCTTTCATGGTGTAGCCAAAGTCCATATGAGTCTTTGGCTTTGTGTCT | 142 (0.002686%) | | | CAAGTCTGGTGCCAGCAGCCGCGGTAATTCCAGCTCCAATAGCGTATATTTAAGTTGTTGCAGTTAAAAAGCTCGTAGTTGAACCTTGGGATGGGTCGGC | 24 (0.000454%) | | | CAAGTTCACCACCGCATGTCGGTACGCTCCAGGCGTCCTTGGCTCGGATTTAGGCCAACCGCGTGCGGTAACACACGGGAGACCAGCTTCCGTCCCGCAT | 18 (0.000341%) | | | CAATAACAGGTCTGTGATGCCCTTAGATGTTCTGGGCCGCACGCGCGCTACACTGATGTATTCAACGAGTTCACACCTTGGCCGACAGGCCCGGGTAATC | 36 (0.000681%) | | | CAATCATACACATGACAACAAGTCATATTCGACTCCAAAACACTAACCAACCTTCTTCTTGCTTCTCAAAGCTTTCATGGTGTAGCCAAAGTCCATATGA | 24 (0.000454%) | | | CAATCATACACATGACATCAAGTCATATTCGACTCCAAAACACTAACCAACCTTCTTCTTGCTTCTCAAAGCTTTCATGGTGTAGCCAAAGTCCATATGA | 178 (0.003367%) | | | CAATCGGTAGGAGCGACGGGCGGTGTGTACAAAGGGCAGGGACGTAGTCAACGCGAGCTGATGACTCGCGCTTACTAGGAATTCCTCGTTGAAGACCAAC | 107 (0.002024%) | | | CAATTACCAGACTCGAAAGAGCCCGGTATTGTTATTTATTGTCACTACCTCCCCGTGTCAGGATTGGGTAATTTGCGCGCCTGCTGCCTTCCTTGGATGT | 26 (0.000492%) | | | CAATTCCCCGCCACATCCTCTCAAACGCAATGGAAAGAGAGAAAGGACGAGGTCTTGACCGTCATCTTTTGCCCGAAGGACGGATGAGCTTTGGCGGGAC | 35 (0.000662%) | | | CACAAAGCCAAAGACTCATATGGACTTTGGCTACACCATGAAAGCTTTGAGAAGCAAGAAGAAGGTTGGTTAGTGTTTTGGAGTCGAATATGACTTGATC | 15 (0.000284%) | | | CACAAAGCCAAAGACTCATATGGACTTTGGCTACACCATGAAAGCTTTGAGAAGCAAGAAGAAGGTTGGTTAGTGTTTTGGAGTCGAATATGACTTGATG | 40 (0.000757%) | | | CACATCCTCTCAAACGCAATGGAAAGAGAGAAAGGACGAGGTCTTGACCGTCATCTTTTGCCCGAAGGACGGATGAGCTTTGGCGGGACTGAATCACTTC | 5 (0.000095%) | | | CACATCTGTTAAAAGATAACGCAGGTGTCCTAAGATGAGCTCAACGAGAACAGAAATCTCGTGTGGAACAAAAGGGTAAAAGCTCGTTTGATTCTGATTT | 7 (0.000132%) | | | CACATGACATCAAGTCATATTCGACTCCAAAACACTAACCAACCTTCTTCTTGCTTCTCAAAGCTTTCATGGTGTAGCCAAAGTCCATATGAGTCTTTGG | 49 (0.000927%) | | | CACATTTCGTTCATCACCCTTGGCCGGCTATCGAACAGCCGGACTCCCATCAAAAGATGGTTGCCAAGAACATCTTCGTTACGGTTTGCTAATTCTCGGA | 5 (0.000095%) | | | CACATTTCGTTCATCACCCTTGGCCGGCTTTCGAACAGCCGGACTCCCATCAAAAGATGGTTGCCAAGAACATCTTCGTTACGGTTTGCTAATTCTCGGA | 4 (0.000076%) | | | CACCACCGCATGTCGGTACGCTCCAGGCGTCCTTGGCTCGGATTTAGGCCAACCGCGTGCGGTAACACACGGGAGACCAGCTTCCGTCCCGCATCAGCAA | 29 (0.000549%) | | | CACCATCCTTTGCTGATGCGGGACGGAAGCTGGTCTCCCGTGTGTTACCGCACGCGGTTGGCCTAAATCCGAGCCAAGGACGCCTGGAGCGTACCGACAT | 7 (0.000132%) | | | CACCCGTTGACTCGCACACATGTCAGACTCCTTGGTCCGTGTTTCAAGACGGGTCGAATGGGGAGCCCACAGGCCGACGCCCGGAGCACGCTGATGCCGA | 12 (0.000227%) | | | CACCCTAACGCCTCGAAGAACTAATGGCAGCCCACGCAAGGCAAGCCCATTCTCCTCGACGATTCAGCAGTTTTTGTCCGAGAACTGCTGAGAAAACTCG | 8 (0.000151%) | | | CACGCCCATCCAACCTAGGCGAGACAAGGGTTCACATTTCGTTCATCACCCTTGGCCGGCTATCGAACAGCCGGACTCCCATCAAAAGATGGTTGCCAAG | 26 (0.000492%) | | | CACGCCCATCCAACCTAGGCGAGACAAGGGTTCACATTTCGTTCATCACCCTTGGCCGGCTTTCGAACAGCCGGACTCCCATCAAAAGATGGTTGCCAAG | 26 (0.000492%) | | | CACGCGCCTAACGGCGTGCCTCGGCATCAGCGTGCTCCGGGCGTCGGCCTGTGGGCTCCCCATTCGACCCGTCTTGAAACACGGACCAAGGAGTCTGACA | 103 (0.001949%) | | | CACGCTTTCACGGTTCGTATTCGTACTGAAAATCAGAATCAAACGAGCTTTTACCCTTTTGTTCCACACGAGATTTCTGTTCTCGTTGAGCTCATCTTAG | 103 (0.001949%) | | | CACGGCAATTCCCCGCCACATCCTCTCAAACGCAATGGAAAGAGAGAAAGGACGAGGTCTTGACCGTCATCTTTTGCCCGAAGGACGGATGAGCTTTGGC | 11 (0.000208%) | | | CACGGTTCGTATTCGTACTGAAAATCAGAATCAAACGAGCTTTTACCCTTTTGTTCCACACGAGATTTCTGTTCTCGTTGAGCTCATCTTAGGACACCTG | 4 (0.000076%) | | | CAGAATCAAACGAGCTTTTACCCTTTTGTTCCACACGAGATTTCTGTTCTCGTTGAGCTCATCTTAGGACACCTGCGTTATCTTTTAACAGATGTGCCGC | 17 (0.000322%) | | | CAGACTCGAAAGAGCCCGGTATTGTTATTTATTGTCACTACCTCCCCGTGTCAGGATTGGGTAATTTGCGCGCCTGCTGCCTTCCTTGGATGTGGTAGCC | 12 (0.000227%) | | | CAGAGCAACGGGCAGAGCCCGCGTCGACCTTTTATCTAATAAATGCGTCCCTTCCATAAGTCGGGGTTTGTTGCACGTATTAGCTCTAGAATTACTACGG | 11 (0.000208%) | | | CAGAGCCCGCGTCGACCTTTTATCTAATAAATGCGTCCCTTCCATAAGTCGGGGTTTGTTGCACGTATTAGCTCTAGAATTACTACGGTTATCCGAGTAG | 13 (0.000246%) | | | CAGAGCGTAGGCTTGCTTTGAGCACTCTAATTTCTTCAAAGTAACAGCGCCGGAGGCACGACCCGGCCAATTAAGACCAGGAGCGTATCGCCGACCGAAG | 29 (0.000549%) | | | CAGAGGCTGTTCACCTTGGAGACCTGATGCGGTTATGAGTACGACCGGGCGTGAGCGGCACTCGGTCCTCCGGATTTTCAAGGGCCGCCGGGGGCGCACC | 10 (0.000189%) | | | CAGAGGTGAAATTCTTGGATTTATGAAAGACGAACAACTGCGAAAGCATTTGCCAAGGATGTTTTCATTAATCAAGAACGAAAGTTGGGGGCTCGAAGAC | 21 (0.000397%) | | | CAGCACGCGCCTAACGGCGTGCCTCGGCATCAGCGTGCTCCGGGCGTCGGCCTGTGGGCTCCCCATTCGACCCGTCTTGAAACACGGACCAAGGAGTCTG | 65 (0.001230%) | | | CAGCAGCCGCGGTAATTCCAGCTCCAATAGCGTATATTTAAGTTGTTGCAGTTAAAAAGCTCGTAGTTGAACCTTGGGATGGGTCGGCCGGTCCGCCTTT | 21 (0.000397%) | | | CAGCCGCGGTAATTCCAGCTCCAATAGCGTATATTTAAGTTGTTGCAGTTAAAAAGCTCGTAGTTGAACCTTGGGATGGGTCGGCCGGTCCGCCTTTGGT | 7 (0.000132%) | | | CAGCGTGCTCCGGGCGTCGGCCTGTGGGCTCCCCATTCGACCCGTCTTGAAACACGGACCAAGGAGTCTGACATGTGTGCGAGTCAACGGGTGAGTAAAC | 7 (0.000132%) | | | CAGGGACAGTCGGGGGCATTCGTATTTCATAGTCAGAGGTGAAATTCTTGGATTTATGAAAGACGAACAACTGCGAAAGCATTTGCCAAGGATGTTTTCA | 37 (0.000700%) | | | CAGGTCTGTGATGCCCTTAGATGTTCTGGGCCGCACGCGCGCTACACTGATGTATTCAACGAGTTCACACCTTGGCCGACAGGCCCGGGTAATCTTTGAA | 15 (0.000284%) | | | CAGGTGGGGAGTTTGGCTGGGGCGGCACATCTGTTAAAAGATAACGCAGGTGTCCTAAGATGAGCTCAACGAGAACAGAAATCTCGTGTGGAACAAAAGG | 31 (0.000586%) | | | CAGTAGTCATATGCTTGTCTCAAAGATTAAGCCATGCATGTGTAAGTATGAACGAATTCAGACTGTGAAACTGCGAATGGCTCATTAAATCAGTTATAGT | 141 (0.002667%) | | | CAGTCGGGGGCATTCGTATTTCATAGTCAGAGGTGAAATTCTTGGATTTATGAAAGACGAACAACTGCGAAAGCATTTGCCAAGGATGTTTTCATTAATC | 5 (0.000095%) | | | CAGTTATAGTTTGTTTGATGGTAACTACTACTCGGATAACCGTAGTAATTCTAGAGCTAATACGTGCAACAAACCCCGACTTATGGAAGGGACGCATTTA | 4 (0.000076%) | | | CATAAGGATACTAAATCCTATTTTCTGGTAAATTTTCATAATTTTTTGACACCTCTAGCTAGGTCATTTGACCTGATACAACATCGGATTTTCATGGTCT | 19 (0.000359%) | | | CATACACATGACATCAAGTCATATTCGACTCCAAAACACTAACCAACCTTCTTCTTGCTTCTCAAAGCTTTCATGGTGTAGCCAAAGTCCATATGAGTCT | 52 (0.000984%) | | | CATAGTCAGAGGTGAAATTCTTGGATTTATGAAAGACGAACAACTGCGAAAGCATTTGCCAAGGATGTTTTCATTAATCAAGAACGAAAGTTGGGGGCTC | 19 (0.000359%) | | | CATATGCTTGTCTCAAAGATTAAGCCATGCATGTGTAAGTATGAACGAATTCAGACTGTGAAACTGCGAATGGCTCATTAAATCAGTTATAGTTTGTTTG | 47 (0.000889%) | | | CATATGGACTTTGGCTACACCATGAAAGCTTTGAGAAGCAAGAAGAAGGTTGGTTAGTGTTTTGGAGTCGAATATGACTTGATGTCATGTGTATGATTGA | 14 (0.000265%) | | | CATATTCGACTCCAAAACACTAACCAACCTTCTTCTTGCTTCTCAAAGCTTTCATGGTGTAGCCAAAGTCCATATGAGTCTTTGGCTTTGTGTCTTCTAA | 92 (0.001740%) | | | CATCAAGTCATATTCGACTCCAAAACACTAACCAACCTTCTTCTTGCTTCTCAAAGCTTTCATGGTGTAGCCAAAGTCCATATGAGTCTTTGGCTTTGTG | 51 (0.000965%) | | | CATCAGAGCAACGGGCAGAGCCCGCGTCGACCTTTTATCTAATAAATGCGTCCCTTCCATAAGTCGGGGTTTGTTGCACGTATTAGCTCTAGAATTACTA | 33 (0.000624%) | | | CATCAGCGTGCTCCGGGCGTCGGCCTGTGGGCTCCCCATTCGACCCGTCTTGAAACACGGACCAAGGAGTCTGACATGTGTGCGAGTCAACGGGTGAGTA | 11 (0.000208%) | | | CATCCAACCTAGGCGAGACAAGGGTTCACATTTCGTTCATCACCCTTGGCCGGCTATCGAACAGCCGGACTCCCATCAAAAGATGGTTGCCAAGAACATC | 5 (0.000095%) | | | CATCCAACCTAGGCGAGACAAGGGTTCACATTTCGTTCATCACCCTTGGCCGGCTTTCGAACAGCCGGACTCCCATCAAAAGATGGTTGCCAAGAACATC | 5 (0.000095%) | | | CATCCTCTCAAACGCAATGGAAAGAGAGAAAGGACGAGGTCTTGACCGTCATCTTTTGCCCGAAGGACGGATGAGCTTTGGCGGGACTGAATCACTTCGA | 3 (0.000057%) | | | CATCCTTTGCTGATGCGGGACGGAAGCTGGTCTCCCGTGTGTTACCGCACGCGGTTGGCCTAAATCCGAGCCAAGGACGCCTGGAGCGTACCGACATGCG | 134 (0.002535%) | | | CATCGACACCTTGCGGCTAGGAACTGGAACGAGACGGGTGGCAAAGATTTCGAGTAGCACTTCATACTACCGTGGGTTTTTTAAACCTTCCGAGTTTTGT | 4 (0.000076%) | | | CATCGACCAGAGGCTGTTCACCTTGGAGACCTGATGCGGTTATGAGTACGACCGGGCGTGAGCGGCACTCGGTCCTCCGGATTTTCAAGGGCCGCCGGGG | 9 (0.000170%) | | | CATGAATCATCAGAGCAACGGGCAGAGCCCGCGTCGACCTTTTATCTAATAAATGCGTCCCTTCCATAAGTCGGGGTTTGTTGCACGTATTAGCTCTAGA | 165 (0.003122%) | | | CATGACATCAAGTCATATTCGACTCCAAAACACTAACCAACCTTCTTCTTGCTTCTCAAAGCTTTCATGGTGTAGCCAAAGTCCATATGAGTCTTTGGCT | 38 (0.000719%) | | | CATGCATGTGTAAGTATGAACGAATTCAGACTGTGAAACTGCGAATGGCTCATTAAATCAGTTATAGTTTGTTTGATGGTAACTACTACTCGGATAACCG | 7 (0.000132%) | | | CATGCTAATGTATCCAGAGCGTAGGCTTGCTTTGAGCACTCTAATTTCTTCAAAGTAACAGCGCCGGAGGCACGACCCGGCCAATTAAGACCAGGAGCGT | 5 (0.000095%) | | | CATGCTTTCCAACGAAGCACGCCCATCCAACCTAGGCGAGACAAGGGTTCACATTTCGTTCATCACCCTTGGCCGGCTATCGAACAGCCGGACTCCCATC | 11 (0.000208%) | | | CATGGGCATCGACACCTTGCGGCTAGGAACTGGAACGAGACGGGTGGCAAAGATTTCGAGTAGCACTTCATACTACCGTGGGTTTTTTAAACCTTCCGAG | 3 (0.000057%) | | | CATGTCGGTACGCTCCAGGCGTCCTTGGCTCGGATTTAGGCCAACCGCGTGCGGTAACACACGGGAGACCAGCTTCCGTCCCGCATCAGCAAAGGATGGT | 4 (0.000076%) | | | CATTAAATCAGTTATAGTTTGTTTGATGGTAACTACTACTCGGATAACCGTAGTAATTCTAGAGCTAATACGTGCAACAAACCCCGACTTATGGAAGGGA | 14 (0.000265%) | | | CATTCAATCGGTAGGAGCGACGGGCGGTGTGTACAAAGGGCAGGGACGTAGTCAACGCGAGCTGATGACTCGCGCTTACTAGGAATTCCTCGTTGAAGAC | 15 (0.000284%) | | | CATTCCAATTACCAGACTCGAAAGAGCCCGGTATTGTTATTTATTGTCACTACCTCCCCGTGTCAGGATTGGGTAATTTGCGCGCCTGCTGCCTTCCTTG | 7 (0.000132%) | | | CATTCGTATTTCATAGTCAGAGGTGAAATTCTTGGATTTATGAAAGACGAACAACTGCGAAAGCATTTGCCAAGGATGTTTTCATTAATCAAGAACGAAA | 8 (0.000151%) | | | CATTGGAGGGCAAGTCTGGTGCCAGCAGCCGCGGTAATTCCAGCTCCAATAGCGTATATTTAAGTTGTTGCAGTTAAAAAGCTCGTAGTTGAACCTTGGG | 30 (0.000568%) | | | CATTGTCAGGTGGGGAGTTTGGCTGGGGCGGCACATCTGTTAAAAGATAACGCAGGTGTCCTAAGATGAGCTCAACGAGAACAGAAATCTCGTGTGGAAC | 8 (0.000151%) | | | CATTGTTCCATCGACCAGAGGCTGTTCACCTTGGAGACCTGATGCGGTTATGAGTACGACCGGGCGTGAGCGGCACTCGGTCCTCCGGATTTTCAAGGGC | 28 (0.000530%) | | | CCAAAGACTCATATGGACTTTGGCTACACCATGAAAGCTTTGAGAAGCAAGAAGAAGGTTGGTTAGTGTTTTGGAGTCGAATATGACTTGATGTCATGTG | 205 (0.003878%) | | | CCAACCTAGGCGAGACAAGGGTTCACATTTCGTTCATCACCCTTGGCCGGCTATCGAACAGCCGGACTCCCATCAAAAGATGGTTGCCAAGAACATCTTC | 18 (0.000341%) | | | CCAACCTAGGCGAGACAAGGGTTCACATTTCGTTCATCACCCTTGGCCGGCTTTCGAACAGCCGGACTCCCATCAAAAGATGGTTGCCAAGAACATCTTC | 27 (0.000511%) | | | CCAACGAAGCACGCCCATCCAACCTAGGCGAGACAAGGGTTCACATTTCGTTCATCACCCTTGGCCGGCTATCGAACAGCCGGACTCCCATCAAAAGATG | 15 (0.000284%) | | | CCAACGAAGCACGCCCATCCAACCTAGGCGAGACAAGGGTTCACATTTCGTTCATCACCCTTGGCCGGCTTTCGAACAGCCGGACTCCCATCAAAAGATG | 9 (0.000170%) | | | CCAATTACCAGACTCGAAAGAGCCCGGTATTGTTATTTATTGTCACTACCTCCCCGTGTCAGGATTGGGTAATTTGCGCGCCTGCTGCCTTCCTTGGATG | 5 (0.000095%) | | | CCACCCTAACGCCTCGAAGAACTAATGGCAGCCCACGCAAGGCAAGCCCATTCTCCTCGACGATTCAGCAGTTTTTGTCCGAGAACTGCTGAGAAAACTC | 7 (0.000132%) | | | CCACCGCATGTCGGTACGCTCCAGGCGTCCTTGGCTCGGATTTAGGCCAACCGCGTGCGGTAACACACGGGAGACCAGCTTCCGTCCCGCATCAGCAAAG | 5 (0.000095%) | | | CCACGCTTTCACGGTTCGTATTCGTACTGAAAATCAGAATCAAACGAGCTTTTACCCTTTTGTTCCACACGAGATTTCTGTTCTCGTTGAGCTCATCTTA | 67 (0.001268%) | | | CCAGACTCGAAAGAGCCCGGTATTGTTATTTATTGTCACTACCTCCCCGTGTCAGGATTGGGTAATTTGCGCGCCTGCTGCCTTCCTTGGATGTGGTAGC | 12 (0.000227%) | | | CCAGAGCGTAGGCTTGCTTTGAGCACTCTAATTTCTTCAAAGTAACAGCGCCGGAGGCACGACCCGGCCAATTAAGACCAGGAGCGTATCGCCGACCGAA | 21 (0.000397%) | | | CCAGAGGCTGTTCACCTTGGAGACCTGATGCGGTTATGAGTACGACCGGGCGTGAGCGGCACTCGGTCCTCCGGATTTTCAAGGGCCGCCGGGGGCGCAC | 15 (0.000284%) | | | CCAGCAGCCGCGGTAATTCCAGCTCCAATAGCGTATATTTAAGTTGTTGCAGTTAAAAAGCTCGTAGTTGAACCTTGGGATGGGTCGGCCGGTCCGCCTT | 14 (0.000265%) | | | CCAGCCGTTCGTTTGCATGTTCCTTGACACTTTTCGTGCCGGGGTTTTGTGATATCCGGAAGCAACGCGCACGACAAGACCGAGATAAAAGCTCCCGATC | 4 (0.000076%) | | | CCAGTAGTCATATGCTTGTCTCAAAGATTAAGCCATGCATGTGTAAGTATGAACGAATTCAGACTGTGAAACTGCGAATGGCTCATTAAATCAGTTATAG | 70 (0.001324%) | | | CCATCCTTTGCTGATGCGGGACGGAAGCTGGTCTCCCGTGTGTTACCGCACGCGGTTGGCCTAAATCCGAGCCAAGGACGCCTGGAGCGTACCGACATGC | 8 (0.000151%) | | | CCATCGACCAGAGGCTGTTCACCTTGGAGACCTGATGCGGTTATGAGTACGACCGGGCGTGAGCGGCACTCGGTCCTCCGGATTTTCAAGGGCCGCCGGG | 9 (0.000170%) | | | CCATGCTAATGTATCCAGAGCGTAGGCTTGCTTTGAGCACTCTAATTTCTTCAAAGTAACAGCGCCGGAGGCACGACCCGGCCAATTAAGACCAGGAGCG | 8 (0.000151%) | | | CCATGCTTTCCAACGAAGCACGCCCATCCAACCTAGGCGAGACAAGGGTTCACATTTCGTTCATCACCCTTGGCCGGCTATCGAACAGCCGGACTCCCAT | 34 (0.000643%) | | | CCATGGGCATCGACACCTTGCGGCTAGGAACTGGAACGAGACGGGTGGCAAAGATTTCGAGTAGCACTTCATACTACCGTGGGTTTTTTAAACCTTCCGA | 6 (0.000114%) | | | CCATTGGAGGGCAAGTCTGGTGCCAGCAGCCGCGGTAATTCCAGCTCCAATAGCGTATATTTAAGTTGTTGCAGTTAAAAAGCTCGTAGTTGAACCTTGG | 19 (0.000359%) | | | CCCATCCAACCTAGGCGAGACAAGGGTTCACATTTCGTTCATCACCCTTGGCCGGCTATCGAACAGCCGGACTCCCATCAAAAGATGGTTGCCAAGAACA | 16 (0.000303%) | | | CCCATCCAACCTAGGCGAGACAAGGGTTCACATTTCGTTCATCACCCTTGGCCGGCTTTCGAACAGCCGGACTCCCATCAAAAGATGGTTGCCAAGAACA | 9 (0.000170%) | | | CCCATGCTAATGTATCCAGAGCGTAGGCTTGCTTTGAGCACTCTAATTTCTTCAAAGTAACAGCGCCGGAGGCACGACCCGGCCAATTAAGACCAGGAGC | 6 (0.000114%) | | | CCCATGGGCATCGACACCTTGCGGCTAGGAACTGGAACGAGACGGGTGGCAAAGATTTCGAGTAGCACTTCATACTACCGTGGGTTTTTTAAACCTTCCG | 14 (0.000265%) | | | CCCCGCCACATCCTCTCAAACGCAATGGAAAGAGAGAAAGGACGAGGTCTTGACCGTCATCTTTTGCCCGAAGGACGGATGAGCTTTGGCGGGACTGAAT | 6 (0.000114%) | | | CCCGAGTGTGAGCGAGGTGTGAGTGTCGCCCATGGGCATCGACACCTTGCGGCTAGGAACTGGAACGAGACGGGTAGCAAAGATTTCGAGTAGCACTTCA | 15 (0.000284%) | | | CCCGAGTGTGAGCGAGGTGTGAGTGTCGCCCATGGGCATCGACACCTTGCGGCTAGGAACTGGAACGAGACGGGTGGCAAAGATTTCGAGTAGCACTTCA | 18 (0.000341%) | | | CCCGCCACATCCTCTCAAACGCAATGGAAAGAGAGAAAGGACGAGGTCTTGACCGTCATCTTTTGCCCGAAGGACGGATGAGCTTTGGCGGGACTGAATC | 9 (0.000170%) | | | CCCGCGTCGACCTTTTATCTAATAAATGCGTCCCTTCCATAAGTCGGGGTTTGTTGCACGTATTAGCTCTAGAATTACTACGGTTATCCGAGTAGTAGTT | 44 (0.000832%) | | | CCCGGCCGTCGGGGCAAGAGCCAGGCCTCGATGAGTAGGAGGGCGCGGCGGTCGCTGCAAAACCTAGGGCGCGAGCCCGGGCGGAGCGGCCGTCGGTGCA | 21 (0.000397%) | | | CCCTAACGCCTCGAAGAACTAATGGCAGCCCACGCAAGGCAAGCCCATTCTCCTCGACGATTCAGCAGTTTTTGTCCGAGAACTGCTGAGAAAACTCGGA | 11 (0.000208%) | | | CCCTCACCATCCTTTGCTGATGCGGGACGGAAGCTGGTCTCCCGTGTGTTACCGCACGCGGTTGGCCTAAATCCGAGCCAAGGACGCCTGGAGCGTACCG | 16 (0.000303%) | | | CCCTGTTGAGCTTGACTCTAGTCCGACTTTGTGAAATGACTTGAGAGGTGTAGGATAAGTGGGAGCTTCGGCGCAAGTGAAATACCACTACTTTTAACGT | 13 (0.000246%) | | | CCCTTAGATGTTCTGGGCCGCACGCGCGCTACACTGATGTATTCAACGAGTTCACACCTTGGCCGACAGGCCCGGGTAATCTTTGAAATTTCATCGTGAT | 4 (0.000076%) | | | CCCTTGCCTACATTGTTCCATCGACCAGAGGCTGTTCACCTTGGAGACCTGATGCGGTTATGAGTACGACCGGGCGTGAGCGGCACTCGGTCCTCCGGAT | 30 (0.000568%) | | | CCGACTTCCCTTGCCTACATTGTTCCATCGACCAGAGGCTGTTCACCTTGGAGACCTGATGCGGTTATGAGTACGACCGGGCGTGAGCGGCACTCGGTCC | 52 (0.000984%) | | | CCGACTTTGTGAAATGACTTGAGAGGTGTAGGATAAGTGGGAGCTTCGGCGCAAGTGAAATACCACTACTTTTAACGTTATTTTACTTACTCCGTGAATC | 21 (0.000397%) | | | CCGAGTGTGAGCGAGGTGTGAGTGTCGCCCATGGGCATCGACACCTTGCGGCTAGGAACTGGAACGAGACGGGTAGCAAAGATTTCGAGTAGCACTTCAT | 10 (0.000189%) | | | CCGAGTGTGAGCGAGGTGTGAGTGTCGCCCATGGGCATCGACACCTTGCGGCTAGGAACTGGAACGAGACGGGTGGCAAAGATTTCGAGTAGCACTTCAT | 18 (0.000341%) | | | CCGCCACATCCTCTCAAACGCAATGGAAAGAGAGAAAGGACGAGGTCTTGACCGTCATCTTTTGCCCGAAGGACGGATGAGCTTTGGCGGGACTGAATCA | 7 (0.000132%) | | | CCGCCTAGGCTGTCCCGAGTGTGAGCGAGGTGTGAGTGTCGCCCATGGGCATCGACACCTTGCGGCTAGGAACTGGAACGAGACGGGTAGCAAAGATTTC | 27 (0.000511%) | | | CCGCCTAGGCTGTCCCGAGTGTGAGCGAGGTGTGAGTGTCGCCCATGGGCATCGACACCTTGCGGCTAGGAACTGGAACGAGACGGGTGGCAAAGATTTC | 62 (0.001173%) | | | CCGCGTCGACCTTTTATCTAATAAATGCGTCCCTTCCATAAGTCGGGGTTTGTTGCACGTATTAGCTCTAGAATTACTACGGTTATCCGAGTAGTAGTTA | 13 (0.000246%) | | | CCGCTAAGGAGTGTGTAACAACTCACCTGCCGAATCAACTAGCCCCGAAAATGGATGGCGCTTAAGCGCGCGACCTATACCCGGCCGTCGGGGCAAGAGC | 19 (0.000359%) | | | CCGGATTGCTCCGTTCCGCATCCGACCAGGACGCATCGCCGGCCCCCATCCGCTTCCCTCCCGACAATTTCAAGCACTCTTTGACTCTCTTTTCAAAGTC | 49 (0.000927%) | | | CCGGCCGTCGGGGCAAGAGCCAGGCCTCGATGAGTAGGAGGGCGCGGCGGTCGCTGCAAAACCTAGGGCGCGAGCCCGGGCGGAGCGGCCGTCGGTGCAG | 10 (0.000189%) | | | CCGGGCGTCGGCCTGTGGGCTCCCCATTCGACCCGTCTTGAAACACGGACCAAGGAGTCTGACATGTGTGCGAGTCAACGGGTGAGTAAACCCGTAAGGC | 13 (0.000246%) | | | CCGGGGAGGCGAATGCCAGCCGTTCGTTTGCATGTTCCTTGACACTTTTCGTGCCGGGGTTTTGTGATATCCGGAAGCAACGCGCACGACAAGACCGAGA | 7 (0.000132%) | | | CCGTAACTTCGGGAAAAGGATTGGCTCTGAGGGCTGGGCTCGGGGGTCCCAGTTCCGAACCCGTCGGCTGTCAGCGGACTGCTCGAGCTGCTTCCGCGGC | 3 (0.000057%) | | | CCGTCGAGTTATCATGAATCATCAGAGCAACGGGCAGAGCCCGCGTCGACCTTTTATCTAATAAATGCGTCCCTTCCATAAGTCGGGGTTTGTTGCACGT | 21 (0.000397%) | | | CCGTCTCCGGGGAGGCGAATGCCAGCCGTTCGTTTGCATGTTCCTTGACACTTTTCGTGCCGGGGTTTTGTGATATCCGGAAGCAACGCGCACGACAAGA | 7 (0.000132%) | | | CCTAACGCCTCGAAGAACTAATGGCAGCCCACGCAAGGCAAGCCCATTCTCCTCGACGATTCAGCAGTTTTTGTCCGAGAACTGCTGAGAAAACTCGGAA | 18 (0.000341%) | | | CCTAACGGCGTGCCTCGGCATCAGCGTGCTCCGGGCGTCGGCCTGTGGGCTCCCCATTCGACCCGTCTTGAAACACGGACCAAGGAGTCTGACATGTGTG | 4 (0.000076%) | | | CCTAAGTAGTGTTTCCTTGTTAGAAGACACAAAGCCAAAGACTCATATGGACTTTGGCTACACCATGAAAGCTTTGAGAAGCAAGAAGAAGGTTGGTTAG | 66 (0.001249%) | | | CCTACATTGTTCCATCGACCAGAGGCTGTTCACCTTGGAGACCTGATGCGGTTATGAGTACGACCGGGCGTGAGCGGCACTCGGTCCTCCGGATTTTCAA | 13 (0.000246%) | | | CCTAGGCGAGACAAGGGTTCACATTTCGTTCATCACCCTTGGCCGGCTATCGAACAGCCGGACTCCCATCAAAAGATGGTTGCCAAGAACATCTTCGTTA | 6 (0.000114%) | | | CCTAGGCGAGACAAGGGTTCACATTTCGTTCATCACCCTTGGCCGGCTTTCGAACAGCCGGACTCCCATCAAAAGATGGTTGCCAAGAACATCTTCGTTA | 12 (0.000227%) | | | CCTAGGCTGTCCCGAGTGTGAGCGAGGTGTGAGTGTCGCCCATGGGCATCGACACCTTGCGGCTAGGAACTGGAACGAGACGGGTAGCAAAGATTTCGAG | 39 (0.000738%) | | | CCTAGGCTGTCCCGAGTGTGAGCGAGGTGTGAGTGTCGCCCATGGGCATCGACACCTTGCGGCTAGGAACTGGAACGAGACGGGTGGCAAAGATTTCGAG | 73 (0.001381%) | | | CCTATACCCGGCCGTCGGGGCAAGAGCCAGGCCTCGATGAGTAGGAGGGCGCGGCGGTCGCTGCAAAACCTAGGGCGCGAGCCCGGGCGGAGCGGCCGTC | 31 (0.000586%) | | | CCTATGATGTTATCCCATGCTAATGTATCCAGAGCGTAGGCTTGCTTTGAGCACTCTAATTTCTTCAAAGTAACAGCGCCGGAGGCACGACCCGGCCAAT | 27 (0.000511%) | | | CCTATTTTCTGGTAAATTTTCATAATTTTTTGACACCTCTAGCTAGGTCATTTGACCTGATACAACATCGGATTTTCATGGTCTAGTTGGGGCTCCGTGG | 6 (0.000114%) | | | CCTCACCATCCTTTGCTGATGCGGGACGGAAGCTGGTCTCCCGTGTGTTACCGCACGCGGTTGGCCTAAATCCGAGCCAAGGACGCCTGGAGCGTACCGA | 57 (0.001078%) | | | CCTCCGCTTATTGATATGCTTAAACTCAGCGGGTAATCCCGCCTGACCTGGGGTCGCTATATGGACTTTGGGTCATCTACAGCTTCCGGACAAGAGCGAC | 4 (0.000076%) | | | CCTCGAAGAACTAATGGCAGCCCACGCAAGGCAAGCCCATTCTCCTCGACGATTCAGCAGTTTTTGTCCGAGAACTGCTGAGAAAACTCGGAAAAAGGCA | 12 (0.000227%) | | | CCTCGGCATCAGCGTGCTCCGGGCGTCGGCCTGTGGGCTCCCCATTCGACCCGTCTTGAAACACGGACCAAGGAGTCTGACATGTGTGCGAGTCAACGGG | 8 (0.000151%) | | | CCTCGTTAAGGGATTTAGATTGTACTCATTCCAATTACCAGACTCGAAAGAGCCCGGTATTGTTATTTATTGTCACTACCTCCCCGTGTCAGGATTGGGT | 218 (0.004124%) | | | CCTGTTGAGCTTGACTCTAGTCCGACTTTGTGAAATGACTTGAGAGGTGTAGGATAAGTGGGAGCTTCGGCGCAAGTGAAATACCACTACTTTTAACGTT | 6 (0.000114%) | | | CCTTAGATGTTCTGGGCCGCACGCGCGCTACACTGATGTATTCAACGAGTTCACACCTTGGCCGACAGGCCCGGGTAATCTTTGAAATTTCATCGTGATG | 5 (0.000095%) | | | CCTTGCCTACATTGTTCCATCGACCAGAGGCTGTTCACCTTGGAGACCTGATGCGGTTATGAGTACGACCGGGCGTGAGCGGCACTCGGTCCTCCGGATT | 8 (0.000151%) | | | CCTTGTTAGAAGACACAAAGCCAAAGACTCATATGGACTTTGGCTACACCATGAAAGCTTTGAGAAGCAAGAAGAAGGTTGGTTAGTGTTTTGGAGTCGA | 43 (0.000813%) | | | CCTTGTTAGTTTCTTTTCCTCCGCTTATTGATATGCTTAAACTCAGCGGGTAATCCCGCCTGACCTGGGGTCGCTATATGGACTTTGGGTCATCTACAGC | 8 (0.000151%) | | | CCTTTGCTGATGCGGGACGGAAGCTGGTCTCCCGTGTGTTACCGCACGCGGTTGGCCTAAATCCGAGCCAAGGACGCCTGGAGCGTACCGACATGCGGTG | 5 (0.000095%) | | | CGAAAGAGCCCGGTATTGTTATTTATTGTCACTACCTCCCCGTGTCAGGATTGGGTAATTTGCGCGCCTGCTGCCTTCCTTGGATGTGGTAGCCGTTTCT | 20 (0.000378%) | | | CGAAATCCGCTAAGGAGTGTGTAACAACTCACCTGCCGAATCAACTAGCCCCGAAAATGGATGGCGCTTAAGCGCGCGACCTATACCCGGCCGTCGGGGC | 17 (0.000322%) | | | CGAAATCCTATGATGTTATCCCATGCTAATGTATCCAGAGCGTAGGCTTGCTTTGAGCACTCTAATTTCTTCAAAGTAACAGCGCCGGAGGCACGACCCG | 426 (0.008059%) | | | CGAAGCACGCCCATCCAACCTAGGCGAGACAAGGGTTCACATTTCGTTCATCACCCTTGGCCGGCTATCGAACAGCCGGACTCCCATCAAAAGATGGTTG | 25 (0.000473%) | | | CGAAGCACGCCCATCCAACCTAGGCGAGACAAGGGTTCACATTTCGTTCATCACCCTTGGCCGGCTTTCGAACAGCCGGACTCCCATCAAAAGATGGTTG | 18 (0.000341%) | | | CGAATATGACTTGATGTCATGTGTATGATTGAGTATAAGA | 542 (0.004101%) | | | CGAATGCCAGCCGTTCGTTTGCATGTTCCTTGACACTTTTCGTGCCGGGGTTTTGTGATATCCGGAAGCAACGCGCACGACAAGACCGAGATAAAAGCTC | 22 (0.000416%) | | | CGAATGGCTCATTAAATCAGTTATAGTTTGTTTGATGGTAACTACTACTCGGATAACCGTAGTAATTCTAGAGCTAATACGTGCAACAAACCCCGACTTA | 11 (0.000208%) | | | CGACCAGAGGCTGTTCACCTTGGAGACCTGATGCGGTTATGAGTACGACCGGGCGTGAGCGGCACTCGGTCCTCCGGATTTTCAAGGGCCGCCGGGGGCG | 19 (0.000359%) | | | CGACCTATACCCGGCCGTCGGGGCAAGAGCCAGGCCTCGATGAGTAGGAGGGCGCGGCGGTCGCTGCAAAACCTAGGGCGCGAGCCCGGGCGGAGCGGCC | 5 (0.000095%) | | | CGACCTTTTATCTAATAAATGCGTCCCTTCCATAAGTCGGGGTTTGTTGCACGTATTAGCTCTAGAATTACTACGGTTATCCGAGTAGTAGTTACCATCA | 23 (0.000435%) | | | CGACGGGCGGTGTGTACAAAGGGCAGGGACGTAGTCAACGCGAGCTGATGACTCGCGCTTACTAGGAATTCCTCGTTGAAGACCAACAATTGCAATGATC | 37 (0.000700%) | | | CGACGTGGGTGGTTCGCCGCCCGCGACGTCGCGAGAAGTCCACTAAACCTTATCATTTAGAGGAAGGAGAAGTCGTAACAAGGTTTCCGTAGGTGAACCT | 98 (0.001854%) | | | CGACTCCAAAACACTAACCAACCTTCTTCTTGCTTCTCAAAGCTTTCATGGTGTAGCCAAAGTCCATATGAGTCTTTGGCTTTGTGTCTTCTAACAAGGA | 55 (0.001041%) | | | CGACTTCCCTTGCCTACATTGTTCCATCGACCAGAGGCTGTTCACCTTGGAGACCTGATGCGGTTATGAGTACGACCGGGCGTGAGCGGCACTCGGTCCT | 34 (0.000643%) | | | CGACTTTGTGAAATGACTTGAGAGGTGTAGGATAAGTGGGAGCTTCGGCGCAAGTGAAATACCACTACTTTTAACGTTATTTTACTTACTCCGTGAATCG | 11 (0.000208%) | | | CGAGACAAGGGTTCACATTTCGTTCATCACCCTTGGCCGGCTATCGAACAGCCGGACTCCCATCAAAAGATGGTTGCCAAGAACATCTTCGTTACGGTTT | 12 (0.000227%) | | | CGAGACAAGGGTTCACATTTCGTTCATCACCCTTGGCCGGCTTTCGAACAGCCGGACTCCCATCAAAAGATGGTTGCCAAGAACATCTTCGTTACGGTTT | 4 (0.000076%) | | | CGAGGTGTGAGTGTCGCCCATGGGCATCGACACCTTGCGGCTAGGAACTGGAACGAGACGGGTAGCAAAGATTTCGAGTAGCACTTCATACTACCGTGGG | 15 (0.000284%) | | | CGAGGTGTGAGTGTCGCCCATGGGCATCGACACCTTGCGGCTAGGAACTGGAACGAGACGGGTGGCAAAGATTTCGAGTAGCACTTCATACTACCGTGGG | 21 (0.000397%) | | | CGAGTGTGAGCGAGGTGTGAGTGTCGCCCATGGGCATCGACACCTTGCGGCTAGGAACTGGAACGAGACGGGTAGCAAAGATTTCGAGTAGCACTTCATA | 33 (0.000624%) | | | CGAGTGTGAGCGAGGTGTGAGTGTCGCCCATGGGCATCGACACCTTGCGGCTAGGAACTGGAACGAGACGGGTGGCAAAGATTTCGAGTAGCACTTCATA | 35 (0.000662%) | | | CGAGTTATCATGAATCATCAGAGCAACGGGCAGAGCCCGCGTCGACCTTTTATCTAATAAATGCGTCCCTTCCATAAGTCGGGGTTTGTTGCACGTATTA | 138 (0.002611%) | | | CGATCACGGCAATTCCCCGCCACATCCTCTCAAACGCAATGGAAAGAGAGAAAGGACGAGGTCTTGACCGTCATCTTTTGCCCGAAGGACGGATGAGCTT | 3 (0.000057%) | | | CGATCCGCCTAGGCTGTCCCGAGTGTGAGCGAGGTGTGAGTGTCGCCCATGGGCATCGACACCTTGCGGCTAGGAACTGGAACGAGACGGGTGGCAAAGA | 6 (0.000114%) | | | CGATCCGGTTAAAATTCCGGAACCGGGACGTGGCGGTTGACGGCAACGTTAGGGAGTCCGGAGACGTCGGCGGGGGCCTCGGGAAGAGTTATCTTTTCTG | 3 (0.000057%) | | | CGATCCGTCGAGTTATCATGAATCATCAGAGCAACGGGCAGAGCCCGCGTCGACCTTTTATCTAATAAATGCGTCCCTTCCATAAGTCGGGGTTTGTTGC | 151 (0.002857%) | | | CGATCCTCGTTAAGGGATTTAGATTGTACTCATTCCAATTACCAGACTCGAAAGAGCCCGGTATTGTTATTTATTGTCACTACCTCCCCGTGTCAGGATT | 6 (0.000114%) | | | CGATCGAAATCCTATGATGTTATCCCATGCTAATGTATCCAGAGCGTAGGCTTGCTTTGAGCACTCTAATTTCTTCAAAGTAACAGCGCCGGAGGCACGA | 8 (0.000151%) | | | CGATCGGGTTGCGGTTTAAGTTGTTATACTCAATCATACA | 76 (0.000575%) | | | CGCAACAAGATCTTAAAGGCGTAAGAATTGTATCCTTGTTAAAAGACACAAAGCCAAAGACTCATATGGACTTTGGCTACACCATGAAAGCTTTGAGAAG | 32 (0.000605%) | | | CGCAATGGAAAGAGAGAAAGGACGAGGTCTTGACCGTCATCTTTTGCCCGAAGGACGGATGAGCTTTGGCGGGACTGAATCACTTCGAGTCACCGTCGAC | 3 (0.000057%) | | | CGCATGTCGGTACGCTCCAGGCGTCCTTGGCTCGGATTTAGGCCAACCGCGTGCGGTAACACACGGGAGACCAGCTTCCGTCCCGCATCAGCAAAGGATG | 8 (0.000151%) | | | CGCCACATCCTCTCAAACGCAATGGAAAGAGAGAAAGGACGAGGTCTTGACCGTCATCTTTTGCCCGAAGGACGGATGAGCTTTGGCGGGACTGAATCAC | 7 (0.000132%) | | | CGCCCATCCAACCTAGGCGAGACAAGGGTTCACATTTCGTTCATCACCCTTGGCCGGCTATCGAACAGCCGGACTCCCATCAAAAGATGGTTGCCAAGAA | 73 (0.001381%) | | | CGCCCATCCAACCTAGGCGAGACAAGGGTTCACATTTCGTTCATCACCCTTGGCCGGCTTTCGAACAGCCGGACTCCCATCAAAAGATGGTTGCCAAGAA | 94 (0.001778%) | | | CGCCCATGGGCATCGACACCTTGCGGCTAGGAACTGGAACGAGACGGGTGGCAAAGATTTCGAGTAGCACTTCATACTACCGTGGGTTTTTTAAACCTTC | 32 (0.000605%) | | | CGCCGCCCGCGACGTCGCGAGAAGTCCACTAAACCTTATCATTTAGAGGAAGGAGAAGTCGTAACAAGGTTTCCGTAGGTGAACCTGCGGAAGGATCGAT | 7 (0.000132%) | | | CGCCGTTACTAAGGGAATCCTTGTTAGTTTCTTTTCCTCCGCTTATTGATATGCTTAAACTCAGCGGGTAATCCCGCCTGACCTGGGGTCGCTATATGGA | 5 (0.000095%) | | | CGCCTAACGGCGTGCCTCGGCATCAGCGTGCTCCGGGCGTCGGCCTGTGGGCTCCCCATTCGACCCGTCTTGAAACACGGACCAAGGAGTCTGACATGTG | 42 (0.000795%) | | | CGCCTAGGCTGTCCCGAGTGTGAGCGAGGTGTGAGTGTCGCCCATGGGCATCGACACCTTGCGGCTAGGAACTGGAACGAGACGGGTAGCAAAGATTTCG | 115 (0.002176%) | | | CGCCTAGGCTGTCCCGAGTGTGAGCGAGGTGTGAGTGTCGCCCATGGGCATCGACACCTTGCGGCTAGGAACTGGAACGAGACGGGTGGCAAAGATTTCG | 207 (0.003916%) | | | CGCCTCGAAGAACTAATGGCAGCCCACGCAAGGCAAGCCCATTCTCCTCGACGATTCAGCAGTTTTTGTCCGAGAACTGCTGAGAAAACTCGGAAAAAGG | 3 (0.000057%) | | | CGCGACCTATACCCGGCCGTCGGGGCAAGAGCCAGGCCTCGATGAGTAGGAGGGCGCGGCGGTCGCTGCAAAACCTAGGGCGCGAGCCCGGGCGGAGCGG | 14 (0.000265%) | | | CGCGCCTAACGGCGTGCCTCGGCATCAGCGTGCTCCGGGCGTCGGCCTGTGGGCTCCCCATTCGACCCGTCTTGAAACACGGACCAAGGAGTCTGACATG | 43 (0.000813%) | | | CGCGGTAATTCCAGCTCCAATAGCGTATATTTAAGTTGTTGCAGTTAAAAAGCTCGTAGTTGAACCTTGGGATGGGTCGGCCGGTCCGCCTTTGGTGTGC | 12 (0.000227%) | | | CGCGTCGACCTTTTATCTAATAAATGCGTCCCTTCCATAAGTCGGGGTTTGTTGCACGTATTAGCTCTAGAATTACTACGGTTATCCGAGTAGTAGTTAC | 5 (0.000095%) | | | CGCTAAGGAGTGTGTAACAACTCACCTGCCGAATCAACTAGCCCCGAAAATGGATGGCGCTTAAGCGCGCGACCTATACCCGGCCGTCGGGGCAAGAGCC | 8 (0.000151%) | | | CGCTTATTGATATGCTTAAACTCAGCGGGTAATCCCGCCTGACCTGGGGTCGCTATATGGACTTTGGGTCATCTACAGCTTCCGGACAAGAGCGACCGAT | 8 (0.000151%) | | | CGCTTTCACGGTTCGTATTCGTACTGAAAATCAGAATCAAACGAGCTTTTACCCTTTTGTTCCACACGAGATTTCTGTTCTCGTTGAGCTCATCTTAGGA | 9 (0.000170%) | | | CGGAAGAGCACACGTCTGAACTCCAGTCACGAGATTCCAT | 17 (0.000129%) | | | CGGACCGGATTGCTCCGTTCCGCATCCGACCAGGACGCATCGCCGGCCCCCATCCGCTTCCCTCCCGACAATTTCAAGCACTCTTTGACTCTCTTTTCAA | 3 (0.000057%) | | | CGGAGAATTAGGGTTCGATTCCGGAGAGGGAGCCTGAGAAACGGCTACCACATCCAAGGAAGGCAGCAGGCGCGCAAATTACCCAATCCTGACACGGGGA | 18 (0.000341%) | | | CGGATCTTAAAGGCGTAAGAATTGTATCCTTGTTAGAAGACACAAAGCCAAAGACTCATATGGACTTTGGCTACACCATGAAAGCTTTGAGAAGCAAGAA | 11 (0.000208%) | | | CGGATTGCTCCGTTCCGCATCCGACCAGGACGCATCGCCGGCCCCCATCCGCTTCCCTCCCGACAATTTCAAGCACTCTTTGACTCTCTTTTCAAAGTCC | 25 (0.000473%) | | | CGGCAATTCCCCGCCACATCCTCTCAAACGCAATGGAAAGAGAGAAAGGACGAGGTCTTGACCGTCATCTTTTGCCCGAAGGACGGATGAGCTTTGGCGG | 172 (0.003254%) | | | CGGCACATCTGTTAAAAGATAACGCAGGTGTCCTAAGATGAGCTCAACGAGAACAGAAATCTCGTGTGGAACAAAAGGGTAAAAGCTCGTTTGATTCTGA | 5 (0.000095%) | | | CGGCATCAGCGTGCTCCGGGCGTCGGCCTGTGGGCTCCCCATTCGACCCGTCTTGAAACACGGACCAAGGAGTCTGACATGTGTGCGAGTCAACGGGTGA | 15 (0.000284%) | | | CGGCCGTCGGGGCAAGAGCCAGGCCTCGATGAGTAGGAGGGCGCGGCGGTCGCTGCAAAACCTAGGGCGCGAGCCCGGGCGGAGCGGCCGTCGGTGCAGA | 46 (0.000870%) | | | CGGCGGACCGGATTGCTCCGTTCCGCATCCGACCAGGACGCATCGCCGGCCCCCATCCGCTTCCCTCCCGACAATTTCAAGCACTCTTTGACTCTCTTTT | 10 (0.000189%) | | | CGGCGTGCCTCGGCATCAGCGTGCTCCGGGCGTCGGCCTGTGGGCTCCCCATTCGACCCGTCTTGAAACACGGACCAAGGAGTCTGACATGTGTGCGAGT | 9 (0.000170%) | | | CGGGCAGAGCCCGCGTCGACCTTTTATCTAATAAATGCGTCCCTTCCATAAGTCGGGGTTTGTTGCACGTATTAGCTCTAGAATTACTACGGTTATCCGA | 66 (0.001249%) | | | CGGGCGGTGTGTACAAAGGGCAGGGACGTAGTCAACGCGAGCTGATGACTCGCGCTTACTAGGAATTCCTCGTTGAAGACCAACAATTGCAATGATCGAT | 32 (0.000605%) | | | CGGGCGTCGGCCTGTGGGCTCCCCATTCGACCCGTCTTGAAACACGGACCAAGGAGTCTGACATGTGTGCGAGTCAACGGGTGAGTAAACCCGTAAGGCG | 54 (0.001022%) | | | CGGGGAGGCGAATGCCAGCCGTTCGTTTGCATGTTCCTTGACACTTTTCGTGCCGGGGTTTTGTGATATCCGGAAGCAACGCGCACGACAAGACCGAGAT | 12 (0.000227%) | | | CGGGGGCATTCGTATTTCATAGTCAGAGGTGAAATTCTTGGATTTATGAAAGACGAACAACTGCGAAAGCATTTGCCAAGGATGTTTTCATTAATCAAGA | 32 (0.000605%) | | | CGGGTGACGGAGAATTAGGGTTCGATTCCGGAGAGGGAGCCTGAGAAACGGCTACCACATCCAAGGAAGGCAGCAGGCGCGCAAATTACCCAATCCTGAC | 7 (0.000132%) | | | CGGGTTGCGGTTTAAGTTGTTATACTCAATCATACACATG | 488 (0.003693%) | | | CGGGTTTACTCACCCGTTGACTCGCACACATGTCAGACTCCTTGGTCCGTGTTTCAAGACGGGTCGAATGGGGAGCCCACAGGCCGACGCCCGGAGCACG | 13 (0.000246%) | | | CGGTAATTCCAGCTCCAATAGCGTATATTTAAGTTGTTGCAGTTAAAAAGCTCGTAGTTGAACCTTGGGATGGGTCGGCCGGTCCGCCTTTGGTGTGCAT | 14 (0.000265%) | | | CGGTACGCTCCAGGCGTCCTTGGCTCGGATTTAGGCCAACCGCGTGCGGTAACACACGGGAGACCAGCTTCCGTCCCGCATCAGCAAAGGATGGTGAGGG | 8 (0.000151%) | | | CGGTAGGAGCGACGGGCGGTGTGTACAAAGGGCAGGGACGTAGTCAACGCGAGCTGATGACTCGCGCTTACTAGGAATTCCTCGTTGAAGACCAACAATT | 51 (0.000965%) | | | CGGTTTAAGTTCTTATACTCAATCATACACATGACATCAA | 519 (0.003927%) | | | CGGTTTAAGTTGTTATACTCAATCATACACATGACAACAAGTCATATTCGACTCCAAAACACTAACCAACCTTCTTCTTGCTTCTCAAAGCTTTCATGGT | 1007 (0.019051%) | | | CGTAACTTCGGGAAAAGGATTGGCTCTGAGGGCTGGGCTCGGGGGTCCCAGTTCCGAACCCGTCGGCTGTCAGCGGACTGCTCGAGCTGCTTCCGCGGCG | 6 (0.000114%) | | | CGTAAGAATTGTATCCTTGTTAGAAGACACAAAGCCAAAGACTCATATGGACTTTGGCTACACCATGAAAGCTTTGAGAAGCAAGAAGAAGGTTGGTTAG | 62 (0.001173%) | | | CGTACTGAAAATCAGAATCAAACGAGCTTTTACCCTTTTGTTCCACACGAGATTTCTGTTCTCGTTGAGCTCATCTTAGGACACCTGCGTTATCTTTTAA | 6 (0.000114%) | | | CGTAGGCTTGCTTTGAGCACTCTAATTTCTTCAAAGTAACAGCGCCGGAGGCACGACCCGGCCAATTAAGACCAGGAGCGTATCGCCGACCGAAGGGACA | 19 (0.000359%) | | | CGTATTCGTACTGAAAATCAGAATCAAACGAGCTTTTACCCTTTTGTTCCACACGAGATTTCTGTTCTCGTTGAGCTCATCTTAGGACACCTGCGTTATC | 24 (0.000454%) | | | CGTATTTCATAGTCAGAGGTGAAATTCTTGGATTTATGAAAGACGAACAACTGCGAAAGCATTTGCCAAGGATGTTTTCATTAATCAAGAACGAAAGTTG | 15 (0.000284%) | | | CGTCCCTCACCATCCTTTGCTGATGCGGGACGGAAGCTGGTCTCCCGTGTGTTACCGCACGCGGTTGGCCTAAATCCGAGCCAAGGACGCCTGGAGCGTA | 12 (0.000227%) | | | CGTCGACCTTTTATCTAATAAATGCGTCCCTTCCATAAGTCGGGGTTTGTTGCACGTATTAGCTCTAGAATTACTACGGTTATCCGAGTAGTAGTTACCA | 5 (0.000095%) | | | CGTCGAGTTATCATGAATCATCAGAGCAACGGGCAGAGCCCGCGTCGACCTTTTATCTAATAAATGCGTCCCTTCCATAAGTCGGGGTTTGTTGCACGTA | 110 (0.002081%) | | | CGTCGGCCTGTGGGCTCCCCATTCGACCCGTCTTGAAACACGGACCAAGGAGTCTGACATGTGTGCGAGTCAACGGGTGAGTAAACCCGTAAGGCGCAAG | 6 (0.000114%) | | | CGTCTCCGGGGAGGCGAATGCCAGCCGTTCGTTTGCATGTTCCTTGACACTTTTCGTGCCGGGGTTTTGTGATATCCGGAAGCAACGCGCACGACAAGAC | 3 (0.000057%) | | | CGTGCCTCGGCATCAGCGTGCTCCGGGCGTCGGCCTGTGGGCTCCCCATTCGACCCGTCTTGAAACACGGACCAAGGAGTCTGACATGTGTGCGAGTCAA | 32 (0.000605%) | | | CGTGCTCCGGGCGTCGGCCTGTGGGCTCCCCATTCGACCCGTCTTGAAACACGGACCAAGGAGTCTGACATGTGTGCGAGTCAACGGGTGAGTAAACCCG | 7 (0.000132%) | | | CGTGGGTGGTTCGCCGCCCGCGACGTCGCGAGAAGTCCACTAAACCTTATCATTTAGAGGAAGGAGAAGTCGTAACAAGGTTTCCGTAGGTGAACCTGCG | 32 (0.000605%) | | | CGTTAAGGGATTTAGATTGTACTCATTCCAATTACCAGACTCGAAAGAGCCCGGTATTGTTATTTATTGTCACTACCTCCCCGTGTCAGGATTGGGTAAT | 64 (0.001211%) | | | CTAAATCCTATTTTCTGGTAAATTTTCATAATTTTTTGACACCTCTAGCTAGGTCATTTGACCTGATACAACATCGGATTTTCATGGTCTAGTTGGGGCT | 22 (0.000416%) | | | CTAACGCCTCGAAGAACTAATGGCAGCCCACGCAAGGCAAGCCCATTCTCCTCGACGATTCAGCAGTTTTTGTCCGAGAACTGCTGAGAAAACTCGGAAA | 32 (0.000605%) | | | CTAACGGCGTGCCTCGGCATCAGCGTGCTCCGGGCGTCGGCCTGTGGGCTCCCCATTCGACCCGTCTTGAAACACGGACCAAGGAGTCTGACATGTGTGC | 100 (0.001892%) | | | CTAAGTAGTGTTTCCTTGTTAGAAGACACAAAGCCAAAGACTCATATGGACTTTGGCTACACCATGAAAGCTTTGAGAAGCAAGAAGAAGGTTGGTTAGT | 83 (0.001570%) | | | CTAATGTATCCAGAGCGTAGGCTTGCTTTGAGCACTCTAATTTCTTCAAAGTAACAGCGCCGGAGGCACGACCCGGCCAATTAAGACCAGGAGCGTATCG | 49 (0.000927%) | | | CTACATTGTTCCATCGACCAGAGGCTGTTCACCTTGGAGACCTGATGCGGTTATGAGTACGACCGGGCGTGAGCGGCACTCGGTCCTCCGGATTTTCAAG | 50 (0.000946%) | | | CTAGGCGAGACAAGGGTTCACATTTCGTTCATCACCCTTGGCCGGCTATCGAACAGCCGGACTCCCATCAAAAGATGGTTGCCAAGAACATCTTCGTTAC | 27 (0.000511%) | | | CTAGGCGAGACAAGGGTTCACATTTCGTTCATCACCCTTGGCCGGCTTTCGAACAGCCGGACTCCCATCAAAAGATGGTTGCCAAGAACATCTTCGTTAC | 34 (0.000643%) | | | CTAGGCTGTCCCGAGTGTGAGCGAGGTGTGAGTGTCGCCCATGGGCATCGACACCTTGCGGCTAGGAACTGGAACGAGACGGGTAGCAAAGATTTCGAGT | 287 (0.005430%) | | | CTAGGCTGTCCCGAGTGTGAGCGAGGTGTGAGTGTCGCCCATGGGCATCGACACCTTGCGGCTAGGAACTGGAACGAGACGGGTGGCAAAGATTTCGAGT | 447 (0.008456%) | | | CTAGTCCGACTTTGTGAAATGACTTGAGAGGTGTAGGATAAGTGGGAGCTTCGGCGCAAGTGAAATACCACTACTTTTAACGTTATTTTACTTACTCCGT | 12 (0.000227%) | | | CTATACCCGGCCGTCGGGGCAAGAGCCAGGCCTCGATGAGTAGGAGGGCGCGGCGGTCGCTGCAAAACCTAGGGCGCGAGCCCGGGCGGAGCGGCCGTCG | 71 (0.001343%) | | | CTATGATGTTATCCCATGCTAATGTATCCAGAGCGTAGGCTTGCTTTGAGCACTCTAATTTCTTCAAAGTAACAGCGCCGGAGGCACGACCCGGCCAATT | 227 (0.004294%) | | | CTATTTTCTGGTAAATTTTCATAATTTTTTGACACCTCTAGCTAGGTCATTTGACCTGATACAACATCGGATTTTCATGGTCTAGTTGGGGCTCCGTGGG | 23 (0.000435%) | | | CTCAAACGCAATGGAAAGAGAGAAAGGACGAGGTCTTGACCGTCATCTTTTGCCCGAAGGACGGATGAGCTTTGGCGGGACTGAATCACTTCGAGTCACC | 17 (0.000322%) | | | CTCAAAGATTAAGCCATGCATGTGTAAGTATGAACGAATTCAGACTGTGAAACTGCGAATGGCTCATTAAATCAGTTATAGTTTGTTTGATGGTAACTAC | 18 (0.000341%) | | | CTCAAAGCTTTCATGGTGTAGCCAAAGTCCATATGAGTCTTTGGCTTTGTGTCTTCTAACAAGGAAACACTACTTAGGCTTATAAGATGCGGTTGCGGTT | 7 (0.000132%) | | | CTCAATCATACACATGACAACAAGTCATATTCGACTCCAAAACACTAACCAACCTTCTTCTTGCTTCTCAAAGCTTTCATGGTGTAGCCAAAGTCCATAT | 19 (0.000359%) | | | CTCAATCATACACATGACATCAAGTCATATTCGACTCCAAAACACTAACCAACCTTCTTCTTGCTTCTCAAAGCTTTCATGGTGTAGCCAAAGTCCATAT | 100 (0.001892%) | | | CTCACCATCCTTTGCTGATGCGGGACGGAAGCTGGTCTCCCGTGTGTTACCGCACGCGGTTGGCCTAAATCCGAGCCAAGGACGCCTGGAGCGTACCGAC | 21 (0.000397%) | | | CTCACCCGTTGACTCGCACACATGTCAGACTCCTTGGTCCGTGTTTCAAGACGGGTCGAATGGGGAGCCCACAGGCCGACGCCCGGAGCACGCTGATGCC | 17 (0.000322%) | | | CTCATATGGACTTTGGCTACACCATGAAAGCTTTGAGAAGCAAGAAGAAGGTTGGTTAGTGTTTTGGAGTCGAATATGACTTGATGTCATGTGTATGATT | 88 (0.001665%) | | | CTCATTAAATCAGTTATAGTTTGTTTGATGGTAACTACTACTCGGATAACCGTAGTAATTCTAGAGCTAATACGTGCAACAAACCCCGACTTATGGAAGG | 10 (0.000189%) | | | CTCATTCCAATTACCAGACTCGAAAGAGCCCGGTATTGTTATTTATTGTCACTACCTCCCCGTGTCAGGATTGGGTAATTTGCGCGCCTGCTGCCTTCCT | 15 (0.000284%) | | | CTCCGCTTATTGATATGCTTAAACTCAGCGGGTAATCCCGCCTGACCTGGGGTCGCTATATGGACTTTGGGTCATCTACAGCTTCCGGACAAGAGCGACC | 5 (0.000095%) | | | CTCCGGGCGTCGGCCTGTGGGCTCCCCATTCGACCCGTCTTGAAACACGGACCAAGGAGTCTGACATGTGTGCGAGTCAACGGGTGAGTAAACCCGTAAG | 21 (0.000397%) | | | CTCCGGGGAGGCGAATGCCAGCCGTTCGTTTGCATGTTCCTTGACACTTTTCGTGCCGGGGTTTTGTGATATCCGGAAGCAACGCGCACGACAAGACCGA | 41 (0.000776%) | | | CTCCGTTCCGCATCCGACCAGGACGCATCGCCGGCCCCCATCCGCTTCCCTCCCGACAATTTCAAGCACTCTTTGACTCTCTTTTCAAAGTCCTTTTCAT | 5 (0.000095%) | | | CTCGAAAGAGCCCGGTATTGTTATTTATTGTCACTACCTCCCCGTGTCAGGATTGGGTAATTTGCGCGCCTGCTGCCTTCCTTGGATGTGGTAGCCGTTT | 32 (0.000605%) | | | CTCGAAGAACTAATGGCAGCCCACGCAAGGCAAGCCCATTCTCCTCGACGATTCAGCAGTTTTTGTCCGAGAACTGCTGAGAAAACTCGGAAAAAGGCAA | 11 (0.000208%) | | | CTCGCCGTTACTAAGGGAATCCTTGTTAGTTTCTTTTCCTCCGCTTATTGATATGCTTAAACTCAGCGGGTAATCCCGCCTGACCTGGGGTCGCTATATG | 44 (0.000832%) | | | CTCGGCATCAGCGTGCTCCGGGCGTCGGCCTGTGGGCTCCCCATTCGACCCGTCTTGAAACACGGACCAAGGAGTCTGACATGTGTGCGAGTCAACGGGT | 21 (0.000397%) | | | CTCGTTAAGGGATTTAGATTGTACTCATTCCAATTACCAGACTCGAAAGAGCCCGGTATTGTTATTTATTGTCACTACCTCCCCGTGTCAGGATTGGGTA | 127 (0.002403%) | | | CTCTAGTCCGACTTTGTGAAATGACTTGAGAGGTGTAGGATAAGTGGGAGCTTCGGCGCAAGTGAAATACCACTACTTTTAACGTTATTTTACTTACTCC | 14 (0.000265%) | | | CTCTCAAACGCAATGGAAAGAGAGAAAGGACGAGGTCTTGACCGTCATCTTTTGCCCGAAGGACGGATGAGCTTTGGCGGGACTGAATCACTTCGAGTCA | 4 (0.000076%) | | | CTCTCTCTCTCTCTCTCTCTCTCTCTCTCTCTCTCTCTCT | 49 (0.000371%) | | | CTGAAAATCAGAATCAAACGAGCTTTTACCCTTTTGTTCCACACGAGATTTCTGTTCTCGTTGAGCTCATCTTAGGACACCTGCGTTATCTTTTAACAGA | 23 (0.000435%) | | | CTGATGCGGGACGGAAGCTGGTCTCCCGTGTGTTACCGCACGCGGTTGGCCTAAATCCGAGCCAAGGACGCCTGGAGCGTACCGACATGCGGTGGTGAAC | 25 (0.000473%) | | | CTGCAGCACGCGCCTAACGGCGTGCCTCGGCATCAGCGTGCTCCGGGCGTCGGCCTGTGGGCTCCCCATTCGACCCGTCTTGAAACACGGACCAAGGAGT | 2141 (0.040504%) | | | CTGCCAGTAGTCATATGCTTGTCTCAAAGATTAAGCCATGCATGTGTAAGTATGAACGAATTCAGACTGTGAAACTGCGAATGGCTCATTAAATCAGTTA | 1952 (0.036928%) | | | CTGCGAATGGCTCATTAAATCAGTTATAGTTTGTTTGATGGTAACTACTACTCGGATAACCGTAGTAATTCTAGAGCTAATACGTGCAACAAACCCCGAC | 29 (0.000549%) | | | CTGGGCCGCACGCGCGCTACACTGATGTATTCAACGAGTTCACACCTTGGCCGACAGGCCCGGGTAATCTTTGAAATTTCATCGTGATGGGGATAGATCG | 66 (0.001249%) | | | CTGGGGCGGCACATCTGTTAAAAGATAACGCAGGTGTCCTAAGATGAGCTCAACGAGAACAGAAATCTCGTGTGGAACAAAAGGGTAAAAGCTCGTTTGA | 11 (0.000208%) | | | CTGGTGCCAGCAGCCGCGGTAATTCCAGCTCCAATAGCGTATATTTAAGTTGTTGCAGTTAAAAAGCTCGTAGTTGAACCTTGGGATGGGTCGGCCGGTC | 69 (0.001305%) | | | CTGTCCCGAGTGTGAGCGAGGTGTGAGTGTCGCCCATGGGCATCGACACCTTGCGGCTAGGAACTGGAACGAGACGGGTAGCAAAGATTTCGAGTAGCAC | 14 (0.000265%) | | | CTGTCCCGAGTGTGAGCGAGGTGTGAGTGTCGCCCATGGGCATCGACACCTTGCGGCTAGGAACTGGAACGAGACGGGTGGCAAAGATTTCGAGTAGCAC | 22 (0.000416%) | | | CTGTGATGCCCTTAGATGTTCTGGGCCGCACGCGCGCTACACTGATGTATTCAACGAGTTCACACCTTGGCCGACAGGCCCGGGTAATCTTTGAAATTTC | 92 (0.001740%) | | | CTGTTAAAAGATAACGCAGGTGTCCTAAGATGAGCTCAACGAGAACAGAAATCTCGTGTGGAACAAAAGGGTAAAAGCTCGTTTGATTCTGATTTTCAGT | 5 (0.000095%) | | | CTGTTGAGCTTGACTCTAGTCCGACTTTGTGAAATGACTTGAGAGGTGTAGGATAAGTGGGAGCTTCGGCGCAAGTGAAATACCACTACTTTTAACGTTA | 13 (0.000246%) | | | CTTAAAAGCCTAAGTAGTGTTTCCTTGTTAGAAGACACAAAGCCAAAGACTCATATGGACTTTGGCTACACCATGAAAGCTTTGAGAAGCAAGAAGAAGG | 210 (0.003973%) | | | CTTAAAGGCGTAAGAATTGTATCCTTGTTAAAAGACACAAAGCCAAAGACTCATATGGACTTTGGCTACACCATGAAAGCTTTGAGAAGCAAGAAGAAGG | 238 (0.004503%) | | | CTTAAAGGCGTAAGAATTGTATCCTTGTTAGAAGACACAAAGCCAAAGACTCATATGGACTTTGGCTACACCATGAAAGCTTTGAGAAGCAAGAAGAAGG | 209 (0.003954%) | | | CTTAAGCGCGCGACCTATACCCGGCCGTCGGGGCAAGAGCCAGGCCTCGATGAGTAGGAGGGCGCGGCGGTCGCTGCAAAACCTAGGGCGCGAGCCCGGG | 52 (0.000984%) | | | CTTAGATGTTCTGGGCCGCACGCGCGCTACACTGATGTATTCAACGAGTTCACACCTTGGCCGACAGGCCCGGGTAATCTTTGAAATTTCATCGTGATGG | 110 (0.002081%) | | | CTTATACTCAATCATACACATGACATCAAGTCATATTCGACTCCAAAACACTAACCAACCTTCTTCTTGCTTCTCAAAGCTTTCATGGTGTAGCCAAAGT | 146 (0.002762%) | | | CTTATTGATATGCTTAAACTCAGCGGGTAATCCCGCCTGACCTGGGGTCGCTATATGGACTTTGGGTCATCTACAGCTTCCGGACAAGAGCGACCGATAA | 23 (0.000435%) | | | CTTCAACGAGGAATTCCTAGTAAGCGCGAGTCATCAGCTCGCGTTGACTACGTCCCTGCCCTTTGTACACACCGCCCGTCGCTCCTACCGATTGAATGAT | 58 (0.001097%) | | | CTTCCCTTGCCTACATTGTTCCATCGACCAGAGGCTGTTCACCTTGGAGACCTGATGCGGTTATGAGTACGACCGGGCGTGAGCGGCACTCGGTCCTCCG | 61 (0.001154%) | | | CTTCTCAAAGCTTTCATGGTGTAGCCAAAGTCCATATGAGTCTTTGGCTTTGTGTCTTCTAACAAGGAAACACTACTTAGGCTTATAAGATGCGGTTGCG | 10 (0.000189%) | | | CTTGACTCTAGTCCGACTTTGTGAAATGACTTGAGAGGTGTAGGATAAGTGGGAGCTTCGGCGCAAGTGAAATACCACTACTTTTAACGTTATTTTACTT | 67 (0.001268%) | | | CTTGATGTCATGTGTATGATTGAGTATAAGAACTTAAACC | 310 (0.002346%) | | | CTTGCCTACATTGTTCCATCGACCAGAGGCTGTTCACCTTGGAGACCTGATGCGGTTATGAGTACGACCGGGCGTGAGCGGCACTCGGTCCTCCGGATTT | 49 (0.000927%) | | | CTTGCTTCTCAAAGCTTTCATGGTGTAGCCAAAGTCCATATGAGTCTTTGGCTTTGTGTCTTCTAACAAGGAAACACTACTTAGGCTTATAAGATGCGGT | 66 (0.001249%) | | | CTTGCTTTGAGCACTCTAATTTCTTCAAAGTAACAGCGCCGGAGGCACGACCCGGCCAATTAAGACCAGGAGCGTATCGCCGACCGAAGGGACAAGCCGA | 69 (0.001305%) | | | CTTGTCTCAAAGATTAAGCCATGCATGTGTAAGTATGAACGAATTCAGACTGTGAAACTGCGAATGGCTCATTAAATCAGTTATAGTTTGTTTGATGGTA | 53 (0.001003%) | | | CTTGTTAGAAGACACAAAGCCAAAGACTCATATGGACTTTGGCTACACCATGAAAGCTTTGAGAAGCAAGAAGAAGGTTGGTTAGTGTTTTGGAGTCGAA | 120 (0.002270%) | | | CTTGTTAGTTTCTTTTCCTCCGCTTATTGATATGCTTAAACTCAGCGGGTAATCCCGCCTGACCTGGGGTCGCTATATGGACTTTGGGTCATCTACAGCT | 28 (0.000530%) | | | CTTTCACGGTTCGTATTCGTACTGAAAATCAGAATCAAACGAGCTTTTACCCTTTTGTTCCACACGAGATTTCTGTTCTCGTTGAGCTCATCTTAGGACA | 55 (0.001041%) | | | CTTTCCAACGAAGCACGCCCATCCAACCTAGGCGAGACAAGGGTTCACATTTCGTTCATCACCCTTGGCCGGCTATCGAACAGCCGGACTCCCATCAAAA | 15 (0.000284%) | | | CTTTGCTGATGCGGGACGGAAGCTGGTCTCCCGTGTGTTACCGCACGCGGTTGGCCTAAATCCGAGCCAAGGACGCCTGGAGCGTACCGACATGCGGTGG | 52 (0.000984%) | | | CTTTTATCTAATAAATGCGTCCCTTCCATAAGTCGGGGTTTGTTGCACGTATTAGCTCTAGAATTACTACGGTTATCCGAGTAGTAGTTACCATCAAACA | 83 (0.001570%) | | | CTTTTCCTCCGCTTATTGATATGCTTAAACTCAGCGGGTAATCCCGCCTGACCTGGGGTCGCTATATGGACTTTGGGTCATCTACAGCTTCCGGACAAGA | 33 (0.000624%) | | | GAAAATCAGAATCAAACGAGCTTTTACCCTTTTGTTCCACACGAGATTTCTGTTCTCGTTGAGCTCATCTTAGGACACCTGCGTTATCTTTTAACAGATG | 8 (0.000151%) | | | GAAACTGCGAATGGCTCATTAAATCAGTTATAGTTTGTTTGATGGTAACTACTACTCGGATAACCGTAGTAATTCTAGAGCTAATACGTGCAACAAACCC | 217 (0.004105%) | | | GAAAGAAGACCCTGTTGAGCTTGACTCTAGTCCGACTTTGTGAAATGACTTGAGAGGTGTAGGATAAGTGGGAGCTTCGGCGCAAGTGAAATACCACTAC | 41 (0.000776%) | | | GAAAGAGAGAAAGGACGAGGTCTTGACCGTCATCTTTTGCCCGAAGGACGGATGAGCTTTGGCGGGACTGAATCACTTCGAGTCACCGTCGACAACTTTT | 3 (0.000057%) | | | GAAAGAGCCCGGTATTGTTATTTATTGTCACTACCTCCCCGTGTCAGGATTGGGTAATTTGCGCGCCTGCTGCCTTCCTTGGATGTGGTAGCCGTTTCTC | 27 (0.000511%) | | | GAAATCCGCTAAGGAGTGTGTAACAACTCACCTGCCGAATCAACTAGCCCCGAAAATGGATGGCGCTTAAGCGCGCGACCTATACCCGGCCGTCGGGGCA | 27 (0.000511%) | | | GAAATCCTATGATGTTATCCCATGCTAATGTATCCAGAGCGTAGGCTTGCTTTGAGCACTCTAATTTCTTCAAAGTAACAGCGCCGGAGGCACGACCCGG | 179 (0.003386%) | | | GAAGACACAAAGCCAAAGACTCATATGGACTTTGGCTACACCATGAAAGCTTTGAGAAGCAAGAAGAAGGTTGGTTAGTGTTTTGGAGTCGAATATGACT | 42 (0.000795%) | | | GAAGACCCTGTTGAGCTTGACTCTAGTCCGACTTTGTGAAATGACTTGAGAGGTGTAGGATAAGTGGGAGCTTCGGCGCAAGTGAAATACCACTACTTTT | 13 (0.000246%) | | | GAAGAGCACACGTCTGAACTCCAGTCACGAGATTCCATCT | 23 (0.000174%) | | | GAAGCACGCCCATCCAACCTAGGCGAGACAAGGGTTCACATTTCGTTCATCACCCTTGGCCGGCTATCGAACAGCCGGACTCCCATCAAAAGATGGTTGC | 18 (0.000341%) | | | GAAGCACGCCCATCCAACCTAGGCGAGACAAGGGTTCACATTTCGTTCATCACCCTTGGCCGGCTTTCGAACAGCCGGACTCCCATCAAAAGATGGTTGC | 10 (0.000189%) | | | GAAGTCGAAATCCGCTAAGGAGTGTGTAACAACTCACCTGCCGAATCAACTAGCCCCGAAAATGGATGGCGCTTAAGCGCGCGACCTATACCCGGCCGTC | 4 (0.000076%) | | | GAATATGACTTGATGTCATGTGTATGATTGAGTATAAGAA | 480 (0.003632%) | | | GAATCAAACGAGCTTTTACCCTTTTGTTCCACACGAGATTTCTGTTCTCGTTGAGCTCATCTTAGGACACCTGCGTTATCTTTTAACAGATGTGCCGCCC | 3 (0.000057%) | | | GAATCATCAGAGCAACGGGCAGAGCCCGCGTCGACCTTTTATCTAATAAATGCGTCCCTTCCATAAGTCGGGGTTTGTTGCACGTATTAGCTCTAGAATT | 123 (0.002327%) | | | GAATCCTTGTTAGTTTCTTTTCCTCCGCTTATTGATATGCTTAAACTCAGCGGGTAATCCCGCCTGACCTGGGGTCGCTATATGGACTTTGGGTCATCTA | 6 (0.000114%) | | | GAATGCCAGCCGTTCGTTTGCATGTTCCTTGACACTTTTCGTGCCGGGGTTTTGTGATATCCGGAAGCAACGCGCACGACAAGACCGAGATAAAAGCTCC | 18 (0.000341%) | | | GAATGGCTCATTAAATCAGTTATAGTTTGTTTGATGGTAACTACTACTCGGATAACCGTAGTAATTCTAGAGCTAATACGTGCAACAAACCCCGACTTAT | 10 (0.000189%) | | | GAATTAGGGTTCGATTCCGGAGAGGGAGCCTGAGAAACGGCTACCACATCCAAGGAAGGCAGCAGGCGCGCAAATTACCCAATCCTGACACGGGGAGGTA | 10 (0.000189%) | | | GAATTGTATCCTTGTTAGAAGACACAAAGCCAAAGACTCATATGGACTTTGGCTACACCATGAAAGCTTTGAGAAGCAAGAAGAAGGTTGGTTAGTGTTT | 22 (0.000416%) | | | GACAAGGGTTCACATTTCGTTCATCACCCTTGGCCGGCTATCGAACAGCCGGACTCCCATCAAAAGATGGTTGCCAAGAACATCTTCGTTACGGTTTGCT | 11 (0.000208%) | | | GACAAGGGTTCACATTTCGTTCATCACCCTTGGCCGGCTTTCGAACAGCCGGACTCCCATCAAAAGATGGTTGCCAAGAACATCTTCGTTACGGTTTGCT | 12 (0.000227%) | | | GACACAAAGCCAAAGACTCATATGGACTTTGGCTACACCATGAAAGCTTTGAGAAGCAAGAAGAAGGTTGGTTAGTGTTTTGGAGTCGAATATGACTTGA | 53 (0.001003%) | | | GACATCAAGTCATATTCGACTCCAAAACACTAACCAACCTTCTTCTTGCTTCTCAAAGCTTTCATGGTGTAGCCAAAGTCCATATGAGTCTTTGGCTTTG | 42 (0.000795%) | | | GACATTGTCAGGTGGGGAGTTTGGCTGGGGCGGCACATCTGTTAAAAGATAACGCAGGTGTCCTAAGATGAGCTCAACGAGAACAGAAATCTCGTGTGGA | 33 (0.000624%) | | | GACCAGAGGCTGTTCACCTTGGAGACCTGATGCGGTTATGAGTACGACCGGGCGTGAGCGGCACTCGGTCCTCCGGATTTTCAAGGGCCGCCGGGGGCGC | 3 (0.000057%) | | | GACCCTGTTGAGCTTGACTCTAGTCCGACTTTGTGAAATGACTTGAGAGGTGTAGGATAAGTGGGAGCTTCGGCGCAAGTGAAATACCACTACTTTTAAC | 3 (0.000057%) | | | GACCGGATTGCTCCGTTCCGCATCCGACCAGGACGCATCGCCGGCCCCCATCCGCTTCCCTCCCGACAATTTCAAGCACTCTTTGACTCTCTTTTCAAAG | 9 (0.000170%) | | | GACCTATACCCGGCCGTCGGGGCAAGAGCCAGGCCTCGATGAGTAGGAGGGCGCGGCGGTCGCTGCAAAACCTAGGGCGCGAGCCCGGGCGGAGCGGCCG | 10 (0.000189%) | | | GACCTTTTATCTAATAAATGCGTCCCTTCCATAAGTCGGGGTTTGTTGCACGTATTAGCTCTAGAATTACTACGGTTATCCGAGTAGTAGTTACCATCAA | 10 (0.000189%) | | | GACGGAGAATTAGGGTTCGATTCCGGAGAGGGAGCCTGAGAAACGGCTACCACATCCAAGGAAGGCAGCAGGCGCGCAAATTACCCAATCCTGACACGGG | 7 (0.000132%) | | | GACGGGCGGTGTGTACAAAGGGCAGGGACGTAGTCAACGCGAGCTGATGACTCGCGCTTACTAGGAATTCCTCGTTGAAGACCAACAATTGCAATGATCG | 11 (0.000208%) | | | GACGTGGGTGGTTCGCCGCCCGCGACGTCGCGAGAAGTCCACTAAACCTTATCATTTAGAGGAAGGAGAAGTCGTAACAAGGTTTCCGTAGGTGAACCTG | 48 (0.000908%) | | | GACTCATATGGACTTTGGCTACACCATGAAAGCTTTGAGAAGCAAGAAGAAGGTTGGTTAGTGTTTTGGAGTCGAATATGACTTGATGTCATGTGTATGA | 4 (0.000076%) | | | GACTCCAAAACACTAACCAACCTTCTTCTTGCTTCTCAAAGCTTTCATGGTGTAGCCAAAGTCCATATGAGTCTTTGGCTTTGTGTCTTCTAACAAGGAA | 265 (0.005013%) | | | GACTCTAGTCCGACTTTGTGAAATGACTTGAGAGGTGTAGGATAAGTGGGAGCTTCGGCGCAAGTGAAATACCACTACTTTTAACGTTATTTTACTTACT | 4 (0.000076%) | | | GACTTCCCTTGCCTACATTGTTCCATCGACCAGAGGCTGTTCACCTTGGAGACCTGATGCGGTTATGAGTACGACCGGGCGTGAGCGGCACTCGGTCCTC | 19 (0.000359%) | | | GACTTGATGTCATGTGTATGATTGAGTATAAGAACTTAAA | 138 (0.001044%) | | | GACTTTGGCTACACCATGAAAGCTTTGAGAAGCAAGAAGAAGGTTGGTTAGTGTTTTGGAGTCGAATATGACTTGATGTCATGTGTATGATTGAGTATAA | 50 (0.000946%) | | | GACTTTGTGAAATGACTTGAGAGGTGTAGGATAAGTGGGAGCTTCGGCGCAAGTGAAATACCACTACTTTTAACGTTATTTTACTTACTCCGTGAATCGG | 6 (0.000114%) | | | GAGAATTAGGGTTCGATTCCGGAGAGGGAGCCTGAGAAACGGCTACCACATCCAAGGAAGGCAGCAGGCGCGCAAATTACCCAATCCTGACACGGGGAGG | 11 (0.000208%) | | | GAGACAAGGGTTCACATTTCGTTCATCACCCTTGGCCGGCTATCGAACAGCCGGACTCCCATCAAAAGATGGTTGCCAAGAACATCTTCGTTACGGTTTG | 6 (0.000114%) | | | GAGACAAGGGTTCACATTTCGTTCATCACCCTTGGCCGGCTTTCGAACAGCCGGACTCCCATCAAAAGATGGTTGCCAAGAACATCTTCGTTACGGTTTG | 8 (0.000151%) | | | GAGCAACGGGCAGAGCCCGCGTCGACCTTTTATCTAATAAATGCGTCCCTTCCATAAGTCGGGGTTTGTTGCACGTATTAGCTCTAGAATTACTACGGTT | 19 (0.000359%) | | | GAGCCCGCGTCGACCTTTTATCTAATAAATGCGTCCCTTCCATAAGTCGGGGTTTGTTGCACGTATTAGCTCTAGAATTACTACGGTTATCCGAGTAGTA | 7 (0.000132%) | | | GAGCGACGGGCGGTGTGTACAAAGGGCAGGGACGTAGTCAACGCGAGCTGATGACTCGCGCTTACTAGGAATTCCTCGTTGAAGACCAACAATTGCAATG | 15 (0.000284%) | | | GAGCGAGGTGTGAGTGTCGCCCATGGGCATCGACACCTTGCGGCTAGGAACTGGAACGAGACGGGTAGCAAAGATTTCGAGTAGCACTTCATACTACCGT | 7 (0.000132%) | | | GAGCGAGGTGTGAGTGTCGCCCATGGGCATCGACACCTTGCGGCTAGGAACTGGAACGAGACGGGTGGCAAAGATTTCGAGTAGCACTTCATACTACCGT | 4 (0.000076%) | | | GAGCGTAGGCTTGCTTTGAGCACTCTAATTTCTTCAAAGTAACAGCGCCGGAGGCACGACCCGGCCAATTAAGACCAGGAGCGTATCGCCGACCGAAGGG | 3 (0.000057%) | | | GAGCTTGACTCTAGTCCGACTTTGTGAAATGACTTGAGAGGTGTAGGATAAGTGGGAGCTTCGGCGCAAGTGAAATACCACTACTTTTAACGTTATTTTA | 4 (0.000076%) | | | GAGGACATTGTCAGGTGGGGAGTTTGGCTGGGGCGGCACATCTGTTAAAAGATAACGCAGGTGTCCTAAGATGAGCTCAACGAGAACAGAAATCTCGTGT | 50 (0.000946%) | | | GAGGCGAATGCCAGCCGTTCGTTTGCATGTTCCTTGACACTTTTCGTGCCGGGGTTTTGTGATATCCGGAAGCAACGCGCACGACAAGACCGAGATAAAA | 8 (0.000151%) | | | GAGGGCAAGTCTGGTGCCAGCAGCCGCGGTAATTCCAGCTCCAATAGCGTATATTTAAGTTGTTGCAGTTAAAAAGCTCGTAGTTGAACCTTGGGATGGG | 19 (0.000359%) | | | GAGGTGTGAGTGTCGCCCATGGGCATCGACACCTTGCGGCTAGGAACTGGAACGAGACGGGTGGCAAAGATTTCGAGTAGCACTTCATACTACCGTGGGT | 14 (0.000265%) | | | GAGTTATCATGAATCATCAGAGCAACGGGCAGAGCCCGCGTCGACCTTTTATCTAATAAATGCGTCCCTTCCATAAGTCGGGGTTTGTTGCACGTATTAG | 69 (0.001305%) | | | GAGTTTGGCTGGGGCGGCACATCTGTTAAAAGATAACGCAGGTGTCCTAAGATGAGCTCAACGAGAACAGAAATCTCGTGTGGAACAAAAGGGTAAAAGC | 3 (0.000057%) | | | GATACTAAATCCTATTTTCTGGTAAATTTTCATAATTTTTTGACACCTCTAGCTAGGTCATTTGACCTGATACAACATCGGATTTTCATGGTCTAGTTGG | 3 (0.000057%) | | | GATAGGCCACGCTTTCACGGTTCGTATTCGTACTGAAAATCAGAATCAAACGAGCTTTTACCCTTTTGTTCCACACGAGATTTCTGTTCTCGTTGAGCTC | 1738 (0.032880%) | | | GATAGTGGCCTACCATGGTGGTAACGGGTGACGGAGAATTAGGGTTCGATTCCGGAGAGGGAGCCTGAGAAACGGCTACCACATCCAAGGAAGGCAGCAG | 4 (0.000076%) | | | GATCAAGTTCACCACCGCATGTCGGTACGCTCCAGGCGTCCTTGGCTCGGATTTAGGCCAACCGCGTGCGGTAACACACGGGAGACCAGCTTCCGTCCCG | 57 (0.001078%) | | | GATCACGGCAATTCCCCGCCACATCCTCTCAAACGCAATGGAAAGAGAGAAAGGACGAGGTCTTGACCGTCATCTTTTGCCCGAAGGACGGATGAGCTTT | 31 (0.000586%) | | | GATCATTCAATCGGTAGGAGCGACGGGCGGTGTGTACAAAGGGCAGGGACGTAGTCAACGCGAGCTGATGACTCGCGCTTACTAGGAATTCCTCGTTGAA | 837 (0.015835%) | | | GATCCATGCTTTCCAACGAAGCACGCCCATCCAACCTAGGCGAGACAAGGGTTCACATTTCGTTCATCACCCTTGGCCGGCTATCGAACAGCCGGACTCC | 106 (0.002005%) | | | GATCCATTGGAGGGCAAGTCTGGTGCCAGCAGCCGCGGTAATTCCAGCTCCAATAGCGTATATTTAAGTTGTTGCAGTTAAAAAGCTCGTAGTTGAACCT | 1540 (0.029134%) | | | GATCCGCCTAGGCTGTCCCGAGTGTGAGCGAGGTGTGAGTGTCGCCCATGGGCATCGACACCTTGCGGCTAGGAACTGGAACGAGACGGGTAGCAAAGAT | 124 (0.002346%) | | | GATCCGCCTAGGCTGTCCCGAGTGTGAGCGAGGTGTGAGTGTCGCCCATGGGCATCGACACCTTGCGGCTAGGAACTGGAACGAGACGGGTGGCAAAGAT | 168 (0.003178%) | | | GATCCGGTTAAAATTCCGGAACCGGGACGTGGCGGTTGACGGCAACGTTAGGGAGTCCGGAGACGTCGGCGGGGGCCTCGGGAAGAGTTATCTTTTCTGT | 135 (0.002554%) | | | GATCCGTAACTTCGGGAAAAGGATTGGCTCTGAGGGCTGGGCTCGGGGGTCCCAGTTCCGAACCCGTCGGCTGTCAGCGGACTGCTCGAGCTGCTTCCGC | 29 (0.000549%) | | | GATCCGTCGAGTTATCATGAATCATCAGAGCAACGGGCAGAGCCCGCGTCGACCTTTTATCTAATAAATGCGTCCCTTCCATAAGTCGGGGTTTGTTGCA | 116 (0.002195%) | | | GATCCTCGTTAAGGGATTTAGATTGTACTCATTCCAATTACCAGACTCGAAAGAGCCCGGTATTGTTATTTATTGTCACTACCTCCCCGTGTCAGGATTG | 96 (0.001816%) | | | GATCGAAATCCTATGATGTTATCCCATGCTAATGTATCCAGAGCGTAGGCTTGCTTTGAGCACTCTAATTTCTTCAAAGTAACAGCGCCGGAGGCACGAC | 410 (0.007756%) | | | GATCGATCAA | 145598 (0.275446%) | | | GATCGATCAAGTTCACCACCGCATGTCGGTACGCTCCAGGCGTCCTTGGCTCGGATTTAGGCCAACCGCGTGCGGTAACACACGGGAGACCAGCTTCCGT | 1241 (0.023478%) | | | GATCGATCACGGCAATTCCCCGCCACATCCTCTCAAACGCAATGGAAAGAGAGAAAGGACGAGGTCTTGACCGTCATCTTTTGCCCGAAGGACGGATGAG | 1028 (0.019448%) | | | GATCGATCCATGCTTTCCAACGAAGCACGCCCATCCAACCTAGGCGAGACAAGGGTTCACATTTCGTTCATCACCCTTGGCCGGCTATCGAACAGCCGGA | 1450 (0.027431%) | | | GATCGATCCGCCTAGGCTGTCCCGAGTGTGAGCGAGGTGTGAGTGTCGCCCATGGGCATCGACACCTTGCGGCTAGGAACTGGAACGAGACGGGTAGCAA | 1037 (0.019618%) | | | GATCGATCCGCCTAGGCTGTCCCGAGTGTGAGCGAGGTGTGAGTGTCGCCCATGGGCATCGACACCTTGCGGCTAGGAACTGGAACGAGACGGGTGGCAA | 1584 (0.029966%) | | | GATCGATCCGGTTAAAATTCCGGAACCGGGACGTGGCGGTTGACGGCAACGTTAGGGAGTCCGGAGACGTCGGCGGGGGCCTCGGGAAGAGTTATCTTTT | 1600 (0.030269%) | | | GATCGATCCGTAACTTCGGGAAAAGGATTGGCTCTGAGGGCTGGGCTCGGGGGTCCCAGTTCCGAACCCGTCGGCTGTCAGCGGACTGCTCGAGCTGCTT | 1276 (0.024140%) | | | GATCGATCCGTCGAGTTATCATGAATCATCAGAGCAACGGGCAGAGCCCGCGTCGACCTTTTATCTAATAAATGCGTCCCTTCCATAAGTCGGGGTTTGT | 1106 (0.020924%) | | | GATCGATCCTCGTTAAGGGATTTAGATTGTACTCATTCCAATTACCAGACTCGAAAGAGCCCGGTATTGTTATTTATTGTCACTACCTCCCCGTGTCAGG | 1484 (0.028075%) | | | GATCGATCGAAATCCTATGATGTTATCCCATGCTAATGTATCCAGAGCGTAGGCTTGCTTTGAGCACTCTAATTTCTTCAAAGTAACAGCGCCGGAGGCA | 1275 (0.024121%) | | | GATCGATCGGCGGACCGGATTGCTCCGTTCCGCATCCGACCAGGACGCATCGCCGGCCCCCATCCGCTTCCCTCCCGACAATTTCAAGCACTCTTTGACT | 868 (0.016421%) | | | GATCGATCGGGTTGCGGTTTAAGTTGTTATACTCAATCAT | 4573 (0.034605%) | | | GATCGATCTT | 168028 (0.317879%) | | | GATCGGAAGAGCACACGTCTGAACTCCAGTCACGAGATTC | 35 (0.000265%) | | | GATCGGCGGACCGGATTGCTCCGTTCCGCATCCGACCAGGACGCATCGCCGGCCCCCATCCGCTTCCCTCCCGACAATTTCAAGCACTCTTTGACTCTCT | 15 (0.000284%) | | | GATCGGGTTGCGGTTTAAGTTGTTATACTCAATCATACAC | 1413 (0.010693%) | | | GATCTTAAAGGCGTAAGAATTGTATCCTTGTTAAAAGACACAAAGCCAAAGACTCATATGGACTTTGGCTACACCATGAAAGCTTTGAGAAGCAAGAAGA | 707 (0.013375%) | | | GATCTTAAAGGCGTAAGAATTGTATCCTTGTTAGAAGACACAAAGCCAAAGACTCATATGGACTTTGGCTACACCATGAAAGCTTTGAGAAGCAAGAAGA | 516 (0.009762%) | | | GATGCCCTTAGATGTTCTGGGCCGCACGCGCGCTACACTGATGTATTCAACGAGTTCACACCTTGGCCGACAGGCCCGGGTAATCTTTGAAATTTCATCG | 8 (0.000151%) | | | GATGTCATGTGTATGATTGAGTATAAGAACTTAAACCGCA | 172 (0.001302%) | | | GATGTTATCCCATGCTAATGTATCCAGAGCGTAGGCTTGCTTTGAGCACTCTAATTTCTTCAAAGTAACAGCGCCGGAGGCACGACCCGGCCAATTAAGA | 10 (0.000189%) | | | GATGTTCTGGGCCGCACGCGCGCTACACTGATGTATTCAACGAGTTCACACCTTGGCCGACAGGCCCGGGTAATCTTTGAAATTTCATCGTGATGGGGAT | 5 (0.000095%) | | | GATTAACAGGGACAGTCGGGGGCATTCGTATTTCATAGTCAGAGGTGAAATTCTTGGATTTATGAAAGACGAACAACTGCGAAAGCATTTGCCAAGGATG | 196 (0.003708%) | | | GATTAAGCCATGCATGTGTAAGTATGAACGAATTCAGACTGTGAAACTGCGAATGGCTCATTAAATCAGTTATAGTTTGTTTGATGGTAACTACTACTCG | 19 (0.000359%) | | | GATTGTACTCATTCCAATTACCAGACTCGAAAGAGCCCGGTATTGTTATTTATTGTCACTACCTCCCCGTGTCAGGATTGGGTAATTTGCGCGCCTGCTG | 7 (0.000132%) | | | GATTTAGATTGTACTCATTCCAATTACCAGACTCGAAAGAGCCCGGTATTGTTATTTATTGTCACTACCTCCCCGTGTCAGGATTGGGTAATTTGCGCGC | 18 (0.000341%) | | | GCAACCGGATCTTAAAGGCGTAAGAATTGTATCCTTGTTAGAAGACACAAAGCCAAAGACTCATATGGACTTTGGCTACACCATGAAAGCTTTGAGAAGC | 3 (0.000057%) | | | GCAACGGGCAGAGCCCGCGTCGACCTTTTATCTAATAAATGCGTCCCTTCCATAAGTCGGGGTTTGTTGCACGTATTAGCTCTAGAATTACTACGGTTAT | 40 (0.000757%) | | | GCAACGTCGCTATGAACGCTTGGCTGCCACAAGCCAGTTATCCCTGTGGTAACTTTTCTGACACCTCTAGCTTCAAATTCCGAAGGTCTAAAGGATCGAT | 61 (0.001154%) | | | GCAAGTCTGGTGCCAGCAGCCGCGGTAATTCCAGCTCCAATAGCGTATATTTAAGTTGTTGCAGTTAAAAAGCTCGTAGTTGAACCTTGGGATGGGTCGG | 26 (0.000492%) | | | GCAATAACAGGTCTGTGATGCCCTTAGATGTTCTGGGCCGCACGCGCGCTACACTGATGTATTCAACGAGTTCACACCTTGGCCGACAGGCCCGGGTAAT | 1004 (0.018994%) | | | GCAATTCCCCGCCACATCCTCTCAAACGCAATGGAAAGAGAGAAAGGACGAGGTCTTGACCGTCATCTTTTGCCCGAAGGACGGATGAGCTTTGGCGGGA | 32 (0.000605%) | | | GCACACGTCTGAACTCCAGTCACGAGATTCCATCTCGTAT | 6 (0.000045%) | | | GCACGCCCATCCAACCTAGGCGAGACAAGGGTTCACATTTCGTTCATCACCCTTGGCCGGCTATCGAACAGCCGGACTCCCATCAAAAGATGGTTGCCAA | 29 (0.000549%) | | | GCACGCCCATCCAACCTAGGCGAGACAAGGGTTCACATTTCGTTCATCACCCTTGGCCGGCTTTCGAACAGCCGGACTCCCATCAAAAGATGGTTGCCAA | 25 (0.000473%) | | | GCACGCGCCTAACGGCGTGCCTCGGCATCAGCGTGCTCCGGGCGTCGGCCTGTGGGCTCCCCATTCGACCCGTCTTGAAACACGGACCAAGGAGTCTGAC | 23 (0.000435%) | | | GCAGAGCCCGCGTCGACCTTTTATCTAATAAATGCGTCCCTTCCATAAGTCGGGGTTTGTTGCACGTATTAGCTCTAGAATTACTACGGTTATCCGAGTA | 18 (0.000341%) | | | GCAGCACGCGCCTAACGGCGTGCCTCGGCATCAGCGTGCTCCGGGCGTCGGCCTGTGGGCTCCCCATTCGACCCGTCTTGAAACACGGACCAAGGAGTCT | 461 (0.008721%) | | | GCAGCCGCGGTAATTCCAGCTCCAATAGCGTATATTTAAGTTGTTGCAGTTAAAAAGCTCGTAGTTGAACCTTGGGATGGGTCGGCCGGTCCGCCTTTGG | 10 (0.000189%) | | | GCATCAGCGTGCTCCGGGCGTCGGCCTGTGGGCTCCCCATTCGACCCGTCTTGAAACACGGACCAAGGAGTCTGACATGTGTGCGAGTCAACGGGTGAGT | 11 (0.000208%) | | | GCATCATAAGGATACTAAATCCTATTTTCTGGTAAATTTTCATAATTTTTTGACACCTCTAGCTAGGTCATTTGACCTGATACAACATCGGATTTTCATG | 11 (0.000208%) | | | GCATGTCGGTACGCTCCAGGCGTCCTTGGCTCGGATTTAGGCCAACCGCGTGCGGTAACACACGGGAGACCAGCTTCCGTCCCGCATCAGCAAAGGATGG | 4 (0.000076%) | | | GCATGTGTAAGTATGAACGAATTCAGACTGTGAAACTGCGAATGGCTCATTAAATCAGTTATAGTTTGTTTGATGGTAACTACTACTCGGATAACCGTAG | 8 (0.000151%) | | | GCCAAAGACTCATATGGACTTTGGCTACACCATGAAAGCTTTGAGAAGCAAGAAGAAGGTTGGTTAGTGTTTTGGAGTCGAATATGACTTGATGTCATGT | 46 (0.000870%) | | | GCCACATCCTCTCAAACGCAATGGAAAGAGAGAAAGGACGAGGTCTTGACCGTCATCTTTTGCCCGAAGGACGGATGAGCTTTGGCGGGACTGAATCACT | 17 (0.000322%) | | | GCCACCCTAACGCCTCGAAGAACTAATGGCAGCCCACGCAAGGCAAGCCCATTCTCCTCGACGATTCAGCAGTTTTTGTCCGAGAACTGCTGAGAAAACT | 868 (0.016421%) | | | GCCACGCTTTCACGGTTCGTATTCGTACTGAAAATCAGAATCAAACGAGCTTTTACCCTTTTGTTCCACACGAGATTTCTGTTCTCGTTGAGCTCATCTT | 167 (0.003159%) | | | GCCAGCAGCCGCGGTAATTCCAGCTCCAATAGCGTATATTTAAGTTGTTGCAGTTAAAAAGCTCGTAGTTGAACCTTGGGATGGGTCGGCCGGTCCGCCT | 30 (0.000568%) | | | GCCAGCCGTTCGTTTGCATGTTCCTTGACACTTTTCGTGCCGGGGTTTTGTGATATCCGGAAGCAACGCGCACGACAAGACCGAGATAAAAGCTCCCGAT | 36 (0.000681%) | | | GCCAGTAGTCATATGCTTGTCTCAAAGATTAAGCCATGCATGTGTAAGTATGAACGAATTCAGACTGTGAAACTGCGAATGGCTCATTAAATCAGTTATA | 233 (0.004408%) | | | GCCATGCATGTGTAAGTATGAACGAATTCAGACTGTGAAACTGCGAATGGCTCATTAAATCAGTTATAGTTTGTTTGATGGTAACTACTACTCGGATAAC | 10 (0.000189%) | | | GCCCATCCAACCTAGGCGAGACAAGGGTTCACATTTCGTTCATCACCCTTGGCCGGCTATCGAACAGCCGGACTCCCATCAAAAGATGGTTGCCAAGAAC | 47 (0.000889%) | | | GCCCATCCAACCTAGGCGAGACAAGGGTTCACATTTCGTTCATCACCCTTGGCCGGCTTTCGAACAGCCGGACTCCCATCAAAAGATGGTTGCCAAGAAC | 45 (0.000851%) | | | GCCCATGGGCATCGACACCTTGCGGCTAGGAACTGGAACGAGACGGGTGGCAAAGATTTCGAGTAGCACTTCATACTACCGTGGGTTTTTTAAACCTTCC | 10 (0.000189%) | | | GCCCGCGTCGACCTTTTATCTAATAAATGCGTCCCTTCCATAAGTCGGGGTTTGTTGCACGTATTAGCTCTAGAATTACTACGGTTATCCGAGTAGTAGT | 3 (0.000057%) | | | GCCCTTAGATGTTCTGGGCCGCACGCGCGCTACACTGATGTATTCAACGAGTTCACACCTTGGCCGACAGGCCCGGGTAATCTTTGAAATTTCATCGTGA | 14 (0.000265%) | | | GCCGACTTCCCTTGCCTACATTGTTCCATCGACCAGAGGCTGTTCACCTTGGAGACCTGATGCGGTTATGAGTACGACCGGGCGTGAGCGGCACTCGGTC | 228 (0.004313%) | | | GCCGCCCGCGACGTCGCGAGAAGTCCACTAAACCTTATCATTTAGAGGAAGGAGAAGTCGTAACAAGGTTTCCGTAGGTGAACCTGCGGAAGGATCGATC | 18 (0.000341%) | | | GCCGCGGTAATTCCAGCTCCAATAGCGTATATTTAAGTTGTTGCAGTTAAAAAGCTCGTAGTTGAACCTTGGGATGGGTCGGCCGGTCCGCCTTTGGTGT | 14 (0.000265%) | | | GCCGTCGGGGCAAGAGCCAGGCCTCGATGAGTAGGAGGGCGCGGCGGTCGCTGCAAAACCTAGGGCGCGAGCCCGGGCGGAGCGGCCGTCGGTGCAGATC | 9 (0.000170%) | | | GCCGTTACTAAGGGAATCCTTGTTAGTTTCTTTTCCTCCGCTTATTGATATGCTTAAACTCAGCGGGTAATCCCGCCTGACCTGGGGTCGCTATATGGAC | 22 (0.000416%) | | | GCCTAACGGCGTGCCTCGGCATCAGCGTGCTCCGGGCGTCGGCCTGTGGGCTCCCCATTCGACCCGTCTTGAAACACGGACCAAGGAGTCTGACATGTGT | 43 (0.000813%) | | | GCCTAAGTAGTGTTTCCTTGTTAGAAGACACAAAGCCAAAGACTCATATGGACTTTGGCTACACCATGAAAGCTTTGAGAAGCAAGAAGAAGGTTGGTTA | 49 (0.000927%) | | | GCCTACATTGTTCCATCGACCAGAGGCTGTTCACCTTGGAGACCTGATGCGGTTATGAGTACGACCGGGCGTGAGCGGCACTCGGTCCTCCGGATTTTCA | 6 (0.000114%) | | | GCCTAGGCTGTCCCGAGTGTGAGCGAGGTGTGAGTGTCGCCCATGGGCATCGACACCTTGCGGCTAGGAACTGGAACGAGACGGGTAGCAAAGATTTCGA | 267 (0.005051%) | | | GCCTAGGCTGTCCCGAGTGTGAGCGAGGTGTGAGTGTCGCCCATGGGCATCGACACCTTGCGGCTAGGAACTGGAACGAGACGGGTGGCAAAGATTTCGA | 379 (0.007170%) | | | GCCTCGAAGAACTAATGGCAGCCCACGCAAGGCAAGCCCATTCTCCTCGACGATTCAGCAGTTTTTGTCCGAGAACTGCTGAGAAAACTCGGAAAAAGGC | 13 (0.000246%) | | | GCCTCGGCATCAGCGTGCTCCGGGCGTCGGCCTGTGGGCTCCCCATTCGACCCGTCTTGAAACACGGACCAAGGAGTCTGACATGTGTGCGAGTCAACGG | 17 (0.000322%) | | | GCGAATGCCAGCCGTTCGTTTGCATGTTCCTTGACACTTTTCGTGCCGGGGTTTTGTGATATCCGGAAGCAACGCGCACGACAAGACCGAGATAAAAGCT | 27 (0.000511%) | | | GCGAATGGCTCATTAAATCAGTTATAGTTTGTTTGATGGTAACTACTACTCGGATAACCGTAGTAATTCTAGAGCTAATACGTGCAACAAACCCCGACTT | 22 (0.000416%) | | | GCGACCTATACCCGGCCGTCGGGGCAAGAGCCAGGCCTCGATGAGTAGGAGGGCGCGGCGGTCGCTGCAAAACCTAGGGCGCGAGCCCGGGCGGAGCGGC | 112 (0.002119%) | | | GCGACGGGCGGTGTGTACAAAGGGCAGGGACGTAGTCAACGCGAGCTGATGACTCGCGCTTACTAGGAATTCCTCGTTGAAGACCAACAATTGCAATGAT | 47 (0.000889%) | | | GCGACGTGGGTGGTTCGCCGCCCGCGACGTCGCGAGAAGTCCACTAAACCTTATCATTTAGAGGAAGGAGAAGTCGTAACAAGGTTTCCGTAGGTGAACC | 2219 (0.041980%) | | | GCGAGACAAGGGTTCACATTTCGTTCATCACCCTTGGCCGGCTATCGAACAGCCGGACTCCCATCAAAAGATGGTTGCCAAGAACATCTTCGTTACGGTT | 31 (0.000586%) | | | GCGAGACAAGGGTTCACATTTCGTTCATCACCCTTGGCCGGCTTTCGAACAGCCGGACTCCCATCAAAAGATGGTTGCCAAGAACATCTTCGTTACGGTT | 28 (0.000530%) | | | GCGAGGTGTGAGTGTCGCCCATGGGCATCGACACCTTGCGGCTAGGAACTGGAACGAGACGGGTAGCAAAGATTTCGAGTAGCACTTCATACTACCGTGG | 26 (0.000492%) | | | GCGAGGTGTGAGTGTCGCCCATGGGCATCGACACCTTGCGGCTAGGAACTGGAACGAGACGGGTGGCAAAGATTTCGAGTAGCACTTCATACTACCGTGG | 44 (0.000832%) | | | GCGCCTAACGGCGTGCCTCGGCATCAGCGTGCTCCGGGCGTCGGCCTGTGGGCTCCCCATTCGACCCGTCTTGAAACACGGACCAAGGAGTCTGACATGT | 24 (0.000454%) | | | GCGCGACCTATACCCGGCCGTCGGGGCAAGAGCCAGGCCTCGATGAGTAGGAGGGCGCGGCGGTCGCTGCAAAACCTAGGGCGCGAGCCCGGGCGGAGCG | 30 (0.000568%) | | | GCGGACCGGATTGCTCCGTTCCGCATCCGACCAGGACGCATCGCCGGCCCCCATCCGCTTCCCTCCCGACAATTTCAAGCACTCTTTGACTCTCTTTTCA | 17 (0.000322%) | | | GCGGAGGACATTGTCAGGTGGGGAGTTTGGCTGGGGCGGCACATCTGTTAAAAGATAACGCAGGTGTCCTAAGATGAGCTCAACGAGAACAGAAATCTCG | 1465 (0.027715%) | | | GCGGCACATCTGTTAAAAGATAACGCAGGTGTCCTAAGATGAGCTCAACGAGAACAGAAATCTCGTGTGGAACAAAAGGGTAAAAGCTCGTTTGATTCTG | 15 (0.000284%) | | | GCGGTAATTCCAGCTCCAATAGCGTATATTTAAGTTGTTGCAGTTAAAAAGCTCGTAGTTGAACCTTGGGATGGGTCGGCCGGTCCGCCTTTGGTGTGCA | 22 (0.000416%) | | | GCGGTTTAAGTTCTTATACTCAATCATACACATGACATCA | 640 (0.004843%) | | | GCGTAAGAATTGTATCCTTGTTAGAAGACACAAAGCCAAAGACTCATATGGACTTTGGCTACACCATGAAAGCTTTGAGAAGCAAGAAGAAGGTTGGTTA | 12 (0.000227%) | | | GCGTAGGCTTGCTTTGAGCACTCTAATTTCTTCAAAGTAACAGCGCCGGAGGCACGACCCGGCCAATTAAGACCAGGAGCGTATCGCCGACCGAAGGGAC | 49 (0.000927%) | | | GCGTCGACCTTTTATCTAATAAATGCGTCCCTTCCATAAGTCGGGGTTTGTTGCACGTATTAGCTCTAGAATTACTACGGTTATCCGAGTAGTAGTTACC | 11 (0.000208%) | | | GCGTCGGCCTGTGGGCTCCCCATTCGACCCGTCTTGAAACACGGACCAAGGAGTCTGACATGTGTGCGAGTCAACGGGTGAGTAAACCCGTAAGGCGCAA | 5 (0.000095%) | | | GCGTGCCTCGGCATCAGCGTGCTCCGGGCGTCGGCCTGTGGGCTCCCCATTCGACCCGTCTTGAAACACGGACCAAGGAGTCTGACATGTGTGCGAGTCA | 5 (0.000095%) | | | GCGTGCTCCGGGCGTCGGCCTGTGGGCTCCCCATTCGACCCGTCTTGAAACACGGACCAAGGAGTCTGACATGTGTGCGAGTCAACGGGTGAGTAAACCC | 21 (0.000397%) | | | GCTAAGGAGTGTGTAACAACTCACCTGCCGAATCAACTAGCCCCGAAAATGGATGGCGCTTAAGCGCGCGACCTATACCCGGCCGTCGGGGCAAGAGCCA | 58 (0.001097%) | | | GCTAATGTATCCAGAGCGTAGGCTTGCTTTGAGCACTCTAATTTCTTCAAAGTAACAGCGCCGGAGGCACGACCCGGCCAATTAAGACCAGGAGCGTATC | 20 (0.000378%) | | | GCTCATTAAATCAGTTATAGTTTGTTTGATGGTAACTACTACTCGGATAACCGTAGTAATTCTAGAGCTAATACGTGCAACAAACCCCGACTTATGGAAG | 12 (0.000227%) | | | GCTCCGGGCGTCGGCCTGTGGGCTCCCCATTCGACCCGTCTTGAAACACGGACCAAGGAGTCTGACATGTGTGCGAGTCAACGGGTGAGTAAACCCGTAA | 7 (0.000132%) | | | GCTCCGTTCCGCATCCGACCAGGACGCATCGCCGGCCCCCATCCGCTTCCCTCCCGACAATTTCAAGCACTCTTTGACTCTCTTTTCAAAGTCCTTTTCA | 7 (0.000132%) | | | GCTCGCCGTTACTAAGGGAATCCTTGTTAGTTTCTTTTCCTCCGCTTATTGATATGCTTAAACTCAGCGGGTAATCCCGCCTGACCTGGGGTCGCTATAT | 59 (0.001116%) | | | GCTGATGCGGGACGGAAGCTGGTCTCCCGTGTGTTACCGCACGCGGTTGGCCTAAATCCGAGCCAAGGACGCCTGGAGCGTACCGACATGCGGTGGTGAA | 7 (0.000132%) | | | GCTGGGGCGGCACATCTGTTAAAAGATAACGCAGGTGTCCTAAGATGAGCTCAACGAGAACAGAAATCTCGTGTGGAACAAAAGGGTAAAAGCTCGTTTG | 10 (0.000189%) | | | GCTGTCCCGAGTGTGAGCGAGGTGTGAGTGTCGCCCATGGGCATCGACACCTTGCGGCTAGGAACTGGAACGAGACGGGTAGCAAAGATTTCGAGTAGCA | 62 (0.001173%) | | | GCTGTCCCGAGTGTGAGCGAGGTGTGAGTGTCGCCCATGGGCATCGACACCTTGCGGCTAGGAACTGGAACGAGACGGGTGGCAAAGATTTCGAGTAGCA | 99 (0.001873%) | | | GCTTAAGCGCGCGACCTATACCCGGCCGTCGGGGCAAGAGCCAGGCCTCGATGAGTAGGAGGGCGCGGCGGTCGCTGCAAAACCTAGGGCGCGAGCCCGG | 1065 (0.020148%) | | | GCTTATTGATATGCTTAAACTCAGCGGGTAATCCCGCCTGACCTGGGGTCGCTATATGGACTTTGGGTCATCTACAGCTTCCGGACAAGAGCGACCGATA | 21 (0.000397%) | | | GCTTCTCAAAGCTTTCATGGTGTAGCCAAAGTCCATATGAGTCTTTGGCTTTGTGTCTTCTAACAAGGAAACACTACTTAGGCTTATAAGATGCGGTTGC | 14 (0.000265%) | | | GCTTGACTCTAGTCCGACTTTGTGAAATGACTTGAGAGGTGTAGGATAAGTGGGAGCTTCGGCGCAAGTGAAATACCACTACTTTTAACGTTATTTTACT | 32 (0.000605%) | | | GCTTGCTTTGAGCACTCTAATTTCTTCAAAGTAACAGCGCCGGAGGCACGACCCGGCCAATTAAGACCAGGAGCGTATCGCCGACCGAAGGGACAAGCCG | 16 (0.000303%) | | | GCTTGTCTCAAAGATTAAGCCATGCATGTGTAAGTATGAACGAATTCAGACTGTGAAACTGCGAATGGCTCATTAAATCAGTTATAGTTTGTTTGATGGT | 19 (0.000359%) | | | GCTTTCACGGTTCGTATTCGTACTGAAAATCAGAATCAAACGAGCTTTTACCCTTTTGTTCCACACGAGATTTCTGTTCTCGTTGAGCTCATCTTAGGAC | 13 (0.000246%) | | | GCTTTCCAACGAAGCACGCCCATCCAACCTAGGCGAGACAAGGGTTCACATTTCGTTCATCACCCTTGGCCGGCTATCGAACAGCCGGACTCCCATCAAA | 76 (0.001438%) | | | GGAAAGAAGACCCTGTTGAGCTTGACTCTAGTCCGACTTTGTGAAATGACTTGAGAGGTGTAGGATAAGTGGGAGCTTCGGCGCAAGTGAAATACCACTA | 760 (0.014378%) | | | GGAAAGAGAGAAAGGACGAGGTCTTGACCGTCATCTTTTGCCCGAAGGACGGATGAGCTTTGGCGGGACTGAATCACTTCGAGTCACCGTCGACAACTTT | 5 (0.000095%) | | | GGAAGAGCACACGTCTGAACTCCAGTCACGAGATTCCATC | 50 (0.000378%) | | | GGAAGTCGAAATCCGCTAAGGAGTGTGTAACAACTCACCTGCCGAATCAACTAGCCCCGAAAATGGATGGCGCTTAAGCGCGCGACCTATACCCGGCCGT | 23 (0.000435%) | | | GGAATCCTTGTTAGTTTCTTTTCCTCCGCTTATTGATATGCTTAAACTCAGCGGGTAATCCCGCCTGACCTGGGGTCGCTATATGGACTTTGGGTCATCT | 18 (0.000341%) | | | GGACAGTCGGGGGCATTCGTATTTCATAGTCAGAGGTGAAATTCTTGGATTTATGAAAGACGAACAACTGCGAAAGCATTTGCCAAGGATGTTTTCATTA | 66 (0.001249%) | | | GGACATTGTCAGGTGGGGAGTTTGGCTGGGGCGGCACATCTGTTAAAAGATAACGCAGGTGTCCTAAGATGAGCTCAACGAGAACAGAAATCTCGTGTGG | 184 (0.003481%) | | | GGACCGGATTGCTCCGTTCCGCATCCGACCAGGACGCATCGCCGGCCCCCATCCGCTTCCCTCCCGACAATTTCAAGCACTCTTTGACTCTCTTTTCAAA | 24 (0.000454%) | | | GGACTTTGGCTACACCATGAAAGCTTTGAGAAGCAAGAAGAAGGTTGGTTAGTGTTTTGGAGTCGAATATGACTTGATGTCATGTGTATGATTGAGTATA | 69 (0.001305%) | | | GGAGAATTAGGGTTCGATTCCGGAGAGGGAGCCTGAGAAACGGCTACCACATCCAAGGAAGGCAGCAGGCGCGCAAATTACCCAATCCTGACACGGGGAG | 102 (0.001930%) | | | GGAGCGACGGGCGGTGTGTACAAAGGGCAGGGACGTAGTCAACGCGAGCTGATGACTCGCGCTTACTAGGAATTCCTCGTTGAAGACCAACAATTGCAAT | 56 (0.001059%) | | | GGAGGACATTGTCAGGTGGGGAGTTTGGCTGGGGCGGCACATCTGTTAAAAGATAACGCAGGTGTCCTAAGATGAGCTCAACGAGAACAGAAATCTCGTG | 411 (0.007775%) | | | GGAGGCGAATGCCAGCCGTTCGTTTGCATGTTCCTTGACACTTTTCGTGCCGGGGTTTTGTGATATCCGGAAGCAACGCGCACGACAAGACCGAGATAAA | 13 (0.000246%) | | | GGAGGGCAAGTCTGGTGCCAGCAGCCGCGGTAATTCCAGCTCCAATAGCGTATATTTAAGTTGTTGCAGTTAAAAAGCTCGTAGTTGAACCTTGGGATGG | 45 (0.000851%) | | | GGAGTTTGGCTGGGGCGGCACATCTGTTAAAAGATAACGCAGGTGTCCTAAGATGAGCTCAACGAGAACAGAAATCTCGTGTGGAACAAAAGGGTAAAAG | 6 (0.000114%) | | | GGATACTAAATCCTATTTTCTGGTAAATTTTCATAATTTTTTGACACCTCTAGCTAGGTCATTTGACCTGATACAACATCGGATTTTCATGGTCTAGTTG | 12 (0.000227%) | | | GGATAGTGGCCTACCATGGTGGTAACGGGTGACGGAGAATTAGGGTTCGATTCCGGAGAGGGAGCCTGAGAAACGGCTACCACATCCAAGGAAGGCAGCA | 715 (0.013527%) | | | GGATCTTAAAGGCGTAAGAATTGTATCCTTGTTAGAAGACACAAAGCCAAAGACTCATATGGACTTTGGCTACACCATGAAAGCTTTGAGAAGCAAGAAG | 22 (0.000416%) | | | GGATTGCTCCGTTCCGCATCCGACCAGGACGCATCGCCGGCCCCCATCCGCTTCCCTCCCGACAATTTCAAGCACTCTTTGACTCTCTTTTCAAAGTCCT | 12 (0.000227%) | | | GGCAAGTCTGGTGCCAGCAGCCGCGGTAATTCCAGCTCCAATAGCGTATATTTAAGTTGTTGCAGTTAAAAAGCTCGTAGTTGAACCTTGGGATGGGTCG | 62 (0.001173%) | | | GGCAATTCCCCGCCACATCCTCTCAAACGCAATGGAAAGAGAGAAAGGACGAGGTCTTGACCGTCATCTTTTGCCCGAAGGACGGATGAGCTTTGGCGGG | 82 (0.001551%) | | | GGCACATCTGTTAAAAGATAACGCAGGTGTCCTAAGATGAGCTCAACGAGAACAGAAATCTCGTGTGGAACAAAAGGGTAAAAGCTCGTTTGATTCTGAT | 20 (0.000378%) | | | GGCAGAGCCCGCGTCGACCTTTTATCTAATAAATGCGTCCCTTCCATAAGTCGGGGTTTGTTGCACGTATTAGCTCTAGAATTACTACGGTTATCCGAGT | 20 (0.000378%) | | | GGCATCAGCGTGCTCCGGGCGTCGGCCTGTGGGCTCCCCATTCGACCCGTCTTGAAACACGGACCAAGGAGTCTGACATGTGTGCGAGTCAACGGGTGAG | 10 (0.000189%) | | | GGCATCGACACCTTGCGGCTAGGAACTGGAACGAGACGGGTGGCAAAGATTTCGAGTAGCACTTCATACTACCGTGGGTTTTTTAAACCTTCCGAGTTTT | 13 (0.000246%) | | | GGCATGCATCATAAGGATACTAAATCCTATTTTCTGGTAAATTTTCATAATTTTTTGACACCTCTAGCTAGGTCATTTGACCTGATACAACATCGGATTT | 3 (0.000057%) | | | GGCATTCGTATTTCATAGTCAGAGGTGAAATTCTTGGATTTATGAAAGACGAACAACTGCGAAAGCATTTGCCAAGGATGTTTTCATTAATCAAGAACGA | 4 (0.000076%) | | | GGCCACGCTTTCACGGTTCGTATTCGTACTGAAAATCAGAATCAAACGAGCTTTTACCCTTTTGTTCCACACGAGATTTCTGTTCTCGTTGAGCTCATCT | 38 (0.000719%) | | | GGCCGCACGCGCGCTACACTGATGTATTCAACGAGTTCACACCTTGGCCGACAGGCCCGGGTAATCTTTGAAATTTCATCGTGATGGGGATAGATCGATC | 5 (0.000095%) | | | GGCCGTCGGGGCAAGAGCCAGGCCTCGATGAGTAGGAGGGCGCGGCGGTCGCTGCAAAACCTAGGGCGCGAGCCCGGGCGGAGCGGCCGTCGGTGCAGAT | 16 (0.000303%) | | | GGCCTACCATGGTGGTAACGGGTGACGGAGAATTAGGGTTCGATTCCGGAGAGGGAGCCTGAGAAACGGCTACCACATCCAAGGAAGGCAGCAGGCGCGC | 20 (0.000378%) | | | GGCGAATGCCAGCCGTTCGTTTGCATGTTCCTTGACACTTTTCGTGCCGGGGTTTTGTGATATCCGGAAGCAACGCGCACGACAAGACCGAGATAAAAGC | 48 (0.000908%) | | | GGCGAGACAAGGGTTCACATTTCGTTCATCACCCTTGGCCGGCTATCGAACAGCCGGACTCCCATCAAAAGATGGTTGCCAAGAACATCTTCGTTACGGT | 18 (0.000341%) | | | GGCGAGACAAGGGTTCACATTTCGTTCATCACCCTTGGCCGGCTTTCGAACAGCCGGACTCCCATCAAAAGATGGTTGCCAAGAACATCTTCGTTACGGT | 17 (0.000322%) | | | GGCGGACCGGATTGCTCCGTTCCGCATCCGACCAGGACGCATCGCCGGCCCCCATCCGCTTCCCTCCCGACAATTTCAAGCACTCTTTGACTCTCTTTTC | 23 (0.000435%) | | | GGCGGCACATCTGTTAAAAGATAACGCAGGTGTCCTAAGATGAGCTCAACGAGAACAGAAATCTCGTGTGGAACAAAAGGGTAAAAGCTCGTTTGATTCT | 5 (0.000095%) | | | GGCGTAAGAATTGTATCCTTGTTAGAAGACACAAAGCCAAAGACTCATATGGACTTTGGCTACACCATGAAAGCTTTGAGAAGCAAGAAGAAGGTTGGTT | 11 (0.000208%) | | | GGCGTGCCTCGGCATCAGCGTGCTCCGGGCGTCGGCCTGTGGGCTCCCCATTCGACCCGTCTTGAAACACGGACCAAGGAGTCTGACATGTGTGCGAGTC | 17 (0.000322%) | | | GGCTCATTAAATCAGTTATAGTTTGTTTGATGGTAACTACTACTCGGATAACCGTAGTAATTCTAGAGCTAATACGTGCAACAAACCCCGACTTATGGAA | 10 (0.000189%) | | | GGCTGGGGCGGCACATCTGTTAAAAGATAACGCAGGTGTCCTAAGATGAGCTCAACGAGAACAGAAATCTCGTGTGGAACAAAAGGGTAAAAGCTCGTTT | 6 (0.000114%) | | | GGCTGTCCCGAGTGTGAGCGAGGTGTGAGTGTCGCCCATGGGCATCGACACCTTGCGGCTAGGAACTGGAACGAGACGGGTAGCAAAGATTTCGAGTAGC | 18 (0.000341%) | | | GGCTGTCCCGAGTGTGAGCGAGGTGTGAGTGTCGCCCATGGGCATCGACACCTTGCGGCTAGGAACTGGAACGAGACGGGTGGCAAAGATTTCGAGTAGC | 20 (0.000378%) | | | GGCTTGCTTTGAGCACTCTAATTTCTTCAAAGTAACAGCGCCGGAGGCACGACCCGGCCAATTAAGACCAGGAGCGTATCGCCGACCGAAGGGACAAGCC | 7 (0.000132%) | | | GGGAATCCTTGTTAGTTTCTTTTCCTCCGCTTATTGATATGCTTAAACTCAGCGGGTAATCCCGCCTGACCTGGGGTCGCTATATGGACTTTGGGTCATC | 61 (0.001154%) | | | GGGACAGTCGGGGGCATTCGTATTTCATAGTCAGAGGTGAAATTCTTGGATTTATGAAAGACGAACAACTGCGAAAGCATTTGCCAAGGATGTTTTCATT | 33 (0.000624%) | | | GGGAGGCGAATGCCAGCCGTTCGTTTGCATGTTCCTTGACACTTTTCGTGCCGGGGTTTTGTGATATCCGGAAGCAACGCGCACGACAAGACCGAGATAA | 14 (0.000265%) | | | GGGAGTTTGGCTGGGGCGGCACATCTGTTAAAAGATAACGCAGGTGTCCTAAGATGAGCTCAACGAGAACAGAAATCTCGTGTGGAACAAAAGGGTAAAA | 42 (0.000795%) | | | GGGATTTAGATTGTACTCATTCCAATTACCAGACTCGAAAGAGCCCGGTATTGTTATTTATTGTCACTACCTCCCCGTGTCAGGATTGGGTAATTTGCGC | 42 (0.000795%) | | | GGGCAAGTCTGGTGCCAGCAGCCGCGGTAATTCCAGCTCCAATAGCGTATATTTAAGTTGTTGCAGTTAAAAAGCTCGTAGTTGAACCTTGGGATGGGTC | 161 (0.003046%) | | | GGGCAGAGCCCGCGTCGACCTTTTATCTAATAAATGCGTCCCTTCCATAAGTCGGGGTTTGTTGCACGTATTAGCTCTAGAATTACTACGGTTATCCGAG | 49 (0.000927%) | | | GGGCATCGACACCTTGCGGCTAGGAACTGGAACGAGACGGGTGGCAAAGATTTCGAGTAGCACTTCATACTACCGTGGGTTTTTTAAACCTTCCGAGTTT | 12 (0.000227%) | | | GGGCATTCGTATTTCATAGTCAGAGGTGAAATTCTTGGATTTATGAAAGACGAACAACTGCGAAAGCATTTGCCAAGGATGTTTTCATTAATCAAGAACG | 35 (0.000662%) | | | GGGCCGCACGCGCGCTACACTGATGTATTCAACGAGTTCACACCTTGGCCGACAGGCCCGGGTAATCTTTGAAATTTCATCGTGATGGGGATAGATCGAT | 7 (0.000132%) | | | GGGCGGCACATCTGTTAAAAGATAACGCAGGTGTCCTAAGATGAGCTCAACGAGAACAGAAATCTCGTGTGGAACAAAAGGGTAAAAGCTCGTTTGATTC | 5 (0.000095%) | | | GGGCGGTGTGTACAAAGGGCAGGGACGTAGTCAACGCGAGCTGATGACTCGCGCTTACTAGGAATTCCTCGTTGAAGACCAACAATTGCAATGATCGATC | 32 (0.000605%) | | | GGGCGTCGGCCTGTGGGCTCCCCATTCGACCCGTCTTGAAACACGGACCAAGGAGTCTGACATGTGTGCGAGTCAACGGGTGAGTAAACCCGTAAGGCGC | 7 (0.000132%) | | | GGGGAGGCGAATGCCAGCCGTTCGTTTGCATGTTCCTTGACACTTTTCGTGCCGGGGTTTTGTGATATCCGGAAGCAACGCGCACGACAAGACCGAGATA | 33 (0.000624%) | | | GGGGAGTTTGGCTGGGGCGGCACATCTGTTAAAAGATAACGCAGGTGTCCTAAGATGAGCTCAACGAGAACAGAAATCTCGTGTGGAACAAAAGGGTAAA | 33 (0.000624%) | | | GGGGCATTCGTATTTCATAGTCAGAGGTGAAATTCTTGGATTTATGAAAGACGAACAACTGCGAAAGCATTTGCCAAGGATGTTTTCATTAATCAAGAAC | 37 (0.000700%) | | | GGGGCGGCACATCTGTTAAAAGATAACGCAGGTGTCCTAAGATGAGCTCAACGAGAACAGAAATCTCGTGTGGAACAAAAGGGTAAAAGCTCGTTTGATT | 5 (0.000095%) | | | GGGGGCATTCGTATTTCATAGTCAGAGGTGAAATTCTTGGATTTATGAAAGACGAACAACTGCGAAAGCATTTGCCAAGGATGTTTTCATTAATCAAGAA | 18 (0.000341%) | | | GGGTGACGGAGAATTAGGGTTCGATTCCGGAGAGGGAGCCTGAGAAACGGCTACCACATCCAAGGAAGGCAGCAGGCGCGCAAATTACCCAATCCTGACA | 17 (0.000322%) | | | GGGTGGTTCGCCGCCCGCGACGTCGCGAGAAGTCCACTAAACCTTATCATTTAGAGGAAGGAGAAGTCGTAACAAGGTTTCCGTAGGTGAACCTGCGGAA | 189 (0.003576%) | | | GGGTTGCGGTTTAAGTTGTTATACTCAATCATACACATGA | 465 (0.003519%) | | | GGGTTTACTCACCCGTTGACTCGCACACATGTCAGACTCCTTGGTCCGTGTTTCAAGACGGGTCGAATGGGGAGCCCACAGGCCGACGCCCGGAGCACGC | 6 (0.000114%) | | | GGTAAATTTTCATAATTTTTTGACACCTCTAGCTAGGTCATTTGACCTGATACAACATCGGATTTTCATGGTCTAGTTGGGGCTCCGTGGGCATATTTGA | 5 (0.000095%) | | | GGTAACGGGTGACGGAGAATTAGGGTTCGATTCCGGAGAGGGAGCCTGAGAAACGGCTACCACATCCAAGGAAGGCAGCAGGCGCGCAAATTACCCAATC | 83 (0.001570%) | | | GGTAATTCCAGCTCCAATAGCGTATATTTAAGTTGTTGCAGTTAAAAAGCTCGTAGTTGAACCTTGGGATGGGTCGGCCGGTCCGCCTTTGGTGTGCATT | 99 (0.001873%) | | | GGTAGGAGCGACGGGCGGTGTGTACAAAGGGCAGGGACGTAGTCAACGCGAGCTGATGACTCGCGCTTACTAGGAATTCCTCGTTGAAGACCAACAATTG | 41 (0.000776%) | | | GGTCTGTGATGCCCTTAGATGTTCTGGGCCGCACGCGCGCTACACTGATGTATTCAACGAGTTCACACCTTGGCCGACAGGCCCGGGTAATCTTTGAAAT | 13 (0.000246%) | | | GGTCTTCAACGAGGAATTCCTAGTAAGCGCGAGTCATCAGCTCGCGTTGACTACGTCCCTGCCCTTTGTACACACCGCCCGTCGCTCCTACCGATTGAAT | 7 (0.000132%) | | | GGTGCCAGCAGCCGCGGTAATTCCAGCTCCAATAGCGTATATTTAAGTTGTTGCAGTTAAAAAGCTCGTAGTTGAACCTTGGGATGGGTCGGCCGGTCCG | 12 (0.000227%) | | | GGTGGGGAGTTTGGCTGGGGCGGCACATCTGTTAAAAGATAACGCAGGTGTCCTAAGATGAGCTCAACGAGAACAGAAATCTCGTGTGGAACAAAAGGGT | 5 (0.000095%) | | | GGTGGTTCGCCGCCCGCGACGTCGCGAGAAGTCCACTAAACCTTATCATTTAGAGGAAGGAGAAGTCGTAACAAGGTTTCCGTAGGTGAACCTGCGGAAG | 28 (0.000530%) | | | GGTGTGAGTGTCGCCCATGGGCATCGACACCTTGCGGCTAGGAACTGGAACGAGACGGGTAGCAAAGATTTCGAGTAGCACTTCATACTACCGTGGGTTT | 3 (0.000057%) | | | GGTGTGAGTGTCGCCCATGGGCATCGACACCTTGCGGCTAGGAACTGGAACGAGACGGGTGGCAAAGATTTCGAGTAGCACTTCATACTACCGTGGGTTT | 9 (0.000170%) | | | GGTTCACATTTCGTTCATCACCCTTGGCCGGCTATCGAACAGCCGGACTCCCATCAAAAGATGGTTGCCAAGAACATCTTCGTTACGGTTTGCTAATTCT | 4 (0.000076%) | | | GGTTCACATTTCGTTCATCACCCTTGGCCGGCTTTCGAACAGCCGGACTCCCATCAAAAGATGGTTGCCAAGAACATCTTCGTTACGGTTTGCTAATTCT | 3 (0.000057%) | | | GGTTCGCCGCCCGCGACGTCGCGAGAAGTCCACTAAACCTTATCATTTAGAGGAAGGAGAAGTCGTAACAAGGTTTCCGTAGGTGAACCTGCGGAAGGAT | 14 (0.000265%) | | | GGTTCGTATTCGTACTGAAAATCAGAATCAAACGAGCTTTTACCCTTTTGTTCCACACGAGATTTCTGTTCTCGTTGAGCTCATCTTAGGACACCTGCGT | 9 (0.000170%) | | | GGTTGCGGTTTAAGTTCTTATACTCAATCATACACATGAC | 9684 (0.073282%) | | | GGTTGCGGTTTAAGTTGTTATACTCAATCATACACATGAC | 963 (0.007287%) | | | GGTTTAAGTTCTTATACTCAATCATACACATGACATCAAG | 1531 (0.011586%) | | | GGTTTAAGTTGTTATACTCAATCATACACATGACAACAAGTCATATTCGACTCCAAAACACTAACCAACCTTCTTCTTGCTTCTCAAAGCTTTCATGGTG | 112 (0.002119%) | | | GGTTTAAGTTGTTATACTCAATCATACACATGACATCAAG | 2183 (0.016519%) | | | GGTTTACTCACCCGTTGACTCGCACACATGTCAGACTCCTTGGTCCGTGTTTCAAGACGGGTCGAATGGGGAGCCCACAGGCCGACGCCCGGAGCACGCT | 17 (0.000322%) | | | GTAAATTTTCATAATTTTTTGACACCTCTAGCTAGGTCATTTGACCTGATACAACATCGGATTTTCATGGTCTAGTTGGGGCTCCGTGGGCATATTTGAT | 16 (0.000303%) | | | GTAACGGGTGACGGAGAATTAGGGTTCGATTCCGGAGAGGGAGCCTGAGAAACGGCTACCACATCCAAGGAAGGCAGCAGGCGCGCAAATTACCCAATCC | 18 (0.000341%) | | | GTAACTTCGGGAAAAGGATTGGCTCTGAGGGCTGGGCTCGGGGGTCCCAGTTCCGAACCCGTCGGCTGTCAGCGGACTGCTCGAGCTGCTTCCGCGGCGA | 12 (0.000227%) | | | GTAAGAATTGTATCCTTGTTAGAAGACACAAAGCCAAAGACTCATATGGACTTTGGCTACACCATGAAAGCTTTGAGAAGCAAGAAGAAGGTTGGTTAGT | 60 (0.001135%) | | | GTAAGTATGAACGAATTCAGACTGTGAAACTGCGAATGGCTCATTAAATCAGTTATAGTTTGTTTGATGGTAACTACTACTCGGATAACCGTAGTAATTC | 23 (0.000435%) | | | GTAATTCCAGCTCCAATAGCGTATATTTAAGTTGTTGCAGTTAAAAAGCTCGTAGTTGAACCTTGGGATGGGTCGGCCGGTCCGCCTTTGGTGTGCATTG | 13 (0.000246%) | | | GTACGCTCCAGGCGTCCTTGGCTCGGATTTAGGCCAACCGCGTGCGGTAACACACGGGAGACCAGCTTCCGTCCCGCATCAGCAAAGGATGGTGAGGGAC | 16 (0.000303%) | | | GTACTCATTCCAATTACCAGACTCGAAAGAGCCCGGTATTGTTATTTATTGTCACTACCTCCCCGTGTCAGGATTGGGTAATTTGCGCGCCTGCTGCCTT | 5 (0.000095%) | | | GTACTGAAAATCAGAATCAAACGAGCTTTTACCCTTTTGTTCCACACGAGATTTCTGTTCTCGTTGAGCTCATCTTAGGACACCTGCGTTATCTTTTAAC | 7 (0.000132%) | | | GTAGGAGCGACGGGCGGTGTGTACAAAGGGCAGGGACGTAGTCAACGCGAGCTGATGACTCGCGCTTACTAGGAATTCCTCGTTGAAGACCAACAATTGC | 118 (0.002232%) | | | GTAGGCTTGCTTTGAGCACTCTAATTTCTTCAAAGTAACAGCGCCGGAGGCACGACCCGGCCAATTAAGACCAGGAGCGTATCGCCGACCGAAGGGACAA | 26 (0.000492%) | | | GTAGTCATATGCTTGTCTCAAAGATTAAGCCATGCATGTGTAAGTATGAACGAATTCAGACTGTGAAACTGCGAATGGCTCATTAAATCAGTTATAGTTT | 102 (0.001930%) | | | GTAGTGTTTCCTTGTTAGAAGACACAAAGCCAAAGACTCATATGGACTTTGGCTACACCATGAAAGCTTTGAGAAGCAAGAAGAAGGTTGGTTAGTGTTT | 15 (0.000284%) | | | GTATCCAGAGCGTAGGCTTGCTTTGAGCACTCTAATTTCTTCAAAGTAACAGCGCCGGAGGCACGACCCGGCCAATTAAGACCAGGAGCGTATCGCCGAC | 17 (0.000322%) | | | GTATCCTTGTTAGAAGACACAAAGCCAAAGACTCATATGGACTTTGGCTACACCATGAAAGCTTTGAGAAGCAAGAAGAAGGTTGGTTAGTGTTTTGGAG | 23 (0.000435%) | | | GTATGAACGAATTCAGACTGTGAAACTGCGAATGGCTCATTAAATCAGTTATAGTTTGTTTGATGGTAACTACTACTCGGATAACCGTAGTAATTCTAGA | 12 (0.000227%) | | | GTATTTCATAGTCAGAGGTGAAATTCTTGGATTTATGAAAGACGAACAACTGCGAAAGCATTTGCCAAGGATGTTTTCATTAATCAAGAACGAAAGTTGG | 6 (0.000114%) | | | GTCAGAGGTGAAATTCTTGGATTTATGAAAGACGAACAACTGCGAAAGCATTTGCCAAGGATGTTTTCATTAATCAAGAACGAAAGTTGGGGGCTCGAAG | 20 (0.000378%) | | | GTCAGGTGGGGAGTTTGGCTGGGGCGGCACATCTGTTAAAAGATAACGCAGGTGTCCTAAGATGAGCTCAACGAGAACAGAAATCTCGTGTGGAACAAAA | 8 (0.000151%) | | | GTCATATGCTTGTCTCAAAGATTAAGCCATGCATGTGTAAGTATGAACGAATTCAGACTGTGAAACTGCGAATGGCTCATTAAATCAGTTATAGTTTGTT | 72 (0.001362%) | | | GTCATATTCGACTCCAAAACACTAACCAACCTTCTTCTTGCTTCTCAAAGCTTTCATGGTGTAGCCAAAGTCCATATGAGTCTTTGGCTTTGTGTCTTCT | 245 (0.004635%) | | | GTCCATATGAGTCTTTGGCTTTGTGTCTTCTAACAAGGAT | 7701 (0.058276%) | | | GTCCCGAGTGTGAGCGAGGTGTGAGTGTCGCCCATGGGCATCGACACCTTGCGGCTAGGAACTGGAACGAGACGGGTAGCAAAGATTTCGAGTAGCACTT | 15 (0.000284%) | | | GTCCCGAGTGTGAGCGAGGTGTGAGTGTCGCCCATGGGCATCGACACCTTGCGGCTAGGAACTGGAACGAGACGGGTGGCAAAGATTTCGAGTAGCACTT | 20 (0.000378%) | | | GTCCCTCACCATCCTTTGCTGATGCGGGACGGAAGCTGGTCTCCCGTGTGTTACCGCACGCGGTTGGCCTAAATCCGAGCCAAGGACGCCTGGAGCGTAC | 12 (0.000227%) | | | GTCCGACTTTGTGAAATGACTTGAGAGGTGTAGGATAAGTGGGAGCTTCGGCGCAAGTGAAATACCACTACTTTTAACGTTATTTTACTTACTCCGTGAA | 19 (0.000359%) | | | GTCGAAATCCGCTAAGGAGTGTGTAACAACTCACCTGCCGAATCAACTAGCCCCGAAAATGGATGGCGCTTAAGCGCGCGACCTATACCCGGCCGTCGGG | 75 (0.001419%) | | | GTCGACCTTTTATCTAATAAATGCGTCCCTTCCATAAGTCGGGGTTTGTTGCACGTATTAGCTCTAGAATTACTACGGTTATCCGAGTAGTAGTTACCAT | 25 (0.000473%) | | | GTCGAGTTATCATGAATCATCAGAGCAACGGGCAGAGCCCGCGTCGACCTTTTATCTAATAAATGCGTCCCTTCCATAAGTCGGGGTTTGTTGCACGTAT | 463 (0.008759%) | | | GTCGCCCATGGGCATCGACACCTTGCGGCTAGGAACTGGAACGAGACGGGTGGCAAAGATTTCGAGTAGCACTTCATACTACCGTGGGTTTTTTAAACCT | 30 (0.000568%) | | | GTCGGCCTGTGGGCTCCCCATTCGACCCGTCTTGAAACACGGACCAAGGAGTCTGACATGTGTGCGAGTCAACGGGTGAGTAAACCCGTAAGGCGCAAGG | 25 (0.000473%) | | | GTCGGGGGCATTCGTATTTCATAGTCAGAGGTGAAATTCTTGGATTTATGAAAGACGAACAACTGCGAAAGCATTTGCCAAGGATGTTTTCATTAATCAA | 8 (0.000151%) | | | GTCGGTACGCTCCAGGCGTCCTTGGCTCGGATTTAGGCCAACCGCGTGCGGTAACACACGGGAGACCAGCTTCCGTCCCGCATCAGCAAAGGATGGTGAG | 10 (0.000189%) | | | GTCGTCCCTCACCATCCTTTGCTGATGCGGGACGGAAGCTGGTCTCCCGTGTGTTACCGCACGCGGTTGGCCTAAATCCGAGCCAAGGACGCCTGGAGCG | 721 (0.013640%) | | | GTCTCAAAGATTAAGCCATGCATGTGTAAGTATGAACGAATTCAGACTGTGAAACTGCGAATGGCTCATTAAATCAGTTATAGTTTGTTTGATGGTAACT | 3 (0.000057%) | | | GTCTCCGGGGAGGCGAATGCCAGCCGTTCGTTTGCATGTTCCTTGACACTTTTCGTGCCGGGGTTTTGTGATATCCGGAAGCAACGCGCACGACAAGACC | 9 (0.000170%) | | | GTCTGAACTCCAGTCACGAGATTCCATCTCGTATGCCGTC | 10 (0.000076%) | | | GTCTGGTGCCAGCAGCCGCGGTAATTCCAGCTCCAATAGCGTATATTTAAGTTGTTGCAGTTAAAAAGCTCGTAGTTGAACCTTGGGATGGGTCGGCCGG | 28 (0.000530%) | | | GTCTGTGATGCCCTTAGATGTTCTGGGCCGCACGCGCGCTACACTGATGTATTCAACGAGTTCACACCTTGGCCGACAGGCCCGGGTAATCTTTGAAATT | 19 (0.000359%) | | | GTCTTCAACGAGGAATTCCTAGTAAGCGCGAGTCATCAGCTCGCGTTGACTACGTCCCTGCCCTTTGTACACACCGCCCGTCGCTCCTACCGATTGAATG | 15 (0.000284%) | | | GTGACGGAGAATTAGGGTTCGATTCCGGAGAGGGAGCCTGAGAAACGGCTACCACATCCAAGGAAGGCAGCAGGCGCGCAAATTACCCAATCCTGACACG | 10 (0.000189%) | | | GTGAGCGAGGTGTGAGTGTCGCCCATGGGCATCGACACCTTGCGGCTAGGAACTGGAACGAGACGGGTAGCAAAGATTTCGAGTAGCACTTCATACTACC | 13 (0.000246%) | | | GTGAGCGAGGTGTGAGTGTCGCCCATGGGCATCGACACCTTGCGGCTAGGAACTGGAACGAGACGGGTGGCAAAGATTTCGAGTAGCACTTCATACTACC | 20 (0.000378%) | | | GTGAGTGTCGCCCATGGGCATCGACACCTTGCGGCTAGGAACTGGAACGAGACGGGTAGCAAAGATTTCGAGTAGCACTTCATACTACCGTGGGTTTTTT | 25 (0.000473%) | | | GTGAGTGTCGCCCATGGGCATCGACACCTTGCGGCTAGGAACTGGAACGAGACGGGTGGCAAAGATTTCGAGTAGCACTTCATACTACCGTGGGTTTTTT | 38 (0.000719%) | | | GTGATGCCCTTAGATGTTCTGGGCCGCACGCGCGCTACACTGATGTATTCAACGAGTTCACACCTTGGCCGACAGGCCCGGGTAATCTTTGAAATTTCAT | 19 (0.000359%) | | | GTGCCAGCAGCCGCGGTAATTCCAGCTCCAATAGCGTATATTTAAGTTGTTGCAGTTAAAAAGCTCGTAGTTGAACCTTGGGATGGGTCGGCCGGTCCGC | 24 (0.000454%) | | | GTGCCTCGGCATCAGCGTGCTCCGGGCGTCGGCCTGTGGGCTCCCCATTCGACCCGTCTTGAAACACGGACCAAGGAGTCTGACATGTGTGCGAGTCAAC | 10 (0.000189%) | | | GTGCTCCGGGCGTCGGCCTGTGGGCTCCCCATTCGACCCGTCTTGAAACACGGACCAAGGAGTCTGACATGTGTGCGAGTCAACGGGTGAGTAAACCCGT | 12 (0.000227%) | | | GTGGCCTACCATGGTGGTAACGGGTGACGGAGAATTAGGGTTCGATTCCGGAGAGGGAGCCTGAGAAACGGCTACCACATCCAAGGAAGGCAGCAGGCGC | 18 (0.000341%) | | | GTGGGGAGTTTGGCTGGGGCGGCACATCTGTTAAAAGATAACGCAGGTGTCCTAAGATGAGCTCAACGAGAACAGAAATCTCGTGTGGAACAAAAGGGTA | 4 (0.000076%) | | | GTGGGTGGTTCGCCGCCCGCGACGTCGCGAGAAGTCCACTAAACCTTATCATTTAGAGGAAGGAGAAGTCGTAACAAGGTTTCCGTAGGTGAACCTGCGG | 67 (0.001268%) | | | GTGGTTCGCCGCCCGCGACGTCGCGAGAAGTCCACTAAACCTTATCATTTAGAGGAAGGAGAAGTCGTAACAAGGTTTCCGTAGGTGAACCTGCGGAAGG | 23 (0.000435%) | | | GTGTAAGTATGAACGAATTCAGACTGTGAAACTGCGAATGGCTCATTAAATCAGTTATAGTTTGTTTGATGGTAACTACTACTCGGATAACCGTAGTAAT | 12 (0.000227%) | | | GTGTATCCTTGTTAGAAGACACAAAGCCAAAGACTCATATGGACTTTGGCTACACCATGAAAGCTTTGAGAAGCAAGAAGAAGGTTGGTTAGTGTTTTGG | 7 (0.000132%) | | | GTGTCGCCCATGGGCATCGACACCTTGCGGCTAGGAACTGGAACGAGACGGGTGGCAAAGATTTCGAGTAGCACTTCATACTACCGTGGGTTTTTTAAAC | 10 (0.000189%) | | | GTGTGAGCGAGGTGTGAGTGTCGCCCATGGGCATCGACACCTTGCGGCTAGGAACTGGAACGAGACGGGTAGCAAAGATTTCGAGTAGCACTTCATACTA | 18 (0.000341%) | | | GTGTGAGCGAGGTGTGAGTGTCGCCCATGGGCATCGACACCTTGCGGCTAGGAACTGGAACGAGACGGGTGGCAAAGATTTCGAGTAGCACTTCATACTA | 34 (0.000643%) | | | GTGTGAGTGTCGCCCATGGGCATCGACACCTTGCGGCTAGGAACTGGAACGAGACGGGTAGCAAAGATTTCGAGTAGCACTTCATACTACCGTGGGTTTT | 15 (0.000284%) | | | GTGTGAGTGTCGCCCATGGGCATCGACACCTTGCGGCTAGGAACTGGAACGAGACGGGTGGCAAAGATTTCGAGTAGCACTTCATACTACCGTGGGTTTT | 20 (0.000378%) | | | GTGTTTCCTTGTTAGAAGACACAAAGCCAAAGACTCATATGGACTTTGGCTACACCATGAAAGCTTTGAGAAGCAAGAAGAAGGTTGGTTAGTGTTTTGG | 44 (0.000832%) | | | GTTAAAAGATAACGCAGGTGTCCTAAGATGAGCTCAACGAGAACAGAAATCTCGTGTGGAACAAAAGGGTAAAAGCTCGTTTGATTCTGATTTTCAGTAC | 23 (0.000435%) | | | GTTAAGGGATTTAGATTGTACTCATTCCAATTACCAGACTCGAAAGAGCCCGGTATTGTTATTTATTGTCACTACCTCCCCGTGTCAGGATTGGGTAATT | 85 (0.001608%) | | | GTTAGAAGACACAAAGCCAAAGACTCATATGGACTTTGGCTACACCATGAAAGCTTTGAGAAGCAAGAAGAAGGTTGGTTAGTGTTTTGGAGTCGAATAT | 84 (0.001589%) | | | GTTAGTTTCTTTTCCTCCGCTTATTGATATGCTTAAACTCAGCGGGTAATCCCGCCTGACCTGGGGTCGCTATATGGACTTTGGGTCATCTACAGCTTCC | 25 (0.000473%) | | | GTTATACTCAATCATACACATGACAACAAGTCATATTCGACTCCAAAACACTAACCAACCTTCTTCTTGCTTCTCAAAGCTTTCATGGTGTAGCCAAAGT | 126 (0.002384%) | | | GTTATAGTTTGTTTGATGGTAACTACTACTCGGATAACCGTAGTAATTCTAGAGCTAATACGTGCAACAAACCCCGACTTATGGAAGGGACGCATTTATT | 18 (0.000341%) | | | GTTATCATGAATCATCAGAGCAACGGGCAGAGCCCGCGTCGACCTTTTATCTAATAAATGCGTCCCTTCCATAAGTCGGGGTTTGTTGCACGTATTAGCT | 277 (0.005240%) | | | GTTATCCCATGCTAATGTATCCAGAGCGTAGGCTTGCTTTGAGCACTCTAATTTCTTCAAAGTAACAGCGCCGGAGGCACGACCCGGCCAATTAAGACCA | 58 (0.001097%) | | | GTTCACATTTCGTTCATCACCCTTGGCCGGCTATCGAACAGCCGGACTCCCATCAAAAGATGGTTGCCAAGAACATCTTCGTTACGGTTTGCTAATTCTC | 9 (0.000170%) | | | GTTCACATTTCGTTCATCACCCTTGGCCGGCTTTCGAACAGCCGGACTCCCATCAAAAGATGGTTGCCAAGAACATCTTCGTTACGGTTTGCTAATTCTC | 14 (0.000265%) | | | GTTCACCACCGCATGTCGGTACGCTCCAGGCGTCCTTGGCTCGGATTTAGGCCAACCGCGTGCGGTAACACACGGGAGACCAGCTTCCGTCCCGCATCAG | 110 (0.002081%) | | | GTTCCATCGACCAGAGGCTGTTCACCTTGGAGACCTGATGCGGTTATGAGTACGACCGGGCGTGAGCGGCACTCGGTCCTCCGGATTTTCAAGGGCCGCC | 41 (0.000776%) | | | GTTCGCCGCCCGCGACGTCGCGAGAAGTCCACTAAACCTTATCATTTAGAGGAAGGAGAAGTCGTAACAAGGTTTCCGTAGGTGAACCTGCGGAAGGATC | 35 (0.000662%) | | | GTTCGCTCGCCGTTACTAAGGGAATCCTTGTTAGTTTCTTTTCCTCCGCTTATTGATATGCTTAAACTCAGCGGGTAATCCCGCCTGACCTGGGGTCGCT | 1065 (0.020148%) | | | GTTCGTATTCGTACTGAAAATCAGAATCAAACGAGCTTTTACCCTTTTGTTCCACACGAGATTTCTGTTCTCGTTGAGCTCATCTTAGGACACCTGCGTT | 14 (0.000265%) | | | GTTCTGGGCCGCACGCGCGCTACACTGATGTATTCAACGAGTTCACACCTTGGCCGACAGGCCCGGGTAATCTTTGAAATTTCATCGTGATGGGGATAGA | 49 (0.000927%) | | | GTTCTTATACTCAATCATACACATGACATCAAGTCATATTCGACTCCAAAACACTAACCAACCTTCTTCTTGCTTCTCAAAGCTTTCATGGTGTAGCCAA | 91 (0.001722%) | | | GTTGAGCTTGACTCTAGTCCGACTTTGTGAAATGACTTGAGAGGTGTAGGATAAGTGGGAGCTTCGGCGCAAGTGAAATACCACTACTTTTAACGTTATT | 36 (0.000681%) | | | GTTGCGGTTTAAGTTCTTATACTCAATCATACACATGACA | 2325 (0.017594%) | | | GTTGCGGTTTAAGTTGTTATACTCAATCATACACATGACA | 2552 (0.019312%) | | | GTTGGTCTTCAACGAGGAATTCCTAGTAAGCGCGAGTCATCAGCTCGCGTTGACTACGTCCCTGCCCTTTGTACACACCGCCCGTCGCTCCTACCGATTG | 198 (0.003746%) | | | GTTGTTATACTCAATCATACACATGACATCAAGTCATATT | 255 (0.001930%) | | | GTTTAAGTTCTTATACTCAATCATACACATGACATCAAGT | 2182 (0.016512%) | | | GTTTAAGTTGTTATACTCAATCATACACATGACAACAAGTCATATTCGACTCCAAAACACTAACCAACCTTCTTCTTGCTTCTCAAAGCTTTCATGGTGT | 133 (0.002516%) | | | GTTTAAGTTGTTATACTCAATCATACACATGACATCAAGT | 1021 (0.007726%) | | | GTTTACTCACCCGTTGACTCGCACACATGTCAGACTCCTTGGTCCGTGTTTCAAGACGGGTCGAATGGGGAGCCCACAGGCCGACGCCCGGAGCACGCTG | 28 (0.000530%) | | | GTTTCCTTGTTAGAAGACACAAAGCCAAAGACTCATATGGACTTTGGCTACACCATGAAAGCTTTGAGAAGCAAGAAGAAGGTTGGTTAGTGTTTTGGAG | 41 (0.000776%) | | | GTTTCTTTTCCTCCGCTTATTGATATGCTTAAACTCAGCGGGTAATCCCGCCTGACCTGGGGTCGCTATATGGACTTTGGGTCATCTACAGCTTCCGGAC | 21 (0.000397%) | | | GTTTGATGGTAACTACTACTCGGATAACCGTAGTAATTCTAGAGCTAATACGTGCAACAAACCCCGACTTATGGAAGGGACGCATTTATTAGATAAAAGG | 27 (0.000511%) | | | GTTTGGCTGGGGCGGCACATCTGTTAAAAGATAACGCAGGTGTCCTAAGATGAGCTCAACGAGAACAGAAATCTCGTGTGGAACAAAAGGGTAAAAGCTC | 45 (0.000851%) | | | GTTTGTTTGATGGTAACTACTACTCGGATAACCGTAGTAATTCTAGAGCTAATACGTGCAACAAACCCCGACTTATGGAAGGGACGCATTTATTAGATAA | 46 (0.000870%) | | | TAAAAGATAACGCAGGTGTCCTAAGATGAGCTCAACGAGAACAGAAATCTCGTGTGGAACAAAAGGGTAAAAGCTCGTTTGATTCTGATTTTCAGTACGA | 17 (0.000322%) | | | TAAAAGCCTAAGTAGTGTTTCCTTGTTAGAAGACACAAAGCCAAAGACTCATATGGACTTTGGCTACACCATGAAAGCTTTGAGAAGCAAGAAGAAGGTT | 33 (0.000624%) | | | TAAAGGCGTAAGAATTGTATCCTTGTTAAAAGACACAAAGCCAAAGACTCATATGGACTTTGGCTACACCATGAAAGCTTTGAGAAGCAAGAAGAAGGTT | 75 (0.001419%) | | | TAAAGGCGTAAGAATTGTATCCTTGTTAGAAGACACAAAGCCAAAGACTCATATGGACTTTGGCTACACCATGAAAGCTTTGAGAAGCAAGAAGAAGGTT | 159 (0.003008%) | | | TAAATCAGTTATAGTTTGTTTGATGGTAACTACTACTCGGATAACCGTAGTAATTCTAGAGCTAATACGTGCAACAAACCCCGACTTATGGAAGGGACGC | 9 (0.000170%) | | | TAAATCCTATTTTCTGGTAAATTTTCATAATTTTTTGACACCTCTAGCTAGGTCATTTGACCTGATACAACATCGGATTTTCATGGTCTAGTTGGGGCTC | 5 (0.000095%) | | | TAAATTTTCATAATTTTTTGACACCTCTAGCTAGGTCATTTGACCTGATACAACATCGGATTTTCATGGTCTAGTTGGGGCTCCGTGGGCATATTTGATG | 6 (0.000114%) | | | TAACAGGGACAGTCGGGGGCATTCGTATTTCATAGTCAGAGGTGAAATTCTTGGATTTATGAAAGACGAACAACTGCGAAAGCATTTGCCAAGGATGTTT | 47 (0.000889%) | | | TAACAGGTCTGTGATGCCCTTAGATGTTCTGGGCCGCACGCGCGCTACACTGATGTATTCAACGAGTTCACACCTTGGCCGACAGGCCCGGGTAATCTTT | 50 (0.000946%) | | | TAACGCCTCGAAGAACTAATGGCAGCCCACGCAAGGCAAGCCCATTCTCCTCGACGATTCAGCAGTTTTTGTCCGAGAACTGCTGAGAAAACTCGGAAAA | 6 (0.000114%) | | | TAACGGCGTGCCTCGGCATCAGCGTGCTCCGGGCGTCGGCCTGTGGGCTCCCCATTCGACCCGTCTTGAAACACGGACCAAGGAGTCTGACATGTGTGCG | 39 (0.000738%) | | | TAACGGGTGACGGAGAATTAGGGTTCGATTCCGGAGAGGGAGCCTGAGAAACGGCTACCACATCCAAGGAAGGCAGCAGGCGCGCAAATTACCCAATCCT | 5 (0.000095%) | | | TAAGAATTGTATCCTTGTTAGAAGACACAAAGCCAAAGACTCATATGGACTTTGGCTACACCATGAAAGCTTTGAGAAGCAAGAAGAAGGTTGGTTAGTG | 84 (0.001589%) | | | TAAGCCATGCATGTGTAAGTATGAACGAATTCAGACTGTGAAACTGCGAATGGCTCATTAAATCAGTTATAGTTTGTTTGATGGTAACTACTACTCGGAT | 8 (0.000151%) | | | TAAGCGCGCGACCTATACCCGGCCGTCGGGGCAAGAGCCAGGCCTCGATGAGTAGGAGGGCGCGGCGGTCGCTGCAAAACCTAGGGCGCGAGCCCGGGCG | 7 (0.000132%) | | | TAAGGATACTAAATCCTATTTTCTGGTAAATTTTCATAATTTTTTGACACCTCTAGCTAGGTCATTTGACCTGATACAACATCGGATTTTCATGGTCTAG | 5 (0.000095%) | | | TAAGGGATTTAGATTGTACTCATTCCAATTACCAGACTCGAAAGAGCCCGGTATTGTTATTTATTGTCACTACCTCCCCGTGTCAGGATTGGGTAATTTG | 43 (0.000813%) | | | TAAGTAGTGTTTCCTTGTTAGAAGACACAAAGCCAAAGACTCATATGGACTTTGGCTACACCATGAAAGCTTTGAGAAGCAAGAAGAAGGTTGGTTAGTG | 43 (0.000813%) | | | TAAGTATGAACGAATTCAGACTGTGAAACTGCGAATGGCTCATTAAATCAGTTATAGTTTGTTTGATGGTAACTACTACTCGGATAACCGTAGTAATTCT | 9 (0.000170%) | | | TAAGTTCTTATACTCAATCATACACATGACATCAAGTCATATTCGACTCCAAAACACTAACCAACCTTCTTCTTGCTTCTCAAAGCTTTCATGGTGTAGC | 22 (0.000416%) | | | TAAGTTGTTATACTCAATCATACACATGACATCAAGTCAT | 81 (0.000613%) | | | TAATGATTAACAGGGACAGTCGGGGGCATTCGTATTTCATAGTCAGAGGTGAAATTCTTGGATTTATGAAAGACGAACAACTGCGAAAGCATTTGCCAAG | 1489 (0.028169%) | | | TAATGTATCCAGAGCGTAGGCTTGCTTTGAGCACTCTAATTTCTTCAAAGTAACAGCGCCGGAGGCACGACCCGGCCAATTAAGACCAGGAGCGTATCGC | 5 (0.000095%) | | | TAATTCCAGCTCCAATAGCGTATATTTAAGTTGTTGCAGTTAAAAAGCTCGTAGTTGAACCTTGGGATGGGTCGGCCGGTCCGCCTTTGGTGTGCATTGG | 7 (0.000132%) | | | TAATTTTTTGACACCTCTAGCTAGGTCATTTGACCTGATACAACATCGGATTTTCATGGTCTAGTTGGGGCTCCGTGGGCATATTTGATGCAAACTTGAC | 9 (0.000170%) | | | TACACATGACATCAAGTCATATTCGACTCCAAAACACTAACCAACCTTCTTCTTGCTTCTCAAAGCTTTCATGGTGTAGCCAAAGTCCATATGAGTCTTT | 18 (0.000341%) | | | TACACCATGAAAGCTTTGAGAAGCAAGAAGAAGGTTGGTTAGTGTTTTGGAGTCGAATATGACTTGATGTCATGTGTATGATTGAGTATAACAACTTAAA | 135 (0.002554%) | | | TACCAGACTCGAAAGAGCCCGGTATTGTTATTTATTGTCACTACCTCCCCGTGTCAGGATTGGGTAATTTGCGCGCCTGCTGCCTTCCTTGGATGTGGTA | 6 (0.000114%) | | | TACCCGGCCGTCGGGGCAAGAGCCAGGCCTCGATGAGTAGGAGGGCGCGGCGGTCGCTGCAAAACCTAGGGCGCGAGCCCGGGCGGAGCGGCCGTCGGTG | 5 (0.000095%) | | | TACGGGTTTACTCACCCGTTGACTCGCACACATGTCAGACTCCTTGGTCCGTGTTTCAAGACGGGTCGAATGGGGAGCCCACAGGCCGACGCCCGGAGCA | 4 (0.000076%) | | | TACTCAATCATACACATGACAACAAGTCATATTCGACTCCAAAACACTAACCAACCTTCTTCTTGCTTCTCAAAGCTTTCATGGTGTAGCCAAAGTCCAT | 20 (0.000378%) | | | TACTCAATCATACACATGACATCAAGTCATATTCGACTCCAAAACACTAACCAACCTTCTTCTTGCTTCTCAAAGCTTTCATGGTGTAGCCAAAGTCCAT | 13 (0.000246%) | | | TAGAAGACACAAAGCCAAAGACTCATATGGACTTTGGCTACACCATGAAAGCTTTGAGAAGCAAGAAGAAGGTTGGTTAGTGTTTTGGAGTCGAATATGA | 44 (0.000832%) | | | TAGATGTTCTGGGCCGCACGCGCGCTACACTGATGTATTCAACGAGTTCACACCTTGGCCGACAGGCCCGGGTAATCTTTGAAATTTCATCGTGATGGGG | 18 (0.000341%) | | | TAGATTGTACTCATTCCAATTACCAGACTCGAAAGAGCCCGGTATTGTTATTTATTGTCACTACCTCCCCGTGTCAGGATTGGGTAATTTGCGCGCCTGC | 5 (0.000095%) | | | TAGGAGCGACGGGCGGTGTGTACAAAGGGCAGGGACGTAGTCAACGCGAGCTGATGACTCGCGCTTACTAGGAATTCCTCGTTGAAGACCAACAATTGCA | 12 (0.000227%) | | | TAGGCCACGCTTTCACGGTTCGTATTCGTACTGAAAATCAGAATCAAACGAGCTTTTACCCTTTTGTTCCACACGAGATTTCTGTTCTCGTTGAGCTCAT | 78 (0.001476%) | | | TAGGCGAGACAAGGGTTCACATTTCGTTCATCACCCTTGGCCGGCTATCGAACAGCCGGACTCCCATCAAAAGATGGTTGCCAAGAACATCTTCGTTACG | 5 (0.000095%) | | | TAGGCGAGACAAGGGTTCACATTTCGTTCATCACCCTTGGCCGGCTTTCGAACAGCCGGACTCCCATCAAAAGATGGTTGCCAAGAACATCTTCGTTACG | 7 (0.000132%) | | | TAGGCTGTCCCGAGTGTGAGCGAGGTGTGAGTGTCGCCCATGGGCATCGACACCTTGCGGCTAGGAACTGGAACGAGACGGGTAGCAAAGATTTCGAGTA | 18 (0.000341%) | | | TAGGCTGTCCCGAGTGTGAGCGAGGTGTGAGTGTCGCCCATGGGCATCGACACCTTGCGGCTAGGAACTGGAACGAGACGGGTGGCAAAGATTTCGAGTA | 24 (0.000454%) | | | TAGGCTTGCTTTGAGCACTCTAATTTCTTCAAAGTAACAGCGCCGGAGGCACGACCCGGCCAATTAAGACCAGGAGCGTATCGCCGACCGAAGGGACAAG | 3 (0.000057%) | | | TAGTCATATGCTTGTCTCAAAGATTAAGCCATGCATGTGTAAGTATGAACGAATTCAGACTGTGAAACTGCGAATGGCTCATTAAATCAGTTATAGTTTG | 56 (0.001059%) | | | TAGTGGCCTACCATGGTGGTAACGGGTGACGGAGAATTAGGGTTCGATTCCGGAGAGGGAGCCTGAGAAACGGCTACCACATCCAAGGAAGGCAGCAGGC | 9 (0.000170%) | | | TAGTGTTTCCTTGTTAGAAGACACAAAGCCAAAGACTCATATGGACTTTGGCTACACCATGAAAGCTTTGAGAAGCAAGAAGAAGGTTGGTTAGTGTTTT | 8 (0.000151%) | | | TAGTTTCTTTTCCTCCGCTTATTGATATGCTTAAACTCAGCGGGTAATCCCGCCTGACCTGGGGTCGCTATATGGACTTTGGGTCATCTACAGCTTCCGG | 9 (0.000170%) | | | TATACCCGGCCGTCGGGGCAAGAGCCAGGCCTCGATGAGTAGGAGGGCGCGGCGGTCGCTGCAAAACCTAGGGCGCGAGCCCGGGCGGAGCGGCCGTCGG | 7 (0.000132%) | | | TATACTCAATCATACACATGACAACAAGTCATATTCGACTCCAAAACACTAACCAACCTTCTTCTTGCTTCTCAAAGCTTTCATGGTGTAGCCAAAGTCC | 13 (0.000246%) | | | TATACTCAATCATACACATGACATCAAGTCATATTCGACTCCAAAACACTAACCAACCTTCTTCTTGCTTCTCAAAGCTTTCATGGTGTAGCCAAAGTCC | 45 (0.000851%) | | | TATAGTTTGTTTGATGGTAACTACTACTCGGATAACCGTAGTAATTCTAGAGCTAATACGTGCAACAAACCCCGACTTATGGAAGGGACGCATTTATTAG | 9 (0.000170%) | | | TATCATGAATCATCAGAGCAACGGGCAGAGCCCGCGTCGACCTTTTATCTAATAAATGCGTCCCTTCCATAAGTCGGGGTTTGTTGCACGTATTAGCTCT | 80 (0.001513%) | | | TATCCAGAGCGTAGGCTTGCTTTGAGCACTCTAATTTCTTCAAAGTAACAGCGCCGGAGGCACGACCCGGCCAATTAAGACCAGGAGCGTATCGCCGACC | 4 (0.000076%) | | | TATCCCATGCTAATGTATCCAGAGCGTAGGCTTGCTTTGAGCACTCTAATTTCTTCAAAGTAACAGCGCCGGAGGCACGACCCGGCCAATTAAGACCAGG | 8 (0.000151%) | | | TATCCTTGTTAGAAGACACAAAGCCAAAGACTCATATGGACTTTGGCTACACCATGAAAGCTTTGAGAAGCAAGAAGAAGGTTGGTTAGTGTTTTGGAGT | 42 (0.000795%) | | | TATCTAATAAATGCGTCCCTTCCATAAGTCGGGGTTTGTTGCACGTATTAGCTCTAGAATTACTACGGTTATCCGAGTAGTAGTTACCATCAAACAAACT | 4 (0.000076%) | | | TATGACTTGATGTCATGTGTATGATTGAGTATAAGAACTT | 322 (0.002437%) | | | TATGATGTTATCCCATGCTAATGTATCCAGAGCGTAGGCTTGCTTTGAGCACTCTAATTTCTTCAAAGTAACAGCGCCGGAGGCACGACCCGGCCAATTA | 35 (0.000662%) | | | TATGCTTGTCTCAAAGATTAAGCCATGCATGTGTAAGTATGAACGAATTCAGACTGTGAAACTGCGAATGGCTCATTAAATCAGTTATAGTTTGTTTGAT | 17 (0.000322%) | | | TATGGACTTTGGCTACACCATGAAAGCTTTGAGAAGCAAGAAGAAGGTTGGTTAGTGTTTTGGAGTCGAATATGACTTGATGTCATGTGTATGATTGAGT | 16 (0.000303%) | | | TATTCGACTCCAAAACACTAACCAACCTTCTTCTTGCTTCTCAAAGCTTTCATGGTGTAGCCAAAGTCCATATGAGTCTTTGGCTTTGTGTCTTCTAACA | 45 (0.000851%) | | | TATTCGTACTGAAAATCAGAATCAAACGAGCTTTTACCCTTTTGTTCCACACGAGATTTCTGTTCTCGTTGAGCTCATCTTAGGACACCTGCGTTATCTT | 3 (0.000057%) | | | TCAAAGATTAAGCCATGCATGTGTAAGTATGAACGAATTCAGACTGTGAAACTGCGAATGGCTCATTAAATCAGTTATAGTTTGTTTGATGGTAACTACT | 5 (0.000095%) | | | TCAAAGCTTTCATGGTGTAGCCAAAGTCCATATGAGTCTTTGGCTTTGTGTCTTCTAACAAGGAAACACTACTTAGGCTTATAAGATGCGGTTGCGGTTT | 16 (0.000303%) | | | TCAACGAGGAATTCCTAGTAAGCGCGAGTCATCAGCTCGCGTTGACTACGTCCCTGCCCTTTGTACACACCGCCCGTCGCTCCTACCGATTGAATGATCG | 3 (0.000057%) | | | TCAAGTCATATTCGACTCCAAAACACTAACCAACCTTCTTCTTGCTTCTCAAAGCTTTCATGGTGTAGCCAAAGTCCATATGAGTCTTTGGCTTTGTGTC | 55 (0.001041%) | | | TCAATCATACACATGACAACAAGTCATATTCGACTCCAAAACACTAACCAACCTTCTTCTTGCTTCTCAAAGCTTTCATGGTGTAGCCAAAGTCCATATG | 14 (0.000265%) | | | TCAATCATACACATGACATCAAGTCATATTCGACTCCAAAACACTAACCAACCTTCTTCTTGCTTCTCAAAGCTTTCATGGTGTAGCCAAAGTCCATATG | 34 (0.000643%) | | | TCAATCGGTAGGAGCGACGGGCGGTGTGTACAAAGGGCAGGGACGTAGTCAACGCGAGCTGATGACTCGCGCTTACTAGGAATTCCTCGTTGAAGACCAA | 49 (0.000927%) | | | TCACATTTCGTTCATCACCCTTGGCCGGCTATCGAACAGCCGGACTCCCATCAAAAGATGGTTGCCAAGAACATCTTCGTTACGGTTTGCTAATTCTCGG | 11 (0.000208%) | | | TCACATTTCGTTCATCACCCTTGGCCGGCTTTCGAACAGCCGGACTCCCATCAAAAGATGGTTGCCAAGAACATCTTCGTTACGGTTTGCTAATTCTCGG | 3 (0.000057%) | | | TCACCACCGCATGTCGGTACGCTCCAGGCGTCCTTGGCTCGGATTTAGGCCAACCGCGTGCGGTAACACACGGGAGACCAGCTTCCGTCCCGCATCAGCA | 15 (0.000284%) | | | TCACCATCCTTTGCTGATGCGGGACGGAAGCTGGTCTCCCGTGTGTTACCGCACGCGGTTGGCCTAAATCCGAGCCAAGGACGCCTGGAGCGTACCGACA | 8 (0.000151%) | | | TCAGAATCAAACGAGCTTTTACCCTTTTGTTCCACACGAGATTTCTGTTCTCGTTGAGCTCATCTTAGGACACCTGCGTTATCTTTTAACAGATGTGCCG | 12 (0.000227%) | | | TCAGAGGTGAAATTCTTGGATTTATGAAAGACGAACAACTGCGAAAGCATTTGCCAAGGATGTTTTCATTAATCAAGAACGAAAGTTGGGGGCTCGAAGA | 4 (0.000076%) | | | TCATAATTTTTTGACACCTCTAGCTAGGTCATTTGACCTGATACAACATCGGATTTTCATGGTCTAGTTGGGGCTCCGTGGGCATATTTGATGCAAACTT | 3 (0.000057%) | | | TCATACACATGACATCAAGTCATATTCGACTCCAAAACACTAACCAACCTTCTTCTTGCTTCTCAAAGCTTTCATGGTGTAGCCAAAGTCCATATGAGTC | 6 (0.000114%) | | | TCATAGTCAGAGGTGAAATTCTTGGATTTATGAAAGACGAACAACTGCGAAAGCATTTGCCAAGGATGTTTTCATTAATCAAGAACGAAAGTTGGGGGCT | 12 (0.000227%) | | | TCATATGGACTTTGGCTACACCATGAAAGCTTTGAGAAGCAAGAAGAAGGTTGGTTAGTGTTTTGGAGTCGAATATGACTTGATGTCATGTGTATGATTG | 33 (0.000624%) | | | TCATATTCGACTCCAAAACACTAACCAACCTTCTTCTTGCTTCTCAAAGCTTTCATGGTGTAGCCAAAGTCCATATGAGTCTTTGGCTTTGTGTCTTCTA | 72 (0.001362%) | | | TCATCAGAGCAACGGGCAGAGCCCGCGTCGACCTTTTATCTAATAAATGCGTCCCTTCCATAAGTCGGGGTTTGTTGCACGTATTAGCTCTAGAATTACT | 4 (0.000076%) | | | TCATGAATCATCAGAGCAACGGGCAGAGCCCGCGTCGACCTTTTATCTAATAAATGCGTCCCTTCCATAAGTCGGGGTTTGTTGCACGTATTAGCTCTAG | 18 (0.000341%) | | | TCATGTGTATGATTGAGTATAAGAACTTAAACCGCAACCG | 4662 (0.035279%) | | | TCCAACCTAGGCGAGACAAGGGTTCACATTTCGTTCATCACCCTTGGCCGGCTATCGAACAGCCGGACTCCCATCAAAAGATGGTTGCCAAGAACATCTT | 3 (0.000057%) | | | TCCAACCTAGGCGAGACAAGGGTTCACATTTCGTTCATCACCCTTGGCCGGCTTTCGAACAGCCGGACTCCCATCAAAAGATGGTTGCCAAGAACATCTT | 5 (0.000095%) | | | TCCAACGAAGCACGCCCATCCAACCTAGGCGAGACAAGGGTTCACATTTCGTTCATCACCCTTGGCCGGCTATCGAACAGCCGGACTCCCATCAAAAGAT | 19 (0.000359%) | | | TCCAACGAAGCACGCCCATCCAACCTAGGCGAGACAAGGGTTCACATTTCGTTCATCACCCTTGGCCGGCTTTCGAACAGCCGGACTCCCATCAAAAGAT | 19 (0.000359%) | | | TCCAATTACCAGACTCGAAAGAGCCCGGTATTGTTATTTATTGTCACTACCTCCCCGTGTCAGGATTGGGTAATTTGCGCGCCTGCTGCCTTCCTTGGAT | 3 (0.000057%) | | | TCCAGAGCGTAGGCTTGCTTTGAGCACTCTAATTTCTTCAAAGTAACAGCGCCGGAGGCACGACCCGGCCAATTAAGACCAGGAGCGTATCGCCGACCGA | 5 (0.000095%) | | | TCCATATGAGTCTTTGGCTTTGTGTCTTCTAACAAGGATA | 265 (0.002005%) | | | TCCATCGACCAGAGGCTGTTCACCTTGGAGACCTGATGCGGTTATGAGTACGACCGGGCGTGAGCGGCACTCGGTCCTCCGGATTTTCAAGGGCCGCCGG | 4 (0.000076%) | | | TCCATGCTTTCCAACGAAGCACGCCCATCCAACCTAGGCGAGACAAGGGTTCACATTTCGTTCATCACCCTTGGCCGGCTATCGAACAGCCGGACTCCCA | 13 (0.000246%) | | | TCCATTGGAGGGCAAGTCTGGTGCCAGCAGCCGCGGTAATTCCAGCTCCAATAGCGTATATTTAAGTTGTTGCAGTTAAAAAGCTCGTAGTTGAACCTTG | 4 (0.000076%) | | | TCCCATGCTAATGTATCCAGAGCGTAGGCTTGCTTTGAGCACTCTAATTTCTTCAAAGTAACAGCGCCGGAGGCACGACCCGGCCAATTAAGACCAGGAG | 11 (0.000208%) | | | TCCCTCACCATCCTTTGCTGATGCGGGACGGAAGCTGGTCTCCCGTGTGTTACCGCACGCGGTTGGCCTAAATCCGAGCCAAGGACGCCTGGAGCGTACC | 3 (0.000057%) | | | TCCCTTGCCTACATTGTTCCATCGACCAGAGGCTGTTCACCTTGGAGACCTGATGCGGTTATGAGTACGACCGGGCGTGAGCGGCACTCGGTCCTCCGGA | 4 (0.000076%) | | | TCCGACTTTGTGAAATGACTTGAGAGGTGTAGGATAAGTGGGAGCTTCGGCGCAAGTGAAATACCACTACTTTTAACGTTATTTTACTTACTCCGTGAAT | 5 (0.000095%) | | | TCCGCCTAGGCTGTCCCGAGTGTGAGCGAGGTGTGAGTGTCGCCCATGGGCATCGACACCTTGCGGCTAGGAACTGGAACGAGACGGGTAGCAAAGATTT | 164 (0.003103%) | | | TCCGCCTAGGCTGTCCCGAGTGTGAGCGAGGTGTGAGTGTCGCCCATGGGCATCGACACCTTGCGGCTAGGAACTGGAACGAGACGGGTGGCAAAGATTT | 289 (0.005467%) | | | TCCGCTTATTGATATGCTTAAACTCAGCGGGTAATCCCGCCTGACCTGGGGTCGCTATATGGACTTTGGGTCATCTACAGCTTCCGGACAAGAGCGACCG | 3 (0.000057%) | | | TCCGGGCGTCGGCCTGTGGGCTCCCCATTCGACCCGTCTTGAAACACGGACCAAGGAGTCTGACATGTGTGCGAGTCAACGGGTGAGTAAACCCGTAAGG | 3 (0.000057%) | | | TCCGGGGAGGCGAATGCCAGCCGTTCGTTTGCATGTTCCTTGACACTTTTCGTGCCGGGGTTTTGTGATATCCGGAAGCAACGCGCACGACAAGACCGAG | 8 (0.000151%) | | | TCCGGTTAAAATTCCGGAACCGGGACGTGGCGGTTGACGGCAACGTTAGGGAGTCCGGAGACGTCGGCGGGGGCCTCGGGAAGAGTTATCTTTTCTGTTT | 20 (0.000378%) | | | TCCGTCGAGTTATCATGAATCATCAGAGCAACGGGCAGAGCCCGCGTCGACCTTTTATCTAATAAATGCGTCCCTTCCATAAGTCGGGGTTTGTTGCACG | 15 (0.000284%) | | | TCCGTCTCCGGGGAGGCGAATGCCAGCCGTTCGTTTGCATGTTCCTTGACACTTTTCGTGCCGGGGTTTTGTGATATCCGGAAGCAACGCGCACGACAAG | 762 (0.014416%) | | | TCCTATGATGTTATCCCATGCTAATGTATCCAGAGCGTAGGCTTGCTTTGAGCACTCTAATTTCTTCAAAGTAACAGCGCCGGAGGCACGACCCGGCCAA | 9 (0.000170%) | | | TCCTCGTTAAGGGATTTAGATTGTACTCATTCCAATTACCAGACTCGAAAGAGCCCGGTATTGTTATTTATTGTCACTACCTCCCCGTGTCAGGATTGGG | 23 (0.000435%) | | | TCCTTGTTAGAAGACACAAAGCCAAAGACTCATATGGACTTTGGCTACACCATGAAAGCTTTGAGAAGCAAGAAGAAGGTTGGTTAGTGTTTTGGAGTCG | 8 (0.000151%) | | | TCGAAAGAGCCCGGTATTGTTATTTATTGTCACTACCTCCCCGTGTCAGGATTGGGTAATTTGCGCGCCTGCTGCCTTCCTTGGATGTGGTAGCCGTTTC | 8 (0.000151%) | | | TCGAAATCCGCTAAGGAGTGTGTAACAACTCACCTGCCGAATCAACTAGCCCCGAAAATGGATGGCGCTTAAGCGCGCGACCTATACCCGGCCGTCGGGG | 58 (0.001097%) | | | TCGAAATCCTATGATGTTATCCCATGCTAATGTATCCAGAGCGTAGGCTTGCTTTGAGCACTCTAATTTCTTCAAAGTAACAGCGCCGGAGGCACGACCC | 227 (0.004294%) | | | TCGAATATGACTTGATGTCATGTGTATGATTGAGTATAAG | 9266 (0.070119%) | | | TCGACCTTTTATCTAATAAATGCGTCCCTTCCATAAGTCGGGGTTTGTTGCACGTATTAGCTCTAGAATTACTACGGTTATCCGAGTAGTAGTTACCATC | 10 (0.000189%) | | | TCGACTCCAAAACACTAACCAACCTTCTTCTTGCTTCTCAAAGCTTTCATGGTGTAGCCAAAGTCCATATGAGTCTTTGGCTTTGTGTCTTCTAACAAGG | 34 (0.000643%) | | | TCGAGTTATCATGAATCATCAGAGCAACGGGCAGAGCCCGCGTCGACCTTTTATCTAATAAATGCGTCCCTTCCATAAGTCGGGGTTTGTTGCACGTATT | 63 (0.001192%) | | | TCGATCACGGCAATTCCCCGCCACATCCTCTCAAACGCAATGGAAAGAGAGAAAGGACGAGGTCTTGACCGTCATCTTTTGCCCGAAGGACGGATGAGCT | 3 (0.000057%) | | | TCGATCCATGCTTTCCAACGAAGCACGCCCATCCAACCTAGGCGAGACAAGGGTTCACATTTCGTTCATCACCCTTGGCCGGCTATCGAACAGCCGGACT | 6 (0.000114%) | | | TCGATCCGGTTAAAATTCCGGAACCGGGACGTGGCGGTTGACGGCAACGTTAGGGAGTCCGGAGACGTCGGCGGGGGCCTCGGGAAGAGTTATCTTTTCT | 5 (0.000095%) | | | TCGATCGAAATCCTATGATGTTATCCCATGCTAATGTATCCAGAGCGTAGGCTTGCTTTGAGCACTCTAATTTCTTCAAAGTAACAGCGCCGGAGGCACG | 3 (0.000057%) | | | TCGATCGGGTTGCGGTTTAAGTTGTTATACTCAATCATAC | 9 (0.000068%) | | | TCGCCGCCCGCGACGTCGCGAGAAGTCCACTAAACCTTATCATTTAGAGGAAGGAGAAGTCGTAACAAGGTTTCCGTAGGTGAACCTGCGGAAGGATCGA | 10 (0.000189%) | | | TCGGAAGAGCACACGTCTGAACTCCAGTCACGAGATTCCA | 6 (0.000045%) | | | TCGGCATCAGCGTGCTCCGGGCGTCGGCCTGTGGGCTCCCCATTCGACCCGTCTTGAAACACGGACCAAGGAGTCTGACATGTGTGCGAGTCAACGGGTG | 17 (0.000322%) | | | TCGGCCTGTGGGCTCCCCATTCGACCCGTCTTGAAACACGGACCAAGGAGTCTGACATGTGTGCGAGTCAACGGGTGAGTAAACCCGTAAGGCGCAAGGA | 15 (0.000284%) | | | TCGGGTTGCGGTTTAAGTTGTTATACTCAATCATACACAT | 144 (0.001090%) | | | TCGGTACGCTCCAGGCGTCCTTGGCTCGGATTTAGGCCAACCGCGTGCGGTAACACACGGGAGACCAGCTTCCGTCCCGCATCAGCAAAGGATGGTGAGG | 3 (0.000057%) | | | TCGGTAGGAGCGACGGGCGGTGTGTACAAAGGGCAGGGACGTAGTCAACGCGAGCTGATGACTCGCGCTTACTAGGAATTCCTCGTTGAAGACCAACAAT | 21 (0.000397%) | | | TCGTACTGAAAATCAGAATCAAACGAGCTTTTACCCTTTTGTTCCACACGAGATTTCTGTTCTCGTTGAGCTCATCTTAGGACACCTGCGTTATCTTTTA | 3 (0.000057%) | | | TCGTATTCGTACTGAAAATCAGAATCAAACGAGCTTTTACCCTTTTGTTCCACACGAGATTTCTGTTCTCGTTGAGCTCATCTTAGGACACCTGCGTTAT | 8 (0.000151%) | | | TCGTCCCTCACCATCCTTTGCTGATGCGGGACGGAAGCTGGTCTCCCGTGTGTTACCGCACGCGGTTGGCCTAAATCCGAGCCAAGGACGCCTGGAGCGT | 5 (0.000095%) | | | TCGTTAAGGGATTTAGATTGTACTCATTCCAATTACCAGACTCGAAAGAGCCCGGTATTGTTATTTATTGTCACTACCTCCCCGTGTCAGGATTGGGTAA | 8 (0.000151%) | | | TCTAATAAATGCGTCCCTTCCATAAGTCGGGGTTTGTTGCACGTATTAGCTCTAGAATTACTACGGTTATCCGAGTAGTAGTTACCATCAAACAAACTAT | 3 (0.000057%) | | | TCTAGTCCGACTTTGTGAAATGACTTGAGAGGTGTAGGATAAGTGGGAGCTTCGGCGCAAGTGAAATACCACTACTTTTAACGTTATTTTACTTACTCCG | 15 (0.000284%) | | | TCTCAAACGCAATGGAAAGAGAGAAAGGACGAGGTCTTGACCGTCATCTTTTGCCCGAAGGACGGATGAGCTTTGGCGGGACTGAATCACTTCGAGTCAC | 7 (0.000132%) | | | TCTCAAAGCTTTCATGGTGTAGCCAAAGTCCATATGAGTCTTTGGCTTTGTGTCTTCTAACAAGGAAACACTACTTAGGCTTATAAGATGCGGTTGCGGT | 4 (0.000076%) | | | TCTCTCTCTCTCTCTCTCTCTCTCTCTCTCTCTCTCTCTC | 80 (0.000605%) | | | TCTGGGCCGCACGCGCGCTACACTGATGTATTCAACGAGTTCACACCTTGGCCGACAGGCCCGGGTAATCTTTGAAATTTCATCGTGATGGGGATAGATC | 4 (0.000076%) | | | TCTGGTAAATTTTCATAATTTTTTGACACCTCTAGCTAGGTCATTTGACCTGATACAACATCGGATTTTCATGGTCTAGTTGGGGCTCCGTGGGCATATT | 3 (0.000057%) | | | TCTGGTGCCAGCAGCCGCGGTAATTCCAGCTCCAATAGCGTATATTTAAGTTGTTGCAGTTAAAAAGCTCGTAGTTGAACCTTGGGATGGGTCGGCCGGT | 16 (0.000303%) | | | TCTGTGATGCCCTTAGATGTTCTGGGCCGCACGCGCGCTACACTGATGTATTCAACGAGTTCACACCTTGGCCGACAGGCCCGGGTAATCTTTGAAATTT | 9 (0.000170%) | | | TCTTAAAAGCCTAAGTAGTGTTTCCTTGTTAGAAGACACAAAGCCAAAGACTCATATGGACTTTGGCTACACCATGAAAGCTTTGAGAAGCAAGAAGAAG | 37 (0.000700%) | | | TCTTAAAGGCGTAAGAATTGTATCCTTGTTAAAAGACACAAAGCCAAAGACTCATATGGACTTTGGCTACACCATGAAAGCTTTGAGAAGCAAGAAGAAG | 102 (0.001930%) | | | TCTTAAAGGCGTAAGAATTGTATCCTTGTTAGAAGACACAAAGCCAAAGACTCATATGGACTTTGGCTACACCATGAAAGCTTTGAGAAGCAAGAAGAAG | 74 (0.001400%) | | | TCTTATACTCAATCATACACATGACATCAAGTCATATTCGACTCCAAAACACTAACCAACCTTCTTCTTGCTTCTCAAAGCTTTCATGGTGTAGCCAAAG | 226 (0.004276%) | | | TCTTCAACGAGGAATTCCTAGTAAGCGCGAGTCATCAGCTCGCGTTGACTACGTCCCTGCCCTTTGTACACACCGCCCGTCGCTCCTACCGATTGAATGA | 14 (0.000265%) | | | TCTTGCTTCTCAAAGCTTTCATGGTGTAGCCAAAGTCCATATGAGTCTTTGGCTTTGTGTCTTCTAACAAGGAAACACTACTTAGGCTTATAAGATGCGG | 5 (0.000095%) | | | TCTTTTCCTCCGCTTATTGATATGCTTAAACTCAGCGGGTAATCCCGCCTGACCTGGGGTCGCTATATGGACTTTGGGTCATCTACAGCTTCCGGACAAG | 3 (0.000057%) | | | TGAAAATCAGAATCAAACGAGCTTTTACCCTTTTGTTCCACACGAGATTTCTGTTCTCGTTGAGCTCATCTTAGGACACCTGCGTTATCTTTTAACAGAT | 43 (0.000813%) | | | TGAATCATCAGAGCAACGGGCAGAGCCCGCGTCGACCTTTTATCTAATAAATGCGTCCCTTCCATAAGTCGGGGTTTGTTGCACGTATTAGCTCTAGAAT | 10 (0.000189%) | | | TGACATCAAGTCATATTCGACTCCAAAACACTAACCAACCTTCTTCTTGCTTCTCAAAGCTTTCATGGTGTAGCCAAAGTCCATATGAGTCTTTGGCTTT | 31 (0.000586%) | | | TGACTCTAGTCCGACTTTGTGAAATGACTTGAGAGGTGTAGGATAAGTGGGAGCTTCGGCGCAAGTGAAATACCACTACTTTTAACGTTATTTTACTTAC | 12 (0.000227%) | | | TGACTTGATGTCATGTGTATGATTGAGTATAAGAACTTAA | 81 (0.000613%) | | | TGAGCGAGGTGTGAGTGTCGCCCATGGGCATCGACACCTTGCGGCTAGGAACTGGAACGAGACGGGTAGCAAAGATTTCGAGTAGCACTTCATACTACCG | 4 (0.000076%) | | | TGAGCGAGGTGTGAGTGTCGCCCATGGGCATCGACACCTTGCGGCTAGGAACTGGAACGAGACGGGTGGCAAAGATTTCGAGTAGCACTTCATACTACCG | 5 (0.000095%) | | | TGAGCTTGACTCTAGTCCGACTTTGTGAAATGACTTGAGAGGTGTAGGATAAGTGGGAGCTTCGGCGCAAGTGAAATACCACTACTTTTAACGTTATTTT | 9 (0.000170%) | | | TGAGTGTCGCCCATGGGCATCGACACCTTGCGGCTAGGAACTGGAACGAGACGGGTAGCAAAGATTTCGAGTAGCACTTCATACTACCGTGGGTTTTTTA | 6 (0.000114%) | | | TGAGTGTCGCCCATGGGCATCGACACCTTGCGGCTAGGAACTGGAACGAGACGGGTGGCAAAGATTTCGAGTAGCACTTCATACTACCGTGGGTTTTTTA | 6 (0.000114%) | | | TGATGCCCTTAGATGTTCTGGGCCGCACGCGCGCTACACTGATGTATTCAACGAGTTCACACCTTGGCCGACAGGCCCGGGTAATCTTTGAAATTTCATC | 3 (0.000057%) | | | TGATGCGGGACGGAAGCTGGTCTCCCGTGTGTTACCGCACGCGGTTGGCCTAAATCCGAGCCAAGGACGCCTGGAGCGTACCGACATGCGGTGGTGAACT | 7 (0.000132%) | | | TGATGGTAACTACTACTCGGATAACCGTAGTAATTCTAGAGCTAATACGTGCAACAAACCCCGACTTATGGAAGGGACGCATTTATTAGATAAAAGGTCG | 4 (0.000076%) | | | TGATGTCATGTGTATGATTGAGTATAAGAACTTAAACCGC | 98 (0.000742%) | | | TGATGTTATCCCATGCTAATGTATCCAGAGCGTAGGCTTGCTTTGAGCACTCTAATTTCTTCAAAGTAACAGCGCCGGAGGCACGACCCGGCCAATTAAG | 28 (0.000530%) | | | TGATTAACAGGGACAGTCGGGGGCATTCGTATTTCATAGTCAGAGGTGAAATTCTTGGATTTATGAAAGACGAACAACTGCGAAAGCATTTGCCAAGGAT | 113 (0.002138%) | | | TGCAGCACGCGCCTAACGGCGTGCCTCGGCATCAGCGTGCTCCGGGCGTCGGCCTGTGGGCTCCCCATTCGACCCGTCTTGAAACACGGACCAAGGAGTC | 80 (0.001513%) | | | TGCATGTGTAAGTATGAACGAATTCAGACTGTGAAACTGCGAATGGCTCATTAAATCAGTTATAGTTTGTTTGATGGTAACTACTACTCGGATAACCGTA | 8 (0.000151%) | | | TGCCAGCAGCCGCGGTAATTCCAGCTCCAATAGCGTATATTTAAGTTGTTGCAGTTAAAAAGCTCGTAGTTGAACCTTGGGATGGGTCGGCCGGTCCGCC | 6 (0.000114%) | | | TGCCAGTAGTCATATGCTTGTCTCAAAGATTAAGCCATGCATGTGTAAGTATGAACGAATTCAGACTGTGAAACTGCGAATGGCTCATTAAATCAGTTAT | 125 (0.002365%) | | | TGCCCTTAGATGTTCTGGGCCGCACGCGCGCTACACTGATGTATTCAACGAGTTCACACCTTGGCCGACAGGCCCGGGTAATCTTTGAAATTTCATCGTG | 8 (0.000151%) | | | TGCCGACTTCCCTTGCCTACATTGTTCCATCGACCAGAGGCTGTTCACCTTGGAGACCTGATGCGGTTATGAGTACGACCGGGCGTGAGCGGCACTCGGT | 95 (0.001797%) | | | TGCCTACATTGTTCCATCGACCAGAGGCTGTTCACCTTGGAGACCTGATGCGGTTATGAGTACGACCGGGCGTGAGCGGCACTCGGTCCTCCGGATTTTC | 12 (0.000227%) | | | TGCGAATGGCTCATTAAATCAGTTATAGTTTGTTTGATGGTAACTACTACTCGGATAACCGTAGTAATTCTAGAGCTAATACGTGCAACAAACCCCGACT | 10 (0.000189%) | | | TGCGGTTTAAGTTCTTATACTCAATCATACACATGACATC | 3686 (0.027893%) | | | TGCTAATGTATCCAGAGCGTAGGCTTGCTTTGAGCACTCTAATTTCTTCAAAGTAACAGCGCCGGAGGCACGACCCGGCCAATTAAGACCAGGAGCGTAT | 11 (0.000208%) | | | TGCTCCGGGCGTCGGCCTGTGGGCTCCCCATTCGACCCGTCTTGAAACACGGACCAAGGAGTCTGACATGTGTGCGAGTCAACGGGTGAGTAAACCCGTA | 5 (0.000095%) | | | TGCTGATGCGGGACGGAAGCTGGTCTCCCGTGTGTTACCGCACGCGGTTGGCCTAAATCCGAGCCAAGGACGCCTGGAGCGTACCGACATGCGGTGGTGA | 3 (0.000057%) | | | TGCTTCTCAAAGCTTTCATGGTGTAGCCAAAGTCCATATGAGTCTTTGGCTTTGTGTCTTCTAACAAGGAAACACTACTTAGGCTTATAAGATGCGGTTG | 3 (0.000057%) | | | TGCTTGTCTCAAAGATTAAGCCATGCATGTGTAAGTATGAACGAATTCAGACTGTGAAACTGCGAATGGCTCATTAAATCAGTTATAGTTTGTTTGATGG | 5 (0.000095%) | | | TGCTTTCCAACGAAGCACGCCCATCCAACCTAGGCGAGACAAGGGTTCACATTTCGTTCATCACCCTTGGCCGGCTATCGAACAGCCGGACTCCCATCAA | 17 (0.000322%) | | | TGGAAAGAGAGAAAGGACGAGGTCTTGACCGTCATCTTTTGCCCGAAGGACGGATGAGCTTTGGCGGGACTGAATCACTTCGAGTCACCGTCGACAACTT | 10 (0.000189%) | | | TGGAAGTCGAAATCCGCTAAGGAGTGTGTAACAACTCACCTGCCGAATCAACTAGCCCCGAAAATGGATGGCGCTTAAGCGCGCGACCTATACCCGGCCG | 936 (0.017707%) | | | TGGACTTTGGCTACACCATGAAAGCTTTGAGAAGCAAGAAGAAGGTTGGTTAGTGTTTTGGAGTCGAATATGACTTGATGTCATGTGTATGATTGAGTAT | 53 (0.001003%) | | | TGGAGGGCAAGTCTGGTGCCAGCAGCCGCGGTAATTCCAGCTCCAATAGCGTATATTTAAGTTGTTGCAGTTAAAAAGCTCGTAGTTGAACCTTGGGATG | 298 (0.005638%) | | | TGGCTGGGGCGGCACATCTGTTAAAAGATAACGCAGGTGTCCTAAGATGAGCTCAACGAGAACAGAAATCTCGTGTGGAACAAAAGGGTAAAAGCTCGTT | 4 (0.000076%) | | | TGGGCATCGACACCTTGCGGCTAGGAACTGGAACGAGACGGGTGGCAAAGATTTCGAGTAGCACTTCATACTACCGTGGGTTTTTTAAACCTTCCGAGTT | 27 (0.000511%) | | | TGGGCCGCACGCGCGCTACACTGATGTATTCAACGAGTTCACACCTTGGCCGACAGGCCCGGGTAATCTTTGAAATTTCATCGTGATGGGGATAGATCGA | 16 (0.000303%) | | | TGGGGAGTTTGGCTGGGGCGGCACATCTGTTAAAAGATAACGCAGGTGTCCTAAGATGAGCTCAACGAGAACAGAAATCTCGTGTGGAACAAAAGGGTAA | 4 (0.000076%) | | | TGGGGCGGCACATCTGTTAAAAGATAACGCAGGTGTCCTAAGATGAGCTCAACGAGAACAGAAATCTCGTGTGGAACAAAAGGGTAAAAGCTCGTTTGAT | 14 (0.000265%) | | | TGGGTGGTTCGCCGCCCGCGACGTCGCGAGAAGTCCACTAAACCTTATCATTTAGAGGAAGGAGAAGTCGTAACAAGGTTTCCGTAGGTGAACCTGCGGA | 7 (0.000132%) | | | TGGTCTTCAACGAGGAATTCCTAGTAAGCGCGAGTCATCAGCTCGCGTTGACTACGTCCCTGCCCTTTGTACACACCGCCCGTCGCTCCTACCGATTGAA | 52 (0.000984%) | | | TGGTGCCAGCAGCCGCGGTAATTCCAGCTCCAATAGCGTATATTTAAGTTGTTGCAGTTAAAAAGCTCGTAGTTGAACCTTGGGATGGGTCGGCCGGTCC | 5 (0.000095%) | | | TGGTTCGCCGCCCGCGACGTCGCGAGAAGTCCACTAAACCTTATCATTTAGAGGAAGGAGAAGTCGTAACAAGGTTTCCGTAGGTGAACCTGCGGAAGGA | 3 (0.000057%) | | | TGTACTCATTCCAATTACCAGACTCGAAAGAGCCCGGTATTGTTATTTATTGTCACTACCTCCCCGTGTCAGGATTGGGTAATTTGCGCGCCTGCTGCCT | 9 (0.000170%) | | | TGTAGCCAAAGTCCATATGAGTCTTTGGCTTTGTGTCTTT | 6910 (0.052290%) | | | TGTATCCAGAGCGTAGGCTTGCTTTGAGCACTCTAATTTCTTCAAAGTAACAGCGCCGGAGGCACGACCCGGCCAATTAAGACCAGGAGCGTATCGCCGA | 26 (0.000492%) | | | TGTATCCTTGTTAGAAGACACAAAGCCAAAGACTCATATGGACTTTGGCTACACCATGAAAGCTTTGAGAAGCAAGAAGAAGGTTGGTTAGTGTTTTGGA | 37 (0.000700%) | | | TGTCAGGTGGGGAGTTTGGCTGGGGCGGCACATCTGTTAAAAGATAACGCAGGTGTCCTAAGATGAGCTCAACGAGAACAGAAATCTCGTGTGGAACAAA | 36 (0.000681%) | | | TGTCATGTGTATGATTGAGTATAAGAACTTAAACCGCAAC | 725 (0.005486%) | | | TGTCCCGAGTGTGAGCGAGGTGTGAGTGTCGCCCATGGGCATCGACACCTTGCGGCTAGGAACTGGAACGAGACGGGTGGCAAAGATTTCGAGTAGCACT | 7 (0.000132%) | | | TGTCGCCCATGGGCATCGACACCTTGCGGCTAGGAACTGGAACGAGACGGGTGGCAAAGATTTCGAGTAGCACTTCATACTACCGTGGGTTTTTTAAACC | 3 (0.000057%) | | | TGTCGGTACGCTCCAGGCGTCCTTGGCTCGGATTTAGGCCAACCGCGTGCGGTAACACACGGGAGACCAGCTTCCGTCCCGCATCAGCAAAGGATGGTGA | 3 (0.000057%) | | | TGTCTCAAAGATTAAGCCATGCATGTGTAAGTATGAACGAATTCAGACTGTGAAACTGCGAATGGCTCATTAAATCAGTTATAGTTTGTTTGATGGTAAC | 4 (0.000076%) | | | TGTGAGCGAGGTGTGAGTGTCGCCCATGGGCATCGACACCTTGCGGCTAGGAACTGGAACGAGACGGGTAGCAAAGATTTCGAGTAGCACTTCATACTAC | 7 (0.000132%) | | | TGTGAGCGAGGTGTGAGTGTCGCCCATGGGCATCGACACCTTGCGGCTAGGAACTGGAACGAGACGGGTGGCAAAGATTTCGAGTAGCACTTCATACTAC | 3 (0.000057%) | | | TGTGAGTGTCGCCCATGGGCATCGACACCTTGCGGCTAGGAACTGGAACGAGACGGGTAGCAAAGATTTCGAGTAGCACTTCATACTACCGTGGGTTTTT | 10 (0.000189%) | | | TGTGAGTGTCGCCCATGGGCATCGACACCTTGCGGCTAGGAACTGGAACGAGACGGGTGGCAAAGATTTCGAGTAGCACTTCATACTACCGTGGGTTTTT | 9 (0.000170%) | | | TGTGATGCCCTTAGATGTTCTGGGCCGCACGCGCGCTACACTGATGTATTCAACGAGTTCACACCTTGGCCGACAGGCCCGGGTAATCTTTGAAATTTCA | 6 (0.000114%) | | | TGTTAAAAGATAACGCAGGTGTCCTAAGATGAGCTCAACGAGAACAGAAATCTCGTGTGGAACAAAAGGGTAAAAGCTCGTTTGATTCTGATTTTCAGTA | 3 (0.000057%) | | | TGTTAGAAGACACAAAGCCAAAGACTCATATGGACTTTGGCTACACCATGAAAGCTTTGAGAAGCAAGAAGAAGGTTGGTTAGTGTTTTGGAGTCGAATA | 39 (0.000738%) | | | TGTTAGTTTCTTTTCCTCCGCTTATTGATATGCTTAAACTCAGCGGGTAATCCCGCCTGACCTGGGGTCGCTATATGGACTTTGGGTCATCTACAGCTTC | 8 (0.000151%) | | | TGTTATCCCATGCTAATGTATCCAGAGCGTAGGCTTGCTTTGAGCACTCTAATTTCTTCAAAGTAACAGCGCCGGAGGCACGACCCGGCCAATTAAGACC | 14 (0.000265%) | | | TGTTCCATCGACCAGAGGCTGTTCACCTTGGAGACCTGATGCGGTTATGAGTACGACCGGGCGTGAGCGGCACTCGGTCCTCCGGATTTTCAAGGGCCGC | 3 (0.000057%) | | | TGTTGAGCTTGACTCTAGTCCGACTTTGTGAAATGACTTGAGAGGTGTAGGATAAGTGGGAGCTTCGGCGCAAGTGAAATACCACTACTTTTAACGTTAT | 15 (0.000284%) | | | TGTTGGTCTTCAACGAGGAATTCCTAGTAAGCGCGAGTCATCAGCTCGCGTTGACTACGTCCCTGCCCTTTGTACACACCGCCCGTCGCTCCTACCGATT | 199 (0.003765%) | | | TGTTTCCTTGTTAGAAGACACAAAGCCAAAGACTCATATGGACTTTGGCTACACCATGAAAGCTTTGAGAAGCAAGAAGAAGGTTGGTTAGTGTTTTGGA | 27 (0.000511%) | | | TTAAAAGATAACGCAGGTGTCCTAAGATGAGCTCAACGAGAACAGAAATCTCGTGTGGAACAAAAGGGTAAAAGCTCGTTTGATTCTGATTTTCAGTACG | 25 (0.000473%) | | | TTAAAAGCCTAAGTAGTGTTTCCTTGTTAGAAGACACAAAGCCAAAGACTCATATGGACTTTGGCTACACCATGAAAGCTTTGAGAAGCAAGAAGAAGGT | 146 (0.002762%) | | | TTAAAGGCGTAAGAATTGTATCCTTGTTAAAAGACACAAAGCCAAAGACTCATATGGACTTTGGCTACACCATGAAAGCTTTGAGAAGCAAGAAGAAGGT | 151 (0.002857%) | | | TTAAAGGCGTAAGAATTGTATCCTTGTTAGAAGACACAAAGCCAAAGACTCATATGGACTTTGGCTACACCATGAAAGCTTTGAGAAGCAAGAAGAAGGT | 116 (0.002195%) | | | TTAAATCAGTTATAGTTTGTTTGATGGTAACTACTACTCGGATAACCGTAGTAATTCTAGAGCTAATACGTGCAACAAACCCCGACTTATGGAAGGGACG | 5 (0.000095%) | | | TTAACAGGGACAGTCGGGGGCATTCGTATTTCATAGTCAGAGGTGAAATTCTTGGATTTATGAAAGACGAACAACTGCGAAAGCATTTGCCAAGGATGTT | 203 (0.003840%) | | | TTAAGCCATGCATGTGTAAGTATGAACGAATTCAGACTGTGAAACTGCGAATGGCTCATTAAATCAGTTATAGTTTGTTTGATGGTAACTACTACTCGGA | 15 (0.000284%) | | | TTAAGCGCGCGACCTATACCCGGCCGTCGGGGCAAGAGCCAGGCCTCGATGAGTAGGAGGGCGCGGCGGTCGCTGCAAAACCTAGGGCGCGAGCCCGGGC | 12 (0.000227%) | | | TTAAGGGATTTAGATTGTACTCATTCCAATTACCAGACTCGAAAGAGCCCGGTATTGTTATTTATTGTCACTACCTCCCCGTGTCAGGATTGGGTAATTT | 17 (0.000322%) | | | TTAAGTTCTTATACTCAATCATACACATGACATCAAGTCATATTCGACTCCAAAACACTAACCAACCTTCTTCTTGCTTCTCAAAGCTTTCATGGTGTAG | 2940 (0.055620%) | | | TTAAGTTGTTATACTCAATCATACACATGACAACAAGTCATATTCGACTCCAAAACACTAACCAACCTTCTTCTTGCTTCTCAAAGCTTTCATGGTGTAG | 64 (0.001211%) | | | TTAAGTTGTTATACTCAATCATACACATGACATCAAGTCA | 457 (0.003458%) | | | TTACCAGACTCGAAAGAGCCCGGTATTGTTATTTATTGTCACTACCTCCCCGTGTCAGGATTGGGTAATTTGCGCGCCTGCTGCCTTCCTTGGATGTGGT | 34 (0.000643%) | | | TTACGGGTTTACTCACCCGTTGACTCGCACACATGTCAGACTCCTTGGTCCGTGTTTCAAGACGGGTCGAATGGGGAGCCCACAGGCCGACGCCCGGAGC | 3214 (0.060803%) | | | TTACTCACCCGTTGACTCGCACACATGTCAGACTCCTTGGTCCGTGTTTCAAGACGGGTCGAATGGGGAGCCCACAGGCCGACGCCCGGAGCACGCTGAT | 9 (0.000170%) | | | TTAGAAGACACAAAGCCAAAGACTCATATGGACTTTGGCTACACCATGAAAGCTTTGAGAAGCAAGAAGAAGGTTGGTTAGTGTTTTGGAGTCGAATATG | 97 (0.001835%) | | | TTAGATGTTCTGGGCCGCACGCGCGCTACACTGATGTATTCAACGAGTTCACACCTTGGCCGACAGGCCCGGGTAATCTTTGAAATTTCATCGTGATGGG | 15 (0.000284%) | | | TTAGATTGTACTCATTCCAATTACCAGACTCGAAAGAGCCCGGTATTGTTATTTATTGTCACTACCTCCCCGTGTCAGGATTGGGTAATTTGCGCGCCTG | 23 (0.000435%) | | | TTAGTTTCTTTTCCTCCGCTTATTGATATGCTTAAACTCAGCGGGTAATCCCGCCTGACCTGGGGTCGCTATATGGACTTTGGGTCATCTACAGCTTCCG | 4 (0.000076%) | | | TTATACTCAATCATACACATGACAACAAGTCATATTCGACTCCAAAACACTAACCAACCTTCTTCTTGCTTCTCAAAGCTTTCATGGTGTAGCCAAAGTC | 295 (0.005581%) | | | TTATACTCAATCATACACATGACATCAAGTCATATTCGACTCCAAAACACTAACCAACCTTCTTCTTGCTTCTCAAAGCTTTCATGGTGTAGCCAAAGTC | 827 (0.015645%) | | | TTATAGTTTGTTTGATGGTAACTACTACTCGGATAACCGTAGTAATTCTAGAGCTAATACGTGCAACAAACCCCGACTTATGGAAGGGACGCATTTATTA | 5 (0.000095%) | | | TTATCATGAATCATCAGAGCAACGGGCAGAGCCCGCGTCGACCTTTTATCTAATAAATGCGTCCCTTCCATAAGTCGGGGTTTGTTGCACGTATTAGCTC | 74 (0.001400%) | | | TTATCCCATGCTAATGTATCCAGAGCGTAGGCTTGCTTTGAGCACTCTAATTTCTTCAAAGTAACAGCGCCGGAGGCACGACCCGGCCAATTAAGACCAG | 3 (0.000057%) | | | TTATCTAATAAATGCGTCCCTTCCATAAGTCGGGGTTTGTTGCACGTATTAGCTCTAGAATTACTACGGTTATCCGAGTAGTAGTTACCATCAAACAAAC | 5 (0.000095%) | | | TTATTGATATGCTTAAACTCAGCGGGTAATCCCGCCTGACCTGGGGTCGCTATATGGACTTTGGGTCATCTACAGCTTCCGGACAAGAGCGACCGATAAA | 13 (0.000246%) | | | TTCAACGAGGAATTCCTAGTAAGCGCGAGTCATCAGCTCGCGTTGACTACGTCCCTGCCCTTTGTACACACCGCCCGTCGCTCCTACCGATTGAATGATC | 12 (0.000227%) | | | TTCAATCGGTAGGAGCGACGGGCGGTGTGTACAAAGGGCAGGGACGTAGTCAACGCGAGCTGATGACTCGCGCTTACTAGGAATTCCTCGTTGAAGACCA | 80 (0.001513%) | | | TTCACATTTCGTTCATCACCCTTGGCCGGCTATCGAACAGCCGGACTCCCATCAAAAGATGGTTGCCAAGAACATCTTCGTTACGGTTTGCTAATTCTCG | 22 (0.000416%) | | | TTCACATTTCGTTCATCACCCTTGGCCGGCTTTCGAACAGCCGGACTCCCATCAAAAGATGGTTGCCAAGAACATCTTCGTTACGGTTTGCTAATTCTCG | 22 (0.000416%) | | | TTCACCACCGCATGTCGGTACGCTCCAGGCGTCCTTGGCTCGGATTTAGGCCAACCGCGTGCGGTAACACACGGGAGACCAGCTTCCGTCCCGCATCAGC | 11 (0.000208%) | | | TTCACGGTTCGTATTCGTACTGAAAATCAGAATCAAACGAGCTTTTACCCTTTTGTTCCACACGAGATTTCTGTTCTCGTTGAGCTCATCTTAGGACACC | 14 (0.000265%) | | | TTCATAGTCAGAGGTGAAATTCTTGGATTTATGAAAGACGAACAACTGCGAAAGCATTTGCCAAGGATGTTTTCATTAATCAAGAACGAAAGTTGGGGGC | 38 (0.000719%) | | | TTCCAACGAAGCACGCCCATCCAACCTAGGCGAGACAAGGGTTCACATTTCGTTCATCACCCTTGGCCGGCTATCGAACAGCCGGACTCCCATCAAAAGA | 115 (0.002176%) | | | TTCCAACGAAGCACGCCCATCCAACCTAGGCGAGACAAGGGTTCACATTTCGTTCATCACCCTTGGCCGGCTTTCGAACAGCCGGACTCCCATCAAAAGA | 2301 (0.043531%) | | | TTCCAATTACCAGACTCGAAAGAGCCCGGTATTGTTATTTATTGTCACTACCTCCCCGTGTCAGGATTGGGTAATTTGCGCGCCTGCTGCCTTCCTTGGA | 11 (0.000208%) | | | TTCCATCGACCAGAGGCTGTTCACCTTGGAGACCTGATGCGGTTATGAGTACGACCGGGCGTGAGCGGCACTCGGTCCTCCGGATTTTCAAGGGCCGCCG | 4 (0.000076%) | | | TTCCCCGCCACATCCTCTCAAACGCAATGGAAAGAGAGAAAGGACGAGGTCTTGACCGTCATCTTTTGCCCGAAGGACGGATGAGCTTTGGCGGGACTGA | 8 (0.000151%) | | | TTCCCTTGCCTACATTGTTCCATCGACCAGAGGCTGTTCACCTTGGAGACCTGATGCGGTTATGAGTACGACCGGGCGTGAGCGGCACTCGGTCCTCCGG | 10 (0.000189%) | | | TTCCTCCGCTTATTGATATGCTTAAACTCAGCGGGTAATCCCGCCTGACCTGGGGTCGCTATATGGACTTTGGGTCATCTACAGCTTCCGGACAAGAGCG | 4 (0.000076%) | | | TTCCTTGTTAGAAGACACAAAGCCAAAGACTCATATGGACTTTGGCTACACCATGAAAGCTTTGAGAAGCAAGAAGAAGGTTGGTTAGTGTTTTGGAGTC | 58 (0.001097%) | | | TTCGACTCCAAAACACTAACCAACCTTCTTCTTGCTTCTCAAAGCTTTCATGGTGTAGCCAAAGTCCATATGAGTCTTTGGCTTTGTGTCTTCTAACAAG | 130 (0.002459%) | | | TTCGCCGCCCGCGACGTCGCGAGAAGTCCACTAAACCTTATCATTTAGAGGAAGGAGAAGTCGTAACAAGGTTTCCGTAGGTGAACCTGCGGAAGGATCG | 7 (0.000132%) | | | TTCGTACTGAAAATCAGAATCAAACGAGCTTTTACCCTTTTGTTCCACACGAGATTTCTGTTCTCGTTGAGCTCATCTTAGGACACCTGCGTTATCTTTT | 25 (0.000473%) | | | TTCGTATTCGTACTGAAAATCAGAATCAAACGAGCTTTTACCCTTTTGTTCCACACGAGATTTCTGTTCTCGTTGAGCTCATCTTAGGACACCTGCGTTA | 15 (0.000284%) | | | TTCGTATTTCATAGTCAGAGGTGAAATTCTTGGATTTATGAAAGACGAACAACTGCGAAAGCATTTGCCAAGGATGTTTTCATTAATCAAGAACGAAAGT | 11 (0.000208%) | | | TTCTCAAAGCTTTCATGGTGTAGCCAAAGTCCATATGAGTCTTTGGCTTTGTGTCTTCTAACAAGGAAACACTACTTAGGCTTATAAGATGCGGTTGCGG | 13 (0.000246%) | | | TTCTTATACTCAATCATACACATGACATCAAGTCATATTCGACTCCAAAACACTAACCAACCTTCTTCTTGCTTCTCAAAGCTTTCATGGTGTAGCCAAA | 96 (0.001816%) | | | TTCTTGCTTCTCAAAGCTTTCATGGTGTAGCCAAAGTCCATATGAGTCTTTGGCTTTGTGTCTTCTAACAAGGAAACACTACTTAGGCTTATAAGATGCG | 347 (0.006565%) | | | TTCTTTTCCTCCGCTTATTGATATGCTTAAACTCAGCGGGTAATCCCGCCTGACCTGGGGTCGCTATATGGACTTTGGGTCATCTACAGCTTCCGGACAA | 37 (0.000700%) | | | TTGACTCTAGTCCGACTTTGTGAAATGACTTGAGAGGTGTAGGATAAGTGGGAGCTTCGGCGCAAGTGAAATACCACTACTTTTAACGTTATTTTACTTA | 16 (0.000303%) | | | TTGAGCTTGACTCTAGTCCGACTTTGTGAAATGACTTGAGAGGTGTAGGATAAGTGGGAGCTTCGGCGCAAGTGAAATACCACTACTTTTAACGTTATTT | 32 (0.000605%) | | | TTGATCGATC | 182040 (0.344387%) | | | TTGATGGTAACTACTACTCGGATAACCGTAGTAATTCTAGAGCTAATACGTGCAACAAACCCCGACTTATGGAAGGGACGCATTTATTAGATAAAAGGTC | 27 (0.000511%) | | | TTGATGTCATGTGTATGATTGAGTATAAGAACTTAAACCG | 399 (0.003019%) | | | TTGCCGACTTCCCTTGCCTACATTGTTCCATCGACCAGAGGCTGTTCACCTTGGAGACCTGATGCGGTTATGAGTACGACCGGGCGTGAGCGGCACTCGG | 145 (0.002743%) | | | TTGCCTACATTGTTCCATCGACCAGAGGCTGTTCACCTTGGAGACCTGATGCGGTTATGAGTACGACCGGGCGTGAGCGGCACTCGGTCCTCCGGATTTT | 4 (0.000076%) | | | TTGCGGTTTAAGTTCTTATACTCAATCATACACATGACAT | 2081 (0.015748%) | | | TTGCGGTTTAAGTTGTTATACTCAATCATACACATGACAT | 664 (0.005025%) | | | TTGCTCCGTTCCGCATCCGACCAGGACGCATCGCCGGCCCCCATCCGCTTCCCTCCCGACAATTTCAAGCACTCTTTGACTCTCTTTTCAAAGTCCTTTT | 8 (0.000151%) | | | TTGCTGATGCGGGACGGAAGCTGGTCTCCCGTGTGTTACCGCACGCGGTTGGCCTAAATCCGAGCCAAGGACGCCTGGAGCGTACCGACATGCGGTGGTG | 5 (0.000095%) | | | TTGCTTTGAGCACTCTAATTTCTTCAAAGTAACAGCGCCGGAGGCACGACCCGGCCAATTAAGACCAGGAGCGTATCGCCGACCGAAGGGACAAGCCGAC | 5 (0.000095%) | | | TTGGAGGGCAAGTCTGGTGCCAGCAGCCGCGGTAATTCCAGCTCCAATAGCGTATATTTAAGTTGTTGCAGTTAAAAAGCTCGTAGTTGAACCTTGGGAT | 347 (0.006565%) | | | TTGGCATGCATCATAAGGATACTAAATCCTATTTTCTGGTAAATTTTCATAATTTTTTGACACCTCTAGCTAGGTCATTTGACCTGATACAACATCGGAT | 387 (0.007321%) | | | TTGGCTGGGGCGGCACATCTGTTAAAAGATAACGCAGGTGTCCTAAGATGAGCTCAACGAGAACAGAAATCTCGTGTGGAACAAAAGGGTAAAAGCTCGT | 9 (0.000170%) | | | TTGGTCTTCAACGAGGAATTCCTAGTAAGCGCGAGTCATCAGCTCGCGTTGACTACGTCCCTGCCCTTTGTACACACCGCCCGTCGCTCCTACCGATTGA | 111 (0.002100%) | | | TTGTACTCATTCCAATTACCAGACTCGAAAGAGCCCGGTATTGTTATTTATTGTCACTACCTCCCCGTGTCAGGATTGGGTAATTTGCGCGCCTGCTGCC | 6 (0.000114%) | | | TTGTATCCTTGTTAGAAGACACAAAGCCAAAGACTCATATGGACTTTGGCTACACCATGAAAGCTTTGAGAAGCAAGAAGAAGGTTGGTTAGTGTTTTGG | 16 (0.000303%) | | | TTGTCAGGTGGGGAGTTTGGCTGGGGCGGCACATCTGTTAAAAGATAACGCAGGTGTCCTAAGATGAGCTCAACGAGAACAGAAATCTCGTGTGGAACAA | 8 (0.000151%) | | | TTGTCTCAAAGATTAAGCCATGCATGTGTAAGTATGAACGAATTCAGACTGTGAAACTGCGAATGGCTCATTAAATCAGTTATAGTTTGTTTGATGGTAA | 6 (0.000114%) | | | TTGTTAGAAGACACAAAGCCAAAGACTCATATGGACTTTGGCTACACCATGAAAGCTTTGAGAAGCAAGAAGAAGGTTGGTTAGTGTTTTGGAGTCGAAT | 71 (0.001343%) | | | TTGTTAGTTTCTTTTCCTCCGCTTATTGATATGCTTAAACTCAGCGGGTAATCCCGCCTGACCTGGGGTCGCTATATGGACTTTGGGTCATCTACAGCTT | 10 (0.000189%) | | | TTGTTCCATCGACCAGAGGCTGTTCACCTTGGAGACCTGATGCGGTTATGAGTACGACCGGGCGTGAGCGGCACTCGGTCCTCCGGATTTTCAAGGGCCG | 4 (0.000076%) | | | TTGTTGGTCTTCAACGAGGAATTCCTAGTAAGCGCGAGTCATCAGCTCGCGTTGACTACGTCCCTGCCCTTTGTACACACCGCCCGTCGCTCCTACCGAT | 211 (0.003992%) | | | TTGTTTGATGGTAACTACTACTCGGATAACCGTAGTAATTCTAGAGCTAATACGTGCAACAAACCCCGACTTATGGAAGGGACGCATTTATTAGATAAAA | 12 (0.000227%) | | | TTTAAGTTCTTATACTCAATCATACACATGACATCAAGTC | 812 (0.006145%) | | | TTTAAGTTGTTATACTCAATCATACACATGACAACAAGTCATATTCGACTCCAAAACACTAACCAACCTTCTTCTTGCTTCTCAAAGCTTTCATGGTGTA | 30 (0.000568%) | | | TTTAAGTTGTTATACTCAATCATACACATGACATCAAGTC | 379 (0.002868%) | | | TTTACTCACCCGTTGACTCGCACACATGTCAGACTCCTTGGTCCGTGTTTCAAGACGGGTCGAATGGGGAGCCCACAGGCCGACGCCCGGAGCACGCTGA | 9 (0.000170%) | | | TTTAGATTGTACTCATTCCAATTACCAGACTCGAAAGAGCCCGGTATTGTTATTTATTGTCACTACCTCCCCGTGTCAGGATTGGGTAATTTGCGCGCCT | 145 (0.002743%) | | | TTTATCTAATAAATGCGTCCCTTCCATAAGTCGGGGTTTGTTGCACGTATTAGCTCTAGAATTACTACGGTTATCCGAGTAGTAGTTACCATCAAACAAA | 7 (0.000132%) | | | TTTCACGGTTCGTATTCGTACTGAAAATCAGAATCAAACGAGCTTTTACCCTTTTGTTCCACACGAGATTTCTGTTCTCGTTGAGCTCATCTTAGGACAC | 57 (0.001078%) | | | TTTCATAATTTTTTGACACCTCTAGCTAGGTCATTTGACCTGATACAACATCGGATTTTCATGGTCTAGTTGGGGCTCCGTGGGCATATTTGATGCAAAC | 9 (0.000170%) | | | TTTCATAGTCAGAGGTGAAATTCTTGGATTTATGAAAGACGAACAACTGCGAAAGCATTTGCCAAGGATGTTTTCATTAATCAAGAACGAAAGTTGGGGG | 9 (0.000170%) | | | TTTCCAACGAAGCACGCCCATCCAACCTAGGCGAGACAAGGGTTCACATTTCGTTCATCACCCTTGGCCGGCTATCGAACAGCCGGACTCCCATCAAAAG | 270 (0.005108%) | | | TTTCCTCCGCTTATTGATATGCTTAAACTCAGCGGGTAATCCCGCCTGACCTGGGGTCGCTATATGGACTTTGGGTCATCTACAGCTTCCGGACAAGAGC | 11 (0.000208%) | | | TTTCCTTGTTAGAAGACACAAAGCCAAAGACTCATATGGACTTTGGCTACACCATGAAAGCTTTGAGAAGCAAGAAGAAGGTTGGTTAGTGTTTTGGAGT | 631 (0.011937%) | | | TTTCGTTCATCACCCTTGGCCGGCTATCGAACAGCCGGACTCCCATCAAAAGATGGTTGCCAAGAACATCTTCGTTACGGTTTGCTAATTCTCGGAATAA | 10 (0.000189%) | | | TTTCTGGTAAATTTTCATAATTTTTTGACACCTCTAGCTAGGTCATTTGACCTGATACAACATCGGATTTTCATGGTCTAGTTGGGGCTCCGTGGGCATA | 7 (0.000132%) | | | TTTCTTTTCCTCCGCTTATTGATATGCTTAAACTCAGCGGGTAATCCCGCCTGACCTGGGGTCGCTATATGGACTTTGGGTCATCTACAGCTTCCGGACA | 12 (0.000227%) | | | TTTGATGGTAACTACTACTCGGATAACCGTAGTAATTCTAGAGCTAATACGTGCAACAAACCCCGACTTATGGAAGGGACGCATTTATTAGATAAAAGGT | 3 (0.000057%) | | | TTTGCCGACTTCCCTTGCCTACATTGTTCCATCGACCAGAGGCTGTTCACCTTGGAGACCTGATGCGGTTATGAGTACGACCGGGCGTGAGCGGCACTCG | 2275 (0.043039%) | | | TTTGCTGATGCGGGACGGAAGCTGGTCTCCCGTGTGTTACCGCACGCGGTTGGCCTAAATCCGAGCCAAGGACGCCTGGAGCGTACCGACATGCGGTGGT | 19 (0.000359%) | | | TTTGGCTGGGGCGGCACATCTGTTAAAAGATAACGCAGGTGTCCTAAGATGAGCTCAACGAGAACAGAAATCTCGTGTGGAACAAAAGGGTAAAAGCTCG | 21 (0.000397%) | | | TTTGTTTGATGGTAACTACTACTCGGATAACCGTAGTAATTCTAGAGCTAATACGTGCAACAAACCCCGACTTATGGAAGGGACGCATTTATTAGATAAA | 7 (0.000132%) | | | TTTTATCTAATAAATGCGTCCCTTCCATAAGTCGGGGTTTGTTGCACGTATTAGCTCTAGAATTACTACGGTTATCCGAGTAGTAGTTACCATCAAACAA | 10 (0.000189%) | | | TTTTCATAATTTTTTGACACCTCTAGCTAGGTCATTTGACCTGATACAACATCGGATTTTCATGGTCTAGTTGGGGCTCCGTGGGCATATTTGATGCAAA | 6 (0.000114%) | | | TTTTCCTCCGCTTATTGATATGCTTAAACTCAGCGGGTAATCCCGCCTGACCTGGGGTCGCTATATGGACTTTGGGTCATCTACAGCTTCCGGACAAGAG | 22 (0.000416%) | | | TTTTCTGGTAAATTTTCATAATTTTTTGACACCTCTAGCTAGGTCATTTGACCTGATACAACATCGGATTTTCATGGTCTAGTTGGGGCTCCGTGGGCAT | 9 (0.000170%) | | | TTTTTTTTTT | 55119 (0.104275%) | | Before filtering: read2: quality Value of each position will be shown on mouse over. Before filtering: read2: base contents Value of each position will be shown on mouse over. Before filtering: read2: KMER counting Darker background means larger counts. The count will be shown on mouse over. | | | | | | | | | | | | | | | | | | | --- | --- | --- | --- | --- | --- | --- | --- | --- | --- | --- | --- | --- | --- | --- | --- | --- | | | AA | AT | AC | AG | TA | TT | TC | TG | CA | CT | CC | CG | GA | GT | GC | GG | | AAA | AAAAA | AAAAT | AAAAC | AAAAG | AAATA | AAATT | AAATC | AAATG | AAACA | AAACT | AAACC | AAACG | AAAGA | AAAGT | AAAGC | AAAGG | | AAT | AATAA | AATAT | AATAC | AATAG | AATTA | AATTT | AATTC | AATTG | AATCA | AATCT | AATCC | AATCG | AATGA | AATGT | AATGC | AATGG | | AAC | AACAA | AACAT | AACAC | AACAG | AACTA | AACTT | AACTC | AACTG | AACCA | AACCT | AACCC | AACCG | AACGA | AACGT | AACGC | AACGG | | AAG | AAGAA | AAGAT | AAGAC | AAGAG | AAGTA | AAGTT | AAGTC | AAGTG | AAGCA | AAGCT | AAGCC | AAGCG | AAGGA | AAGGT | AAGGC | AAGGG | | ATA | ATAAA | ATAAT | ATAAC | ATAAG | ATATA | ATATT | ATATC | ATATG | ATACA | ATACT | ATACC | ATACG | ATAGA | ATAGT | ATAGC | ATAGG | | ATT | ATTAA | ATTAT | ATTAC | ATTAG | ATTTA | ATTTT | ATTTC | ATTTG | ATTCA | ATTCT | ATTCC | ATTCG | ATTGA | ATTGT | ATTGC | ATTGG | | ATC | ATCAA | ATCAT | ATCAC | ATCAG | ATCTA | ATCTT | ATCTC | ATCTG | ATCCA | ATCCT | ATCCC | ATCCG | ATCGA | ATCGT | ATCGC | ATCGG | | ATG | ATGAA | ATGAT | ATGAC | ATGAG | ATGTA | ATGTT | ATGTC | ATGTG | ATGCA | ATGCT | ATGCC | ATGCG | ATGGA | ATGGT | ATGGC | ATGGG | | ACA | ACAAA | ACAAT | ACAAC | ACAAG | ACATA | ACATT | ACATC | ACATG | ACACA | ACACT | ACACC | ACACG | ACAGA | ACAGT | ACAGC | ACAGG | | ACT | ACTAA | ACTAT | ACTAC | ACTAG | ACTTA | ACTTT | ACTTC | ACTTG | ACTCA | ACTCT | ACTCC | ACTCG | ACTGA | ACTGT | ACTGC | ACTGG | | ACC | ACCAA | ACCAT | ACCAC | ACCAG | ACCTA | ACCTT | ACCTC | ACCTG | ACCCA | ACCCT | ACCCC | ACCCG | ACCGA | ACCGT | ACCGC | ACCGG | | ACG | ACGAA | ACGAT | ACGAC | ACGAG | ACGTA | ACGTT | ACGTC | ACGTG | ACGCA | ACGCT | ACGCC | ACGCG | ACGGA | ACGGT | ACGGC | ACGGG | | AGA | AGAAA | AGAAT | AGAAC | AGAAG | AGATA | AGATT | AGATC | AGATG | AGACA | AGACT | AGACC | AGACG | AGAGA | AGAGT | AGAGC | AGAGG | | AGT | AGTAA | AGTAT | AGTAC | AGTAG | AGTTA | AGTTT | AGTTC | AGTTG | AGTCA | AGTCT | AGTCC | AGTCG | AGTGA | AGTGT | AGTGC | AGTGG | | AGC | AGCAA | AGCAT | AGCAC | AGCAG | AGCTA | AGCTT | AGCTC | AGCTG | AGCCA | AGCCT | AGCCC | AGCCG | AGCGA | AGCGT | AGCGC | AGCGG | | AGG | AGGAA | AGGAT | AGGAC | AGGAG | AGGTA | AGGTT | AGGTC | AGGTG | AGGCA | AGGCT | AGGCC | AGGCG | AGGGA | AGGGT | AGGGC | AGGGG | | TAA | TAAAA | TAAAT | TAAAC | TAAAG | TAATA | TAATT | TAATC | TAATG | TAACA | TAACT | TAACC | TAACG | TAAGA | TAAGT | TAAGC | TAAGG | | TAT | TATAA | TATAT | TATAC | TATAG | TATTA | TATTT | TATTC | TATTG | TATCA | TATCT | TATCC | TATCG | TATGA | TATGT | TATGC | TATGG | | TAC | TACAA | TACAT | TACAC | TACAG | TACTA | TACTT | TACTC | TACTG | TACCA | TACCT | TACCC | TACCG | TACGA | TACGT | TACGC | TACGG | | TAG | TAGAA | TAGAT | TAGAC | TAGAG | TAGTA | TAGTT | TAGTC | TAGTG | TAGCA | TAGCT | TAGCC | TAGCG | TAGGA | TAGGT | TAGGC | TAGGG | | TTA | TTAAA | TTAAT | TTAAC | TTAAG | TTATA | TTATT | TTATC | TTATG | TTACA | TTACT | TTACC | TTACG | TTAGA | TTAGT | TTAGC | TTAGG | | TTT | TTTAA | TTTAT | TTTAC | TTTAG | TTTTA | TTTTT | TTTTC | TTTTG | TTTCA | TTTCT | TTTCC | TTTCG | TTTGA | TTTGT | TTTGC | TTTGG | | TTC | TTCAA | TTCAT | TTCAC | TTCAG | TTCTA | TTCTT | TTCTC | TTCTG | TTCCA | TTCCT | TTCCC | TTCCG | TTCGA | TTCGT | TTCGC | TTCGG | | TTG | TTGAA | TTGAT | TTGAC | TTGAG | TTGTA | TTGTT | TTGTC | TTGTG | TTGCA | TTGCT | TTGCC | TTGCG | TTGGA | TTGGT | TTGGC | TTGGG | | TCA | TCAAA | TCAAT | TCAAC | TCAAG | TCATA | TCATT | TCATC | TCATG | TCACA | TCACT | TCACC | TCACG | TCAGA | TCAGT | TCAGC | TCAGG | | TCT | TCTAA | TCTAT | TCTAC | TCTAG | TCTTA | TCTTT | TCTTC | TCTTG | TCTCA | TCTCT | TCTCC | TCTCG | TCTGA | TCTGT | TCTGC | TCTGG | | TCC | TCCAA | TCCAT | TCCAC | TCCAG | TCCTA | TCCTT | TCCTC | TCCTG | TCCCA | TCCCT | TCCCC | TCCCG | TCCGA | TCCGT | TCCGC | TCCGG | | TCG | TCGAA | TCGAT | TCGAC | TCGAG | TCGTA | TCGTT | TCGTC | TCGTG | TCGCA | TCGCT | TCGCC | TCGCG | TCGGA | TCGGT | TCGGC | TCGGG | | TGA | TGAAA | TGAAT | TGAAC | TGAAG | TGATA | TGATT | TGATC | TGATG | TGACA | TGACT | TGACC | TGACG | TGAGA | TGAGT | TGAGC | TGAGG | | TGT | TGTAA | TGTAT | TGTAC | TGTAG | TGTTA | TGTTT | TGTTC | TGTTG | TGTCA | TGTCT | TGTCC | TGTCG | TGTGA | TGTGT | TGTGC | TGTGG | | TGC | TGCAA | TGCAT | TGCAC | TGCAG | TGCTA | TGCTT | TGCTC | TGCTG | TGCCA | TGCCT | TGCCC | TGCCG | TGCGA | TGCGT | TGCGC | TGCGG | | TGG | TGGAA | TGGAT | TGGAC | TGGAG | TGGTA | TGGTT | TGGTC | TGGTG | TGGCA | TGGCT | TGGCC | TGGCG | TGGGA | TGGGT | TGGGC | TGGGG | | CAA | CAAAA | CAAAT | CAAAC | CAAAG | CAATA | CAATT | CAATC | CAATG | CAACA | CAACT | CAACC | CAACG | CAAGA | CAAGT | CAAGC | CAAGG | | CAT | CATAA | CATAT | CATAC | CATAG | CATTA | CATTT | CATTC | CATTG | CATCA | CATCT | CATCC | CATCG | CATGA | CATGT | CATGC | CATGG | | CAC | CACAA | CACAT | CACAC | CACAG | CACTA | CACTT | CACTC | CACTG | CACCA | CACCT | CACCC | CACCG | CACGA | CACGT | CACGC | CACGG | | CAG | CAGAA | CAGAT | CAGAC | CAGAG | CAGTA | CAGTT | CAGTC | CAGTG | CAGCA | CAGCT | CAGCC | CAGCG | CAGGA | CAGGT | CAGGC | CAGGG | | CTA | CTAAA | CTAAT | CTAAC | CTAAG | CTATA | CTATT | CTATC | CTATG | CTACA | CTACT | CTACC | CTACG | CTAGA | CTAGT | CTAGC | CTAGG | | CTT | CTTAA | CTTAT | CTTAC | CTTAG | CTTTA | CTTTT | CTTTC | CTTTG | CTTCA | CTTCT | CTTCC | CTTCG | CTTGA | CTTGT | CTTGC | CTTGG | | CTC | CTCAA | CTCAT | CTCAC | CTCAG | CTCTA | CTCTT | CTCTC | CTCTG | CTCCA | CTCCT | CTCCC | CTCCG | CTCGA | CTCGT | CTCGC | CTCGG | | CTG | CTGAA | CTGAT | CTGAC | CTGAG | CTGTA | CTGTT | CTGTC | CTGTG | CTGCA | CTGCT | CTGCC | CTGCG | CTGGA | CTGGT | CTGGC | CTGGG | | CCA | CCAAA | CCAAT | CCAAC | CCAAG | CCATA | CCATT | CCATC | CCATG | CCACA | CCACT | CCACC | CCACG | CCAGA | CCAGT | CCAGC | CCAGG | | CCT | CCTAA | CCTAT | CCTAC | CCTAG | CCTTA | CCTTT | CCTTC | CCTTG | CCTCA | CCTCT | CCTCC | CCTCG | CCTGA | CCTGT | CCTGC | CCTGG | | CCC | CCCAA | CCCAT | CCCAC | CCCAG | CCCTA | CCCTT | CCCTC | CCCTG | CCCCA | CCCCT | CCCCC | CCCCG | CCCGA | CCCGT | CCCGC | CCCGG | | CCG | CCGAA | CCGAT | CCGAC | CCGAG | CCGTA | CCGTT | CCGTC | CCGTG | CCGCA | CCGCT | CCGCC | CCGCG | CCGGA | CCGGT | CCGGC | CCGGG | | CGA | CGAAA | CGAAT | CGAAC | CGAAG | CGATA | CGATT | CGATC | CGATG | CGACA | CGACT | CGACC | CGACG | CGAGA | CGAGT | CGAGC | CGAGG | | CGT | CGTAA | CGTAT | CGTAC | CGTAG | CGTTA | CGTTT | CGTTC | CGTTG | CGTCA | CGTCT | CGTCC | CGTCG | CGTGA | CGTGT | CGTGC | CGTGG | | CGC | CGCAA | CGCAT | CGCAC | CGCAG | CGCTA | CGCTT | CGCTC | CGCTG | CGCCA | CGCCT | CGCCC | CGCCG | CGCGA | CGCGT | CGCGC | CGCGG | | CGG | CGGAA | CGGAT | CGGAC | CGGAG | CGGTA | CGGTT | CGGTC | CGGTG | CGGCA | CGGCT | CGGCC | CGGCG | CGGGA | CGGGT | CGGGC | CGGGG | | GAA | GAAAA | GAAAT | GAAAC | GAAAG | GAATA | GAATT | GAATC | GAATG | GAACA | GAACT | GAACC | GAACG | GAAGA | GAAGT | GAAGC | GAAGG | | GAT | GATAA | GATAT | GATAC | GATAG | GATTA | GATTT | GATTC | GATTG | GATCA | GATCT | GATCC | GATCG | GATGA | GATGT | GATGC | GATGG | | GAC | GACAA | GACAT | GACAC | GACAG | GACTA | GACTT | GACTC | GACTG | GACCA | GACCT | GACCC | GACCG | GACGA | GACGT | GACGC | GACGG | | GAG | GAGAA | GAGAT | GAGAC | GAGAG | GAGTA | GAGTT | GAGTC | GAGTG | GAGCA | GAGCT | GAGCC | GAGCG | GAGGA | GAGGT | GAGGC | GAGGG | | GTA | GTAAA | GTAAT | GTAAC | GTAAG | GTATA | GTATT | GTATC | GTATG | GTACA | GTACT | GTACC | GTACG | GTAGA | GTAGT | GTAGC | GTAGG | | GTT | GTTAA | GTTAT | GTTAC | GTTAG | GTTTA | GTTTT | GTTTC | GTTTG | GTTCA | GTTCT | GTTCC | GTTCG | GTTGA | GTTGT | GTTGC | GTTGG | | GTC | GTCAA | GTCAT | GTCAC | GTCAG | GTCTA | GTCTT | GTCTC | GTCTG | GTCCA | GTCCT | GTCCC | GTCCG | GTCGA | GTCGT | GTCGC | GTCGG | | GTG | GTGAA | GTGAT | GTGAC | GTGAG | GTGTA | GTGTT | GTGTC | GTGTG | GTGCA | GTGCT | GTGCC | GTGCG | GTGGA | GTGGT | GTGGC | GTGGG | | GCA | GCAAA | GCAAT | GCAAC | GCAAG | GCATA | GCATT | GCATC | GCATG | GCACA | GCACT | GCACC | GCACG | GCAGA | GCAGT | GCAGC | GCAGG | | GCT | GCTAA | GCTAT | GCTAC | GCTAG | GCTTA | GCTTT | GCTTC | GCTTG | GCTCA | GCTCT | GCTCC | GCTCG | GCTGA | GCTGT | GCTGC | GCTGG | | GCC | GCCAA | GCCAT | GCCAC | GCCAG | GCCTA | GCCTT | GCCTC | GCCTG | GCCCA | GCCCT | GCCCC | GCCCG | GCCGA | GCCGT | GCCGC | GCCGG | | GCG | GCGAA | GCGAT | GCGAC | GCGAG | GCGTA | GCGTT | GCGTC | GCGTG | GCGCA | GCGCT | GCGCC | GCGCG | GCGGA | GCGGT | GCGGC | GCGGG | | GGA | GGAAA | GGAAT | GGAAC | GGAAG | GGATA | GGATT | GGATC | GGATG | GGACA | GGACT | GGACC | GGACG | GGAGA | GGAGT | GGAGC | GGAGG | | GGT | GGTAA | GGTAT | GGTAC | GGTAG | GGTTA | GGTTT | GGTTC | GGTTG | GGTCA | GGTCT | GGTCC | GGTCG | GGTGA | GGTGT | GGTGC | GGTGG | | GGC | GGCAA | GGCAT | GGCAC | GGCAG | GGCTA | GGCTT | GGCTC | GGCTG | GGCCA | GGCCT | GGCCC | GGCCG | GGCGA | GGCGT | GGCGC | GGCGG | | GGG | GGGAA | GGGAT | GGGAC | GGGAG | GGGTA | GGGTT | GGGTC | GGGTG | GGGCA | GGGCT | GGGCC | GGGCG | GGGGA | GGGGT | GGGGC | GGGGG | Before filtering: read2: overrepresented sequences Sampling rate: 1 / 20 | | | | | --- | --- | --- | | overrepresented sequence | count (% of bases) | distribution: cycle 1 ~ cycle 150 | | AAAAAAAAAAAAAAAAAAAA | 6570 (0.024859%) | | | AAAAGATAACGCAGGTGTCCTAAGATGAGCTCAACGAGAACAGAAATCTCGTGTGGAACAAAAGGGTAAAAGCTCGTTTGATTCTGATTTTCAGTACGAA | 7 (0.000132%) | | | AAAAGTGTCGGAGTTTTTTCAGCAGTTCTCGGACAAAAATTGCTGAGTGGCCGAGAAGAATGGGCGTGTCATGCGTGGGCTGACATGGATTCTTCGAGGC | 5 (0.000095%) | | | AAAATCATCAATCGTTCCAACTAATCTACCGAAGTACTCGGCTAAGAAGAAAGAAGACGGACGAATCCGAGCCAAAGCCGTACAAGCGCGAGATACCTTC | 22 (0.000416%) | | | AAACACAGGCCCCGGAACTCATCATCGAGCGTAACATCGCCCGTGAATTAACTGAGAAGGATAGGTGGTAGGTAGTTCGATGCGCGAGCATGGAGCCTAC | 11 (0.000208%) | | | AAACACTTGGTGATATGAACACAAACGTTCAATATGACAAACCCATGCCAAGTAAAGAGAAAATGAAAACTGGTGATTGTTGCGGAAATCGTCCAGGATT | 293 (0.005543%) | | | AAACGTTCAATATGACAAACCCATGCCAAGTAAAGAGAAAATGAAAACTGGTGATTGTTGCGGAAATCGTCCAGGATTCCTCGACCAGGACTTGAAATCG | 4 (0.000076%) | | | AAAGCCAAAGACTCATATGGACTTTGGCTACACCATGAAAGCTTTGAGAAGCAAGAAGAAGGTTGGTTAGTGTTTTGGAGTCGAATATGACTTGATGTCA | 279 (0.005278%) | | | AAAGGCGTAAGAATTGTATCCTTGTTAGAAGACACAAAGC | 199 (0.001506%) | | | AAAGGGTGTTGGTCGATTAAGACAGCAGGACGGTGGTCATGGAAGTCGAAATCCGCTAAGGAGTGTGTAACAACTCACCTGCCGAATCAACTAGCCCCGA | 16 (0.000303%) | | | AAAGTTGGGAATTCGTTAAGGAGCTGTTGCTTTGTTAGTGTAGAAACACTTGTGTAGAATTGGGGATTGTTTTTTTTGGAGTGATTTAGGGGAGGGTCGA | 11 (0.000208%) | | | AAATACGGGCGAGAGACCGATAGCGAACAAGTACCGCGAGGTAAAGATGAAAAGGACTTTGAAAAGAGAGTCAAAGAGTGCTTGAAATTGTCGGGAGGGA | 6 (0.000114%) | | | AAATCATCAATCGTTCCAACTAATCTACCGAAGTACTCGGCTAAGAAGAAAGAAGACGGACGAATCCGAGCCAAAGCCGTACAAGCGCGAGATACCTTCG | 9 (0.000170%) | | | AAATCCTATGATGTTATCCCATGCTAATGTATCCAGAGCGTAGGCTTGCTTTGAGCACTCTAATTTCTTCAAAGTAACAGCGCCGGAGGCACGACCCGGC | 35 (0.000662%) | | | AAATCGTCGTCCCTCACCATCCTTTGCTGATGCGGGACGGAAGCTGGTCTCCCGTGTGTTACCGCACGCGGTTGGCCTAAATCCGAGCCAAGGACGCCTG | 28 (0.000530%) | | | AAATTAGGTACGAAACACAGGCCCCGGAACTCATCATCGAGCGTAACATCGCCCGTGAATTAACTGAGAAGGATAGGTGGTAGGTAGTTCGATGCGCGAG | 10 (0.000189%) | | | AAATTTCTGCCCTATCAACTTTCGATGGTAGGATAGTGGCCTACCATGGTGGTAACGGGTGACGGAGAATTAGGGTTCGATTCCGGAGAGGGAGCCTGAG | 37 (0.000700%) | | | AACACAGGCCCCGGAACTCATCATCGAGCGTAACATCGCCCGTGAATTAACTGAGAAGGATAGGTGGTAGGTAGTTCGATGCGCGAGCATGGAGCCTACG | 23 (0.000435%) | | | AACAGGGACAGTCGGGGGCATTCGTATTTCATAGTCAGAGGTGAAATTCTTGGATTTATGAAAGACGAACAACTGCGAAAGCATTTGCCAAGGATGTTTT | 11 (0.000208%) | | | AACCCACAAAGGGTGTTGGTCGATTAAGACAGCAGGACGGTGGTCATGGAAGTCGAAATCCGCTAAGGAGTGTGTAACAACTCACCTGCCGAATCAACTA | 4 (0.000076%) | | | AACCGGGACGTGGCGGTTGACGGCAACGTTAGGGAGTCCGGAGACGTCGGCGGGGGCCTCGGGAAGAGTTATCTTTTCTGTTTAACAGCCTGCCCACCCT | 3 (0.000057%) | | | AACCTAGGCGAGACAAGGGTTCACATTTCGTTCATCACCCTTGGCCGGCTATCGAACAGCCGGACTCCCATCAAAAGATGGTTGCCAAGAACATCTTCGT | 3 (0.000057%) | | | AACGAAGCACGCCCATCCAACCTAGGCGAGACAAGGGTTCACATTTCGTTCATCACCCTTGGCCGGCTATCGAACAGCCGGACTCCCATCAAAAGATGGT | 21 (0.000397%) | | | AACGGCGTGCCTCGGCATCAGCGTGCTCCGGGCGTCGGCCTGTGGGCTCCCCATTCGACCCGTCTTGAAACACGGACCAAGGAGTCTGACATGTGTGCGA | 7 (0.000132%) | | | AACGGGCAGAGCCCGCGTCGACCTTTTATCTAATAAATGCGTCCCTTCCATAAGTCGGGGTTTGTTGCACGTATTAGCTCTAGAATTACTACGGTTATCC | 8 (0.000151%) | | | AACGTTAGGGAGTCCGGAGACGTCGGCGGGGGCCTCGGGAAGAGTTATCTTTTCTGTTTAACAGCCTGCCCACCCTGGAAACGGCTCAGCCGGAGGTAGG | 3 (0.000057%) | | | AACTAGCTACGTGGAGGCATCCCTTCACGGCCGGCTTCTTAGAGGGACTATGGCCGTTTAGGCCAAGGAAGTTTGAGGCAATAACAGGTCTGTGATGCCC | 4 (0.000076%) | | | AACTTTCGATGGTAGGATAGTGGCCTACCATGGTGGTAACGGGTGACGGAGAATTAGGGTTCGATTCCGGAGAGGGAGCCTGAGAAACGGCTACCACATC | 4 (0.000076%) | | | AAGAATTGTATCCTTGTTAGAAGACACAAAGCCAAAGACTCATATGGACTTTGGCTACACCATGAAAGCTTTGAGAAGCAAGAAGAAGGTTGGTTAGTGT | 17 (0.000322%) | | | AAGACACAAAGCCAAAGACTCATATGGACTTTGGCTACACCATGAAAGCTTTGAGAAGCAAGAAGAAGGTTGGTTAGTGTTTTGGAGTCGAATATGACTT | 528 (0.009989%) | | | AAGACAGCAGGACGGTGGTCATGGAAGTCGAAATCCGCTAAGGAGTGTGTAACAACTCACCTGCCGAATCAACTAGCCCCGAAAATGGATGGCGCTTAAG | 14 (0.000265%) | | | AAGAGCGTCGTGTAGGGAAAGAGTGTTCAGAGCCGTGTAG | 11 (0.000083%) | | | AAGATCGATC | 204027 (0.385983%) | | | AAGCACGCCCATCCAACCTAGGCGAGACAAGGGTTCACATTTCGTTCATCACCCTTGGCCGGCTATCGAACAGCCGGACTCCCATCAAAAGATGGTTGCC | 5 (0.000095%) | | | AAGCCAAAGACTCATATGGACTTTGGCTACACCATGAAAGCTTTGAGAAGCAAGAAGAAGGTTGGTTAGTGTTTTGGAGTCGAATATGACTTGATGTCAT | 9 (0.000170%) | | | AAGCCTAAGTAGTGTTTCCTTGTTAGAACATACAAAGCCAAAGACTCATACGGACTTTGGCTACACCATGAAAGCTTTGAGAAGCTAGAAGAAGGTTGGT | 1229 (0.023251%) | | | AAGCCTAAGTAGTGTTTCCTTGTTAGAAGACACAAAGCCAAAGACTCATATGGACTTTGGCTACACCATGAAAGCTTTGAGAAGCAAGAAGAAGGTTGGT | 1683 (0.031839%) | | | AAGGAGCTGTTGCTTTGTTAGTGTAGAAACACTTGTGTAGAATTGGGGATTGTTTTTTTTGGAGTGATTTAGGGGAGGGTCGAATCTTAGCGACAAAGGG | 5 (0.000095%) | | | AAGGCGTAAGAATTGTATCCTTGTTAGAAGACACAAAGCC | 61 (0.000462%) | | | AAGGGTGTTGGTCGATTAAGACAGCAGGACGGTGGTCATGGAAGTCGAAATCCGCTAAGGAGTGTGTAACAACTCACCTGCCGAATCAACTAGCCCCGAA | 3 (0.000057%) | | | AAGGGTTCACATTTCGTTCATCACCCTTGGCCGGCTTTCGAACAGCCGGACTCCCATCAAAAGATGGTTGCCAAGAACATCTTCGTTACGGTTTGCTAAT | 3 (0.000057%) | | | AAGGTATCTCGCGCTTGTACGGCTTTGGCTCGGATTCGTCCGTCTTCTTTCTTCTTAGCCGAGTACTTCGGTAGATTAGTTGGAACGATTGATGATTTTG | 13 (0.000246%) | | | AAGTAGTGTTTCCTTGTTAGAAGACACAAAGCCAAAGACTCATATGGACTTTGGCTACACCATGAAAGCTTTGAGAAGCAAGAAGAAGGTTGGTTAGTGT | 27 (0.000511%) | | | AAGTCATATTCGACTCCAAAACACTAACCAACCTTCTTCTTGCTTCTCAAAGCTTTCATGGTGTAGCCAAAGTCCATATGAGTCTTTGGCTTTGTGTCTT | 128 (0.002422%) | | | AAGTCTGGTGCCAGCAGCCGCGGTAATTCCAGCTCCAATAGCGTATATTTAAGTTGTTGCAGTTAAAAAGCTCGTAGTTGAACCTTGGGATGGGTCGGCC | 6 (0.000114%) | | | AAGTTCTTATACTCAATCATACACATGACATCAAGTCATATTCGACTCCAAAACACTAACCAACCTTCTTCTTGCTTCTCAAAGCTTTCATGGTGTAGCC | 29 (0.000549%) | | | AAGTTGTTATACTCAATCATACACATGACAACAAGTCATATTCGACTCCAAAACACTAACCAACCTTCTTCTTGCTTCTCAAAGCTTTCATGGTGTAGCC | 31 (0.000586%) | | | AAGTTGTTATACTCAATCATACACATGACATCAAGTCATA | 126 (0.000953%) | | | AATACGGGCGAGAGACCGATAGCGAACAAGTACCGCGAGGTAAAGATGAAAAGGACTTTGAAAAGAGAGTCAAAGAGTGCTTGAAATTGTCGGGAGGGAA | 5 (0.000095%) | | | AATCAGCTTCCTTGCGCCTTACGGGTTTACTCACCCGTTGACTCGCACACATGTCAGACTCCTTGGTCCGTGTTTCAAGACGGGTCGAATGGGGAGCCCA | 173 (0.003273%) | | | AATCATACACATGACATCAAGTCATATTCGACTCCAAAACACTAACCAACCTTCTTCTTGCTTCTCAAAGCTTTCATGGTGTAGCCAAAGTCCATATGAG | 53 (0.001003%) | | | AATCATCAATCGTTCCAACTAATCTACCGAAGTACTCGGCTAAGAAGAAAGAAGACGGACGAATCCGAGCCAAAGCCGTACAAGCGCGAGATACCTTCGG | 3 (0.000057%) | | | AATCATCAGAGCAACGGGCAGAGCCCGCGTCGACCTTTTATCTAATAAATGCGTCCCTTCCATAAGTCGGGGTTTGTTGCACGTATTAGCTCTAGAATTA | 44 (0.000832%) | | | AATCCTATGATGTTATCCCATGCTAATGTATCCAGAGCGTAGGCTTGCTTTGAGCACTCTAATTTCTTCAAAGTAACAGCGCCGGAGGCACGACCCGGCC | 39 (0.000738%) | | | AATCGGTAGGAGCGACGGGCGGTGTGTACAAAGGGCAGGGACGTAGTCAACGCGAGCTGATGACTCGCGCTTACTAGGAATTCCTCGTTGAAGACCAACA | 157 (0.002970%) | | | AATCGTCGTCCCTCACCATCCTTTGCTGATGCGGGACGGAAGCTGGTCTCCCGTGTGTTACCGCACGCGGTTGGCCTAAATCCGAGCCAAGGACGCCTGG | 7 (0.000132%) | | | AATCGTTCCAACTAATCTACCGAAGTACTCGGCTAAGAAGAAAGAAGACGGACGAATCCGAGCCAAAGCCGTACAAGCGCGAGATACCTTCGGGACAGCC | 14 (0.000265%) | | | AATGATTAACAGGGACAGTCGGGGGCATTCGTATTTCATAGTCAGAGGTGAAATTCTTGGATTTATGAAAGACGAACAACTGCGAAAGCATTTGCCAAGG | 571 (0.010802%) | | | AATGCTTTGTTTTAATTAAACAGTCGGATTCCCCTTGTCCGTACCAGTTCTGAGCTGACTGTTCGACGCCCGGGGAAAGCTCCCGAGAGAGCCGTTCCCA | 686 (0.012978%) | | | AATGTATCCAGAGCGTAGGCTTGCTTTGAGCACTCTAATTTCTTCAAAGTAACAGCGCCGGAGGCACGACCCGGCCAATTAAGACCAGGAGCGTATCGCC | 5 (0.000095%) | | | AATTAGGTACGAAACACAGGCCCCGGAACTCATCATCGAGCGTAACATCGCCCGTGAATTAACTGAGAAGGATAGGTGGTAGGTAGTTCGATGCGCGAGC | 4 (0.000076%) | | | AATTCCAGCTCCAATAGCGTATATTTAAGTTGTTGCAGTTAAAAAGCTCGTAGTTGAACCTTGGGATGGGTCGGCCGGTCCGCCTTTGGTGTGCATTGGT | 15 (0.000284%) | | | AATTGTTGGTCTTCAACGAGGAATTCCTAGTAAGCGCGAGTCATCAGCTCGCGTTGACTACGTCCCTGCCCTTTGTACACACCGCCCGTCGCTCCTACCG | 1386 (0.026221%) | | | AATTTCTGCCCTATCAACTTTCGATGGTAGGATAGTGGCCTACCATGGTGGTAACGGGTGACGGAGAATTAGGGTTCGATTCCGGAGAGGGAGCCTGAGA | 14 (0.000265%) | | | ACAAACGTTCAATATGACAAACCCATGCCAAGTAAAGAGAAAATGAAAACTGGTGATTGTTGCGGAAATCGTCCAGGATTCCTCGACCAGGACTTGAAAT | 3 (0.000057%) | | | ACAAAGCCAAAGACTCATATGGACTTTGGCTACACCATGAAAGCTTTGAGAAGCAAGAAGAAGGTTGGTTAGTGTTTTGGAGTCGAATATGACTTGATGT | 150 (0.002838%) | | | ACAAAGGGTGTTGGTCGATTAAGACAGCAGGACGGTGGTCATGGAAGTCGAAATCCGCTAAGGAGTGTGTAACAACTCACCTGCCGAATCAACTAGCCCC | 10 (0.000189%) | | | ACAAATCGTCGTCCCTCACCATCCTTTGCTGATGCGGGACGGAAGCTGGTCTCCCGTGTGTTACCGCACGCGGTTGGCCTAAATCCGAGCCAAGGACGCC | 8 (0.000151%) | | | ACAAGGGTTCACATTTCGTTCATCACCCTTGGCCGGCTATCGAACAGCCGGACTCCCATCAAAAGATGGTTGCCAAGAACATCTTCGTTACGGTTTGCTA | 4 (0.000076%) | | | ACACAAAGCCAAAGACTCATATGGACTTTGGCTACACCATGAAAGCTTTGAGAAGCAAGAAGAAGGTTGGTTAGTGTTTTGGAGTCGAATATGACTTGAT | 148 (0.002800%) | | | ACACAGGCCCCGGAACTCATCATCGAGCGTAACATCGCCCGTGAATTAACTGAGAAGGATAGGTGGTAGGTAGTTCGATGCGCGAGCATGGAGCCTACGA | 3 (0.000057%) | | | ACACATGACATCAAGTCATATTCGACTCCAAAACACTAACCAACCTTCTTCTTGCTTCTCAAAGCTTTCATGGTGTAGCCAAAGTCCATATGAGTCTTTG | 21 (0.000397%) | | | ACATCAAGTCATATTCGACTCCAAAACACTAACCAACCTTCTTCTTGCTTCTCAAAGCTTTCATGGTGTAGCCAAAGTCCATATGAGTCTTTGGCTTTGT | 43 (0.000813%) | | | ACATGACATCAAGTCATATTCGACTCCAAAACACTAACCAACCTTCTTCTTGCTTCTCAAAGCTTTCATGGTGTAGCCAAAGTCCATATGAGTCTTTGGC | 67 (0.001268%) | | | ACATTGTCAGGTGGGGAGTTTGGCTGGGGCGGCACATCTGTTAAAAGATAACGCAGGTGTCCTAAGATGAGCTCAACGAGAACAGAAATCTCGTGTGGAA | 14 (0.000265%) | | | ACCAGGGGTTGAAATCGTCGACCAGGTCCGAGACTTCATCGACCGGGTCCGAGGATTCGTCGACCAGGACGGCCGGATGTCCGAGAAAAAAAAATGTTGC | 7 (0.000132%) | | | ACCATCCTTTGCTGATGCGGGACGGAAGCTGGTCTCCCGTGTGTTACCGCACGCGGTTGGCCTAAATCCGAGCCAAGGACGCCTGGAGCGTACCGACATG | 10 (0.000189%) | | | ACCCGCCGAAGCGAGCCTTGGGACCAAAAACAGGGGTTGTACCCCGCCTCCGATTCACGGAGTAAGTAAAATAACGTTAAAAGTAGTGGTATTTCACTTG | 5 (0.000095%) | | | ACCGGACACCACGCGACGTGCGGTGCTCTTCCAGCCGCTGGACCCTACCTCCGGCTGAGCCGTTTCCAGGGTGGGCAGGCTGTTAAACAGAAAAGATAAC | 5 (0.000095%) | | | ACCGGGACGTGGCGGTTGACGGCAACGTTAGGGAGTCCGGAGACGTCGGCGGGGGCCTCGGGAAGAGTTATCTTTTCTGTTTAACAGCCTGCCCACCCTG | 4 (0.000076%) | | | ACCTAGGCGAGACAAGGGTTCACATTTCGTTCATCACCCTTGGCCGGCTATCGAACAGCCGGACTCCCATCAAAAGATGGTTGCCAAGAACATCTTCGTT | 3 (0.000057%) | | | ACCTCAGCCTGCTAACTAGCTACGTGGAGGCATCCCTTCACGGCCGGCTTCTTAGAGGGACTATGGCCGTTTAGGCCAAGGAAGTTTGAGGCAATAACAG | 15 (0.000284%) | | | ACGAAACACAGGCCCCGGAACTCATCATCGAGCGTAACATCGCCCGTGAATTAACTGAGAAGGATAGGTGGTAGGTAGTTCGATGCGCGAGCATGGAGCC | 31 (0.000586%) | | | ACGAAGCACGCCCATCCAACCTAGGCGAGACAAGGGTTCACATTTCGTTCATCACCCTTGGCCGGCTATCGAACAGCCGGACTCCCATCAAAAGATGGTT | 9 (0.000170%) | | | ACGCCCATCCAACCTAGGCGAGACAAGGGTTCACATTTCGTTCATCACCCTTGGCCGGCTTTCGAACAGCCGGACTCCCATCAAAAGATGGTTGCCAAGA | 2021 (0.038234%) | | | ACGCGCCTAACGGCGTGCCTCGGCATCAGCGTGCTCCGGGCGTCGGCCTGTGGGCTCCCCATTCGACCCGTCTTGAAACACGGACCAAGGAGTCTGACAT | 23 (0.000435%) | | | ACGCTTTCACGGTTCGTATTCGTACTGAAAATCAGAATCAAACGAGCTTTTACCCTTTTGTTCCACACGAGATTTCTGTTCTCGTTGAGCTCATCTTAGG | 7 (0.000132%) | | | ACGGCAACGTTAGGGAGTCCGGAGACGTCGGCGGGGGCCTCGGGAAGAGTTATCTTTTCTGTTTAACAGCCTGCCCACCCTGGAAACGGCTCAGCCGGAG | 4 (0.000076%) | | | ACGGCGTGCCTCGGCATCAGCGTGCTCCGGGCGTCGGCCTGTGGGCTCCCCATTCGACCCGTCTTGAAACACGGACCAAGGAGTCTGACATGTGTGCGAG | 5 (0.000095%) | | | ACGGCTTTGGCTCGGATTCGTCCGTCTTCTTTCTTCTTAGCCGAGTACTTCGGTAGATTAGTTGGAACGATTGATGATTTTGAGTTAATTGAACGTTCGG | 11 (0.000208%) | | | ACGGGCGAGAGACCGATAGCGAACAAGTACCGCGAGGTAAAGATGAAAAGGACTTTGAAAAGAGAGTCAAAGAGTGCTTGAAATTGTCGGGAGGGAAGCG | 3 (0.000057%) | | | ACGGTGGTCATGGAAGTCGAAATCCGCTAAGGAGTGTGTAACAACTCACCTGCCGAATCAACTAGCCCCGAAAATGGATGGCGCTTAAGCGCGCGACCTA | 4 (0.000076%) | | | ACGTGGGTGGTTCGCCGCCCGCGACGTCGCGAGAAGTCCACTAAACCTTATCATTTAGAGGAAGGAGAAGTCGTAACAAGGTTTCCGTAGGTGAACCTGC | 5 (0.000095%) | | | ACTACCTACCACCTATCCTTCTCAGTTAATTCACGGGCGATGTTACGCTCGATGATGAGTTCCGGGGCCTGTGTTTCGTACCTAATTTGAAGGAATTGTT | 3 (0.000057%) | | | ACTAGCTACGTGGAGGCATCCCTTCACGGCCGGCTTCTTAGAGGGACTATGGCCGTTTAGGCCAAGGAAGTTTGAGGCAATAACAGGTCTGTGATGCCCT | 3 (0.000057%) | | | ACTCAATCATACACATGACATCAAGTCATATTCGACTCCAAAACACTAACCAACCTTCTTCTTGCTTCTCAAAGCTTTCATGGTGTAGCCAAAGTCCATA | 25 (0.000473%) | | | ACTCCAAAACACTAACCAACCTTCTTCTTGCTTCTCAAAGCTTTCATGGTGTAGCCAAAGTCCATATGAGTCTTTGGCTTTGTGTCTTCTAACAAGGAAA | 415 (0.007851%) | | | ACTCCAAAACACTAACCAACCTTCTTCTTGCTTCTCAAAGCTTTCATGGTGTAGCCAAAGTCCATATGAGTCTTTGGCTTTGTGTCTTCTAACAAGGATA | 10 (0.000189%) | | | ACTCGGTCCTCCGGATTTTCAAGGGCCGCCGGGGGCGCACCGGACACCACGCGACGTGCGGTGCTCTTCCAGCCGCTGGACCCTACCTCCGGCTGAGCCG | 15 (0.000284%) | | | ACTTCCCTTGCCTACATTGTTCCATCGACCAGAGGCTGTTCACCTTGGAGACCTGATGCGGTTATGAGTACGACCGGGCGTGAGCGGCACTCGGTCCTCC | 19 (0.000359%) | | | ACTTGGTGATATGAACACAAACGTTCAATATGACAAACCCATGCCAAGTAAAGAGAAAATGAAAACTGGTGATTGTTGCGGAAATCGTCCAGGATTCCTC | 5 (0.000095%) | | | AGAAGACACAAAGCCAAAGACTCATATGGACTTTGGCTACACCATGAAAGCTTTGAGAAGCAAGAAGAAGGTTGGTTAGTGTTTTGGAGTCGAATATGAC | 70 (0.001324%) | | | AGAATTGTATCCTTGTTAGAAGACACAAAGCCAAAGACTCATATGGACTTTGGCTACACCATGAAAGCTTTGAGAAGCAAGAAGAAGGTTGGTTAGTGTT | 4 (0.000076%) | | | AGACAAGGGTTCACATTTCGTTCATCACCCTTGGCCGGCTTTCGAACAGCCGGACTCCCATCAAAAGATGGTTGCCAAGAACATCTTCGTTACGGTTTGC | 3 (0.000057%) | | | AGACACAAAGCCAAAGACTCATATGGACTTTGGCTACACCATGAAAGCTTTGAGAAGCAAGAAGAAGGTTGGTTAGTGTTTTGGAGTCGAATATGACTTG | 41 (0.000776%) | | | AGACAGCAGGACGGTGGTCATGGAAGTCGAAATCCGCTAAGGAGTGTGTAACAACTCACCTGCCGAATCAACTAGCCCCGAAAATGGATGGCGCTTAAGC | 10 (0.000189%) | | | AGACGTCGGCGGGGGCCTCGGGAAGAGTTATCTTTTCTGTTTAACAGCCTGCCCACCCTGGAAACGGCTCAGCCGGAGGTAGGGTCCAGCGGCTGGAAGA | 3 (0.000057%) | | | AGAGCAACGGGCAGAGCCCGCGTCGACCTTTTATCTAATAAATGCGTCCCTTCCATAAGTCGGGGTTTGTTGCACGTATTAGCTCTAGAATTACTACGGT | 38 (0.000719%) | | | AGAGCCCGCGTCGACCTTTTATCTAATAAATGCGTCCCTTCCATAAGTCGGGGTTTGTTGCACGTATTAGCTCTAGAATTACTACGGTTATCCGAGTAGT | 9 (0.000170%) | | | AGAGCGTAGGCTTGCTTTGAGCACTCTAATTTCTTCAAAGTAACAGCGCCGGAGGCACGACCCGGCCAATTAAGACCAGGAGCGTATCGCCGACCGAAGG | 14 (0.000265%) | | | AGAGCGTCGTGTAGGGAAAGAGTGTTCAGAGCCGTGTAGA | 13 (0.000098%) | | | AGAGGCTGTTCACCTTGGAGACCTGATGCGGTTATGAGTACGACCGGGCGTGAGCGGCACTCGGTCCTCCGGATTTTCAAGGGCCGCCGGGGGCGCACCG | 148 (0.002800%) | | | AGATCGGAAGAGCGTCGTGTAGGGAAAGAGTGTTCAGAGC | 19671 (0.148856%) | | | AGCAACGGGCAGAGCCCGCGTCGACCTTTTATCTAATAAATGCGTCCCTTCCATAAGTCGGGGTTTGTTGCACGTATTAGCTCTAGAATTACTACGGTTA | 3 (0.000057%) | | | AGCACGCCCATCCAACCTAGGCGAGACAAGGGTTCACATTTCGTTCATCACCCTTGGCCGGCTATCGAACAGCCGGACTCCCATCAAAAGATGGTTGCCA | 12 (0.000227%) | | | AGCACGCGCCTAACGGCGTGCCTCGGCATCAGCGTGCTCCGGGCGTCGGCCTGTGGGCTCCCCATTCGACCCGTCTTGAAACACGGACCAAGGAGTCTGA | 30 (0.000568%) | | | AGCAGCCGCGGTAATTCCAGCTCCAATAGCGTATATTTAAGTTGTTGCAGTTAAAAAGCTCGTAGTTGAACCTTGGGATGGGTCGGCCGGTCCGCCTTTG | 8 (0.000151%) | | | AGCAGGACGGTGGTCATGGAAGTCGAAATCCGCTAAGGAGTGTGTAACAACTCACCTGCCGAATCAACTAGCCCCGAAAATGGATGGCGCTTAAGCGCGC | 6 (0.000114%) | | | AGCCAAAGACTCATATGGACTTTGGCTACACCATGAAAGCTTTGAGAAGCAAGAAGAAGGTTGGTTAGTGTTTTGGAGTCGAATATGACTTGATGTCATG | 10 (0.000189%) | | | AGCCCGCGTCGACCTTTTATCTAATAAATGCGTCCCTTCCATAAGTCGGGGTTTGTTGCACGTATTAGCTCTAGAATTACTACGGTTATCCGAGTAGTAG | 3 (0.000057%) | | | AGCCGCGGTAATTCCAGCTCCAATAGCGTATATTTAAGTTGTTGCAGTTAAAAAGCTCGTAGTTGAACCTTGGGATGGGTCGGCCGGTCCGCCTTTGGTG | 7 (0.000132%) | | | AGCCTAAGTAGTGTTTCCTTGTTAGAAGACACAAAGCCAAAGACTCATATGGACTTTGGCTACACCATGAAAGCTTTGAGAAGCAAGAAGAAGGTTGGTT | 36 (0.000681%) | | | AGCCTGCTAACTAGCTACGTGGAGGCATCCCTTCACGGCCGGCTTCTTAGAGGGACTATGGCCGTTTAGGCCAAGGAAGTTTGAGGCAATAACAGGTCTG | 8 (0.000151%) | | | AGCGACGGGCGGTGTGTACAAAGGGCAGGGACGTAGTCAACGCGAGCTGATGACTCGCGCTTACTAGGAATTCCTCGTTGAAGACCAACAATTGCAATGA | 4 (0.000076%) | | | AGCGAGGTGTGAGTGTCGCCCATGGGCATCGACACCTTGCGGCTAGGAACTGGAACGAGACGGGTGGCAAAGATTTCGAGTAGCACTTCATACTACCGTG | 4 (0.000076%) | | | AGCGGCACTCGGTCCTCCGGATTTTCAAGGGCCGCCGGGGGCGCACCGGACACCACGCGACGTGCGGTGCTCTTCCAGCCGCTGGACCCTACCTCCGGCT | 6 (0.000114%) | | | AGCGTAGGCTTGCTTTGAGCACTCTAATTTCTTCAAAGTAACAGCGCCGGAGGCACGACCCGGCCAATTAAGACCAGGAGCGTATCGCCGACCGAAGGGA | 7 (0.000132%) | | | AGCGTCGTGTAGGGAAAGAGTGTTCAGAGCCGTGTAGATC | 8 (0.000061%) | | | AGCTAGTGTTCGTAGGCTCCATGCTCGCGCATCGAACTACCTACCACCTATCCTTCTCAGTTAATTCACGGGCGATGTTACGCTCGATGATGAGTTCCGG | 10 (0.000189%) | | | AGCTCCAATAGCGTATATTTAAGTTGTTGCAGTTAAAAAGCTCGTAGTTGAACCTTGGGATGGGTCGGCCGGTCCGCCTTTGGTGTGCATTGGTCGGCTT | 5 (0.000095%) | | | AGCTCGTGTAAAGTTGGGAATTCGTTAAGGAGCTGTTGCTTTGTTAGTGTAGAAACACTTGTGTAGAATTGGGGATTGTTTTTTTTGGAGTGATTTAGGG | 10 (0.000189%) | | | AGCTTCCTTGCGCCTTACGGGTTTACTCACCCGTTGACTCGCACACATGTCAGACTCCTTGGTCCGTGTTTCAAGACGGGTCGAATGGGGAGCCCACAGG | 30 (0.000568%) | | | AGGACATTGTCAGGTGGGGAGTTTGGCTGGGGCGGCACATCTGTTAAAAGATAACGCAGGTGTCCTAAGATGAGCTCAACGAGAACAGAAATCTCGTGTG | 179 (0.003386%) | | | AGGACGGTGGTCATGGAAGTCGAAATCCGCTAAGGAGTGTGTAACAACTCACCTGCCGAATCAACTAGCCCCGAAAATGGATGGCGCTTAAGCGCGCGAC | 27 (0.000511%) | | | AGGAGCGACGGGCGGTGTGTACAAAGGGCAGGGACGTAGTCAACGCGAGCTGATGACTCGCGCTTACTAGGAATTCCTCGTTGAAGACCAACAATTGCAA | 17 (0.000322%) | | | AGGAGCTGTTGCTTTGTTAGTGTAGAAACACTTGTGTAGAATTGGGGATTGTTTTTTTTGGAGTGATTTAGGGGAGGGTCGAATCTTAGCGACAAAGGGC | 11 (0.000208%) | | | AGGCGAGACAAGGGTTCACATTTCGTTCATCACCCTTGGCCGGCTATCGAACAGCCGGACTCCCATCAAAAGATGGTTGCCAAGAACATCTTCGTTACGG | 5 (0.000095%) | | | AGGCGTAAGAATTGTATCCTTGTTAGAAGACACAAAGCCA | 116 (0.000878%) | | | AGGCTCCATGCTCGCGCATCGAACTACCTACCACCTATCCTTCTCAGTTAATTCACGGGCGATGTTACGCTCGATGATGAGTTCCGGGGCCTGTGTTTCG | 5 (0.000095%) | | | AGGCTGTCCCGAGTGTGAGCGAGGTGTGAGTGTCGCCCATGGGCATCGACACCTTGCGGCTAGGAACTGGAACGAGACGGGTAGCAAAGATTTCGAGTAG | 1796 (0.033977%) | | | AGGCTGTCCCGAGTGTGAGCGAGGTGTGAGTGTCGCCCATGGGCATCGACACCTTGCGGCTAGGAACTGGAACGAGACGGGTGGCAAAGATTTCGAGTAG | 15 (0.000284%) | | | AGGCTGTTCACCTTGGAGACCTGATGCGGTTATGAGTACGACCGGGCGTGAGCGGCACTCGGTCCTCCGGATTTTCAAGGGCCGCCGGGGGCGCACCGGA | 33 (0.000624%) | | | AGGGAAAGAGTGTTCAGAGCCGTGTAGATCTCGGTGGTCG | 52 (0.000393%) | | | AGGGACAGTCGGGGGCATTCGTATTTCATAGTCAGAGGTGAAATTCTTGGATTTATGAAAGACGAACAACTGCGAAAGCATTTGCCAAGGATGTTTTCAT | 10 (0.000189%) | | | AGGGAGTCCGGAGACGTCGGCGGGGGCCTCGGGAAGAGTTATCTTTTCTGTTTAACAGCCTGCCCACCCTGGAAACGGCTCAGCCGGAGGTAGGGTCCAG | 6 (0.000114%) | | | AGGGCAAGTCTGGTGCCAGCAGCCGCGGTAATTCCAGCTCCAATAGCGTATATTTAAGTTGTTGCAGTTAAAAAGCTCGTAGTTGAACCTTGGGATGGGT | 22 (0.000416%) | | | AGGGTGTTGGTCGATTAAGACAGCAGGACGGTGGTCATGGAAGTCGAAATCCGCTAAGGAGTGTGTAACAACTCACCTGCCGAATCAACTAGCCCCGAAA | 13 (0.000246%) | | | AGGTACGAAACACAGGCCCCGGAACTCATCATCGAGCGTAACATCGCCCGTGAATTAACTGAGAAGGATAGGTGGTAGGTAGTTCGATGCGCGAGCATGG | 3 (0.000057%) | | | AGGTATCTCGCGCTTGTACGGCTTTGGCTCGGATTCGTCCGTCTTCTTTCTTCTTAGCCGAGTACTTCGGTAGATTAGTTGGAACGATTGATGATTTTGA | 6 (0.000114%) | | | AGGTGTGAGTGTCGCCCATGGGCATCGACACCTTGCGGCTAGGAACTGGAACGAGACGGGTGGCAAAGATTTCGAGTAGCACTTCATACTACCGTGGGTT | 3 (0.000057%) | | | AGTAGTCATATGCTTGTCTCAAAGATTAAGCCATGCATGTGTAAGTATGAACGAATTCAGACTGTGAAACTGCGAATGGCTCATTAAATCAGTTATAGTT | 2227 (0.042131%) | | | AGTAGTGTTTCCTTGTTAGAAGACACAAAGCCAAAGACTCATATGGACTTTGGCTACACCATGAAAGCTTTGAGAAGCAAGAAGAAGGTTGGTTAGTGTT | 23 (0.000435%) | | | AGTATAAGAACTTAAACCGCAACCGCATCTTATAAGCCTAAGTAGTGTTTCCTTGTTAGAAGACACAAAGCCAAAGACTCATATGGACTTTGGCTACACC | 6 (0.000114%) | | | AGTCATATGCTTGTCTCAAAGATTAAGCCATGCATGTGTAAGTATGAACGAATTCAGACTGTGAAACTGCGAATGGCTCATTAAATCAGTTATAGTTTGT | 14 (0.000265%) | | | AGTCATATTCGACTCCAAAACACTAACCAACCTTCTTCTTGCTTCTCAAAGCTTTCATGGTGTAGCCAAAGTCCATATGAGTCTTTGGCTTTGTGTCTTC | 5 (0.000095%) | | | AGTCGAATATGACTTGATGTCATGTGTATGATTGAGTATA | 1222 (0.009247%) | | | AGTCGGGGGCATTCGTATTTCATAGTCAGAGGTGAAATTCTTGGATTTATGAAAGACGAACAACTGCGAAAGCATTTGCCAAGGATGTTTTCATTAATCA | 3 (0.000057%) | | | AGTGTATCCTTGTTAGAAGACACAAAGCCAAAGACTCATA | 8087 (0.061197%) | | | AGTGTTCGTAGGCTCCATGCTCGCGCATCGAACTACCTACCACCTATCCTTCTCAGTTAATTCACGGGCGATGTTACGCTCGATGATGAGTTCCGGGGCC | 262 (0.004957%) | | | AGTGTTTCCTTGTTAGAAGACACAAAGCCAAAGACTCATATGGACTTTGGCTACACCATGAAAGCTTTGAGAAGCAAGAAGAAGGTTGGTTAGTGTTTTG | 12 (0.000227%) | | | AGTGTTTTGGAGTCGAATATGACTTGATGTCATGTGTATG | 204 (0.001544%) | | | AGTTATCATGAATCATCAGAGCAACGGGCAGAGCCCGCGTCGACCTTTTATCTAATAAATGCGTCCCTTCCATAAGTCGGGGTTTGTTGCACGTATTAGC | 19 (0.000359%) | | | AGTTCTCGGACAAAAATTGCTGAGTGGCCGAGAAGAATGGGCGTGTCATGCGTGGGCTGACATGGATTCTTCGAGGCCTAGGGGTGGCGGTATATAACTT | 5 (0.000095%) | | | AGTTCTTATACTCAATCATACACATGACATCAAGTCATATTCGACTCCAAAACACTAACCAACCTTCTTCTTGCTTCTCAAAGCTTTCATGGTGTAGCCA | 19 (0.000359%) | | | AGTTGTTATACTCAATCATACACATGACAACAAGTCATATTCGACTCCAAAACACTAACCAACCTTCTTCTTGCTTCTCAAAGCTTTCATGGTGTAGCCA | 15 (0.000284%) | | | AGTTGTTATACTCAATCATACACATGACATCAAGTCATAT | 98 (0.000742%) | | | AGTTTTTTCAGCAGTTCTCGGACAAAAATTGCTGAGTGGCCGAGAAGAATGGGCGTGTCATGCGTGGGCTGACATGGATTCTTCGAGGCCTAGGGGTGGC | 3 (0.000057%) | | | ATAAAAGTGTCGGAGTTTTTTCAGCAGTTCTCGGACAAAAATTGCTGAGTGGCCGAGAAGAATGGGCGTGTCATGCGTGGGCTGACATGGATTCTTCGAG | 153 (0.002894%) | | | ATAAGAACTTAAACCGCAACCGCATCTTATAAGCCTAAGTAGTGTTTCCTTGTTAGAAGACACAAAGCCAAAGACTCATATGGACTTTGGCTACACCATG | 26 (0.000492%) | | | ATACACATGACATCAAGTCATATTCGACTCCAAAACACTAACCAACCTTCTTCTTGCTTCTCAAAGCTTTCATGGTGTAGCCAAAGTCCATATGAGTCTT | 272 (0.005146%) | | | ATACGGGCGAGAGACCGATAGCGAACAAGTACCGCGAGGTAAAGATGAAAAGGACTTTGAAAAGAGAGTCAAAGAGTGCTTGAAATTGTCGGGAGGGAAG | 17 (0.000322%) | | | ATACTCAATCATACACATGACATCAAGTCATATTCGACTCCAAAACACTAACCAACCTTCTTCTTGCTTCTCAAAGCTTTCATGGTGTAGCCAAAGTCCA | 30 (0.000568%) | | | ATAGCTAGTGTTCGTAGGCTCCATGCTCGCGCATCGAACTACCTACCACCTATCCTTCTCAGTTAATTCACGGGCGATGTTACGCTCGATGATGAGTTCC | 5 (0.000095%) | | | ATATGCTTGTCTCAAAGATTAAGCCATGCATGTGTAAGTATGAACGAATTCAGACTGTGAAACTGCGAATGGCTCATTAAATCAGTTATAGTTTGTTTGA | 55 (0.001041%) | | | ATATTCGACTCCAAAACACTAACCAACCTTCTTCTTGCTTCTCAAAGCTTTCATGGTGTAGCCAAAGTCCATATGAGTCTTTGGCTTTGTGTCTTCTAAC | 151 (0.002857%) | | | ATATTCGACTCCAAAACACTAACCAACCTTCTTCTTGCTTCTCAAAGCTTTCATGGTGTAGCCAAAGTCCATATGAGTCTTTGGCTTTGTGTCTTTTAAC | 59 (0.001116%) | | | ATATTTAAGTTGTTGCAGTTAAAAAGCTCGTAGTTGAACCTTGGGATGGGTCGGCCGGTCCGCCTTTGGTGTGCATTGGTCGGCTTGTCCCTTCGGTCGG | 7 (0.000132%) | | | ATCAACTTTCGATGGTAGGATAGTGGCCTACCATGGTGGTAACGGGTGACGGAGAATTAGGGTTCGATTCCGGAGAGGGAGCCTGAGAAACGGCTACCAC | 9 (0.000170%) | | | ATCAAGTCATATTCGACTCCAAAACACTAACCAACCTTCTTCTTGCTTCTCAAAGCTTTCATGGTGTAGCCAAAGTCCATATGAGTCTTTGGCTTTGTGT | 1473 (0.027867%) | | | ATCAATCGTTCCAACTAATCTACCGAAGTACTCGGCTAAGAAGAAAGAAGACGGACGAATCCGAGCCAAAGCCGTACAAGCGCGAGATACCTTCGGGACA | 4 (0.000076%) | | | ATCAGAGCAACGGGCAGAGCCCGCGTCGACCTTTTATCTAATAAATGCGTCCCTTCCATAAGTCGGGGTTTGTTGCACGTATTAGCTCTAGAATTACTAC | 32 (0.000605%) | | | ATCAGCGTGCTCCGGGCGTCGGCCTGTGGGCTCCCCATTCGACCCGTCTTGAAACACGGACCAAGGAGTCTGACATGTGTGCGAGTCAACGGGTGAGTAA | 5 (0.000095%) | | | ATCAGCTTCCTTGCGCCTTACGGGTTTACTCACCCGTTGACTCGCACACATGTCAGACTCCTTGGTCCGTGTTTCAAGACGGGTCGAATGGGGAGCCCAC | 47 (0.000889%) | | | ATCATACACATGACATCAAGTCATATTCGACTCCAAAACACTAACCAACCTTCTTCTTGCTTCTCAAAGCTTTCATGGTGTAGCCAAAGTCCATATGAGT | 29 (0.000549%) | | | ATCATCAGAGCAACGGGCAGAGCCCGCGTCGACCTTTTATCTAATAAATGCGTCCCTTCCATAAGTCGGGGTTTGTTGCACGTATTAGCTCTAGAATTAC | 15 (0.000284%) | | | ATCATGAATCATCAGAGCAACGGGCAGAGCCCGCGTCGACCTTTTATCTAATAAATGCGTCCCTTCCATAAGTCGGGGTTTGTTGCACGTATTAGCTCTA | 62 (0.001173%) | | | ATCATTCAAATTTCTGCCCTATCAACTTTCGATGGTAGGATAGTGGCCTACCATGGTGGTAACGGGTGACGGAGAATTAGGGTTCGATTCCGGAGAGGGA | 4 (0.000076%) | | | ATCCAACCTAGGCGAGACAAGGGTTCACATTTCGTTCATCACCCTTGGCCGGCTATCGAACAGCCGGACTCCCATCAAAAGATGGTTGCCAAGAACATCT | 25 (0.000473%) | | | ATCCAACCTAGGCGAGACAAGGGTTCACATTTCGTTCATCACCCTTGGCCGGCTTTCGAACAGCCGGACTCCCATCAAAAGATGGTTGCCAAGAACATCT | 27 (0.000511%) | | | ATCCAGAGCGTAGGCTTGCTTTGAGCACTCTAATTTCTTCAAAGTAACAGCGCCGGAGGCACGACCCGGCCAATTAAGACCAGGAGCGTATCGCCGACCG | 15 (0.000284%) | | | ATCCATTACATTTTATCGGTCGCTCTTGTCCGGAAGCTGTAGATGACCCAAAGTCCATATAGCGACCCCAGGTCAGGCGGGATTACCCGCTGAGTTTAAG | 3 (0.000057%) | | | ATCCCATGCTAATGTATCCAGAGCGTAGGCTTGCTTTGAGCACTCTAATTTCTTCAAAGTAACAGCGCCGGAGGCACGACCCGGCCAATTAAGACCAGGA | 11 (0.000208%) | | | ATCCCGCCAATCAGCTTCCTTGCGCCTTACGGGTTTACTCACCCGTTGACTCGCACACATGTCAGACTCCTTGGTCCGTGTTTCAAGACGGGTCGAATGG | 10 (0.000189%) | | | ATCCGTCGAGTTATCATGAATCATCAGAGCAACGGGCAGAGCCCGCGTCGACCTTTTATCTAATAAATGCGTCCCTTCCATAAGTCGGGGTTTGTTGCAC | 9 (0.000170%) | | | ATCCTATGATGTTATCCCATGCTAATGTATCCAGAGCGTAGGCTTGCTTTGAGCACTCTAATTTCTTCAAAGTAACAGCGCCGGAGGCACGACCCGGCCA | 37 (0.000700%) | | | ATCCTTGTTAGAAGACACAAAGCCAAAGACTCATATGGACTTTGGCTACACCATGAAAGCTTTGAGAAGCAAGAAGAAGGTTGGTTAGTGTTTTGGAGTC | 10 (0.000189%) | | | ATCCTTTGCTGATGCGGGACGGAAGCTGGTCTCCCGTGTGTTACCGCACGCGGTTGGCCTAAATCCGAGCCAAGGACGCCTGGAGCGTACCGACATGCGG | 4 (0.000076%) | | | ATCGAAATCCTATGATGTTATCCCATGCTAATGTATCCAGAGCGTAGGCTTGCTTTGAGCACTCTAATTTCTTCAAAGTAACAGCGCCGGAGGCACGACC | 16 (0.000303%) | | | ATCGAACTACCTACCACCTATCCTTCTCAGTTAATTCACGGGCGATGTTACGCTCGATGATGAGTTCCGGGGCCTGTGTTTCGTACCTAATTTGAAGGAA | 5 (0.000095%) | | | ATCGACACCTTGCGGCTAGGAACTGGAACGAGACGGGTGGCAAAGATTTCGAGTAGCACTTCATACTACCGTGGGTTTTTTAAACCTTCCGAGTTTTGTT | 7 (0.000132%) | | | ATCGACCCGCCGAAGCGAGCCTTGGGACCAAAAACAGGGGTTGTACCCCGCCTCCGATTCACGGAGTAAGTAAAATAACGTTAAAAGTAGTGGTATTTCA | 4 (0.000076%) | | | ATCGATCCATTACATTTTATCGGTCGCTCTTGTCCGGAAGCTGTAGATGACCCAAAGTCCATATAGCGACCCCAGGTCAGGCGGGATTACCCGCTGAGTT | 8 (0.000151%) | | | ATCGATCCCGCCAATCAGCTTCCTTGCGCCTTACGGGTTTACTCACCCGTTGACTCGCACACATGTCAGACTCCTTGGTCCGTGTTTCAAGACGGGTCGA | 1263 (0.023894%) | | | ATCGATCCGTAACTTCGGGAAAAGGATTGGCTCTGAGGGCTGGGCTCGGGGGTCCCAGTTCCGAACCCGTCGGCTGTCAGCGGACTGCTCGAGCTGCTTC | 32 (0.000605%) | | | ATCGATCCGTCGAGTTATCATGAATCATCAGAGCAACGGGCAGAGCCCGCGTCGACCTTTTATCTAATAAATGCGTCCCTTCCATAAGTCGGGGTTTGTT | 23 (0.000435%) | | | ATCGATCGACCCGCCGAAGCGAGCCTTGGGACCAAAAACAGGGGTTGTACCCCGCCTCCGATTCACGGAGTAAGTAAAATAACGTTAAAAGTAGTGGTAT | 12 (0.000227%) | | | ATCGATCTCATGTGTATGATTGAGTATAAGAACTTAAACCGCAACCGCATCTTATAAGCCTAAGTAGTGTTTCCTTGTTAGAAGACACAAAGCCAAAGAC | 12 (0.000227%) | | | ATCGATCTTAAAGGCGTAAGAATTGTATCCTTGTTAAAAGACACAAAGCCAAAGACTCATATGGACTTTGGCTACACCATGAAAGCTTTGAGAAGCAAGA | 17 (0.000322%) | | | ATCGATCTTCTGAGAAGGGTTCGAGTGTGAGCATGCCTGTCGGGACCCGAAAGATGGTGAACTATGCCTGAGCGGGGTAAAGCCAGAGGAAACTCTGGTG | 18 (0.000341%) | | | ATCGGAAGAGCGTCGTGTAGGGAAAGAGTGTTCAGAGCCG | 19 (0.000144%) | | | ATCGGTAGGAGCGACGGGCGGTGTGTACAAAGGGCAGGGACGTAGTCAACGCGAGCTGATGACTCGCGCTTACTAGGAATTCCTCGTTGAAGACCAACAA | 83 (0.001570%) | | | ATCGTCGTCCCTCACCATCCTTTGCTGATGCGGGACGGAAGCTGGTCTCCCGTGTGTTACCGCACGCGGTTGGCCTAAATCCGAGCCAAGGACGCCTGGA | 4 (0.000076%) | | | ATCTCATGTGTATGATTGAGTATAAGAACTTAAACCGCAACCGCATCTTATAAGCCTAAGTAGTGTTTCCTTGTTAGAAGACACAAAGCCAAAGACTCAT | 7 (0.000132%) | | | ATCTCGCGCTTGTACGGCTTTGGCTCGGATTCGTCCGTCTTCTTTCTTCTTAGCCGAGTACTTCGGTAGATTAGTTGGAACGATTGATGATTTTGAGTTA | 4 (0.000076%) | | | ATCTTAAAGGCGTAAGAATTGTATCCTTGTTAAAAGACACAAAGCCAAAGACTCATATGGACTTTGGCTACACCATGAAAGCTTTGAGAAGCAAGAAGAA | 47 (0.000889%) | | | ATCTTAAAGGCGTAAGAATTGTATCCTTGTTAGAAGACAC | 1325 (0.010027%) | | | ATCTTCTGAGAAGGGTTCGAGTGTGAGCATGCCTGTCGGGACCCGAAAGATGGTGAACTATGCCTGAGCGGGGTAAAGCCAGAGGAAACTCTGGTGGAAG | 3 (0.000057%) | | | ATGAACACAAACGTTCAATATGACAAACCCATGCCAAGTAAAGAGAAAATGAAAACTGGTGATTGTTGCGGAAATCGTCCAGGATTCCTCGACCAGGACT | 11 (0.000208%) | | | ATGAATCATCAGAGCAACGGGCAGAGCCCGCGTCGACCTTTTATCTAATAAATGCGTCCCTTCCATAAGTCGGGGTTTGTTGCACGTATTAGCTCTAGAA | 71 (0.001343%) | | | ATGACATCAAGTCATATTCGACTCCAAAACACTAACCAACCTTCTTCTTGCTTCTCAAAGCTTTCATGGTGTAGCCAAAGTCCATATGAGTCTTTGGCTT | 117 (0.002213%) | | | ATGATGTTATCCCATGCTAATGTATCCAGAGCGTAGGCTTGCTTTGAGCACTCTAATTTCTTCAAAGTAACAGCGCCGGAGGCACGACCCGGCCAATTAA | 63 (0.001192%) | | | ATGATTAACAGGGACAGTCGGGGGCATTCGTATTTCATAGTCAGAGGTGAAATTCTTGGATTTATGAAAGACGAACAACTGCGAAAGCATTTGCCAAGGA | 93 (0.001759%) | | | ATGATTGAGTATAAGAACTTAAACCGCAACCGCATCTTATAAGCCTAAGTAGTGTTTCCTTGTTAGAAGACACAAAGCCAAAGACTCATATGGACTTTGG | 29 (0.000549%) | | | ATGCTAATGTATCCAGAGCGTAGGCTTGCTTTGAGCACTCTAATTTCTTCAAAGTAACAGCGCCGGAGGCACGACCCGGCCAATTAAGACCAGGAGCGTA | 14 (0.000265%) | | | ATGCTTGTCTCAAAGATTAAGCCATGCATGTGTAAGTATGAACGAATTCAGACTGTGAAACTGCGAATGGCTCATTAAATCAGTTATAGTTTGTTTGATG | 14 (0.000265%) | | | ATGCTTTGTTTTAATTAAACAGTCGGATTCCCCTTGTCCGTACCAGTTCTGAGCTGACTGTTCGACGCCCGGGGAAAGCTCCCGAGAGAGCCGTTCCCAG | 4 (0.000076%) | | | ATGGAAGTCGAAATCCGCTAAGGAGTGTGTAACAACTCACCTGCCGAATCAACTAGCCCCGAAAATGGATGGCGCTTAAGCGCGCGACCTATACCCGGCC | 33 (0.000624%) | | | ATGGCCTCTGTGCTGGCGACGCATCATTCAAATTTCTGCCCTATCAACTTTCGATGGTAGGATAGTGGCCTACCATGGTGGTAACGGGTGACGGAGAATT | 381 (0.007208%) | | | ATGGGCATCGACACCTTGCGGCTAGGAACTGGAACGAGACGGGTGGCAAAGATTTCGAGTAGCACTTCATACTACCGTGGGTTTTTTAAACCTTCCGAGT | 8 (0.000151%) | | | ATGTATCCAGAGCGTAGGCTTGCTTTGAGCACTCTAATTTCTTCAAAGTAACAGCGCCGGAGGCACGACCCGGCCAATTAAGACCAGGAGCGTATCGCCG | 5 (0.000095%) | | | ATGTGTAAGTATGAACGAATTCAGACTGTGAAACTGCGAATGGCTCATTAAATCAGTTATAGTTTGTTTGATGGTAACTACTACTCGGATAACCGTAGTA | 188 (0.003557%) | | | ATGTGTATGATTGAGTATAAGAACTTAAACCGCAACCGCATCTTATAAGCCTAAGTAGTGTTTCCTTGTTAGAAGACACAAAGCCAAAGACTCATATGGA | 13 (0.000246%) | | | ATGTTATCCCATGCTAATGTATCCAGAGCGTAGGCTTGCTTTGAGCACTCTAATTTCTTCAAAGTAACAGCGCCGGAGGCACGACCCGGCCAATTAAGAC | 13 (0.000246%) | | | ATTAACAGGGACAGTCGGGGGCATTCGTATTTCATAGTCAGAGGTGAAATTCTTGGATTTATGAAAGACGAACAACTGCGAAAGCATTTGCCAAGGATGT | 99 (0.001873%) | | | ATTAAGACAGCAGGACGGTGGTCATGGAAGTCGAAATCCGCTAAGGAGTGTGTAACAACTCACCTGCCGAATCAACTAGCCCCGAAAATGGATGGCGCTT | 21 (0.000397%) | | | ATTAGGTACGAAACACAGGCCCCGGAACTCATCATCGAGCGTAACATCGCCCGTGAATTAACTGAGAAGGATAGGTGGTAGGTAGTTCGATGCGCGAGCA | 13 (0.000246%) | | | ATTAGTCTTTCGCCCCTATACCCAAGTCAGACGAACGATTTGCACGTCAGTATCGCTGCGGGCTTCCACCAGAGTTTCCTCTGGCTTTACCCCGCTCAGG | 24 (0.000454%) | | | ATTCAAATTTCTGCCCTATCAACTTTCGATGGTAGGATAGTGGCCTACCATGGTGGTAACGGGTGACGGAGAATTAGGGTTCGATTCCGGAGAGGGAGCC | 12 (0.000227%) | | | ATTCAATCGGTAGGAGCGACGGGCGGTGTGTACAAAGGGCAGGGACGTAGTCAACGCGAGCTGATGACTCGCGCTTACTAGGAATTCCTCGTTGAAGACC | 1057 (0.019997%) | | | ATTCCAGCTCCAATAGCGTATATTTAAGTTGTTGCAGTTAAAAAGCTCGTAGTTGAACCTTGGGATGGGTCGGCCGGTCCGCCTTTGGTGTGCATTGGTC | 4 (0.000076%) | | | ATTCGACTCCAAAACACTAACCAACCTTCTTCTTGCTTCTCAAAGCTTTCATGGTGTAGCCAAAGTCCATATGAGTCTTTGGCTTTGTGTCTTCTAACAA | 93 (0.001759%) | | | ATTCGACTCCAAAACACTAACCAACCTTCTTCTTGCTTCTCAAAGCTTTCATGGTGTAGCCAAAGTCCATATGAGTCTTTGGCTTTGTGTCTTTTAACAA | 201 (0.003803%) | | | ATTCGTACTGAAAATCAGAATCAAACGAGCTTTTACCCTTTTGTTCCACACGAGATTTCTGTTCTCGTTGAGCTCATCTTAGGACACCTGCGTTATCTTT | 12 (0.000227%) | | | ATTCGTATTTCATAGTCAGAGGTGAAATTCTTGGATTTATGAAAGACGAACAACTGCGAAAGCATTTGCCAAGGATGTTTTCATTAATCAAGAACGAAAG | 8 (0.000151%) | | | ATTGAGTATAAGAACTTAAACCGCAACCGCATCTTATAAGCCTAAGTAGTGTTTCCTTGTTAGAAGACACAAAGCCAAAGACTCATATGGACTTTGGCTA | 29 (0.000549%) | | | ATTGGAGGGCAAGTCTGGTGCCAGCAGCCGCGGTAATTCCAGCTCCAATAGCGTATATTTAAGTTGTTGCAGTTAAAAAGCTCGTAGTTGAACCTTGGGA | 1509 (0.028548%) | | | ATTGTATCCTTGTTAGAAGACACAAAGCCAAAGACTCATATGGACTTTGGCTACACCATGAAAGCTTTGAGAAGCAAGAAGAAGGTTGGTTAGTGTTTTG | 26 (0.000492%) | | | ATTGTCAGGTGGGGAGTTTGGCTGGGGCGGCACATCTGTTAAAAGATAACGCAGGTGTCCTAAGATGAGCTCAACGAGAACAGAAATCTCGTGTGGAACA | 24 (0.000454%) | | | ATTGTTCCATCGACCAGAGGCTGTTCACCTTGGAGACCTGATGCGGTTATGAGTACGACCGGGCGTGAGCGGCACTCGGTCCTCCGGATTTTCAAGGGCC | 11 (0.000208%) | | | ATTGTTGGTCTTCAACGAGGAATTCCTAGTAAGCGCGAGTCATCAGCTCGCGTTGACTACGTCCCTGCCCTTTGTACACACCGCCCGTCGCTCCTACCGA | 41 (0.000776%) | | | ATTTCATAGTCAGAGGTGAAATTCTTGGATTTATGAAAGACGAACAACTGCGAAAGCATTTGCCAAGGATGTTTTCATTAATCAAGAACGAAAGTTGGGG | 6 (0.000114%) | | | ATTTCGTTCATCACCCTTGGCCGGCTATCGAACAGCCGGACTCCCATCAAAAGATGGTTGCCAAGAACATCTTCGTTACGGTTTGCTAATTCTCGGAATA | 19 (0.000359%) | | | ATTTCGTTCATCACCCTTGGCCGGCTTTCGAACAGCCGGACTCCCATCAAAAGATGGTTGCCAAGAACATCTTCGTTACGGTTTGCTAATTCTCGGAATA | 12 (0.000227%) | | | ATTTCTGCCCTATCAACTTTCGATGGTAGGATAGTGGCCTACCATGGTGGTAACGGGTGACGGAGAATTAGGGTTCGATTCCGGAGAGGGAGCCTGAGAA | 7 (0.000132%) | | | ATTTGCACGTCAGTATCGCTGCGGGCTTCCACCAGAGTTTCCTCTGGCTTTACCCCGCTCAGGCATAGTTCACCATCTTTCGGGTCCCGACAGGCATGCT | 33 (0.000624%) | | | ATTTTATCGGTCGCTCTTGTCCGGAAGCTGTAGATGACCCAAAGTCCATATAGCGACCCCAGGTCAGGCGGGATTACCCGCTGAGTTTAAGCATATCAAT | 16 (0.000303%) | | | ATTTTCAAGGGCCGCCGGGGGCGCACCGGACACCACGCGACGTGCGGTGCTCTTCCAGCCGCTGGACCCTACCTCCGGCTGAGCCGTTTCCAGGGTGGGC | 7 (0.000132%) | | | CAAAATCATCAATCGTTCCAACTAATCTACCGAAGTACTCGGCTAAGAAGAAAGAAGACGGACGAATCCGAGCCAAAGCCGTACAAGCGCGAGATACCTT | 29 (0.000549%) | | | CAAACGTTCAATATGACAAACCCATGCCAAGTAAAGAGAAAATGAAAACTGGTGATTGTTGCGGAAATCGTCCAGGATTCCTCGACCAGGACTTGAAATC | 28 (0.000530%) | | | CAAAGCCAAAGACTCATATGGACTTTGGCTACACCATGAAAGCTTTGAGAAGCAAGAAGAAGGTTGGTTAGTGTTTTGGAGTCGAATATGACTTGATGTC | 113 (0.002138%) | | | CAAAGGGTGTTGGTCGATTAAGACAGCAGGACGGTGGTCATGGAAGTCGAAATCCGCTAAGGAGTGTGTAACAACTCACCTGCCGAATCAACTAGCCCCG | 31 (0.000586%) | | | CAAATCGTCGTCCCTCACCATCCTTTGCTGATGCGGGACGGAAGCTGGTCTCCCGTGTGTTACCGCACGCGGTTGGCCTAAATCCGAGCCAAGGACGCCT | 5 (0.000095%) | | | CAAATTAGGTACGAAACACAGGCCCCGGAACTCATCATCGAGCGTAACATCGCCCGTGAATTAACTGAGAAGGATAGGTGGTAGGTAGTTCGATGCGCGA | 8 (0.000151%) | | | CAAATTTCTGCCCTATCAACTTTCGATGGTAGGATAGTGGCCTACCATGGTGGTAACGGGTGACGGAGAATTAGGGTTCGATTCCGGAGAGGGAGCCTGA | 113 (0.002138%) | | | CAACCTAGGCGAGACAAGGGTTCACATTTCGTTCATCACCCTTGGCCGGCTATCGAACAGCCGGACTCCCATCAAAAGATGGTTGCCAAGAACATCTTCG | 4 (0.000076%) | | | CAACGAAGCACGCCCATCCAACCTAGGCGAGACAAGGGTTCACATTTCGTTCATCACCCTTGGCCGGCTATCGAACAGCCGGACTCCCATCAAAAGATGG | 71 (0.001343%) | | | CAACGGGCAGAGCCCGCGTCGACCTTTTATCTAATAAATGCGTCCCTTCCATAAGTCGGGGTTTGTTGCACGTATTAGCTCTAGAATTACTACGGTTATC | 22 (0.000416%) | | | CAACGTTAGGGAGTCCGGAGACGTCGGCGGGGGCCTCGGGAAGAGTTATCTTTTCTGTTTAACAGCCTGCCCACCCTGGAAACGGCTCAGCCGGAGGTAG | 4 (0.000076%) | | | CAACTTTCGATGGTAGGATAGTGGCCTACCATGGTGGTAACGGGTGACGGAGAATTAGGGTTCGATTCCGGAGAGGGAGCCTGAGAAACGGCTACCACAT | 22 (0.000416%) | | | CAAGGGCCGCCGGGGGCGCACCGGACACCACGCGACGTGCGGTGCTCTTCCAGCCGCTGGACCCTACCTCCGGCTGAGCCGTTTCCAGGGTGGGCAGGCT | 8 (0.000151%) | | | CAAGGGTTCACATTTCGTTCATCACCCTTGGCCGGCTATCGAACAGCCGGACTCCCATCAAAAGATGGTTGCCAAGAACATCTTCGTTACGGTTTGCTAA | 26 (0.000492%) | | | CAAGGGTTCACATTTCGTTCATCACCCTTGGCCGGCTTTCGAACAGCCGGACTCCCATCAAAAGATGGTTGCCAAGAACATCTTCGTTACGGTTTGCTAA | 22 (0.000416%) | | | CAAGTCATATTCGACTCCAAAACACTAACCAACCTTCTTCTTGCTTCTCAAAGCTTTCATGGTGTAGCCAAAGTCCATATGAGTCTTTGGCTTTGTGTCT | 150 (0.002838%) | | | CAAGTCTGGTGCCAGCAGCCGCGGTAATTCCAGCTCCAATAGCGTATATTTAAGTTGTTGCAGTTAAAAAGCTCGTAGTTGAACCTTGGGATGGGTCGGC | 21 (0.000397%) | | | CAATAGCGTATATTTAAGTTGTTGCAGTTAAAAAGCTCGTAGTTGAACCTTGGGATGGGTCGGCCGGTCCGCCTTTGGTGTGCATTGGTCGGCTTGTCCC | 9 (0.000170%) | | | CAATCAGCTTCCTTGCGCCTTACGGGTTTACTCACCCGTTGACTCGCACACATGTCAGACTCCTTGGTCCGTGTTTCAAGACGGGTCGAATGGGGAGCCC | 156 (0.002951%) | | | CAATCATACACATGACATCAAGTCATATTCGACTCCAAAACACTAACCAACCTTCTTCTTGCTTCTCAAAGCTTTCATGGTGTAGCCAAAGTCCATATGA | 142 (0.002686%) | | | CAATCGTTCCAACTAATCTACCGAAGTACTCGGCTAAGAAGAAAGAAGACGGACGAATCCGAGCCAAAGCCGTACAAGCGCGAGATACCTTCGGGACAGC | 3 (0.000057%) | | | CACAAACGTTCAATATGACAAACCCATGCCAAGTAAAGAGAAAATGAAAACTGGTGATTGTTGCGGAAATCGTCCAGGATTCCTCGACCAGGACTTGAAA | 16 (0.000303%) | | | CACAAAGCCAAAGACTCATATGGACTTTGGCTACACCATGAAAGCTTTGAGAAGCAAGAAGAAGGTTGGTTAGTGTTTTGGAGTCGAATATGACTTGATG | 46 (0.000870%) | | | CACAAAGGGTGTTGGTCGATTAAGACAGCAGGACGGTGGTCATGGAAGTCGAAATCCGCTAAGGAGTGTGTAACAACTCACCTGCCGAATCAACTAGCCC | 49 (0.000927%) | | | CACAAATCGTCGTCCCTCACCATCCTTTGCTGATGCGGGACGGAAGCTGGTCTCCCGTGTGTTACCGCACGCGGTTGGCCTAAATCCGAGCCAAGGACGC | 52 (0.000984%) | | | CACACATGTCAGACTCCTTGGTCCGTGTTTCAAGACGGGTCGAATGGGGAGCCCACAGGCCGACGCCCGGAGCACGCTGATGCCGAGGCACGCCGTTAGG | 7 (0.000132%) | | | CACAGGCCCCGGAACTCATCATCGAGCGTAACATCGCCCGTGAATTAACTGAGAAGGATAGGTGGTAGGTAGTTCGATGCGCGAGCATGGAGCCTACGAA | 26 (0.000492%) | | | CACATCTGTTAAAAGATAACGCAGGTGTCCTAAGATGAGCTCAACGAGAACAGAAATCTCGTGTGGAACAAAAGGGTAAAAGCTCGTTTGATTCTGATTT | 10 (0.000189%) | | | CACATGACATCAAGTCATATTCGACTCCAAAACACTAACCAACCTTCTTCTTGCTTCTCAAAGCTTTCATGGTGTAGCCAAAGTCCATATGAGTCTTTGG | 41 (0.000776%) | | | CACATTTCGTTCATCACCCTTGGCCGGCTATCGAACAGCCGGACTCCCATCAAAAGATGGTTGCCAAGAACATCTTCGTTACGGTTTGCTAATTCTCGGA | 9 (0.000170%) | | | CACATTTCGTTCATCACCCTTGGCCGGCTTTCGAACAGCCGGACTCCCATCAAAAGATGGTTGCCAAGAACATCTTCGTTACGGTTTGCTAATTCTCGGA | 7 (0.000132%) | | | CACCATCCTTTGCTGATGCGGGACGGAAGCTGGTCTCCCGTGTGTTACCGCACGCGGTTGGCCTAAATCCGAGCCAAGGACGCCTGGAGCGTACCGACAT | 5 (0.000095%) | | | CACCCGTTGACTCGCACACATGTCAGACTCCTTGGTCCGTGTTTCAAGACGGGTCGAATGGGGAGCCCACAGGCCGACGCCCGGAGCACGCTGATGCCGA | 12 (0.000227%) | | | CACCGGACACCACGCGACGTGCGGTGCTCTTCCAGCCGCTGGACCCTACCTCCGGCTGAGCCGTTTCCAGGGTGGGCAGGCTGTTAAACAGAAAAGATAA | 3 (0.000057%) | | | CACGCCCATCCAACCTAGGCGAGACAAGGGTTCACATTTCGTTCATCACCCTTGGCCGGCTATCGAACAGCCGGACTCCCATCAAAAGATGGTTGCCAAG | 21 (0.000397%) | | | CACGCGCCTAACGGCGTGCCTCGGCATCAGCGTGCTCCGGGCGTCGGCCTGTGGGCTCCCCATTCGACCCGTCTTGAAACACGGACCAAGGAGTCTGACA | 90 (0.001703%) | | | CACGCTTTCACGGTTCGTATTCGTACTGAAAATCAGAATCAAACGAGCTTTTACCCTTTTGTTCCACACGAGATTTCTGTTCTCGTTGAGCTCATCTTAG | 2001 (0.037855%) | | | CACGGTTCGTATTCGTACTGAAAATCAGAATCAAACGAGCTTTTACCCTTTTGTTCCACACGAGATTTCTGTTCTCGTTGAGCTCATCTTAGGACACCTG | 6 (0.000114%) | | | CACTTGGTGATATGAACACAAACGTTCAATATGACAAACCCATGCCAAGTAAAGAGAAAATGAAAACTGGTGATTGTTGCGGAAATCGTCCAGGATTCCT | 3 (0.000057%) | | | CAGAGCAACGGGCAGAGCCCGCGTCGACCTTTTATCTAATAAATGCGTCCCTTCCATAAGTCGGGGTTTGTTGCACGTATTAGCTCTAGAATTACTACGG | 9 (0.000170%) | | | CAGAGCCCGCGTCGACCTTTTATCTAATAAATGCGTCCCTTCCATAAGTCGGGGTTTGTTGCACGTATTAGCTCTAGAATTACTACGGTTATCCGAGTAG | 11 (0.000208%) | | | CAGAGCGTAGGCTTGCTTTGAGCACTCTAATTTCTTCAAAGTAACAGCGCCGGAGGCACGACCCGGCCAATTAAGACCAGGAGCGTATCGCCGACCGAAG | 34 (0.000643%) | | | CAGCACGCGCCTAACGGCGTGCCTCGGCATCAGCGTGCTCCGGGCGTCGGCCTGTGGGCTCCCCATTCGACCCGTCTTGAAACACGGACCAAGGAGTCTG | 74 (0.001400%) | | | CAGCAGCCGCGGTAATTCCAGCTCCAATAGCGTATATTTAAGTTGTTGCAGTTAAAAAGCTCGTAGTTGAACCTTGGGATGGGTCGGCCGGTCCGCCTTT | 26 (0.000492%) | | | CAGCAGGACGGTGGTCATGGAAGTCGAAATCCGCTAAGGAGTGTGTAACAACTCACCTGCCGAATCAACTAGCCCCGAAAATGGATGGCGCTTAAGCGCG | 5 (0.000095%) | | | CAGCAGTTCTCGGACAAAAATTGCTGAGTGGCCGAGAAGAATGGGCGTGTCATGCGTGGGCTGACATGGATTCTTCGAGGCCTAGGGGTGGCGGTATATA | 4 (0.000076%) | | | CAGCCGCGGTAATTCCAGCTCCAATAGCGTATATTTAAGTTGTTGCAGTTAAAAAGCTCGTAGTTGAACCTTGGGATGGGTCGGCCGGTCCGCCTTTGGT | 9 (0.000170%) | | | CAGCCTGCTAACTAGCTACGTGGAGGCATCCCTTCACGGCCGGCTTCTTAGAGGGACTATGGCCGTTTAGGCCAAGGAAGTTTGAGGCAATAACAGGTCT | 26 (0.000492%) | | | CAGCGTGCTCCGGGCGTCGGCCTGTGGGCTCCCCATTCGACCCGTCTTGAAACACGGACCAAGGAGTCTGACATGTGTGCGAGTCAACGGGTGAGTAAAC | 6 (0.000114%) | | | CAGCTCCAATAGCGTATATTTAAGTTGTTGCAGTTAAAAAGCTCGTAGTTGAACCTTGGGATGGGTCGGCCGGTCCGCCTTTGGTGTGCATTGGTCGGCT | 5 (0.000095%) | | | CAGCTTCCTTGCGCCTTACGGGTTTACTCACCCGTTGACTCGCACACATGTCAGACTCCTTGGTCCGTGTTTCAAGACGGGTCGAATGGGGAGCCCACAG | 108 (0.002043%) | | | CAGGACGGTGGTCATGGAAGTCGAAATCCGCTAAGGAGTGTGTAACAACTCACCTGCCGAATCAACTAGCCCCGAAAATGGATGGCGCTTAAGCGCGCGA | 21 (0.000397%) | | | CAGGGACAGTCGGGGGCATTCGTATTTCATAGTCAGAGGTGAAATTCTTGGATTTATGAAAGACGAACAACTGCGAAAGCATTTGCCAAGGATGTTTTCA | 34 (0.000643%) | | | CAGGGGTTGAAATCGTCGACCAGGTCCGAGACTTCATCGACCGGGTCCGAGGATTCGTCGACCAGGACGGCCGGATGTCCGAGAAAAAAAAATGTTGCCG | 18 (0.000341%) | | | CAGGTGGGGAGTTTGGCTGGGGCGGCACATCTGTTAAAAGATAACGCAGGTGTCCTAAGATGAGCTCAACGAGAACAGAAATCTCGTGTGGAACAAAAGG | 27 (0.000511%) | | | CAGTCGGGGGCATTCGTATTTCATAGTCAGAGGTGAAATTCTTGGATTTATGAAAGACGAACAACTGCGAAAGCATTTGCCAAGGATGTTTTCATTAATC | 9 (0.000170%) | | | CAGTTCTCGGACAAAAATTGCTGAGTGGCCGAGAAGAATGGGCGTGTCATGCGTGGGCTGACATGGATTCTTCGAGGCCTAGGGGTGGCGGTATATAACT | 5 (0.000095%) | | | CATACACATGACATCAAGTCATATTCGACTCCAAAACACTAACCAACCTTCTTCTTGCTTCTCAAAGCTTTCATGGTGTAGCCAAAGTCCATATGAGTCT | 26 (0.000492%) | | | CATATGAGTCTTTGGCTTTGTGTCTTCTAACAAGGATACA | 6766 (0.051200%) | | | CATATGCTTGTCTCAAAGATTAAGCCATGCATGTGTAAGTATGAACGAATTCAGACTGTGAAACTGCGAATGGCTCATTAAATCAGTTATAGTTTGTTTG | 47 (0.000889%) | | | CATATTCGACTCCAAAACACTAACCAACCTTCTTCTTGCTTCTCAAAGCTTTCATGGTGTAGCCAAAGTCCATATGAGTCTTTGGCTTTGTGTCTTCTAA | 75 (0.001419%) | | | CATATTCGACTCCAAAACACTAACCAACCTTCTTCTTGCTTCTCAAAGCTTTCATGGTGTAGCCAAAGTCCATATGAGTCTTTGGCTTTGTGTCTTTTAA | 320 (0.006054%) | | | CATCAAGTCATATTCGACTCCAAAACACTAACCAACCTTCTTCTTGCTTCTCAAAGCTTTCATGGTGTAGCCAAAGTCCATATGAGTCTTTGGCTTTGTG | 45 (0.000851%) | | | CATCAATCGTTCCAACTAATCTACCGAAGTACTCGGCTAAGAAGAAAGAAGACGGACGAATCCGAGCCAAAGCCGTACAAGCGCGAGATACCTTCGGGAC | 10 (0.000189%) | | | CATCAGAGCAACGGGCAGAGCCCGCGTCGACCTTTTATCTAATAAATGCGTCCCTTCCATAAGTCGGGGTTTGTTGCACGTATTAGCTCTAGAATTACTA | 47 (0.000889%) | | | CATCAGCGTGCTCCGGGCGTCGGCCTGTGGGCTCCCCATTCGACCCGTCTTGAAACACGGACCAAGGAGTCTGACATGTGTGCGAGTCAACGGGTGAGTA | 12 (0.000227%) | | | CATCATCGAGCGTAACATCGCCCGTGAATTAACTGAGAAGGATAGGTGGTAGGTAGTTCGATGCGCGAGCATGGAGCCTACGAACACTAGCTATCCGATC | 16 (0.000303%) | | | CATCATTCAAATTTCTGCCCTATCAACTTTCGATGGTAGGATAGTGGCCTACCATGGTGGTAACGGGTGACGGAGAATTAGGGTTCGATTCCGGAGAGGG | 25 (0.000473%) | | | CATCCAACCTAGGCGAGACAAGGGTTCACATTTCGTTCATCACCCTTGGCCGGCTTTCGAACAGCCGGACTCCCATCAAAAGATGGTTGCCAAGAACATC | 4 (0.000076%) | | | CATCCTTTGCTGATGCGGGACGGAAGCTGGTCTCCCGTGTGTTACCGCACGCGGTTGGCCTAAATCCGAGCCAAGGACGCCTGGAGCGTACCGACATGCG | 113 (0.002138%) | | | CATCGAACTACCTACCACCTATCCTTCTCAGTTAATTCACGGGCGATGTTACGCTCGATGATGAGTTCCGGGGCCTGTGTTTCGTACCTAATTTGAAGGA | 10 (0.000189%) | | | CATCGACACCTTGCGGCTAGGAACTGGAACGAGACGGGTGGCAAAGATTTCGAGTAGCACTTCATACTACCGTGGGTTTTTTAAACCTTCCGAGTTTTGT | 6 (0.000114%) | | | CATGAATCATCAGAGCAACGGGCAGAGCCCGCGTCGACCTTTTATCTAATAAATGCGTCCCTTCCATAAGTCGGGGTTTGTTGCACGTATTAGCTCTAGA | 169 (0.003197%) | | | CATGACATCAAGTCATATTCGACTCCAAAACACTAACCAACCTTCTTCTTGCTTCTCAAAGCTTTCATGGTGTAGCCAAAGTCCATATGAGTCTTTGGCT | 39 (0.000738%) | | | CATGCTAATGTATCCAGAGCGTAGGCTTGCTTTGAGCACTCTAATTTCTTCAAAGTAACAGCGCCGGAGGCACGACCCGGCCAATTAAGACCAGGAGCGT | 4 (0.000076%) | | | CATGGAAGTCGAAATCCGCTAAGGAGTGTGTAACAACTCACCTGCCGAATCAACTAGCCCCGAAAATGGATGGCGCTTAAGCGCGCGACCTATACCCGGC | 49 (0.000927%) | | | CATGGCCTCTGTGCTGGCGACGCATCATTCAAATTTCTGCCCTATCAACTTTCGATGGTAGGATAGTGGCCTACCATGGTGGTAACGGGTGACGGAGAAT | 257 (0.004862%) | | | CATGTGTATGATTGAGTATAAGAACTTAAACCGCAACCGCATCTTATAAGCCTAAGTAGTGTTTCCTTGTTAGAAGACACAAAGCCAAAGACTCATATGG | 17 (0.000322%) | | | CATTCAAATTTCTGCCCTATCAACTTTCGATGGTAGGATAGTGGCCTACCATGGTGGTAACGGGTGACGGAGAATTAGGGTTCGATTCCGGAGAGGGAGC | 8 (0.000151%) | | | CATTGTCAGGTGGGGAGTTTGGCTGGGGCGGCACATCTGTTAAAAGATAACGCAGGTGTCCTAAGATGAGCTCAACGAGAACAGAAATCTCGTGTGGAAC | 11 (0.000208%) | | | CATTGTTCCATCGACCAGAGGCTGTTCACCTTGGAGACCTGATGCGGTTATGAGTACGACCGGGCGTGAGCGGCACTCGGTCCTCCGGATTTTCAAGGGC | 26 (0.000492%) | | | CCAAAGACTCATATGGACTTTGGCTACACCATGAAAGCTTTGAGAAGCAAGAAGAAGGTTGGTTAGTGTTTTGGAGTCGAATATGACTTGATGTCATGTG | 131 (0.002478%) | | | CCAACCTAGGCGAGACAAGGGTTCACATTTCGTTCATCACCCTTGGCCGGCTATCGAACAGCCGGACTCCCATCAAAAGATGGTTGCCAAGAACATCTTC | 23 (0.000435%) | | | CCAACGAAGCACGCCCATCCAACCTAGGCGAGACAAGGGTTCACATTTCGTTCATCACCCTTGGCCGGCTATCGAACAGCCGGACTCCCATCAAAAGATG | 6 (0.000114%) | | | CCAATCAGCTTCCTTGCGCCTTACGGGTTTACTCACCCGTTGACTCGCACACATGTCAGACTCCTTGGTCCGTGTTTCAAGACGGGTCGAATGGGGAGCC | 14 (0.000265%) | | | CCACAAAGGGTGTTGGTCGATTAAGACAGCAGGACGGTGGTCATGGAAGTCGAAATCCGCTAAGGAGTGTGTAACAACTCACCTGCCGAATCAACTAGCC | 6 (0.000114%) | | | CCAGAGCGTAGGCTTGCTTTGAGCACTCTAATTTCTTCAAAGTAACAGCGCCGGAGGCACGACCCGGCCAATTAAGACCAGGAGCGTATCGCCGACCGAA | 17 (0.000322%) | | | CCAGCAGCCGCGGTAATTCCAGCTCCAATAGCGTATATTTAAGTTGTTGCAGTTAAAAAGCTCGTAGTTGAACCTTGGGATGGGTCGGCCGGTCCGCCTT | 9 (0.000170%) | | | CCAGCTCCAATAGCGTATATTTAAGTTGTTGCAGTTAAAAAGCTCGTAGTTGAACCTTGGGATGGGTCGGCCGGTCCGCCTTTGGTGTGCATTGGTCGGC | 14 (0.000265%) | | | CCAGGGGTTGAAATCGTCGACCAGGTCCGAGACTTCATCGACCGGGTCCGAGGATTCGTCGACCAGGACGGCCGGATGTCCGAGAAAAAAAAATGTTGCC | 3 (0.000057%) | | | CCATCCAACCTAGGCGAGACAAGGGTTCACATTTCGTTCATCACCCTTGGCCGGCTTTCGAACAGCCGGACTCCCATCAAAAGATGGTTGCCAAGAACAT | 3 (0.000057%) | | | CCATCCTTTGCTGATGCGGGACGGAAGCTGGTCTCCCGTGTGTTACCGCACGCGGTTGGCCTAAATCCGAGCCAAGGACGCCTGGAGCGTACCGACATGC | 14 (0.000265%) | | | CCATGCTAATGTATCCAGAGCGTAGGCTTGCTTTGAGCACTCTAATTTCTTCAAAGTAACAGCGCCGGAGGCACGACCCGGCCAATTAAGACCAGGAGCG | 14 (0.000265%) | | | CCATGCTCGCGCATCGAACTACCTACCACCTATCCTTCTCAGTTAATTCACGGGCGATGTTACGCTCGATGATGAGTTCCGGGGCCTGTGTTTCGTACCT | 4 (0.000076%) | | | CCATGGGCATCGACACCTTGCGGCTAGGAACTGGAACGAGACGGGTGGCAAAGATTTCGAGTAGCACTTCATACTACCGTGGGTTTTTTAAACCTTCCGA | 10 (0.000189%) | | | CCCACAAAGGGTGTTGGTCGATTAAGACAGCAGGACGGTGGTCATGGAAGTCGAAATCCGCTAAGGAGTGTGTAACAACTCACCTGCCGAATCAACTAGC | 9 (0.000170%) | | | CCCATCCAACCTAGGCGAGACAAGGGTTCACATTTCGTTCATCACCCTTGGCCGGCTATCGAACAGCCGGACTCCCATCAAAAGATGGTTGCCAAGAACA | 17 (0.000322%) | | | CCCATCCAACCTAGGCGAGACAAGGGTTCACATTTCGTTCATCACCCTTGGCCGGCTTTCGAACAGCCGGACTCCCATCAAAAGATGGTTGCCAAGAACA | 15 (0.000284%) | | | CCCATGCTAATGTATCCAGAGCGTAGGCTTGCTTTGAGCACTCTAATTTCTTCAAAGTAACAGCGCCGGAGGCACGACCCGGCCAATTAAGACCAGGAGC | 17 (0.000322%) | | | CCCATGGGCATCGACACCTTGCGGCTAGGAACTGGAACGAGACGGGTGGCAAAGATTTCGAGTAGCACTTCATACTACCGTGGGTTTTTTAAACCTTCCG | 7 (0.000132%) | | | CCCGAAGGTATCTCGCGCTTGTACGGCTTTGGCTCGGATTCGTCCGTCTTCTTTCTTCTTAGCCGAGTACTTCGGTAGATTAGTTGGAACGATTGATGAT | 48 (0.000908%) | | | CCCGAGTGTGAGCGAGGTGTGAGTGTCGCCCATGGGCATCGACACCTTGCGGCTAGGAACTGGAACGAGACGGGTAGCAAAGATTTCGAGTAGCACTTCA | 14 (0.000265%) | | | CCCGAGTGTGAGCGAGGTGTGAGTGTCGCCCATGGGCATCGACACCTTGCGGCTAGGAACTGGAACGAGACGGGTGGCAAAGATTTCGAGTAGCACTTCA | 19 (0.000359%) | | | CCCGCCAATCAGCTTCCTTGCGCCTTACGGGTTTACTCACCCGTTGACTCGCACACATGTCAGACTCCTTGGTCCGTGTTTCAAGACGGGTCGAATGGGG | 14 (0.000265%) | | | CCCGCCGAAGCGAGCCTTGGGACCAAAAACAGGGGTTGTACCCCGCCTCCGATTCACGGAGTAAGTAAAATAACGTTAAAAGTAGTGGTATTTCACTTGC | 6 (0.000114%) | | | CCCGCGTCGACCTTTTATCTAATAAATGCGTCCCTTCCATAAGTCGGGGTTTGTTGCACGTATTAGCTCTAGAATTACTACGGTTATCCGAGTAGTAGTT | 36 (0.000681%) | | | CCCTCACCATCCTTTGCTGATGCGGGACGGAAGCTGGTCTCCCGTGTGTTACCGCACGCGGTTGGCCTAAATCCGAGCCAAGGACGCCTGGAGCGTACCG | 19 (0.000359%) | | | CCCTTGCCTACATTGTTCCATCGACCAGAGGCTGTTCACCTTGGAGACCTGATGCGGTTATGAGTACGACCGGGCGTGAGCGGCACTCGGTCCTCCGGAT | 25 (0.000473%) | | | CCGAAGCGAGCCTTGGGACCAAAAACAGGGGTTGTACCCCGCCTCCGATTCACGGAGTAAGTAAAATAACGTTAAAAGTAGTGGTATTTCACTTGCGCCG | 12 (0.000227%) | | | CCGAAGGTATCTCGCGCTTGTACGGCTTTGGCTCGGATTCGTCCGTCTTCTTTCTTCTTAGCCGAGTACTTCGGTAGATTAGTTGGAACGATTGATGATT | 6 (0.000114%) | | | CCGACTTCCCTTGCCTACATTGTTCCATCGACCAGAGGCTGTTCACCTTGGAGACCTGATGCGGTTATGAGTACGACCGGGCGTGAGCGGCACTCGGTCC | 49 (0.000927%) | | | CCGAGTGTGAGCGAGGTGTGAGTGTCGCCCATGGGCATCGACACCTTGCGGCTAGGAACTGGAACGAGACGGGTAGCAAAGATTTCGAGTAGCACTTCAT | 12 (0.000227%) | | | CCGAGTGTGAGCGAGGTGTGAGTGTCGCCCATGGGCATCGACACCTTGCGGCTAGGAACTGGAACGAGACGGGTGGCAAAGATTTCGAGTAGCACTTCAT | 18 (0.000341%) | | | CCGCCAATCAGCTTCCTTGCGCCTTACGGGTTTACTCACCCGTTGACTCGCACACATGTCAGACTCCTTGGTCCGTGTTTCAAGACGGGTCGAATGGGGA | 57 (0.001078%) | | | CCGCCGGGGGCGCACCGGACACCACGCGACGTGCGGTGCTCTTCCAGCCGCTGGACCCTACCTCCGGCTGAGCCGTTTCCAGGGTGGGCAGGCTGTTAAA | 5 (0.000095%) | | | CCGCCTAGGCTGTCCCGAGTGTGAGCGAGGTGTGAGTGTCGCCCATGGGCATCGACACCTTGCGGCTAGGAACTGGAACGAGACGGGTGGCAAAGATTTC | 69 (0.001305%) | | | CCGCGGTAATTCCAGCTCCAATAGCGTATATTTAAGTTGTTGCAGTTAAAAAGCTCGTAGTTGAACCTTGGGATGGGTCGGCCGGTCCGCCTTTGGTGTG | 9 (0.000170%) | | | CCGGAACCGGGACGTGGCGGTTGACGGCAACGTTAGGGAGTCCGGAGACGTCGGCGGGGGCCTCGGGAAGAGTTATCTTTTCTGTTTAACAGCCTGCCCA | 110 (0.002081%) | | | CCGGACACCACGCGACGTGCGGTGCTCTTCCAGCCGCTGGACCCTACCTCCGGCTGAGCCGTTTCCAGGGTGGGCAGGCTGTTAAACAGAAAAGATAACT | 27 (0.000511%) | | | CCGGAGACGTCGGCGGGGGCCTCGGGAAGAGTTATCTTTTCTGTTTAACAGCCTGCCCACCCTGGAAACGGCTCAGCCGGAGGTAGGGTCCAGCGGCTGG | 7 (0.000132%) | | | CCGGATTTTCAAGGGCCGCCGGGGGCGCACCGGACACCACGCGACGTGCGGTGCTCTTCCAGCCGCTGGACCCTACCTCCGGCTGAGCCGTTTCCAGGGT | 5 (0.000095%) | | | CCGGGACGTGGCGGTTGACGGCAACGTTAGGGAGTCCGGAGACGTCGGCGGGGGCCTCGGGAAGAGTTATCTTTTCTGTTTAACAGCCTGCCCACCCTGG | 32 (0.000605%) | | | CCGGGCGTCGGCCTGTGGGCTCCCCATTCGACCCGTCTTGAAACACGGACCAAGGAGTCTGACATGTGTGCGAGTCAACGGGTGAGTAAACCCGTAAGGC | 4 (0.000076%) | | | CCGGGGGCGCACCGGACACCACGCGACGTGCGGTGCTCTTCCAGCCGCTGGACCCTACCTCCGGCTGAGCCGTTTCCAGGGTGGGCAGGCTGTTAAACAG | 6 (0.000114%) | | | CCGTCGAGTTATCATGAATCATCAGAGCAACGGGCAGAGCCCGCGTCGACCTTTTATCTAATAAATGCGTCCCTTCCATAAGTCGGGGTTTGTTGCACGT | 20 (0.000378%) | | | CCGTTGACTCGCACACATGTCAGACTCCTTGGTCCGTGTTTCAAGACGGGTCGAATGGGGAGCCCACAGGCCGACGCCCGGAGCACGCTGATGCCGAGGC | 3 (0.000057%) | | | CCTAACGGCGTGCCTCGGCATCAGCGTGCTCCGGGCGTCGGCCTGTGGGCTCCCCATTCGACCCGTCTTGAAACACGGACCAAGGAGTCTGACATGTGTG | 8 (0.000151%) | | | CCTAAGTAGTGTTTCCTTGTTAGAAGACACAAAGCCAAAGACTCATATGGACTTTGGCTACACCATGAAAGCTTTGAGAAGCAAGAAGAAGGTTGGTTAG | 59 (0.001116%) | | | CCTACATTGTTCCATCGACCAGAGGCTGTTCACCTTGGAGACCTGATGCGGTTATGAGTACGACCGGGCGTGAGCGGCACTCGGTCCTCCGGATTTTCAA | 9 (0.000170%) | | | CCTAGGCGAGACAAGGGTTCACATTTCGTTCATCACCCTTGGCCGGCTATCGAACAGCCGGACTCCCATCAAAAGATGGTTGCCAAGAACATCTTCGTTA | 8 (0.000151%) | | | CCTAGGCTGTCCCGAGTGTGAGCGAGGTGTGAGTGTCGCCCATGGGCATCGACACCTTGCGGCTAGGAACTGGAACGAGACGGGTGGCAAAGATTTCGAG | 48 (0.000908%) | | | CCTATGATGTTATCCCATGCTAATGTATCCAGAGCGTAGGCTTGCTTTGAGCACTCTAATTTCTTCAAAGTAACAGCGCCGGAGGCACGACCCGGCCAAT | 43 (0.000813%) | | | CCTCACCATCCTTTGCTGATGCGGGACGGAAGCTGGTCTCCCGTGTGTTACCGCACGCGGTTGGCCTAAATCCGAGCCAAGGACGCCTGGAGCGTACCGA | 78 (0.001476%) | | | CCTCAGCCTGCTAACTAGCTACGTGGAGGCATCCCTTCACGGCCGGCTTCTTAGAGGGACTATGGCCGTTTAGGCCAAGGAAGTTTGAGGCAATAACAGG | 107 (0.002024%) | | | CCTCCGGATTTTCAAGGGCCGCCGGGGGCGCACCGGACACCACGCGACGTGCGGTGCTCTTCCAGCCGCTGGACCCTACCTCCGGCTGAGCCGTTTCCAG | 3 (0.000057%) | | | CCTCGGCATCAGCGTGCTCCGGGCGTCGGCCTGTGGGCTCCCCATTCGACCCGTCTTGAAACACGGACCAAGGAGTCTGACATGTGTGCGAGTCAACGGG | 4 (0.000076%) | | | CCTCTGTGCTGGCGACGCATCATTCAAATTTCTGCCCTATCAACTTTCGATGGTAGGATAGTGGCCTACCATGGTGGTAACGGGTGACGGAGAATTAGGG | 106 (0.002005%) | | | CCTTGCCTACATTGTTCCATCGACCAGAGGCTGTTCACCTTGGAGACCTGATGCGGTTATGAGTACGACCGGGCGTGAGCGGCACTCGGTCCTCCGGATT | 10 (0.000189%) | | | CCTTGCGCCTTACGGGTTTACTCACCCGTTGACTCGCACACATGTCAGACTCCTTGGTCCGTGTTTCAAGACGGGTCGAATGGGGAGCCCACAGGCCGAC | 7 (0.000132%) | | | CCTTGTTAGAAGACACAAAGCCAAAGACTCATATGGACTTTGGCTACACCATGAAAGCTTTGAGAAGCAAGAAGAAGGTTGGTTAGTGTTTTGGAGTCGA | 46 (0.000870%) | | | CCTTTGCTGATGCGGGACGGAAGCTGGTCTCCCGTGTGTTACCGCACGCGGTTGGCCTAAATCCGAGCCAAGGACGCCTGGAGCGTACCGACATGCGGTG | 4 (0.000076%) | | | CGAAACACAGGCCCCGGAACTCATCATCGAGCGTAACATCGCCCGTGAATTAACTGAGAAGGATAGGTGGTAGGTAGTTCGATGCGCGAGCATGGAGCCT | 37 (0.000700%) | | | CGAAATCCTATGATGTTATCCCATGCTAATGTATCCAGAGCGTAGGCTTGCTTTGAGCACTCTAATTTCTTCAAAGTAACAGCGCCGGAGGCACGACCCG | 444 (0.008400%) | | | CGAACTACCTACCACCTATCCTTCTCAGTTAATTCACGGGCGATGTTACGCTCGATGATGAGTTCCGGGGCCTGTGTTTCGTACCTAATTTGAAGGAATT | 5 (0.000095%) | | | CGAAGCACGCCCATCCAACCTAGGCGAGACAAGGGTTCACATTTCGTTCATCACCCTTGGCCGGCTATCGAACAGCCGGACTCCCATCAAAAGATGGTTG | 27 (0.000511%) | | | CGAAGGTATCTCGCGCTTGTACGGCTTTGGCTCGGATTCGTCCGTCTTCTTTCTTCTTAGCCGAGTACTTCGGTAGATTAGTTGGAACGATTGATGATTT | 19 (0.000359%) | | | CGACCAGGGGTTGAAATCGTCGACCAGGTCCGAGACTTCATCGACCGGGTCCGAGGATTCGTCGACCAGGACGGCCGGATGTCCGAGAAAAAAAAATGTT | 27 (0.000511%) | | | CGACGCATCATTCAAATTTCTGCCCTATCAACTTTCGATGGTAGGATAGTGGCCTACCATGGTGGTAACGGGTGACGGAGAATTAGGGTTCGATTCCGGA | 23 (0.000435%) | | | CGACGGGCGGTGTGTACAAAGGGCAGGGACGTAGTCAACGCGAGCTGATGACTCGCGCTTACTAGGAATTCCTCGTTGAAGACCAACAATTGCAATGATC | 16 (0.000303%) | | | CGACGTGGGTGGTTCGCCGCCCGCGACGTCGCGAGAAGTCCACTAAACCTTATCATTTAGAGGAAGGAGAAGTCGTAACAAGGTTTCCGTAGGTGAACCT | 124 (0.002346%) | | | CGACTCCAAAACACTAACCAACCTTCTTCTTGCTTCTCAAAGCTTTCATGGTGTAGCCAAAGTCCATATGAGTCTTTGGCTTTGTGTCTTCTAACAAGGA | 58 (0.001097%) | | | CGACTCCAAAACACTAACCAACCTTCTTCTTGCTTCTCAAAGCTTTCATGGTGTAGCCAAAGTCCATATGAGTCTTTGGCTTTGTGTCTTTTAACAAGGA | 30 (0.000568%) | | | CGACTTCCCTTGCCTACATTGTTCCATCGACCAGAGGCTGTTCACCTTGGAGACCTGATGCGGTTATGAGTACGACCGGGCGTGAGCGGCACTCGGTCCT | 24 (0.000454%) | | | CGAGACAAGGGTTCACATTTCGTTCATCACCCTTGGCCGGCTATCGAACAGCCGGACTCCCATCAAAAGATGGTTGCCAAGAACATCTTCGTTACGGTTT | 6 (0.000114%) | | | CGAGACAAGGGTTCACATTTCGTTCATCACCCTTGGCCGGCTTTCGAACAGCCGGACTCCCATCAAAAGATGGTTGCCAAGAACATCTTCGTTACGGTTT | 7 (0.000132%) | | | CGAGCTCGTGTAAAGTTGGGAATTCGTTAAGGAGCTGTTGCTTTGTTAGTGTAGAAACACTTGTGTAGAATTGGGGATTGTTTTTTTTGGAGTGATTTAG | 741 (0.014018%) | | | CGAGGTGTGAGTGTCGCCCATGGGCATCGACACCTTGCGGCTAGGAACTGGAACGAGACGGGTGGCAAAGATTTCGAGTAGCACTTCATACTACCGTGGG | 22 (0.000416%) | | | CGAGTGTGAGCATGCCTGTCGGGACCCGAAAGATGGTGAACTATGCCTGAGCGGGGTAAAGCCAGAGGAAACTCTGGTGGAAGCCCGCAGCGATACTGAC | 4 (0.000076%) | | | CGAGTGTGAGCGAGGTGTGAGTGTCGCCCATGGGCATCGACACCTTGCGGCTAGGAACTGGAACGAGACGGGTAGCAAAGATTTCGAGTAGCACTTCATA | 23 (0.000435%) | | | CGAGTGTGAGCGAGGTGTGAGTGTCGCCCATGGGCATCGACACCTTGCGGCTAGGAACTGGAACGAGACGGGTGGCAAAGATTTCGAGTAGCACTTCATA | 34 (0.000643%) | | | CGAGTTATCATGAATCATCAGAGCAACGGGCAGAGCCCGCGTCGACCTTTTATCTAATAAATGCGTCCCTTCCATAAGTCGGGGTTTGTTGCACGTATTA | 131 (0.002478%) | | | CGATCCATTACATTTTATCGGTCGCTCTTGTCCGGAAGCTGTAGATGACCCAAAGTCCATATAGCGACCCCAGGTCAGGCGGGATTACCCGCTGAGTTTA | 6 (0.000114%) | | | CGATCCCGCCAATCAGCTTCCTTGCGCCTTACGGGTTTACTCACCCGTTGACTCGCACACATGTCAGACTCCTTGGTCCGTGTTTCAAGACGGGTCGAAT | 4 (0.000076%) | | | CGATCCGTCGAGTTATCATGAATCATCAGAGCAACGGGCAGAGCCCGCGTCGACCTTTTATCTAATAAATGCGTCCCTTCCATAAGTCGGGGTTTGTTGC | 163 (0.003084%) | | | CGATCGACCCGCCGAAGCGAGCCTTGGGACCAAAAACAGGGGTTGTACCCCGCCTCCGATTCACGGAGTAAGTAAAATAACGTTAAAAGTAGTGGTATTT | 3 (0.000057%) | | | CGATTAAGACAGCAGGACGGTGGTCATGGAAGTCGAAATCCGCTAAGGAGTGTGTAACAACTCACCTGCCGAATCAACTAGCCCCGAAAATGGATGGCGC | 14 (0.000265%) | | | CGATTAGTCTTTCGCCCCTATACCCAAGTCAGACGAACGATTTGCACGTCAGTATCGCTGCGGGCTTCCACCAGAGTTTCCTCTGGCTTTACCCCGCTCA | 11 (0.000208%) | | | CGATTTGCACGTCAGTATCGCTGCGGGCTTCCACCAGAGTTTCCTCTGGCTTTACCCCGCTCAGGCATAGTTCACCATCTTTCGGGTCCCGACAGGCATG | 503 (0.009516%) | | | CGCACACATGTCAGACTCCTTGGTCCGTGTTTCAAGACGGGTCGAATGGGGAGCCCACAGGCCGACGCCCGGAGCACGCTGATGCCGAGGCACGCCGTTA | 20 (0.000378%) | | | CGCACCGGACACCACGCGACGTGCGGTGCTCTTCCAGCCGCTGGACCCTACCTCCGGCTGAGCCGTTTCCAGGGTGGGCAGGCTGTTAAACAGAAAAGAT | 4 (0.000076%) | | | CGCATCATTCAAATTTCTGCCCTATCAACTTTCGATGGTAGGATAGTGGCCTACCATGGTGGTAACGGGTGACGGAGAATTAGGGTTCGATTCCGGAGAG | 4 (0.000076%) | | | CGCCAATCAGCTTCCTTGCGCCTTACGGGTTTACTCACCCGTTGACTCGCACACATGTCAGACTCCTTGGTCCGTGTTTCAAGACGGGTCGAATGGGGAG | 1196 (0.022626%) | | | CGCCCATCCAACCTAGGCGAGACAAGGGTTCACATTTCGTTCATCACCCTTGGCCGGCTATCGAACAGCCGGACTCCCATCAAAAGATGGTTGCCAAGAA | 76 (0.001438%) | | | CGCCCATCCAACCTAGGCGAGACAAGGGTTCACATTTCGTTCATCACCCTTGGCCGGCTTTCGAACAGCCGGACTCCCATCAAAAGATGGTTGCCAAGAA | 89 (0.001684%) | | | CGCCCATGGGCATCGACACCTTGCGGCTAGGAACTGGAACGAGACGGGTGGCAAAGATTTCGAGTAGCACTTCATACTACCGTGGGTTTTTTAAACCTTC | 30 (0.000568%) | | | CGCCGAAGCGAGCCTTGGGACCAAAAACAGGGGTTGTACCCCGCCTCCGATTCACGGAGTAAGTAAAATAACGTTAAAAGTAGTGGTATTTCACTTGCGC | 4 (0.000076%) | | | CGCCGCCCGCGACGTCGCGAGAAGTCCACTAAACCTTATCATTTAGAGGAAGGAGAAGTCGTAACAAGGTTTCCGTAGGTGAACCTGCGGAAGGATCGAT | 16 (0.000303%) | | | CGCCGGGGGCGCACCGGACACCACGCGACGTGCGGTGCTCTTCCAGCCGCTGGACCCTACCTCCGGCTGAGCCGTTTCCAGGGTGGGCAGGCTGTTAAAC | 17 (0.000322%) | | | CGCCTAACGGCGTGCCTCGGCATCAGCGTGCTCCGGGCGTCGGCCTGTGGGCTCCCCATTCGACCCGTCTTGAAACACGGACCAAGGAGTCTGACATGTG | 30 (0.000568%) | | | CGCCTAGGCTGTCCCGAGTGTGAGCGAGGTGTGAGTGTCGCCCATGGGCATCGACACCTTGCGGCTAGGAACTGGAACGAGACGGGTGGCAAAGATTTCG | 183 (0.003462%) | | | CGCCTTACGGGTTTACTCACCCGTTGACTCGCACACATGTCAGACTCCTTGGTCCGTGTTTCAAGACGGGTCGAATGGGGAGCCCACAGGCCGACGCCCG | 9 (0.000170%) | | | CGCGCATCGAACTACCTACCACCTATCCTTCTCAGTTAATTCACGGGCGATGTTACGCTCGATGATGAGTTCCGGGGCCTGTGTTTCGTACCTAATTTGA | 12 (0.000227%) | | | CGCGCCTAACGGCGTGCCTCGGCATCAGCGTGCTCCGGGCGTCGGCCTGTGGGCTCCCCATTCGACCCGTCTTGAAACACGGACCAAGGAGTCTGACATG | 23 (0.000435%) | | | CGCGCTTGTACGGCTTTGGCTCGGATTCGTCCGTCTTCTTTCTTCTTAGCCGAGTACTTCGGTAGATTAGTTGGAACGATTGATGATTTTGAGTTAATTG | 26 (0.000492%) | | | CGCGGTAATTCCAGCTCCAATAGCGTATATTTAAGTTGTTGCAGTTAAAAAGCTCGTAGTTGAACCTTGGGATGGGTCGGCCGGTCCGCCTTTGGTGTGC | 11 (0.000208%) | | | CGCTTGTACGGCTTTGGCTCGGATTCGTCCGTCTTCTTTCTTCTTAGCCGAGTACTTCGGTAGATTAGTTGGAACGATTGATGATTTTGAGTTAATTGAA | 5 (0.000095%) | | | CGCTTTCACGGTTCGTATTCGTACTGAAAATCAGAATCAAACGAGCTTTTACCCTTTTGTTCCACACGAGATTTCTGTTCTCGTTGAGCTCATCTTAGGA | 9 (0.000170%) | | | CGGAACCGGGACGTGGCGGTTGACGGCAACGTTAGGGAGTCCGGAGACGTCGGCGGGGGCCTCGGGAAGAGTTATCTTTTCTGTTTAACAGCCTGCCCAC | 90 (0.001703%) | | | CGGAAGAGCGTCGTGTAGGGAAAGAGTGTTCAGAGCCGTG | 9 (0.000068%) | | | CGGACACCACGCGACGTGCGGTGCTCTTCCAGCCGCTGGACCCTACCTCCGGCTGAGCCGTTTCCAGGGTGGGCAGGCTGTTAAACAGAAAAGATAACTC | 36 (0.000681%) | | | CGGAGACGTCGGCGGGGGCCTCGGGAAGAGTTATCTTTTCTGTTTAACAGCCTGCCCACCCTGGAAACGGCTCAGCCGGAGGTAGGGTCCAGCGGCTGGA | 41 (0.000776%) | | | CGGAGGACATTGTCAGGTGGGGAGTTTGGCTGGGGCGGCACATCTGTTAAAAGATAACGCAGGTGTCCTAAGATGAGCTCAACGAGAACAGAAATCTCGT | 53 (0.001003%) | | | CGGATAGCTAGTGTTCGTAGGCTCCATGCTCGCGCATCGAACTACCTACCACCTATCCTTCTCAGTTAATTCACGGGCGATGTTACGCTCGATGATGAGT | 695 (0.013148%) | | | CGGATTTTCAAGGGCCGCCGGGGGCGCACCGGACACCACGCGACGTGCGGTGCTCTTCCAGCCGCTGGACCCTACCTCCGGCTGAGCCGTTTCCAGGGTG | 38 (0.000719%) | | | CGGCAACGTTAGGGAGTCCGGAGACGTCGGCGGGGGCCTCGGGAAGAGTTATCTTTTCTGTTTAACAGCCTGCCCACCCTGGAAACGGCTCAGCCGGAGG | 30 (0.000568%) | | | CGGCAATTCCCCGCCACATCCTCTCAAACGCAATGGAAAGAGAGAAAGGACGAGGTCTTGACCGTCATCTTTTGCCCGAAGGACGGATGAGCTTTGGCGG | 1085 (0.020526%) | | | CGGCACATCTGTTAAAAGATAACGCAGGTGTCCTAAGATGAGCTCAACGAGAACAGAAATCTCGTGTGGAACAAAAGGGTAAAAGCTCGTTTGATTCTGA | 11 (0.000208%) | | | CGGCACTCGGTCCTCCGGATTTTCAAGGGCCGCCGGGGGCGCACCGGACACCACGCGACGTGCGGTGCTCTTCCAGCCGCTGGACCCTACCTCCGGCTGA | 60 (0.001135%) | | | CGGCATCAGCGTGCTCCGGGCGTCGGCCTGTGGGCTCCCCATTCGACCCGTCTTGAAACACGGACCAAGGAGTCTGACATGTGTGCGAGTCAACGGGTGA | 14 (0.000265%) | | | CGGCCTGTGGGCTCCCCATTCGACCCGTCTTGAAACACGGACCAAGGAGTCTGACATGTGTGCGAGTCAACGGGTGAGTAAACCCGTAAGGCGCAAGGAA | 9 (0.000170%) | | | CGGCGACGTGGGTGGTTCGCCGCCCGCGACGTCGCGAGAAGTCCACTAAACCTTATCATTTAGAGGAAGGAGAAGTCGTAACAAGGTTTCCGTAGGTGAA | 464 (0.008778%) | | | CGGCGTGCCTCGGCATCAGCGTGCTCCGGGCGTCGGCCTGTGGGCTCCCCATTCGACCCGTCTTGAAACACGGACCAAGGAGTCTGACATGTGTGCGAGT | 14 (0.000265%) | | | CGGGACGTGGCGGTTGACGGCAACGTTAGGGAGTCCGGAGACGTCGGCGGGGGCCTCGGGAAGAGTTATCTTTTCTGTTTAACAGCCTGCCCACCCTGGA | 41 (0.000776%) | | | CGGGCAGAGCCCGCGTCGACCTTTTATCTAATAAATGCGTCCCTTCCATAAGTCGGGGTTTGTTGCACGTATTAGCTCTAGAATTACTACGGTTATCCGA | 51 (0.000965%) | | | CGGGCGAGAGACCGATAGCGAACAAGTACCGCGAGGTAAAGATGAAAAGGACTTTGAAAAGAGAGTCAAAGAGTGCTTGAAATTGTCGGGAGGGAAGCGG | 7 (0.000132%) | | | CGGGCGGTGTGTACAAAGGGCAGGGACGTAGTCAACGCGAGCTGATGACTCGCGCTTACTAGGAATTCCTCGTTGAAGACCAACAATTGCAATGATCGAT | 33 (0.000624%) | | | CGGGCGTCGGCCTGTGGGCTCCCCATTCGACCCGTCTTGAAACACGGACCAAGGAGTCTGACATGTGTGCGAGTCAACGGGTGAGTAAACCCGTAAGGCG | 32 (0.000605%) | | | CGGGGGCATTCGTATTTCATAGTCAGAGGTGAAATTCTTGGATTTATGAAAGACGAACAACTGCGAAAGCATTTGCCAAGGATGTTTTCATTAATCAAGA | 37 (0.000700%) | | | CGGGGGCGCACCGGACACCACGCGACGTGCGGTGCTCTTCCAGCCGCTGGACCCTACCTCCGGCTGAGCCGTTTCCAGGGTGGGCAGGCTGTTAAACAGA | 39 (0.000738%) | | | CGGGTTTACTCACCCGTTGACTCGCACACATGTCAGACTCCTTGGTCCGTGTTTCAAGACGGGTCGAATGGGGAGCCCACAGGCCGACGCCCGGAGCACG | 22 (0.000416%) | | | CGGTAATTCCAGCTCCAATAGCGTATATTTAAGTTGTTGCAGTTAAAAAGCTCGTAGTTGAACCTTGGGATGGGTCGGCCGGTCCGCCTTTGGTGTGCAT | 16 (0.000303%) | | | CGGTAGGAGCGACGGGCGGTGTGTACAAAGGGCAGGGACGTAGTCAACGCGAGCTGATGACTCGCGCTTACTAGGAATTCCTCGTTGAAGACCAACAATT | 65 (0.001230%) | | | CGGTCCTCCGGATTTTCAAGGGCCGCCGGGGGCGCACCGGACACCACGCGACGTGCGGTGCTCTTCCAGCCGCTGGACCCTACCTCCGGCTGAGCCGTTT | 12 (0.000227%) | | | CGGTGGTCATGGAAGTCGAAATCCGCTAAGGAGTGTGTAACAACTCACCTGCCGAATCAACTAGCCCCGAAAATGGATGGCGCTTAAGCGCGCGACCTAT | 21 (0.000397%) | | | CGGTTGACGGCAACGTTAGGGAGTCCGGAGACGTCGGCGGGGGCCTCGGGAAGAGTTATCTTTTCTGTTTAACAGCCTGCCCACCCTGGAAACGGCTCAG | 6 (0.000114%) | | | CGGTTTAAGTTCTTATACTCAATCATACACATGACATCAAGTCATATTCGACTCCAAAACACTAACCAACCTTCTTCTTGCTTCTCAAAGCTTTCATGGT | 70 (0.001324%) | | | CGGTTTAAGTTGTTATACTCAATCATACACATGACAACAAGTCATATTCGACTCCAAAACACTAACCAACCTTCTTCTTGCTTCTCAAAGCTTTCATGGT | 105 (0.001986%) | | | CGTAAGAATTGTATCCTTGTTAAAAGACACAAAGCCAAAGACTCATATGGACTTTGGCTACACCATGAAAGCTTTGAGAAGCAAGAAGAAGGTTGGTTAG | 539 (0.010197%) | | | CGTAAGAATTGTATCCTTGTTAGAAGACACAAAGCCAAAG | 236 (0.001786%) | | | CGTATATTTAAGTTGTTGCAGTTAAAAAGCTCGTAGTTGAACCTTGGGATGGGTCGGCCGGTCCGCCTTTGGTGTGCATTGGTCGGCTTGTCCCTTCGGT | 8 (0.000151%) | | | CGTATTCGTACTGAAAATCAGAATCAAACGAGCTTTTACCCTTTTGTTCCACACGAGATTTCTGTTCTCGTTGAGCTCATCTTAGGACACCTGCGTTATC | 23 (0.000435%) | | | CGTATTTCATAGTCAGAGGTGAAATTCTTGGATTTATGAAAGACGAACAACTGCGAAAGCATTTGCCAAGGATGTTTTCATTAATCAAGAACGAAAGTTG | 10 (0.000189%) | | | CGTCCCTCACCATCCTTTGCTGATGCGGGACGGAAGCTGGTCTCCCGTGTGTTACCGCACGCGGTTGGCCTAAATCCGAGCCAAGGACGCCTGGAGCGTA | 19 (0.000359%) | | | CGTCGAGTTATCATGAATCATCAGAGCAACGGGCAGAGCCCGCGTCGACCTTTTATCTAATAAATGCGTCCCTTCCATAAGTCGGGGTTTGTTGCACGTA | 103 (0.001949%) | | | CGTCGTCCCTCACCATCCTTTGCTGATGCGGGACGGAAGCTGGTCTCCCGTGTGTTACCGCACGCGGTTGGCCTAAATCCGAGCCAAGGACGCCTGGAGC | 4 (0.000076%) | | | CGTCGTGTAGGGAAAGAGTGTTCAGAGCCGTGTAGATCTC | 7 (0.000053%) | | | CGTGCCTCGGCATCAGCGTGCTCCGGGCGTCGGCCTGTGGGCTCCCCATTCGACCCGTCTTGAAACACGGACCAAGGAGTCTGACATGTGTGCGAGTCAA | 21 (0.000397%) | | | CGTGCTCCGGGCGTCGGCCTGTGGGCTCCCCATTCGACCCGTCTTGAAACACGGACCAAGGAGTCTGACATGTGTGCGAGTCAACGGGTGAGTAAACCCG | 4 (0.000076%) | | | CGTGGCGGTTGACGGCAACGTTAGGGAGTCCGGAGACGTCGGCGGGGGCCTCGGGAAGAGTTATCTTTTCTGTTTAACAGCCTGCCCACCCTGGAAACGG | 7 (0.000132%) | | | CGTGGGTGGTTCGCCGCCCGCGACGTCGCGAGAAGTCCACTAAACCTTATCATTTAGAGGAAGGAGAAGTCGTAACAAGGTTTCCGTAGGTGAACCTGCG | 33 (0.000624%) | | | CGTGTAGGGAAAGAGTGTTCAGAGCCGTGTAGATCTCGGT | 11 (0.000083%) | | | CGTTAAGGAGCTGTTGCTTTGTTAGTGTAGAAACACTTGTGTAGAATTGGGGATTGTTTTTTTTGGAGTGATTTAGGGGAGGGTCGAATCTTAGCGACAA | 197 (0.003727%) | | | CGTTAGGGAGTCCGGAGACGTCGGCGGGGGCCTCGGGAAGAGTTATCTTTTCTGTTTAACAGCCTGCCCACCCTGGAAACGGCTCAGCCGGAGGTAGGGT | 7 (0.000132%) | | | CGTTCATCACCCTTGGCCGGCTTTCGAACAGCCGGACTCCCATCAAAAGATGGTTGCCAAGAACATCTTCGTTACGGTTTGCTAATTCTCGGAATAACAT | 22 (0.000416%) | | | CGTTGACTCGCACACATGTCAGACTCCTTGGTCCGTGTTTCAAGACGGGTCGAATGGGGAGCCCACAGGCCGACGCCCGGAGCACGCTGATGCCGAGGCA | 13 (0.000246%) | | | CTAAATACGGGCGAGAGACCGATAGCGAACAAGTACCGCGAGGTAAAGATGAAAAGGACTTTGAAAAGAGAGTCAAAGAGTGCTTGAAATTGTCGGGAGG | 705 (0.013337%) | | | CTAACGGCGTGCCTCGGCATCAGCGTGCTCCGGGCGTCGGCCTGTGGGCTCCCCATTCGACCCGTCTTGAAACACGGACCAAGGAGTCTGACATGTGTGC | 100 (0.001892%) | | | CTAACTAGCTACGTGGAGGCATCCCTTCACGGCCGGCTTCTTAGAGGGACTATGGCCGTTTAGGCCAAGGAAGTTTGAGGCAATAACAGGTCTGTGATGC | 51 (0.000965%) | | | CTAAGTAGTGTTTCCTTGTTAGAAGACACAAAGCCAAAGACTCATATGGACTTTGGCTACACCATGAAAGCTTTGAGAAGCAAGAAGAAGGTTGGTTAGT | 56 (0.001059%) | | | CTAATGTATCCAGAGCGTAGGCTTGCTTTGAGCACTCTAATTTCTTCAAAGTAACAGCGCCGGAGGCACGACCCGGCCAATTAAGACCAGGAGCGTATCG | 45 (0.000851%) | | | CTACATTGTTCCATCGACCAGAGGCTGTTCACCTTGGAGACCTGATGCGGTTATGAGTACGACCGGGCGTGAGCGGCACTCGGTCCTCCGGATTTTCAAG | 35 (0.000662%) | | | CTACCTACCACCTATCCTTCTCAGTTAATTCACGGGCGATGTTACGCTCGATGATGAGTTCCGGGGCCTGTGTTTCGTACCTAATTTGAAGGAATTGTTG | 11 (0.000208%) | | | CTAGCTACGTGGAGGCATCCCTTCACGGCCGGCTTCTTAGAGGGACTATGGCCGTTTAGGCCAAGGAAGTTTGAGGCAATAACAGGTCTGTGATGCCCTT | 25 (0.000473%) | | | CTAGGCGAGACAAGGGTTCACATTTCGTTCATCACCCTTGGCCGGCTATCGAACAGCCGGACTCCCATCAAAAGATGGTTGCCAAGAACATCTTCGTTAC | 32 (0.000605%) | | | CTAGGCGAGACAAGGGTTCACATTTCGTTCATCACCCTTGGCCGGCTTTCGAACAGCCGGACTCCCATCAAAAGATGGTTGCCAAGAACATCTTCGTTAC | 61 (0.001154%) | | | CTAGGCTGTCCCGAGTGTGAGCGAGGTGTGAGTGTCGCCCATGGGCATCGACACCTTGCGGCTAGGAACTGGAACGAGACGGGTGGCAAAGATTTCGAGT | 411 (0.007775%) | | | CTAGTGTTCGTAGGCTCCATGCTCGCGCATCGAACTACCTACCACCTATCCTTCTCAGTTAATTCACGGGCGATGTTACGCTCGATGATGAGTTCCGGGG | 12 (0.000227%) | | | CTATGATGTTATCCCATGCTAATGTATCCAGAGCGTAGGCTTGCTTTGAGCACTCTAATTTCTTCAAAGTAACAGCGCCGGAGGCACGACCCGGCCAATT | 211 (0.003992%) | | | CTCAAAATCATCAATCGTTCCAACTAATCTACCGAAGTACTCGGCTAAGAAGAAAGAAGACGGACGAATCCGAGCCAAAGCCGTACAAGCGCGAGATACC | 1987 (0.037591%) | | | CTCAAAGATTAAGCCATGCATGTGTAAGTATGAACGAATTCAGACTGTGAAACTGCGAATGGCTCATTAAATCAGTTATAGTTTGTTTGATGGTAACTAC | 13 (0.000246%) | | | CTCAATCATACACATGACATCAAGTCATATTCGACTCCAAAACACTAACCAACCTTCTTCTTGCTTCTCAAAGCTTTCATGGTGTAGCCAAAGTCCATAT | 73 (0.001381%) | | | CTCACCATCCTTTGCTGATGCGGGACGGAAGCTGGTCTCCCGTGTGTTACCGCACGCGGTTGGCCTAAATCCGAGCCAAGGACGCCTGGAGCGTACCGAC | 16 (0.000303%) | | | CTCACCCGTTGACTCGCACACATGTCAGACTCCTTGGTCCGTGTTTCAAGACGGGTCGAATGGGGAGCCCACAGGCCGACGCCCGGAGCACGCTGATGCC | 13 (0.000246%) | | | CTCAGCCTGCTAACTAGCTACGTGGAGGCATCCCTTCACGGCCGGCTTCTTAGAGGGACTATGGCCGTTTAGGCCAAGGAAGTTTGAGGCAATAACAGGT | 37 (0.000700%) | | | CTCATGTGTATGATTGAGTATAAGAACTTAAACCGCAACCGCATCTTATAAGCCTAAGTAGTGTTTCCTTGTTAGAAGACACAAAGCCAAAGACTCATAT | 105 (0.001986%) | | | CTCCAAAACACTAACCAACCTTCTTCTTGCTTCTCAAAGCTTTCATGGTGTAGCCAAAGTCCATATGAGTCTTTGGCTTTGTGTCTTCTAACAAGGATAC | 19 (0.000359%) | | | CTCCAATAGCGTATATTTAAGTTGTTGCAGTTAAAAAGCTCGTAGTTGAACCTTGGGATGGGTCGGCCGGTCCGCCTTTGGTGTGCATTGGTCGGCTTGT | 32 (0.000605%) | | | CTCCATGCTCGCGCATCGAACTACCTACCACCTATCCTTCTCAGTTAATTCACGGGCGATGTTACGCTCGATGATGAGTTCCGGGGCCTGTGTTTCGTAC | 11 (0.000208%) | | | CTCCGGATTTTCAAGGGCCGCCGGGGGCGCACCGGACACCACGCGACGTGCGGTGCTCTTCCAGCCGCTGGACCCTACCTCCGGCTGAGCCGTTTCCAGG | 3 (0.000057%) | | | CTCCGGGCGTCGGCCTGTGGGCTCCCCATTCGACCCGTCTTGAAACACGGACCAAGGAGTCTGACATGTGTGCGAGTCAACGGGTGAGTAAACCCGTAAG | 10 (0.000189%) | | | CTCGCACACATGTCAGACTCCTTGGTCCGTGTTTCAAGACGGGTCGAATGGGGAGCCCACAGGCCGACGCCCGGAGCACGCTGATGCCGAGGCACGCCGT | 22 (0.000416%) | | | CTCGCGCTTGTACGGCTTTGGCTCGGATTCGTCCGTCTTCTTTCTTCTTAGCCGAGTACTTCGGTAGATTAGTTGGAACGATTGATGATTTTGAGTTAAT | 11 (0.000208%) | | | CTCGGCATCAGCGTGCTCCGGGCGTCGGCCTGTGGGCTCCCCATTCGACCCGTCTTGAAACACGGACCAAGGAGTCTGACATGTGTGCGAGTCAACGGGT | 29 (0.000549%) | | | CTCGGTCCTCCGGATTTTCAAGGGCCGCCGGGGGCGCACCGGACACCACGCGACGTGCGGTGCTCTTCCAGCCGCTGGACCCTACCTCCGGCTGAGCCGT | 22 (0.000416%) | | | CTCGTGTAAAGTTGGGAATTCGTTAAGGAGCTGTTGCTTTGTTAGTGTAGAAACACTTGTGTAGAATTGGGGATTGTTTTTTTTGGAGTGATTTAGGGGA | 39 (0.000738%) | | | CTCTGTGCTGGCGACGCATCATTCAAATTTCTGCCCTATCAACTTTCGATGGTAGGATAGTGGCCTACCATGGTGGTAACGGGTGACGGAGAATTAGGGT | 57 (0.001078%) | | | CTGAGAAGGGTTCGAGTGTGAGCATGCCTGTCGGGACCCGAAAGATGGTGAACTATGCCTGAGCGGGGTAAAGCCAGAGGAAACTCTGGTGGAAGCCCGC | 27 (0.000511%) | | | CTGATGCGGGACGGAAGCTGGTCTCCCGTGTGTTACCGCACGCGGTTGGCCTAAATCCGAGCCAAGGACGCCTGGAGCGTACCGACATGCGGTGGTGAAC | 29 (0.000549%) | | | CTGCAGCACGCGCCTAACGGCGTGCCTCGGCATCAGCGTGCTCCGGGCGTCGGCCTGTGGGCTCCCCATTCGACCCGTCTTGAAACACGGACCAAGGAGT | 2138 (0.040447%) | | | CTGCCCTATCAACTTTCGATGGTAGGATAGTGGCCTACCATGGTGGTAACGGGTGACGGAGAATTAGGGTTCGATTCCGGAGAGGGAGCCTGAGAAACGG | 14 (0.000265%) | | | CTGCTAACTAGCTACGTGGAGGCATCCCTTCACGGCCGGCTTCTTAGAGGGACTATGGCCGTTTAGGCCAAGGAAGTTTGAGGCAATAACAGGTCTGTGA | 56 (0.001059%) | | | CTGGCGACGCATCATTCAAATTTCTGCCCTATCAACTTTCGATGGTAGGATAGTGGCCTACCATGGTGGTAACGGGTGACGGAGAATTAGGGTTCGATTC | 24 (0.000454%) | | | CTGGGGCGGCACATCTGTTAAAAGATAACGCAGGTGTCCTAAGATGAGCTCAACGAGAACAGAAATCTCGTGTGGAACAAAAGGGTAAAAGCTCGTTTGA | 27 (0.000511%) | | | CTGGGTGTCACAAATCGTCGTCCCTCACCATCCTTTGCTGATGCGGGACGGAAGCTGGTCTCCCGTGTGTTACCGCACGCGGTTGGCCTAAATCCGAGCC | 826 (0.015626%) | | | CTGGTGCCAGCAGCCGCGGTAATTCCAGCTCCAATAGCGTATATTTAAGTTGTTGCAGTTAAAAAGCTCGTAGTTGAACCTTGGGATGGGTCGGCCGGTC | 31 (0.000586%) | | | CTGTCCCGAAGGTATCTCGCGCTTGTACGGCTTTGGCTCGGATTCGTCCGTCTTCTTTCTTCTTAGCCGAGTACTTCGGTAGATTAGTTGGAACGATTGA | 61 (0.001154%) | | | CTGTCCCGAGTGTGAGCGAGGTGTGAGTGTCGCCCATGGGCATCGACACCTTGCGGCTAGGAACTGGAACGAGACGGGTAGCAAAGATTTCGAGTAGCAC | 23 (0.000435%) | | | CTGTCCCGAGTGTGAGCGAGGTGTGAGTGTCGCCCATGGGCATCGACACCTTGCGGCTAGGAACTGGAACGAGACGGGTGGCAAAGATTTCGAGTAGCAC | 28 (0.000530%) | | | CTGTGCTGGCGACGCATCATTCAAATTTCTGCCCTATCAACTTTCGATGGTAGGATAGTGGCCTACCATGGTGGTAACGGGTGACGGAGAATTAGGGTTC | 33 (0.000624%) | | | CTGTTAAAAGATAACGCAGGTGTCCTAAGATGAGCTCAACGAGAACAGAAATCTCGTGTGGAACAAAAGGGTAAAAGCTCGTTTGATTCTGATTTTCAGT | 5 (0.000095%) | | | CTGTTGCTTTGTTAGTGTAGAAACACTTGTGTAGAATTGGGGATTGTTTTTTTTGGAGTGATTTAGGGGAGGGTCGAATCTTAGCGACAAAGGGCTGAAT | 19 (0.000359%) | | | CTTAAAGGCGTAAGAATTGTATCCTTGTTAGAAGACACAA | 1035 (0.007832%) | | | CTTACGGGTTTACTCACCCGTTGACTCGCACACATGTCAGACTCCTTGGTCCGTGTTTCAAGACGGGTCGAATGGGGAGCCCACAGGCCGACGCCCGGAG | 13 (0.000246%) | | | CTTATACTCAATCATACACATGACATCAAGTCATATTCGACTCCAAAACACTAACCAACCTTCTTCTTGCTTCTCAAAGCTTTCATGGTGTAGCCAAAGT | 148 (0.002800%) | | | CTTCAAATTAGGTACGAAACACAGGCCCCGGAACTCATCATCGAGCGTAACATCGCCCGTGAATTAACTGAGAAGGATAGGTGGTAGGTAGTTCGATGCG | 771 (0.014586%) | | | CTTCAACGAGGAATTCCTAGTAAGCGCGAGTCATCAGCTCGCGTTGACTACGTCCCTGCCCTTTGTACACACCGCCCGTCGCTCCTACCGATTGAATGAT | 59 (0.001116%) | | | CTTCCCTTGCCTACATTGTTCCATCGACCAGAGGCTGTTCACCTTGGAGACCTGATGCGGTTATGAGTACGACCGGGCGTGAGCGGCACTCGGTCCTCCG | 53 (0.001003%) | | | CTTCCTTGCGCCTTACGGGTTTACTCACCCGTTGACTCGCACACATGTCAGACTCCTTGGTCCGTGTTTCAAGACGGGTCGAATGGGGAGCCCACAGGCC | 100 (0.001892%) | | | CTTCTGAGAAGGGTTCGAGTGTGAGCATGCCTGTCGGGACCCGAAAGATGGTGAACTATGCCTGAGCGGGGTAAAGCCAGAGGAAACTCTGGTGGAAGCC | 14 (0.000265%) | | | CTTGCCTACATTGTTCCATCGACCAGAGGCTGTTCACCTTGGAGACCTGATGCGGTTATGAGTACGACCGGGCGTGAGCGGCACTCGGTCCTCCGGATTT | 33 (0.000624%) | | | CTTGCGCCTTACGGGTTTACTCACCCGTTGACTCGCACACATGTCAGACTCCTTGGTCCGTGTTTCAAGACGGGTCGAATGGGGAGCCCACAGGCCGACG | 16 (0.000303%) | | | CTTGGAGACCTGATGCGGTTATGAGTACGACCGGGCGTGAGCGGCACTCGGTCCTCCGGATTTTCAAGGGCCGCCGGGGGCGCACCGGACACCACGCGAC | 131 (0.002478%) | | | CTTGGTGATATGAACACAAACGTTCAATATGACAAACCCATGCCAAGTAAAGAGAAAATGAAAACTGGTGATTGTTGCGGAAATCGTCCAGGATTCCTCG | 16 (0.000303%) | | | CTTGTACGGCTTTGGCTCGGATTCGTCCGTCTTCTTTCTTCTTAGCCGAGTACTTCGGTAGATTAGTTGGAACGATTGATGATTTTGAGTTAATTGAACG | 14 (0.000265%) | | | CTTGTCTCAAAGATTAAGCCATGCATGTGTAAGTATGAACGAATTCAGACTGTGAAACTGCGAATGGCTCATTAAATCAGTTATAGTTTGTTTGATGGTA | 69 (0.001305%) | | | CTTGTTAGAAGACACAAAGCCAAAGACTCATATGGACTTTGGCTACACCATGAAAGCTTTGAGAAGCAAGAAGAAGGTTGGTTAGTGTTTTGGAGTCGAA | 109 (0.002062%) | | | CTTTCACGGTTCGTATTCGTACTGAAAATCAGAATCAAACGAGCTTTTACCCTTTTGTTCCACACGAGATTTCTGTTCTCGTTGAGCTCATCTTAGGACA | 35 (0.000662%) | | | CTTTCGATGGTAGGATAGTGGCCTACCATGGTGGTAACGGGTGACGGAGAATTAGGGTTCGATTCCGGAGAGGGAGCCTGAGAAACGGCTACCACATCCA | 8 (0.000151%) | | | CTTTGCTGATGCGGGACGGAAGCTGGTCTCCCGTGTGTTACCGCACGCGGTTGGCCTAAATCCGAGCCAAGGACGCCTGGAGCGTACCGACATGCGGTGG | 60 (0.001135%) | | | CTTTGGCTCGGATTCGTCCGTCTTCTTTCTTCTTAGCCGAGTACTTCGGTAGATTAGTTGGAACGATTGATGATTTTGAGTTAATTGAACGTTCGGCGTA | 6 (0.000114%) | | | CTTTGTTTTAATTAAACAGTCGGATTCCCCTTGTCCGTACCAGTTCTGAGCTGACTGTTCGACGCCCGGGGAAAGCTCCCGAGAGAGCCGTTCCCAGTCC | 77 (0.001457%) | | | GAAACACAGGCCCCGGAACTCATCATCGAGCGTAACATCGCCCGTGAATTAACTGAGAAGGATAGGTGGTAGGTAGTTCGATGCGCGAGCATGGAGCCTA | 72 (0.001362%) | | | GAAATCCTATGATGTTATCCCATGCTAATGTATCCAGAGCGTAGGCTTGCTTTGAGCACTCTAATTTCTTCAAAGTAACAGCGCCGGAGGCACGACCCGG | 175 (0.003311%) | | | GAACCCACAAAGGGTGTTGGTCGATTAAGACAGCAGGACGGTGGTCATGGAAGTCGAAATCCGCTAAGGAGTGTGTAACAACTCACCTGCCGAATCAACT | 884 (0.016724%) | | | GAACCGGGACGTGGCGGTTGACGGCAACGTTAGGGAGTCCGGAGACGTCGGCGGGGGCCTCGGGAAGAGTTATCTTTTCTGTTTAACAGCCTGCCCACCC | 7 (0.000132%) | | | GAAGACACAAAGCCAAAGACTCATATGGACTTTGGCTACACCATGAAAGCTTTGAGAAGCAAGAAGAAGGTTGGTTAGTGTTTTGGAGTCGAATATGACT | 35 (0.000662%) | | | GAAGAGCGTCGTGTAGGGAAAGAGTGTTCAGAGCCGTGTA | 21 (0.000159%) | | | GAAGCACGCCCATCCAACCTAGGCGAGACAAGGGTTCACATTTCGTTCATCACCCTTGGCCGGCTATCGAACAGCCGGACTCCCATCAAAAGATGGTTGC | 13 (0.000246%) | | | GAAGGTATCTCGCGCTTGTACGGCTTTGGCTCGGATTCGTCCGTCTTCTTTCTTCTTAGCCGAGTACTTCGGTAGATTAGTTGGAACGATTGATGATTTT | 10 (0.000189%) | | | GAAGTCGAAATCCGCTAAGGAGTGTGTAACAACTCACCTGCCGAATCAACTAGCCCCGAAAATGGATGGCGCTTAAGCGCGCGACCTATACCCGGCCGTC | 4 (0.000076%) | | | GAATCATCAGAGCAACGGGCAGAGCCCGCGTCGACCTTTTATCTAATAAATGCGTCCCTTCCATAAGTCGGGGTTTGTTGCACGTATTAGCTCTAGAATT | 104 (0.001967%) | | | GAATTGTATCCTTGTTAGAAGACACAAAGCCAAAGACTCATATGGACTTTGGCTACACCATGAAAGCTTTGAGAAGCAAGAAGAAGGTTGGTTAGTGTTT | 18 (0.000341%) | | | GACAAGGGTTCACATTTCGTTCATCACCCTTGGCCGGCTATCGAACAGCCGGACTCCCATCAAAAGATGGTTGCCAAGAACATCTTCGTTACGGTTTGCT | 14 (0.000265%) | | | GACAAGGGTTCACATTTCGTTCATCACCCTTGGCCGGCTTTCGAACAGCCGGACTCCCATCAAAAGATGGTTGCCAAGAACATCTTCGTTACGGTTTGCT | 9 (0.000170%) | | | GACACAAAGCCAAAGACTCATATGGACTTTGGCTACACCATGAAAGCTTTGAGAAGCAAGAAGAAGGTTGGTTAGTGTTTTGGAGTCGAATATGACTTGA | 49 (0.000927%) | | | GACAGCAGGACGGTGGTCATGGAAGTCGAAATCCGCTAAGGAGTGTGTAACAACTCACCTGCCGAATCAACTAGCCCCGAAAATGGATGGCGCTTAAGCG | 7 (0.000132%) | | | GACAGTCGGGGGCATTCGTATTTCATAGTCAGAGGTGAAATTCTTGGATTTATGAAAGACGAACAACTGCGAAAGCATTTGCCAAGGATGTTTTCATTAA | 7 (0.000132%) | | | GACATCAAGTCATATTCGACTCCAAAACACTAACCAACCTTCTTCTTGCTTCTCAAAGCTTTCATGGTGTAGCCAAAGTCCATATGAGTCTTTGGCTTTG | 45 (0.000851%) | | | GACATTGTCAGGTGGGGAGTTTGGCTGGGGCGGCACATCTGTTAAAAGATAACGCAGGTGTCCTAAGATGAGCTCAACGAGAACAGAAATCTCGTGTGGA | 36 (0.000681%) | | | GACCAGGGGTTGAAATCGTCGACCAGGTCCGAGACTTCATCGACCGGGTCCGAGGATTCGTCGACCAGGACGGCCGGATGTCCGAGAAAAAAAAATGTTG | 6 (0.000114%) | | | GACCCGCCGAAGCGAGCCTTGGGACCAAAAACAGGGGTTGTACCCCGCCTCCGATTCACGGAGTAAGTAAAATAACGTTAAAAGTAGTGGTATTTCACTT | 4 (0.000076%) | | | GACCTCAGCCTGCTAACTAGCTACGTGGAGGCATCCCTTCACGGCCGGCTTCTTAGAGGGACTATGGCCGTTTAGGCCAAGGAAGTTTGAGGCAATAACA | 6 (0.000114%) | | | GACGCATCATTCAAATTTCTGCCCTATCAACTTTCGATGGTAGGATAGTGGCCTACCATGGTGGTAACGGGTGACGGAGAATTAGGGTTCGATTCCGGAG | 21 (0.000397%) | | | GACGGGCGGTGTGTACAAAGGGCAGGGACGTAGTCAACGCGAGCTGATGACTCGCGCTTACTAGGAATTCCTCGTTGAAGACCAACAATTGCAATGATCG | 17 (0.000322%) | | | GACGGTGGTCATGGAAGTCGAAATCCGCTAAGGAGTGTGTAACAACTCACCTGCCGAATCAACTAGCCCCGAAAATGGATGGCGCTTAAGCGCGCGACCT | 9 (0.000170%) | | | GACGTGGGTGGTTCGCCGCCCGCGACGTCGCGAGAAGTCCACTAAACCTTATCATTTAGAGGAAGGAGAAGTCGTAACAAGGTTTCCGTAGGTGAACCTG | 47 (0.000889%) | | | GACTCCAAAACACTAACCAACCTTCTTCTTGCTTCTCAAAGCTTTCATGGTGTAGCCAAAGTCCATATGAGTCTTTGGCTTTGTGTCTTCTAACAAGGAT | 30 (0.000568%) | | | GACTTCCCTTGCCTACATTGTTCCATCGACCAGAGGCTGTTCACCTTGGAGACCTGATGCGGTTATGAGTACGACCGGGCGTGAGCGGCACTCGGTCCTC | 22 (0.000416%) | | | GAGAAGGGTTCGAGTGTGAGCATGCCTGTCGGGACCCGAAAGATGGTGAACTATGCCTGAGCGGGGTAAAGCCAGAGGAAACTCTGGTGGAAGCCCGCAG | 8 (0.000151%) | | | GAGACAAGGGTTCACATTTCGTTCATCACCCTTGGCCGGCTATCGAACAGCCGGACTCCCATCAAAAGATGGTTGCCAAGAACATCTTCGTTACGGTTTG | 8 (0.000151%) | | | GAGACCTCAGCCTGCTAACTAGCTACGTGGAGGCATCCCTTCACGGCCGGCTTCTTAGAGGGACTATGGCCGTTTAGGCCAAGGAAGTTTGAGGCAATAA | 972 (0.018389%) | | | GAGACGTCGGCGGGGGCCTCGGGAAGAGTTATCTTTTCTGTTTAACAGCCTGCCCACCCTGGAAACGGCTCAGCCGGAGGTAGGGTCCAGCGGCTGGAAG | 17 (0.000322%) | | | GAGCAACGGGCAGAGCCCGCGTCGACCTTTTATCTAATAAATGCGTCCCTTCCATAAGTCGGGGTTTGTTGCACGTATTAGCTCTAGAATTACTACGGTT | 14 (0.000265%) | | | GAGCCCGCGTCGACCTTTTATCTAATAAATGCGTCCCTTCCATAAGTCGGGGTTTGTTGCACGTATTAGCTCTAGAATTACTACGGTTATCCGAGTAGTA | 6 (0.000114%) | | | GAGCGACGGGCGGTGTGTACAAAGGGCAGGGACGTAGTCAACGCGAGCTGATGACTCGCGCTTACTAGGAATTCCTCGTTGAAGACCAACAATTGCAATG | 9 (0.000170%) | | | GAGCGAGGTGTGAGTGTCGCCCATGGGCATCGACACCTTGCGGCTAGGAACTGGAACGAGACGGGTGGCAAAGATTTCGAGTAGCACTTCATACTACCGT | 10 (0.000189%) | | | GAGCGGCACTCGGTCCTCCGGATTTTCAAGGGCCGCCGGGGGCGCACCGGACACCACGCGACGTGCGGTGCTCTTCCAGCCGCTGGACCCTACCTCCGGC | 13 (0.000246%) | | | GAGCGTAGGCTTGCTTTGAGCACTCTAATTTCTTCAAAGTAACAGCGCCGGAGGCACGACCCGGCCAATTAAGACCAGGAGCGTATCGCCGACCGAAGGG | 4 (0.000076%) | | | GAGCGTCGTGTAGGGAAAGAGTGTTCAGAGCCGTGTAGAT | 17 (0.000129%) | | | GAGCTCGTGTAAAGTTGGGAATTCGTTAAGGAGCTGTTGCTTTGTTAGTGTAGAAACACTTGTGTAGAATTGGGGATTGTTTTTTTTGGAGTGATTTAGG | 4 (0.000076%) | | | GAGCTGTTGCTTTGTTAGTGTAGAAACACTTGTGTAGAATTGGGGATTGTTTTTTTTGGAGTGATTTAGGGGAGGGTCGAATCTTAGCGACAAAGGGCTG | 7 (0.000132%) | | | GAGGACATTGTCAGGTGGGGAGTTTGGCTGGGGCGGCACATCTGTTAAAAGATAACGCAGGTGTCCTAAGATGAGCTCAACGAGAACAGAAATCTCGTGT | 53 (0.001003%) | | | GAGGCTGTTCACCTTGGAGACCTGATGCGGTTATGAGTACGACCGGGCGTGAGCGGCACTCGGTCCTCCGGATTTTCAAGGGCCGCCGGGGGCGCACCGG | 10 (0.000189%) | | | GAGGGCAAGTCTGGTGCCAGCAGCCGCGGTAATTCCAGCTCCAATAGCGTATATTTAAGTTGTTGCAGTTAAAAAGCTCGTAGTTGAACCTTGGGATGGG | 28 (0.000530%) | | | GAGGTGTGAGTGTCGCCCATGGGCATCGACACCTTGCGGCTAGGAACTGGAACGAGACGGGTAGCAAAGATTTCGAGTAGCACTTCATACTACCGTGGGT | 68 (0.001286%) | | | GAGGTGTGAGTGTCGCCCATGGGCATCGACACCTTGCGGCTAGGAACTGGAACGAGACGGGTGGCAAAGATTTCGAGTAGCACTTCATACTACCGTGGGT | 7 (0.000132%) | | | GAGTATAAGAACTTAAACCGCAACCGCATCTTATAAGCCTAAGTAGTGTTTCCTTGTTAGAAGACACAAAGCCAAAGACTCATATGGACTTTGGCTACAC | 7 (0.000132%) | | | GAGTCGAATATGACTTGATGTCATGTGTATGATTGAGTAT | 1492 (0.011290%) | | | GAGTGTCGCCCATGGGCATCGACACCTTGCGGCTAGGAACTGGAACGAGACGGGTGGCAAAGATTTCGAGTAGCACTTCATACTACCGTGGGTTTTTTAA | 4 (0.000076%) | | | GAGTGTGAGCGAGGTGTGAGTGTCGCCCATGGGCATCGACACCTTGCGGCTAGGAACTGGAACGAGACGGGTGGCAAAGATTTCGAGTAGCACTTCATAC | 5 (0.000095%) | | | GAGTTATCATGAATCATCAGAGCAACGGGCAGAGCCCGCGTCGACCTTTTATCTAATAAATGCGTCCCTTCCATAAGTCGGGGTTTGTTGCACGTATTAG | 62 (0.001173%) | | | GATAGCTAGTGTTCGTAGGCTCCATGCTCGCGCATCGAACTACCTACCACCTATCCTTCTCAGTTAATTCACGGGCGATGTTACGCTCGATGATGAGTTC | 23 (0.000435%) | | | GATCCATTACATTTTATCGGTCGCTCTTGTCCGGAAGCTGTAGATGACCCAAAGTCCATATAGCGACCCCAGGTCAGGCGGGATTACCCGCTGAGTTTAA | 28 (0.000530%) | | | GATCCCGCCAATCAGCTTCCTTGCGCCTTACGGGTTTACTCACCCGTTGACTCGCACACATGTCAGACTCCTTGGTCCGTGTTTCAAGACGGGTCGAATG | 562 (0.010632%) | | | GATCCGTCGAGTTATCATGAATCATCAGAGCAACGGGCAGAGCCCGCGTCGACCTTTTATCTAATAAATGCGTCCCTTCCATAAGTCGGGGTTTGTTGCA | 117 (0.002213%) | | | GATCGAAATCCTATGATGTTATCCCATGCTAATGTATCCAGAGCGTAGGCTTGCTTTGAGCACTCTAATTTCTTCAAAGTAACAGCGCCGGAGGCACGAC | 1570 (0.029702%) | | | GATCGACCCGCCGAAGCGAGCCTTGGGACCAAAAACAGGGGTTGTACCCCGCCTCCGATTCACGGAGTAAGTAAAATAACGTTAAAAGTAGTGGTATTTC | 33 (0.000624%) | | | GATCGATCAA | 169469 (0.320605%) | | | GATCGATCCATTACATTTTATCGGTCGCTCTTGTCCGGAAGCTGTAGATGACCCAAAGTCCATATAGCGACCCCAGGTCAGGCGGGATTACCCGCTGAGT | 1200 (0.022702%) | | | GATCGATCCGTAACTTCGGGAAAAGGATTGGCTCTGAGGGCTGGGCTCGGGGGTCCCAGTTCCGAACCCGTCGGCTGTCAGCGGACTGCTCGAGCTGCTT | 1226 (0.023194%) | | | GATCGATCCGTCGAGTTATCATGAATCATCAGAGCAACGGGCAGAGCCCGCGTCGACCTTTTATCTAATAAATGCGTCCCTTCCATAAGTCGGGGTTTGT | 1171 (0.022153%) | | | GATCGATCGACCCGCCGAAGCGAGCCTTGGGACCAAAAACAGGGGTTGTACCCCGCCTCCGATTCACGGAGTAAGTAAAATAACGTTAAAAGTAGTGGTA | 948 (0.017934%) | | | GATCGATCTCATGTGTATGATTGAGTATAAGAACTTAAACCGCAACCGCATCTTATAAGCCTAAGTAGTGTTTCCTTGTTAGAAGACACAAAGCCAAAGA | 207 (0.003916%) | | | GATCGATCTT | 189995 (0.359437%) | | | GATCGATCTTAAAGGCGTAAGAATTGTATCCTTGTTAAAAGACACAAAGCCAAAGACTCATATGGACTTTGGCTACACCATGAAAGCTTTGAGAAGCAAG | 488 (0.009232%) | | | GATCGATCTTAAAGGCGTAAGAATTGTATCCTTGTTAGAA | 4489 (0.033970%) | | | GATCGATCTTCTGAGAAGGGTTCGAGTGTGAGCATGCCTGTCGGGACCCGAAAGATGGTGAACTATGCCTGAGCGGGGTAAAGCCAGAGGAAACTCTGGT | 1117 (0.021132%) | | | GATCGGAAGAGCGTCGTGTAGGGAAAGAGTGTTCAGAGCC | 7 (0.000053%) | | | GATCTCATGTGTATGATTGAGTATAAGAACTTAAACCGCAACCGCATCTTATAAGCCTAAGTAGTGTTTCCTTGTTAGAAGACACAAAGCCAAAGACTCA | 56 (0.001059%) | | | GATCTTAAAGGCGTAAGAATTGTATCCTTGTTAAAAGACACAAAGCCAAAGACTCATATGGACTTTGGCTACACCATGAAAGCTTTGAGAAGCAAGAAGA | 233 (0.004408%) | | | GATCTTCTGAGAAGGGTTCGAGTGTGAGCATGCCTGTCGGGACCCGAAAGATGGTGAACTATGCCTGAGCGGGGTAAAGCCAGAGGAAACTCTGGTGGAA | 137 (0.002592%) | | | GATGTTATCCCATGCTAATGTATCCAGAGCGTAGGCTTGCTTTGAGCACTCTAATTTCTTCAAAGTAACAGCGCCGGAGGCACGACCCGGCCAATTAAGA | 10 (0.000189%) | | | GATTAACAGGGACAGTCGGGGGCATTCGTATTTCATAGTCAGAGGTGAAATTCTTGGATTTATGAAAGACGAACAACTGCGAAAGCATTTGCCAAGGATG | 171 (0.003235%) | | | GATTAAGACAGCAGGACGGTGGTCATGGAAGTCGAAATCCGCTAAGGAGTGTGTAACAACTCACCTGCCGAATCAACTAGCCCCGAAAATGGATGGCGCT | 41 (0.000776%) | | | GATTAGTCTTTCGCCCCTATACCCAAGTCAGACGAACGATTTGCACGTCAGTATCGCTGCGGGCTTCCACCAGAGTTTCCTCTGGCTTTACCCCGCTCAG | 11 (0.000208%) | | | GATTGAGTATAAGAACTTAAACCGCAACCGCATCTTATAAGCCTAAGTAGTGTTTCCTTGTTAGAAGACACAAAGCCAAAGACTCATATGGACTTTGGCT | 38 (0.000719%) | | | GATTTGCACGTCAGTATCGCTGCGGGCTTCCACCAGAGTTTCCTCTGGCTTTACCCCGCTCAGGCATAGTTCACCATCTTTCGGGTCCCGACAGGCATGC | 26 (0.000492%) | | | GATTTTCAAGGGCCGCCGGGGGCGCACCGGACACCACGCGACGTGCGGTGCTCTTCCAGCCGCTGGACCCTACCTCCGGCTGAGCCGTTTCCAGGGTGGG | 14 (0.000265%) | | | GCAACGGGCAGAGCCCGCGTCGACCTTTTATCTAATAAATGCGTCCCTTCCATAAGTCGGGGTTTGTTGCACGTATTAGCTCTAGAATTACTACGGTTAT | 38 (0.000719%) | | | GCAACGTTAGGGAGTCCGGAGACGTCGGCGGGGGCCTCGGGAAGAGTTATCTTTTCTGTTTAACAGCCTGCCCACCCTGGAAACGGCTCAGCCGGAGGTA | 19 (0.000359%) | | | GCAAGTCTGGTGCCAGCAGCCGCGGTAATTCCAGCTCCAATAGCGTATATTTAAGTTGTTGCAGTTAAAAAGCTCGTAGTTGAACCTTGGGATGGGTCGG | 12 (0.000227%) | | | GCACACATGTCAGACTCCTTGGTCCGTGTTTCAAGACGGGTCGAATGGGGAGCCCACAGGCCGACGCCCGGAGCACGCTGATGCCGAGGCACGCCGTTAG | 4 (0.000076%) | | | GCACCGGACACCACGCGACGTGCGGTGCTCTTCCAGCCGCTGGACCCTACCTCCGGCTGAGCCGTTTCCAGGGTGGGCAGGCTGTTAAACAGAAAAGATA | 31 (0.000586%) | | | GCACGCCCATCCAACCTAGGCGAGACAAGGGTTCACATTTCGTTCATCACCCTTGGCCGGCTATCGAACAGCCGGACTCCCATCAAAAGATGGTTGCCAA | 41 (0.000776%) | | | GCACGCGCCTAACGGCGTGCCTCGGCATCAGCGTGCTCCGGGCGTCGGCCTGTGGGCTCCCCATTCGACCCGTCTTGAAACACGGACCAAGGAGTCTGAC | 25 (0.000473%) | | | GCACTCGGTCCTCCGGATTTTCAAGGGCCGCCGGGGGCGCACCGGACACCACGCGACGTGCGGTGCTCTTCCAGCCGCTGGACCCTACCTCCGGCTGAGC | 4 (0.000076%) | | | GCAGAGCCCGCGTCGACCTTTTATCTAATAAATGCGTCCCTTCCATAAGTCGGGGTTTGTTGCACGTATTAGCTCTAGAATTACTACGGTTATCCGAGTA | 12 (0.000227%) | | | GCAGCACGCGCCTAACGGCGTGCCTCGGCATCAGCGTGCTCCGGGCGTCGGCCTGTGGGCTCCCCATTCGACCCGTCTTGAAACACGGACCAAGGAGTCT | 427 (0.008078%) | | | GCAGCCGCGGTAATTCCAGCTCCAATAGCGTATATTTAAGTTGTTGCAGTTAAAAAGCTCGTAGTTGAACCTTGGGATGGGTCGGCCGGTCCGCCTTTGG | 13 (0.000246%) | | | GCAGGACGGTGGTCATGGAAGTCGAAATCCGCTAAGGAGTGTGTAACAACTCACCTGCCGAATCAACTAGCCCCGAAAATGGATGGCGCTTAAGCGCGCG | 42 (0.000795%) | | | GCATCAGCGTGCTCCGGGCGTCGGCCTGTGGGCTCCCCATTCGACCCGTCTTGAAACACGGACCAAGGAGTCTGACATGTGTGCGAGTCAACGGGTGAGT | 10 (0.000189%) | | | GCATCATTCAAATTTCTGCCCTATCAACTTTCGATGGTAGGATAGTGGCCTACCATGGTGGTAACGGGTGACGGAGAATTAGGGTTCGATTCCGGAGAGG | 42 (0.000795%) | | | GCATCGAACTACCTACCACCTATCCTTCTCAGTTAATTCACGGGCGATGTTACGCTCGATGATGAGTTCCGGGGCCTGTGTTTCGTACCTAATTTGAAGG | 5 (0.000095%) | | | GCATCGACACCTTGCGGCTAGGAACTGGAACGAGACGGGTGGCAAAGATTTCGAGTAGCACTTCATACTACCGTGGGTTTTTTAAACCTTCCGAGTTTTG | 6 (0.000114%) | | | GCATGGCCTCTGTGCTGGCGACGCATCATTCAAATTTCTGCCCTATCAACTTTCGATGGTAGGATAGTGGCCTACCATGGTGGTAACGGGTGACGGAGAA | 1694 (0.032047%) | | | GCCAAAGACTCATATGGACTTTGGCTACACCATGAAAGCTTTGAGAAGCAAGAAGAAGGTTGGTTAGTGTTTTGGAGTCGAATATGACTTGATGTCATGT | 48 (0.000908%) | | | GCCAATCAGCTTCCTTGCGCCTTACGGGTTTACTCACCCGTTGACTCGCACACATGTCAGACTCCTTGGTCCGTGTTTCAAGACGGGTCGAATGGGGAGC | 120 (0.002270%) | | | GCCAGCAGCCGCGGTAATTCCAGCTCCAATAGCGTATATTTAAGTTGTTGCAGTTAAAAAGCTCGTAGTTGAACCTTGGGATGGGTCGGCCGGTCCGCCT | 13 (0.000246%) | | | GCCCATCCAACCTAGGCGAGACAAGGGTTCACATTTCGTTCATCACCCTTGGCCGGCTATCGAACAGCCGGACTCCCATCAAAAGATGGTTGCCAAGAAC | 58 (0.001097%) | | | GCCCATCCAACCTAGGCGAGACAAGGGTTCACATTTCGTTCATCACCCTTGGCCGGCTTTCGAACAGCCGGACTCCCATCAAAAGATGGTTGCCAAGAAC | 48 (0.000908%) | | | GCCCATGGGCATCGACACCTTGCGGCTAGGAACTGGAACGAGACGGGTGGCAAAGATTTCGAGTAGCACTTCATACTACCGTGGGTTTTTTAAACCTTCC | 5 (0.000095%) | | | GCCCGCGTCGACCTTTTATCTAATAAATGCGTCCCTTCCATAAGTCGGGGTTTGTTGCACGTATTAGCTCTAGAATTACTACGGTTATCCGAGTAGTAGT | 4 (0.000076%) | | | GCCCTATCAACTTTCGATGGTAGGATAGTGGCCTACCATGGTGGTAACGGGTGACGGAGAATTAGGGTTCGATTCCGGAGAGGGAGCCTGAGAAACGGCT | 16 (0.000303%) | | | GCCGAAGCGAGCCTTGGGACCAAAAACAGGGGTTGTACCCCGCCTCCGATTCACGGAGTAAGTAAAATAACGTTAAAAGTAGTGGTATTTCACTTGCGCC | 39 (0.000738%) | | | GCCGACTTCCCTTGCCTACATTGTTCCATCGACCAGAGGCTGTTCACCTTGGAGACCTGATGCGGTTATGAGTACGACCGGGCGTGAGCGGCACTCGGTC | 257 (0.004862%) | | | GCCGCCCGCGACGTCGCGAGAAGTCCACTAAACCTTATCATTTAGAGGAAGGAGAAGTCGTAACAAGGTTTCCGTAGGTGAACCTGCGGAAGGATCGATC | 18 (0.000341%) | | | GCCGCCGGGGGCGCACCGGACACCACGCGACGTGCGGTGCTCTTCCAGCCGCTGGACCCTACCTCCGGCTGAGCCGTTTCCAGGGTGGGCAGGCTGTTAA | 10 (0.000189%) | | | GCCGCGGTAATTCCAGCTCCAATAGCGTATATTTAAGTTGTTGCAGTTAAAAAGCTCGTAGTTGAACCTTGGGATGGGTCGGCCGGTCCGCCTTTGGTGT | 8 (0.000151%) | | | GCCGGGGGCGCACCGGACACCACGCGACGTGCGGTGCTCTTCCAGCCGCTGGACCCTACCTCCGGCTGAGCCGTTTCCAGGGTGGGCAGGCTGTTAAACA | 25 (0.000473%) | | | GCCTAACGGCGTGCCTCGGCATCAGCGTGCTCCGGGCGTCGGCCTGTGGGCTCCCCATTCGACCCGTCTTGAAACACGGACCAAGGAGTCTGACATGTGT | 43 (0.000813%) | | | GCCTAAGTAGTGTTTCCTTGTTAGAACATACAAAGCCAAAGACTCATACGGACTTTGGCTACACCATGAAAGCTTTGAGAAGCTAGAAGAAGGTTGGTTA | 48 (0.000908%) | | | GCCTAAGTAGTGTTTCCTTGTTAGAAGACACAAAGCCAAAGACTCATATGGACTTTGGCTACACCATGAAAGCTTTGAGAAGCAAGAAGAAGGTTGGTTA | 40 (0.000757%) | | | GCCTACATTGTTCCATCGACCAGAGGCTGTTCACCTTGGAGACCTGATGCGGTTATGAGTACGACCGGGCGTGAGCGGCACTCGGTCCTCCGGATTTTCA | 8 (0.000151%) | | | GCCTAGGCTGTCCCGAGTGTGAGCGAGGTGTGAGTGTCGCCCATGGGCATCGACACCTTGCGGCTAGGAACTGGAACGAGACGGGTGGCAAAGATTTCGA | 359 (0.006792%) | | | GCCTCGGCATCAGCGTGCTCCGGGCGTCGGCCTGTGGGCTCCCCATTCGACCCGTCTTGAAACACGGACCAAGGAGTCTGACATGTGTGCGAGTCAACGG | 13 (0.000246%) | | | GCCTCTGTGCTGGCGACGCATCATTCAAATTTCTGCCCTATCAACTTTCGATGGTAGGATAGTGGCCTACCATGGTGGTAACGGGTGACGGAGAATTAGG | 12 (0.000227%) | | | GCCTGCTAACTAGCTACGTGGAGGCATCCCTTCACGGCCGGCTTCTTAGAGGGACTATGGCCGTTTAGGCCAAGGAAGTTTGAGGCAATAACAGGTCTGT | 4 (0.000076%) | | | GCCTTACGGGTTTACTCACCCGTTGACTCGCACACATGTCAGACTCCTTGGTCCGTGTTTCAAGACGGGTCGAATGGGGAGCCCACAGGCCGACGCCCGG | 22 (0.000416%) | | | GCGACGCATCATTCAAATTTCTGCCCTATCAACTTTCGATGGTAGGATAGTGGCCTACCATGGTGGTAACGGGTGACGGAGAATTAGGGTTCGATTCCGG | 37 (0.000700%) | | | GCGACGGGCGGTGTGTACAAAGGGCAGGGACGTAGTCAACGCGAGCTGATGACTCGCGCTTACTAGGAATTCCTCGTTGAAGACCAACAATTGCAATGAT | 41 (0.000776%) | | | GCGACGTGGGTGGTTCGCCGCCCGCGACGTCGCGAGAAGTCCACTAAACCTTATCATTTAGAGGAAGGAGAAGTCGTAACAAGGTTTCCGTAGGTGAACC | 388 (0.007340%) | | | GCGAGACAAGGGTTCACATTTCGTTCATCACCCTTGGCCGGCTATCGAACAGCCGGACTCCCATCAAAAGATGGTTGCCAAGAACATCTTCGTTACGGTT | 31 (0.000586%) | | | GCGAGACAAGGGTTCACATTTCGTTCATCACCCTTGGCCGGCTTTCGAACAGCCGGACTCCCATCAAAAGATGGTTGCCAAGAACATCTTCGTTACGGTT | 39 (0.000738%) | | | GCGAGGTGTGAGTGTCGCCCATGGGCATCGACACCTTGCGGCTAGGAACTGGAACGAGACGGGTGGCAAAGATTTCGAGTAGCACTTCATACTACCGTGG | 42 (0.000795%) | | | GCGCACCGGACACCACGCGACGTGCGGTGCTCTTCCAGCCGCTGGACCCTACCTCCGGCTGAGCCGTTTCCAGGGTGGGCAGGCTGTTAAACAGAAAAGA | 23 (0.000435%) | | | GCGCATCGAACTACCTACCACCTATCCTTCTCAGTTAATTCACGGGCGATGTTACGCTCGATGATGAGTTCCGGGGCCTGTGTTTCGTACCTAATTTGAA | 8 (0.000151%) | | | GCGCCTAACGGCGTGCCTCGGCATCAGCGTGCTCCGGGCGTCGGCCTGTGGGCTCCCCATTCGACCCGTCTTGAAACACGGACCAAGGAGTCTGACATGT | 19 (0.000359%) | | | GCGCCTTACGGGTTTACTCACCCGTTGACTCGCACACATGTCAGACTCCTTGGTCCGTGTTTCAAGACGGGTCGAATGGGGAGCCCACAGGCCGACGCCC | 14 (0.000265%) | | | GCGCTTGTACGGCTTTGGCTCGGATTCGTCCGTCTTCTTTCTTCTTAGCCGAGTACTTCGGTAGATTAGTTGGAACGATTGATGATTTTGAGTTAATTGA | 6 (0.000114%) | | | GCGGAGGACATTGTCAGGTGGGGAGTTTGGCTGGGGCGGCACATCTGTTAAAAGATAACGCAGGTGTCCTAAGATGAGCTCAACGAGAACAGAAATCTCG | 1497 (0.028321%) | | | GCGGCACATCTGTTAAAAGATAACGCAGGTGTCCTAAGATGAGCTCAACGAGAACAGAAATCTCGTGTGGAACAAAAGGGTAAAAGCTCGTTTGATTCTG | 18 (0.000341%) | | | GCGGCACTCGGTCCTCCGGATTTTCAAGGGCCGCCGGGGGCGCACCGGACACCACGCGACGTGCGGTGCTCTTCCAGCCGCTGGACCCTACCTCCGGCTG | 56 (0.001059%) | | | GCGGCGACGTGGGTGGTTCGCCGCCCGCGACGTCGCGAGAAGTCCACTAAACCTTATCATTTAGAGGAAGGAGAAGTCGTAACAAGGTTTCCGTAGGTGA | 1266 (0.023950%) | | | GCGGTAATTCCAGCTCCAATAGCGTATATTTAAGTTGTTGCAGTTAAAAAGCTCGTAGTTGAACCTTGGGATGGGTCGGCCGGTCCGCCTTTGGTGTGCA | 15 (0.000284%) | | | GCGGTTGACGGCAACGTTAGGGAGTCCGGAGACGTCGGCGGGGGCCTCGGGAAGAGTTATCTTTTCTGTTTAACAGCCTGCCCACCCTGGAAACGGCTCA | 7 (0.000132%) | | | GCGGTTTAAGTTCTTATACTCAATCATACACATGACATCAAGTCATATTCGACTCCAAAACACTAACCAACCTTCTTCTTGCTTCTCAAAGCTTTCATGG | 140 (0.002649%) | | | GCGGTTTAAGTTGTTATACTCAATCATACACATGACAACAAGTCATATTCGACTCCAAAACACTAACCAACCTTCTTCTTGCTTCTCAAAGCTTTCATGG | 52 (0.000984%) | | | GCGTAAGAATTGTATCCTTGTTAGAAGACACAAAGCCAAA | 190 (0.001438%) | | | GCGTCGTGTAGGGAAAGAGTGTTCAGAGCCGTGTAGATCT | 8 (0.000061%) | | | GCGTGCCTCGGCATCAGCGTGCTCCGGGCGTCGGCCTGTGGGCTCCCCATTCGACCCGTCTTGAAACACGGACCAAGGAGTCTGACATGTGTGCGAGTCA | 8 (0.000151%) | | | GCGTGCTCCGGGCGTCGGCCTGTGGGCTCCCCATTCGACCCGTCTTGAAACACGGACCAAGGAGTCTGACATGTGTGCGAGTCAACGGGTGAGTAAACCC | 3 (0.000057%) | | | GCTAACTAGCTACGTGGAGGCATCCCTTCACGGCCGGCTTCTTAGAGGGACTATGGCCGTTTAGGCCAAGGAAGTTTGAGGCAATAACAGGTCTGTGATG | 28 (0.000530%) | | | GCTAATGTATCCAGAGCGTAGGCTTGCTTTGAGCACTCTAATTTCTTCAAAGTAACAGCGCCGGAGGCACGACCCGGCCAATTAAGACCAGGAGCGTATC | 23 (0.000435%) | | | GCTAGTGTTCGTAGGCTCCATGCTCGCGCATCGAACTACCTACCACCTATCCTTCTCAGTTAATTCACGGGCGATGTTACGCTCGATGATGAGTTCCGGG | 23 (0.000435%) | | | GCTCCAATAGCGTATATTTAAGTTGTTGCAGTTAAAAAGCTCGTAGTTGAACCTTGGGATGGGTCGGCCGGTCCGCCTTTGGTGTGCATTGGTCGGCTTG | 12 (0.000227%) | | | GCTCCATGCTCGCGCATCGAACTACCTACCACCTATCCTTCTCAGTTAATTCACGGGCGATGTTACGCTCGATGATGAGTTCCGGGGCCTGTGTTTCGTA | 11 (0.000208%) | | | GCTCCGGGCGTCGGCCTGTGGGCTCCCCATTCGACCCGTCTTGAAACACGGACCAAGGAGTCTGACATGTGTGCGAGTCAACGGGTGAGTAAACCCGTAA | 9 (0.000170%) | | | GCTCGCGCATCGAACTACCTACCACCTATCCTTCTCAGTTAATTCACGGGCGATGTTACGCTCGATGATGAGTTCCGGGGCCTGTGTTTCGTACCTAATT | 6 (0.000114%) | | | GCTCGTGTAAAGTTGGGAATTCGTTAAGGAGCTGTTGCTTTGTTAGTGTAGAAACACTTGTGTAGAATTGGGGATTGTTTTTTTTGGAGTGATTTAGGGG | 5 (0.000095%) | | | GCTGATGCGGGACGGAAGCTGGTCTCCCGTGTGTTACCGCACGCGGTTGGCCTAAATCCGAGCCAAGGACGCCTGGAGCGTACCGACATGCGGTGGTGAA | 6 (0.000114%) | | | GCTGGCGACGCATCATTCAAATTTCTGCCCTATCAACTTTCGATGGTAGGATAGTGGCCTACCATGGTGGTAACGGGTGACGGAGAATTAGGGTTCGATT | 32 (0.000605%) | | | GCTGGGGCGGCACATCTGTTAAAAGATAACGCAGGTGTCCTAAGATGAGCTCAACGAGAACAGAAATCTCGTGTGGAACAAAAGGGTAAAAGCTCGTTTG | 8 (0.000151%) | | | GCTGTCCCGAAGGTATCTCGCGCTTGTACGGCTTTGGCTCGGATTCGTCCGTCTTCTTTCTTCTTAGCCGAGTACTTCGGTAGATTAGTTGGAACGATTG | 57 (0.001078%) | | | GCTGTCCCGAGTGTGAGCGAGGTGTGAGTGTCGCCCATGGGCATCGACACCTTGCGGCTAGGAACTGGAACGAGACGGGTAGCAAAGATTTCGAGTAGCA | 68 (0.001286%) | | | GCTGTCCCGAGTGTGAGCGAGGTGTGAGTGTCGCCCATGGGCATCGACACCTTGCGGCTAGGAACTGGAACGAGACGGGTGGCAAAGATTTCGAGTAGCA | 105 (0.001986%) | | | GCTGTTGCTTTGTTAGTGTAGAAACACTTGTGTAGAATTGGGGATTGTTTTTTTTGGAGTGATTTAGGGGAGGGTCGAATCTTAGCGACAAAGGGCTGAA | 3 (0.000057%) | | | GCTTCCTTGCGCCTTACGGGTTTACTCACCCGTTGACTCGCACACATGTCAGACTCCTTGGTCCGTGTTTCAAGACGGGTCGAATGGGGAGCCCACAGGC | 42 (0.000795%) | | | GCTTGTACGGCTTTGGCTCGGATTCGTCCGTCTTCTTTCTTCTTAGCCGAGTACTTCGGTAGATTAGTTGGAACGATTGATGATTTTGAGTTAATTGAAC | 21 (0.000397%) | | | GCTTGTCTCAAAGATTAAGCCATGCATGTGTAAGTATGAACGAATTCAGACTGTGAAACTGCGAATGGCTCATTAAATCAGTTATAGTTTGTTTGATGGT | 13 (0.000246%) | | | GCTTTCACGGTTCGTATTCGTACTGAAAATCAGAATCAAACGAGCTTTTACCCTTTTGTTCCACACGAGATTTCTGTTCTCGTTGAGCTCATCTTAGGAC | 16 (0.000303%) | | | GCTTTGGCTCGGATTCGTCCGTCTTCTTTCTTCTTAGCCGAGTACTTCGGTAGATTAGTTGGAACGATTGATGATTTTGAGTTAATTGAACGTTCGGCGT | 5 (0.000095%) | | | GCTTTGTTTTAATTAAACAGTCGGATTCCCCTTGTCCGTACCAGTTCTGAGCTGACTGTTCGACGCCCGGGGAAAGCTCCCGAGAGAGCCGTTCCCAGTC | 13 (0.000246%) | | | GGAAAGAGTGTTCAGAGCCGTGTAGATCTCGGTGGTCGCC | 15 (0.000114%) | | | GGAACCGGGACGTGGCGGTTGACGGCAACGTTAGGGAGTCCGGAGACGTCGGCGGGGGCCTCGGGAAGAGTTATCTTTTCTGTTTAACAGCCTGCCCACC | 51 (0.000965%) | | | GGAAGAGCGTCGTGTAGGGAAAGAGTGTTCAGAGCCGTGT | 48 (0.000363%) | | | GGAAGTCGAAATCCGCTAAGGAGTGTGTAACAACTCACCTGCCGAATCAACTAGCCCCGAAAATGGATGGCGCTTAAGCGCGCGACCTATACCCGGCCGT | 30 (0.000568%) | | | GGACAGTCGGGGGCATTCGTATTTCATAGTCAGAGGTGAAATTCTTGGATTTATGAAAGACGAACAACTGCGAAAGCATTTGCCAAGGATGTTTTCATTA | 64 (0.001211%) | | | GGACATTGTCAGGTGGGGAGTTTGGCTGGGGCGGCACATCTGTTAAAAGATAACGCAGGTGTCCTAAGATGAGCTCAACGAGAACAGAAATCTCGTGTGG | 179 (0.003386%) | | | GGACGGTGGTCATGGAAGTCGAAATCCGCTAAGGAGTGTGTAACAACTCACCTGCCGAATCAACTAGCCCCGAAAATGGATGGCGCTTAAGCGCGCGACC | 47 (0.000889%) | | | GGACGTGGCGGTTGACGGCAACGTTAGGGAGTCCGGAGACGTCGGCGGGGGCCTCGGGAAGAGTTATCTTTTCTGTTTAACAGCCTGCCCACCCTGGAAA | 3 (0.000057%) | | | GGAGACGTCGGCGGGGGCCTCGGGAAGAGTTATCTTTTCTGTTTAACAGCCTGCCCACCCTGGAAACGGCTCAGCCGGAGGTAGGGTCCAGCGGCTGGAA | 21 (0.000397%) | | | GGAGCGACGGGCGGTGTGTACAAAGGGCAGGGACGTAGTCAACGCGAGCTGATGACTCGCGCTTACTAGGAATTCCTCGTTGAAGACCAACAATTGCAAT | 33 (0.000624%) | | | GGAGCTGTTGCTTTGTTAGTGTAGAAACACTTGTGTAGAATTGGGGATTGTTTTTTTTGGAGTGATTTAGGGGAGGGTCGAATCTTAGCGACAAAGGGCT | 16 (0.000303%) | | | GGAGGACATTGTCAGGTGGGGAGTTTGGCTGGGGCGGCACATCTGTTAAAAGATAACGCAGGTGTCCTAAGATGAGCTCAACGAGAACAGAAATCTCGTG | 373 (0.007057%) | | | GGAGGGCAAGTCTGGTGCCAGCAGCCGCGGTAATTCCAGCTCCAATAGCGTATATTTAAGTTGTTGCAGTTAAAAAGCTCGTAGTTGAACCTTGGGATGG | 74 (0.001400%) | | | GGAGTAATGATTAACAGGGACAGTCGGGGGCATTCGTATTTCATAGTCAGAGGTGAAATTCTTGGATTTATGAAAGACGAACAACTGCGAAAGCATTTGC | 1017 (0.019240%) | | | GGAGTCCGGAGACGTCGGCGGGGGCCTCGGGAAGAGTTATCTTTTCTGTTTAACAGCCTGCCCACCCTGGAAACGGCTCAGCCGGAGGTAGGGTCCAGCG | 4 (0.000076%) | | | GGAGTTTGGCTGGGGCGGCACATCTGTTAAAAGATAACGCAGGTGTCCTAAGATGAGCTCAACGAGAACAGAAATCTCGTGTGGAACAAAAGGGTAAAAG | 6 (0.000114%) | | | GGAGTTTTTTCAGCAGTTCTCGGACAAAAATTGCTGAGTGGCCGAGAAGAATGGGCGTGTCATGCGTGGGCTGACATGGATTCTTCGAGGCCTAGGGGTG | 4 (0.000076%) | | | GGATAGCTAGTGTTCGTAGGCTCCATGCTCGCGCATCGAACTACCTACCACCTATCCTTCTCAGTTAATTCACGGGCGATGTTACGCTCGATGATGAGTT | 33 (0.000624%) | | | GGATTTTCAAGGGCCGCCGGGGGCGCACCGGACACCACGCGACGTGCGGTGCTCTTCCAGCCGCTGGACCCTACCTCCGGCTGAGCCGTTTCCAGGGTGG | 48 (0.000908%) | | | GGCAACGTTAGGGAGTCCGGAGACGTCGGCGGGGGCCTCGGGAAGAGTTATCTTTTCTGTTTAACAGCCTGCCCACCCTGGAAACGGCTCAGCCGGAGGT | 9 (0.000170%) | | | GGCAAGTCTGGTGCCAGCAGCCGCGGTAATTCCAGCTCCAATAGCGTATATTTAAGTTGTTGCAGTTAAAAAGCTCGTAGTTGAACCTTGGGATGGGTCG | 69 (0.001305%) | | | GGCAATTCCCCGCCACATCCTCTCAAACGCAATGGAAAGAGAGAAAGGACGAGGTCTTGACCGTCATCTTTTGCCCGAAGGACGGATGAGCTTTGGCGGG | 71 (0.001343%) | | | GGCACATCTGTTAAAAGATAACGCAGGTGTCCTAAGATGAGCTCAACGAGAACAGAAATCTCGTGTGGAACAAAAGGGTAAAAGCTCGTTTGATTCTGAT | 14 (0.000265%) | | | GGCACTCGGTCCTCCGGATTTTCAAGGGCCGCCGGGGGCGCACCGGACACCACGCGACGTGCGGTGCTCTTCCAGCCGCTGGACCCTACCTCCGGCTGAG | 14 (0.000265%) | | | GGCAGAGCCCGCGTCGACCTTTTATCTAATAAATGCGTCCCTTCCATAAGTCGGGGTTTGTTGCACGTATTAGCTCTAGAATTACTACGGTTATCCGAGT | 28 (0.000530%) | | | GGCATCAGCGTGCTCCGGGCGTCGGCCTGTGGGCTCCCCATTCGACCCGTCTTGAAACACGGACCAAGGAGTCTGACATGTGTGCGAGTCAACGGGTGAG | 5 (0.000095%) | | | GGCATCGACACCTTGCGGCTAGGAACTGGAACGAGACGGGTGGCAAAGATTTCGAGTAGCACTTCATACTACCGTGGGTTTTTTAAACCTTCCGAGTTTT | 15 (0.000284%) | | | GGCATTCGTATTTCATAGTCAGAGGTGAAATTCTTGGATTTATGAAAGACGAACAACTGCGAAAGCATTTGCCAAGGATGTTTTCATTAATCAAGAACGA | 9 (0.000170%) | | | GGCCGCCGGGGGCGCACCGGACACCACGCGACGTGCGGTGCTCTTCCAGCCGCTGGACCCTACCTCCGGCTGAGCCGTTTCCAGGGTGGGCAGGCTGTTA | 8 (0.000151%) | | | GGCCTCTGTGCTGGCGACGCATCATTCAAATTTCTGCCCTATCAACTTTCGATGGTAGGATAGTGGCCTACCATGGTGGTAACGGGTGACGGAGAATTAG | 157 (0.002970%) | | | GGCCTGTGGGCTCCCCATTCGACCCGTCTTGAAACACGGACCAAGGAGTCTGACATGTGTGCGAGTCAACGGGTGAGTAAACCCGTAAGGCGCAAGGAAG | 5 (0.000095%) | | | GGCGACGCATCATTCAAATTTCTGCCCTATCAACTTTCGATGGTAGGATAGTGGCCTACCATGGTGGTAACGGGTGACGGAGAATTAGGGTTCGATTCCG | 21 (0.000397%) | | | GGCGACGTGGGTGGTTCGCCGCCCGCGACGTCGCGAGAAGTCCACTAAACCTTATCATTTAGAGGAAGGAGAAGTCGTAACAAGGTTTCCGTAGGTGAAC | 378 (0.007151%) | | | GGCGAGACAAGGGTTCACATTTCGTTCATCACCCTTGGCCGGCTATCGAACAGCCGGACTCCCATCAAAAGATGGTTGCCAAGAACATCTTCGTTACGGT | 14 (0.000265%) | | | GGCGAGACAAGGGTTCACATTTCGTTCATCACCCTTGGCCGGCTTTCGAACAGCCGGACTCCCATCAAAAGATGGTTGCCAAGAACATCTTCGTTACGGT | 17 (0.000322%) | | | GGCGCACCGGACACCACGCGACGTGCGGTGCTCTTCCAGCCGCTGGACCCTACCTCCGGCTGAGCCGTTTCCAGGGTGGGCAGGCTGTTAAACAGAAAAG | 30 (0.000568%) | | | GGCGGCACATCTGTTAAAAGATAACGCAGGTGTCCTAAGATGAGCTCAACGAGAACAGAAATCTCGTGTGGAACAAAAGGGTAAAAGCTCGTTTGATTCT | 3 (0.000057%) | | | GGCGGTTGACGGCAACGTTAGGGAGTCCGGAGACGTCGGCGGGGGCCTCGGGAAGAGTTATCTTTTCTGTTTAACAGCCTGCCCACCCTGGAAACGGCTC | 18 (0.000341%) | | | GGCGTAAGAATTGTATCCTTGTTAGAAGACACAAAGCCAA | 62 (0.000469%) | | | GGCGTGCCTCGGCATCAGCGTGCTCCGGGCGTCGGCCTGTGGGCTCCCCATTCGACCCGTCTTGAAACACGGACCAAGGAGTCTGACATGTGTGCGAGTC | 19 (0.000359%) | | | GGCTGGGGCGGCACATCTGTTAAAAGATAACGCAGGTGTCCTAAGATGAGCTCAACGAGAACAGAAATCTCGTGTGGAACAAAAGGGTAAAAGCTCGTTT | 6 (0.000114%) | | | GGCTGTCCCGAGTGTGAGCGAGGTGTGAGTGTCGCCCATGGGCATCGACACCTTGCGGCTAGGAACTGGAACGAGACGGGTAGCAAAGATTTCGAGTAGC | 9 (0.000170%) | | | GGCTGTCCCGAGTGTGAGCGAGGTGTGAGTGTCGCCCATGGGCATCGACACCTTGCGGCTAGGAACTGGAACGAGACGGGTGGCAAAGATTTCGAGTAGC | 22 (0.000416%) | | | GGCTTTGGCTCGGATTCGTCCGTCTTCTTTCTTCTTAGCCGAGTACTTCGGTAGATTAGTTGGAACGATTGATGATTTTGAGTTAATTGAACGTTCGGCG | 8 (0.000151%) | | | GGGAAAGAGTGTTCAGAGCCGTGTAGATCTCGGTGGTCGC | 16 (0.000121%) | | | GGGACAGTCGGGGGCATTCGTATTTCATAGTCAGAGGTGAAATTCTTGGATTTATGAAAGACGAACAACTGCGAAAGCATTTGCCAAGGATGTTTTCATT | 26 (0.000492%) | | | GGGACGTGGCGGTTGACGGCAACGTTAGGGAGTCCGGAGACGTCGGCGGGGGCCTCGGGAAGAGTTATCTTTTCTGTTTAACAGCCTGCCCACCCTGGAA | 21 (0.000397%) | | | GGGAGTCCGGAGACGTCGGCGGGGGCCTCGGGAAGAGTTATCTTTTCTGTTTAACAGCCTGCCCACCCTGGAAACGGCTCAGCCGGAGGTAGGGTCCAGC | 3 (0.000057%) | | | GGGAGTTTGGCTGGGGCGGCACATCTGTTAAAAGATAACGCAGGTGTCCTAAGATGAGCTCAACGAGAACAGAAATCTCGTGTGGAACAAAAGGGTAAAA | 38 (0.000719%) | | | GGGCAAGTCTGGTGCCAGCAGCCGCGGTAATTCCAGCTCCAATAGCGTATATTTAAGTTGTTGCAGTTAAAAAGCTCGTAGTTGAACCTTGGGATGGGTC | 144 (0.002724%) | | | GGGCAGAGCCCGCGTCGACCTTTTATCTAATAAATGCGTCCCTTCCATAAGTCGGGGTTTGTTGCACGTATTAGCTCTAGAATTACTACGGTTATCCGAG | 55 (0.001041%) | | | GGGCATCGACACCTTGCGGCTAGGAACTGGAACGAGACGGGTGGCAAAGATTTCGAGTAGCACTTCATACTACCGTGGGTTTTTTAAACCTTCCGAGTTT | 13 (0.000246%) | | | GGGCATTCGTATTTCATAGTCAGAGGTGAAATTCTTGGATTTATGAAAGACGAACAACTGCGAAAGCATTTGCCAAGGATGTTTTCATTAATCAAGAACG | 32 (0.000605%) | | | GGGCCGCCGGGGGCGCACCGGACACCACGCGACGTGCGGTGCTCTTCCAGCCGCTGGACCCTACCTCCGGCTGAGCCGTTTCCAGGGTGGGCAGGCTGTT | 3 (0.000057%) | | | GGGCGCACCGGACACCACGCGACGTGCGGTGCTCTTCCAGCCGCTGGACCCTACCTCCGGCTGAGCCGTTTCCAGGGTGGGCAGGCTGTTAAACAGAAAA | 6 (0.000114%) | | | GGGCGGCACATCTGTTAAAAGATAACGCAGGTGTCCTAAGATGAGCTCAACGAGAACAGAAATCTCGTGTGGAACAAAAGGGTAAAAGCTCGTTTGATTC | 4 (0.000076%) | | | GGGCGGTGTGTACAAAGGGCAGGGACGTAGTCAACGCGAGCTGATGACTCGCGCTTACTAGGAATTCCTCGTTGAAGACCAACAATTGCAATGATCGATC | 20 (0.000378%) | | | GGGCGTCGGCCTGTGGGCTCCCCATTCGACCCGTCTTGAAACACGGACCAAGGAGTCTGACATGTGTGCGAGTCAACGGGTGAGTAAACCCGTAAGGCGC | 8 (0.000151%) | | | GGGGAGTTTGGCTGGGGCGGCACATCTGTTAAAAGATAACGCAGGTGTCCTAAGATGAGCTCAACGAGAACAGAAATCTCGTGTGGAACAAAAGGGTAAA | 32 (0.000605%) | | | GGGGCATTCGTATTTCATAGTCAGAGGTGAAATTCTTGGATTTATGAAAGACGAACAACTGCGAAAGCATTTGCCAAGGATGTTTTCATTAATCAAGAAC | 27 (0.000511%) | | | GGGGCGCACCGGACACCACGCGACGTGCGGTGCTCTTCCAGCCGCTGGACCCTACCTCCGGCTGAGCCGTTTCCAGGGTGGGCAGGCTGTTAAACAGAAA | 10 (0.000189%) | | | GGGGCGGCACATCTGTTAAAAGATAACGCAGGTGTCCTAAGATGAGCTCAACGAGAACAGAAATCTCGTGTGGAACAAAAGGGTAAAAGCTCGTTTGATT | 7 (0.000132%) | | | GGGGGCATTCGTATTTCATAGTCAGAGGTGAAATTCTTGGATTTATGAAAGACGAACAACTGCGAAAGCATTTGCCAAGGATGTTTTCATTAATCAAGAA | 17 (0.000322%) | | | GGGGGCGCACCGGACACCACGCGACGTGCGGTGCTCTTCCAGCCGCTGGACCCTACCTCCGGCTGAGCCGTTTCCAGGGTGGGCAGGCTGTTAAACAGAA | 23 (0.000435%) | | | GGGGTTGAAATCGTCGACCAGGTCCGAGACTTCATCGACCGGGTCCGAGGATTCGTCGACCAGGACGGCCGGATGTCCGAGAAAAAAAAATGTTGCCGAA | 5 (0.000095%) | | | GGGTGGTTCGCCGCCCGCGACGTCGCGAGAAGTCCACTAAACCTTATCATTTAGAGGAAGGAGAAGTCGTAACAAGGTTTCCGTAGGTGAACCTGCGGAA | 138 (0.002611%) | | | GGGTGTCACAAATCGTCGTCCCTCACCATCCTTTGCTGATGCGGGACGGAAGCTGGTCTCCCGTGTGTTACCGCACGCGGTTGGCCTAAATCCGAGCCAA | 7 (0.000132%) | | | GGGTGTTGGTCGATTAAGACAGCAGGACGGTGGTCATGGAAGTCGAAATCCGCTAAGGAGTGTGTAACAACTCACCTGCCGAATCAACTAGCCCCGAAAA | 24 (0.000454%) | | | GGGTTCACATTTCGTTCATCACCCTTGGCCGGCTATCGAACAGCCGGACTCCCATCAAAAGATGGTTGCCAAGAACATCTTCGTTACGGTTTGCTAATTC | 3 (0.000057%) | | | GGGTTCACATTTCGTTCATCACCCTTGGCCGGCTTTCGAACAGCCGGACTCCCATCAAAAGATGGTTGCCAAGAACATCTTCGTTACGGTTTGCTAATTC | 3 (0.000057%) | | | GGGTTGAAATCGTCGACCAGGTCCGAGACTTCATCGACCGGGTCCGAGGATTCGTCGACCAGGACGGCCGGATGTCCGAGAAAAAAAAATGTTGCCGAAT | 3 (0.000057%) | | | GGGTTGCGGTTTAAGTTGTTATACTCAATCATACACATGACAACAAGTCATATTCGACTCCAAAACACTAACCAACCTTCTTCTTGCTTCTCAAAGCTTT | 474 (0.008967%) | | | GGGTTTACTCACCCGTTGACTCGCACACATGTCAGACTCCTTGGTCCGTGTTTCAAGACGGGTCGAATGGGGAGCCCACAGGCCGACGCCCGGAGCACGC | 7 (0.000132%) | | | GGTAATTCCAGCTCCAATAGCGTATATTTAAGTTGTTGCAGTTAAAAAGCTCGTAGTTGAACCTTGGGATGGGTCGGCCGGTCCGCCTTTGGTGTGCATT | 108 (0.002043%) | | | GGTACGAAACACAGGCCCCGGAACTCATCATCGAGCGTAACATCGCCCGTGAATTAACTGAGAAGGATAGGTGGTAGGTAGTTCGATGCGCGAGCATGGA | 3 (0.000057%) | | | GGTAGGAGCGACGGGCGGTGTGTACAAAGGGCAGGGACGTAGTCAACGCGAGCTGATGACTCGCGCTTACTAGGAATTCCTCGTTGAAGACCAACAATTG | 41 (0.000776%) | | | GGTATCTCGCGCTTGTACGGCTTTGGCTCGGATTCGTCCGTCTTCTTTCTTCTTAGCCGAGTACTTCGGTAGATTAGTTGGAACGATTGATGATTTTGAG | 20 (0.000378%) | | | GGTCCTCCGGATTTTCAAGGGCCGCCGGGGGCGCACCGGACACCACGCGACGTGCGGTGCTCTTCCAGCCGCTGGACCCTACCTCCGGCTGAGCCGTTTC | 6 (0.000114%) | | | GGTCGACCAGGGGTTGAAATCGTCGACCAGGTCCGAGACTTCATCGACCGGGTCCGAGGATTCGTCGACCAGGACGGCCGGATGTCCGAGAAAAAAAAAT | 774 (0.014643%) | | | GGTCGATTAAGACAGCAGGACGGTGGTCATGGAAGTCGAAATCCGCTAAGGAGTGTGTAACAACTCACCTGCCGAATCAACTAGCCCCGAAAATGGATGG | 43 (0.000813%) | | | GGTCTTCAACGAGGAATTCCTAGTAAGCGCGAGTCATCAGCTCGCGTTGACTACGTCCCTGCCCTTTGTACACACCGCCCGTCGCTCCTACCGATTGAAT | 9 (0.000170%) | | | GGTGATATGAACACAAACGTTCAATATGACAAACCCATGCCAAGTAAAGAGAAAATGAAAACTGGTGATTGTTGCGGAAATCGTCCAGGATTCCTCGACC | 3 (0.000057%) | | | GGTGCCAGCAGCCGCGGTAATTCCAGCTCCAATAGCGTATATTTAAGTTGTTGCAGTTAAAAAGCTCGTAGTTGAACCTTGGGATGGGTCGGCCGGTCCG | 9 (0.000170%) | | | GGTGGGGAGTTTGGCTGGGGCGGCACATCTGTTAAAAGATAACGCAGGTGTCCTAAGATGAGCTCAACGAGAACAGAAATCTCGTGTGGAACAAAAGGGT | 3 (0.000057%) | | | GGTGGTCATGGAAGTCGAAATCCGCTAAGGAGTGTGTAACAACTCACCTGCCGAATCAACTAGCCCCGAAAATGGATGGCGCTTAAGCGCGCGACCTATA | 9 (0.000170%) | | | GGTGGTTCGCCGCCCGCGACGTCGCGAGAAGTCCACTAAACCTTATCATTTAGAGGAAGGAGAAGTCGTAACAAGGTTTCCGTAGGTGAACCTGCGGAAG | 14 (0.000265%) | | | GGTGTCACAAATCGTCGTCCCTCACCATCCTTTGCTGATGCGGGACGGAAGCTGGTCTCCCGTGTGTTACCGCACGCGGTTGGCCTAAATCCGAGCCAAG | 6 (0.000114%) | | | GGTGTGAGTGTCGCCCATGGGCATCGACACCTTGCGGCTAGGAACTGGAACGAGACGGGTAGCAAAGATTTCGAGTAGCACTTCATACTACCGTGGGTTT | 3 (0.000057%) | | | GGTGTGAGTGTCGCCCATGGGCATCGACACCTTGCGGCTAGGAACTGGAACGAGACGGGTGGCAAAGATTTCGAGTAGCACTTCATACTACCGTGGGTTT | 3 (0.000057%) | | | GGTGTTGGTCGATTAAGACAGCAGGACGGTGGTCATGGAAGTCGAAATCCGCTAAGGAGTGTGTAACAACTCACCTGCCGAATCAACTAGCCCCGAAAAT | 4 (0.000076%) | | | GGTTCACATTTCGTTCATCACCCTTGGCCGGCTATCGAACAGCCGGACTCCCATCAAAAGATGGTTGCCAAGAACATCTTCGTTACGGTTTGCTAATTCT | 4 (0.000076%) | | | GGTTCACATTTCGTTCATCACCCTTGGCCGGCTTTCGAACAGCCGGACTCCCATCAAAAGATGGTTGCCAAGAACATCTTCGTTACGGTTTGCTAATTCT | 3 (0.000057%) | | | GGTTCGCCGCCCGCGACGTCGCGAGAAGTCCACTAAACCTTATCATTTAGAGGAAGGAGAAGTCGTAACAAGGTTTCCGTAGGTGAACCTGCGGAAGGAT | 15 (0.000284%) | | | GGTTCGTATTCGTACTGAAAATCAGAATCAAACGAGCTTTTACCCTTTTGTTCCACACGAGATTTCTGTTCTCGTTGAGCTCATCTTAGGACACCTGCGT | 12 (0.000227%) | | | GGTTGAAATCGTCGACCAGGTCCGAGACTTCATCGACCGGGTCCGAGGATTCGTCGACCAGGACGGCCGGATGTCCGAGAAAAAAAAATGTTGCCGAATA | 14 (0.000265%) | | | GGTTGACGGCAACGTTAGGGAGTCCGGAGACGTCGGCGGGGGCCTCGGGAAGAGTTATCTTTTCTGTTTAACAGCCTGCCCACCCTGGAAACGGCTCAGC | 14 (0.000265%) | | | GGTTGCGGTTTAAGTTCTTATACTCAATCATACACATGACATCAAGTCATATTCGACTCCAAAACACTAACCAACCTTCTTCTTGCTTCTCAAAGCTTTC | 1252 (0.023686%) | | | GGTTGCGGTTTAAGTTGTTATACTCAATCATACACATGACAACAAGTCATATTCGACTCCAAAACACTAACCAACCTTCTTCTTGCTTCTCAAAGCTTTC | 32 (0.000605%) | | | GGTTTAAGTTCTTATACTCAATCATACACATGACATCAAGTCATATTCGACTCCAAAACACTAACCAACCTTCTTCTTGCTTCTCAAAGCTTTCATGGTG | 241 (0.004559%) | | | GGTTTAAGTTGTTATACTCAATCATACACATGACAACAAGTCATATTCGACTCCAAAACACTAACCAACCTTCTTCTTGCTTCTCAAAGCTTTCATGGTG | 99 (0.001873%) | | | GGTTTACTCACCCGTTGACTCGCACACATGTCAGACTCCTTGGTCCGTGTTTCAAGACGGGTCGAATGGGGAGCCCACAGGCCGACGCCCGGAGCACGCT | 26 (0.000492%) | | | GTAAAGTTGGGAATTCGTTAAGGAGCTGTTGCTTTGTTAGTGTAGAAACACTTGTGTAGAATTGGGGATTGTTTTTTTTGGAGTGATTTAGGGGAGGGTC | 57 (0.001078%) | | | GTAAGAATTGTATCCTTGTTAGAAGACACAAAGCCAAAGA | 342 (0.002588%) | | | GTAATTCCAGCTCCAATAGCGTATATTTAAGTTGTTGCAGTTAAAAAGCTCGTAGTTGAACCTTGGGATGGGTCGGCCGGTCCGCCTTTGGTGTGCATTG | 12 (0.000227%) | | | GTACGAAACACAGGCCCCGGAACTCATCATCGAGCGTAACATCGCCCGTGAATTAACTGAGAAGGATAGGTGGTAGGTAGTTCGATGCGCGAGCATGGAG | 20 (0.000378%) | | | GTACGGCTTTGGCTCGGATTCGTCCGTCTTCTTTCTTCTTAGCCGAGTACTTCGGTAGATTAGTTGGAACGATTGATGATTTTGAGTTAATTGAACGTTC | 7 (0.000132%) | | | GTAGGAGCGACGGGCGGTGTGTACAAAGGGCAGGGACGTAGTCAACGCGAGCTGATGACTCGCGCTTACTAGGAATTCCTCGTTGAAGACCAACAATTGC | 90 (0.001703%) | | | GTAGGCTCCATGCTCGCGCATCGAACTACCTACCACCTATCCTTCTCAGTTAATTCACGGGCGATGTTACGCTCGATGATGAGTTCCGGGGCCTGTGTTT | 19 (0.000359%) | | | GTAGGGAAAGAGTGTTCAGAGCCGTGTAGATCTCGGTGGT | 39 (0.000295%) | | | GTAGTCATATGCTTGTCTCAAAGATTAAGCCATGCATGTGTAAGTATGAACGAATTCAGACTGTGAAACTGCGAATGGCTCATTAAATCAGTTATAGTTT | 83 (0.001570%) | | | GTATAAGAACTTAAACCGCAACCGCATCTTATAAGCCTAAGTAGTGTTTCCTTGTTAGAAGACACAAAGCCAAAGACTCATATGGACTTTGGCTACACCA | 10 (0.000189%) | | | GTATATTTAAGTTGTTGCAGTTAAAAAGCTCGTAGTTGAACCTTGGGATGGGTCGGCCGGTCCGCCTTTGGTGTGCATTGGTCGGCTTGTCCCTTCGGTC | 15 (0.000284%) | | | GTATCCAGAGCGTAGGCTTGCTTTGAGCACTCTAATTTCTTCAAAGTAACAGCGCCGGAGGCACGACCCGGCCAATTAAGACCAGGAGCGTATCGCCGAC | 17 (0.000322%) | | | GTATCCTTGTTAGAAGACACAAAGCCAAAGACTCATATGGACTTTGGCTACACCATGAAAGCTTTGAGAAGCAAGAAGAAGGTTGGTTAGTGTTTTGGAG | 16 (0.000303%) | | | GTATGATTGAGTATAAGAACTTAAACCGCAACCGCATCTTATAAGCCTAAGTAGTGTTTCCTTGTTAGAAGACACAAAGCCAAAGACTCATATGGACTTT | 51 (0.000965%) | | | GTATTTCATAGTCAGAGGTGAAATTCTTGGATTTATGAAAGACGAACAACTGCGAAAGCATTTGCCAAGGATGTTTTCATTAATCAAGAACGAAAGTTGG | 4 (0.000076%) | | | GTCACAAATCGTCGTCCCTCACCATCCTTTGCTGATGCGGGACGGAAGCTGGTCTCCCGTGTGTTACCGCACGCGGTTGGCCTAAATCCGAGCCAAGGAC | 5 (0.000095%) | | | GTCAGGTGGGGAGTTTGGCTGGGGCGGCACATCTGTTAAAAGATAACGCAGGTGTCCTAAGATGAGCTCAACGAGAACAGAAATCTCGTGTGGAACAAAA | 15 (0.000284%) | | | GTCATATGCTTGTCTCAAAGATTAAGCCATGCATGTGTAAGTATGAACGAATTCAGACTGTGAAACTGCGAATGGCTCATTAAATCAGTTATAGTTTGTT | 43 (0.000813%) | | | GTCATATTCGACTCCAAAACACTAACCAACCTTCTTCTTGCTTCTCAAAGCTTTCATGGTGTAGCCAAAGTCCATATGAGTCTTTGGCTTTGTGTCTTCT | 202 (0.003821%) | | | GTCCCGAAGGTATCTCGCGCTTGTACGGCTTTGGCTCGGATTCGTCCGTCTTCTTTCTTCTTAGCCGAGTACTTCGGTAGATTAGTTGGAACGATTGATG | 3 (0.000057%) | | | GTCCCGAGTGTGAGCGAGGTGTGAGTGTCGCCCATGGGCATCGACACCTTGCGGCTAGGAACTGGAACGAGACGGGTAGCAAAGATTTCGAGTAGCACTT | 20 (0.000378%) | | | GTCCCGAGTGTGAGCGAGGTGTGAGTGTCGCCCATGGGCATCGACACCTTGCGGCTAGGAACTGGAACGAGACGGGTGGCAAAGATTTCGAGTAGCACTT | 19 (0.000359%) | | | GTCCCTCACCATCCTTTGCTGATGCGGGACGGAAGCTGGTCTCCCGTGTGTTACCGCACGCGGTTGGCCTAAATCCGAGCCAAGGACGCCTGGAGCGTAC | 18 (0.000341%) | | | GTCCGGAGACGTCGGCGGGGGCCTCGGGAAGAGTTATCTTTTCTGTTTAACAGCCTGCCCACCCTGGAAACGGCTCAGCCGGAGGTAGGGTCCAGCGGCT | 9 (0.000170%) | | | GTCCTCCGGATTTTCAAGGGCCGCCGGGGGCGCACCGGACACCACGCGACGTGCGGTGCTCTTCCAGCCGCTGGACCCTACCTCCGGCTGAGCCGTTTCC | 5 (0.000095%) | | | GTCGAATATGACTTGATGTCATGTGTATGATTGAGTATAA | 486 (0.003678%) | | | GTCGACCAGGGGTTGAAATCGTCGACCAGGTCCGAGACTTCATCGACCGGGTCCGAGGATTCGTCGACCAGGACGGCCGGATGTCCGAGAAAAAAAAATG | 53 (0.001003%) | | | GTCGAGTTATCATGAATCATCAGAGCAACGGGCAGAGCCCGCGTCGACCTTTTATCTAATAAATGCGTCCCTTCCATAAGTCGGGGTTTGTTGCACGTAT | 423 (0.008002%) | | | GTCGATTAAGACAGCAGGACGGTGGTCATGGAAGTCGAAATCCGCTAAGGAGTGTGTAACAACTCACCTGCCGAATCAACTAGCCCCGAAAATGGATGGC | 29 (0.000549%) | | | GTCGCCCATGGGCATCGACACCTTGCGGCTAGGAACTGGAACGAGACGGGTGGCAAAGATTTCGAGTAGCACTTCATACTACCGTGGGTTTTTTAAACCT | 23 (0.000435%) | | | GTCGGAGTTTTTTCAGCAGTTCTCGGACAAAAATTGCTGAGTGGCCGAGAAGAATGGGCGTGTCATGCGTGGGCTGACATGGATTCTTCGAGGCCTAGGG | 3 (0.000057%) | | | GTCGGGGGCATTCGTATTTCATAGTCAGAGGTGAAATTCTTGGATTTATGAAAGACGAACAACTGCGAAAGCATTTGCCAAGGATGTTTTCATTAATCAA | 6 (0.000114%) | | | GTCGTCCCTCACCATCCTTTGCTGATGCGGGACGGAAGCTGGTCTCCCGTGTGTTACCGCACGCGGTTGGCCTAAATCCGAGCCAAGGACGCCTGGAGCG | 32 (0.000605%) | | | GTCGTGTAGGGAAAGAGTGTTCAGAGCCGTGTAGATCTCG | 16 (0.000121%) | | | GTCTGGTGCCAGCAGCCGCGGTAATTCCAGCTCCAATAGCGTATATTTAAGTTGTTGCAGTTAAAAAGCTCGTAGTTGAACCTTGGGATGGGTCGGCCGG | 23 (0.000435%) | | | GTCTTCAACGAGGAATTCCTAGTAAGCGCGAGTCATCAGCTCGCGTTGACTACGTCCCTGCCCTTTGTACACACCGCCCGTCGCTCCTACCGATTGAATG | 16 (0.000303%) | | | GTGAGCGAGGTGTGAGTGTCGCCCATGGGCATCGACACCTTGCGGCTAGGAACTGGAACGAGACGGGTGGCAAAGATTTCGAGTAGCACTTCATACTACC | 30 (0.000568%) | | | GTGAGCGGCACTCGGTCCTCCGGATTTTCAAGGGCCGCCGGGGGCGCACCGGACACCACGCGACGTGCGGTGCTCTTCCAGCCGCTGGACCCTACCTCCG | 363 (0.006867%) | | | GTGAGTGTCGCCCATGGGCATCGACACCTTGCGGCTAGGAACTGGAACGAGACGGGTGGCAAAGATTTCGAGTAGCACTTCATACTACCGTGGGTTTTTT | 46 (0.000870%) | | | GTGATATGAACACAAACGTTCAATATGACAAACCCATGCCAAGTAAAGAGAAAATGAAAACTGGTGATTGTTGCGGAAATCGTCCAGGATTCCTCGACCA | 3 (0.000057%) | | | GTGCCAGCAGCCGCGGTAATTCCAGCTCCAATAGCGTATATTTAAGTTGTTGCAGTTAAAAAGCTCGTAGTTGAACCTTGGGATGGGTCGGCCGGTCCGC | 26 (0.000492%) | | | GTGCCTCGGCATCAGCGTGCTCCGGGCGTCGGCCTGTGGGCTCCCCATTCGACCCGTCTTGAAACACGGACCAAGGAGTCTGACATGTGTGCGAGTCAAC | 8 (0.000151%) | | | GTGCTCCGGGCGTCGGCCTGTGGGCTCCCCATTCGACCCGTCTTGAAACACGGACCAAGGAGTCTGACATGTGTGCGAGTCAACGGGTGAGTAAACCCGT | 13 (0.000246%) | | | GTGCTGGCGACGCATCATTCAAATTTCTGCCCTATCAACTTTCGATGGTAGGATAGTGGCCTACCATGGTGGTAACGGGTGACGGAGAATTAGGGTTCGA | 40 (0.000757%) | | | GTGGCGGTTGACGGCAACGTTAGGGAGTCCGGAGACGTCGGCGGGGGCCTCGGGAAGAGTTATCTTTTCTGTTTAACAGCCTGCCCACCCTGGAAACGGC | 10 (0.000189%) | | | GTGGGGAGTTTGGCTGGGGCGGCACATCTGTTAAAAGATAACGCAGGTGTCCTAAGATGAGCTCAACGAGAACAGAAATCTCGTGTGGAACAAAAGGGTA | 9 (0.000170%) | | | GTGGGTGGTTCGCCGCCCGCGACGTCGCGAGAAGTCCACTAAACCTTATCATTTAGAGGAAGGAGAAGTCGTAACAAGGTTTCCGTAGGTGAACCTGCGG | 86 (0.001627%) | | | GTGGTCATGGAAGTCGAAATCCGCTAAGGAGTGTGTAACAACTCACCTGCCGAATCAACTAGCCCCGAAAATGGATGGCGCTTAAGCGCGCGACCTATAC | 12 (0.000227%) | | | GTGGTTCGCCGCCCGCGACGTCGCGAGAAGTCCACTAAACCTTATCATTTAGAGGAAGGAGAAGTCGTAACAAGGTTTCCGTAGGTGAACCTGCGGAAGG | 13 (0.000246%) | | | GTGTAAAGTTGGGAATTCGTTAAGGAGCTGTTGCTTTGTTAGTGTAGAAACACTTGTGTAGAATTGGGGATTGTTTTTTTTGGAGTGATTTAGGGGAGGG | 18 (0.000341%) | | | GTGTAGGGAAAGAGTGTTCAGAGCCGTGTAGATCTCGGTG | 25 (0.000189%) | | | GTGTATCCTTGTTAGAAGACACAAAGCCAAAGACTCATAT | 69 (0.000522%) | | | GTGTATGATTGAGTATAAGAACTTAAACCGCAACCGCATCTTATAAGCCTAAGTAGTGTTTCCTTGTTAGAAGACACAAAGCCAAAGACTCATATGGACT | 11 (0.000208%) | | | GTGTCACAAATCGTCGTCCCTCACCATCCTTTGCTGATGCGGGACGGAAGCTGGTCTCCCGTGTGTTACCGCACGCGGTTGGCCTAAATCCGAGCCAAGG | 7 (0.000132%) | | | GTGTCGCCCATGGGCATCGACACCTTGCGGCTAGGAACTGGAACGAGACGGGTGGCAAAGATTTCGAGTAGCACTTCATACTACCGTGGGTTTTTTAAAC | 6 (0.000114%) | | | GTGTGAGCATGCCTGTCGGGACCCGAAAGATGGTGAACTATGCCTGAGCGGGGTAAAGCCAGAGGAAACTCTGGTGGAAGCCCGCAGCGATACTGACGTG | 3 (0.000057%) | | | GTGTGAGCGAGGTGTGAGTGTCGCCCATGGGCATCGACACCTTGCGGCTAGGAACTGGAACGAGACGGGTAGCAAAGATTTCGAGTAGCACTTCATACTA | 22 (0.000416%) | | | GTGTGAGCGAGGTGTGAGTGTCGCCCATGGGCATCGACACCTTGCGGCTAGGAACTGGAACGAGACGGGTGGCAAAGATTTCGAGTAGCACTTCATACTA | 30 (0.000568%) | | | GTGTGAGTGTCGCCCATGGGCATCGACACCTTGCGGCTAGGAACTGGAACGAGACGGGTAGCAAAGATTTCGAGTAGCACTTCATACTACCGTGGGTTTT | 16 (0.000303%) | | | GTGTGAGTGTCGCCCATGGGCATCGACACCTTGCGGCTAGGAACTGGAACGAGACGGGTGGCAAAGATTTCGAGTAGCACTTCATACTACCGTGGGTTTT | 8 (0.000151%) | | | GTGTTCGTAGGCTCCATGCTCGCGCATCGAACTACCTACCACCTATCCTTCTCAGTTAATTCACGGGCGATGTTACGCTCGATGATGAGTTCCGGGGCCT | 7 (0.000132%) | | | GTGTTGGTCGATTAAGACAGCAGGACGGTGGTCATGGAAGTCGAAATCCGCTAAGGAGTGTGTAACAACTCACCTGCCGAATCAACTAGCCCCGAAAATG | 4 (0.000076%) | | | GTGTTTCCTTGTTAGAAGACACAAAGCCAAAGACTCATATGGACTTTGGCTACACCATGAAAGCTTTGAGAAGCAAGAAGAAGGTTGGTTAGTGTTTTGG | 28 (0.000530%) | | | GTGTTTTGGAGTCGAATATGACTTGATGTCATGTGTATGA | 215 (0.001627%) | | | GTTAAAAGATAACGCAGGTGTCCTAAGATGAGCTCAACGAGAACAGAAATCTCGTGTGGAACAAAAGGGTAAAAGCTCGTTTGATTCTGATTTTCAGTAC | 32 (0.000605%) | | | GTTAAGGAGCTGTTGCTTTGTTAGTGTAGAAACACTTGTGTAGAATTGGGGATTGTTTTTTTTGGAGTGATTTAGGGGAGGGTCGAATCTTAGCGACAAA | 40 (0.000757%) | | | GTTAGAAGACACAAAGCCAAAGACTCATATGGACTTTGGCTACACCATGAAAGCTTTGAGAAGCAAGAAGAAGGTTGGTTAGTGTTTTGGAGTCGAATAT | 80 (0.001513%) | | | GTTAGGGAGTCCGGAGACGTCGGCGGGGGCCTCGGGAAGAGTTATCTTTTCTGTTTAACAGCCTGCCCACCCTGGAAACGGCTCAGCCGGAGGTAGGGTC | 16 (0.000303%) | | | GTTAGTGTTTTGGAGTCGAATATGACTTGATGTCATGTGT | 16246 (0.122938%) | | | GTTATCATGAATCATCAGAGCAACGGGCAGAGCCCGCGTCGACCTTTTATCTAATAAATGCGTCCCTTCCATAAGTCGGGGTTTGTTGCACGTATTAGCT | 282 (0.005335%) | | | GTTATCCCATGCTAATGTATCCAGAGCGTAGGCTTGCTTTGAGCACTCTAATTTCTTCAAAGTAACAGCGCCGGAGGCACGACCCGGCCAATTAAGACCA | 47 (0.000889%) | | | GTTCAATATGACAAACCCATGCCAAGTAAAGAGAAAATGAAAACTGGTGATTGTTGCGGAAATCGTCCAGGATTCCTCGACCAGGACTTGAAATCGTCGA | 25 (0.000473%) | | | GTTCACATTTCGTTCATCACCCTTGGCCGGCTATCGAACAGCCGGACTCCCATCAAAAGATGGTTGCCAAGAACATCTTCGTTACGGTTTGCTAATTCTC | 15 (0.000284%) | | | GTTCACATTTCGTTCATCACCCTTGGCCGGCTTTCGAACAGCCGGACTCCCATCAAAAGATGGTTGCCAAGAACATCTTCGTTACGGTTTGCTAATTCTC | 18 (0.000341%) | | | GTTCATCACCCTTGGCCGGCTTTCGAACAGCCGGACTCCCATCAAAAGATGGTTGCCAAGAACATCTTCGTTACGGTTTGCTAATTCTCGGAATAACATC | 9 (0.000170%) | | | GTTCGAGTGTGAGCATGCCTGTCGGGACCCGAAAGATGGTGAACTATGCCTGAGCGGGGTAAAGCCAGAGGAAACTCTGGTGGAAGCCCGCAGCGATACT | 4 (0.000076%) | | | GTTCGATTAGTCTTTCGCCCCTATACCCAAGTCAGACGAACGATTTGCACGTCAGTATCGCTGCGGGCTTCCACCAGAGTTTCCTCTGGCTTTACCCCGC | 784 (0.014832%) | | | GTTCGCCGCCCGCGACGTCGCGAGAAGTCCACTAAACCTTATCATTTAGAGGAAGGAGAAGTCGTAACAAGGTTTCCGTAGGTGAACCTGCGGAAGGATC | 33 (0.000624%) | | | GTTCGTAGGCTCCATGCTCGCGCATCGAACTACCTACCACCTATCCTTCTCAGTTAATTCACGGGCGATGTTACGCTCGATGATGAGTTCCGGGGCCTGT | 17 (0.000322%) | | | GTTCGTATTCGTACTGAAAATCAGAATCAAACGAGCTTTTACCCTTTTGTTCCACACGAGATTTCTGTTCTCGTTGAGCTCATCTTAGGACACCTGCGTT | 12 (0.000227%) | | | GTTCTTATACTCAATCATACACATGACATCAAGTCATATTCGACTCCAAAACACTAACCAACCTTCTTCTTGCTTCTCAAAGCTTTCATGGTGTAGCCAA | 86 (0.001627%) | | | GTTGAAATCGTCGACCAGGTCCGAGACTTCATCGACCGGGTCCGAGGATTCGTCGACCAGGACGGCCGGATGTCCGAGAAAAAAAAATGTTGCCGAATAA | 46 (0.000870%) | | | GTTGACGGCAACGTTAGGGAGTCCGGAGACGTCGGCGGGGGCCTCGGGAAGAGTTATCTTTTCTGTTTAACAGCCTGCCCACCCTGGAAACGGCTCAGCC | 38 (0.000719%) | | | GTTGACTCGCACACATGTCAGACTCCTTGGTCCGTGTTTCAAGACGGGTCGAATGGGGAGCCCACAGGCCGACGCCCGGAGCACGCTGATGCCGAGGCAC | 47 (0.000889%) | | | GTTGCGGTTTAAGTTCTTATACTCAATCATACACATGACATCAAGTCATATTCGACTCCAAAACACTAACCAACCTTCTTCTTGCTTCTCAAAGCTTTCA | 277 (0.005240%) | | | GTTGCGGTTTAAGTTGTTATACTCAATCATACACATGACAACAAGTCATATTCGACTCCAAAACACTAACCAACCTTCTTCTTGCTTCTCAAAGCTTTCA | 148 (0.002800%) | | | GTTGGTCGATTAAGACAGCAGGACGGTGGTCATGGAAGTCGAAATCCGCTAAGGAGTGTGTAACAACTCACCTGCCGAATCAACTAGCCCCGAAAATGGA | 17 (0.000322%) | | | GTTGGTCTTCAACGAGGAATTCCTAGTAAGCGCGAGTCATCAGCTCGCGTTGACTACGTCCCTGCCCTTTGTACACACCGCCCGTCGCTCCTACCGATTG | 174 (0.003292%) | | | GTTGTTATACTCAATCATACACATGACAACAAGTCATATTCGACTCCAAAACACTAACCAACCTTCTTCTTGCTTCTCAAAGCTTTCATGGTGTAGCCAA | 22 (0.000416%) | | | GTTGTTATACTCAATCATACACATGACATCAAGTCATATT | 256 (0.001937%) | | | GTTTAAGTTCTTATACTCAATCATACACATGACATCAAGTCATATTCGACTCCAAAACACTAACCAACCTTCTTCTTGCTTCTCAAAGCTTTCATGGTGT | 409 (0.007738%) | | | GTTTAAGTTGTTATACTCAATCATACACATGACAACAAGTCATATTCGACTCCAAAACACTAACCAACCTTCTTCTTGCTTCTCAAAGCTTTCATGGTGT | 118 (0.002232%) | | | GTTTACTCACCCGTTGACTCGCACACATGTCAGACTCCTTGGTCCGTGTTTCAAGACGGGTCGAATGGGGAGCCCACAGGCCGACGCCCGGAGCACGCTG | 26 (0.000492%) | | | GTTTGGCTGGGGCGGCACATCTGTTAAAAGATAACGCAGGTGTCCTAAGATGAGCTCAACGAGAACAGAAATCTCGTGTGGAACAAAAGGGTAAAAGCTC | 32 (0.000605%) | | | GTTTTAATTAAACAGTCGGATTCCCCTTGTCCGTACCAGTTCTGAGCTGACTGTTCGACGCCCGGGGAAAGCTCCCGAGAGAGCCGTTCCCAGTCCGTCC | 25 (0.000473%) | | | GTTTTGGAGTCGAATATGACTTGATGTCATGTGTATGATT | 475 (0.003594%) | | | GTTTTTTCAGCAGTTCTCGGACAAAAATTGCTGAGTGGCCGAGAAGAATGGGCGTGTCATGCGTGGGCTGACATGGATTCTTCGAGGCCTAGGGGTGGCG | 19 (0.000359%) | | | TAAAAGATAACGCAGGTGTCCTAAGATGAGCTCAACGAGAACAGAAATCTCGTGTGGAACAAAAGGGTAAAAGCTCGTTTGATTCTGATTTTCAGTACGA | 22 (0.000416%) | | | TAAAAGTGTCGGAGTTTTTTCAGCAGTTCTCGGACAAAAATTGCTGAGTGGCCGAGAAGAATGGGCGTGTCATGCGTGGGCTGACATGGATTCTTCGAGG | 6 (0.000114%) | | | TAAAGGCGTAAGAATTGTATCCTTGTTAGAAGACACAAAG | 576 (0.004359%) | | | TAAAGTTGGGAATTCGTTAAGGAGCTGTTGCTTTGTTAGTGTAGAAACACTTGTGTAGAATTGGGGATTGTTTTTTTTGGAGTGATTTAGGGGAGGGTCG | 7 (0.000132%) | | | TAAATACGGGCGAGAGACCGATAGCGAACAAGTACCGCGAGGTAAAGATGAAAAGGACTTTGAAAAGAGAGTCAAAGAGTGCTTGAAATTGTCGGGAGGG | 22 (0.000416%) | | | TAACAGGGACAGTCGGGGGCATTCGTATTTCATAGTCAGAGGTGAAATTCTTGGATTTATGAAAGACGAACAACTGCGAAAGCATTTGCCAAGGATGTTT | 55 (0.001041%) | | | TAACGGCGTGCCTCGGCATCAGCGTGCTCCGGGCGTCGGCCTGTGGGCTCCCCATTCGACCCGTCTTGAAACACGGACCAAGGAGTCTGACATGTGTGCG | 55 (0.001041%) | | | TAACTAGCTACGTGGAGGCATCCCTTCACGGCCGGCTTCTTAGAGGGACTATGGCCGTTTAGGCCAAGGAAGTTTGAGGCAATAACAGGTCTGTGATGCC | 16 (0.000303%) | | | TAAGAACTTAAACCGCAACCGCATCTTATAAGCCTAAGTAGTGTTTCCTTGTTAGAAGACACAAAGCCAAAGACTCATATGGACTTTGGCTACACCATGA | 26 (0.000492%) | | | TAAGAATTGTATCCTTGTTAGAAGACACAAAGCCAAAGACTCATATGGACTTTGGCTACACCATGAAAGCTTTGAGAAGCAAGAAGAAGGTTGGTTAGTG | 1088 (0.020583%) | | | TAAGACAGCAGGACGGTGGTCATGGAAGTCGAAATCCGCTAAGGAGTGTGTAACAACTCACCTGCCGAATCAACTAGCCCCGAAAATGGATGGCGCTTAA | 55 (0.001041%) | | | TAAGGAGCTGTTGCTTTGTTAGTGTAGAAACACTTGTGTAGAATTGGGGATTGTTTTTTTTGGAGTGATTTAGGGGAGGGTCGAATCTTAGCGACAAAGG | 24 (0.000454%) | | | TAAGTAGTGTTTCCTTGTTAGAAGACACAAAGCCAAAGACTCATATGGACTTTGGCTACACCATGAAAGCTTTGAGAAGCAAGAAGAAGGTTGGTTAGTG | 44 (0.000832%) | | | TAAGTTCTTATACTCAATCATACACATGACATCAAGTCATATTCGACTCCAAAACACTAACCAACCTTCTTCTTGCTTCTCAAAGCTTTCATGGTGTAGC | 41 (0.000776%) | | | TAAGTTGTTATACTCAATCATACACATGACAACAAGTCATATTCGACTCCAAAACACTAACCAACCTTCTTCTTGCTTCTCAAAGCTTTCATGGTGTAGC | 10 (0.000189%) | | | TAAGTTGTTATACTCAATCATACACATGACATCAAGTCAT | 9616 (0.072767%) | | | TAATTCCAGCTCCAATAGCGTATATTTAAGTTGTTGCAGTTAAAAAGCTCGTAGTTGAACCTTGGGATGGGTCGGCCGGTCCGCCTTTGGTGTGCATTGG | 10 (0.000189%) | | | TACACATGACATCAAGTCATATTCGACTCCAAAACACTAACCAACCTTCTTCTTGCTTCTCAAAGCTTTCATGGTGTAGCCAAAGTCCATATGAGTCTTT | 23 (0.000435%) | | | TACATTGTTCCATCGACCAGAGGCTGTTCACCTTGGAGACCTGATGCGGTTATGAGTACGACCGGGCGTGAGCGGCACTCGGTCCTCCGGATTTTCAAGG | 3 (0.000057%) | | | TACATTTTATCGGTCGCTCTTGTCCGGAAGCTGTAGATGACCCAAAGTCCATATAGCGACCCCAGGTCAGGCGGGATTACCCGCTGAGTTTAAGCATATC | 14 (0.000265%) | | | TACCTACCACCTATCCTTCTCAGTTAATTCACGGGCGATGTTACGCTCGATGATGAGTTCCGGGGCCTGTGTTTCGTACCTAATTTGAAGGAATTGTTGA | 3 (0.000057%) | | | TACGAAACACAGGCCCCGGAACTCATCATCGAGCGTAACATCGCCCGTGAATTAACTGAGAAGGATAGGTGGTAGGTAGTTCGATGCGCGAGCATGGAGC | 6 (0.000114%) | | | TACGGCTTTGGCTCGGATTCGTCCGTCTTCTTTCTTCTTAGCCGAGTACTTCGGTAGATTAGTTGGAACGATTGATGATTTTGAGTTAATTGAACGTTCG | 3 (0.000057%) | | | TACGGGTTTACTCACCCGTTGACTCGCACACATGTCAGACTCCTTGGTCCGTGTTTCAAGACGGGTCGAATGGGGAGCCCACAGGCCGACGCCCGGAGCA | 6 (0.000114%) | | | TACTCAATCATACACATGACATCAAGTCATATTCGACTCCAAAACACTAACCAACCTTCTTCTTGCTTCTCAAAGCTTTCATGGTGTAGCCAAAGTCCAT | 23 (0.000435%) | | | TACTCACCCGTTGACTCGCACACATGTCAGACTCCTTGGTCCGTGTTTCAAGACGGGTCGAATGGGGAGCCCACAGGCCGACGCCCGGAGCACGCTGATG | 6 (0.000114%) | | | TAGAAGACACAAAGCCAAAGACTCATATGGACTTTGGCTACACCATGAAAGCTTTGAGAAGCAAGAAGAAGGTTGGTTAGTGTTTTGGAGTCGAATATGA | 38 (0.000719%) | | | TAGCTAGTGTTCGTAGGCTCCATGCTCGCGCATCGAACTACCTACCACCTATCCTTCTCAGTTAATTCACGGGCGATGTTACGCTCGATGATGAGTTCCG | 4 (0.000076%) | | | TAGGAGCGACGGGCGGTGTGTACAAAGGGCAGGGACGTAGTCAACGCGAGCTGATGACTCGCGCTTACTAGGAATTCCTCGTTGAAGACCAACAATTGCA | 12 (0.000227%) | | | TAGGCGAGACAAGGGTTCACATTTCGTTCATCACCCTTGGCCGGCTATCGAACAGCCGGACTCCCATCAAAAGATGGTTGCCAAGAACATCTTCGTTACG | 4 (0.000076%) | | | TAGGCGAGACAAGGGTTCACATTTCGTTCATCACCCTTGGCCGGCTTTCGAACAGCCGGACTCCCATCAAAAGATGGTTGCCAAGAACATCTTCGTTACG | 3 (0.000057%) | | | TAGGCTGTCCCGAAGGTATCTCGCGCTTGTACGGCTTTGGCTCGGATTCGTCCGTCTTCTTTCTTCTTAGCCGAGTACTTCGGTAGATTAGTTGGAACGA | 1809 (0.034223%) | | | TAGGCTGTCCCGAGTGTGAGCGAGGTGTGAGTGTCGCCCATGGGCATCGACACCTTGCGGCTAGGAACTGGAACGAGACGGGTGGCAAAGATTTCGAGTA | 41 (0.000776%) | | | TAGGGAAAGAGTGTTCAGAGCCGTGTAGATCTCGGTGGTC | 13 (0.000098%) | | | TAGGGAGTCCGGAGACGTCGGCGGGGGCCTCGGGAAGAGTTATCTTTTCTGTTTAACAGCCTGCCCACCCTGGAAACGGCTCAGCCGGAGGTAGGGTCCA | 7 (0.000132%) | | | TAGGTACGAAACACAGGCCCCGGAACTCATCATCGAGCGTAACATCGCCCGTGAATTAACTGAGAAGGATAGGTGGTAGGTAGTTCGATGCGCGAGCATG | 16 (0.000303%) | | | TAGTCATATGCTTGTCTCAAAGATTAAGCCATGCATGTGTAAGTATGAACGAATTCAGACTGTGAAACTGCGAATGGCTCATTAAATCAGTTATAGTTTG | 32 (0.000605%) | | | TAGTGTTCGTAGGCTCCATGCTCGCGCATCGAACTACCTACCACCTATCCTTCTCAGTTAATTCACGGGCGATGTTACGCTCGATGATGAGTTCCGGGGC | 5 (0.000095%) | | | TAGTGTTTCCTTGTTAGAAGACACAAAGCCAAAGACTCATATGGACTTTGGCTACACCATGAAAGCTTTGAGAAGCAAGAAGAAGGTTGGTTAGTGTTTT | 23 (0.000435%) | | | TAGTGTTTTGGAGTCGAATATGACTTGATGTCATGTGTAT | 264 (0.001998%) | | | TATAAGAACTTAAACCGCAACCGCATCTTATAAGCCTAAGTAGTGTTTCCTTGTTAGAAGACACAAAGCCAAAGACTCATATGGACTTTGGCTACACCAT | 16 (0.000303%) | | | TATACTCAATCATACACATGACATCAAGTCATATTCGACTCCAAAACACTAACCAACCTTCTTCTTGCTTCTCAAAGCTTTCATGGTGTAGCCAAAGTCC | 36 (0.000681%) | | | TATCAACTTTCGATGGTAGGATAGTGGCCTACCATGGTGGTAACGGGTGACGGAGAATTAGGGTTCGATTCCGGAGAGGGAGCCTGAGAAACGGCTACCA | 3 (0.000057%) | | | TATCATGAATCATCAGAGCAACGGGCAGAGCCCGCGTCGACCTTTTATCTAATAAATGCGTCCCTTCCATAAGTCGGGGTTTGTTGCACGTATTAGCTCT | 64 (0.001211%) | | | TATCCAGAGCGTAGGCTTGCTTTGAGCACTCTAATTTCTTCAAAGTAACAGCGCCGGAGGCACGACCCGGCCAATTAAGACCAGGAGCGTATCGCCGACC | 7 (0.000132%) | | | TATCCTTGTTAGAAGACACAAAGCCAAAGACTCATATGGACTTTGGCTACACCATGAAAGCTTTGAGAAGCAAGAAGAAGGTTGGTTAGTGTTTTGGAGT | 35 (0.000662%) | | | TATGAACACAAACGTTCAATATGACAAACCCATGCCAAGTAAAGAGAAAATGAAAACTGGTGATTGTTGCGGAAATCGTCCAGGATTCCTCGACCAGGAC | 4 (0.000076%) | | | TATGATGTTATCCCATGCTAATGTATCCAGAGCGTAGGCTTGCTTTGAGCACTCTAATTTCTTCAAAGTAACAGCGCCGGAGGCACGACCCGGCCAATTA | 25 (0.000473%) | | | TATGATTGAGTATAAGAACTTAAACCGCAACCGCATCTTATAAGCCTAAGTAGTGTTTCCTTGTTAGAAGACACAAAGCCAAAGACTCATATGGACTTTG | 16 (0.000303%) | | | TATGCTTGTCTCAAAGATTAAGCCATGCATGTGTAAGTATGAACGAATTCAGACTGTGAAACTGCGAATGGCTCATTAAATCAGTTATAGTTTGTTTGAT | 7 (0.000132%) | | | TATTCGACTCCAAAACACTAACCAACCTTCTTCTTGCTTCTCAAAGCTTTCATGGTGTAGCCAAAGTCCATATGAGTCTTTGGCTTTGTGTCTTCTAACA | 48 (0.000908%) | | | TATTCGACTCCAAAACACTAACCAACCTTCTTCTTGCTTCTCAAAGCTTTCATGGTGTAGCCAAAGTCCATATGAGTCTTTGGCTTTGTGTCTTTTAACA | 8 (0.000151%) | | | TATTCGTACTGAAAATCAGAATCAAACGAGCTTTTACCCTTTTGTTCCACACGAGATTTCTGTTCTCGTTGAGCTCATCTTAGGACACCTGCGTTATCTT | 4 (0.000076%) | | | TATTTCATAGTCAGAGGTGAAATTCTTGGATTTATGAAAGACGAACAACTGCGAAAGCATTTGCCAAGGATGTTTTCATTAATCAAGAACGAAAGTTGGG | 3 (0.000057%) | | | TCAAAATCATCAATCGTTCCAACTAATCTACCGAAGTACTCGGCTAAGAAGAAAGAAGACGGACGAATCCGAGCCAAAGCCGTACAAGCGCGAGATACCT | 17 (0.000322%) | | | TCAAAGATTAAGCCATGCATGTGTAAGTATGAACGAATTCAGACTGTGAAACTGCGAATGGCTCATTAAATCAGTTATAGTTTGTTTGATGGTAACTACT | 8 (0.000151%) | | | TCAAATTAGGTACGAAACACAGGCCCCGGAACTCATCATCGAGCGTAACATCGCCCGTGAATTAACTGAGAAGGATAGGTGGTAGGTAGTTCGATGCGCG | 3 (0.000057%) | | | TCAAATTTCTGCCCTATCAACTTTCGATGGTAGGATAGTGGCCTACCATGGTGGTAACGGGTGACGGAGAATTAGGGTTCGATTCCGGAGAGGGAGCCTG | 16 (0.000303%) | | | TCAACTTTCGATGGTAGGATAGTGGCCTACCATGGTGGTAACGGGTGACGGAGAATTAGGGTTCGATTCCGGAGAGGGAGCCTGAGAAACGGCTACCACA | 3 (0.000057%) | | | TCAAGGGCCGCCGGGGGCGCACCGGACACCACGCGACGTGCGGTGCTCTTCCAGCCGCTGGACCCTACCTCCGGCTGAGCCGTTTCCAGGGTGGGCAGGC | 8 (0.000151%) | | | TCAAGTCATATTCGACTCCAAAACACTAACCAACCTTCTTCTTGCTTCTCAAAGCTTTCATGGTGTAGCCAAAGTCCATATGAGTCTTTGGCTTTGTGTC | 39 (0.000738%) | | | TCAATCATACACATGACATCAAGTCATATTCGACTCCAAAACACTAACCAACCTTCTTCTTGCTTCTCAAAGCTTTCATGGTGTAGCCAAAGTCCATATG | 34 (0.000643%) | | | TCAATCGGTAGGAGCGACGGGCGGTGTGTACAAAGGGCAGGGACGTAGTCAACGCGAGCTGATGACTCGCGCTTACTAGGAATTCCTCGTTGAAGACCAA | 45 (0.000851%) | | | TCACATTTCGTTCATCACCCTTGGCCGGCTATCGAACAGCCGGACTCCCATCAAAAGATGGTTGCCAAGAACATCTTCGTTACGGTTTGCTAATTCTCGG | 10 (0.000189%) | | | TCACATTTCGTTCATCACCCTTGGCCGGCTTTCGAACAGCCGGACTCCCATCAAAAGATGGTTGCCAAGAACATCTTCGTTACGGTTTGCTAATTCTCGG | 3 (0.000057%) | | | TCACCATCCTTTGCTGATGCGGGACGGAAGCTGGTCTCCCGTGTGTTACCGCACGCGGTTGGCCTAAATCCGAGCCAAGGACGCCTGGAGCGTACCGACA | 8 (0.000151%) | | | TCAGCCTGCTAACTAGCTACGTGGAGGCATCCCTTCACGGCCGGCTTCTTAGAGGGACTATGGCCGTTTAGGCCAAGGAAGTTTGAGGCAATAACAGGTC | 5 (0.000095%) | | | TCAGCGTGCTCCGGGCGTCGGCCTGTGGGCTCCCCATTCGACCCGTCTTGAAACACGGACCAAGGAGTCTGACATGTGTGCGAGTCAACGGGTGAGTAAA | 4 (0.000076%) | | | TCAGCTTCCTTGCGCCTTACGGGTTTACTCACCCGTTGACTCGCACACATGTCAGACTCCTTGGTCCGTGTTTCAAGACGGGTCGAATGGGGAGCCCACA | 26 (0.000492%) | | | TCATACACATGACATCAAGTCATATTCGACTCCAAAACACTAACCAACCTTCTTCTTGCTTCTCAAAGCTTTCATGGTGTAGCCAAAGTCCATATGAGTC | 5 (0.000095%) | | | TCATATGCTTGTCTCAAAGATTAAGCCATGCATGTGTAAGTATGAACGAATTCAGACTGTGAAACTGCGAATGGCTCATTAAATCAGTTATAGTTTGTTT | 6 (0.000114%) | | | TCATATTCGACTCCAAAACACTAACCAACCTTCTTCTTGCTTCTCAAAGCTTTCATGGTGTAGCCAAAGTCCATATGAGTCTTTGGCTTTGTGTCTTCTA | 68 (0.001286%) | | | TCATCAGAGCAACGGGCAGAGCCCGCGTCGACCTTTTATCTAATAAATGCGTCCCTTCCATAAGTCGGGGTTTGTTGCACGTATTAGCTCTAGAATTACT | 5 (0.000095%) | | | TCATCATCGAGCGTAACATCGCCCGTGAATTAACTGAGAAGGATAGGTGGTAGGTAGTTCGATGCGCGAGCATGGAGCCTACGAACACTAGCTATCCGAT | 227 (0.004294%) | | | TCATGAATCATCAGAGCAACGGGCAGAGCCCGCGTCGACCTTTTATCTAATAAATGCGTCCCTTCCATAAGTCGGGGTTTGTTGCACGTATTAGCTCTAG | 21 (0.000397%) | | | TCATGTGTATGATTGAGTATAAGAACTTAAACCGCAACCGCATCTTATAAGCCTAAGTAGTGTTTCCTTGTTAGAAGACACAAAGCCAAAGACTCATATG | 8 (0.000151%) | | | TCATTCAAATTTCTGCCCTATCAACTTTCGATGGTAGGATAGTGGCCTACCATGGTGGTAACGGGTGACGGAGAATTAGGGTTCGATTCCGGAGAGGGAG | 3 (0.000057%) | | | TCCAACCTAGGCGAGACAAGGGTTCACATTTCGTTCATCACCCTTGGCCGGCTATCGAACAGCCGGACTCCCATCAAAAGATGGTTGCCAAGAACATCTT | 3 (0.000057%) | | | TCCAACCTAGGCGAGACAAGGGTTCACATTTCGTTCATCACCCTTGGCCGGCTTTCGAACAGCCGGACTCCCATCAAAAGATGGTTGCCAAGAACATCTT | 4 (0.000076%) | | | TCCAACGAAGCACGCCCATCCAACCTAGGCGAGACAAGGGTTCACATTTCGTTCATCACCCTTGGCCGGCTATCGAACAGCCGGACTCCCATCAAAAGAT | 17 (0.000322%) | | | TCCAGAGCGTAGGCTTGCTTTGAGCACTCTAATTTCTTCAAAGTAACAGCGCCGGAGGCACGACCCGGCCAATTAAGACCAGGAGCGTATCGCCGACCGA | 4 (0.000076%) | | | TCCAGCTCCAATAGCGTATATTTAAGTTGTTGCAGTTAAAAAGCTCGTAGTTGAACCTTGGGATGGGTCGGCCGGTCCGCCTTTGGTGTGCATTGGTCGG | 50 (0.000946%) | | | TCCATGCTCGCGCATCGAACTACCTACCACCTATCCTTCTCAGTTAATTCACGGGCGATGTTACGCTCGATGATGAGTTCCGGGGCCTGTGTTTCGTACC | 3 (0.000057%) | | | TCCCATGCTAATGTATCCAGAGCGTAGGCTTGCTTTGAGCACTCTAATTTCTTCAAAGTAACAGCGCCGGAGGCACGACCCGGCCAATTAAGACCAGGAG | 3 (0.000057%) | | | TCCCGAAGGTATCTCGCGCTTGTACGGCTTTGGCTCGGATTCGTCCGTCTTCTTTCTTCTTAGCCGAGTACTTCGGTAGATTAGTTGGAACGATTGATGA | 3 (0.000057%) | | | TCCCGCCAATCAGCTTCCTTGCGCCTTACGGGTTTACTCACCCGTTGACTCGCACACATGTCAGACTCCTTGGTCCGTGTTTCAAGACGGGTCGAATGGG | 5 (0.000095%) | | | TCCCTTGCCTACATTGTTCCATCGACCAGAGGCTGTTCACCTTGGAGACCTGATGCGGTTATGAGTACGACCGGGCGTGAGCGGCACTCGGTCCTCCGGA | 5 (0.000095%) | | | TCCGCCTAGGCTGTCCCGAGTGTGAGCGAGGTGTGAGTGTCGCCCATGGGCATCGACACCTTGCGGCTAGGAACTGGAACGAGACGGGTGGCAAAGATTT | 2010 (0.038026%) | | | TCCGGAACCGGGACGTGGCGGTTGACGGCAACGTTAGGGAGTCCGGAGACGTCGGCGGGGGCCTCGGGAAGAGTTATCTTTTCTGTTTAACAGCCTGCCC | 21 (0.000397%) | | | TCCGGATTTTCAAGGGCCGCCGGGGGCGCACCGGACACCACGCGACGTGCGGTGCTCTTCCAGCCGCTGGACCCTACCTCCGGCTGAGCCGTTTCCAGGG | 5 (0.000095%) | | | TCCGGGCGTCGGCCTGTGGGCTCCCCATTCGACCCGTCTTGAAACACGGACCAAGGAGTCTGACATGTGTGCGAGTCAACGGGTGAGTAAACCCGTAAGG | 3 (0.000057%) | | | TCCGTCGAGTTATCATGAATCATCAGAGCAACGGGCAGAGCCCGCGTCGACCTTTTATCTAATAAATGCGTCCCTTCCATAAGTCGGGGTTTGTTGCACG | 22 (0.000416%) | | | TCCTATGATGTTATCCCATGCTAATGTATCCAGAGCGTAGGCTTGCTTTGAGCACTCTAATTTCTTCAAAGTAACAGCGCCGGAGGCACGACCCGGCCAA | 5 (0.000095%) | | | TCCTTGCGCCTTACGGGTTTACTCACCCGTTGACTCGCACACATGTCAGACTCCTTGGTCCGTGTTTCAAGACGGGTCGAATGGGGAGCCCACAGGCCGA | 6 (0.000114%) | | | TCCTTGTTAGAAGACACAAAGCCAAAGACTCATATGGACTTTGGCTACACCATGAAAGCTTTGAGAAGCAAGAAGAAGGTTGGTTAGTGTTTTGGAGTCG | 240 (0.004540%) | | | TCGAAATCCTATGATGTTATCCCATGCTAATGTATCCAGAGCGTAGGCTTGCTTTGAGCACTCTAATTTCTTCAAAGTAACAGCGCCGGAGGCACGACCC | 238 (0.004503%) | | | TCGACCAGGGGTTGAAATCGTCGACCAGGTCCGAGACTTCATCGACCGGGTCCGAGGATTCGTCGACCAGGACGGCCGGATGTCCGAGAAAAAAAAATGT | 9 (0.000170%) | | | TCGACCCGCCGAAGCGAGCCTTGGGACCAAAAACAGGGGTTGTACCCCGCCTCCGATTCACGGAGTAAGTAAAATAACGTTAAAAGTAGTGGTATTTCAC | 5 (0.000095%) | | | TCGACTCCAAAACACTAACCAACCTTCTTCTTGCTTCTCAAAGCTTTCATGGTGTAGCCAAAGTCCATATGAGTCTTTGGCTTTGTGTCTTCTAACAAGG | 43 (0.000813%) | | | TCGACTCCAAAACACTAACCAACCTTCTTCTTGCTTCTCAAAGCTTTCATGGTGTAGCCAAAGTCCATATGAGTCTTTGGCTTTGTGTCTTTTAACAAGG | 14 (0.000265%) | | | TCGAGTTATCATGAATCATCAGAGCAACGGGCAGAGCCCGCGTCGACCTTTTATCTAATAAATGCGTCCCTTCCATAAGTCGGGGTTTGTTGCACGTATT | 66 (0.001249%) | | | TCGATCCATTACATTTTATCGGTCGCTCTTGTCCGGAAGCTGTAGATGACCCAAAGTCCATATAGCGACCCCAGGTCAGGCGGGATTACCCGCTGAGTTT | 5 (0.000095%) | | | TCGATCCGTCGAGTTATCATGAATCATCAGAGCAACGGGCAGAGCCCGCGTCGACCTTTTATCTAATAAATGCGTCCCTTCCATAAGTCGGGGTTTGTTG | 4 (0.000076%) | | | TCGATCGACCCGCCGAAGCGAGCCTTGGGACCAAAAACAGGGGTTGTACCCCGCCTCCGATTCACGGAGTAAGTAAAATAACGTTAAAAGTAGTGGTATT | 5 (0.000095%) | | | TCGATCTCATGTGTATGATTGAGTATAAGAACTTAAACCGCAACCGCATCTTATAAGCCTAAGTAGTGTTTCCTTGTTAGAAGACACAAAGCCAAAGACT | 3 (0.000057%) | | | TCGATTAGTCTTTCGCCCCTATACCCAAGTCAGACGAACGATTTGCACGTCAGTATCGCTGCGGGCTTCCACCAGAGTTTCCTCTGGCTTTACCCCGCTC | 5 (0.000095%) | | | TCGCACACATGTCAGACTCCTTGGTCCGTGTTTCAAGACGGGTCGAATGGGGAGCCCACAGGCCGACGCCCGGAGCACGCTGATGCCGAGGCACGCCGTT | 6 (0.000114%) | | | TCGCCCATGGGCATCGACACCTTGCGGCTAGGAACTGGAACGAGACGGGTGGCAAAGATTTCGAGTAGCACTTCATACTACCGTGGGTTTTTTAAACCTT | 4 (0.000076%) | | | TCGCCGCCCGCGACGTCGCGAGAAGTCCACTAAACCTTATCATTTAGAGGAAGGAGAAGTCGTAACAAGGTTTCCGTAGGTGAACCTGCGGAAGGATCGA | 6 (0.000114%) | | | TCGCGCTTGTACGGCTTTGGCTCGGATTCGTCCGTCTTCTTTCTTCTTAGCCGAGTACTTCGGTAGATTAGTTGGAACGATTGATGATTTTGAGTTAATT | 3 (0.000057%) | | | TCGGCATCAGCGTGCTCCGGGCGTCGGCCTGTGGGCTCCCCATTCGACCCGTCTTGAAACACGGACCAAGGAGTCTGACATGTGTGCGAGTCAACGGGTG | 25 (0.000473%) | | | TCGGCCTGTGGGCTCCCCATTCGACCCGTCTTGAAACACGGACCAAGGAGTCTGACATGTGTGCGAGTCAACGGGTGAGTAAACCCGTAAGGCGCAAGGA | 44 (0.000832%) | | | TCGGTAGGAGCGACGGGCGGTGTGTACAAAGGGCAGGGACGTAGTCAACGCGAGCTGATGACTCGCGCTTACTAGGAATTCCTCGTTGAAGACCAACAAT | 31 (0.000586%) | | | TCGGTCCTCCGGATTTTCAAGGGCCGCCGGGGGCGCACCGGACACCACGCGACGTGCGGTGCTCTTCCAGCCGCTGGACCCTACCTCCGGCTGAGCCGTT | 10 (0.000189%) | | | TCGTACTGAAAATCAGAATCAAACGAGCTTTTACCCTTTTGTTCCACACGAGATTTCTGTTCTCGTTGAGCTCATCTTAGGACACCTGCGTTATCTTTTA | 3 (0.000057%) | | | TCGTAGGCTCCATGCTCGCGCATCGAACTACCTACCACCTATCCTTCTCAGTTAATTCACGGGCGATGTTACGCTCGATGATGAGTTCCGGGGCCTGTGT | 6 (0.000114%) | | | TCGTATTCGTACTGAAAATCAGAATCAAACGAGCTTTTACCCTTTTGTTCCACACGAGATTTCTGTTCTCGTTGAGCTCATCTTAGGACACCTGCGTTAT | 9 (0.000170%) | | | TCGTCCCTCACCATCCTTTGCTGATGCGGGACGGAAGCTGGTCTCCCGTGTGTTACCGCACGCGGTTGGCCTAAATCCGAGCCAAGGACGCCTGGAGCGT | 3 (0.000057%) | | | TCGTTCATCACCCTTGGCCGGCTTTCGAACAGCCGGACTCCCATCAAAAGATGGTTGCCAAGAACATCTTCGTTACGGTTTGCTAATTCTCGGAATAACA | 4 (0.000076%) | | | TCTCGCGCTTGTACGGCTTTGGCTCGGATTCGTCCGTCTTCTTTCTTCTTAGCCGAGTACTTCGGTAGATTAGTTGGAACGATTGATGATTTTGAGTTAA | 10 (0.000189%) | | | TCTGAGAAGGGTTCGAGTGTGAGCATGCCTGTCGGGACCCGAAAGATGGTGAACTATGCCTGAGCGGGGTAAAGCCAGAGGAAACTCTGGTGGAAGCCCG | 4 (0.000076%) | | | TCTGCCCTATCAACTTTCGATGGTAGGATAGTGGCCTACCATGGTGGTAACGGGTGACGGAGAATTAGGGTTCGATTCCGGAGAGGGAGCCTGAGAAACG | 8 (0.000151%) | | | TCTGGTGCCAGCAGCCGCGGTAATTCCAGCTCCAATAGCGTATATTTAAGTTGTTGCAGTTAAAAAGCTCGTAGTTGAACCTTGGGATGGGTCGGCCGGT | 17 (0.000322%) | | | TCTGTGCTGGCGACGCATCATTCAAATTTCTGCCCTATCAACTTTCGATGGTAGGATAGTGGCCTACCATGGTGGTAACGGGTGACGGAGAATTAGGGTT | 21 (0.000397%) | | | TCTTAAAGGCGTAAGAATTGTATCCTTGTTAGAAGACACA | 415 (0.003140%) | | | TCTTATACTCAATCATACACATGACATCAAGTCATATTCGACTCCAAAACACTAACCAACCTTCTTCTTGCTTCTCAAAGCTTTCATGGTGTAGCCAAAG | 208 (0.003935%) | | | TCTTCAACGAGGAATTCCTAGTAAGCGCGAGTCATCAGCTCGCGTTGACTACGTCCCTGCCCTTTGTACACACCGCCCGTCGCTCCTACCGATTGAATGA | 19 (0.000359%) | | | TGAACACAAACGTTCAATATGACAAACCCATGCCAAGTAAAGAGAAAATGAAAACTGGTGATTGTTGCGGAAATCGTCCAGGATTCCTCGACCAGGACTT | 20 (0.000378%) | | | TGAATCATCAGAGCAACGGGCAGAGCCCGCGTCGACCTTTTATCTAATAAATGCGTCCCTTCCATAAGTCGGGGTTTGTTGCACGTATTAGCTCTAGAAT | 8 (0.000151%) | | | TGACATCAAGTCATATTCGACTCCAAAACACTAACCAACCTTCTTCTTGCTTCTCAAAGCTTTCATGGTGTAGCCAAAGTCCATATGAGTCTTTGGCTTT | 38 (0.000719%) | | | TGACGGCAACGTTAGGGAGTCCGGAGACGTCGGCGGGGGCCTCGGGAAGAGTTATCTTTTCTGTTTAACAGCCTGCCCACCCTGGAAACGGCTCAGCCGG | 15 (0.000284%) | | | TGACTCGCACACATGTCAGACTCCTTGGTCCGTGTTTCAAGACGGGTCGAATGGGGAGCCCACAGGCCGACGCCCGGAGCACGCTGATGCCGAGGCACGC | 6 (0.000114%) | | | TGAGAAGGGTTCGAGTGTGAGCATGCCTGTCGGGACCCGAAAGATGGTGAACTATGCCTGAGCGGGGTAAAGCCAGAGGAAACTCTGGTGGAAGCCCGCA | 8 (0.000151%) | | | TGAGCGAGGTGTGAGTGTCGCCCATGGGCATCGACACCTTGCGGCTAGGAACTGGAACGAGACGGGTGGCAAAGATTTCGAGTAGCACTTCATACTACCG | 4 (0.000076%) | | | TGAGCGGCACTCGGTCCTCCGGATTTTCAAGGGCCGCCGGGGGCGCACCGGACACCACGCGACGTGCGGTGCTCTTCCAGCCGCTGGACCCTACCTCCGG | 3 (0.000057%) | | | TGAGTATAAGAACTTAAACCGCAACCGCATCTTATAAGCCTAAGTAGTGTTTCCTTGTTAGAAGACACAAAGCCAAAGACTCATATGGACTTTGGCTACA | 21 (0.000397%) | | | TGAGTGTCGCCCATGGGCATCGACACCTTGCGGCTAGGAACTGGAACGAGACGGGTGGCAAAGATTTCGAGTAGCACTTCATACTACCGTGGGTTTTTTA | 5 (0.000095%) | | | TGATGTTATCCCATGCTAATGTATCCAGAGCGTAGGCTTGCTTTGAGCACTCTAATTTCTTCAAAGTAACAGCGCCGGAGGCACGACCCGGCCAATTAAG | 22 (0.000416%) | | | TGATTAACAGGGACAGTCGGGGGCATTCGTATTTCATAGTCAGAGGTGAAATTCTTGGATTTATGAAAGACGAACAACTGCGAAAGCATTTGCCAAGGAT | 109 (0.002062%) | | | TGATTGAGTATAAGAACTTAAACCGCAACCGCATCTTATAAGCCTAAGTAGTGTTTCCTTGTTAGAAGACACAAAGCCAAAGACTCATATGGACTTTGGC | 7 (0.000132%) | | | TGCAGCACGCGCCTAACGGCGTGCCTCGGCATCAGCGTGCTCCGGGCGTCGGCCTGTGGGCTCCCCATTCGACCCGTCTTGAAACACGGACCAAGGAGTC | 61 (0.001154%) | | | TGCCGACTTCCCTTGCCTACATTGTTCCATCGACCAGAGGCTGTTCACCTTGGAGACCTGATGCGGTTATGAGTACGACCGGGCGTGAGCGGCACTCGGT | 2363 (0.044704%) | | | TGCCTACATTGTTCCATCGACCAGAGGCTGTTCACCTTGGAGACCTGATGCGGTTATGAGTACGACCGGGCGTGAGCGGCACTCGGTCCTCCGGATTTTC | 8 (0.000151%) | | | TGCGCCTTACGGGTTTACTCACCCGTTGACTCGCACACATGTCAGACTCCTTGGTCCGTGTTTCAAGACGGGTCGAATGGGGAGCCCACAGGCCGACGCC | 15 (0.000284%) | | | TGCGGTTTAAGTTCTTATACTCAATCATACACATGACATCAAGTCATATTCGACTCCAAAACACTAACCAACCTTCTTCTTGCTTCTCAAAGCTTTCATG | 405 (0.007662%) | | | TGCGGTTTAAGTTGTTATACTCAATCATACACATGACAACAAGTCATATTCGACTCCAAAACACTAACCAACCTTCTTCTTGCTTCTCAAAGCTTTCATG | 110 (0.002081%) | | | TGCTAACTAGCTACGTGGAGGCATCCCTTCACGGCCGGCTTCTTAGAGGGACTATGGCCGTTTAGGCCAAGGAAGTTTGAGGCAATAACAGGTCTGTGAT | 19 (0.000359%) | | | TGCTAATGTATCCAGAGCGTAGGCTTGCTTTGAGCACTCTAATTTCTTCAAAGTAACAGCGCCGGAGGCACGACCCGGCCAATTAAGACCAGGAGCGTAT | 10 (0.000189%) | | | TGCTCGCGCATCGAACTACCTACCACCTATCCTTCTCAGTTAATTCACGGGCGATGTTACGCTCGATGATGAGTTCCGGGGCCTGTGTTTCGTACCTAAT | 4 (0.000076%) | | | TGCTGATGCGGGACGGAAGCTGGTCTCCCGTGTGTTACCGCACGCGGTTGGCCTAAATCCGAGCCAAGGACGCCTGGAGCGTACCGACATGCGGTGGTGA | 5 (0.000095%) | | | TGCTGGCGACGCATCATTCAAATTTCTGCCCTATCAACTTTCGATGGTAGGATAGTGGCCTACCATGGTGGTAACGGGTGACGGAGAATTAGGGTTCGAT | 5 (0.000095%) | | | TGCTTGTCTCAAAGATTAAGCCATGCATGTGTAAGTATGAACGAATTCAGACTGTGAAACTGCGAATGGCTCATTAAATCAGTTATAGTTTGTTTGATGG | 8 (0.000151%) | | | TGCTTTGTTTTAATTAAACAGTCGGATTCCCCTTGTCCGTACCAGTTCTGAGCTGACTGTTCGACGCCCGGGGAAAGCTCCCGAGAGAGCCGTTCCCAGT | 3 (0.000057%) | | | TGGAAGTCGAAATCCGCTAAGGAGTGTGTAACAACTCACCTGCCGAATCAACTAGCCCCGAAAATGGATGGCGCTTAAGCGCGCGACCTATACCCGGCCG | 88 (0.001665%) | | | TGGAGGGCAAGTCTGGTGCCAGCAGCCGCGGTAATTCCAGCTCCAATAGCGTATATTTAAGTTGTTGCAGTTAAAAAGCTCGTAGTTGAACCTTGGGATG | 296 (0.005600%) | | | TGGCCTCTGTGCTGGCGACGCATCATTCAAATTTCTGCCCTATCAACTTTCGATGGTAGGATAGTGGCCTACCATGGTGGTAACGGGTGACGGAGAATTA | 23 (0.000435%) | | | TGGCGACGCATCATTCAAATTTCTGCCCTATCAACTTTCGATGGTAGGATAGTGGCCTACCATGGTGGTAACGGGTGACGGAGAATTAGGGTTCGATTCC | 22 (0.000416%) | | | TGGCGGTTGACGGCAACGTTAGGGAGTCCGGAGACGTCGGCGGGGGCCTCGGGAAGAGTTATCTTTTCTGTTTAACAGCCTGCCCACCCTGGAAACGGCT | 4 (0.000076%) | | | TGGCTGGGGCGGCACATCTGTTAAAAGATAACGCAGGTGTCCTAAGATGAGCTCAACGAGAACAGAAATCTCGTGTGGAACAAAAGGGTAAAAGCTCGTT | 3 (0.000057%) | | | TGGGCATCGACACCTTGCGGCTAGGAACTGGAACGAGACGGGTGGCAAAGATTTCGAGTAGCACTTCATACTACCGTGGGTTTTTTAAACCTTCCGAGTT | 23 (0.000435%) | | | TGGGGAGTTTGGCTGGGGCGGCACATCTGTTAAAAGATAACGCAGGTGTCCTAAGATGAGCTCAACGAGAACAGAAATCTCGTGTGGAACAAAAGGGTAA | 5 (0.000095%) | | | TGGGGCGGCACATCTGTTAAAAGATAACGCAGGTGTCCTAAGATGAGCTCAACGAGAACAGAAATCTCGTGTGGAACAAAAGGGTAAAAGCTCGTTTGAT | 17 (0.000322%) | | | TGGGTGGTTCGCCGCCCGCGACGTCGCGAGAAGTCCACTAAACCTTATCATTTAGAGGAAGGAGAAGTCGTAACAAGGTTTCCGTAGGTGAACCTGCGGA | 10 (0.000189%) | | | TGGGTGTCACAAATCGTCGTCCCTCACCATCCTTTGCTGATGCGGGACGGAAGCTGGTCTCCCGTGTGTTACCGCACGCGGTTGGCCTAAATCCGAGCCA | 3 (0.000057%) | | | TGGTCATGGAAGTCGAAATCCGCTAAGGAGTGTGTAACAACTCACCTGCCGAATCAACTAGCCCCGAAAATGGATGGCGCTTAAGCGCGCGACCTATACC | 23 (0.000435%) | | | TGGTCGATTAAGACAGCAGGACGGTGGTCATGGAAGTCGAAATCCGCTAAGGAGTGTGTAACAACTCACCTGCCGAATCAACTAGCCCCGAAAATGGATG | 9 (0.000170%) | | | TGGTCTTCAACGAGGAATTCCTAGTAAGCGCGAGTCATCAGCTCGCGTTGACTACGTCCCTGCCCTTTGTACACACCGCCCGTCGCTCCTACCGATTGAA | 37 (0.000700%) | | | TGGTGATATGAACACAAACGTTCAATATGACAAACCCATGCCAAGTAAAGAGAAAATGAAAACTGGTGATTGTTGCGGAAATCGTCCAGGATTCCTCGAC | 3 (0.000057%) | | | TGGTGCCAGCAGCCGCGGTAATTCCAGCTCCAATAGCGTATATTTAAGTTGTTGCAGTTAAAAAGCTCGTAGTTGAACCTTGGGATGGGTCGGCCGGTCC | 5 (0.000095%) | | | TGGTTCGCCGCCCGCGACGTCGCGAGAAGTCCACTAAACCTTATCATTTAGAGGAAGGAGAAGTCGTAACAAGGTTTCCGTAGGTGAACCTGCGGAAGGA | 6 (0.000114%) | | | TGTAAAGTTGGGAATTCGTTAAGGAGCTGTTGCTTTGTTAGTGTAGAAACACTTGTGTAGAATTGGGGATTGTTTTTTTTGGAGTGATTTAGGGGAGGGT | 15 (0.000284%) | | | TGTACGGCTTTGGCTCGGATTCGTCCGTCTTCTTTCTTCTTAGCCGAGTACTTCGGTAGATTAGTTGGAACGATTGATGATTTTGAGTTAATTGAACGTT | 3 (0.000057%) | | | TGTAGGGAAAGAGTGTTCAGAGCCGTGTAGATCTCGGTGG | 7 (0.000053%) | | | TGTATCCAGAGCGTAGGCTTGCTTTGAGCACTCTAATTTCTTCAAAGTAACAGCGCCGGAGGCACGACCCGGCCAATTAAGACCAGGAGCGTATCGCCGA | 27 (0.000511%) | | | TGTATCCTTGTTAGAAGACACAAAGCCAAAGACTCATATGGACTTTGGCTACACCATGAAAGCTTTGAGAAGCAAGAAGAAGGTTGGTTAGTGTTTTGGA | 680 (0.012864%) | | | TGTCAGGTGGGGAGTTTGGCTGGGGCGGCACATCTGTTAAAAGATAACGCAGGTGTCCTAAGATGAGCTCAACGAGAACAGAAATCTCGTGTGGAACAAA | 49 (0.000927%) | | | TGTCCCGAAGGTATCTCGCGCTTGTACGGCTTTGGCTCGGATTCGTCCGTCTTCTTTCTTCTTAGCCGAGTACTTCGGTAGATTAGTTGGAACGATTGAT | 4 (0.000076%) | | | TGTCCCGAGTGTGAGCGAGGTGTGAGTGTCGCCCATGGGCATCGACACCTTGCGGCTAGGAACTGGAACGAGACGGGTGGCAAAGATTTCGAGTAGCACT | 3 (0.000057%) | | | TGTCGCCCATGGGCATCGACACCTTGCGGCTAGGAACTGGAACGAGACGGGTGGCAAAGATTTCGAGTAGCACTTCATACTACCGTGGGTTTTTTAAACC | 7 (0.000132%) | | | TGTCTCAAAGATTAAGCCATGCATGTGTAAGTATGAACGAATTCAGACTGTGAAACTGCGAATGGCTCATTAAATCAGTTATAGTTTGTTTGATGGTAAC | 3 (0.000057%) | | | TGTGAGCGAGGTGTGAGTGTCGCCCATGGGCATCGACACCTTGCGGCTAGGAACTGGAACGAGACGGGTGGCAAAGATTTCGAGTAGCACTTCATACTAC | 5 (0.000095%) | | | TGTGAGTGTCGCCCATGGGCATCGACACCTTGCGGCTAGGAACTGGAACGAGACGGGTAGCAAAGATTTCGAGTAGCACTTCATACTACCGTGGGTTTTT | 7 (0.000132%) | | | TGTGAGTGTCGCCCATGGGCATCGACACCTTGCGGCTAGGAACTGGAACGAGACGGGTGGCAAAGATTTCGAGTAGCACTTCATACTACCGTGGGTTTTT | 6 (0.000114%) | | | TGTGCTGGCGACGCATCATTCAAATTTCTGCCCTATCAACTTTCGATGGTAGGATAGTGGCCTACCATGGTGGTAACGGGTGACGGAGAATTAGGGTTCG | 10 (0.000189%) | | | TGTTAGAAGACACAAAGCCAAAGACTCATATGGACTTTGGCTACACCATGAAAGCTTTGAGAAGCAAGAAGAAGGTTGGTTAGTGTTTTGGAGTCGAATA | 29 (0.000549%) | | | TGTTATACTCAATCATACACATGACAACAAGTCATATTCGACTCCAAAACACTAACCAACCTTCTTCTTGCTTCTCAAAGCTTTCATGGTGTAGCCAAAG | 6 (0.000114%) | | | TGTTATCCCATGCTAATGTATCCAGAGCGTAGGCTTGCTTTGAGCACTCTAATTTCTTCAAAGTAACAGCGCCGGAGGCACGACCCGGCCAATTAAGACC | 10 (0.000189%) | | | TGTTCCATCGACCAGAGGCTGTTCACCTTGGAGACCTGATGCGGTTATGAGTACGACCGGGCGTGAGCGGCACTCGGTCCTCCGGATTTTCAAGGGCCGC | 5 (0.000095%) | | | TGTTGCTTTGTTAGTGTAGAAACACTTGTGTAGAATTGGGGATTGTTTTTTTTGGAGTGATTTAGGGGAGGGTCGAATCTTAGCGACAAAGGGCTGAATC | 16 (0.000303%) | | | TGTTGGTCGATTAAGACAGCAGGACGGTGGTCATGGAAGTCGAAATCCGCTAAGGAGTGTGTAACAACTCACCTGCCGAATCAACTAGCCCCGAAAATGG | 13 (0.000246%) | | | TGTTGGTCTTCAACGAGGAATTCCTAGTAAGCGCGAGTCATCAGCTCGCGTTGACTACGTCCCTGCCCTTTGTACACACCGCCCGTCGCTCCTACCGATT | 193 (0.003651%) | | | TGTTTTGGAGTCGAATATGACTTGATGTCATGTGTATGAT | 272 (0.002058%) | | | TTAAAAGATAACGCAGGTGTCCTAAGATGAGCTCAACGAGAACAGAAATCTCGTGTGGAACAAAAGGGTAAAAGCTCGTTTGATTCTGATTTTCAGTACG | 20 (0.000378%) | | | TTAAAGGCGTAAGAATTGTATCCTTGTTAGAAGACACAAA | 465 (0.003519%) | | | TTAACAGGGACAGTCGGGGGCATTCGTATTTCATAGTCAGAGGTGAAATTCTTGGATTTATGAAAGACGAACAACTGCGAAAGCATTTGCCAAGGATGTT | 189 (0.003576%) | | | TTAAGACAGCAGGACGGTGGTCATGGAAGTCGAAATCCGCTAAGGAGTGTGTAACAACTCACCTGCCGAATCAACTAGCCCCGAAAATGGATGGCGCTTA | 78 (0.001476%) | | | TTAAGGAGCTGTTGCTTTGTTAGTGTAGAAACACTTGTGTAGAATTGGGGATTGTTTTTTTTGGAGTGATTTAGGGGAGGGTCGAATCTTAGCGACAAAG | 18 (0.000341%) | | | TTAAGTTCTTATACTCAATCATACACATGACATCAAGTCATATTCGACTCCAAAACACTAACCAACCTTCTTCTTGCTTCTCAAAGCTTTCATGGTGTAG | 176 (0.003330%) | | | TTAAGTTGTTATACTCAATCATACACATGACAACAAGTCATATTCGACTCCAAAACACTAACCAACCTTCTTCTTGCTTCTCAAAGCTTTCATGGTGTAG | 61 (0.001154%) | | | TTACATTTTATCGGTCGCTCTTGTCCGGAAGCTGTAGATGACCCAAAGTCCATATAGCGACCCCAGGTCAGGCGGGATTACCCGCTGAGTTTAAGCATAT | 12 (0.000227%) | | | TTACGGGTTTACTCACCCGTTGACTCGCACACATGTCAGACTCCTTGGTCCGTGTTTCAAGACGGGTCGAATGGGGAGCCCACAGGCCGACGCCCGGAGC | 30 (0.000568%) | | | TTACTCACCCGTTGACTCGCACACATGTCAGACTCCTTGGTCCGTGTTTCAAGACGGGTCGAATGGGGAGCCCACAGGCCGACGCCCGGAGCACGCTGAT | 11 (0.000208%) | | | TTAGAAGACACAAAGCCAAAGACTCATATGGACTTTGGCTACACCATGAAAGCTTTGAGAAGCAAGAAGAAGGTTGGTTAGTGTTTTGGAGTCGAATATG | 92 (0.001740%) | | | TTAGGTACGAAACACAGGCCCCGGAACTCATCATCGAGCGTAACATCGCCCGTGAATTAACTGAGAAGGATAGGTGGTAGGTAGTTCGATGCGCGAGCAT | 9 (0.000170%) | | | TTAGTGTTTTGGAGTCGAATATGACTTGATGTCATGTGTA | 357 (0.002702%) | | | TTATACTCAATCATACACATGACAACAAGTCATATTCGACTCCAAAACACTAACCAACCTTCTTCTTGCTTCTCAAAGCTTTCATGGTGTAGCCAAAGTC | 318 (0.006016%) | | | TTATACTCAATCATACACATGACATCAAGTCATATTCGACTCCAAAACACTAACCAACCTTCTTCTTGCTTCTCAAAGCTTTCATGGTGTAGCCAAAGTC | 684 (0.012940%) | | | TTATCATGAATCATCAGAGCAACGGGCAGAGCCCGCGTCGACCTTTTATCTAATAAATGCGTCCCTTCCATAAGTCGGGGTTTGTTGCACGTATTAGCTC | 67 (0.001268%) | | | TTATCCCATGCTAATGTATCCAGAGCGTAGGCTTGCTTTGAGCACTCTAATTTCTTCAAAGTAACAGCGCCGGAGGCACGACCCGGCCAATTAAGACCAG | 5 (0.000095%) | | | TTATCGGTCGCTCTTGTCCGGAAGCTGTAGATGACCCAAAGTCCATATAGCGACCCCAGGTCAGGCGGGATTACCCGCTGAGTTTAAGCATATCAATAAG | 8 (0.000151%) | | | TTCAAATTAGGTACGAAACACAGGCCCCGGAACTCATCATCGAGCGTAACATCGCCCGTGAATTAACTGAGAAGGATAGGTGGTAGGTAGTTCGATGCGC | 38 (0.000719%) | | | TTCAAATTTCTGCCCTATCAACTTTCGATGGTAGGATAGTGGCCTACCATGGTGGTAACGGGTGACGGAGAATTAGGGTTCGATTCCGGAGAGGGAGCCT | 26 (0.000492%) | | | TTCAACGAGGAATTCCTAGTAAGCGCGAGTCATCAGCTCGCGTTGACTACGTCCCTGCCCTTTGTACACACCGCCCGTCGCTCCTACCGATTGAATGATC | 5 (0.000095%) | | | TTCAAGGGCCGCCGGGGGCGCACCGGACACCACGCGACGTGCGGTGCTCTTCCAGCCGCTGGACCCTACCTCCGGCTGAGCCGTTTCCAGGGTGGGCAGG | 14 (0.000265%) | | | TTCAATATGACAAACCCATGCCAAGTAAAGAGAAAATGAAAACTGGTGATTGTTGCGGAAATCGTCCAGGATTCCTCGACCAGGACTTGAAATCGTCGAG | 3 (0.000057%) | | | TTCAATCGGTAGGAGCGACGGGCGGTGTGTACAAAGGGCAGGGACGTAGTCAACGCGAGCTGATGACTCGCGCTTACTAGGAATTCCTCGTTGAAGACCA | 51 (0.000965%) | | | TTCAATGTCTGTTGGTGCCAAGAGGGAAAAGGGCTATTAAGCTATATAGGGGGGTGGGTGTTGAGGGAGTCTGGGCAGTCCGTGGGGAACCCCCTTTTTC | 459 (0.008683%) | | | TTCACATTTCGTTCATCACCCTTGGCCGGCTATCGAACAGCCGGACTCCCATCAAAAGATGGTTGCCAAGAACATCTTCGTTACGGTTTGCTAATTCTCG | 18 (0.000341%) | | | TTCACATTTCGTTCATCACCCTTGGCCGGCTTTCGAACAGCCGGACTCCCATCAAAAGATGGTTGCCAAGAACATCTTCGTTACGGTTTGCTAATTCTCG | 14 (0.000265%) | | | TTCACGGTTCGTATTCGTACTGAAAATCAGAATCAAACGAGCTTTTACCCTTTTGTTCCACACGAGATTTCTGTTCTCGTTGAGCTCATCTTAGGACACC | 17 (0.000322%) | | | TTCCAACGAAGCACGCCCATCCAACCTAGGCGAGACAAGGGTTCACATTTCGTTCATCACCCTTGGCCGGCTATCGAACAGCCGGACTCCCATCAAAAGA | 106 (0.002005%) | | | TTCCAGCTCCAATAGCGTATATTTAAGTTGTTGCAGTTAAAAAGCTCGTAGTTGAACCTTGGGATGGGTCGGCCGGTCCGCCTTTGGTGTGCATTGGTCG | 5 (0.000095%) | | | TTCCCTTGCCTACATTGTTCCATCGACCAGAGGCTGTTCACCTTGGAGACCTGATGCGGTTATGAGTACGACCGGGCGTGAGCGGCACTCGGTCCTCCGG | 6 (0.000114%) | | | TTCCGGAACCGGGACGTGGCGGTTGACGGCAACGTTAGGGAGTCCGGAGACGTCGGCGGGGGCCTCGGGAAGAGTTATCTTTTCTGTTTAACAGCCTGCC | 2316 (0.043815%) | | | TTCCTTGCGCCTTACGGGTTTACTCACCCGTTGACTCGCACACATGTCAGACTCCTTGGTCCGTGTTTCAAGACGGGTCGAATGGGGAGCCCACAGGCCG | 11 (0.000208%) | | | TTCGACTCCAAAACACTAACCAACCTTCTTCTTGCTTCTCAAAGCTTTCATGGTGTAGCCAAAGTCCATATGAGTCTTTGGCTTTGTGTCTTCTAACAAG | 120 (0.002270%) | | | TTCGACTCCAAAACACTAACCAACCTTCTTCTTGCTTCTCAAAGCTTTCATGGTGTAGCCAAAGTCCATATGAGTCTTTGGCTTTGTGTCTTTTAACAAG | 74 (0.001400%) | | | TTCGAGTGTGAGCATGCCTGTCGGGACCCGAAAGATGGTGAACTATGCCTGAGCGGGGTAAAGCCAGAGGAAACTCTGGTGGAAGCCCGCAGCGATACTG | 5 (0.000095%) | | | TTCGATTAGTCTTTCGCCCCTATACCCAAGTCAGACGAACGATTTGCACGTCAGTATCGCTGCGGGCTTCCACCAGAGTTTCCTCTGGCTTTACCCCGCT | 10 (0.000189%) | | | TTCGCCGCCCGCGACGTCGCGAGAAGTCCACTAAACCTTATCATTTAGAGGAAGGAGAAGTCGTAACAAGGTTTCCGTAGGTGAACCTGCGGAAGGATCG | 5 (0.000095%) | | | TTCGTACTGAAAATCAGAATCAAACGAGCTTTTACCCTTTTGTTCCACACGAGATTTCTGTTCTCGTTGAGCTCATCTTAGGACACCTGCGTTATCTTTT | 20 (0.000378%) | | | TTCGTAGGCTCCATGCTCGCGCATCGAACTACCTACCACCTATCCTTCTCAGTTAATTCACGGGCGATGTTACGCTCGATGATGAGTTCCGGGGCCTGTG | 11 (0.000208%) | | | TTCGTATTCGTACTGAAAATCAGAATCAAACGAGCTTTTACCCTTTTGTTCCACACGAGATTTCTGTTCTCGTTGAGCTCATCTTAGGACACCTGCGTTA | 12 (0.000227%) | | | TTCGTATTTCATAGTCAGAGGTGAAATTCTTGGATTTATGAAAGACGAACAACTGCGAAAGCATTTGCCAAGGATGTTTTCATTAATCAAGAACGAAAGT | 24 (0.000454%) | | | TTCGTTCATCACCCTTGGCCGGCTATCGAACAGCCGGACTCCCATCAAAAGATGGTTGCCAAGAACATCTTCGTTACGGTTTGCTAATTCTCGGAATAAC | 18 (0.000341%) | | | TTCGTTCATCACCCTTGGCCGGCTTTCGAACAGCCGGACTCCCATCAAAAGATGGTTGCCAAGAACATCTTCGTTACGGTTTGCTAATTCTCGGAATAAC | 12 (0.000227%) | | | TTCTGAGAAGGGTTCGAGTGTGAGCATGCCTGTCGGGACCCGAAAGATGGTGAACTATGCCTGAGCGGGGTAAAGCCAGAGGAAACTCTGGTGGAAGCCC | 9 (0.000170%) | | | TTCTGCCCTATCAACTTTCGATGGTAGGATAGTGGCCTACCATGGTGGTAACGGGTGACGGAGAATTAGGGTTCGATTCCGGAGAGGGAGCCTGAGAAAC | 11 (0.000208%) | | | TTCTTATACTCAATCATACACATGACATCAAGTCATATTCGACTCCAAAACACTAACCAACCTTCTTCTTGCTTCTCAAAGCTTTCATGGTGTAGCCAAA | 99 (0.001873%) | | | TTGAAATCGTCGACCAGGTCCGAGACTTCATCGACCGGGTCCGAGGATTCGTCGACCAGGACGGCCGGATGTCCGAGAAAAAAAAATGTTGCCGAATAAC | 47 (0.000889%) | | | TTGACGGCAACGTTAGGGAGTCCGGAGACGTCGGCGGGGGCCTCGGGAAGAGTTATCTTTTCTGTTTAACAGCCTGCCCACCCTGGAAACGGCTCAGCCG | 6 (0.000114%) | | | TTGACTCGCACACATGTCAGACTCCTTGGTCCGTGTTTCAAGACGGGTCGAATGGGGAGCCCACAGGCCGACGCCCGGAGCACGCTGATGCCGAGGCACG | 6 (0.000114%) | | | TTGAGTATAAGAACTTAAACCGCAACCGCATCTTATAAGCCTAAGTAGTGTTTCCTTGTTAGAAGACACAAAGCCAAAGACTCATATGGACTTTGGCTAC | 7 (0.000132%) | | | TTGCCTACATTGTTCCATCGACCAGAGGCTGTTCACCTTGGAGACCTGATGCGGTTATGAGTACGACCGGGCGTGAGCGGCACTCGGTCCTCCGGATTTT | 6 (0.000114%) | | | TTGCGCCTTACGGGTTTACTCACCCGTTGACTCGCACACATGTCAGACTCCTTGGTCCGTGTTTCAAGACGGGTCGAATGGGGAGCCCACAGGCCGACGC | 22 (0.000416%) | | | TTGCGGTTTAAGTTCTTATACTCAATCATACACATGACATCAAGTCATATTCGACTCCAAAACACTAACCAACCTTCTTCTTGCTTCTCAAAGCTTTCAT | 449 (0.008494%) | | | TTGCGGTTTAAGTTGTTATACTCAATCATACACATGACAACAAGTCATATTCGACTCCAAAACACTAACCAACCTTCTTCTTGCTTCTCAAAGCTTTCAT | 79 (0.001495%) | | | TTGCTGATGCGGGACGGAAGCTGGTCTCCCGTGTGTTACCGCACGCGGTTGGCCTAAATCCGAGCCAAGGACGCCTGGAGCGTACCGACATGCGGTGGTG | 3 (0.000057%) | | | TTGGAGACCTGATGCGGTTATGAGTACGACCGGGCGTGAGCGGCACTCGGTCCTCCGGATTTTCAAGGGCCGCCGGGGGCGCACCGGACACCACGCGACG | 25 (0.000473%) | | | TTGGAGGGCAAGTCTGGTGCCAGCAGCCGCGGTAATTCCAGCTCCAATAGCGTATATTTAAGTTGTTGCAGTTAAAAAGCTCGTAGTTGAACCTTGGGAT | 338 (0.006394%) | | | TTGGAGTCGAATATGACTTGATGTCATGTGTATGATTGAG | 284 (0.002149%) | | | TTGGCTCGGATTCGTCCGTCTTCTTTCTTCTTAGCCGAGTACTTCGGTAGATTAGTTGGAACGATTGATGATTTTGAGTTAATTGAACGTTCGGCGTATG | 5 (0.000095%) | | | TTGGCTGGGGCGGCACATCTGTTAAAAGATAACGCAGGTGTCCTAAGATGAGCTCAACGAGAACAGAAATCTCGTGTGGAACAAAAGGGTAAAAGCTCGT | 25 (0.000473%) | | | TTGGTCGATTAAGACAGCAGGACGGTGGTCATGGAAGTCGAAATCCGCTAAGGAGTGTGTAACAACTCACCTGCCGAATCAACTAGCCCCGAAAATGGAT | 6 (0.000114%) | | | TTGGTCTTCAACGAGGAATTCCTAGTAAGCGCGAGTCATCAGCTCGCGTTGACTACGTCCCTGCCCTTTGTACACACCGCCCGTCGCTCCTACCGATTGA | 104 (0.001967%) | | | TTGGTGATATGAACACAAACGTTCAATATGACAAACCCATGCCAAGTAAAGAGAAAATGAAAACTGGTGATTGTTGCGGAAATCGTCCAGGATTCCTCGA | 3 (0.000057%) | | | TTGTACGGCTTTGGCTCGGATTCGTCCGTCTTCTTTCTTCTTAGCCGAGTACTTCGGTAGATTAGTTGGAACGATTGATGATTTTGAGTTAATTGAACGT | 10 (0.000189%) | | | TTGTATCCTTGTTAGAAGACACAAAGCCAAAGACTCATATGGACTTTGGCTACACCATGAAAGCTTTGAGAAGCAAGAAGAAGGTTGGTTAGTGTTTTGG | 14 (0.000265%) | | | TTGTCAGGTGGGGAGTTTGGCTGGGGCGGCACATCTGTTAAAAGATAACGCAGGTGTCCTAAGATGAGCTCAACGAGAACAGAAATCTCGTGTGGAACAA | 11 (0.000208%) | | | TTGTCTCAAAGATTAAGCCATGCATGTGTAAGTATGAACGAATTCAGACTGTGAAACTGCGAATGGCTCATTAAATCAGTTATAGTTTGTTTGATGGTAA | 10 (0.000189%) | | | TTGTTAGAAGACACAAAGCCAAAGACTCATATGGACTTTGGCTACACCATGAAAGCTTTGAGAAGCAAGAAGAAGGTTGGTTAGTGTTTTGGAGTCGAAT | 56 (0.001059%) | | | TTGTTATACTCAATCATACACATGACAACAAGTCATATTCGACTCCAAAACACTAACCAACCTTCTTCTTGCTTCTCAAAGCTTTCATGGTGTAGCCAAA | 45 (0.000851%) | | | TTGTTGGTCTTCAACGAGGAATTCCTAGTAAGCGCGAGTCATCAGCTCGCGTTGACTACGTCCCTGCCCTTTGTACACACCGCCCGTCGCTCCTACCGAT | 180 (0.003405%) | | | TTTAAGTTCTTATACTCAATCATACACATGACATCAAGTCATATTCGACTCCAAAACACTAACCAACCTTCTTCTTGCTTCTCAAAGCTTTCATGGTGTA | 113 (0.002138%) | | | TTTAAGTTGTTATACTCAATCATACACATGACAACAAGTCATATTCGACTCCAAAACACTAACCAACCTTCTTCTTGCTTCTCAAAGCTTTCATGGTGTA | 53 (0.001003%) | | | TTTAATTAAACAGTCGGATTCCCCTTGTCCGTACCAGTTCTGAGCTGACTGTTCGACGCCCGGGGAAAGCTCCCGAGAGAGCCGTTCCCAGTCCGTCCCC | 17 (0.000322%) | | | TTTACTCACCCGTTGACTCGCACACATGTCAGACTCCTTGGTCCGTGTTTCAAGACGGGTCGAATGGGGAGCCCACAGGCCGACGCCCGGAGCACGCTGA | 13 (0.000246%) | | | TTTATCGGTCGCTCTTGTCCGGAAGCTGTAGATGACCCAAAGTCCATATAGCGACCCCAGGTCAGGCGGGATTACCCGCTGAGTTTAAGCATATCAATAA | 15 (0.000284%) | | | TTTCAAGGGCCGCCGGGGGCGCACCGGACACCACGCGACGTGCGGTGCTCTTCCAGCCGCTGGACCCTACCTCCGGCTGAGCCGTTTCCAGGGTGGGCAG | 29 (0.000549%) | | | TTTCACGGTTCGTATTCGTACTGAAAATCAGAATCAAACGAGCTTTTACCCTTTTGTTCCACACGAGATTTCTGTTCTCGTTGAGCTCATCTTAGGACAC | 54 (0.001022%) | | | TTTCAGCAGTTCTCGGACAAAAATTGCTGAGTGGCCGAGAAGAATGGGCGTGTCATGCGTGGGCTGACATGGATTCTTCGAGGCCTAGGGGTGGCGGTAT | 10 (0.000189%) | | | TTTCCAACGAAGCACGCCCATCCAACCTAGGCGAGACAAGGGTTCACATTTCGTTCATCACCCTTGGCCGGCTATCGAACAGCCGGACTCCCATCAAAAG | 1826 (0.034545%) | | | TTTCGTTCATCACCCTTGGCCGGCTATCGAACAGCCGGACTCCCATCAAAAGATGGTTGCCAAGAACATCTTCGTTACGGTTTGCTAATTCTCGGAATAA | 9 (0.000170%) | | | TTTCGTTCATCACCCTTGGCCGGCTTTCGAACAGCCGGACTCCCATCAAAAGATGGTTGCCAAGAACATCTTCGTTACGGTTTGCTAATTCTCGGAATAA | 9 (0.000170%) | | | TTTCTGCCCTATCAACTTTCGATGGTAGGATAGTGGCCTACCATGGTGGTAACGGGTGACGGAGAATTAGGGTTCGATTCCGGAGAGGGAGCCTGAGAAA | 14 (0.000265%) | | | TTTGCTGATGCGGGACGGAAGCTGGTCTCCCGTGTGTTACCGCACGCGGTTGGCCTAAATCCGAGCCAAGGACGCCTGGAGCGTACCGACATGCGGTGGT | 27 (0.000511%) | | | TTTGGAGTCGAATATGACTTGATGTCATGTGTATGATTGA | 368 (0.002785%) | | | TTTGGCTCGGATTCGTCCGTCTTCTTTCTTCTTAGCCGAGTACTTCGGTAGATTAGTTGGAACGATTGATGATTTTGAGTTAATTGAACGTTCGGCGTAT | 27 (0.000511%) | | | TTTGGCTGGGGCGGCACATCTGTTAAAAGATAACGCAGGTGTCCTAAGATGAGCTCAACGAGAACAGAAATCTCGTGTGGAACAAAAGGGTAAAAGCTCG | 20 (0.000378%) | | | TTTGGCTTTGTGTCTTCTAACAAGGAAACACTACTTAGGC | 6916 (0.052335%) | | | TTTGTTTTAATTAAACAGTCGGATTCCCCTTGTCCGTACCAGTTCTGAGCTGACTGTTCGACGCCCGGGGAAAGCTCCCGAGAGAGCCGTTCCCAGTCCG | 21 (0.000397%) | | | TTTTAATTAAACAGTCGGATTCCCCTTGTCCGTACCAGTTCTGAGCTGACTGTTCGACGCCCGGGGAAAGCTCCCGAGAGAGCCGTTCCCAGTCCGTCCC | 17 (0.000322%) | | | TTTTATCGGTCGCTCTTGTCCGGAAGCTGTAGATGACCCAAAGTCCATATAGCGACCCCAGGTCAGGCGGGATTACCCGCTGAGTTTAAGCATATCAATA | 3 (0.000057%) | | | TTTTCAAGGGCCGCCGGGGGCGCACCGGACACCACGCGACGTGCGGTGCTCTTCCAGCCGCTGGACCCTACCTCCGGCTGAGCCGTTTCCAGGGTGGGCA | 24 (0.000454%) | | | TTTTCAGCAGTTCTCGGACAAAAATTGCTGAGTGGCCGAGAAGAATGGGCGTGTCATGCGTGGGCTGACATGGATTCTTCGAGGCCTAGGGGTGGCGGTA | 11 (0.000208%) | | | TTTTGGAGTCGAATATGACTTGATGTCATGTGTATGATTG | 535 (0.004049%) | | | TTTTTCAGCAGTTCTCGGACAAAAATTGCTGAGTGGCCGAGAAGAATGGGCGTGTCATGCGTGGGCTGACATGGATTCTTCGAGGCCTAGGGGTGGCGGT | 31 (0.000586%) | | | TTTTTTTTTT | 54385 (0.102887%) | | After filtering After filtering: read1: quality Value of each position will be shown on mouse over. After filtering: read1: base contents Value of each position will be shown on mouse over. After filtering: read1: KMER counting Darker background means larger counts. The count will be shown on mouse over. | | | | | | | | | | | | | | | | | | | --- | --- | --- | --- | --- | --- | --- | --- | --- | --- | --- | --- | --- | --- | --- | --- | --- | | | AA | AT | AC | AG | TA | TT | TC | TG | CA | CT | CC | CG | GA | GT | GC | GG | | AAA | AAAAA | AAAAT | AAAAC | AAAAG | AAATA | AAATT | AAATC | AAATG | AAACA | AAACT | AAACC | AAACG | AAAGA | AAAGT | AAAGC | AAAGG | | AAT | AATAA | AATAT | AATAC | AATAG | AATTA | AATTT | AATTC | AATTG | AATCA | AATCT | AATCC | AATCG | AATGA | AATGT | AATGC | AATGG | | AAC | AACAA | AACAT | AACAC | AACAG | AACTA | AACTT | AACTC | AACTG | AACCA | AACCT | AACCC | AACCG | AACGA | AACGT | AACGC | AACGG | | AAG | AAGAA | AAGAT | AAGAC | AAGAG | AAGTA | AAGTT | AAGTC | AAGTG | AAGCA | AAGCT | AAGCC | AAGCG | AAGGA | AAGGT | AAGGC | AAGGG | | ATA | ATAAA | ATAAT | ATAAC | ATAAG | ATATA | ATATT | ATATC | ATATG | ATACA | ATACT | ATACC | ATACG | ATAGA | ATAGT | ATAGC | ATAGG | | ATT | ATTAA | ATTAT | ATTAC | ATTAG | ATTTA | ATTTT | ATTTC | ATTTG | ATTCA | ATTCT | ATTCC | ATTCG | ATTGA | ATTGT | ATTGC | ATTGG | | ATC | ATCAA | ATCAT | ATCAC | ATCAG | ATCTA | ATCTT | ATCTC | ATCTG | ATCCA | ATCCT | ATCCC | ATCCG | ATCGA | ATCGT | ATCGC | ATCGG | | ATG | ATGAA | ATGAT | ATGAC | ATGAG | ATGTA | ATGTT | ATGTC | ATGTG | ATGCA | ATGCT | ATGCC | ATGCG | ATGGA | ATGGT | ATGGC | ATGGG | | ACA | ACAAA | ACAAT | ACAAC | ACAAG | ACATA | ACATT | ACATC | ACATG | ACACA | ACACT | ACACC | ACACG | ACAGA | ACAGT | ACAGC | ACAGG | | ACT | ACTAA | ACTAT | ACTAC | ACTAG | ACTTA | ACTTT | ACTTC | ACTTG | ACTCA | ACTCT | ACTCC | ACTCG | ACTGA | ACTGT | ACTGC | ACTGG | | ACC | ACCAA | ACCAT | ACCAC | ACCAG | ACCTA | ACCTT | ACCTC | ACCTG | ACCCA | ACCCT | ACCCC | ACCCG | ACCGA | ACCGT | ACCGC | ACCGG | | ACG | ACGAA | ACGAT | ACGAC | ACGAG | ACGTA | ACGTT | ACGTC | ACGTG | ACGCA | ACGCT | ACGCC | ACGCG | ACGGA | ACGGT | ACGGC | ACGGG | | AGA | AGAAA | AGAAT | AGAAC | AGAAG | AGATA | AGATT | AGATC | AGATG | AGACA | AGACT | AGACC | AGACG | AGAGA | AGAGT | AGAGC | AGAGG | | AGT | AGTAA | AGTAT | AGTAC | AGTAG | AGTTA | AGTTT | AGTTC | AGTTG | AGTCA | AGTCT | AGTCC | AGTCG | AGTGA | AGTGT | AGTGC | AGTGG | | AGC | AGCAA | AGCAT | AGCAC | AGCAG | AGCTA | AGCTT | AGCTC | AGCTG | AGCCA | AGCCT | AGCCC | AGCCG | AGCGA | AGCGT | AGCGC | AGCGG | | AGG | AGGAA | AGGAT | AGGAC | AGGAG | AGGTA | AGGTT | AGGTC | AGGTG | AGGCA | AGGCT | AGGCC | AGGCG | AGGGA | AGGGT | AGGGC | AGGGG | | TAA | TAAAA | TAAAT | TAAAC | TAAAG | TAATA | TAATT | TAATC | TAATG | TAACA | TAACT | TAACC | TAACG | TAAGA | TAAGT | TAAGC | TAAGG | | TAT | TATAA | TATAT | TATAC | TATAG | TATTA | TATTT | TATTC | TATTG | TATCA | TATCT | TATCC | TATCG | TATGA | TATGT | TATGC | TATGG | | TAC | TACAA | TACAT | TACAC | TACAG | TACTA | TACTT | TACTC | TACTG | TACCA | TACCT | TACCC | TACCG | TACGA | TACGT | TACGC | TACGG | | TAG | TAGAA | TAGAT | TAGAC | TAGAG | TAGTA | TAGTT | TAGTC | TAGTG | TAGCA | TAGCT | TAGCC | TAGCG | TAGGA | TAGGT | TAGGC | TAGGG | | TTA | TTAAA | TTAAT | TTAAC | TTAAG | TTATA | TTATT | TTATC | TTATG | TTACA | TTACT | TTACC | TTACG | TTAGA | TTAGT | TTAGC | TTAGG | | TTT | TTTAA | TTTAT | TTTAC | TTTAG | TTTTA | TTTTT | TTTTC | TTTTG | TTTCA | TTTCT | TTTCC | TTTCG | TTTGA | TTTGT | TTTGC | TTTGG | | TTC | TTCAA | TTCAT | TTCAC | TTCAG | TTCTA | TTCTT | TTCTC | TTCTG | TTCCA | TTCCT | TTCCC | TTCCG | TTCGA | TTCGT | TTCGC | TTCGG | | TTG | TTGAA | TTGAT | TTGAC | TTGAG | TTGTA | TTGTT | TTGTC | TTGTG | TTGCA | TTGCT | TTGCC | TTGCG | TTGGA | TTGGT | TTGGC | TTGGG | | TCA | TCAAA | TCAAT | TCAAC | TCAAG | TCATA | TCATT | TCATC | TCATG | TCACA | TCACT | TCACC | TCACG | TCAGA | TCAGT | TCAGC | TCAGG | | TCT | TCTAA | TCTAT | TCTAC | TCTAG | TCTTA | TCTTT | TCTTC | TCTTG | TCTCA | TCTCT | TCTCC | TCTCG | TCTGA | TCTGT | TCTGC | TCTGG | | TCC | TCCAA | TCCAT | TCCAC | TCCAG | TCCTA | TCCTT | TCCTC | TCCTG | TCCCA | TCCCT | TCCCC | TCCCG | TCCGA | TCCGT | TCCGC | TCCGG | | TCG | TCGAA | TCGAT | TCGAC | TCGAG | TCGTA | TCGTT | TCGTC | TCGTG | TCGCA | TCGCT | TCGCC | TCGCG | TCGGA | TCGGT | TCGGC | TCGGG | | TGA | TGAAA | TGAAT | TGAAC | TGAAG | TGATA | TGATT | TGATC | TGATG | TGACA | TGACT | TGACC | TGACG | TGAGA | TGAGT | TGAGC | TGAGG | | TGT | TGTAA | TGTAT | TGTAC | TGTAG | TGTTA | TGTTT | TGTTC | TGTTG | TGTCA | TGTCT | TGTCC | TGTCG | TGTGA | TGTGT | TGTGC | TGTGG | | TGC | TGCAA | TGCAT | TGCAC | TGCAG | TGCTA | TGCTT | TGCTC | TGCTG | TGCCA | TGCCT | TGCCC | TGCCG | TGCGA | TGCGT | TGCGC | TGCGG | | TGG | TGGAA | TGGAT | TGGAC | TGGAG | TGGTA | TGGTT | TGGTC | TGGTG | TGGCA | TGGCT | TGGCC | TGGCG | TGGGA | TGGGT | TGGGC | TGGGG | | CAA | CAAAA | CAAAT | CAAAC | CAAAG | CAATA | CAATT | CAATC | CAATG | CAACA | CAACT | CAACC | CAACG | CAAGA | CAAGT | CAAGC | CAAGG | | CAT | CATAA | CATAT | CATAC | CATAG | CATTA | CATTT | CATTC | CATTG | CATCA | CATCT | CATCC | CATCG | CATGA | CATGT | CATGC | CATGG | | CAC | CACAA | CACAT | CACAC | CACAG | CACTA | CACTT | CACTC | CACTG | CACCA | CACCT | CACCC | CACCG | CACGA | CACGT | CACGC | CACGG | | CAG | CAGAA | CAGAT | CAGAC | CAGAG | CAGTA | CAGTT | CAGTC | CAGTG | CAGCA | CAGCT | CAGCC | CAGCG | CAGGA | CAGGT | CAGGC | CAGGG | | CTA | CTAAA | CTAAT | CTAAC | CTAAG | CTATA | CTATT | CTATC | CTATG | CTACA | CTACT | CTACC | CTACG | CTAGA | CTAGT | CTAGC | CTAGG | | CTT | CTTAA | CTTAT | CTTAC | CTTAG | CTTTA | CTTTT | CTTTC | CTTTG | CTTCA | CTTCT | CTTCC | CTTCG | CTTGA | CTTGT | CTTGC | CTTGG | | CTC | CTCAA | CTCAT | CTCAC | CTCAG | CTCTA | CTCTT | CTCTC | CTCTG | CTCCA | CTCCT | CTCCC | CTCCG | CTCGA | CTCGT | CTCGC | CTCGG | | CTG | CTGAA | CTGAT | CTGAC | CTGAG | CTGTA | CTGTT | CTGTC | CTGTG | CTGCA | CTGCT | CTGCC | CTGCG | CTGGA | CTGGT | CTGGC | CTGGG | | CCA | CCAAA | CCAAT | CCAAC | CCAAG | CCATA | CCATT | CCATC | CCATG | CCACA | CCACT | CCACC | CCACG | CCAGA | CCAGT | CCAGC | CCAGG | | CCT | CCTAA | CCTAT | CCTAC | CCTAG | CCTTA | CCTTT | CCTTC | CCTTG | CCTCA | CCTCT | CCTCC | CCTCG | CCTGA | CCTGT | CCTGC | CCTGG | | CCC | CCCAA | CCCAT | CCCAC | CCCAG | CCCTA | CCCTT | CCCTC | CCCTG | CCCCA | CCCCT | CCCCC | CCCCG | CCCGA | CCCGT | CCCGC | CCCGG | | CCG | CCGAA | CCGAT | CCGAC | CCGAG | CCGTA | CCGTT | CCGTC | CCGTG | CCGCA | CCGCT | CCGCC | CCGCG | CCGGA | CCGGT | CCGGC | CCGGG | | CGA | CGAAA | CGAAT | CGAAC | CGAAG | CGATA | CGATT | CGATC | CGATG | CGACA | CGACT | CGACC | CGACG | CGAGA | CGAGT | CGAGC | CGAGG | | CGT | CGTAA | CGTAT | CGTAC | CGTAG | CGTTA | CGTTT | CGTTC | CGTTG | CGTCA | CGTCT | CGTCC | CGTCG | CGTGA | CGTGT | CGTGC | CGTGG | | CGC | CGCAA | CGCAT | CGCAC | CGCAG | CGCTA | CGCTT | CGCTC | CGCTG | CGCCA | CGCCT | CGCCC | CGCCG | CGCGA | CGCGT | CGCGC | CGCGG | | CGG | CGGAA | CGGAT | CGGAC | CGGAG | CGGTA | CGGTT | CGGTC | CGGTG | CGGCA | CGGCT | CGGCC | CGGCG | CGGGA | CGGGT | CGGGC | CGGGG | | GAA | GAAAA | GAAAT | GAAAC | GAAAG | GAATA | GAATT | GAATC | GAATG | GAACA | GAACT | GAACC | GAACG | GAAGA | GAAGT | GAAGC | GAAGG | | GAT | GATAA | GATAT | GATAC | GATAG | GATTA | GATTT | GATTC | GATTG | GATCA | GATCT | GATCC | GATCG | GATGA | GATGT | GATGC | GATGG | | GAC | GACAA | GACAT | GACAC | GACAG | GACTA | GACTT | GACTC | GACTG | GACCA | GACCT | GACCC | GACCG | GACGA | GACGT | GACGC | GACGG | | GAG | GAGAA | GAGAT | GAGAC | GAGAG | GAGTA | GAGTT | GAGTC | GAGTG | GAGCA | GAGCT | GAGCC | GAGCG | GAGGA | GAGGT | GAGGC | GAGGG | | GTA | GTAAA | GTAAT | GTAAC | GTAAG | GTATA | GTATT | GTATC | GTATG | GTACA | GTACT | GTACC | GTACG | GTAGA | GTAGT | GTAGC | GTAGG | | GTT | GTTAA | GTTAT | GTTAC | GTTAG | GTTTA | GTTTT | GTTTC | GTTTG | GTTCA | GTTCT | GTTCC | GTTCG | GTTGA | GTTGT | GTTGC | GTTGG | | GTC | GTCAA | GTCAT | GTCAC | GTCAG | GTCTA | GTCTT | GTCTC | GTCTG | GTCCA | GTCCT | GTCCC | GTCCG | GTCGA | GTCGT | GTCGC | GTCGG | | GTG | GTGAA | GTGAT | GTGAC | GTGAG | GTGTA | GTGTT | GTGTC | GTGTG | GTGCA | GTGCT | GTGCC | GTGCG | GTGGA | GTGGT | GTGGC | GTGGG | | GCA | GCAAA | GCAAT | GCAAC | GCAAG | GCATA | GCATT | GCATC | GCATG | GCACA | GCACT | GCACC | GCACG | GCAGA | GCAGT | GCAGC | GCAGG | | GCT | GCTAA | GCTAT | GCTAC | GCTAG | GCTTA | GCTTT | GCTTC | GCTTG | GCTCA | GCTCT | GCTCC | GCTCG | GCTGA | GCTGT | GCTGC | GCTGG | | GCC | GCCAA | GCCAT | GCCAC | GCCAG | GCCTA | GCCTT | GCCTC | GCCTG | GCCCA | GCCCT | GCCCC | GCCCG | GCCGA | GCCGT | GCCGC | GCCGG | | GCG | GCGAA | GCGAT | GCGAC | GCGAG | GCGTA | GCGTT | GCGTC | GCGTG | GCGCA | GCGCT | GCGCC | GCGCG | GCGGA | GCGGT | GCGGC | GCGGG | | GGA | GGAAA | GGAAT | GGAAC | GGAAG | GGATA | GGATT | GGATC | GGATG | GGACA | GGACT | GGACC | GGACG | GGAGA | GGAGT | GGAGC | GGAGG | | GGT | GGTAA | GGTAT | GGTAC | GGTAG | GGTTA | GGTTT | GGTTC | GGTTG | GGTCA | GGTCT | GGTCC | GGTCG | GGTGA | GGTGT | GGTGC | GGTGG | | GGC | GGCAA | GGCAT | GGCAC | GGCAG | GGCTA | GGCTT | GGCTC | GGCTG | GGCCA | GGCCT | GGCCC | GGCCG | GGCGA | GGCGT | GGCGC | GGCGG | | GGG | GGGAA | GGGAT | GGGAC | GGGAG | GGGTA | GGGTT | GGGTC | GGGTG | GGGCA | GGGCT | GGGCC | GGGCG | GGGGA | GGGGT | GGGGC | GGGGG | After filtering: read1: overrepresented sequences Sampling rate: 1 / 20 | | | | | --- | --- | --- | | overrepresented sequence | count (% of bases) | distribution: cycle 1 ~ cycle 150 | | AAAAAAAAAA | 57742 (0.110660%) | | | AAAAAGCAACGTCGCTATGAACGCTTGGCTGCCACAAGCCAGTTATCCCTGTGGTAACTTTTCTGACACCTCTAGCTTCAAATTCCGAAGGTCTAAAGGA | 331 (0.006343%) | | | AAAAGATAACGCAGGTGTCCTAAGATGAGCTCAACGAGAACAGAAATCTCGTGTGGAACAAAAGGGTAAAAGCTCGTTTGATTCTGATTTTCAGTACGAA | 10 (0.000192%) | | | AAAAGCAACGTCGCTATGAACGCTTGGCTGCCACAAGCCAGTTATCCCTGTGGTAACTTTTCTGACACCTCTAGCTTCAAATTCCGAAGGTCTAAAGGAT | 26 (0.000498%) | | | AAAAGCCTAAGTAGTGTTTCCTTGTTAGAAGACACAAAGCCAAAGACTCATATGGACTTTGGCTACACCATGAAAGCTTTGAGAAGCAAGAAGAAGGTTG | 22 (0.000422%) | | | AAACCGCAACCGGATCTTAAAGGCGTAAGAATTGTATCCTTGTTAGAAGACACAAAGCCAAAGACTCATATGGACTTTGGCTACACCATGAAAGCTTTGA | 64 (0.001227%) | | | AAACGAGCTTTTACCCTTTTGTTCCACACGAGATTTCTGTTCTCGTTGAGCTCATCTTAGGACACCTGCGTTATCTTTTAACAGATGTGCCGCCCCAGCC | 5 (0.000096%) | | | AAACTGCGAATGGCTCATTAAATCAGTTATAGTTTGTTTGATGGTAACTACTACTCGGATAACCGTAGTAATTCTAGAGCTAATACGTGCAACAAACCCC | 4 (0.000077%) | | | AAAGAAGACCCTGTTGAGCTTGACTCTAGTCCGACTTTGTGAAATGACTTGAGAGGTGTAGGATAAGTGGGAGCTTCGGCGCAAGTGAAATACCACTACT | 11 (0.000211%) | | | AAAGACTCATATGGACTTTGGCTACACCATGAAAGCTTTGAGAAGCAAGAAGAAGGTTGGTTAGTGTTTTGGAGTCGAATATGACTTGATGTCATGTGTA | 76 (0.001457%) | | | AAAGAGAGAAAGGACGAGGTCTTGACCGTCATCTTTTGCCCGAAGGACGGATGAGCTTTGGCGGGACTGAATCACTTCGAGTCACCGTCGACAACTTTTC | 5 (0.000096%) | | | AAAGATTAAGCCATGCATGTGTAAGTATGAACGAATTCAGACTGTGAAACTGCGAATGGCTCATTAAATCAGTTATAGTTTGTTTGATGGTAACTACTAC | 13 (0.000249%) | | | AAAGCAACGTCGCTATGAACGCTTGGCTGCCACAAGCCAGTTATCCCTGTGGTAACTTTTCTGACACCTCTAGCTTCAAATTCCGAAGGTCTAAAGGATC | 275 (0.005270%) | | | AAAGCCAAAGACTCATATGGACTTTGGCTACACCATGAAAGCTTTGAGAAGCAAGAAGAAGGTTGGTTAGTGTTTTGGAGTCGAATATGACTTGATGTCA | 307 (0.005883%) | | | AAAGCCTAAGTAGTGTTTCCTTGTTAGAAGACACAAAGCCAAAGACTCATATGGACTTTGGCTACACCATGAAAGCTTTGAGAAGCAAGAAGAAGGTTGG | 7 (0.000134%) | | | AAAGCTTTCATGGTGTAGCCAAAGTCCATATGAGTCTTTGGCTTTGTGTCTTCTAACAAGGAAACACTACTTAGGCTTATAAGATGCGGTTGCGGTTTAA | 7 (0.000134%) | | | AAAGGCGTAAGAATTGTATCCTTGTTAAAAGACACAAAGCCAAAGACTCATATGGACTTTGGCTACACCATGAAAGCTTTGAGAAGCAAGAAGAAGGTTG | 10 (0.000192%) | | | AAAGGCGTAAGAATTGTATCCTTGTTAGAAGACACAAAGCCAAAGACTCATATGGACTTTGGCTACACCATGAAAGCTTTGAGAAGCAAGAAGAAGGTTG | 13 (0.000249%) | | | AAATCAGAATCAAACGAGCTTTTACCCTTTTGTTCCACACGAGATTTCTGTTCTCGTTGAGCTCATCTTAGGACACCTGCGTTATCTTTTAACAGATGTG | 4 (0.000077%) | | | AAATCAGTTATAGTTTGTTTGATGGTAACTACTACTCGGATAACCGTAGTAATTCTAGAGCTAATACGTGCAACAAACCCCGACTTATGGAAGGGACGCA | 5 (0.000096%) | | | AAATCCGCTAAGGAGTGTGTAACAACTCACCTGCCGAATCAACTAGCCCCGAAAATGGATGGCGCTTAAGCGCGCGACCTATACCCGGCCGTCGGGGCAA | 8 (0.000153%) | | | AAATCCTATGATGTTATCCCATGCTAATGTATCCAGAGCGTAGGCTTGCTTTGAGCACTCTAATTTCTTCAAAGTAACAGCGCCGGAGGCACGACCCGGC | 27 (0.000517%) | | | AAATTTTCATAATTTTTTGACACCTCTAGCTAGGTCATTTGACCTGATACAACATCGGATTTTCATGGTCTAGTTGGGGCTCCGTGGGCATATTTGATGC | 17 (0.000326%) | | | AACAGGGACAGTCGGGGGCATTCGTATTTCATAGTCAGAGGTGAAATTCTTGGATTTATGAAAGACGAACAACTGCGAAAGCATTTGCCAAGGATGTTTT | 19 (0.000364%) | | | AACAGGTCTGTGATGCCCTTAGATGTTCTGGGCCGCACGCGCGCTACACTGATGTATTCAACGAGTTCACACCTTGGCCGACAGGCCCGGGTAATCTTTG | 14 (0.000268%) | | | AACCTAGGCGAGACAAGGGTTCACATTTCGTTCATCACCCTTGGCCGGCTATCGAACAGCCGGACTCCCATCAAAAGATGGTTGCCAAGAACATCTTCGT | 3 (0.000057%) | | | AACCTAGGCGAGACAAGGGTTCACATTTCGTTCATCACCCTTGGCCGGCTTTCGAACAGCCGGACTCCCATCAAAAGATGGTTGCCAAGAACATCTTCGT | 5 (0.000096%) | | | AACGAAGCACGCCCATCCAACCTAGGCGAGACAAGGGTTCACATTTCGTTCATCACCCTTGGCCGGCTATCGAACAGCCGGACTCCCATCAAAAGATGGT | 21 (0.000402%) | | | AACGAAGCACGCCCATCCAACCTAGGCGAGACAAGGGTTCACATTTCGTTCATCACCCTTGGCCGGCTTTCGAACAGCCGGACTCCCATCAAAAGATGGT | 19 (0.000364%) | | | AACGCCTCGAAGAACTAATGGCAGCCCACGCAAGGCAAGCCCATTCTCCTCGACGATTCAGCAGTTTTTGTCCGAGAACTGCTGAGAAAACTCGGAAAAA | 11 (0.000211%) | | | AACGGCGTGCCTCGGCATCAGCGTGCTCCGGGCGTCGGCCTGTGGGCTCCCCATTCGACCCGTCTTGAAACACGGACCAAGGAGTCTGACATGTGTGCGA | 4 (0.000077%) | | | AACGGGCAGAGCCCGCGTCGACCTTTTATCTAATAAATGCGTCCCTTCCATAAGTCGGGGTTTGTTGCACGTATTAGCTCTAGAATTACTACGGTTATCC | 14 (0.000268%) | | | AACGGGTGACGGAGAATTAGGGTTCGATTCCGGAGAGGGAGCCTGAGAAACGGCTACCACATCCAAGGAAGGCAGCAGGCGCGCAAATTACCCAATCCTG | 4 (0.000077%) | | | AAGAAGACCCTGTTGAGCTTGACTCTAGTCCGACTTTGTGAAATGACTTGAGAGGTGTAGGATAAGTGGGAGCTTCGGCGCAAGTGAAATACCACTACTT | 11 (0.000211%) | | | AAGAATTGTATCCTTGTTAGAAGACACAAAGCCAAAGACTCATATGGACTTTGGCTACACCATGAAAGCTTTGAGAAGCAAGAAGAAGGTTGGTTAGTGT | 22 (0.000422%) | | | AAGACACAAAGCCAAAGACTCATATGGACTTTGGCTACACCATGAAAGCTTTGAGAAGCAAGAAGAAGGTTGGTTAGTGTTTTGGAGTCGAATATGACTT | 617 (0.011824%) | | | AAGACCCTGTTGAGCTTGACTCTAGTCCGACTTTGTGAAATGACTTGAGAGGTGTAGGATAAGTGGGAGCTTCGGCGCAAGTGAAATACCACTACTTTTA | 4 (0.000077%) | | | AAGACTCATATGGACTTTGGCTACACCATGAAAGCTTTGAGAAGCAAGAAGAAGGTTGGTTAGTGTTTTGGAGTCGAATATGACTTGATGTCATGTGTAT | 25 (0.000479%) | | | AAGAGCCCGGTATTGTTATTTATTGTCACTACCTCCCCGTGTCAGGATTGGGTAATTTGCGCGCCTGCTGCCTTCCTTGGATGTGGTAGCCGTTTCTCAG | 9 (0.000172%) | | | AAGATCGATC | 199249 (0.381850%) | | | AAGCAACGTCGCTATGAACGCTTGGCTGCCACAAGCCAGTTATCCCTGTGGTAACTTTTCTGACACCTCTAGCTTCAAATTCCGAAGGTCTAAAGGATCG | 130 (0.002491%) | | | AAGCCAAAGACTCATATGGACTTTGGCTACACCATGAAAGCTTTGAGAAGCAAGAAGAAGGTTGGTTAGTGTTTTGGAGTCGAATATGACTTGATGTCAT | 12 (0.000230%) | | | AAGCCATGCATGTGTAAGTATGAACGAATTCAGACTGTGAAACTGCGAATGGCTCATTAAATCAGTTATAGTTTGTTTGATGGTAACTACTACTCGGATA | 5 (0.000096%) | | | AAGCCTAAGTAGTGTTTCCTTGTTAGAAGACACAAAGCCAAAGACTCATATGGACTTTGGCTACACCATGAAAGCTTTGAGAAGCAAGAAGAAGGTTGGT | 1195 (0.022902%) | | | AAGCGCGCGACCTATACCCGGCCGTCGGGGCAAGAGCCAGGCCTCGATGAGTAGGAGGGCGCGGCGGTCGCTGCAAAACCTAGGGCGCGAGCCCGGGCGG | 18 (0.000345%) | | | AAGCTTTCATGGTGTAGCCAAAGTCCATATGAGTCTTTGGCTTTGTGTCTTCTAACAAGGAAACACTACTTAGGCTTATAAGATGCGGTTGCGGTTTAAG | 3 (0.000057%) | | | AAGGATACTAAATCCTATTTTCTGGTAAATTTTCATAATTTTTTGACACCTCTAGCTAGGTCATTTGACCTGATACAACATCGGATTTTCATGGTCTAGT | 7 (0.000134%) | | | AAGGCGTAAGAATTGTATCCTTGTTAAAAGACACAAAGCCAAAGACTCATATGGACTTTGGCTACACCATGAAAGCTTTGAGAAGCAAGAAGAAGGTTGG | 5 (0.000096%) | | | AAGGCGTAAGAATTGTATCCTTGTTAGAAGACACAAAGCCAAAGACTCATATGGACTTTGGCTACACCATGAAAGCTTTGAGAAGCAAGAAGAAGGTTGG | 7 (0.000134%) | | | AAGGGATTTAGATTGTACTCATTCCAATTACCAGACTCGAAAGAGCCCGGTATTGTTATTTATTGTCACTACCTCCCCGTGTCAGGATTGGGTAATTTGC | 51 (0.000977%) | | | AAGGGTTCACATTTCGTTCATCACCCTTGGCCGGCTATCGAACAGCCGGACTCCCATCAAAAGATGGTTGCCAAGAACATCTTCGTTACGGTTTGCTAAT | 4 (0.000077%) | | | AAGTAGTGTTTCCTTGTTAGAAGACACAAAGCCAAAGACTCATATGGACTTTGGCTACACCATGAAAGCTTTGAGAAGCAAGAAGAAGGTTGGTTAGTGT | 28 (0.000537%) | | | AAGTATGAACGAATTCAGACTGTGAAACTGCGAATGGCTCATTAAATCAGTTATAGTTTGTTTGATGGTAACTACTACTCGGATAACCGTAGTAATTCTA | 3 (0.000057%) | | | AAGTCATATTCGACTCCAAAACACTAACCAACCTTCTTCTTGCTTCTCAAAGCTTTCATGGTGTAGCCAAAGTCCATATGAGTCTTTGGCTTTGTGTCTT | 138 (0.002645%) | | | AAGTCTGGTGCCAGCAGCCGCGGTAATTCCAGCTCCAATAGCGTATATTTAAGTTGTTGCAGTTAAAAAGCTCGTAGTTGAACCTTGGGATGGGTCGGCC | 5 (0.000096%) | | | AAGTTCACCACCGCATGTCGGTACGCTCCAGGCGTCCTTGGCTCGGATTTAGGCCAACCGCGTGCGGTAACACACGGGAGACCAGCTTCCGTCCCGCATC | 16 (0.000307%) | | | AAGTTCTTATACTCAATCATACACATGACATCAAGTCATATTCGACTCCAAAACACTAACCAACCTTCTTCTTGCTTCTCAAAGCTTTCATGGTGTAGCC | 45 (0.000862%) | | | AAGTTGTTATACTCAATCATACACATGACATCAAGTCATA | 115 (0.000882%) | | | AATAACAGGTCTGTGATGCCCTTAGATGTTCTGGGCCGCACGCGCGCTACACTGATGTATTCAACGAGTTCACACCTTGGCCGACAGGCCCGGGTAATCT | 14 (0.000268%) | | | AATATGACTTGATGTCATGTGTATGATTGAGTATAAGAAC | 1373 (0.010525%) | | | AATCAAACGAGCTTTTACCCTTTTGTTCCACACGAGATTTCTGTTCTCGTTGAGCTCATCTTAGGACACCTGCGTTATCTTTTAACAGATGTGCCGCCCC | 28 (0.000537%) | | | AATCAGTTATAGTTTGTTTGATGGTAACTACTACTCGGATAACCGTAGTAATTCTAGAGCTAATACGTGCAACAAACCCCGACTTATGGAAGGGACGCAT | 5 (0.000096%) | | | AATCATACACATGACAACAAGTCATATTCGACTCCAAAACACTAACCAACCTTCTTCTTGCTTCTCAAAGCTTTCATGGTGTAGCCAAAGTCCATATGAG | 16 (0.000307%) | | | AATCATACACATGACATCAAGTCATATTCGACTCCAAAACACTAACCAACCTTCTTCTTGCTTCTCAAAGCTTTCATGGTGTAGCCAAAGTCCATATGAG | 70 (0.001342%) | | | AATCATCAGAGCAACGGGCAGAGCCCGCGTCGACCTTTTATCTAATAAATGCGTCCCTTCCATAAGTCGGGGTTTGTTGCACGTATTAGCTCTAGAATTA | 45 (0.000862%) | | | AATCCGCTAAGGAGTGTGTAACAACTCACCTGCCGAATCAACTAGCCCCGAAAATGGATGGCGCTTAAGCGCGCGACCTATACCCGGCCGTCGGGGCAAG | 14 (0.000268%) | | | AATCCTATGATGTTATCCCATGCTAATGTATCCAGAGCGTAGGCTTGCTTTGAGCACTCTAATTTCTTCAAAGTAACAGCGCCGGAGGCACGACCCGGCC | 34 (0.000652%) | | | AATCCTTGTTAGTTTCTTTTCCTCCGCTTATTGATATGCTTAAACTCAGCGGGTAATCCCGCCTGACCTGGGGTCGCTATATGGACTTTGGGTCATCTAC | 7 (0.000134%) | | | AATCGGTAGGAGCGACGGGCGGTGTGTACAAAGGGCAGGGACGTAGTCAACGCGAGCTGATGACTCGCGCTTACTAGGAATTCCTCGTTGAAGACCAACA | 68 (0.001303%) | | | AATGATTAACAGGGACAGTCGGGGGCATTCGTATTTCATAGTCAGAGGTGAAATTCTTGGATTTATGAAAGACGAACAACTGCGAAAGCATTTGCCAAGG | 89 (0.001706%) | | | AATGTATCCAGAGCGTAGGCTTGCTTTGAGCACTCTAATTTCTTCAAAGTAACAGCGCCGGAGGCACGACCCGGCCAATTAAGACCAGGAGCGTATCGCC | 8 (0.000153%) | | | AATTACCAGACTCGAAAGAGCCCGGTATTGTTATTTATTGTCACTACCTCCCCGTGTCAGGATTGGGTAATTTGCGCGCCTGCTGCCTTCCTTGGATGTG | 21 (0.000402%) | | | AATTAGGGTTCGATTCCGGAGAGGGAGCCTGAGAAACGGCTACCACATCCAAGGAAGGCAGCAGGCGCGCAAATTACCCAATCCTGACACGGGGAGGTAG | 17 (0.000326%) | | | AATTCCAGCTCCAATAGCGTATATTTAAGTTGTTGCAGTTAAAAAGCTCGTAGTTGAACCTTGGGATGGGTCGGCCGGTCCGCCTTTGGTGTGCATTGGT | 10 (0.000192%) | | | AATTCCCCGCCACATCCTCTCAAACGCAATGGAAAGAGAGAAAGGACGAGGTCTTGACCGTCATCTTTTGCCCGAAGGACGGATGAGCTTTGGCGGGACT | 13 (0.000249%) | | | AATTGTATCCTTGTTAGAAGACACAAAGCCAAAGACTCATATGGACTTTGGCTACACCATGAAAGCTTTGAGAAGCAAGAAGAAGGTTGGTTAGTGTTTT | 4 (0.000077%) | | | AATTGTTGGTCTTCAACGAGGAATTCCTAGTAAGCGCGAGTCATCAGCTCGCGTTGACTACGTCCCTGCCCTTTGTACACACCGCCCGTCGCTCCTACCG | 1354 (0.025949%) | | | AATTTTCATAATTTTTTGACACCTCTAGCTAGGTCATTTGACCTGATACAACATCGGATTTTCATGGTCTAGTTGGGGCTCCGTGGGCATATTTGATGCA | 3 (0.000057%) | | | ACAAAGCCAAAGACTCATATGGACTTTGGCTACACCATGAAAGCTTTGAGAAGCAAGAAGAAGGTTGGTTAGTGTTTTGGAGTCGAATATGACTTGATGT | 142 (0.002721%) | | | ACAAGGGTTCACATTTCGTTCATCACCCTTGGCCGGCTTTCGAACAGCCGGACTCCCATCAAAAGATGGTTGCCAAGAACATCTTCGTTACGGTTTGCTA | 3 (0.000057%) | | | ACACAAAGCCAAAGACTCATATGGACTTTGGCTACACCATGAAAGCTTTGAGAAGCAAGAAGAAGGTTGGTTAGTGTTTTGGAGTCGAATATGACTTGAT | 171 (0.003277%) | | | ACACATGACATCAAGTCATATTCGACTCCAAAACACTAACCAACCTTCTTCTTGCTTCTCAAAGCTTTCATGGTGTAGCCAAAGTCCATATGAGTCTTTG | 32 (0.000613%) | | | ACACCATGAAAGCTTTGAGAAGCAAGAAGAAGGTTGGTTAGTGTTTTGGAGTCGAATATGACTTGATGTCATGTGTATGATTGAGTATAACAACTTAAAC | 4 (0.000077%) | | | ACAGGTCTGTGATGCCCTTAGATGTTCTGGGCCGCACGCGCGCTACACTGATGTATTCAACGAGTTCACACCTTGGCCGACAGGCCCGGGTAATCTTTGA | 4 (0.000077%) | | | ACATCAAGTCATATTCGACTCCAAAACACTAACCAACCTTCTTCTTGCTTCTCAAAGCTTTCATGGTGTAGCCAAAGTCCATATGAGTCTTTGGCTTTGT | 48 (0.000920%) | | | ACATCTGTTAAAAGATAACGCAGGTGTCCTAAGATGAGCTCAACGAGAACAGAAATCTCGTGTGGAACAAAAGGGTAAAAGCTCGTTTGATTCTGATTTT | 3 (0.000057%) | | | ACATGACATCAAGTCATATTCGACTCCAAAACACTAACCAACCTTCTTCTTGCTTCTCAAAGCTTTCATGGTGTAGCCAAAGTCCATATGAGTCTTTGGC | 67 (0.001284%) | | | ACATTGTCAGGTGGGGAGTTTGGCTGGGGCGGCACATCTGTTAAAAGATAACGCAGGTGTCCTAAGATGAGCTCAACGAGAACAGAAATCTCGTGTGGAA | 7 (0.000134%) | | | ACCACCGCATGTCGGTACGCTCCAGGCGTCCTTGGCTCGGATTTAGGCCAACCGCGTGCGGTAACACACGGGAGACCAGCTTCCGTCCCGCATCAGCAAA | 5 (0.000096%) | | | ACCAGAGGCTGTTCACCTTGGAGACCTGATGCGGTTATGAGTACGACCGGGCGTGAGCGGCACTCGGTCCTCCGGATTTTCAAGGGCCGCCGGGGGCGCA | 14 (0.000268%) | | | ACCATCCTTTGCTGATGCGGGACGGAAGCTGGTCTCCCGTGTGTTACCGCACGCGGTTGGCCTAAATCCGAGCCAAGGACGCCTGGAGCGTACCGACATG | 9 (0.000172%) | | | ACCCTAACGCCTCGAAGAACTAATGGCAGCCCACGCAAGGCAAGCCCATTCTCCTCGACGATTCAGCAGTTTTTGTCCGAGAACTGCTGAGAAAACTCGG | 5 (0.000096%) | | | ACCCTGTTGAGCTTGACTCTAGTCCGACTTTGTGAAATGACTTGAGAGGTGTAGGATAAGTGGGAGCTTCGGCGCAAGTGAAATACCACTACTTTTAACG | 11 (0.000211%) | | | ACCGCATGTCGGTACGCTCCAGGCGTCCTTGGCTCGGATTTAGGCCAACCGCGTGCGGTAACACACGGGAGACCAGCTTCCGTCCCGCATCAGCAAAGGA | 5 (0.000096%) | | | ACCGGATTGCTCCGTTCCGCATCCGACCAGGACGCATCGCCGGCCCCCATCCGCTTCCCTCCCGACAATTTCAAGCACTCTTTGACTCTCTTTTCAAAGT | 6 (0.000115%) | | | ACCTAGGCGAGACAAGGGTTCACATTTCGTTCATCACCCTTGGCCGGCTATCGAACAGCCGGACTCCCATCAAAAGATGGTTGCCAAGAACATCTTCGTT | 6 (0.000115%) | | | ACCTAGGCGAGACAAGGGTTCACATTTCGTTCATCACCCTTGGCCGGCTTTCGAACAGCCGGACTCCCATCAAAAGATGGTTGCCAAGAACATCTTCGTT | 7 (0.000134%) | | | ACGAAGCACGCCCATCCAACCTAGGCGAGACAAGGGTTCACATTTCGTTCATCACCCTTGGCCGGCTATCGAACAGCCGGACTCCCATCAAAAGATGGTT | 9 (0.000172%) | | | ACGAAGCACGCCCATCCAACCTAGGCGAGACAAGGGTTCACATTTCGTTCATCACCCTTGGCCGGCTTTCGAACAGCCGGACTCCCATCAAAAGATGGTT | 7 (0.000134%) | | | ACGCGCCTAACGGCGTGCCTCGGCATCAGCGTGCTCCGGGCGTCGGCCTGTGGGCTCCCCATTCGACCCGTCTTGAAACACGGACCAAGGAGTCTGACAT | 21 (0.000402%) | | | ACGCTTTCACGGTTCGTATTCGTACTGAAAATCAGAATCAAACGAGCTTTTACCCTTTTGTTCCACACGAGATTTCTGTTCTCGTTGAGCTCATCTTAGG | 11 (0.000211%) | | | ACGGGCAGAGCCCGCGTCGACCTTTTATCTAATAAATGCGTCCCTTCCATAAGTCGGGGTTTGTTGCACGTATTAGCTCTAGAATTACTACGGTTATCCG | 4 (0.000077%) | | | ACGGGCGGTGTGTACAAAGGGCAGGGACGTAGTCAACGCGAGCTGATGACTCGCGCTTACTAGGAATTCCTCGTTGAAGACCAACAATTGCAATGATCGA | 3 (0.000057%) | | | ACGTGGGTGGTTCGCCGCCCGCGACGTCGCGAGAAGTCCACTAAACCTTATCATTTAGAGGAAGGAGAAGTCGTAACAAGGTTTCCGTAGGTGAACCTGC | 4 (0.000077%) | | | ACTAAATCCTATTTTCTGGTAAATTTTCATAATTTTTTGACACCTCTAGCTAGGTCATTTGACCTGATACAACATCGGATTTTCATGGTCTAGTTGGGGC | 20 (0.000383%) | | | ACTCAATCATACACATGACAACAAGTCATATTCGACTCCAAAACACTAACCAACCTTCTTCTTGCTTCTCAAAGCTTTCATGGTGTAGCCAAAGTCCATA | 6 (0.000115%) | | | ACTCAATCATACACATGACATCAAGTCATATTCGACTCCAAAACACTAACCAACCTTCTTCTTGCTTCTCAAAGCTTTCATGGTGTAGCCAAAGTCCATA | 25 (0.000479%) | | | ACTCATATGGACTTTGGCTACACCATGAAAGCTTTGAGAAGCAAGAAGAAGGTTGGTTAGTGTTTTGGAGTCGAATATGACTTGATGTCATGTGTATGAT | 74 (0.001418%) | | | ACTCATTCCAATTACCAGACTCGAAAGAGCCCGGTATTGTTATTTATTGTCACTACCTCCCCGTGTCAGGATTGGGTAATTTGCGCGCCTGCTGCCTTCC | 43 (0.000824%) | | | ACTCCAAAACACTAACCAACCTTCTTCTTGCTTCTCAAAGCTTTCATGGTGTAGCCAAAGTCCATATGAGTCTTTGGCTTTGTGTCTTCTAACAAGGAAA | 177 (0.003392%) | | | ACTCGAAAGAGCCCGGTATTGTTATTTATTGTCACTACCTCCCCGTGTCAGGATTGGGTAATTTGCGCGCCTGCTGCCTTCCTTGGATGTGGTAGCCGTT | 4 (0.000077%) | | | ACTGAAAATCAGAATCAAACGAGCTTTTACCCTTTTGTTCCACACGAGATTTCTGTTCTCGTTGAGCTCATCTTAGGACACCTGCGTTATCTTTTAACAG | 6 (0.000115%) | | | ACTTCCCTTGCCTACATTGTTCCATCGACCAGAGGCTGTTCACCTTGGAGACCTGATGCGGTTATGAGTACGACCGGGCGTGAGCGGCACTCGGTCCTCC | 23 (0.000441%) | | | ACTTGATGTCATGTGTATGATTGAGTATAAGAACTTAAAC | 319 (0.002445%) | | | ACTTTGTGAAATGACTTGAGAGGTGTAGGATAAGTGGGAGCTTCGGCGCAAGTGAAATACCACTACTTTTAACGTTATTTTACTTACTCCGTGAATCGGA | 5 (0.000096%) | | | AGAAGACACAAAGCCAAAGACTCATATGGACTTTGGCTACACCATGAAAGCTTTGAGAAGCAAGAAGAAGGTTGGTTAGTGTTTTGGAGTCGAATATGAC | 71 (0.001361%) | | | AGAAGACCCTGTTGAGCTTGACTCTAGTCCGACTTTGTGAAATGACTTGAGAGGTGTAGGATAAGTGGGAGCTTCGGCGCAAGTGAAATACCACTACTTT | 31 (0.000594%) | | | AGAATTAGGGTTCGATTCCGGAGAGGGAGCCTGAGAAACGGCTACCACATCCAAGGAAGGCAGCAGGCGCGCAAATTACCCAATCCTGACACGGGGAGGT | 4 (0.000077%) | | | AGAATTGTATCCTTGTTAGAAGACACAAAGCCAAAGACTCATATGGACTTTGGCTACACCATGAAAGCTTTGAGAAGCAAGAAGAAGGTTGGTTAGTGTT | 7 (0.000134%) | | | AGACACAAAGCCAAAGACTCATATGGACTTTGGCTACACCATGAAAGCTTTGAGAAGCAAGAAGAAGGTTGGTTAGTGTTTTGGAGTCGAATATGACTTG | 57 (0.001092%) | | | AGACCCTGTTGAGCTTGACTCTAGTCCGACTTTGTGAAATGACTTGAGAGGTGTAGGATAAGTGGGAGCTTCGGCGCAAGTGAAATACCACTACTTTTAA | 5 (0.000096%) | | | AGACTCATATGGACTTTGGCTACACCATGAAAGCTTTGAGAAGCAAGAAGAAGGTTGGTTAGTGTTTTGGAGTCGAATATGACTTGATGTCATGTGTATG | 8 (0.000153%) | | | AGACTCGAAAGAGCCCGGTATTGTTATTTATTGTCACTACCTCCCCGTGTCAGGATTGGGTAATTTGCGCGCCTGCTGCCTTCCTTGGATGTGGTAGCCG | 6 (0.000115%) | | | AGAGCAACGGGCAGAGCCCGCGTCGACCTTTTATCTAATAAATGCGTCCCTTCCATAAGTCGGGGTTTGTTGCACGTATTAGCTCTAGAATTACTACGGT | 30 (0.000575%) | | | AGAGCCCGCGTCGACCTTTTATCTAATAAATGCGTCCCTTCCATAAGTCGGGGTTTGTTGCACGTATTAGCTCTAGAATTACTACGGTTATCCGAGTAGT | 10 (0.000192%) | | | AGAGCGTAGGCTTGCTTTGAGCACTCTAATTTCTTCAAAGTAACAGCGCCGGAGGCACGACCCGGCCAATTAAGACCAGGAGCGTATCGCCGACCGAAGG | 10 (0.000192%) | | | AGAGGTGAAATTCTTGGATTTATGAAAGACGAACAACTGCGAAAGCATTTGCCAAGGATGTTTTCATTAATCAAGAACGAAAGTTGGGGGCTCGAAGACG | 4 (0.000077%) | | | AGATCGGAAGAGCACACGTCTGAACTCCAGTCACGAGATT | 52 (0.000399%) | | | AGATGTTCTGGGCCGCACGCGCGCTACACTGATGTATTCAACGAGTTCACACCTTGGCCGACAGGCCCGGGTAATCTTTGAAATTTCATCGTGATGGGGA | 4 (0.000077%) | | | AGATTAAGCCATGCATGTGTAAGTATGAACGAATTCAGACTGTGAAACTGCGAATGGCTCATTAAATCAGTTATAGTTTGTTTGATGGTAACTACTACTC | 5 (0.000096%) | | | AGATTGTACTCATTCCAATTACCAGACTCGAAAGAGCCCGGTATTGTTATTTATTGTCACTACCTCCCCGTGTCAGGATTGGGTAATTTGCGCGCCTGCT | 5 (0.000096%) | | | AGCAACGGGCAGAGCCCGCGTCGACCTTTTATCTAATAAATGCGTCCCTTCCATAAGTCGGGGTTTGTTGCACGTATTAGCTCTAGAATTACTACGGTTA | 3 (0.000057%) | | | AGCAACGTCGCTATGAACGCTTGGCTGCCACAAGCCAGTTATCCCTGTGGTAACTTTTCTGACACCTCTAGCTTCAAATTCCGAAGGTCTAAAGGATCGA | 16 (0.000307%) | | | AGCACGCCCATCCAACCTAGGCGAGACAAGGGTTCACATTTCGTTCATCACCCTTGGCCGGCTATCGAACAGCCGGACTCCCATCAAAAGATGGTTGCCA | 17 (0.000326%) | | | AGCACGCCCATCCAACCTAGGCGAGACAAGGGTTCACATTTCGTTCATCACCCTTGGCCGGCTTTCGAACAGCCGGACTCCCATCAAAAGATGGTTGCCA | 20 (0.000383%) | | | AGCACGCGCCTAACGGCGTGCCTCGGCATCAGCGTGCTCCGGGCGTCGGCCTGTGGGCTCCCCATTCGACCCGTCTTGAAACACGGACCAAGGAGTCTGA | 36 (0.000690%) | | | AGCAGCCGCGGTAATTCCAGCTCCAATAGCGTATATTTAAGTTGTTGCAGTTAAAAAGCTCGTAGTTGAACCTTGGGATGGGTCGGCCGGTCCGCCTTTG | 8 (0.000153%) | | | AGCCAAAGACTCATATGGACTTTGGCTACACCATGAAAGCTTTGAGAAGCAAGAAGAAGGTTGGTTAGTGTTTTGGAGTCGAATATGACTTGATGTCATG | 11 (0.000211%) | | | AGCCGCGGTAATTCCAGCTCCAATAGCGTATATTTAAGTTGTTGCAGTTAAAAAGCTCGTAGTTGAACCTTGGGATGGGTCGGCCGGTCCGCCTTTGGTG | 6 (0.000115%) | | | AGCCTAAGTAGTGTTTCCTTGTTAGAAGACACAAAGCCAAAGACTCATATGGACTTTGGCTACACCATGAAAGCTTTGAGAAGCAAGAAGAAGGTTGGTT | 35 (0.000671%) | | | AGCGACGGGCGGTGTGTACAAAGGGCAGGGACGTAGTCAACGCGAGCTGATGACTCGCGCTTACTAGGAATTCCTCGTTGAAGACCAACAATTGCAATGA | 6 (0.000115%) | | | AGGACATTGTCAGGTGGGGAGTTTGGCTGGGGCGGCACATCTGTTAAAAGATAACGCAGGTGTCCTAAGATGAGCTCAACGAGAACAGAAATCTCGTGTG | 154 (0.002951%) | | | AGGAGCGACGGGCGGTGTGTACAAAGGGCAGGGACGTAGTCAACGCGAGCTGATGACTCGCGCTTACTAGGAATTCCTCGTTGAAGACCAACAATTGCAA | 13 (0.000249%) | | | AGGCCACGCTTTCACGGTTCGTATTCGTACTGAAAATCAGAATCAAACGAGCTTTTACCCTTTTGTTCCACACGAGATTTCTGTTCTCGTTGAGCTCATC | 20 (0.000383%) | | | AGGCGAGACAAGGGTTCACATTTCGTTCATCACCCTTGGCCGGCTATCGAACAGCCGGACTCCCATCAAAAGATGGTTGCCAAGAACATCTTCGTTACGG | 3 (0.000057%) | | | AGGCGAGACAAGGGTTCACATTTCGTTCATCACCCTTGGCCGGCTTTCGAACAGCCGGACTCCCATCAAAAGATGGTTGCCAAGAACATCTTCGTTACGG | 4 (0.000077%) | | | AGGCGTAAGAATTGTATCCTTGTTAAAAGACACAAAGCCAAAGACTCATATGGACTTTGGCTACACCATGAAAGCTTTGAGAAGCAAGAAGAAGGTTGGT | 13 (0.000249%) | | | AGGCGTAAGAATTGTATCCTTGTTAGAAGACACAAAGCCAAAGACTCATATGGACTTTGGCTACACCATGAAAGCTTTGAGAAGCAAGAAGAAGGTTGGT | 11 (0.000211%) | | | AGGCTGTCCCGAGTGTGAGCGAGGTGTGAGTGTCGCCCATGGGCATCGACACCTTGCGGCTAGGAACTGGAACGAGACGGGTAGCAAAGATTTCGAGTAG | 11 (0.000211%) | | | AGGCTGTCCCGAGTGTGAGCGAGGTGTGAGTGTCGCCCATGGGCATCGACACCTTGCGGCTAGGAACTGGAACGAGACGGGTGGCAAAGATTTCGAGTAG | 11 (0.000211%) | | | AGGCTTGCTTTGAGCACTCTAATTTCTTCAAAGTAACAGCGCCGGAGGCACGACCCGGCCAATTAAGACCAGGAGCGTATCGCCGACCGAAGGGACAAGC | 4 (0.000077%) | | | AGGGAATCCTTGTTAGTTTCTTTTCCTCCGCTTATTGATATGCTTAAACTCAGCGGGTAATCCCGCCTGACCTGGGGTCGCTATATGGACTTTGGGTCAT | 216 (0.004140%) | | | AGGGACAGTCGGGGGCATTCGTATTTCATAGTCAGAGGTGAAATTCTTGGATTTATGAAAGACGAACAACTGCGAAAGCATTTGCCAAGGATGTTTTCAT | 6 (0.000115%) | | | AGGGATTTAGATTGTACTCATTCCAATTACCAGACTCGAAAGAGCCCGGTATTGTTATTTATTGTCACTACCTCCCCGTGTCAGGATTGGGTAATTTGCG | 10 (0.000192%) | | | AGGGCAAGTCTGGTGCCAGCAGCCGCGGTAATTCCAGCTCCAATAGCGTATATTTAAGTTGTTGCAGTTAAAAAGCTCGTAGTTGAACCTTGGGATGGGT | 27 (0.000517%) | | | AGGTCTGTGATGCCCTTAGATGTTCTGGGCCGCACGCGCGCTACACTGATGTATTCAACGAGTTCACACCTTGGCCGACAGGCCCGGGTAATCTTTGAAA | 3 (0.000057%) | | | AGGTGGGGAGTTTGGCTGGGGCGGCACATCTGTTAAAAGATAACGCAGGTGTCCTAAGATGAGCTCAACGAGAACAGAAATCTCGTGTGGAACAAAAGGG | 6 (0.000115%) | | | AGGTGTGAGTGTCGCCCATGGGCATCGACACCTTGCGGCTAGGAACTGGAACGAGACGGGTGGCAAAGATTTCGAGTAGCACTTCATACTACCGTGGGTT | 3 (0.000057%) | | | AGTAGTCATATGCTTGTCTCAAAGATTAAGCCATGCATGTGTAAGTATGAACGAATTCAGACTGTGAAACTGCGAATGGCTCATTAAATCAGTTATAGTT | 89 (0.001706%) | | | AGTAGTGTTTCCTTGTTAGAAGACACAAAGCCAAAGACTCATATGGACTTTGGCTACACCATGAAAGCTTTGAGAAGCAAGAAGAAGGTTGGTTAGTGTT | 12 (0.000230%) | | | AGTCAGAGGTGAAATTCTTGGATTTATGAAAGACGAACAACTGCGAAAGCATTTGCCAAGGATGTTTTCATTAATCAAGAACGAAAGTTGGGGGCTCGAA | 9 (0.000172%) | | | AGTCATATGCTTGTCTCAAAGATTAAGCCATGCATGTGTAAGTATGAACGAATTCAGACTGTGAAACTGCGAATGGCTCATTAAATCAGTTATAGTTTGT | 18 (0.000345%) | | | AGTCGAAATCCGCTAAGGAGTGTGTAACAACTCACCTGCCGAATCAACTAGCCCCGAAAATGGATGGCGCTTAAGCGCGCGACCTATACCCGGCCGTCGG | 7 (0.000134%) | | | AGTCGGGGGCATTCGTATTTCATAGTCAGAGGTGAAATTCTTGGATTTATGAAAGACGAACAACTGCGAAAGCATTTGCCAAGGATGTTTTCATTAATCA | 4 (0.000077%) | | | AGTGTATCCTTGTTAGAAGACACAAAGCCAAAGACTCATATGGACTTTGGCTACACCATGAAAGCTTTGAGAAGCAAGAAGAAGGTTGGTTAGTGTTTTG | 756 (0.014488%) | | | AGTGTTTCCTTGTTAGAAGACACAAAGCCAAAGACTCATATGGACTTTGGCTACACCATGAAAGCTTTGAGAAGCAAGAAGAAGGTTGGTTAGTGTTTTG | 13 (0.000249%) | | | AGTTATCATGAATCATCAGAGCAACGGGCAGAGCCCGCGTCGACCTTTTATCTAATAAATGCGTCCCTTCCATAAGTCGGGGTTTGTTGCACGTATTAGC | 6 (0.000115%) | | | AGTTCACCACCGCATGTCGGTACGCTCCAGGCGTCCTTGGCTCGGATTTAGGCCAACCGCGTGCGGTAACACACGGGAGACCAGCTTCCGTCCCGCATCA | 7 (0.000134%) | | | AGTTCTTATACTCAATCATACACATGACATCAAGTCATATTCGACTCCAAAACACTAACCAACCTTCTTCTTGCTTCTCAAAGCTTTCATGGTGTAGCCA | 26 (0.000498%) | | | AGTTGTTATACTCAATCATACACATGACATCAAGTCATAT | 83 (0.000636%) | | | AGTTTGGCTGGGGCGGCACATCTGTTAAAAGATAACGCAGGTGTCCTAAGATGAGCTCAACGAGAACAGAAATCTCGTGTGGAACAAAAGGGTAAAAGCT | 5 (0.000096%) | | | ATAACAGGTCTGTGATGCCCTTAGATGTTCTGGGCCGCACGCGCGCTACACTGATGTATTCAACGAGTTCACACCTTGGCCGACAGGCCCGGGTAATCTT | 15 (0.000287%) | | | ATAAGGATACTAAATCCTATTTTCTGGTAAATTTTCATAATTTTTTGACACCTCTAGCTAGGTCATTTGACCTGATACAACATCGGATTTTCATGGTCTA | 20 (0.000383%) | | | ATAATTTTTTGACACCTCTAGCTAGGTCATTTGACCTGATACAACATCGGATTTTCATGGTCTAGTTGGGGCTCCGTGGGCATATTTGATGCAAACTTGA | 8 (0.000153%) | | | ATACACATGACATCAAGTCATATTCGACTCCAAAACACTAACCAACCTTCTTCTTGCTTCTCAAAGCTTTCATGGTGTAGCCAAAGTCCATATGAGTCTT | 293 (0.005615%) | | | ATACCCGGCCGTCGGGGCAAGAGCCAGGCCTCGATGAGTAGGAGGGCGCGGCGGTCGCTGCAAAACCTAGGGCGCGAGCCCGGGCGGAGCGGCCGTCGGT | 8 (0.000153%) | | | ATACTAAATCCTATTTTCTGGTAAATTTTCATAATTTTTTGACACCTCTAGCTAGGTCATTTGACCTGATACAACATCGGATTTTCATGGTCTAGTTGGG | 5 (0.000096%) | | | ATACTCAATCATACACATGACAACAAGTCATATTCGACTCCAAAACACTAACCAACCTTCTTCTTGCTTCTCAAAGCTTTCATGGTGTAGCCAAAGTCCA | 13 (0.000249%) | | | ATACTCAATCATACACATGACATCAAGTCATATTCGACTCCAAAACACTAACCAACCTTCTTCTTGCTTCTCAAAGCTTTCATGGTGTAGCCAAAGTCCA | 22 (0.000422%) | | | ATAGGCCACGCTTTCACGGTTCGTATTCGTACTGAAAATCAGAATCAAACGAGCTTTTACCCTTTTGTTCCACACGAGATTTCTGTTCTCGTTGAGCTCA | 79 (0.001514%) | | | ATAGTCAGAGGTGAAATTCTTGGATTTATGAAAGACGAACAACTGCGAAAGCATTTGCCAAGGATGTTTTCATTAATCAAGAACGAAAGTTGGGGGCTCG | 13 (0.000249%) | | | ATAGTGGCCTACCATGGTGGTAACGGGTGACGGAGAATTAGGGTTCGATTCCGGAGAGGGAGCCTGAGAAACGGCTACCACATCCAAGGAAGGCAGCAGG | 16 (0.000307%) | | | ATAGTTTGTTTGATGGTAACTACTACTCGGATAACCGTAGTAATTCTAGAGCTAATACGTGCAACAAACCCCGACTTATGGAAGGGACGCATTTATTAGA | 3 (0.000057%) | | | ATATGACTTGATGTCATGTGTATGATTGAGTATAAGAACT | 118 (0.000905%) | | | ATATGCTTGTCTCAAAGATTAAGCCATGCATGTGTAAGTATGAACGAATTCAGACTGTGAAACTGCGAATGGCTCATTAAATCAGTTATAGTTTGTTTGA | 59 (0.001131%) | | | ATATGGACTTTGGCTACACCATGAAAGCTTTGAGAAGCAAGAAGAAGGTTGGTTAGTGTTTTGGAGTCGAATATGACTTGATGTCATGTGTATGATTGAG | 19 (0.000364%) | | | ATATTCGACTCCAAAACACTAACCAACCTTCTTCTTGCTTCTCAAAGCTTTCATGGTGTAGCCAAAGTCCATATGAGTCTTTGGCTTTGTGTCTTCTAAC | 145 (0.002779%) | | | ATCAAACGAGCTTTTACCCTTTTGTTCCACACGAGATTTCTGTTCTCGTTGAGCTCATCTTAGGACACCTGCGTTATCTTTTAACAGATGTGCCGCCCCA | 5 (0.000096%) | | | ATCAAGTCATATTCGACTCCAAAACACTAACCAACCTTCTTCTTGCTTCTCAAAGCTTTCATGGTGTAGCCAAAGTCCATATGAGTCTTTGGCTTTGTGT | 1466 (0.028095%) | | | ATCAAGTTCACCACCGCATGTCGGTACGCTCCAGGCGTCCTTGGCTCGGATTTAGGCCAACCGCGTGCGGTAACACACGGGAGACCAGCTTCCGTCCCGC | 76 (0.001457%) | | | ATCAGAATCAAACGAGCTTTTACCCTTTTGTTCCACACGAGATTTCTGTTCTCGTTGAGCTCATCTTAGGACACCTGCGTTATCTTTTAACAGATGTGCC | 10 (0.000192%) | | | ATCAGAGCAACGGGCAGAGCCCGCGTCGACCTTTTATCTAATAAATGCGTCCCTTCCATAAGTCGGGGTTTGTTGCACGTATTAGCTCTAGAATTACTAC | 26 (0.000498%) | | | ATCAGCGTGCTCCGGGCGTCGGCCTGTGGGCTCCCCATTCGACCCGTCTTGAAACACGGACCAAGGAGTCTGACATGTGTGCGAGTCAACGGGTGAGTAA | 6 (0.000115%) | | | ATCAGTTATAGTTTGTTTGATGGTAACTACTACTCGGATAACCGTAGTAATTCTAGAGCTAATACGTGCAACAAACCCCGACTTATGGAAGGGACGCATT | 8 (0.000153%) | | | ATCATAAGGATACTAAATCCTATTTTCTGGTAAATTTTCATAATTTTTTGACACCTCTAGCTAGGTCATTTGACCTGATACAACATCGGATTTTCATGGT | 4 (0.000077%) | | | ATCATACACATGACAACAAGTCATATTCGACTCCAAAACACTAACCAACCTTCTTCTTGCTTCTCAAAGCTTTCATGGTGTAGCCAAAGTCCATATGAGT | 3 (0.000057%) | | | ATCATACACATGACATCAAGTCATATTCGACTCCAAAACACTAACCAACCTTCTTCTTGCTTCTCAAAGCTTTCATGGTGTAGCCAAAGTCCATATGAGT | 35 (0.000671%) | | | ATCATCAGAGCAACGGGCAGAGCCCGCGTCGACCTTTTATCTAATAAATGCGTCCCTTCCATAAGTCGGGGTTTGTTGCACGTATTAGCTCTAGAATTAC | 14 (0.000268%) | | | ATCATGAATCATCAGAGCAACGGGCAGAGCCCGCGTCGACCTTTTATCTAATAAATGCGTCCCTTCCATAAGTCGGGGTTTGTTGCACGTATTAGCTCTA | 61 (0.001169%) | | | ATCATTCAATCGGTAGGAGCGACGGGCGGTGTGTACAAAGGGCAGGGACGTAGTCAACGCGAGCTGATGACTCGCGCTTACTAGGAATTCCTCGTTGAAG | 15 (0.000287%) | | | ATCCAACCTAGGCGAGACAAGGGTTCACATTTCGTTCATCACCCTTGGCCGGCTATCGAACAGCCGGACTCCCATCAAAAGATGGTTGCCAAGAACATCT | 21 (0.000402%) | | | ATCCAACCTAGGCGAGACAAGGGTTCACATTTCGTTCATCACCCTTGGCCGGCTTTCGAACAGCCGGACTCCCATCAAAAGATGGTTGCCAAGAACATCT | 27 (0.000517%) | | | ATCCAGAGCGTAGGCTTGCTTTGAGCACTCTAATTTCTTCAAAGTAACAGCGCCGGAGGCACGACCCGGCCAATTAAGACCAGGAGCGTATCGCCGACCG | 12 (0.000230%) | | | ATCCATGCTTTCCAACGAAGCACGCCCATCCAACCTAGGCGAGACAAGGGTTCACATTTCGTTCATCACCCTTGGCCGGCTATCGAACAGCCGGACTCCC | 5 (0.000096%) | | | ATCCATTGGAGGGCAAGTCTGGTGCCAGCAGCCGCGGTAATTCCAGCTCCAATAGCGTATATTTAAGTTGTTGCAGTTAAAAAGCTCGTAGTTGAACCTT | 20 (0.000383%) | | | ATCCCATGCTAATGTATCCAGAGCGTAGGCTTGCTTTGAGCACTCTAATTTCTTCAAAGTAACAGCGCCGGAGGCACGACCCGGCCAATTAAGACCAGGA | 19 (0.000364%) | | | ATCCGCCTAGGCTGTCCCGAGTGTGAGCGAGGTGTGAGTGTCGCCCATGGGCATCGACACCTTGCGGCTAGGAACTGGAACGAGACGGGTAGCAAAGATT | 90 (0.001725%) | | | ATCCGCCTAGGCTGTCCCGAGTGTGAGCGAGGTGTGAGTGTCGCCCATGGGCATCGACACCTTGCGGCTAGGAACTGGAACGAGACGGGTGGCAAAGATT | 100 (0.001916%) | | | ATCCGCTAAGGAGTGTGTAACAACTCACCTGCCGAATCAACTAGCCCCGAAAATGGATGGCGCTTAAGCGCGCGACCTATACCCGGCCGTCGGGGCAAGA | 4 (0.000077%) | | | ATCCGGTTAAAATTCCGGAACCGGGACGTGGCGGTTGACGGCAACGTTAGGGAGTCCGGAGACGTCGGCGGGGGCCTCGGGAAGAGTTATCTTTTCTGTT | 19 (0.000364%) | | | ATCCGTAACTTCGGGAAAAGGATTGGCTCTGAGGGCTGGGCTCGGGGGTCCCAGTTCCGAACCCGTCGGCTGTCAGCGGACTGCTCGAGCTGCTTCCGCG | 3 (0.000057%) | | | ATCCGTCGAGTTATCATGAATCATCAGAGCAACGGGCAGAGCCCGCGTCGACCTTTTATCTAATAAATGCGTCCCTTCCATAAGTCGGGGTTTGTTGCAC | 4 (0.000077%) | | | ATCCTATGATGTTATCCCATGCTAATGTATCCAGAGCGTAGGCTTGCTTTGAGCACTCTAATTTCTTCAAAGTAACAGCGCCGGAGGCACGACCCGGCCA | 52 (0.000997%) | | | ATCCTATTTTCTGGTAAATTTTCATAATTTTTTGACACCTCTAGCTAGGTCATTTGACCTGATACAACATCGGATTTTCATGGTCTAGTTGGGGCTCCGT | 3 (0.000057%) | | | ATCCTCGTTAAGGGATTTAGATTGTACTCATTCCAATTACCAGACTCGAAAGAGCCCGGTATTGTTATTTATTGTCACTACCTCCCCGTGTCAGGATTGG | 17 (0.000326%) | | | ATCCTTGTTAGAAGACACAAAGCCAAAGACTCATATGGACTTTGGCTACACCATGAAAGCTTTGAGAAGCAAGAAGAAGGTTGGTTAGTGTTTTGGAGTC | 10 (0.000192%) | | | ATCCTTGTTAGTTTCTTTTCCTCCGCTTATTGATATGCTTAAACTCAGCGGGTAATCCCGCCTGACCTGGGGTCGCTATATGGACTTTGGGTCATCTACA | 20 (0.000383%) | | | ATCCTTTGCTGATGCGGGACGGAAGCTGGTCTCCCGTGTGTTACCGCACGCGGTTGGCCTAAATCCGAGCCAAGGACGCCTGGAGCGTACCGACATGCGG | 8 (0.000153%) | | | ATCGAAATCCTATGATGTTATCCCATGCTAATGTATCCAGAGCGTAGGCTTGCTTTGAGCACTCTAATTTCTTCAAAGTAACAGCGCCGGAGGCACGACC | 11 (0.000211%) | | | ATCGACACCTTGCGGCTAGGAACTGGAACGAGACGGGTGGCAAAGATTTCGAGTAGCACTTCATACTACCGTGGGTTTTTTAAACCTTCCGAGTTTTGTT | 10 (0.000192%) | | | ATCGACCAGAGGCTGTTCACCTTGGAGACCTGATGCGGTTATGAGTACGACCGGGCGTGAGCGGCACTCGGTCCTCCGGATTTTCAAGGGCCGCCGGGGG | 24 (0.000460%) | | | ATCGATCAAGTTCACCACCGCATGTCGGTACGCTCCAGGCGTCCTTGGCTCGGATTTAGGCCAACCGCGTGCGGTAACACACGGGAGACCAGCTTCCGTC | 7 (0.000134%) | | | ATCGATCACGGCAATTCCCCGCCACATCCTCTCAAACGCAATGGAAAGAGAGAAAGGACGAGGTCTTGACCGTCATCTTTTGCCCGAAGGACGGATGAGC | 105 (0.002012%) | | | ATCGATCCATGCTTTCCAACGAAGCACGCCCATCCAACCTAGGCGAGACAAGGGTTCACATTTCGTTCATCACCCTTGGCCGGCTATCGAACAGCCGGAC | 25 (0.000479%) | | | ATCGATCCGCCTAGGCTGTCCCGAGTGTGAGCGAGGTGTGAGTGTCGCCCATGGGCATCGACACCTTGCGGCTAGGAACTGGAACGAGACGGGTAGCAAA | 13 (0.000249%) | | | ATCGATCCGCCTAGGCTGTCCCGAGTGTGAGCGAGGTGTGAGTGTCGCCCATGGGCATCGACACCTTGCGGCTAGGAACTGGAACGAGACGGGTGGCAAA | 15 (0.000287%) | | | ATCGATCCGGTTAAAATTCCGGAACCGGGACGTGGCGGTTGACGGCAACGTTAGGGAGTCCGGAGACGTCGGCGGGGGCCTCGGGAAGAGTTATCTTTTC | 15 (0.000287%) | | | ATCGATCCGTAACTTCGGGAAAAGGATTGGCTCTGAGGGCTGGGCTCGGGGGTCCCAGTTCCGAACCCGTCGGCTGTCAGCGGACTGCTCGAGCTGCTTC | 33 (0.000632%) | | | ATCGATCCGTCGAGTTATCATGAATCATCAGAGCAACGGGCAGAGCCCGCGTCGACCTTTTATCTAATAAATGCGTCCCTTCCATAAGTCGGGGTTTGTT | 17 (0.000326%) | | | ATCGATCCTCGTTAAGGGATTTAGATTGTACTCATTCCAATTACCAGACTCGAAAGAGCCCGGTATTGTTATTTATTGTCACTACCTCCCCGTGTCAGGA | 41 (0.000786%) | | | ATCGATCGAAATCCTATGATGTTATCCCATGCTAATGTATCCAGAGCGTAGGCTTGCTTTGAGCACTCTAATTTCTTCAAAGTAACAGCGCCGGAGGCAC | 21 (0.000402%) | | | ATCGATCGGCGGACCGGATTGCTCCGTTCCGCATCCGACCAGGACGCATCGCCGGCCCCCATCCGCTTCCCTCCCGACAATTTCAAGCACTCTTTGACTC | 14 (0.000268%) | | | ATCGATCGGGTTGCGGTTTAAGTTGTTATACTCAATCATA | 19 (0.000146%) | | | ATCGGGTTGCGGTTTAAGTTGTTATACTCAATCATACACA | 120 (0.000920%) | | | ATCGGTAGGAGCGACGGGCGGTGTGTACAAAGGGCAGGGACGTAGTCAACGCGAGCTGATGACTCGCGCTTACTAGGAATTCCTCGTTGAAGACCAACAA | 64 (0.001227%) | | | ATCTAATAAATGCGTCCCTTCCATAAGTCGGGGTTTGTTGCACGTATTAGCTCTAGAATTACTACGGTTATCCGAGTAGTAGTTACCATCAAACAAACTA | 7 (0.000134%) | | | ATCTTAAAAGCCTAAGTAGTGTTTCCTTGTTAGAAGACACAAAGCCAAAGACTCATATGGACTTTGGCTACACCATGAAAGCTTTGAGAAGCAAGAAGAA | 610 (0.011690%) | | | ATCTTAAAGGCGTAAGAATTGTATCCTTGTTAAAAGACACAAAGCCAAAGACTCATATGGACTTTGGCTACACCATGAAAGCTTTGAGAAGCAAGAAGAA | 38 (0.000728%) | | | ATCTTAAAGGCGTAAGAATTGTATCCTTGTTAGAAGACACAAAGCCAAAGACTCATATGGACTTTGGCTACACCATGAAAGCTTTGAGAAGCAAGAAGAA | 27 (0.000517%) | | | ATGAATCATCAGAGCAACGGGCAGAGCCCGCGTCGACCTTTTATCTAATAAATGCGTCCCTTCCATAAGTCGGGGTTTGTTGCACGTATTAGCTCTAGAA | 86 (0.001648%) | | | ATGACATCAAGTCATATTCGACTCCAAAACACTAACCAACCTTCTTCTTGCTTCTCAAAGCTTTCATGGTGTAGCCAAAGTCCATATGAGTCTTTGGCTT | 110 (0.002108%) | | | ATGACTTGATGTCATGTGTATGATTGAGTATAAGAACTTA | 268 (0.002054%) | | | ATGATGTTATCCCATGCTAATGTATCCAGAGCGTAGGCTTGCTTTGAGCACTCTAATTTCTTCAAAGTAACAGCGCCGGAGGCACGACCCGGCCAATTAA | 65 (0.001246%) | | | ATGATTAACAGGGACAGTCGGGGGCATTCGTATTTCATAGTCAGAGGTGAAATTCTTGGATTTATGAAAGACGAACAACTGCGAAAGCATTTGCCAAGGA | 112 (0.002146%) | | | ATGCATCATAAGGATACTAAATCCTATTTTCTGGTAAATTTTCATAATTTTTTGACACCTCTAGCTAGGTCATTTGACCTGATACAACATCGGATTTTCA | 3 (0.000057%) | | | ATGCATGTGTAAGTATGAACGAATTCAGACTGTGAAACTGCGAATGGCTCATTAAATCAGTTATAGTTTGTTTGATGGTAACTACTACTCGGATAACCGT | 4 (0.000077%) | | | ATGCCAGCCGTTCGTTTGCATGTTCCTTGACACTTTTCGTGCCGGGGTTTTGTGATATCCGGAAGCAACGCGCACGACAAGACCGAGATAAAAGCTCCCG | 8 (0.000153%) | | | ATGCCCTTAGATGTTCTGGGCCGCACGCGCGCTACACTGATGTATTCAACGAGTTCACACCTTGGCCGACAGGCCCGGGTAATCTTTGAAATTTCATCGT | 10 (0.000192%) | | | ATGCTAATGTATCCAGAGCGTAGGCTTGCTTTGAGCACTCTAATTTCTTCAAAGTAACAGCGCCGGAGGCACGACCCGGCCAATTAAGACCAGGAGCGTA | 6 (0.000115%) | | | ATGCTTGTCTCAAAGATTAAGCCATGCATGTGTAAGTATGAACGAATTCAGACTGTGAAACTGCGAATGGCTCATTAAATCAGTTATAGTTTGTTTGATG | 20 (0.000383%) | | | ATGCTTTCCAACGAAGCACGCCCATCCAACCTAGGCGAGACAAGGGTTCACATTTCGTTCATCACCCTTGGCCGGCTATCGAACAGCCGGACTCCCATCA | 290 (0.005558%) | | | ATGGACTTTGGCTACACCATGAAAGCTTTGAGAAGCAAGAAGAAGGTTGGTTAGTGTTTTGGAGTCGAATATGACTTGATGTCATGTGTATGATTGAGTA | 74 (0.001418%) | | | ATGGCTCATTAAATCAGTTATAGTTTGTTTGATGGTAACTACTACTCGGATAACCGTAGTAATTCTAGAGCTAATACGTGCAACAAACCCCGACTTATGG | 3 (0.000057%) | | | ATGGGCATCGACACCTTGCGGCTAGGAACTGGAACGAGACGGGTGGCAAAGATTTCGAGTAGCACTTCATACTACCGTGGGTTTTTTAAACCTTCCGAGT | 5 (0.000096%) | | | ATGTATCCAGAGCGTAGGCTTGCTTTGAGCACTCTAATTTCTTCAAAGTAACAGCGCCGGAGGCACGACCCGGCCAATTAAGACCAGGAGCGTATCGCCG | 4 (0.000077%) | | | ATGTCATGTGTATGATTGAGTATAAGAACTTAAACCGCAA | 135 (0.001035%) | | | ATGTCGGTACGCTCCAGGCGTCCTTGGCTCGGATTTAGGCCAACCGCGTGCGGTAACACACGGGAGACCAGCTTCCGTCCCGCATCAGCAAAGGATGGTG | 7 (0.000134%) | | | ATGTGTAAGTATGAACGAATTCAGACTGTGAAACTGCGAATGGCTCATTAAATCAGTTATAGTTTGTTTGATGGTAACTACTACTCGGATAACCGTAGTA | 6 (0.000115%) | | | ATGTTATCCCATGCTAATGTATCCAGAGCGTAGGCTTGCTTTGAGCACTCTAATTTCTTCAAAGTAACAGCGCCGGAGGCACGACCCGGCCAATTAAGAC | 17 (0.000326%) | | | ATGTTCTGGGCCGCACGCGCGCTACACTGATGTATTCAACGAGTTCACACCTTGGCCGACAGGCCCGGGTAATCTTTGAAATTTCATCGTGATGGGGATA | 7 (0.000134%) | | | ATTAAATCAGTTATAGTTTGTTTGATGGTAACTACTACTCGGATAACCGTAGTAATTCTAGAGCTAATACGTGCAACAAACCCCGACTTATGGAAGGGAC | 33 (0.000632%) | | | ATTAACAGGGACAGTCGGGGGCATTCGTATTTCATAGTCAGAGGTGAAATTCTTGGATTTATGAAAGACGAACAACTGCGAAAGCATTTGCCAAGGATGT | 110 (0.002108%) | | | ATTAAGCCATGCATGTGTAAGTATGAACGAATTCAGACTGTGAAACTGCGAATGGCTCATTAAATCAGTTATAGTTTGTTTGATGGTAACTACTACTCGG | 18 (0.000345%) | | | ATTACCAGACTCGAAAGAGCCCGGTATTGTTATTTATTGTCACTACCTCCCCGTGTCAGGATTGGGTAATTTGCGCGCCTGCTGCCTTCCTTGGATGTGG | 7 (0.000134%) | | | ATTAGGGTTCGATTCCGGAGAGGGAGCCTGAGAAACGGCTACCACATCCAAGGAAGGCAGCAGGCGCGCAAATTACCCAATCCTGACACGGGGAGGTAGT | 5 (0.000096%) | | | ATTCAATCGGTAGGAGCGACGGGCGGTGTGTACAAAGGGCAGGGACGTAGTCAACGCGAGCTGATGACTCGCGCTTACTAGGAATTCCTCGTTGAAGACC | 318 (0.006094%) | | | ATTCCAATTACCAGACTCGAAAGAGCCCGGTATTGTTATTTATTGTCACTACCTCCCCGTGTCAGGATTGGGTAATTTGCGCGCCTGCTGCCTTCCTTGG | 22 (0.000422%) | | | ATTCCAGCTCCAATAGCGTATATTTAAGTTGTTGCAGTTAAAAAGCTCGTAGTTGAACCTTGGGATGGGTCGGCCGGTCCGCCTTTGGTGTGCATTGGTC | 12 (0.000230%) | | | ATTCCCCGCCACATCCTCTCAAACGCAATGGAAAGAGAGAAAGGACGAGGTCTTGACCGTCATCTTTTGCCCGAAGGACGGATGAGCTTTGGCGGGACTG | 16 (0.000307%) | | | ATTCGACTCCAAAACACTAACCAACCTTCTTCTTGCTTCTCAAAGCTTTCATGGTGTAGCCAAAGTCCATATGAGTCTTTGGCTTTGTGTCTTCTAACAA | 126 (0.002415%) | | | ATTCGTACTGAAAATCAGAATCAAACGAGCTTTTACCCTTTTGTTCCACACGAGATTTCTGTTCTCGTTGAGCTCATCTTAGGACACCTGCGTTATCTTT | 5 (0.000096%) | | | ATTCGTATTTCATAGTCAGAGGTGAAATTCTTGGATTTATGAAAGACGAACAACTGCGAAAGCATTTGCCAAGGATGTTTTCATTAATCAAGAACGAAAG | 9 (0.000172%) | | | ATTGGAGGGCAAGTCTGGTGCCAGCAGCCGCGGTAATTCCAGCTCCAATAGCGTATATTTAAGTTGTTGCAGTTAAAAAGCTCGTAGTTGAACCTTGGGA | 205 (0.003929%) | | | ATTGTACTCATTCCAATTACCAGACTCGAAAGAGCCCGGTATTGTTATTTATTGTCACTACCTCCCCGTGTCAGGATTGGGTAATTTGCGCGCCTGCTGC | 17 (0.000326%) | | | ATTGTATCCTTGTTAGAAGACACAAAGCCAAAGACTCATATGGACTTTGGCTACACCATGAAAGCTTTGAGAAGCAAGAAGAAGGTTGGTTAGTGTTTTG | 38 (0.000728%) | | | ATTGTCAGGTGGGGAGTTTGGCTGGGGCGGCACATCTGTTAAAAGATAACGCAGGTGTCCTAAGATGAGCTCAACGAGAACAGAAATCTCGTGTGGAACA | 27 (0.000517%) | | | ATTGTTCCATCGACCAGAGGCTGTTCACCTTGGAGACCTGATGCGGTTATGAGTACGACCGGGCGTGAGCGGCACTCGGTCCTCCGGATTTTCAAGGGCC | 13 (0.000249%) | | | ATTGTTGGTCTTCAACGAGGAATTCCTAGTAAGCGCGAGTCATCAGCTCGCGTTGACTACGTCCCTGCCCTTTGTACACACCGCCCGTCGCTCCTACCGA | 27 (0.000517%) | | | ATTTAGATTGTACTCATTCCAATTACCAGACTCGAAAGAGCCCGGTATTGTTATTTATTGTCACTACCTCCCCGTGTCAGGATTGGGTAATTTGCGCGCC | 9 (0.000172%) | | | ATTTCATAGTCAGAGGTGAAATTCTTGGATTTATGAAAGACGAACAACTGCGAAAGCATTTGCCAAGGATGTTTTCATTAATCAAGAACGAAAGTTGGGG | 9 (0.000172%) | | | ATTTCGTTCATCACCCTTGGCCGGCTATCGAACAGCCGGACTCCCATCAAAAGATGGTTGCCAAGAACATCTTCGTTACGGTTTGCTAATTCTCGGAATA | 18 (0.000345%) | | | ATTTTCATAATTTTTTGACACCTCTAGCTAGGTCATTTGACCTGATACAACATCGGATTTTCATGGTCTAGTTGGGGCTCCGTGGGCATATTTGATGCAA | 5 (0.000096%) | | | ATTTTCTGGTAAATTTTCATAATTTTTTGACACCTCTAGCTAGGTCATTTGACCTGATACAACATCGGATTTTCATGGTCTAGTTGGGGCTCCGTGGGCA | 10 (0.000192%) | | | CAAAAAGCAACGTCGCTATGAACGCTTGGCTGCCACAAGCCAGTTATCCCTGTGGTAACTTTTCTGACACCTCTAGCTTCAAATTCCGAAGGTCTAAAGG | 1216 (0.023304%) | | | CAAACGAGCTTTTACCCTTTTGTTCCACACGAGATTTCTGTTCTCGTTGAGCTCATCTTAGGACACCTGCGTTATCTTTTAACAGATGTGCCGCCCCAGC | 5 (0.000096%) | | | CAAAGACTCATATGGACTTTGGCTACACCATGAAAGCTTTGAGAAGCAAGAAGAAGGTTGGTTAGTGTTTTGGAGTCGAATATGACTTGATGTCATGTGT | 117 (0.002242%) | | | CAAAGATTAAGCCATGCATGTGTAAGTATGAACGAATTCAGACTGTGAAACTGCGAATGGCTCATTAAATCAGTTATAGTTTGTTTGATGGTAACTACTA | 54 (0.001035%) | | | CAAAGCCAAAGACTCATATGGACTTTGGCTACACCATGAAAGCTTTGAGAAGCAAGAAGAAGGTTGGTTAGTGTTTTGGAGTCGAATATGACTTGATGTC | 109 (0.002089%) | | | CAAAGCTTTCATGGTGTAGCCAAAGTCCATATGAGTCTTTGGCTTTGTGTCTTCTAACAAGGAAACACTACTTAGGCTTATAAGATGCGGTTGCGGTTTA | 5 (0.000096%) | | | CAACCGGATCTTAAAGGCGTAAGAATTGTATCCTTGTTAGAAGACACAAAGCCAAAGACTCATATGGACTTTGGCTACACCATGAAAGCTTTGAGAAGCA | 4 (0.000077%) | | | CAACCTAGGCGAGACAAGGGTTCACATTTCGTTCATCACCCTTGGCCGGCTATCGAACAGCCGGACTCCCATCAAAAGATGGTTGCCAAGAACATCTTCG | 5 (0.000096%) | | | CAACCTAGGCGAGACAAGGGTTCACATTTCGTTCATCACCCTTGGCCGGCTTTCGAACAGCCGGACTCCCATCAAAAGATGGTTGCCAAGAACATCTTCG | 10 (0.000192%) | | | CAACGAAGCACGCCCATCCAACCTAGGCGAGACAAGGGTTCACATTTCGTTCATCACCCTTGGCCGGCTATCGAACAGCCGGACTCCCATCAAAAGATGG | 69 (0.001322%) | | | CAACGAAGCACGCCCATCCAACCTAGGCGAGACAAGGGTTCACATTTCGTTCATCACCCTTGGCCGGCTTTCGAACAGCCGGACTCCCATCAAAAGATGG | 75 (0.001437%) | | | CAACGAGGAATTCCTAGTAAGCGCGAGTCATCAGCTCGCGTTGACTACGTCCCTGCCCTTTGTACACACCGCCCGTCGCTCCTACCGATTGAATGATCGA | 7 (0.000134%) | | | CAACGGGCAGAGCCCGCGTCGACCTTTTATCTAATAAATGCGTCCCTTCCATAAGTCGGGGTTTGTTGCACGTATTAGCTCTAGAATTACTACGGTTATC | 31 (0.000594%) | | | CAACGTCGCTATGAACGCTTGGCTGCCACAAGCCAGTTATCCCTGTGGTAACTTTTCTGACACCTCTAGCTTCAAATTCCGAAGGTCTAAAGGATCGATC | 32 (0.000613%) | | | CAAGGGTTCACATTTCGTTCATCACCCTTGGCCGGCTATCGAACAGCCGGACTCCCATCAAAAGATGGTTGCCAAGAACATCTTCGTTACGGTTTGCTAA | 21 (0.000402%) | | | CAAGGGTTCACATTTCGTTCATCACCCTTGGCCGGCTTTCGAACAGCCGGACTCCCATCAAAAGATGGTTGCCAAGAACATCTTCGTTACGGTTTGCTAA | 35 (0.000671%) | | | CAAGTCATATTCGACTCCAAAACACTAACCAACCTTCTTCTTGCTTCTCAAAGCTTTCATGGTGTAGCCAAAGTCCATATGAGTCTTTGGCTTTGTGTCT | 123 (0.002357%) | | | CAAGTCTGGTGCCAGCAGCCGCGGTAATTCCAGCTCCAATAGCGTATATTTAAGTTGTTGCAGTTAAAAAGCTCGTAGTTGAACCTTGGGATGGGTCGGC | 21 (0.000402%) | | | CAAGTTCACCACCGCATGTCGGTACGCTCCAGGCGTCCTTGGCTCGGATTTAGGCCAACCGCGTGCGGTAACACACGGGAGACCAGCTTCCGTCCCGCAT | 19 (0.000364%) | | | CAATAACAGGTCTGTGATGCCCTTAGATGTTCTGGGCCGCACGCGCGCTACACTGATGTATTCAACGAGTTCACACCTTGGCCGACAGGCCCGGGTAATC | 61 (0.001169%) | | | CAATCATACACATGACAACAAGTCATATTCGACTCCAAAACACTAACCAACCTTCTTCTTGCTTCTCAAAGCTTTCATGGTGTAGCCAAAGTCCATATGA | 34 (0.000652%) | | | CAATCATACACATGACATCAAGTCATATTCGACTCCAAAACACTAACCAACCTTCTTCTTGCTTCTCAAAGCTTTCATGGTGTAGCCAAAGTCCATATGA | 163 (0.003124%) | | | CAATCGGTAGGAGCGACGGGCGGTGTGTACAAAGGGCAGGGACGTAGTCAACGCGAGCTGATGACTCGCGCTTACTAGGAATTCCTCGTTGAAGACCAAC | 88 (0.001686%) | | | CAATTACCAGACTCGAAAGAGCCCGGTATTGTTATTTATTGTCACTACCTCCCCGTGTCAGGATTGGGTAATTTGCGCGCCTGCTGCCTTCCTTGGATGT | 19 (0.000364%) | | | CAATTCCCCGCCACATCCTCTCAAACGCAATGGAAAGAGAGAAAGGACGAGGTCTTGACCGTCATCTTTTGCCCGAAGGACGGATGAGCTTTGGCGGGAC | 27 (0.000517%) | | | CACAAAGCCAAAGACTCATATGGACTTTGGCTACACCATGAAAGCTTTGAGAAGCAAGAAGAAGGTTGGTTAGTGTTTTGGAGTCGAATATGACTTGATC | 9 (0.000172%) | | | CACAAAGCCAAAGACTCATATGGACTTTGGCTACACCATGAAAGCTTTGAGAAGCAAGAAGAAGGTTGGTTAGTGTTTTGGAGTCGAATATGACTTGATG | 36 (0.000690%) | | | CACATCCTCTCAAACGCAATGGAAAGAGAGAAAGGACGAGGTCTTGACCGTCATCTTTTGCCCGAAGGACGGATGAGCTTTGGCGGGACTGAATCACTTC | 4 (0.000077%) | | | CACATCTGTTAAAAGATAACGCAGGTGTCCTAAGATGAGCTCAACGAGAACAGAAATCTCGTGTGGAACAAAAGGGTAAAAGCTCGTTTGATTCTGATTT | 9 (0.000172%) | | | CACATGACATCAAGTCATATTCGACTCCAAAACACTAACCAACCTTCTTCTTGCTTCTCAAAGCTTTCATGGTGTAGCCAAAGTCCATATGAGTCTTTGG | 41 (0.000786%) | | | CACATTTCGTTCATCACCCTTGGCCGGCTATCGAACAGCCGGACTCCCATCAAAAGATGGTTGCCAAGAACATCTTCGTTACGGTTTGCTAATTCTCGGA | 5 (0.000096%) | | | CACATTTCGTTCATCACCCTTGGCCGGCTTTCGAACAGCCGGACTCCCATCAAAAGATGGTTGCCAAGAACATCTTCGTTACGGTTTGCTAATTCTCGGA | 11 (0.000211%) | | | CACCACCGCATGTCGGTACGCTCCAGGCGTCCTTGGCTCGGATTTAGGCCAACCGCGTGCGGTAACACACGGGAGACCAGCTTCCGTCCCGCATCAGCAA | 21 (0.000402%) | | | CACCATCCTTTGCTGATGCGGGACGGAAGCTGGTCTCCCGTGTGTTACCGCACGCGGTTGGCCTAAATCCGAGCCAAGGACGCCTGGAGCGTACCGACAT | 7 (0.000134%) | | | CACCCGTTGACTCGCACACATGTCAGACTCCTTGGTCCGTGTTTCAAGACGGGTCGAATGGGGAGCCCACAGGCCGACGCCCGGAGCACGCTGATGCCGA | 6 (0.000115%) | | | CACCCTAACGCCTCGAAGAACTAATGGCAGCCCACGCAAGGCAAGCCCATTCTCCTCGACGATTCAGCAGTTTTTGTCCGAGAACTGCTGAGAAAACTCG | 11 (0.000211%) | | | CACCGCATGTCGGTACGCTCCAGGCGTCCTTGGCTCGGATTTAGGCCAACCGCGTGCGGTAACACACGGGAGACCAGCTTCCGTCCCGCATCAGCAAAGG | 7 (0.000134%) | | | CACGCCCATCCAACCTAGGCGAGACAAGGGTTCACATTTCGTTCATCACCCTTGGCCGGCTATCGAACAGCCGGACTCCCATCAAAAGATGGTTGCCAAG | 26 (0.000498%) | | | CACGCCCATCCAACCTAGGCGAGACAAGGGTTCACATTTCGTTCATCACCCTTGGCCGGCTTTCGAACAGCCGGACTCCCATCAAAAGATGGTTGCCAAG | 33 (0.000632%) | | | CACGCGCCTAACGGCGTGCCTCGGCATCAGCGTGCTCCGGGCGTCGGCCTGTGGGCTCCCCATTCGACCCGTCTTGAAACACGGACCAAGGAGTCTGACA | 76 (0.001457%) | | | CACGCTTTCACGGTTCGTATTCGTACTGAAAATCAGAATCAAACGAGCTTTTACCCTTTTGTTCCACACGAGATTTCTGTTCTCGTTGAGCTCATCTTAG | 111 (0.002127%) | | | CACGGCAATTCCCCGCCACATCCTCTCAAACGCAATGGAAAGAGAGAAAGGACGAGGTCTTGACCGTCATCTTTTGCCCGAAGGACGGATGAGCTTTGGC | 7 (0.000134%) | | | CACGGTTCGTATTCGTACTGAAAATCAGAATCAAACGAGCTTTTACCCTTTTGTTCCACACGAGATTTCTGTTCTCGTTGAGCTCATCTTAGGACACCTG | 10 (0.000192%) | | | CAGAATCAAACGAGCTTTTACCCTTTTGTTCCACACGAGATTTCTGTTCTCGTTGAGCTCATCTTAGGACACCTGCGTTATCTTTTAACAGATGTGCCGC | 18 (0.000345%) | | | CAGACTCGAAAGAGCCCGGTATTGTTATTTATTGTCACTACCTCCCCGTGTCAGGATTGGGTAATTTGCGCGCCTGCTGCCTTCCTTGGATGTGGTAGCC | 19 (0.000364%) | | | CAGAGCAACGGGCAGAGCCCGCGTCGACCTTTTATCTAATAAATGCGTCCCTTCCATAAGTCGGGGTTTGTTGCACGTATTAGCTCTAGAATTACTACGG | 16 (0.000307%) | | | CAGAGCCCGCGTCGACCTTTTATCTAATAAATGCGTCCCTTCCATAAGTCGGGGTTTGTTGCACGTATTAGCTCTAGAATTACTACGGTTATCCGAGTAG | 7 (0.000134%) | | | CAGAGCGTAGGCTTGCTTTGAGCACTCTAATTTCTTCAAAGTAACAGCGCCGGAGGCACGACCCGGCCAATTAAGACCAGGAGCGTATCGCCGACCGAAG | 39 (0.000747%) | | | CAGAGGCTGTTCACCTTGGAGACCTGATGCGGTTATGAGTACGACCGGGCGTGAGCGGCACTCGGTCCTCCGGATTTTCAAGGGCCGCCGGGGGCGCACC | 12 (0.000230%) | | | CAGAGGTGAAATTCTTGGATTTATGAAAGACGAACAACTGCGAAAGCATTTGCCAAGGATGTTTTCATTAATCAAGAACGAAAGTTGGGGGCTCGAAGAC | 18 (0.000345%) | | | CAGCACGCGCCTAACGGCGTGCCTCGGCATCAGCGTGCTCCGGGCGTCGGCCTGTGGGCTCCCCATTCGACCCGTCTTGAAACACGGACCAAGGAGTCTG | 75 (0.001437%) | | | CAGCAGCCGCGGTAATTCCAGCTCCAATAGCGTATATTTAAGTTGTTGCAGTTAAAAAGCTCGTAGTTGAACCTTGGGATGGGTCGGCCGGTCCGCCTTT | 27 (0.000517%) | | | CAGCCGCGGTAATTCCAGCTCCAATAGCGTATATTTAAGTTGTTGCAGTTAAAAAGCTCGTAGTTGAACCTTGGGATGGGTCGGCCGGTCCGCCTTTGGT | 10 (0.000192%) | | | CAGCGTGCTCCGGGCGTCGGCCTGTGGGCTCCCCATTCGACCCGTCTTGAAACACGGACCAAGGAGTCTGACATGTGTGCGAGTCAACGGGTGAGTAAAC | 9 (0.000172%) | | | CAGGGACAGTCGGGGGCATTCGTATTTCATAGTCAGAGGTGAAATTCTTGGATTTATGAAAGACGAACAACTGCGAAAGCATTTGCCAAGGATGTTTTCA | 29 (0.000556%) | | | CAGGTCTGTGATGCCCTTAGATGTTCTGGGCCGCACGCGCGCTACACTGATGTATTCAACGAGTTCACACCTTGGCCGACAGGCCCGGGTAATCTTTGAA | 11 (0.000211%) | | | CAGGTGGGGAGTTTGGCTGGGGCGGCACATCTGTTAAAAGATAACGCAGGTGTCCTAAGATGAGCTCAACGAGAACAGAAATCTCGTGTGGAACAAAAGG | 36 (0.000690%) | | | CAGTAGTCATATGCTTGTCTCAAAGATTAAGCCATGCATGTGTAAGTATGAACGAATTCAGACTGTGAAACTGCGAATGGCTCATTAAATCAGTTATAGT | 136 (0.002606%) | | | CAGTCGGGGGCATTCGTATTTCATAGTCAGAGGTGAAATTCTTGGATTTATGAAAGACGAACAACTGCGAAAGCATTTGCCAAGGATGTTTTCATTAATC | 4 (0.000077%) | | | CATAAGGATACTAAATCCTATTTTCTGGTAAATTTTCATAATTTTTTGACACCTCTAGCTAGGTCATTTGACCTGATACAACATCGGATTTTCATGGTCT | 24 (0.000460%) | | | CATACACATGACATCAAGTCATATTCGACTCCAAAACACTAACCAACCTTCTTCTTGCTTCTCAAAGCTTTCATGGTGTAGCCAAAGTCCATATGAGTCT | 47 (0.000901%) | | | CATAGTCAGAGGTGAAATTCTTGGATTTATGAAAGACGAACAACTGCGAAAGCATTTGCCAAGGATGTTTTCATTAATCAAGAACGAAAGTTGGGGGCTC | 17 (0.000326%) | | | CATATGCTTGTCTCAAAGATTAAGCCATGCATGTGTAAGTATGAACGAATTCAGACTGTGAAACTGCGAATGGCTCATTAAATCAGTTATAGTTTGTTTG | 52 (0.000997%) | | | CATATGGACTTTGGCTACACCATGAAAGCTTTGAGAAGCAAGAAGAAGGTTGGTTAGTGTTTTGGAGTCGAATATGACTTGATGTCATGTGTATGATTGA | 12 (0.000230%) | | | CATATTCGACTCCAAAACACTAACCAACCTTCTTCTTGCTTCTCAAAGCTTTCATGGTGTAGCCAAAGTCCATATGAGTCTTTGGCTTTGTGTCTTCTAA | 80 (0.001533%) | | | CATCAAGTCATATTCGACTCCAAAACACTAACCAACCTTCTTCTTGCTTCTCAAAGCTTTCATGGTGTAGCCAAAGTCCATATGAGTCTTTGGCTTTGTG | 50 (0.000958%) | | | CATCAGAGCAACGGGCAGAGCCCGCGTCGACCTTTTATCTAATAAATGCGTCCCTTCCATAAGTCGGGGTTTGTTGCACGTATTAGCTCTAGAATTACTA | 44 (0.000843%) | | | CATCAGCGTGCTCCGGGCGTCGGCCTGTGGGCTCCCCATTCGACCCGTCTTGAAACACGGACCAAGGAGTCTGACATGTGTGCGAGTCAACGGGTGAGTA | 10 (0.000192%) | | | CATCCAACCTAGGCGAGACAAGGGTTCACATTTCGTTCATCACCCTTGGCCGGCTATCGAACAGCCGGACTCCCATCAAAAGATGGTTGCCAAGAACATC | 6 (0.000115%) | | | CATCCAACCTAGGCGAGACAAGGGTTCACATTTCGTTCATCACCCTTGGCCGGCTTTCGAACAGCCGGACTCCCATCAAAAGATGGTTGCCAAGAACATC | 6 (0.000115%) | | | CATCCTTTGCTGATGCGGGACGGAAGCTGGTCTCCCGTGTGTTACCGCACGCGGTTGGCCTAAATCCGAGCCAAGGACGCCTGGAGCGTACCGACATGCG | 126 (0.002415%) | | | CATCGACACCTTGCGGCTAGGAACTGGAACGAGACGGGTGGCAAAGATTTCGAGTAGCACTTCATACTACCGTGGGTTTTTTAAACCTTCCGAGTTTTGT | 5 (0.000096%) | | | CATCGACCAGAGGCTGTTCACCTTGGAGACCTGATGCGGTTATGAGTACGACCGGGCGTGAGCGGCACTCGGTCCTCCGGATTTTCAAGGGCCGCCGGGG | 4 (0.000077%) | | | CATGAATCATCAGAGCAACGGGCAGAGCCCGCGTCGACCTTTTATCTAATAAATGCGTCCCTTCCATAAGTCGGGGTTTGTTGCACGTATTAGCTCTAGA | 163 (0.003124%) | | | CATGACATCAAGTCATATTCGACTCCAAAACACTAACCAACCTTCTTCTTGCTTCTCAAAGCTTTCATGGTGTAGCCAAAGTCCATATGAGTCTTTGGCT | 33 (0.000632%) | | | CATGCATCATAAGGATACTAAATCCTATTTTCTGGTAAATTTTCATAATTTTTTGACACCTCTAGCTAGGTCATTTGACCTGATACAACATCGGATTTTC | 3 (0.000057%) | | | CATGCATGTGTAAGTATGAACGAATTCAGACTGTGAAACTGCGAATGGCTCATTAAATCAGTTATAGTTTGTTTGATGGTAACTACTACTCGGATAACCG | 7 (0.000134%) | | | CATGCTAATGTATCCAGAGCGTAGGCTTGCTTTGAGCACTCTAATTTCTTCAAAGTAACAGCGCCGGAGGCACGACCCGGCCAATTAAGACCAGGAGCGT | 8 (0.000153%) | | | CATGCTTTCCAACGAAGCACGCCCATCCAACCTAGGCGAGACAAGGGTTCACATTTCGTTCATCACCCTTGGCCGGCTATCGAACAGCCGGACTCCCATC | 15 (0.000287%) | | | CATGGGCATCGACACCTTGCGGCTAGGAACTGGAACGAGACGGGTGGCAAAGATTTCGAGTAGCACTTCATACTACCGTGGGTTTTTTAAACCTTCCGAG | 7 (0.000134%) | | | CATGTCGGTACGCTCCAGGCGTCCTTGGCTCGGATTTAGGCCAACCGCGTGCGGTAACACACGGGAGACCAGCTTCCGTCCCGCATCAGCAAAGGATGGT | 10 (0.000192%) | | | CATGTGTAAGTATGAACGAATTCAGACTGTGAAACTGCGAATGGCTCATTAAATCAGTTATAGTTTGTTTGATGGTAACTACTACTCGGATAACCGTAGT | 3 (0.000057%) | | | CATTAAATCAGTTATAGTTTGTTTGATGGTAACTACTACTCGGATAACCGTAGTAATTCTAGAGCTAATACGTGCAACAAACCCCGACTTATGGAAGGGA | 12 (0.000230%) | | | CATTCAATCGGTAGGAGCGACGGGCGGTGTGTACAAAGGGCAGGGACGTAGTCAACGCGAGCTGATGACTCGCGCTTACTAGGAATTCCTCGTTGAAGAC | 13 (0.000249%) | | | CATTCCAATTACCAGACTCGAAAGAGCCCGGTATTGTTATTTATTGTCACTACCTCCCCGTGTCAGGATTGGGTAATTTGCGCGCCTGCTGCCTTCCTTG | 3 (0.000057%) | | | CATTCGTATTTCATAGTCAGAGGTGAAATTCTTGGATTTATGAAAGACGAACAACTGCGAAAGCATTTGCCAAGGATGTTTTCATTAATCAAGAACGAAA | 3 (0.000057%) | | | CATTGGAGGGCAAGTCTGGTGCCAGCAGCCGCGGTAATTCCAGCTCCAATAGCGTATATTTAAGTTGTTGCAGTTAAAAAGCTCGTAGTTGAACCTTGGG | 28 (0.000537%) | | | CATTGTCAGGTGGGGAGTTTGGCTGGGGCGGCACATCTGTTAAAAGATAACGCAGGTGTCCTAAGATGAGCTCAACGAGAACAGAAATCTCGTGTGGAAC | 14 (0.000268%) | | | CATTGTTCCATCGACCAGAGGCTGTTCACCTTGGAGACCTGATGCGGTTATGAGTACGACCGGGCGTGAGCGGCACTCGGTCCTCCGGATTTTCAAGGGC | 25 (0.000479%) | | | CCAAAGACTCATATGGACTTTGGCTACACCATGAAAGCTTTGAGAAGCAAGAAGAAGGTTGGTTAGTGTTTTGGAGTCGAATATGACTTGATGTCATGTG | 161 (0.003085%) | | | CCAACCTAGGCGAGACAAGGGTTCACATTTCGTTCATCACCCTTGGCCGGCTATCGAACAGCCGGACTCCCATCAAAAGATGGTTGCCAAGAACATCTTC | 24 (0.000460%) | | | CCAACCTAGGCGAGACAAGGGTTCACATTTCGTTCATCACCCTTGGCCGGCTTTCGAACAGCCGGACTCCCATCAAAAGATGGTTGCCAAGAACATCTTC | 18 (0.000345%) | | | CCAACGAAGCACGCCCATCCAACCTAGGCGAGACAAGGGTTCACATTTCGTTCATCACCCTTGGCCGGCTATCGAACAGCCGGACTCCCATCAAAAGATG | 13 (0.000249%) | | | CCAACGAAGCACGCCCATCCAACCTAGGCGAGACAAGGGTTCACATTTCGTTCATCACCCTTGGCCGGCTTTCGAACAGCCGGACTCCCATCAAAAGATG | 11 (0.000211%) | | | CCAATTACCAGACTCGAAAGAGCCCGGTATTGTTATTTATTGTCACTACCTCCCCGTGTCAGGATTGGGTAATTTGCGCGCCTGCTGCCTTCCTTGGATG | 7 (0.000134%) | | | CCACATCCTCTCAAACGCAATGGAAAGAGAGAAAGGACGAGGTCTTGACCGTCATCTTTTGCCCGAAGGACGGATGAGCTTTGGCGGGACTGAATCACTT | 3 (0.000057%) | | | CCACCCTAACGCCTCGAAGAACTAATGGCAGCCCACGCAAGGCAAGCCCATTCTCCTCGACGATTCAGCAGTTTTTGTCCGAGAACTGCTGAGAAAACTC | 5 (0.000096%) | | | CCACCGCATGTCGGTACGCTCCAGGCGTCCTTGGCTCGGATTTAGGCCAACCGCGTGCGGTAACACACGGGAGACCAGCTTCCGTCCCGCATCAGCAAAG | 7 (0.000134%) | | | CCACGCTTTCACGGTTCGTATTCGTACTGAAAATCAGAATCAAACGAGCTTTTACCCTTTTGTTCCACACGAGATTTCTGTTCTCGTTGAGCTCATCTTA | 56 (0.001073%) | | | CCAGACTCGAAAGAGCCCGGTATTGTTATTTATTGTCACTACCTCCCCGTGTCAGGATTGGGTAATTTGCGCGCCTGCTGCCTTCCTTGGATGTGGTAGC | 13 (0.000249%) | | | CCAGAGCGTAGGCTTGCTTTGAGCACTCTAATTTCTTCAAAGTAACAGCGCCGGAGGCACGACCCGGCCAATTAAGACCAGGAGCGTATCGCCGACCGAA | 19 (0.000364%) | | | CCAGAGGCTGTTCACCTTGGAGACCTGATGCGGTTATGAGTACGACCGGGCGTGAGCGGCACTCGGTCCTCCGGATTTTCAAGGGCCGCCGGGGGCGCAC | 13 (0.000249%) | | | CCAGCAGCCGCGGTAATTCCAGCTCCAATAGCGTATATTTAAGTTGTTGCAGTTAAAAAGCTCGTAGTTGAACCTTGGGATGGGTCGGCCGGTCCGCCTT | 18 (0.000345%) | | | CCAGCCGTTCGTTTGCATGTTCCTTGACACTTTTCGTGCCGGGGTTTTGTGATATCCGGAAGCAACGCGCACGACAAGACCGAGATAAAAGCTCCCGATC | 5 (0.000096%) | | | CCAGTAGTCATATGCTTGTCTCAAAGATTAAGCCATGCATGTGTAAGTATGAACGAATTCAGACTGTGAAACTGCGAATGGCTCATTAAATCAGTTATAG | 92 (0.001763%) | | | CCATCCTTTGCTGATGCGGGACGGAAGCTGGTCTCCCGTGTGTTACCGCACGCGGTTGGCCTAAATCCGAGCCAAGGACGCCTGGAGCGTACCGACATGC | 8 (0.000153%) | | | CCATGCTAATGTATCCAGAGCGTAGGCTTGCTTTGAGCACTCTAATTTCTTCAAAGTAACAGCGCCGGAGGCACGACCCGGCCAATTAAGACCAGGAGCG | 5 (0.000096%) | | | CCATGCTTTCCAACGAAGCACGCCCATCCAACCTAGGCGAGACAAGGGTTCACATTTCGTTCATCACCCTTGGCCGGCTATCGAACAGCCGGACTCCCAT | 27 (0.000517%) | | | CCATGGGCATCGACACCTTGCGGCTAGGAACTGGAACGAGACGGGTGGCAAAGATTTCGAGTAGCACTTCATACTACCGTGGGTTTTTTAAACCTTCCGA | 8 (0.000153%) | | | CCATTGGAGGGCAAGTCTGGTGCCAGCAGCCGCGGTAATTCCAGCTCCAATAGCGTATATTTAAGTTGTTGCAGTTAAAAAGCTCGTAGTTGAACCTTGG | 12 (0.000230%) | | | CCCATCCAACCTAGGCGAGACAAGGGTTCACATTTCGTTCATCACCCTTGGCCGGCTATCGAACAGCCGGACTCCCATCAAAAGATGGTTGCCAAGAACA | 15 (0.000287%) | | | CCCATCCAACCTAGGCGAGACAAGGGTTCACATTTCGTTCATCACCCTTGGCCGGCTTTCGAACAGCCGGACTCCCATCAAAAGATGGTTGCCAAGAACA | 21 (0.000402%) | | | CCCATGCTAATGTATCCAGAGCGTAGGCTTGCTTTGAGCACTCTAATTTCTTCAAAGTAACAGCGCCGGAGGCACGACCCGGCCAATTAAGACCAGGAGC | 13 (0.000249%) | | | CCCATGGGCATCGACACCTTGCGGCTAGGAACTGGAACGAGACGGGTGGCAAAGATTTCGAGTAGCACTTCATACTACCGTGGGTTTTTTAAACCTTCCG | 11 (0.000211%) | | | CCCCGCCACATCCTCTCAAACGCAATGGAAAGAGAGAAAGGACGAGGTCTTGACCGTCATCTTTTGCCCGAAGGACGGATGAGCTTTGGCGGGACTGAAT | 6 (0.000115%) | | | CCCGAGTGTGAGCGAGGTGTGAGTGTCGCCCATGGGCATCGACACCTTGCGGCTAGGAACTGGAACGAGACGGGTAGCAAAGATTTCGAGTAGCACTTCA | 10 (0.000192%) | | | CCCGAGTGTGAGCGAGGTGTGAGTGTCGCCCATGGGCATCGACACCTTGCGGCTAGGAACTGGAACGAGACGGGTGGCAAAGATTTCGAGTAGCACTTCA | 24 (0.000460%) | | | CCCGCCACATCCTCTCAAACGCAATGGAAAGAGAGAAAGGACGAGGTCTTGACCGTCATCTTTTGCCCGAAGGACGGATGAGCTTTGGCGGGACTGAATC | 6 (0.000115%) | | | CCCGCGTCGACCTTTTATCTAATAAATGCGTCCCTTCCATAAGTCGGGGTTTGTTGCACGTATTAGCTCTAGAATTACTACGGTTATCCGAGTAGTAGTT | 40 (0.000767%) | | | CCCGGCCGTCGGGGCAAGAGCCAGGCCTCGATGAGTAGGAGGGCGCGGCGGTCGCTGCAAAACCTAGGGCGCGAGCCCGGGCGGAGCGGCCGTCGGTGCA | 19 (0.000364%) | | | CCCTAACGCCTCGAAGAACTAATGGCAGCCCACGCAAGGCAAGCCCATTCTCCTCGACGATTCAGCAGTTTTTGTCCGAGAACTGCTGAGAAAACTCGGA | 10 (0.000192%) | | | CCCTCACCATCCTTTGCTGATGCGGGACGGAAGCTGGTCTCCCGTGTGTTACCGCACGCGGTTGGCCTAAATCCGAGCCAAGGACGCCTGGAGCGTACCG | 20 (0.000383%) | | | CCCTGTTGAGCTTGACTCTAGTCCGACTTTGTGAAATGACTTGAGAGGTGTAGGATAAGTGGGAGCTTCGGCGCAAGTGAAATACCACTACTTTTAACGT | 13 (0.000249%) | | | CCCTTAGATGTTCTGGGCCGCACGCGCGCTACACTGATGTATTCAACGAGTTCACACCTTGGCCGACAGGCCCGGGTAATCTTTGAAATTTCATCGTGAT | 5 (0.000096%) | | | CCCTTGCCTACATTGTTCCATCGACCAGAGGCTGTTCACCTTGGAGACCTGATGCGGTTATGAGTACGACCGGGCGTGAGCGGCACTCGGTCCTCCGGAT | 35 (0.000671%) | | | CCGACTTCCCTTGCCTACATTGTTCCATCGACCAGAGGCTGTTCACCTTGGAGACCTGATGCGGTTATGAGTACGACCGGGCGTGAGCGGCACTCGGTCC | 48 (0.000920%) | | | CCGACTTTGTGAAATGACTTGAGAGGTGTAGGATAAGTGGGAGCTTCGGCGCAAGTGAAATACCACTACTTTTAACGTTATTTTACTTACTCCGTGAATC | 19 (0.000364%) | | | CCGAGTGTGAGCGAGGTGTGAGTGTCGCCCATGGGCATCGACACCTTGCGGCTAGGAACTGGAACGAGACGGGTAGCAAAGATTTCGAGTAGCACTTCAT | 12 (0.000230%) | | | CCGAGTGTGAGCGAGGTGTGAGTGTCGCCCATGGGCATCGACACCTTGCGGCTAGGAACTGGAACGAGACGGGTGGCAAAGATTTCGAGTAGCACTTCAT | 25 (0.000479%) | | | CCGCCACATCCTCTCAAACGCAATGGAAAGAGAGAAAGGACGAGGTCTTGACCGTCATCTTTTGCCCGAAGGACGGATGAGCTTTGGCGGGACTGAATCA | 11 (0.000211%) | | | CCGCCTAGGCTGTCCCGAGTGTGAGCGAGGTGTGAGTGTCGCCCATGGGCATCGACACCTTGCGGCTAGGAACTGGAACGAGACGGGTAGCAAAGATTTC | 31 (0.000594%) | | | CCGCCTAGGCTGTCCCGAGTGTGAGCGAGGTGTGAGTGTCGCCCATGGGCATCGACACCTTGCGGCTAGGAACTGGAACGAGACGGGTGGCAAAGATTTC | 62 (0.001188%) | | | CCGCGGTAATTCCAGCTCCAATAGCGTATATTTAAGTTGTTGCAGTTAAAAAGCTCGTAGTTGAACCTTGGGATGGGTCGGCCGGTCCGCCTTTGGTGTG | 6 (0.000115%) | | | CCGCGTCGACCTTTTATCTAATAAATGCGTCCCTTCCATAAGTCGGGGTTTGTTGCACGTATTAGCTCTAGAATTACTACGGTTATCCGAGTAGTAGTTA | 12 (0.000230%) | | | CCGCTAAGGAGTGTGTAACAACTCACCTGCCGAATCAACTAGCCCCGAAAATGGATGGCGCTTAAGCGCGCGACCTATACCCGGCCGTCGGGGCAAGAGC | 32 (0.000613%) | | | CCGGATTGCTCCGTTCCGCATCCGACCAGGACGCATCGCCGGCCCCCATCCGCTTCCCTCCCGACAATTTCAAGCACTCTTTGACTCTCTTTTCAAAGTC | 34 (0.000652%) | | | CCGGCCGTCGGGGCAAGAGCCAGGCCTCGATGAGTAGGAGGGCGCGGCGGTCGCTGCAAAACCTAGGGCGCGAGCCCGGGCGGAGCGGCCGTCGGTGCAG | 18 (0.000345%) | | | CCGGGCGTCGGCCTGTGGGCTCCCCATTCGACCCGTCTTGAAACACGGACCAAGGAGTCTGACATGTGTGCGAGTCAACGGGTGAGTAAACCCGTAAGGC | 13 (0.000249%) | | | CCGGGGAGGCGAATGCCAGCCGTTCGTTTGCATGTTCCTTGACACTTTTCGTGCCGGGGTTTTGTGATATCCGGAAGCAACGCGCACGACAAGACCGAGA | 14 (0.000268%) | | | CCGTCGAGTTATCATGAATCATCAGAGCAACGGGCAGAGCCCGCGTCGACCTTTTATCTAATAAATGCGTCCCTTCCATAAGTCGGGGTTTGTTGCACGT | 28 (0.000537%) | | | CCGTCTCCGGGGAGGCGAATGCCAGCCGTTCGTTTGCATGTTCCTTGACACTTTTCGTGCCGGGGTTTTGTGATATCCGGAAGCAACGCGCACGACAAGA | 10 (0.000192%) | | | CCTAACGCCTCGAAGAACTAATGGCAGCCCACGCAAGGCAAGCCCATTCTCCTCGACGATTCAGCAGTTTTTGTCCGAGAACTGCTGAGAAAACTCGGAA | 18 (0.000345%) | | | CCTAACGGCGTGCCTCGGCATCAGCGTGCTCCGGGCGTCGGCCTGTGGGCTCCCCATTCGACCCGTCTTGAAACACGGACCAAGGAGTCTGACATGTGTG | 4 (0.000077%) | | | CCTAAGTAGTGTTTCCTTGTTAGAAGACACAAAGCCAAAGACTCATATGGACTTTGGCTACACCATGAAAGCTTTGAGAAGCAAGAAGAAGGTTGGTTAG | 53 (0.001016%) | | | CCTACATTGTTCCATCGACCAGAGGCTGTTCACCTTGGAGACCTGATGCGGTTATGAGTACGACCGGGCGTGAGCGGCACTCGGTCCTCCGGATTTTCAA | 13 (0.000249%) | | | CCTAGGCGAGACAAGGGTTCACATTTCGTTCATCACCCTTGGCCGGCTATCGAACAGCCGGACTCCCATCAAAAGATGGTTGCCAAGAACATCTTCGTTA | 11 (0.000211%) | | | CCTAGGCGAGACAAGGGTTCACATTTCGTTCATCACCCTTGGCCGGCTTTCGAACAGCCGGACTCCCATCAAAAGATGGTTGCCAAGAACATCTTCGTTA | 12 (0.000230%) | | | CCTAGGCTGTCCCGAGTGTGAGCGAGGTGTGAGTGTCGCCCATGGGCATCGACACCTTGCGGCTAGGAACTGGAACGAGACGGGTAGCAAAGATTTCGAG | 45 (0.000862%) | | | CCTAGGCTGTCCCGAGTGTGAGCGAGGTGTGAGTGTCGCCCATGGGCATCGACACCTTGCGGCTAGGAACTGGAACGAGACGGGTGGCAAAGATTTCGAG | 59 (0.001131%) | | | CCTATACCCGGCCGTCGGGGCAAGAGCCAGGCCTCGATGAGTAGGAGGGCGCGGCGGTCGCTGCAAAACCTAGGGCGCGAGCCCGGGCGGAGCGGCCGTC | 19 (0.000364%) | | | CCTATGATGTTATCCCATGCTAATGTATCCAGAGCGTAGGCTTGCTTTGAGCACTCTAATTTCTTCAAAGTAACAGCGCCGGAGGCACGACCCGGCCAAT | 24 (0.000460%) | | | CCTATTTTCTGGTAAATTTTCATAATTTTTTGACACCTCTAGCTAGGTCATTTGACCTGATACAACATCGGATTTTCATGGTCTAGTTGGGGCTCCGTGG | 4 (0.000077%) | | | CCTCACCATCCTTTGCTGATGCGGGACGGAAGCTGGTCTCCCGTGTGTTACCGCACGCGGTTGGCCTAAATCCGAGCCAAGGACGCCTGGAGCGTACCGA | 81 (0.001552%) | | | CCTCCGCTTATTGATATGCTTAAACTCAGCGGGTAATCCCGCCTGACCTGGGGTCGCTATATGGACTTTGGGTCATCTACAGCTTCCGGACAAGAGCGAC | 3 (0.000057%) | | | CCTCGAAGAACTAATGGCAGCCCACGCAAGGCAAGCCCATTCTCCTCGACGATTCAGCAGTTTTTGTCCGAGAACTGCTGAGAAAACTCGGAAAAAGGCA | 12 (0.000230%) | | | CCTCGGCATCAGCGTGCTCCGGGCGTCGGCCTGTGGGCTCCCCATTCGACCCGTCTTGAAACACGGACCAAGGAGTCTGACATGTGTGCGAGTCAACGGG | 7 (0.000134%) | | | CCTCGTTAAGGGATTTAGATTGTACTCATTCCAATTACCAGACTCGAAAGAGCCCGGTATTGTTATTTATTGTCACTACCTCCCCGTGTCAGGATTGGGT | 209 (0.004005%) | | | CCTGTTGAGCTTGACTCTAGTCCGACTTTGTGAAATGACTTGAGAGGTGTAGGATAAGTGGGAGCTTCGGCGCAAGTGAAATACCACTACTTTTAACGTT | 7 (0.000134%) | | | CCTTAGATGTTCTGGGCCGCACGCGCGCTACACTGATGTATTCAACGAGTTCACACCTTGGCCGACAGGCCCGGGTAATCTTTGAAATTTCATCGTGATG | 7 (0.000134%) | | | CCTTGCCTACATTGTTCCATCGACCAGAGGCTGTTCACCTTGGAGACCTGATGCGGTTATGAGTACGACCGGGCGTGAGCGGCACTCGGTCCTCCGGATT | 12 (0.000230%) | | | CCTTGTTAGAAGACACAAAGCCAAAGACTCATATGGACTTTGGCTACACCATGAAAGCTTTGAGAAGCAAGAAGAAGGTTGGTTAGTGTTTTGGAGTCGA | 49 (0.000939%) | | | CCTTGTTAGTTTCTTTTCCTCCGCTTATTGATATGCTTAAACTCAGCGGGTAATCCCGCCTGACCTGGGGTCGCTATATGGACTTTGGGTCATCTACAGC | 11 (0.000211%) | | | CCTTTGCTGATGCGGGACGGAAGCTGGTCTCCCGTGTGTTACCGCACGCGGTTGGCCTAAATCCGAGCCAAGGACGCCTGGAGCGTACCGACATGCGGTG | 7 (0.000134%) | | | CGAAAGAGCCCGGTATTGTTATTTATTGTCACTACCTCCCCGTGTCAGGATTGGGTAATTTGCGCGCCTGCTGCCTTCCTTGGATGTGGTAGCCGTTTCT | 19 (0.000364%) | | | CGAAATCCGCTAAGGAGTGTGTAACAACTCACCTGCCGAATCAACTAGCCCCGAAAATGGATGGCGCTTAAGCGCGCGACCTATACCCGGCCGTCGGGGC | 9 (0.000172%) | | | CGAAATCCTATGATGTTATCCCATGCTAATGTATCCAGAGCGTAGGCTTGCTTTGAGCACTCTAATTTCTTCAAAGTAACAGCGCCGGAGGCACGACCCG | 477 (0.009141%) | | | CGAAGCACGCCCATCCAACCTAGGCGAGACAAGGGTTCACATTTCGTTCATCACCCTTGGCCGGCTATCGAACAGCCGGACTCCCATCAAAAGATGGTTG | 23 (0.000441%) | | | CGAAGCACGCCCATCCAACCTAGGCGAGACAAGGGTTCACATTTCGTTCATCACCCTTGGCCGGCTTTCGAACAGCCGGACTCCCATCAAAAGATGGTTG | 22 (0.000422%) | | | CGAATATGACTTGATGTCATGTGTATGATTGAGTATAAGA | 518 (0.003971%) | | | CGAATGCCAGCCGTTCGTTTGCATGTTCCTTGACACTTTTCGTGCCGGGGTTTTGTGATATCCGGAAGCAACGCGCACGACAAGACCGAGATAAAAGCTC | 23 (0.000441%) | | | CGAATGGCTCATTAAATCAGTTATAGTTTGTTTGATGGTAACTACTACTCGGATAACCGTAGTAATTCTAGAGCTAATACGTGCAACAAACCCCGACTTA | 9 (0.000172%) | | | CGACCAGAGGCTGTTCACCTTGGAGACCTGATGCGGTTATGAGTACGACCGGGCGTGAGCGGCACTCGGTCCTCCGGATTTTCAAGGGCCGCCGGGGGCG | 17 (0.000326%) | | | CGACCTTTTATCTAATAAATGCGTCCCTTCCATAAGTCGGGGTTTGTTGCACGTATTAGCTCTAGAATTACTACGGTTATCCGAGTAGTAGTTACCATCA | 25 (0.000479%) | | | CGACGGGCGGTGTGTACAAAGGGCAGGGACGTAGTCAACGCGAGCTGATGACTCGCGCTTACTAGGAATTCCTCGTTGAAGACCAACAATTGCAATGATC | 27 (0.000517%) | | | CGACGTGGGTGGTTCGCCGCCCGCGACGTCGCGAGAAGTCCACTAAACCTTATCATTTAGAGGAAGGAGAAGTCGTAACAAGGTTTCCGTAGGTGAACCT | 97 (0.001859%) | | | CGACTCCAAAACACTAACCAACCTTCTTCTTGCTTCTCAAAGCTTTCATGGTGTAGCCAAAGTCCATATGAGTCTTTGGCTTTGTGTCTTCTAACAAGGA | 58 (0.001112%) | | | CGACTTCCCTTGCCTACATTGTTCCATCGACCAGAGGCTGTTCACCTTGGAGACCTGATGCGGTTATGAGTACGACCGGGCGTGAGCGGCACTCGGTCCT | 34 (0.000652%) | | | CGACTTTGTGAAATGACTTGAGAGGTGTAGGATAAGTGGGAGCTTCGGCGCAAGTGAAATACCACTACTTTTAACGTTATTTTACTTACTCCGTGAATCG | 15 (0.000287%) | | | CGAGACAAGGGTTCACATTTCGTTCATCACCCTTGGCCGGCTATCGAACAGCCGGACTCCCATCAAAAGATGGTTGCCAAGAACATCTTCGTTACGGTTT | 7 (0.000134%) | | | CGAGACAAGGGTTCACATTTCGTTCATCACCCTTGGCCGGCTTTCGAACAGCCGGACTCCCATCAAAAGATGGTTGCCAAGAACATCTTCGTTACGGTTT | 6 (0.000115%) | | | CGAGGTGTGAGTGTCGCCCATGGGCATCGACACCTTGCGGCTAGGAACTGGAACGAGACGGGTAGCAAAGATTTCGAGTAGCACTTCATACTACCGTGGG | 10 (0.000192%) | | | CGAGGTGTGAGTGTCGCCCATGGGCATCGACACCTTGCGGCTAGGAACTGGAACGAGACGGGTGGCAAAGATTTCGAGTAGCACTTCATACTACCGTGGG | 22 (0.000422%) | | | CGAGTGTGAGCGAGGTGTGAGTGTCGCCCATGGGCATCGACACCTTGCGGCTAGGAACTGGAACGAGACGGGTAGCAAAGATTTCGAGTAGCACTTCATA | 29 (0.000556%) | | | CGAGTGTGAGCGAGGTGTGAGTGTCGCCCATGGGCATCGACACCTTGCGGCTAGGAACTGGAACGAGACGGGTGGCAAAGATTTCGAGTAGCACTTCATA | 31 (0.000594%) | | | CGAGTTATCATGAATCATCAGAGCAACGGGCAGAGCCCGCGTCGACCTTTTATCTAATAAATGCGTCCCTTCCATAAGTCGGGGTTTGTTGCACGTATTA | 158 (0.003028%) | | | CGATCCGCCTAGGCTGTCCCGAGTGTGAGCGAGGTGTGAGTGTCGCCCATGGGCATCGACACCTTGCGGCTAGGAACTGGAACGAGACGGGTAGCAAAGA | 4 (0.000077%) | | | CGATCCGCCTAGGCTGTCCCGAGTGTGAGCGAGGTGTGAGTGTCGCCCATGGGCATCGACACCTTGCGGCTAGGAACTGGAACGAGACGGGTGGCAAAGA | 9 (0.000172%) | | | CGATCCGGTTAAAATTCCGGAACCGGGACGTGGCGGTTGACGGCAACGTTAGGGAGTCCGGAGACGTCGGCGGGGGCCTCGGGAAGAGTTATCTTTTCTG | 3 (0.000057%) | | | CGATCCGTAACTTCGGGAAAAGGATTGGCTCTGAGGGCTGGGCTCGGGGGTCCCAGTTCCGAACCCGTCGGCTGTCAGCGGACTGCTCGAGCTGCTTCCG | 3 (0.000057%) | | | CGATCCGTCGAGTTATCATGAATCATCAGAGCAACGGGCAGAGCCCGCGTCGACCTTTTATCTAATAAATGCGTCCCTTCCATAAGTCGGGGTTTGTTGC | 183 (0.003507%) | | | CGATCCTCGTTAAGGGATTTAGATTGTACTCATTCCAATTACCAGACTCGAAAGAGCCCGGTATTGTTATTTATTGTCACTACCTCCCCGTGTCAGGATT | 7 (0.000134%) | | | CGATCGAAATCCTATGATGTTATCCCATGCTAATGTATCCAGAGCGTAGGCTTGCTTTGAGCACTCTAATTTCTTCAAAGTAACAGCGCCGGAGGCACGA | 5 (0.000096%) | | | CGATCGGCGGACCGGATTGCTCCGTTCCGCATCCGACCAGGACGCATCGCCGGCCCCCATCCGCTTCCCTCCCGACAATTTCAAGCACTCTTTGACTCTC | 3 (0.000057%) | | | CGATCGGGTTGCGGTTTAAGTTGTTATACTCAATCATACA | 75 (0.000575%) | | | CGCAACAAGATCTTAAAGGCGTAAGAATTGTATCCTTGTTAAAAGACACAAAGCCAAAGACTCATATGGACTTTGGCTACACCATGAAAGCTTTGAGAAG | 52 (0.000997%) | | | CGCAACCGGATCTTAAAGGCGTAAGAATTGTATCCTTGTTAGAAGACACAAAGCCAAAGACTCATATGGACTTTGGCTACACCATGAAAGCTTTGAGAAG | 5 (0.000096%) | | | CGCATGTCGGTACGCTCCAGGCGTCCTTGGCTCGGATTTAGGCCAACCGCGTGCGGTAACACACGGGAGACCAGCTTCCGTCCCGCATCAGCAAAGGATG | 11 (0.000211%) | | | CGCCCATCCAACCTAGGCGAGACAAGGGTTCACATTTCGTTCATCACCCTTGGCCGGCTATCGAACAGCCGGACTCCCATCAAAAGATGGTTGCCAAGAA | 72 (0.001380%) | | | CGCCCATCCAACCTAGGCGAGACAAGGGTTCACATTTCGTTCATCACCCTTGGCCGGCTTTCGAACAGCCGGACTCCCATCAAAAGATGGTTGCCAAGAA | 82 (0.001571%) | | | CGCCCATGGGCATCGACACCTTGCGGCTAGGAACTGGAACGAGACGGGTGGCAAAGATTTCGAGTAGCACTTCATACTACCGTGGGTTTTTTAAACCTTC | 21 (0.000402%) | | | CGCCGCCCGCGACGTCGCGAGAAGTCCACTAAACCTTATCATTTAGAGGAAGGAGAAGTCGTAACAAGGTTTCCGTAGGTGAACCTGCGGAAGGATCGAT | 11 (0.000211%) | | | CGCCGTTACTAAGGGAATCCTTGTTAGTTTCTTTTCCTCCGCTTATTGATATGCTTAAACTCAGCGGGTAATCCCGCCTGACCTGGGGTCGCTATATGGA | 6 (0.000115%) | | | CGCCTAACGGCGTGCCTCGGCATCAGCGTGCTCCGGGCGTCGGCCTGTGGGCTCCCCATTCGACCCGTCTTGAAACACGGACCAAGGAGTCTGACATGTG | 37 (0.000709%) | | | CGCCTAGGCTGTCCCGAGTGTGAGCGAGGTGTGAGTGTCGCCCATGGGCATCGACACCTTGCGGCTAGGAACTGGAACGAGACGGGTAGCAAAGATTTCG | 111 (0.002127%) | | | CGCCTAGGCTGTCCCGAGTGTGAGCGAGGTGTGAGTGTCGCCCATGGGCATCGACACCTTGCGGCTAGGAACTGGAACGAGACGGGTGGCAAAGATTTCG | 170 (0.003258%) | | | CGCCTCGAAGAACTAATGGCAGCCCACGCAAGGCAAGCCCATTCTCCTCGACGATTCAGCAGTTTTTGTCCGAGAACTGCTGAGAAAACTCGGAAAAAGG | 6 (0.000115%) | | | CGCGACCTATACCCGGCCGTCGGGGCAAGAGCCAGGCCTCGATGAGTAGGAGGGCGCGGCGGTCGCTGCAAAACCTAGGGCGCGAGCCCGGGCGGAGCGG | 12 (0.000230%) | | | CGCGCCTAACGGCGTGCCTCGGCATCAGCGTGCTCCGGGCGTCGGCCTGTGGGCTCCCCATTCGACCCGTCTTGAAACACGGACCAAGGAGTCTGACATG | 46 (0.000882%) | | | CGCGCGACCTATACCCGGCCGTCGGGGCAAGAGCCAGGCCTCGATGAGTAGGAGGGCGCGGCGGTCGCTGCAAAACCTAGGGCGCGAGCCCGGGCGGAGC | 13 (0.000249%) | | | CGCGGTAATTCCAGCTCCAATAGCGTATATTTAAGTTGTTGCAGTTAAAAAGCTCGTAGTTGAACCTTGGGATGGGTCGGCCGGTCCGCCTTTGGTGTGC | 17 (0.000326%) | | | CGCGTCGACCTTTTATCTAATAAATGCGTCCCTTCCATAAGTCGGGGTTTGTTGCACGTATTAGCTCTAGAATTACTACGGTTATCCGAGTAGTAGTTAC | 8 (0.000153%) | | | CGCTAAGGAGTGTGTAACAACTCACCTGCCGAATCAACTAGCCCCGAAAATGGATGGCGCTTAAGCGCGCGACCTATACCCGGCCGTCGGGGCAAGAGCC | 15 (0.000287%) | | | CGCTTATTGATATGCTTAAACTCAGCGGGTAATCCCGCCTGACCTGGGGTCGCTATATGGACTTTGGGTCATCTACAGCTTCCGGACAAGAGCGACCGAT | 6 (0.000115%) | | | CGCTTTCACGGTTCGTATTCGTACTGAAAATCAGAATCAAACGAGCTTTTACCCTTTTGTTCCACACGAGATTTCTGTTCTCGTTGAGCTCATCTTAGGA | 8 (0.000153%) | | | CGGACCGGATTGCTCCGTTCCGCATCCGACCAGGACGCATCGCCGGCCCCCATCCGCTTCCCTCCCGACAATTTCAAGCACTCTTTGACTCTCTTTTCAA | 3 (0.000057%) | | | CGGAGAATTAGGGTTCGATTCCGGAGAGGGAGCCTGAGAAACGGCTACCACATCCAAGGAAGGCAGCAGGCGCGCAAATTACCCAATCCTGACACGGGGA | 19 (0.000364%) | | | CGGATCTTAAAGGCGTAAGAATTGTATCCTTGTTAGAAGACACAAAGCCAAAGACTCATATGGACTTTGGCTACACCATGAAAGCTTTGAGAAGCAAGAA | 7 (0.000134%) | | | CGGATTGCTCCGTTCCGCATCCGACCAGGACGCATCGCCGGCCCCCATCCGCTTCCCTCCCGACAATTTCAAGCACTCTTTGACTCTCTTTTCAAAGTCC | 24 (0.000460%) | | | CGGCAATTCCCCGCCACATCCTCTCAAACGCAATGGAAAGAGAGAAAGGACGAGGTCTTGACCGTCATCTTTTGCCCGAAGGACGGATGAGCTTTGGCGG | 188 (0.003603%) | | | CGGCACATCTGTTAAAAGATAACGCAGGTGTCCTAAGATGAGCTCAACGAGAACAGAAATCTCGTGTGGAACAAAAGGGTAAAAGCTCGTTTGATTCTGA | 12 (0.000230%) | | | CGGCATCAGCGTGCTCCGGGCGTCGGCCTGTGGGCTCCCCATTCGACCCGTCTTGAAACACGGACCAAGGAGTCTGACATGTGTGCGAGTCAACGGGTGA | 15 (0.000287%) | | | CGGCCGTCGGGGCAAGAGCCAGGCCTCGATGAGTAGGAGGGCGCGGCGGTCGCTGCAAAACCTAGGGCGCGAGCCCGGGCGGAGCGGCCGTCGGTGCAGA | 46 (0.000882%) | | | CGGCGGACCGGATTGCTCCGTTCCGCATCCGACCAGGACGCATCGCCGGCCCCCATCCGCTTCCCTCCCGACAATTTCAAGCACTCTTTGACTCTCTTTT | 15 (0.000287%) | | | CGGCGTGCCTCGGCATCAGCGTGCTCCGGGCGTCGGCCTGTGGGCTCCCCATTCGACCCGTCTTGAAACACGGACCAAGGAGTCTGACATGTGTGCGAGT | 9 (0.000172%) | | | CGGGCAGAGCCCGCGTCGACCTTTTATCTAATAAATGCGTCCCTTCCATAAGTCGGGGTTTGTTGCACGTATTAGCTCTAGAATTACTACGGTTATCCGA | 47 (0.000901%) | | | CGGGCGGTGTGTACAAAGGGCAGGGACGTAGTCAACGCGAGCTGATGACTCGCGCTTACTAGGAATTCCTCGTTGAAGACCAACAATTGCAATGATCGAT | 43 (0.000824%) | | | CGGGCGTCGGCCTGTGGGCTCCCCATTCGACCCGTCTTGAAACACGGACCAAGGAGTCTGACATGTGTGCGAGTCAACGGGTGAGTAAACCCGTAAGGCG | 46 (0.000882%) | | | CGGGGAGGCGAATGCCAGCCGTTCGTTTGCATGTTCCTTGACACTTTTCGTGCCGGGGTTTTGTGATATCCGGAAGCAACGCGCACGACAAGACCGAGAT | 14 (0.000268%) | | | CGGGGGCATTCGTATTTCATAGTCAGAGGTGAAATTCTTGGATTTATGAAAGACGAACAACTGCGAAAGCATTTGCCAAGGATGTTTTCATTAATCAAGA | 22 (0.000422%) | | | CGGGTGACGGAGAATTAGGGTTCGATTCCGGAGAGGGAGCCTGAGAAACGGCTACCACATCCAAGGAAGGCAGCAGGCGCGCAAATTACCCAATCCTGAC | 3 (0.000057%) | | | CGGGTTGCGGTTTAAGTTGTTATACTCAATCATACACATG | 497 (0.003810%) | | | CGGGTTTACTCACCCGTTGACTCGCACACATGTCAGACTCCTTGGTCCGTGTTTCAAGACGGGTCGAATGGGGAGCCCACAGGCCGACGCCCGGAGCACG | 18 (0.000345%) | | | CGGTAATTCCAGCTCCAATAGCGTATATTTAAGTTGTTGCAGTTAAAAAGCTCGTAGTTGAACCTTGGGATGGGTCGGCCGGTCCGCCTTTGGTGTGCAT | 19 (0.000364%) | | | CGGTACGCTCCAGGCGTCCTTGGCTCGGATTTAGGCCAACCGCGTGCGGTAACACACGGGAGACCAGCTTCCGTCCCGCATCAGCAAAGGATGGTGAGGG | 4 (0.000077%) | | | CGGTAGGAGCGACGGGCGGTGTGTACAAAGGGCAGGGACGTAGTCAACGCGAGCTGATGACTCGCGCTTACTAGGAATTCCTCGTTGAAGACCAACAATT | 54 (0.001035%) | | | CGGTTTAAGTTCTTATACTCAATCATACACATGACATCAA | 555 (0.004255%) | | | CGGTTTAAGTTGTTATACTCAATCATACACATGACAACAAGTCATATTCGACTCCAAAACACTAACCAACCTTCTTCTTGCTTCTCAAAGCTTTCATGGT | 896 (0.017171%) | | | CGTAACTTCGGGAAAAGGATTGGCTCTGAGGGCTGGGCTCGGGGGTCCCAGTTCCGAACCCGTCGGCTGTCAGCGGACTGCTCGAGCTGCTTCCGCGGCG | 3 (0.000057%) | | | CGTAAGAATTGTATCCTTGTTAGAAGACACAAAGCCAAAGACTCATATGGACTTTGGCTACACCATGAAAGCTTTGAGAAGCAAGAAGAAGGTTGGTTAG | 58 (0.001112%) | | | CGTAGGCTTGCTTTGAGCACTCTAATTTCTTCAAAGTAACAGCGCCGGAGGCACGACCCGGCCAATTAAGACCAGGAGCGTATCGCCGACCGAAGGGACA | 17 (0.000326%) | | | CGTATTCGTACTGAAAATCAGAATCAAACGAGCTTTTACCCTTTTGTTCCACACGAGATTTCTGTTCTCGTTGAGCTCATCTTAGGACACCTGCGTTATC | 27 (0.000517%) | | | CGTATTTCATAGTCAGAGGTGAAATTCTTGGATTTATGAAAGACGAACAACTGCGAAAGCATTTGCCAAGGATGTTTTCATTAATCAAGAACGAAAGTTG | 7 (0.000134%) | | | CGTCCCTCACCATCCTTTGCTGATGCGGGACGGAAGCTGGTCTCCCGTGTGTTACCGCACGCGGTTGGCCTAAATCCGAGCCAAGGACGCCTGGAGCGTA | 11 (0.000211%) | | | CGTCGAGTTATCATGAATCATCAGAGCAACGGGCAGAGCCCGCGTCGACCTTTTATCTAATAAATGCGTCCCTTCCATAAGTCGGGGTTTGTTGCACGTA | 104 (0.001993%) | | | CGTCGGCCTGTGGGCTCCCCATTCGACCCGTCTTGAAACACGGACCAAGGAGTCTGACATGTGTGCGAGTCAACGGGTGAGTAAACCCGTAAGGCGCAAG | 6 (0.000115%) | | | CGTCTCCGGGGAGGCGAATGCCAGCCGTTCGTTTGCATGTTCCTTGACACTTTTCGTGCCGGGGTTTTGTGATATCCGGAAGCAACGCGCACGACAAGAC | 3 (0.000057%) | | | CGTGCCTCGGCATCAGCGTGCTCCGGGCGTCGGCCTGTGGGCTCCCCATTCGACCCGTCTTGAAACACGGACCAAGGAGTCTGACATGTGTGCGAGTCAA | 21 (0.000402%) | | | CGTGCTCCGGGCGTCGGCCTGTGGGCTCCCCATTCGACCCGTCTTGAAACACGGACCAAGGAGTCTGACATGTGTGCGAGTCAACGGGTGAGTAAACCCG | 9 (0.000172%) | | | CGTGGGTGGTTCGCCGCCCGCGACGTCGCGAGAAGTCCACTAAACCTTATCATTTAGAGGAAGGAGAAGTCGTAACAAGGTTTCCGTAGGTGAACCTGCG | 27 (0.000517%) | | | CGTTAAGGGATTTAGATTGTACTCATTCCAATTACCAGACTCGAAAGAGCCCGGTATTGTTATTTATTGTCACTACCTCCCCGTGTCAGGATTGGGTAAT | 61 (0.001169%) | | | CTAAATCCTATTTTCTGGTAAATTTTCATAATTTTTTGACACCTCTAGCTAGGTCATTTGACCTGATACAACATCGGATTTTCATGGTCTAGTTGGGGCT | 39 (0.000747%) | | | CTAACGCCTCGAAGAACTAATGGCAGCCCACGCAAGGCAAGCCCATTCTCCTCGACGATTCAGCAGTTTTTGTCCGAGAACTGCTGAGAAAACTCGGAAA | 28 (0.000537%) | | | CTAACGGCGTGCCTCGGCATCAGCGTGCTCCGGGCGTCGGCCTGTGGGCTCCCCATTCGACCCGTCTTGAAACACGGACCAAGGAGTCTGACATGTGTGC | 94 (0.001801%) | | | CTAAGTAGTGTTTCCTTGTTAGAAGACACAAAGCCAAAGACTCATATGGACTTTGGCTACACCATGAAAGCTTTGAGAAGCAAGAAGAAGGTTGGTTAGT | 77 (0.001476%) | | | CTAATGTATCCAGAGCGTAGGCTTGCTTTGAGCACTCTAATTTCTTCAAAGTAACAGCGCCGGAGGCACGACCCGGCCAATTAAGACCAGGAGCGTATCG | 37 (0.000709%) | | | CTACATTGTTCCATCGACCAGAGGCTGTTCACCTTGGAGACCTGATGCGGTTATGAGTACGACCGGGCGTGAGCGGCACTCGGTCCTCCGGATTTTCAAG | 48 (0.000920%) | | | CTAGGCGAGACAAGGGTTCACATTTCGTTCATCACCCTTGGCCGGCTATCGAACAGCCGGACTCCCATCAAAAGATGGTTGCCAAGAACATCTTCGTTAC | 25 (0.000479%) | | | CTAGGCGAGACAAGGGTTCACATTTCGTTCATCACCCTTGGCCGGCTTTCGAACAGCCGGACTCCCATCAAAAGATGGTTGCCAAGAACATCTTCGTTAC | 25 (0.000479%) | | | CTAGGCTGTCCCGAGTGTGAGCGAGGTGTGAGTGTCGCCCATGGGCATCGACACCTTGCGGCTAGGAACTGGAACGAGACGGGTAGCAAAGATTTCGAGT | 274 (0.005251%) | | | CTAGGCTGTCCCGAGTGTGAGCGAGGTGTGAGTGTCGCCCATGGGCATCGACACCTTGCGGCTAGGAACTGGAACGAGACGGGTGGCAAAGATTTCGAGT | 414 (0.007934%) | | | CTAGTCCGACTTTGTGAAATGACTTGAGAGGTGTAGGATAAGTGGGAGCTTCGGCGCAAGTGAAATACCACTACTTTTAACGTTATTTTACTTACTCCGT | 17 (0.000326%) | | | CTATACCCGGCCGTCGGGGCAAGAGCCAGGCCTCGATGAGTAGGAGGGCGCGGCGGTCGCTGCAAAACCTAGGGCGCGAGCCCGGGCGGAGCGGCCGTCG | 64 (0.001227%) | | | CTATGATGTTATCCCATGCTAATGTATCCAGAGCGTAGGCTTGCTTTGAGCACTCTAATTTCTTCAAAGTAACAGCGCCGGAGGCACGACCCGGCCAATT | 220 (0.004216%) | | | CTATTTTCTGGTAAATTTTCATAATTTTTTGACACCTCTAGCTAGGTCATTTGACCTGATACAACATCGGATTTTCATGGTCTAGTTGGGGCTCCGTGGG | 13 (0.000249%) | | | CTCAAACGCAATGGAAAGAGAGAAAGGACGAGGTCTTGACCGTCATCTTTTGCCCGAAGGACGGATGAGCTTTGGCGGGACTGAATCACTTCGAGTCACC | 4 (0.000077%) | | | CTCAAAGATTAAGCCATGCATGTGTAAGTATGAACGAATTCAGACTGTGAAACTGCGAATGGCTCATTAAATCAGTTATAGTTTGTTTGATGGTAACTAC | 24 (0.000460%) | | | CTCAAAGCTTTCATGGTGTAGCCAAAGTCCATATGAGTCTTTGGCTTTGTGTCTTCTAACAAGGAAACACTACTTAGGCTTATAAGATGCGGTTGCGGTT | 9 (0.000172%) | | | CTCAATCATACACATGACAACAAGTCATATTCGACTCCAAAACACTAACCAACCTTCTTCTTGCTTCTCAAAGCTTTCATGGTGTAGCCAAAGTCCATAT | 13 (0.000249%) | | | CTCAATCATACACATGACATCAAGTCATATTCGACTCCAAAACACTAACCAACCTTCTTCTTGCTTCTCAAAGCTTTCATGGTGTAGCCAAAGTCCATAT | 100 (0.001916%) | | | CTCACCATCCTTTGCTGATGCGGGACGGAAGCTGGTCTCCCGTGTGTTACCGCACGCGGTTGGCCTAAATCCGAGCCAAGGACGCCTGGAGCGTACCGAC | 9 (0.000172%) | | | CTCACCCGTTGACTCGCACACATGTCAGACTCCTTGGTCCGTGTTTCAAGACGGGTCGAATGGGGAGCCCACAGGCCGACGCCCGGAGCACGCTGATGCC | 15 (0.000287%) | | | CTCATATGGACTTTGGCTACACCATGAAAGCTTTGAGAAGCAAGAAGAAGGTTGGTTAGTGTTTTGGAGTCGAATATGACTTGATGTCATGTGTATGATT | 83 (0.001591%) | | | CTCATTAAATCAGTTATAGTTTGTTTGATGGTAACTACTACTCGGATAACCGTAGTAATTCTAGAGCTAATACGTGCAACAAACCCCGACTTATGGAAGG | 6 (0.000115%) | | | CTCATTCCAATTACCAGACTCGAAAGAGCCCGGTATTGTTATTTATTGTCACTACCTCCCCGTGTCAGGATTGGGTAATTTGCGCGCCTGCTGCCTTCCT | 18 (0.000345%) | | | CTCCGCTTATTGATATGCTTAAACTCAGCGGGTAATCCCGCCTGACCTGGGGTCGCTATATGGACTTTGGGTCATCTACAGCTTCCGGACAAGAGCGACC | 5 (0.000096%) | | | CTCCGGGCGTCGGCCTGTGGGCTCCCCATTCGACCCGTCTTGAAACACGGACCAAGGAGTCTGACATGTGTGCGAGTCAACGGGTGAGTAAACCCGTAAG | 15 (0.000287%) | | | CTCCGGGGAGGCGAATGCCAGCCGTTCGTTTGCATGTTCCTTGACACTTTTCGTGCCGGGGTTTTGTGATATCCGGAAGCAACGCGCACGACAAGACCGA | 38 (0.000728%) | | | CTCGAAAGAGCCCGGTATTGTTATTTATTGTCACTACCTCCCCGTGTCAGGATTGGGTAATTTGCGCGCCTGCTGCCTTCCTTGGATGTGGTAGCCGTTT | 41 (0.000786%) | | | CTCGAAGAACTAATGGCAGCCCACGCAAGGCAAGCCCATTCTCCTCGACGATTCAGCAGTTTTTGTCCGAGAACTGCTGAGAAAACTCGGAAAAAGGCAA | 17 (0.000326%) | | | CTCGCCGTTACTAAGGGAATCCTTGTTAGTTTCTTTTCCTCCGCTTATTGATATGCTTAAACTCAGCGGGTAATCCCGCCTGACCTGGGGTCGCTATATG | 34 (0.000652%) | | | CTCGGCATCAGCGTGCTCCGGGCGTCGGCCTGTGGGCTCCCCATTCGACCCGTCTTGAAACACGGACCAAGGAGTCTGACATGTGTGCGAGTCAACGGGT | 22 (0.000422%) | | | CTCGTTAAGGGATTTAGATTGTACTCATTCCAATTACCAGACTCGAAAGAGCCCGGTATTGTTATTTATTGTCACTACCTCCCCGTGTCAGGATTGGGTA | 151 (0.002894%) | | | CTCTAGTCCGACTTTGTGAAATGACTTGAGAGGTGTAGGATAAGTGGGAGCTTCGGCGCAAGTGAAATACCACTACTTTTAACGTTATTTTACTTACTCC | 30 (0.000575%) | | | CTCTCTCTCTCTCTCTCTCTCTCTCTCTCTCTCTCTCTCT | 47 (0.000360%) | | | CTGAAAATCAGAATCAAACGAGCTTTTACCCTTTTGTTCCACACGAGATTTCTGTTCTCGTTGAGCTCATCTTAGGACACCTGCGTTATCTTTTAACAGA | 21 (0.000402%) | | | CTGATGCGGGACGGAAGCTGGTCTCCCGTGTGTTACCGCACGCGGTTGGCCTAAATCCGAGCCAAGGACGCCTGGAGCGTACCGACATGCGGTGGTGAAC | 22 (0.000422%) | | | CTGCAGCACGCGCCTAACGGCGTGCCTCGGCATCAGCGTGCTCCGGGCGTCGGCCTGTGGGCTCCCCATTCGACCCGTCTTGAAACACGGACCAAGGAGT | 2112 (0.040475%) | | | CTGCCAGTAGTCATATGCTTGTCTCAAAGATTAAGCCATGCATGTGTAAGTATGAACGAATTCAGACTGTGAAACTGCGAATGGCTCATTAAATCAGTTA | 2028 (0.038866%) | | | CTGCGAATGGCTCATTAAATCAGTTATAGTTTGTTTGATGGTAACTACTACTCGGATAACCGTAGTAATTCTAGAGCTAATACGTGCAACAAACCCCGAC | 30 (0.000575%) | | | CTGGGCCGCACGCGCGCTACACTGATGTATTCAACGAGTTCACACCTTGGCCGACAGGCCCGGGTAATCTTTGAAATTTCATCGTGATGGGGATAGATCG | 75 (0.001437%) | | | CTGGGGCGGCACATCTGTTAAAAGATAACGCAGGTGTCCTAAGATGAGCTCAACGAGAACAGAAATCTCGTGTGGAACAAAAGGGTAAAAGCTCGTTTGA | 13 (0.000249%) | | | CTGGTGCCAGCAGCCGCGGTAATTCCAGCTCCAATAGCGTATATTTAAGTTGTTGCAGTTAAAAAGCTCGTAGTTGAACCTTGGGATGGGTCGGCCGGTC | 49 (0.000939%) | | | CTGTCCCGAGTGTGAGCGAGGTGTGAGTGTCGCCCATGGGCATCGACACCTTGCGGCTAGGAACTGGAACGAGACGGGTAGCAAAGATTTCGAGTAGCAC | 10 (0.000192%) | | | CTGTCCCGAGTGTGAGCGAGGTGTGAGTGTCGCCCATGGGCATCGACACCTTGCGGCTAGGAACTGGAACGAGACGGGTGGCAAAGATTTCGAGTAGCAC | 48 (0.000920%) | | | CTGTGATGCCCTTAGATGTTCTGGGCCGCACGCGCGCTACACTGATGTATTCAACGAGTTCACACCTTGGCCGACAGGCCCGGGTAATCTTTGAAATTTC | 114 (0.002185%) | | | CTGTTAAAAGATAACGCAGGTGTCCTAAGATGAGCTCAACGAGAACAGAAATCTCGTGTGGAACAAAAGGGTAAAAGCTCGTTTGATTCTGATTTTCAGT | 5 (0.000096%) | | | CTGTTGAGCTTGACTCTAGTCCGACTTTGTGAAATGACTTGAGAGGTGTAGGATAAGTGGGAGCTTCGGCGCAAGTGAAATACCACTACTTTTAACGTTA | 12 (0.000230%) | | | CTTAAAAGCCTAAGTAGTGTTTCCTTGTTAGAAGACACAAAGCCAAAGACTCATATGGACTTTGGCTACACCATGAAAGCTTTGAGAAGCAAGAAGAAGG | 196 (0.003756%) | | | CTTAAAGGCGTAAGAATTGTATCCTTGTTAAAAGACACAAAGCCAAAGACTCATATGGACTTTGGCTACACCATGAAAGCTTTGAGAAGCAAGAAGAAGG | 219 (0.004197%) | | | CTTAAAGGCGTAAGAATTGTATCCTTGTTAGAAGACACAAAGCCAAAGACTCATATGGACTTTGGCTACACCATGAAAGCTTTGAGAAGCAAGAAGAAGG | 225 (0.004312%) | | | CTTAAGCGCGCGACCTATACCCGGCCGTCGGGGCAAGAGCCAGGCCTCGATGAGTAGGAGGGCGCGGCGGTCGCTGCAAAACCTAGGGCGCGAGCCCGGG | 65 (0.001246%) | | | CTTAGATGTTCTGGGCCGCACGCGCGCTACACTGATGTATTCAACGAGTTCACACCTTGGCCGACAGGCCCGGGTAATCTTTGAAATTTCATCGTGATGG | 132 (0.002530%) | | | CTTATACTCAATCATACACATGACATCAAGTCATATTCGACTCCAAAACACTAACCAACCTTCTTCTTGCTTCTCAAAGCTTTCATGGTGTAGCCAAAGT | 127 (0.002434%) | | | CTTATTGATATGCTTAAACTCAGCGGGTAATCCCGCCTGACCTGGGGTCGCTATATGGACTTTGGGTCATCTACAGCTTCCGGACAAGAGCGACCGATAA | 37 (0.000709%) | | | CTTCAACGAGGAATTCCTAGTAAGCGCGAGTCATCAGCTCGCGTTGACTACGTCCCTGCCCTTTGTACACACCGCCCGTCGCTCCTACCGATTGAATGAT | 63 (0.001207%) | | | CTTCCCTTGCCTACATTGTTCCATCGACCAGAGGCTGTTCACCTTGGAGACCTGATGCGGTTATGAGTACGACCGGGCGTGAGCGGCACTCGGTCCTCCG | 57 (0.001092%) | | | CTTCTCAAAGCTTTCATGGTGTAGCCAAAGTCCATATGAGTCTTTGGCTTTGTGTCTTCTAACAAGGAAACACTACTTAGGCTTATAAGATGCGGTTGCG | 9 (0.000172%) | | | CTTGACTCTAGTCCGACTTTGTGAAATGACTTGAGAGGTGTAGGATAAGTGGGAGCTTCGGCGCAAGTGAAATACCACTACTTTTAACGTTATTTTACTT | 47 (0.000901%) | | | CTTGATGTCATGTGTATGATTGAGTATAAGAACTTAAACC | 258 (0.001978%) | | | CTTGCCTACATTGTTCCATCGACCAGAGGCTGTTCACCTTGGAGACCTGATGCGGTTATGAGTACGACCGGGCGTGAGCGGCACTCGGTCCTCCGGATTT | 55 (0.001054%) | | | CTTGCTTCTCAAAGCTTTCATGGTGTAGCCAAAGTCCATATGAGTCTTTGGCTTTGTGTCTTCTAACAAGGAAACACTACTTAGGCTTATAAGATGCGGT | 45 (0.000862%) | | | CTTGCTTTGAGCACTCTAATTTCTTCAAAGTAACAGCGCCGGAGGCACGACCCGGCCAATTAAGACCAGGAGCGTATCGCCGACCGAAGGGACAAGCCGA | 59 (0.001131%) | | | CTTGTCTCAAAGATTAAGCCATGCATGTGTAAGTATGAACGAATTCAGACTGTGAAACTGCGAATGGCTCATTAAATCAGTTATAGTTTGTTTGATGGTA | 52 (0.000997%) | | | CTTGTTAGAAGACACAAAGCCAAAGACTCATATGGACTTTGGCTACACCATGAAAGCTTTGAGAAGCAAGAAGAAGGTTGGTTAGTGTTTTGGAGTCGAA | 116 (0.002223%) | | | CTTGTTAGTTTCTTTTCCTCCGCTTATTGATATGCTTAAACTCAGCGGGTAATCCCGCCTGACCTGGGGTCGCTATATGGACTTTGGGTCATCTACAGCT | 24 (0.000460%) | | | CTTTCACGGTTCGTATTCGTACTGAAAATCAGAATCAAACGAGCTTTTACCCTTTTGTTCCACACGAGATTTCTGTTCTCGTTGAGCTCATCTTAGGACA | 40 (0.000767%) | | | CTTTCCAACGAAGCACGCCCATCCAACCTAGGCGAGACAAGGGTTCACATTTCGTTCATCACCCTTGGCCGGCTATCGAACAGCCGGACTCCCATCAAAA | 15 (0.000287%) | | | CTTTGCTGATGCGGGACGGAAGCTGGTCTCCCGTGTGTTACCGCACGCGGTTGGCCTAAATCCGAGCCAAGGACGCCTGGAGCGTACCGACATGCGGTGG | 69 (0.001322%) | | | CTTTTATCTAATAAATGCGTCCCTTCCATAAGTCGGGGTTTGTTGCACGTATTAGCTCTAGAATTACTACGGTTATCCGAGTAGTAGTTACCATCAAACA | 76 (0.001457%) | | | CTTTTCCTCCGCTTATTGATATGCTTAAACTCAGCGGGTAATCCCGCCTGACCTGGGGTCGCTATATGGACTTTGGGTCATCTACAGCTTCCGGACAAGA | 48 (0.000920%) | | | GAAAATCAGAATCAAACGAGCTTTTACCCTTTTGTTCCACACGAGATTTCTGTTCTCGTTGAGCTCATCTTAGGACACCTGCGTTATCTTTTAACAGATG | 13 (0.000249%) | | | GAAACTGCGAATGGCTCATTAAATCAGTTATAGTTTGTTTGATGGTAACTACTACTCGGATAACCGTAGTAATTCTAGAGCTAATACGTGCAACAAACCC | 176 (0.003373%) | | | GAAAGAAGACCCTGTTGAGCTTGACTCTAGTCCGACTTTGTGAAATGACTTGAGAGGTGTAGGATAAGTGGGAGCTTCGGCGCAAGTGAAATACCACTAC | 33 (0.000632%) | | | GAAAGAGCCCGGTATTGTTATTTATTGTCACTACCTCCCCGTGTCAGGATTGGGTAATTTGCGCGCCTGCTGCCTTCCTTGGATGTGGTAGCCGTTTCTC | 29 (0.000556%) | | | GAAATCCGCTAAGGAGTGTGTAACAACTCACCTGCCGAATCAACTAGCCCCGAAAATGGATGGCGCTTAAGCGCGCGACCTATACCCGGCCGTCGGGGCA | 25 (0.000479%) | | | GAAATCCTATGATGTTATCCCATGCTAATGTATCCAGAGCGTAGGCTTGCTTTGAGCACTCTAATTTCTTCAAAGTAACAGCGCCGGAGGCACGACCCGG | 216 (0.004140%) | | | GAAGACACAAAGCCAAAGACTCATATGGACTTTGGCTACACCATGAAAGCTTTGAGAAGCAAGAAGAAGGTTGGTTAGTGTTTTGGAGTCGAATATGACT | 36 (0.000690%) | | | GAAGACCCTGTTGAGCTTGACTCTAGTCCGACTTTGTGAAATGACTTGAGAGGTGTAGGATAAGTGGGAGCTTCGGCGCAAGTGAAATACCACTACTTTT | 14 (0.000268%) | | | GAAGCACGCCCATCCAACCTAGGCGAGACAAGGGTTCACATTTCGTTCATCACCCTTGGCCGGCTATCGAACAGCCGGACTCCCATCAAAAGATGGTTGC | 12 (0.000230%) | | | GAAGCACGCCCATCCAACCTAGGCGAGACAAGGGTTCACATTTCGTTCATCACCCTTGGCCGGCTTTCGAACAGCCGGACTCCCATCAAAAGATGGTTGC | 12 (0.000230%) | | | GAAGTCGAAATCCGCTAAGGAGTGTGTAACAACTCACCTGCCGAATCAACTAGCCCCGAAAATGGATGGCGCTTAAGCGCGCGACCTATACCCGGCCGTC | 3 (0.000057%) | | | GAATATGACTTGATGTCATGTGTATGATTGAGTATAAGAA | 432 (0.003312%) | | | GAATCAAACGAGCTTTTACCCTTTTGTTCCACACGAGATTTCTGTTCTCGTTGAGCTCATCTTAGGACACCTGCGTTATCTTTTAACAGATGTGCCGCCC | 3 (0.000057%) | | | GAATCATCAGAGCAACGGGCAGAGCCCGCGTCGACCTTTTATCTAATAAATGCGTCCCTTCCATAAGTCGGGGTTTGTTGCACGTATTAGCTCTAGAATT | 106 (0.002031%) | | | GAATCCTTGTTAGTTTCTTTTCCTCCGCTTATTGATATGCTTAAACTCAGCGGGTAATCCCGCCTGACCTGGGGTCGCTATATGGACTTTGGGTCATCTA | 12 (0.000230%) | | | GAATGCCAGCCGTTCGTTTGCATGTTCCTTGACACTTTTCGTGCCGGGGTTTTGTGATATCCGGAAGCAACGCGCACGACAAGACCGAGATAAAAGCTCC | 13 (0.000249%) | | | GAATGGCTCATTAAATCAGTTATAGTTTGTTTGATGGTAACTACTACTCGGATAACCGTAGTAATTCTAGAGCTAATACGTGCAACAAACCCCGACTTAT | 6 (0.000115%) | | | GAATTAGGGTTCGATTCCGGAGAGGGAGCCTGAGAAACGGCTACCACATCCAAGGAAGGCAGCAGGCGCGCAAATTACCCAATCCTGACACGGGGAGGTA | 7 (0.000134%) | | | GAATTGTATCCTTGTTAGAAGACACAAAGCCAAAGACTCATATGGACTTTGGCTACACCATGAAAGCTTTGAGAAGCAAGAAGAAGGTTGGTTAGTGTTT | 17 (0.000326%) | | | GACAAGGGTTCACATTTCGTTCATCACCCTTGGCCGGCTATCGAACAGCCGGACTCCCATCAAAAGATGGTTGCCAAGAACATCTTCGTTACGGTTTGCT | 12 (0.000230%) | | | GACAAGGGTTCACATTTCGTTCATCACCCTTGGCCGGCTTTCGAACAGCCGGACTCCCATCAAAAGATGGTTGCCAAGAACATCTTCGTTACGGTTTGCT | 7 (0.000134%) | | | GACACAAAGCCAAAGACTCATATGGACTTTGGCTACACCATGAAAGCTTTGAGAAGCAAGAAGAAGGTTGGTTAGTGTTTTGGAGTCGAATATGACTTGA | 59 (0.001131%) | | | GACAGTCGGGGGCATTCGTATTTCATAGTCAGAGGTGAAATTCTTGGATTTATGAAAGACGAACAACTGCGAAAGCATTTGCCAAGGATGTTTTCATTAA | 4 (0.000077%) | | | GACATCAAGTCATATTCGACTCCAAAACACTAACCAACCTTCTTCTTGCTTCTCAAAGCTTTCATGGTGTAGCCAAAGTCCATATGAGTCTTTGGCTTTG | 53 (0.001016%) | | | GACATTGTCAGGTGGGGAGTTTGGCTGGGGCGGCACATCTGTTAAAAGATAACGCAGGTGTCCTAAGATGAGCTCAACGAGAACAGAAATCTCGTGTGGA | 38 (0.000728%) | | | GACCAGAGGCTGTTCACCTTGGAGACCTGATGCGGTTATGAGTACGACCGGGCGTGAGCGGCACTCGGTCCTCCGGATTTTCAAGGGCCGCCGGGGGCGC | 5 (0.000096%) | | | GACCGGATTGCTCCGTTCCGCATCCGACCAGGACGCATCGCCGGCCCCCATCCGCTTCCCTCCCGACAATTTCAAGCACTCTTTGACTCTCTTTTCAAAG | 12 (0.000230%) | | | GACCTATACCCGGCCGTCGGGGCAAGAGCCAGGCCTCGATGAGTAGGAGGGCGCGGCGGTCGCTGCAAAACCTAGGGCGCGAGCCCGGGCGGAGCGGCCG | 11 (0.000211%) | | | GACCTTTTATCTAATAAATGCGTCCCTTCCATAAGTCGGGGTTTGTTGCACGTATTAGCTCTAGAATTACTACGGTTATCCGAGTAGTAGTTACCATCAA | 8 (0.000153%) | | | GACGGAGAATTAGGGTTCGATTCCGGAGAGGGAGCCTGAGAAACGGCTACCACATCCAAGGAAGGCAGCAGGCGCGCAAATTACCCAATCCTGACACGGG | 4 (0.000077%) | | | GACGGGCGGTGTGTACAAAGGGCAGGGACGTAGTCAACGCGAGCTGATGACTCGCGCTTACTAGGAATTCCTCGTTGAAGACCAACAATTGCAATGATCG | 13 (0.000249%) | | | GACGTGGGTGGTTCGCCGCCCGCGACGTCGCGAGAAGTCCACTAAACCTTATCATTTAGAGGAAGGAGAAGTCGTAACAAGGTTTCCGTAGGTGAACCTG | 53 (0.001016%) | | | GACTCATATGGACTTTGGCTACACCATGAAAGCTTTGAGAAGCAAGAAGAAGGTTGGTTAGTGTTTTGGAGTCGAATATGACTTGATGTCATGTGTATGA | 7 (0.000134%) | | | GACTCCAAAACACTAACCAACCTTCTTCTTGCTTCTCAAAGCTTTCATGGTGTAGCCAAAGTCCATATGAGTCTTTGGCTTTGTGTCTTCTAACAAGGAA | 245 (0.004695%) | | | GACTTCCCTTGCCTACATTGTTCCATCGACCAGAGGCTGTTCACCTTGGAGACCTGATGCGGTTATGAGTACGACCGGGCGTGAGCGGCACTCGGTCCTC | 14 (0.000268%) | | | GACTTGATGTCATGTGTATGATTGAGTATAAGAACTTAAA | 130 (0.000997%) | | | GACTTTGGCTACACCATGAAAGCTTTGAGAAGCAAGAAGAAGGTTGGTTAGTGTTTTGGAGTCGAATATGACTTGATGTCATGTGTATGATTGAGTATAA | 47 (0.000901%) | | | GAGAATTAGGGTTCGATTCCGGAGAGGGAGCCTGAGAAACGGCTACCACATCCAAGGAAGGCAGCAGGCGCGCAAATTACCCAATCCTGACACGGGGAGG | 4 (0.000077%) | | | GAGACAAGGGTTCACATTTCGTTCATCACCCTTGGCCGGCTATCGAACAGCCGGACTCCCATCAAAAGATGGTTGCCAAGAACATCTTCGTTACGGTTTG | 12 (0.000230%) | | | GAGACAAGGGTTCACATTTCGTTCATCACCCTTGGCCGGCTTTCGAACAGCCGGACTCCCATCAAAAGATGGTTGCCAAGAACATCTTCGTTACGGTTTG | 3 (0.000057%) | | | GAGCAACGGGCAGAGCCCGCGTCGACCTTTTATCTAATAAATGCGTCCCTTCCATAAGTCGGGGTTTGTTGCACGTATTAGCTCTAGAATTACTACGGTT | 18 (0.000345%) | | | GAGCCCGCGTCGACCTTTTATCTAATAAATGCGTCCCTTCCATAAGTCGGGGTTTGTTGCACGTATTAGCTCTAGAATTACTACGGTTATCCGAGTAGTA | 7 (0.000134%) | | | GAGCCCGGTATTGTTATTTATTGTCACTACCTCCCCGTGTCAGGATTGGGTAATTTGCGCGCCTGCTGCCTTCCTTGGATGTGGTAGCCGTTTCTCAGGC | 3 (0.000057%) | | | GAGCGACGGGCGGTGTGTACAAAGGGCAGGGACGTAGTCAACGCGAGCTGATGACTCGCGCTTACTAGGAATTCCTCGTTGAAGACCAACAATTGCAATG | 11 (0.000211%) | | | GAGCGAGGTGTGAGTGTCGCCCATGGGCATCGACACCTTGCGGCTAGGAACTGGAACGAGACGGGTAGCAAAGATTTCGAGTAGCACTTCATACTACCGT | 6 (0.000115%) | | | GAGCGAGGTGTGAGTGTCGCCCATGGGCATCGACACCTTGCGGCTAGGAACTGGAACGAGACGGGTGGCAAAGATTTCGAGTAGCACTTCATACTACCGT | 7 (0.000134%) | | | GAGGACATTGTCAGGTGGGGAGTTTGGCTGGGGCGGCACATCTGTTAAAAGATAACGCAGGTGTCCTAAGATGAGCTCAACGAGAACAGAAATCTCGTGT | 76 (0.001457%) | | | GAGGCGAATGCCAGCCGTTCGTTTGCATGTTCCTTGACACTTTTCGTGCCGGGGTTTTGTGATATCCGGAAGCAACGCGCACGACAAGACCGAGATAAAA | 8 (0.000153%) | | | GAGGGCAAGTCTGGTGCCAGCAGCCGCGGTAATTCCAGCTCCAATAGCGTATATTTAAGTTGTTGCAGTTAAAAAGCTCGTAGTTGAACCTTGGGATGGG | 14 (0.000268%) | | | GAGGTGTGAGTGTCGCCCATGGGCATCGACACCTTGCGGCTAGGAACTGGAACGAGACGGGTAGCAAAGATTTCGAGTAGCACTTCATACTACCGTGGGT | 8 (0.000153%) | | | GAGGTGTGAGTGTCGCCCATGGGCATCGACACCTTGCGGCTAGGAACTGGAACGAGACGGGTGGCAAAGATTTCGAGTAGCACTTCATACTACCGTGGGT | 7 (0.000134%) | | | GAGTGTCGCCCATGGGCATCGACACCTTGCGGCTAGGAACTGGAACGAGACGGGTAGCAAAGATTTCGAGTAGCACTTCATACTACCGTGGGTTTTTTAA | 4 (0.000077%) | | | GAGTGTCGCCCATGGGCATCGACACCTTGCGGCTAGGAACTGGAACGAGACGGGTGGCAAAGATTTCGAGTAGCACTTCATACTACCGTGGGTTTTTTAA | 4 (0.000077%) | | | GAGTGTGAGCGAGGTGTGAGTGTCGCCCATGGGCATCGACACCTTGCGGCTAGGAACTGGAACGAGACGGGTGGCAAAGATTTCGAGTAGCACTTCATAC | 4 (0.000077%) | | | GAGTTATCATGAATCATCAGAGCAACGGGCAGAGCCCGCGTCGACCTTTTATCTAATAAATGCGTCCCTTCCATAAGTCGGGGTTTGTTGCACGTATTAG | 78 (0.001495%) | | | GAGTTTGGCTGGGGCGGCACATCTGTTAAAAGATAACGCAGGTGTCCTAAGATGAGCTCAACGAGAACAGAAATCTCGTGTGGAACAAAAGGGTAAAAGC | 6 (0.000115%) | | | GATAGGCCACGCTTTCACGGTTCGTATTCGTACTGAAAATCAGAATCAAACGAGCTTTTACCCTTTTGTTCCACACGAGATTTCTGTTCTCGTTGAGCTC | 1713 (0.032829%) | | | GATAGTGGCCTACCATGGTGGTAACGGGTGACGGAGAATTAGGGTTCGATTCCGGAGAGGGAGCCTGAGAAACGGCTACCACATCCAAGGAAGGCAGCAG | 3 (0.000057%) | | | GATCAAGTTCACCACCGCATGTCGGTACGCTCCAGGCGTCCTTGGCTCGGATTTAGGCCAACCGCGTGCGGTAACACACGGGAGACCAGCTTCCGTCCCG | 47 (0.000901%) | | | GATCACGGCAATTCCCCGCCACATCCTCTCAAACGCAATGGAAAGAGAGAAAGGACGAGGTCTTGACCGTCATCTTTTGCCCGAAGGACGGATGAGCTTT | 31 (0.000594%) | | | GATCATTCAATCGGTAGGAGCGACGGGCGGTGTGTACAAAGGGCAGGGACGTAGTCAACGCGAGCTGATGACTCGCGCTTACTAGGAATTCCTCGTTGAA | 762 (0.014603%) | | | GATCCATGCTTTCCAACGAAGCACGCCCATCCAACCTAGGCGAGACAAGGGTTCACATTTCGTTCATCACCCTTGGCCGGCTATCGAACAGCCGGACTCC | 78 (0.001495%) | | | GATCCATTGGAGGGCAAGTCTGGTGCCAGCAGCCGCGGTAATTCCAGCTCCAATAGCGTATATTTAAGTTGTTGCAGTTAAAAAGCTCGTAGTTGAACCT | 1486 (0.028478%) | | | GATCCGCCTAGGCTGTCCCGAGTGTGAGCGAGGTGTGAGTGTCGCCCATGGGCATCGACACCTTGCGGCTAGGAACTGGAACGAGACGGGTAGCAAAGAT | 96 (0.001840%) | | | GATCCGCCTAGGCTGTCCCGAGTGTGAGCGAGGTGTGAGTGTCGCCCATGGGCATCGACACCTTGCGGCTAGGAACTGGAACGAGACGGGTGGCAAAGAT | 142 (0.002721%) | | | GATCCGGTTAAAATTCCGGAACCGGGACGTGGCGGTTGACGGCAACGTTAGGGAGTCCGGAGACGTCGGCGGGGGCCTCGGGAAGAGTTATCTTTTCTGT | 145 (0.002779%) | | | GATCCGTAACTTCGGGAAAAGGATTGGCTCTGAGGGCTGGGCTCGGGGGTCCCAGTTCCGAACCCGTCGGCTGTCAGCGGACTGCTCGAGCTGCTTCCGC | 21 (0.000402%) | | | GATCCGTCGAGTTATCATGAATCATCAGAGCAACGGGCAGAGCCCGCGTCGACCTTTTATCTAATAAATGCGTCCCTTCCATAAGTCGGGGTTTGTTGCA | 123 (0.002357%) | | | GATCCTCGTTAAGGGATTTAGATTGTACTCATTCCAATTACCAGACTCGAAAGAGCCCGGTATTGTTATTTATTGTCACTACCTCCCCGTGTCAGGATTG | 88 (0.001686%) | | | GATCGAAATCCTATGATGTTATCCCATGCTAATGTATCCAGAGCGTAGGCTTGCTTTGAGCACTCTAATTTCTTCAAAGTAACAGCGCCGGAGGCACGAC | 354 (0.006784%) | | | GATCGATCAA | 145668 (0.279165%) | | | GATCGATCAAGTTCACCACCGCATGTCGGTACGCTCCAGGCGTCCTTGGCTCGGATTTAGGCCAACCGCGTGCGGTAACACACGGGAGACCAGCTTCCGT | 1310 (0.025105%) | | | GATCGATCACGGCAATTCCCCGCCACATCCTCTCAAACGCAATGGAAAGAGAGAAAGGACGAGGTCTTGACCGTCATCTTTTGCCCGAAGGACGGATGAG | 933 (0.017880%) | | | GATCGATCCATGCTTTCCAACGAAGCACGCCCATCCAACCTAGGCGAGACAAGGGTTCACATTTCGTTCATCACCCTTGGCCGGCTATCGAACAGCCGGA | 1366 (0.026179%) | | | GATCGATCCGCCTAGGCTGTCCCGAGTGTGAGCGAGGTGTGAGTGTCGCCCATGGGCATCGACACCTTGCGGCTAGGAACTGGAACGAGACGGGTAGCAA | 1052 (0.020161%) | | | GATCGATCCGCCTAGGCTGTCCCGAGTGTGAGCGAGGTGTGAGTGTCGCCCATGGGCATCGACACCTTGCGGCTAGGAACTGGAACGAGACGGGTGGCAA | 1547 (0.029647%) | | | GATCGATCCGGTTAAAATTCCGGAACCGGGACGTGGCGGTTGACGGCAACGTTAGGGAGTCCGGAGACGTCGGCGGGGGCCTCGGGAAGAGTTATCTTTT | 1624 (0.031123%) | | | GATCGATCCGTAACTTCGGGAAAAGGATTGGCTCTGAGGGCTGGGCTCGGGGGTCCCAGTTCCGAACCCGTCGGCTGTCAGCGGACTGCTCGAGCTGCTT | 1344 (0.025757%) | | | GATCGATCCGTCGAGTTATCATGAATCATCAGAGCAACGGGCAGAGCCCGCGTCGACCTTTTATCTAATAAATGCGTCCCTTCCATAAGTCGGGGTTTGT | 1141 (0.021867%) | | | GATCGATCCTCGTTAAGGGATTTAGATTGTACTCATTCCAATTACCAGACTCGAAAGAGCCCGGTATTGTTATTTATTGTCACTACCTCCCCGTGTCAGG | 1505 (0.028843%) | | | GATCGATCGAAATCCTATGATGTTATCCCATGCTAATGTATCCAGAGCGTAGGCTTGCTTTGAGCACTCTAATTTCTTCAAAGTAACAGCGCCGGAGGCA | 1296 (0.024837%) | | | GATCGATCGGCGGACCGGATTGCTCCGTTCCGCATCCGACCAGGACGCATCGCCGGCCCCCATCCGCTTCCCTCCCGACAATTTCAAGCACTCTTTGACT | 907 (0.017382%) | | | GATCGATCGGGTTGCGGTTTAAGTTGTTATACTCAATCAT | 4328 (0.033178%) | | | GATCGATCTT | 165258 (0.316708%) | | | GATCGGCGGACCGGATTGCTCCGTTCCGCATCCGACCAGGACGCATCGCCGGCCCCCATCCGCTTCCCTCCCGACAATTTCAAGCACTCTTTGACTCTCT | 9 (0.000172%) | | | GATCGGGTTGCGGTTTAAGTTGTTATACTCAATCATACAC | 1247 (0.009559%) | | | GATCTTAAAGGCGTAAGAATTGTATCCTTGTTAAAAGACACAAAGCCAAAGACTCATATGGACTTTGGCTACACCATGAAAGCTTTGAGAAGCAAGAAGA | 696 (0.013338%) | | | GATCTTAAAGGCGTAAGAATTGTATCCTTGTTAGAAGACACAAAGCCAAAGACTCATATGGACTTTGGCTACACCATGAAAGCTTTGAGAAGCAAGAAGA | 521 (0.009985%) | | | GATGCCCTTAGATGTTCTGGGCCGCACGCGCGCTACACTGATGTATTCAACGAGTTCACACCTTGGCCGACAGGCCCGGGTAATCTTTGAAATTTCATCG | 9 (0.000172%) | | | GATGTCATGTGTATGATTGAGTATAAGAACTTAAACCGCA | 128 (0.000981%) | | | GATGTTATCCCATGCTAATGTATCCAGAGCGTAGGCTTGCTTTGAGCACTCTAATTTCTTCAAAGTAACAGCGCCGGAGGCACGACCCGGCCAATTAAGA | 12 (0.000230%) | | | GATGTTCTGGGCCGCACGCGCGCTACACTGATGTATTCAACGAGTTCACACCTTGGCCGACAGGCCCGGGTAATCTTTGAAATTTCATCGTGATGGGGAT | 12 (0.000230%) | | | GATTAACAGGGACAGTCGGGGGCATTCGTATTTCATAGTCAGAGGTGAAATTCTTGGATTTATGAAAGACGAACAACTGCGAAAGCATTTGCCAAGGATG | 183 (0.003507%) | | | GATTAAGCCATGCATGTGTAAGTATGAACGAATTCAGACTGTGAAACTGCGAATGGCTCATTAAATCAGTTATAGTTTGTTTGATGGTAACTACTACTCG | 22 (0.000422%) | | | GATTGTACTCATTCCAATTACCAGACTCGAAAGAGCCCGGTATTGTTATTTATTGTCACTACCTCCCCGTGTCAGGATTGGGTAATTTGCGCGCCTGCTG | 7 (0.000134%) | | | GATTTAGATTGTACTCATTCCAATTACCAGACTCGAAAGAGCCCGGTATTGTTATTTATTGTCACTACCTCCCCGTGTCAGGATTGGGTAATTTGCGCGC | 19 (0.000364%) | | | GCAACCGGATCTTAAAGGCGTAAGAATTGTATCCTTGTTAGAAGACACAAAGCCAAAGACTCATATGGACTTTGGCTACACCATGAAAGCTTTGAGAAGC | 5 (0.000096%) | | | GCAACGGGCAGAGCCCGCGTCGACCTTTTATCTAATAAATGCGTCCCTTCCATAAGTCGGGGTTTGTTGCACGTATTAGCTCTAGAATTACTACGGTTAT | 45 (0.000862%) | | | GCAACGTCGCTATGAACGCTTGGCTGCCACAAGCCAGTTATCCCTGTGGTAACTTTTCTGACACCTCTAGCTTCAAATTCCGAAGGTCTAAAGGATCGAT | 73 (0.001399%) | | | GCAAGTCTGGTGCCAGCAGCCGCGGTAATTCCAGCTCCAATAGCGTATATTTAAGTTGTTGCAGTTAAAAAGCTCGTAGTTGAACCTTGGGATGGGTCGG | 18 (0.000345%) | | | GCAATAACAGGTCTGTGATGCCCTTAGATGTTCTGGGCCGCACGCGCGCTACACTGATGTATTCAACGAGTTCACACCTTGGCCGACAGGCCCGGGTAAT | 1018 (0.019509%) | | | GCAATTCCCCGCCACATCCTCTCAAACGCAATGGAAAGAGAGAAAGGACGAGGTCTTGACCGTCATCTTTTGCCCGAAGGACGGATGAGCTTTGGCGGGA | 30 (0.000575%) | | | GCACATCTGTTAAAAGATAACGCAGGTGTCCTAAGATGAGCTCAACGAGAACAGAAATCTCGTGTGGAACAAAAGGGTAAAAGCTCGTTTGATTCTGATT | 6 (0.000115%) | | | GCACGCCCATCCAACCTAGGCGAGACAAGGGTTCACATTTCGTTCATCACCCTTGGCCGGCTATCGAACAGCCGGACTCCCATCAAAAGATGGTTGCCAA | 39 (0.000747%) | | | GCACGCCCATCCAACCTAGGCGAGACAAGGGTTCACATTTCGTTCATCACCCTTGGCCGGCTTTCGAACAGCCGGACTCCCATCAAAAGATGGTTGCCAA | 27 (0.000517%) | | | GCACGCGCCTAACGGCGTGCCTCGGCATCAGCGTGCTCCGGGCGTCGGCCTGTGGGCTCCCCATTCGACCCGTCTTGAAACACGGACCAAGGAGTCTGAC | 19 (0.000364%) | | | GCAGAGCCCGCGTCGACCTTTTATCTAATAAATGCGTCCCTTCCATAAGTCGGGGTTTGTTGCACGTATTAGCTCTAGAATTACTACGGTTATCCGAGTA | 26 (0.000498%) | | | GCAGCACGCGCCTAACGGCGTGCCTCGGCATCAGCGTGCTCCGGGCGTCGGCCTGTGGGCTCCCCATTCGACCCGTCTTGAAACACGGACCAAGGAGTCT | 425 (0.008145%) | | | GCAGCCGCGGTAATTCCAGCTCCAATAGCGTATATTTAAGTTGTTGCAGTTAAAAAGCTCGTAGTTGAACCTTGGGATGGGTCGGCCGGTCCGCCTTTGG | 15 (0.000287%) | | | GCATCAGCGTGCTCCGGGCGTCGGCCTGTGGGCTCCCCATTCGACCCGTCTTGAAACACGGACCAAGGAGTCTGACATGTGTGCGAGTCAACGGGTGAGT | 10 (0.000192%) | | | GCATCATAAGGATACTAAATCCTATTTTCTGGTAAATTTTCATAATTTTTTGACACCTCTAGCTAGGTCATTTGACCTGATACAACATCGGATTTTCATG | 9 (0.000172%) | | | GCATGTGTAAGTATGAACGAATTCAGACTGTGAAACTGCGAATGGCTCATTAAATCAGTTATAGTTTGTTTGATGGTAACTACTACTCGGATAACCGTAG | 7 (0.000134%) | | | GCCAAAGACTCATATGGACTTTGGCTACACCATGAAAGCTTTGAGAAGCAAGAAGAAGGTTGGTTAGTGTTTTGGAGTCGAATATGACTTGATGTCATGT | 57 (0.001092%) | | | GCCACATCCTCTCAAACGCAATGGAAAGAGAGAAAGGACGAGGTCTTGACCGTCATCTTTTGCCCGAAGGACGGATGAGCTTTGGCGGGACTGAATCACT | 25 (0.000479%) | | | GCCACCCTAACGCCTCGAAGAACTAATGGCAGCCCACGCAAGGCAAGCCCATTCTCCTCGACGATTCAGCAGTTTTTGTCCGAGAACTGCTGAGAAAACT | 881 (0.016884%) | | | GCCACGCTTTCACGGTTCGTATTCGTACTGAAAATCAGAATCAAACGAGCTTTTACCCTTTTGTTCCACACGAGATTTCTGTTCTCGTTGAGCTCATCTT | 127 (0.002434%) | | | GCCAGCAGCCGCGGTAATTCCAGCTCCAATAGCGTATATTTAAGTTGTTGCAGTTAAAAAGCTCGTAGTTGAACCTTGGGATGGGTCGGCCGGTCCGCCT | 27 (0.000517%) | | | GCCAGCCGTTCGTTTGCATGTTCCTTGACACTTTTCGTGCCGGGGTTTTGTGATATCCGGAAGCAACGCGCACGACAAGACCGAGATAAAAGCTCCCGAT | 28 (0.000537%) | | | GCCAGTAGTCATATGCTTGTCTCAAAGATTAAGCCATGCATGTGTAAGTATGAACGAATTCAGACTGTGAAACTGCGAATGGCTCATTAAATCAGTTATA | 263 (0.005040%) | | | GCCATGCATGTGTAAGTATGAACGAATTCAGACTGTGAAACTGCGAATGGCTCATTAAATCAGTTATAGTTTGTTTGATGGTAACTACTACTCGGATAAC | 14 (0.000268%) | | | GCCCATCCAACCTAGGCGAGACAAGGGTTCACATTTCGTTCATCACCCTTGGCCGGCTATCGAACAGCCGGACTCCCATCAAAAGATGGTTGCCAAGAAC | 53 (0.001016%) | | | GCCCATCCAACCTAGGCGAGACAAGGGTTCACATTTCGTTCATCACCCTTGGCCGGCTTTCGAACAGCCGGACTCCCATCAAAAGATGGTTGCCAAGAAC | 36 (0.000690%) | | | GCCCATGGGCATCGACACCTTGCGGCTAGGAACTGGAACGAGACGGGTGGCAAAGATTTCGAGTAGCACTTCATACTACCGTGGGTTTTTTAAACCTTCC | 16 (0.000307%) | | | GCCCGCGTCGACCTTTTATCTAATAAATGCGTCCCTTCCATAAGTCGGGGTTTGTTGCACGTATTAGCTCTAGAATTACTACGGTTATCCGAGTAGTAGT | 5 (0.000096%) | | | GCCCTTAGATGTTCTGGGCCGCACGCGCGCTACACTGATGTATTCAACGAGTTCACACCTTGGCCGACAGGCCCGGGTAATCTTTGAAATTTCATCGTGA | 16 (0.000307%) | | | GCCGACTTCCCTTGCCTACATTGTTCCATCGACCAGAGGCTGTTCACCTTGGAGACCTGATGCGGTTATGAGTACGACCGGGCGTGAGCGGCACTCGGTC | 218 (0.004178%) | | | GCCGCCCGCGACGTCGCGAGAAGTCCACTAAACCTTATCATTTAGAGGAAGGAGAAGTCGTAACAAGGTTTCCGTAGGTGAACCTGCGGAAGGATCGATC | 14 (0.000268%) | | | GCCGCGGTAATTCCAGCTCCAATAGCGTATATTTAAGTTGTTGCAGTTAAAAAGCTCGTAGTTGAACCTTGGGATGGGTCGGCCGGTCCGCCTTTGGTGT | 7 (0.000134%) | | | GCCGTCGGGGCAAGAGCCAGGCCTCGATGAGTAGGAGGGCGCGGCGGTCGCTGCAAAACCTAGGGCGCGAGCCCGGGCGGAGCGGCCGTCGGTGCAGATC | 7 (0.000134%) | | | GCCGTTACTAAGGGAATCCTTGTTAGTTTCTTTTCCTCCGCTTATTGATATGCTTAAACTCAGCGGGTAATCCCGCCTGACCTGGGGTCGCTATATGGAC | 16 (0.000307%) | | | GCCTAACGGCGTGCCTCGGCATCAGCGTGCTCCGGGCGTCGGCCTGTGGGCTCCCCATTCGACCCGTCTTGAAACACGGACCAAGGAGTCTGACATGTGT | 48 (0.000920%) | | | GCCTAAGTAGTGTTTCCTTGTTAGAAGACACAAAGCCAAAGACTCATATGGACTTTGGCTACACCATGAAAGCTTTGAGAAGCAAGAAGAAGGTTGGTTA | 38 (0.000728%) | | | GCCTAGGCTGTCCCGAGTGTGAGCGAGGTGTGAGTGTCGCCCATGGGCATCGACACCTTGCGGCTAGGAACTGGAACGAGACGGGTAGCAAAGATTTCGA | 257 (0.004925%) | | | GCCTAGGCTGTCCCGAGTGTGAGCGAGGTGTGAGTGTCGCCCATGGGCATCGACACCTTGCGGCTAGGAACTGGAACGAGACGGGTGGCAAAGATTTCGA | 378 (0.007244%) | | | GCCTCGAAGAACTAATGGCAGCCCACGCAAGGCAAGCCCATTCTCCTCGACGATTCAGCAGTTTTTGTCCGAGAACTGCTGAGAAAACTCGGAAAAAGGC | 14 (0.000268%) | | | GCCTCGGCATCAGCGTGCTCCGGGCGTCGGCCTGTGGGCTCCCCATTCGACCCGTCTTGAAACACGGACCAAGGAGTCTGACATGTGTGCGAGTCAACGG | 10 (0.000192%) | | | GCGAATGCCAGCCGTTCGTTTGCATGTTCCTTGACACTTTTCGTGCCGGGGTTTTGTGATATCCGGAAGCAACGCGCACGACAAGACCGAGATAAAAGCT | 30 (0.000575%) | | | GCGAATGGCTCATTAAATCAGTTATAGTTTGTTTGATGGTAACTACTACTCGGATAACCGTAGTAATTCTAGAGCTAATACGTGCAACAAACCCCGACTT | 20 (0.000383%) | | | GCGACCTATACCCGGCCGTCGGGGCAAGAGCCAGGCCTCGATGAGTAGGAGGGCGCGGCGGTCGCTGCAAAACCTAGGGCGCGAGCCCGGGCGGAGCGGC | 84 (0.001610%) | | | GCGACGGGCGGTGTGTACAAAGGGCAGGGACGTAGTCAACGCGAGCTGATGACTCGCGCTTACTAGGAATTCCTCGTTGAAGACCAACAATTGCAATGAT | 44 (0.000843%) | | | GCGACGTGGGTGGTTCGCCGCCCGCGACGTCGCGAGAAGTCCACTAAACCTTATCATTTAGAGGAAGGAGAAGTCGTAACAAGGTTTCCGTAGGTGAACC | 2275 (0.043599%) | | | GCGAGACAAGGGTTCACATTTCGTTCATCACCCTTGGCCGGCTATCGAACAGCCGGACTCCCATCAAAAGATGGTTGCCAAGAACATCTTCGTTACGGTT | 41 (0.000786%) | | | GCGAGACAAGGGTTCACATTTCGTTCATCACCCTTGGCCGGCTTTCGAACAGCCGGACTCCCATCAAAAGATGGTTGCCAAGAACATCTTCGTTACGGTT | 32 (0.000613%) | | | GCGAGGTGTGAGTGTCGCCCATGGGCATCGACACCTTGCGGCTAGGAACTGGAACGAGACGGGTAGCAAAGATTTCGAGTAGCACTTCATACTACCGTGG | 31 (0.000594%) | | | GCGAGGTGTGAGTGTCGCCCATGGGCATCGACACCTTGCGGCTAGGAACTGGAACGAGACGGGTGGCAAAGATTTCGAGTAGCACTTCATACTACCGTGG | 53 (0.001016%) | | | GCGCCTAACGGCGTGCCTCGGCATCAGCGTGCTCCGGGCGTCGGCCTGTGGGCTCCCCATTCGACCCGTCTTGAAACACGGACCAAGGAGTCTGACATGT | 25 (0.000479%) | | | GCGCGACCTATACCCGGCCGTCGGGGCAAGAGCCAGGCCTCGATGAGTAGGAGGGCGCGGCGGTCGCTGCAAAACCTAGGGCGCGAGCCCGGGCGGAGCG | 34 (0.000652%) | | | GCGCGCGACCTATACCCGGCCGTCGGGGCAAGAGCCAGGCCTCGATGAGTAGGAGGGCGCGGCGGTCGCTGCAAAACCTAGGGCGCGAGCCCGGGCGGAG | 5 (0.000096%) | | | GCGGACCGGATTGCTCCGTTCCGCATCCGACCAGGACGCATCGCCGGCCCCCATCCGCTTCCCTCCCGACAATTTCAAGCACTCTTTGACTCTCTTTTCA | 19 (0.000364%) | | | GCGGAGGACATTGTCAGGTGGGGAGTTTGGCTGGGGCGGCACATCTGTTAAAAGATAACGCAGGTGTCCTAAGATGAGCTCAACGAGAACAGAAATCTCG | 1582 (0.030318%) | | | GCGGCACATCTGTTAAAAGATAACGCAGGTGTCCTAAGATGAGCTCAACGAGAACAGAAATCTCGTGTGGAACAAAAGGGTAAAAGCTCGTTTGATTCTG | 9 (0.000172%) | | | GCGGTAATTCCAGCTCCAATAGCGTATATTTAAGTTGTTGCAGTTAAAAAGCTCGTAGTTGAACCTTGGGATGGGTCGGCCGGTCCGCCTTTGGTGTGCA | 22 (0.000422%) | | | GCGGTTTAAGTTCTTATACTCAATCATACACATGACATCA | 651 (0.004990%) | | | GCGTAAGAATTGTATCCTTGTTAGAAGACACAAAGCCAAAGACTCATATGGACTTTGGCTACACCATGAAAGCTTTGAGAAGCAAGAAGAAGGTTGGTTA | 19 (0.000364%) | | | GCGTAGGCTTGCTTTGAGCACTCTAATTTCTTCAAAGTAACAGCGCCGGAGGCACGACCCGGCCAATTAAGACCAGGAGCGTATCGCCGACCGAAGGGAC | 52 (0.000997%) | | | GCGTCGACCTTTTATCTAATAAATGCGTCCCTTCCATAAGTCGGGGTTTGTTGCACGTATTAGCTCTAGAATTACTACGGTTATCCGAGTAGTAGTTACC | 12 (0.000230%) | | | GCGTCGGCCTGTGGGCTCCCCATTCGACCCGTCTTGAAACACGGACCAAGGAGTCTGACATGTGTGCGAGTCAACGGGTGAGTAAACCCGTAAGGCGCAA | 8 (0.000153%) | | | GCGTGCCTCGGCATCAGCGTGCTCCGGGCGTCGGCCTGTGGGCTCCCCATTCGACCCGTCTTGAAACACGGACCAAGGAGTCTGACATGTGTGCGAGTCA | 9 (0.000172%) | | | GCGTGCTCCGGGCGTCGGCCTGTGGGCTCCCCATTCGACCCGTCTTGAAACACGGACCAAGGAGTCTGACATGTGTGCGAGTCAACGGGTGAGTAAACCC | 12 (0.000230%) | | | GCTAAGGAGTGTGTAACAACTCACCTGCCGAATCAACTAGCCCCGAAAATGGATGGCGCTTAAGCGCGCGACCTATACCCGGCCGTCGGGGCAAGAGCCA | 56 (0.001073%) | | | GCTAATGTATCCAGAGCGTAGGCTTGCTTTGAGCACTCTAATTTCTTCAAAGTAACAGCGCCGGAGGCACGACCCGGCCAATTAAGACCAGGAGCGTATC | 22 (0.000422%) | | | GCTCATTAAATCAGTTATAGTTTGTTTGATGGTAACTACTACTCGGATAACCGTAGTAATTCTAGAGCTAATACGTGCAACAAACCCCGACTTATGGAAG | 12 (0.000230%) | | | GCTCCGGGCGTCGGCCTGTGGGCTCCCCATTCGACCCGTCTTGAAACACGGACCAAGGAGTCTGACATGTGTGCGAGTCAACGGGTGAGTAAACCCGTAA | 3 (0.000057%) | | | GCTCCGTTCCGCATCCGACCAGGACGCATCGCCGGCCCCCATCCGCTTCCCTCCCGACAATTTCAAGCACTCTTTGACTCTCTTTTCAAAGTCCTTTTCA | 5 (0.000096%) | | | GCTCGCCGTTACTAAGGGAATCCTTGTTAGTTTCTTTTCCTCCGCTTATTGATATGCTTAAACTCAGCGGGTAATCCCGCCTGACCTGGGGTCGCTATAT | 49 (0.000939%) | | | GCTGATGCGGGACGGAAGCTGGTCTCCCGTGTGTTACCGCACGCGGTTGGCCTAAATCCGAGCCAAGGACGCCTGGAGCGTACCGACATGCGGTGGTGAA | 10 (0.000192%) | | | GCTGGGGCGGCACATCTGTTAAAAGATAACGCAGGTGTCCTAAGATGAGCTCAACGAGAACAGAAATCTCGTGTGGAACAAAAGGGTAAAAGCTCGTTTG | 11 (0.000211%) | | | GCTGTCCCGAGTGTGAGCGAGGTGTGAGTGTCGCCCATGGGCATCGACACCTTGCGGCTAGGAACTGGAACGAGACGGGTAGCAAAGATTTCGAGTAGCA | 69 (0.001322%) | | | GCTGTCCCGAGTGTGAGCGAGGTGTGAGTGTCGCCCATGGGCATCGACACCTTGCGGCTAGGAACTGGAACGAGACGGGTGGCAAAGATTTCGAGTAGCA | 104 (0.001993%) | | | GCTTAAGCGCGCGACCTATACCCGGCCGTCGGGGCAAGAGCCAGGCCTCGATGAGTAGGAGGGCGCGGCGGTCGCTGCAAAACCTAGGGCGCGAGCCCGG | 1050 (0.020123%) | | | GCTTATTGATATGCTTAAACTCAGCGGGTAATCCCGCCTGACCTGGGGTCGCTATATGGACTTTGGGTCATCTACAGCTTCCGGACAAGAGCGACCGATA | 31 (0.000594%) | | | GCTTCTCAAAGCTTTCATGGTGTAGCCAAAGTCCATATGAGTCTTTGGCTTTGTGTCTTCTAACAAGGAAACACTACTTAGGCTTATAAGATGCGGTTGC | 12 (0.000230%) | | | GCTTGACTCTAGTCCGACTTTGTGAAATGACTTGAGAGGTGTAGGATAAGTGGGAGCTTCGGCGCAAGTGAAATACCACTACTTTTAACGTTATTTTACT | 22 (0.000422%) | | | GCTTGCTTTGAGCACTCTAATTTCTTCAAAGTAACAGCGCCGGAGGCACGACCCGGCCAATTAAGACCAGGAGCGTATCGCCGACCGAAGGGACAAGCCG | 10 (0.000192%) | | | GCTTGTCTCAAAGATTAAGCCATGCATGTGTAAGTATGAACGAATTCAGACTGTGAAACTGCGAATGGCTCATTAAATCAGTTATAGTTTGTTTGATGGT | 18 (0.000345%) | | | GCTTTCACGGTTCGTATTCGTACTGAAAATCAGAATCAAACGAGCTTTTACCCTTTTGTTCCACACGAGATTTCTGTTCTCGTTGAGCTCATCTTAGGAC | 19 (0.000364%) | | | GCTTTCCAACGAAGCACGCCCATCCAACCTAGGCGAGACAAGGGTTCACATTTCGTTCATCACCCTTGGCCGGCTATCGAACAGCCGGACTCCCATCAAA | 65 (0.001246%) | | | GGAAAGAAGACCCTGTTGAGCTTGACTCTAGTCCGACTTTGTGAAATGACTTGAGAGGTGTAGGATAAGTGGGAGCTTCGGCGCAAGTGAAATACCACTA | 747 (0.014316%) | | | GGAAAGAGAGAAAGGACGAGGTCTTGACCGTCATCTTTTGCCCGAAGGACGGATGAGCTTTGGCGGGACTGAATCACTTCGAGTCACCGTCGACAACTTT | 11 (0.000211%) | | | GGAAGTCGAAATCCGCTAAGGAGTGTGTAACAACTCACCTGCCGAATCAACTAGCCCCGAAAATGGATGGCGCTTAAGCGCGCGACCTATACCCGGCCGT | 25 (0.000479%) | | | GGAATCCTTGTTAGTTTCTTTTCCTCCGCTTATTGATATGCTTAAACTCAGCGGGTAATCCCGCCTGACCTGGGGTCGCTATATGGACTTTGGGTCATCT | 30 (0.000575%) | | | GGACAGTCGGGGGCATTCGTATTTCATAGTCAGAGGTGAAATTCTTGGATTTATGAAAGACGAACAACTGCGAAAGCATTTGCCAAGGATGTTTTCATTA | 75 (0.001437%) | | | GGACATTGTCAGGTGGGGAGTTTGGCTGGGGCGGCACATCTGTTAAAAGATAACGCAGGTGTCCTAAGATGAGCTCAACGAGAACAGAAATCTCGTGTGG | 176 (0.003373%) | | | GGACCGGATTGCTCCGTTCCGCATCCGACCAGGACGCATCGCCGGCCCCCATCCGCTTCCCTCCCGACAATTTCAAGCACTCTTTGACTCTCTTTTCAAA | 23 (0.000441%) | | | GGACTTTGGCTACACCATGAAAGCTTTGAGAAGCAAGAAGAAGGTTGGTTAGTGTTTTGGAGTCGAATATGACTTGATGTCATGTGTATGATTGAGTATA | 80 (0.001533%) | | | GGAGAATTAGGGTTCGATTCCGGAGAGGGAGCCTGAGAAACGGCTACCACATCCAAGGAAGGCAGCAGGCGCGCAAATTACCCAATCCTGACACGGGGAG | 88 (0.001686%) | | | GGAGCGACGGGCGGTGTGTACAAAGGGCAGGGACGTAGTCAACGCGAGCTGATGACTCGCGCTTACTAGGAATTCCTCGTTGAAGACCAACAATTGCAAT | 54 (0.001035%) | | | GGAGGACATTGTCAGGTGGGGAGTTTGGCTGGGGCGGCACATCTGTTAAAAGATAACGCAGGTGTCCTAAGATGAGCTCAACGAGAACAGAAATCTCGTG | 450 (0.008624%) | | | GGAGGCGAATGCCAGCCGTTCGTTTGCATGTTCCTTGACACTTTTCGTGCCGGGGTTTTGTGATATCCGGAAGCAACGCGCACGACAAGACCGAGATAAA | 15 (0.000287%) | | | GGAGGGCAAGTCTGGTGCCAGCAGCCGCGGTAATTCCAGCTCCAATAGCGTATATTTAAGTTGTTGCAGTTAAAAAGCTCGTAGTTGAACCTTGGGATGG | 58 (0.001112%) | | | GGAGTTTGGCTGGGGCGGCACATCTGTTAAAAGATAACGCAGGTGTCCTAAGATGAGCTCAACGAGAACAGAAATCTCGTGTGGAACAAAAGGGTAAAAG | 6 (0.000115%) | | | GGATACTAAATCCTATTTTCTGGTAAATTTTCATAATTTTTTGACACCTCTAGCTAGGTCATTTGACCTGATACAACATCGGATTTTCATGGTCTAGTTG | 14 (0.000268%) | | | GGATAGTGGCCTACCATGGTGGTAACGGGTGACGGAGAATTAGGGTTCGATTCCGGAGAGGGAGCCTGAGAAACGGCTACCACATCCAAGGAAGGCAGCA | 728 (0.013952%) | | | GGATCTTAAAGGCGTAAGAATTGTATCCTTGTTAGAAGACACAAAGCCAAAGACTCATATGGACTTTGGCTACACCATGAAAGCTTTGAGAAGCAAGAAG | 17 (0.000326%) | | | GGATTGCTCCGTTCCGCATCCGACCAGGACGCATCGCCGGCCCCCATCCGCTTCCCTCCCGACAATTTCAAGCACTCTTTGACTCTCTTTTCAAAGTCCT | 10 (0.000192%) | | | GGATTTAGATTGTACTCATTCCAATTACCAGACTCGAAAGAGCCCGGTATTGTTATTTATTGTCACTACCTCCCCGTGTCAGGATTGGGTAATTTGCGCG | 5 (0.000096%) | | | GGCAAGTCTGGTGCCAGCAGCCGCGGTAATTCCAGCTCCAATAGCGTATATTTAAGTTGTTGCAGTTAAAAAGCTCGTAGTTGAACCTTGGGATGGGTCG | 66 (0.001265%) | | | GGCAATTCCCCGCCACATCCTCTCAAACGCAATGGAAAGAGAGAAAGGACGAGGTCTTGACCGTCATCTTTTGCCCGAAGGACGGATGAGCTTTGGCGGG | 83 (0.001591%) | | | GGCACATCTGTTAAAAGATAACGCAGGTGTCCTAAGATGAGCTCAACGAGAACAGAAATCTCGTGTGGAACAAAAGGGTAAAAGCTCGTTTGATTCTGAT | 25 (0.000479%) | | | GGCAGAGCCCGCGTCGACCTTTTATCTAATAAATGCGTCCCTTCCATAAGTCGGGGTTTGTTGCACGTATTAGCTCTAGAATTACTACGGTTATCCGAGT | 21 (0.000402%) | | | GGCATCAGCGTGCTCCGGGCGTCGGCCTGTGGGCTCCCCATTCGACCCGTCTTGAAACACGGACCAAGGAGTCTGACATGTGTGCGAGTCAACGGGTGAG | 11 (0.000211%) | | | GGCATCGACACCTTGCGGCTAGGAACTGGAACGAGACGGGTGGCAAAGATTTCGAGTAGCACTTCATACTACCGTGGGTTTTTTAAACCTTCCGAGTTTT | 9 (0.000172%) | | | GGCATGCATCATAAGGATACTAAATCCTATTTTCTGGTAAATTTTCATAATTTTTTGACACCTCTAGCTAGGTCATTTGACCTGATACAACATCGGATTT | 3 (0.000057%) | | | GGCATTCGTATTTCATAGTCAGAGGTGAAATTCTTGGATTTATGAAAGACGAACAACTGCGAAAGCATTTGCCAAGGATGTTTTCATTAATCAAGAACGA | 4 (0.000077%) | | | GGCCACGCTTTCACGGTTCGTATTCGTACTGAAAATCAGAATCAAACGAGCTTTTACCCTTTTGTTCCACACGAGATTTCTGTTCTCGTTGAGCTCATCT | 30 (0.000575%) | | | GGCCGCACGCGCGCTACACTGATGTATTCAACGAGTTCACACCTTGGCCGACAGGCCCGGGTAATCTTTGAAATTTCATCGTGATGGGGATAGATCGATC | 3 (0.000057%) | | | GGCCGTCGGGGCAAGAGCCAGGCCTCGATGAGTAGGAGGGCGCGGCGGTCGCTGCAAAACCTAGGGCGCGAGCCCGGGCGGAGCGGCCGTCGGTGCAGAT | 22 (0.000422%) | | | GGCCTACCATGGTGGTAACGGGTGACGGAGAATTAGGGTTCGATTCCGGAGAGGGAGCCTGAGAAACGGCTACCACATCCAAGGAAGGCAGCAGGCGCGC | 17 (0.000326%) | | | GGCGAATGCCAGCCGTTCGTTTGCATGTTCCTTGACACTTTTCGTGCCGGGGTTTTGTGATATCCGGAAGCAACGCGCACGACAAGACCGAGATAAAAGC | 43 (0.000824%) | | | GGCGAGACAAGGGTTCACATTTCGTTCATCACCCTTGGCCGGCTATCGAACAGCCGGACTCCCATCAAAAGATGGTTGCCAAGAACATCTTCGTTACGGT | 19 (0.000364%) | | | GGCGAGACAAGGGTTCACATTTCGTTCATCACCCTTGGCCGGCTTTCGAACAGCCGGACTCCCATCAAAAGATGGTTGCCAAGAACATCTTCGTTACGGT | 20 (0.000383%) | | | GGCGGACCGGATTGCTCCGTTCCGCATCCGACCAGGACGCATCGCCGGCCCCCATCCGCTTCCCTCCCGACAATTTCAAGCACTCTTTGACTCTCTTTTC | 12 (0.000230%) | | | GGCGGCACATCTGTTAAAAGATAACGCAGGTGTCCTAAGATGAGCTCAACGAGAACAGAAATCTCGTGTGGAACAAAAGGGTAAAAGCTCGTTTGATTCT | 10 (0.000192%) | | | GGCGTAAGAATTGTATCCTTGTTAGAAGACACAAAGCCAAAGACTCATATGGACTTTGGCTACACCATGAAAGCTTTGAGAAGCAAGAAGAAGGTTGGTT | 10 (0.000192%) | | | GGCGTGCCTCGGCATCAGCGTGCTCCGGGCGTCGGCCTGTGGGCTCCCCATTCGACCCGTCTTGAAACACGGACCAAGGAGTCTGACATGTGTGCGAGTC | 14 (0.000268%) | | | GGCTCATTAAATCAGTTATAGTTTGTTTGATGGTAACTACTACTCGGATAACCGTAGTAATTCTAGAGCTAATACGTGCAACAAACCCCGACTTATGGAA | 9 (0.000172%) | | | GGCTGGGGCGGCACATCTGTTAAAAGATAACGCAGGTGTCCTAAGATGAGCTCAACGAGAACAGAAATCTCGTGTGGAACAAAAGGGTAAAAGCTCGTTT | 3 (0.000057%) | | | GGCTGTCCCGAGTGTGAGCGAGGTGTGAGTGTCGCCCATGGGCATCGACACCTTGCGGCTAGGAACTGGAACGAGACGGGTAGCAAAGATTTCGAGTAGC | 12 (0.000230%) | | | GGCTGTCCCGAGTGTGAGCGAGGTGTGAGTGTCGCCCATGGGCATCGACACCTTGCGGCTAGGAACTGGAACGAGACGGGTGGCAAAGATTTCGAGTAGC | 17 (0.000326%) | | | GGCTTGCTTTGAGCACTCTAATTTCTTCAAAGTAACAGCGCCGGAGGCACGACCCGGCCAATTAAGACCAGGAGCGTATCGCCGACCGAAGGGACAAGCC | 7 (0.000134%) | | | GGGAATCCTTGTTAGTTTCTTTTCCTCCGCTTATTGATATGCTTAAACTCAGCGGGTAATCCCGCCTGACCTGGGGTCGCTATATGGACTTTGGGTCATC | 53 (0.001016%) | | | GGGACAGTCGGGGGCATTCGTATTTCATAGTCAGAGGTGAAATTCTTGGATTTATGAAAGACGAACAACTGCGAAAGCATTTGCCAAGGATGTTTTCATT | 24 (0.000460%) | | | GGGAGGCGAATGCCAGCCGTTCGTTTGCATGTTCCTTGACACTTTTCGTGCCGGGGTTTTGTGATATCCGGAAGCAACGCGCACGACAAGACCGAGATAA | 16 (0.000307%) | | | GGGAGTTTGGCTGGGGCGGCACATCTGTTAAAAGATAACGCAGGTGTCCTAAGATGAGCTCAACGAGAACAGAAATCTCGTGTGGAACAAAAGGGTAAAA | 29 (0.000556%) | | | GGGATTTAGATTGTACTCATTCCAATTACCAGACTCGAAAGAGCCCGGTATTGTTATTTATTGTCACTACCTCCCCGTGTCAGGATTGGGTAATTTGCGC | 42 (0.000805%) | | | GGGCAAGTCTGGTGCCAGCAGCCGCGGTAATTCCAGCTCCAATAGCGTATATTTAAGTTGTTGCAGTTAAAAAGCTCGTAGTTGAACCTTGGGATGGGTC | 156 (0.002990%) | | | GGGCAGAGCCCGCGTCGACCTTTTATCTAATAAATGCGTCCCTTCCATAAGTCGGGGTTTGTTGCACGTATTAGCTCTAGAATTACTACGGTTATCCGAG | 45 (0.000862%) | | | GGGCATCGACACCTTGCGGCTAGGAACTGGAACGAGACGGGTGGCAAAGATTTCGAGTAGCACTTCATACTACCGTGGGTTTTTTAAACCTTCCGAGTTT | 12 (0.000230%) | | | GGGCATTCGTATTTCATAGTCAGAGGTGAAATTCTTGGATTTATGAAAGACGAACAACTGCGAAAGCATTTGCCAAGGATGTTTTCATTAATCAAGAACG | 44 (0.000843%) | | | GGGCCGCACGCGCGCTACACTGATGTATTCAACGAGTTCACACCTTGGCCGACAGGCCCGGGTAATCTTTGAAATTTCATCGTGATGGGGATAGATCGAT | 7 (0.000134%) | | | GGGCGGCACATCTGTTAAAAGATAACGCAGGTGTCCTAAGATGAGCTCAACGAGAACAGAAATCTCGTGTGGAACAAAAGGGTAAAAGCTCGTTTGATTC | 11 (0.000211%) | | | GGGCGGTGTGTACAAAGGGCAGGGACGTAGTCAACGCGAGCTGATGACTCGCGCTTACTAGGAATTCCTCGTTGAAGACCAACAATTGCAATGATCGATC | 25 (0.000479%) | | | GGGCGTCGGCCTGTGGGCTCCCCATTCGACCCGTCTTGAAACACGGACCAAGGAGTCTGACATGTGTGCGAGTCAACGGGTGAGTAAACCCGTAAGGCGC | 5 (0.000096%) | | | GGGGAGGCGAATGCCAGCCGTTCGTTTGCATGTTCCTTGACACTTTTCGTGCCGGGGTTTTGTGATATCCGGAAGCAACGCGCACGACAAGACCGAGATA | 36 (0.000690%) | | | GGGGAGTTTGGCTGGGGCGGCACATCTGTTAAAAGATAACGCAGGTGTCCTAAGATGAGCTCAACGAGAACAGAAATCTCGTGTGGAACAAAAGGGTAAA | 34 (0.000652%) | | | GGGGCATTCGTATTTCATAGTCAGAGGTGAAATTCTTGGATTTATGAAAGACGAACAACTGCGAAAGCATTTGCCAAGGATGTTTTCATTAATCAAGAAC | 37 (0.000709%) | | | GGGGGCATTCGTATTTCATAGTCAGAGGTGAAATTCTTGGATTTATGAAAGACGAACAACTGCGAAAGCATTTGCCAAGGATGTTTTCATTAATCAAGAA | 18 (0.000345%) | | | GGGTGACGGAGAATTAGGGTTCGATTCCGGAGAGGGAGCCTGAGAAACGGCTACCACATCCAAGGAAGGCAGCAGGCGCGCAAATTACCCAATCCTGACA | 13 (0.000249%) | | | GGGTGGTTCGCCGCCCGCGACGTCGCGAGAAGTCCACTAAACCTTATCATTTAGAGGAAGGAGAAGTCGTAACAAGGTTTCCGTAGGTGAACCTGCGGAA | 162 (0.003105%) | | | GGGTTCACATTTCGTTCATCACCCTTGGCCGGCTTTCGAACAGCCGGACTCCCATCAAAAGATGGTTGCCAAGAACATCTTCGTTACGGTTTGCTAATTC | 3 (0.000057%) | | | GGGTTGCGGTTTAAGTTGTTATACTCAATCATACACATGA | 417 (0.003197%) | | | GGGTTTACTCACCCGTTGACTCGCACACATGTCAGACTCCTTGGTCCGTGTTTCAAGACGGGTCGAATGGGGAGCCCACAGGCCGACGCCCGGAGCACGC | 5 (0.000096%) | | | GGTAAATTTTCATAATTTTTTGACACCTCTAGCTAGGTCATTTGACCTGATACAACATCGGATTTTCATGGTCTAGTTGGGGCTCCGTGGGCATATTTGA | 5 (0.000096%) | | | GGTAACGGGTGACGGAGAATTAGGGTTCGATTCCGGAGAGGGAGCCTGAGAAACGGCTACCACATCCAAGGAAGGCAGCAGGCGCGCAAATTACCCAATC | 90 (0.001725%) | | | GGTAATTCCAGCTCCAATAGCGTATATTTAAGTTGTTGCAGTTAAAAAGCTCGTAGTTGAACCTTGGGATGGGTCGGCCGGTCCGCCTTTGGTGTGCATT | 126 (0.002415%) | | | GGTAGGAGCGACGGGCGGTGTGTACAAAGGGCAGGGACGTAGTCAACGCGAGCTGATGACTCGCGCTTACTAGGAATTCCTCGTTGAAGACCAACAATTG | 43 (0.000824%) | | | GGTCTGTGATGCCCTTAGATGTTCTGGGCCGCACGCGCGCTACACTGATGTATTCAACGAGTTCACACCTTGGCCGACAGGCCCGGGTAATCTTTGAAAT | 11 (0.000211%) | | | GGTCTTCAACGAGGAATTCCTAGTAAGCGCGAGTCATCAGCTCGCGTTGACTACGTCCCTGCCCTTTGTACACACCGCCCGTCGCTCCTACCGATTGAAT | 10 (0.000192%) | | | GGTGACGGAGAATTAGGGTTCGATTCCGGAGAGGGAGCCTGAGAAACGGCTACCACATCCAAGGAAGGCAGCAGGCGCGCAAATTACCCAATCCTGACAC | 3 (0.000057%) | | | GGTGCCAGCAGCCGCGGTAATTCCAGCTCCAATAGCGTATATTTAAGTTGTTGCAGTTAAAAAGCTCGTAGTTGAACCTTGGGATGGGTCGGCCGGTCCG | 14 (0.000268%) | | | GGTGGTTCGCCGCCCGCGACGTCGCGAGAAGTCCACTAAACCTTATCATTTAGAGGAAGGAGAAGTCGTAACAAGGTTTCCGTAGGTGAACCTGCGGAAG | 21 (0.000402%) | | | GGTGTGAGTGTCGCCCATGGGCATCGACACCTTGCGGCTAGGAACTGGAACGAGACGGGTAGCAAAGATTTCGAGTAGCACTTCATACTACCGTGGGTTT | 6 (0.000115%) | | | GGTGTGAGTGTCGCCCATGGGCATCGACACCTTGCGGCTAGGAACTGGAACGAGACGGGTGGCAAAGATTTCGAGTAGCACTTCATACTACCGTGGGTTT | 7 (0.000134%) | | | GGTTCACATTTCGTTCATCACCCTTGGCCGGCTATCGAACAGCCGGACTCCCATCAAAAGATGGTTGCCAAGAACATCTTCGTTACGGTTTGCTAATTCT | 9 (0.000172%) | | | GGTTCACATTTCGTTCATCACCCTTGGCCGGCTTTCGAACAGCCGGACTCCCATCAAAAGATGGTTGCCAAGAACATCTTCGTTACGGTTTGCTAATTCT | 5 (0.000096%) | | | GGTTCGCCGCCCGCGACGTCGCGAGAAGTCCACTAAACCTTATCATTTAGAGGAAGGAGAAGTCGTAACAAGGTTTCCGTAGGTGAACCTGCGGAAGGAT | 14 (0.000268%) | | | GGTTCGTATTCGTACTGAAAATCAGAATCAAACGAGCTTTTACCCTTTTGTTCCACACGAGATTTCTGTTCTCGTTGAGCTCATCTTAGGACACCTGCGT | 3 (0.000057%) | | | GGTTGCGGTTTAAGTTCTTATACTCAATCATACACATGAC | 9307 (0.071346%) | | | GGTTGCGGTTTAAGTTGTTATACTCAATCATACACATGAC | 901 (0.006907%) | | | GGTTTAAGTTCTTATACTCAATCATACACATGACATCAAG | 1481 (0.011353%) | | | GGTTTAAGTTGTTATACTCAATCATACACATGACAACAAGTCATATTCGACTCCAAAACACTAACCAACCTTCTTCTTGCTTCTCAAAGCTTTCATGGTG | 123 (0.002357%) | | | GGTTTAAGTTGTTATACTCAATCATACACATGACATCAAG | 2143 (0.016428%) | | | GGTTTACTCACCCGTTGACTCGCACACATGTCAGACTCCTTGGTCCGTGTTTCAAGACGGGTCGAATGGGGAGCCCACAGGCCGACGCCCGGAGCACGCT | 20 (0.000383%) | | | GTAAATTTTCATAATTTTTTGACACCTCTAGCTAGGTCATTTGACCTGATACAACATCGGATTTTCATGGTCTAGTTGGGGCTCCGTGGGCATATTTGAT | 17 (0.000326%) | | | GTAACGGGTGACGGAGAATTAGGGTTCGATTCCGGAGAGGGAGCCTGAGAAACGGCTACCACATCCAAGGAAGGCAGCAGGCGCGCAAATTACCCAATCC | 19 (0.000364%) | | | GTAACTTCGGGAAAAGGATTGGCTCTGAGGGCTGGGCTCGGGGGTCCCAGTTCCGAACCCGTCGGCTGTCAGCGGACTGCTCGAGCTGCTTCCGCGGCGA | 10 (0.000192%) | | | GTAAGAATTGTATCCTTGTTAGAAGACACAAAGCCAAAGACTCATATGGACTTTGGCTACACCATGAAAGCTTTGAGAAGCAAGAAGAAGGTTGGTTAGT | 58 (0.001112%) | | | GTAAGTATGAACGAATTCAGACTGTGAAACTGCGAATGGCTCATTAAATCAGTTATAGTTTGTTTGATGGTAACTACTACTCGGATAACCGTAGTAATTC | 30 (0.000575%) | | | GTAATTCCAGCTCCAATAGCGTATATTTAAGTTGTTGCAGTTAAAAAGCTCGTAGTTGAACCTTGGGATGGGTCGGCCGGTCCGCCTTTGGTGTGCATTG | 19 (0.000364%) | | | GTACGCTCCAGGCGTCCTTGGCTCGGATTTAGGCCAACCGCGTGCGGTAACACACGGGAGACCAGCTTCCGTCCCGCATCAGCAAAGGATGGTGAGGGAC | 12 (0.000230%) | | | GTACTCATTCCAATTACCAGACTCGAAAGAGCCCGGTATTGTTATTTATTGTCACTACCTCCCCGTGTCAGGATTGGGTAATTTGCGCGCCTGCTGCCTT | 3 (0.000057%) | | | GTACTGAAAATCAGAATCAAACGAGCTTTTACCCTTTTGTTCCACACGAGATTTCTGTTCTCGTTGAGCTCATCTTAGGACACCTGCGTTATCTTTTAAC | 4 (0.000077%) | | | GTAGGAGCGACGGGCGGTGTGTACAAAGGGCAGGGACGTAGTCAACGCGAGCTGATGACTCGCGCTTACTAGGAATTCCTCGTTGAAGACCAACAATTGC | 114 (0.002185%) | | | GTAGGCTTGCTTTGAGCACTCTAATTTCTTCAAAGTAACAGCGCCGGAGGCACGACCCGGCCAATTAAGACCAGGAGCGTATCGCCGACCGAAGGGACAA | 33 (0.000632%) | | | GTAGTCATATGCTTGTCTCAAAGATTAAGCCATGCATGTGTAAGTATGAACGAATTCAGACTGTGAAACTGCGAATGGCTCATTAAATCAGTTATAGTTT | 98 (0.001878%) | | | GTAGTGTTTCCTTGTTAGAAGACACAAAGCCAAAGACTCATATGGACTTTGGCTACACCATGAAAGCTTTGAGAAGCAAGAAGAAGGTTGGTTAGTGTTT | 18 (0.000345%) | | | GTATCCAGAGCGTAGGCTTGCTTTGAGCACTCTAATTTCTTCAAAGTAACAGCGCCGGAGGCACGACCCGGCCAATTAAGACCAGGAGCGTATCGCCGAC | 16 (0.000307%) | | | GTATCCTTGTTAGAAGACACAAAGCCAAAGACTCATATGGACTTTGGCTACACCATGAAAGCTTTGAGAAGCAAGAAGAAGGTTGGTTAGTGTTTTGGAG | 22 (0.000422%) | | | GTATGAACGAATTCAGACTGTGAAACTGCGAATGGCTCATTAAATCAGTTATAGTTTGTTTGATGGTAACTACTACTCGGATAACCGTAGTAATTCTAGA | 10 (0.000192%) | | | GTATTCGTACTGAAAATCAGAATCAAACGAGCTTTTACCCTTTTGTTCCACACGAGATTTCTGTTCTCGTTGAGCTCATCTTAGGACACCTGCGTTATCT | 4 (0.000077%) | | | GTATTTCATAGTCAGAGGTGAAATTCTTGGATTTATGAAAGACGAACAACTGCGAAAGCATTTGCCAAGGATGTTTTCATTAATCAAGAACGAAAGTTGG | 5 (0.000096%) | | | GTCAGAGGTGAAATTCTTGGATTTATGAAAGACGAACAACTGCGAAAGCATTTGCCAAGGATGTTTTCATTAATCAAGAACGAAAGTTGGGGGCTCGAAG | 20 (0.000383%) | | | GTCAGGTGGGGAGTTTGGCTGGGGCGGCACATCTGTTAAAAGATAACGCAGGTGTCCTAAGATGAGCTCAACGAGAACAGAAATCTCGTGTGGAACAAAA | 12 (0.000230%) | | | GTCATATGCTTGTCTCAAAGATTAAGCCATGCATGTGTAAGTATGAACGAATTCAGACTGTGAAACTGCGAATGGCTCATTAAATCAGTTATAGTTTGTT | 57 (0.001092%) | | | GTCATATTCGACTCCAAAACACTAACCAACCTTCTTCTTGCTTCTCAAAGCTTTCATGGTGTAGCCAAAGTCCATATGAGTCTTTGGCTTTGTGTCTTCT | 260 (0.004983%) | | | GTCCATATGAGTCTTTGGCTTTGTGTCTTCTAACAAGGAT | 7093 (0.054373%) | | | GTCCCGAGTGTGAGCGAGGTGTGAGTGTCGCCCATGGGCATCGACACCTTGCGGCTAGGAACTGGAACGAGACGGGTAGCAAAGATTTCGAGTAGCACTT | 16 (0.000307%) | | | GTCCCGAGTGTGAGCGAGGTGTGAGTGTCGCCCATGGGCATCGACACCTTGCGGCTAGGAACTGGAACGAGACGGGTGGCAAAGATTTCGAGTAGCACTT | 28 (0.000537%) | | | GTCCCTCACCATCCTTTGCTGATGCGGGACGGAAGCTGGTCTCCCGTGTGTTACCGCACGCGGTTGGCCTAAATCCGAGCCAAGGACGCCTGGAGCGTAC | 9 (0.000172%) | | | GTCCGACTTTGTGAAATGACTTGAGAGGTGTAGGATAAGTGGGAGCTTCGGCGCAAGTGAAATACCACTACTTTTAACGTTATTTTACTTACTCCGTGAA | 12 (0.000230%) | | | GTCGAAATCCGCTAAGGAGTGTGTAACAACTCACCTGCCGAATCAACTAGCCCCGAAAATGGATGGCGCTTAAGCGCGCGACCTATACCCGGCCGTCGGG | 73 (0.001399%) | | | GTCGACCTTTTATCTAATAAATGCGTCCCTTCCATAAGTCGGGGTTTGTTGCACGTATTAGCTCTAGAATTACTACGGTTATCCGAGTAGTAGTTACCAT | 23 (0.000441%) | | | GTCGAGTTATCATGAATCATCAGAGCAACGGGCAGAGCCCGCGTCGACCTTTTATCTAATAAATGCGTCCCTTCCATAAGTCGGGGTTTGTTGCACGTAT | 462 (0.008854%) | | | GTCGCCCATGGGCATCGACACCTTGCGGCTAGGAACTGGAACGAGACGGGTGGCAAAGATTTCGAGTAGCACTTCATACTACCGTGGGTTTTTTAAACCT | 30 (0.000575%) | | | GTCGGCCTGTGGGCTCCCCATTCGACCCGTCTTGAAACACGGACCAAGGAGTCTGACATGTGTGCGAGTCAACGGGTGAGTAAACCCGTAAGGCGCAAGG | 28 (0.000537%) | | | GTCGGGGGCATTCGTATTTCATAGTCAGAGGTGAAATTCTTGGATTTATGAAAGACGAACAACTGCGAAAGCATTTGCCAAGGATGTTTTCATTAATCAA | 6 (0.000115%) | | | GTCGGTACGCTCCAGGCGTCCTTGGCTCGGATTTAGGCCAACCGCGTGCGGTAACACACGGGAGACCAGCTTCCGTCCCGCATCAGCAAAGGATGGTGAG | 8 (0.000153%) | | | GTCGTCCCTCACCATCCTTTGCTGATGCGGGACGGAAGCTGGTCTCCCGTGTGTTACCGCACGCGGTTGGCCTAAATCCGAGCCAAGGACGCCTGGAGCG | 683 (0.013089%) | | | GTCTCCGGGGAGGCGAATGCCAGCCGTTCGTTTGCATGTTCCTTGACACTTTTCGTGCCGGGGTTTTGTGATATCCGGAAGCAACGCGCACGACAAGACC | 8 (0.000153%) | | | GTCTGGTGCCAGCAGCCGCGGTAATTCCAGCTCCAATAGCGTATATTTAAGTTGTTGCAGTTAAAAAGCTCGTAGTTGAACCTTGGGATGGGTCGGCCGG | 18 (0.000345%) | | | GTCTGTGATGCCCTTAGATGTTCTGGGCCGCACGCGCGCTACACTGATGTATTCAACGAGTTCACACCTTGGCCGACAGGCCCGGGTAATCTTTGAAATT | 21 (0.000402%) | | | GTCTTCAACGAGGAATTCCTAGTAAGCGCGAGTCATCAGCTCGCGTTGACTACGTCCCTGCCCTTTGTACACACCGCCCGTCGCTCCTACCGATTGAATG | 13 (0.000249%) | | | GTGACGGAGAATTAGGGTTCGATTCCGGAGAGGGAGCCTGAGAAACGGCTACCACATCCAAGGAAGGCAGCAGGCGCGCAAATTACCCAATCCTGACACG | 8 (0.000153%) | | | GTGAGCGAGGTGTGAGTGTCGCCCATGGGCATCGACACCTTGCGGCTAGGAACTGGAACGAGACGGGTAGCAAAGATTTCGAGTAGCACTTCATACTACC | 22 (0.000422%) | | | GTGAGCGAGGTGTGAGTGTCGCCCATGGGCATCGACACCTTGCGGCTAGGAACTGGAACGAGACGGGTGGCAAAGATTTCGAGTAGCACTTCATACTACC | 23 (0.000441%) | | | GTGAGTGTCGCCCATGGGCATCGACACCTTGCGGCTAGGAACTGGAACGAGACGGGTAGCAAAGATTTCGAGTAGCACTTCATACTACCGTGGGTTTTTT | 25 (0.000479%) | | | GTGAGTGTCGCCCATGGGCATCGACACCTTGCGGCTAGGAACTGGAACGAGACGGGTGGCAAAGATTTCGAGTAGCACTTCATACTACCGTGGGTTTTTT | 50 (0.000958%) | | | GTGATGCCCTTAGATGTTCTGGGCCGCACGCGCGCTACACTGATGTATTCAACGAGTTCACACCTTGGCCGACAGGCCCGGGTAATCTTTGAAATTTCAT | 22 (0.000422%) | | | GTGCCAGCAGCCGCGGTAATTCCAGCTCCAATAGCGTATATTTAAGTTGTTGCAGTTAAAAAGCTCGTAGTTGAACCTTGGGATGGGTCGGCCGGTCCGC | 26 (0.000498%) | | | GTGCCTCGGCATCAGCGTGCTCCGGGCGTCGGCCTGTGGGCTCCCCATTCGACCCGTCTTGAAACACGGACCAAGGAGTCTGACATGTGTGCGAGTCAAC | 15 (0.000287%) | | | GTGCTCCGGGCGTCGGCCTGTGGGCTCCCCATTCGACCCGTCTTGAAACACGGACCAAGGAGTCTGACATGTGTGCGAGTCAACGGGTGAGTAAACCCGT | 17 (0.000326%) | | | GTGGCCTACCATGGTGGTAACGGGTGACGGAGAATTAGGGTTCGATTCCGGAGAGGGAGCCTGAGAAACGGCTACCACATCCAAGGAAGGCAGCAGGCGC | 9 (0.000172%) | | | GTGGGGAGTTTGGCTGGGGCGGCACATCTGTTAAAAGATAACGCAGGTGTCCTAAGATGAGCTCAACGAGAACAGAAATCTCGTGTGGAACAAAAGGGTA | 8 (0.000153%) | | | GTGGGTGGTTCGCCGCCCGCGACGTCGCGAGAAGTCCACTAAACCTTATCATTTAGAGGAAGGAGAAGTCGTAACAAGGTTTCCGTAGGTGAACCTGCGG | 76 (0.001457%) | | | GTGGTTCGCCGCCCGCGACGTCGCGAGAAGTCCACTAAACCTTATCATTTAGAGGAAGGAGAAGTCGTAACAAGGTTTCCGTAGGTGAACCTGCGGAAGG | 17 (0.000326%) | | | GTGTAAGTATGAACGAATTCAGACTGTGAAACTGCGAATGGCTCATTAAATCAGTTATAGTTTGTTTGATGGTAACTACTACTCGGATAACCGTAGTAAT | 8 (0.000153%) | | | GTGTATCCTTGTTAGAAGACACAAAGCCAAAGACTCATATGGACTTTGGCTACACCATGAAAGCTTTGAGAAGCAAGAAGAAGGTTGGTTAGTGTTTTGG | 5 (0.000096%) | | | GTGTCGCCCATGGGCATCGACACCTTGCGGCTAGGAACTGGAACGAGACGGGTGGCAAAGATTTCGAGTAGCACTTCATACTACCGTGGGTTTTTTAAAC | 10 (0.000192%) | | | GTGTGAGCGAGGTGTGAGTGTCGCCCATGGGCATCGACACCTTGCGGCTAGGAACTGGAACGAGACGGGTAGCAAAGATTTCGAGTAGCACTTCATACTA | 34 (0.000652%) | | | GTGTGAGCGAGGTGTGAGTGTCGCCCATGGGCATCGACACCTTGCGGCTAGGAACTGGAACGAGACGGGTGGCAAAGATTTCGAGTAGCACTTCATACTA | 35 (0.000671%) | | | GTGTGAGTGTCGCCCATGGGCATCGACACCTTGCGGCTAGGAACTGGAACGAGACGGGTAGCAAAGATTTCGAGTAGCACTTCATACTACCGTGGGTTTT | 14 (0.000268%) | | | GTGTGAGTGTCGCCCATGGGCATCGACACCTTGCGGCTAGGAACTGGAACGAGACGGGTGGCAAAGATTTCGAGTAGCACTTCATACTACCGTGGGTTTT | 24 (0.000460%) | | | GTGTTTCCTTGTTAGAAGACACAAAGCCAAAGACTCATATGGACTTTGGCTACACCATGAAAGCTTTGAGAAGCAAGAAGAAGGTTGGTTAGTGTTTTGG | 35 (0.000671%) | | | GTTAAAAGATAACGCAGGTGTCCTAAGATGAGCTCAACGAGAACAGAAATCTCGTGTGGAACAAAAGGGTAAAAGCTCGTTTGATTCTGATTTTCAGTAC | 24 (0.000460%) | | | GTTAAGGGATTTAGATTGTACTCATTCCAATTACCAGACTCGAAAGAGCCCGGTATTGTTATTTATTGTCACTACCTCCCCGTGTCAGGATTGGGTAATT | 104 (0.001993%) | | | GTTAGAAGACACAAAGCCAAAGACTCATATGGACTTTGGCTACACCATGAAAGCTTTGAGAAGCAAGAAGAAGGTTGGTTAGTGTTTTGGAGTCGAATAT | 81 (0.001552%) | | | GTTAGTTTCTTTTCCTCCGCTTATTGATATGCTTAAACTCAGCGGGTAATCCCGCCTGACCTGGGGTCGCTATATGGACTTTGGGTCATCTACAGCTTCC | 25 (0.000479%) | | | GTTATACTCAATCATACACATGACAACAAGTCATATTCGACTCCAAAACACTAACCAACCTTCTTCTTGCTTCTCAAAGCTTTCATGGTGTAGCCAAAGT | 122 (0.002338%) | | | GTTATAGTTTGTTTGATGGTAACTACTACTCGGATAACCGTAGTAATTCTAGAGCTAATACGTGCAACAAACCCCGACTTATGGAAGGGACGCATTTATT | 11 (0.000211%) | | | GTTATCATGAATCATCAGAGCAACGGGCAGAGCCCGCGTCGACCTTTTATCTAATAAATGCGTCCCTTCCATAAGTCGGGGTTTGTTGCACGTATTAGCT | 298 (0.005711%) | | | GTTATCCCATGCTAATGTATCCAGAGCGTAGGCTTGCTTTGAGCACTCTAATTTCTTCAAAGTAACAGCGCCGGAGGCACGACCCGGCCAATTAAGACCA | 58 (0.001112%) | | | GTTCACATTTCGTTCATCACCCTTGGCCGGCTATCGAACAGCCGGACTCCCATCAAAAGATGGTTGCCAAGAACATCTTCGTTACGGTTTGCTAATTCTC | 13 (0.000249%) | | | GTTCACATTTCGTTCATCACCCTTGGCCGGCTTTCGAACAGCCGGACTCCCATCAAAAGATGGTTGCCAAGAACATCTTCGTTACGGTTTGCTAATTCTC | 15 (0.000287%) | | | GTTCACCACCGCATGTCGGTACGCTCCAGGCGTCCTTGGCTCGGATTTAGGCCAACCGCGTGCGGTAACACACGGGAGACCAGCTTCCGTCCCGCATCAG | 95 (0.001821%) | | | GTTCCATCGACCAGAGGCTGTTCACCTTGGAGACCTGATGCGGTTATGAGTACGACCGGGCGTGAGCGGCACTCGGTCCTCCGGATTTTCAAGGGCCGCC | 41 (0.000786%) | | | GTTCGCCGCCCGCGACGTCGCGAGAAGTCCACTAAACCTTATCATTTAGAGGAAGGAGAAGTCGTAACAAGGTTTCCGTAGGTGAACCTGCGGAAGGATC | 21 (0.000402%) | | | GTTCGCTCGCCGTTACTAAGGGAATCCTTGTTAGTTTCTTTTCCTCCGCTTATTGATATGCTTAAACTCAGCGGGTAATCCCGCCTGACCTGGGGTCGCT | 1057 (0.020257%) | | | GTTCGTATTCGTACTGAAAATCAGAATCAAACGAGCTTTTACCCTTTTGTTCCACACGAGATTTCTGTTCTCGTTGAGCTCATCTTAGGACACCTGCGTT | 12 (0.000230%) | | | GTTCTGGGCCGCACGCGCGCTACACTGATGTATTCAACGAGTTCACACCTTGGCCGACAGGCCCGGGTAATCTTTGAAATTTCATCGTGATGGGGATAGA | 42 (0.000805%) | | | GTTCTTATACTCAATCATACACATGACATCAAGTCATATTCGACTCCAAAACACTAACCAACCTTCTTCTTGCTTCTCAAAGCTTTCATGGTGTAGCCAA | 107 (0.002051%) | | | GTTGAGCTTGACTCTAGTCCGACTTTGTGAAATGACTTGAGAGGTGTAGGATAAGTGGGAGCTTCGGCGCAAGTGAAATACCACTACTTTTAACGTTATT | 46 (0.000882%) | | | GTTGCGGTTTAAGTTCTTATACTCAATCATACACATGACA | 2282 (0.017493%) | | | GTTGCGGTTTAAGTTGTTATACTCAATCATACACATGACA | 2448 (0.018766%) | | | GTTGGTCTTCAACGAGGAATTCCTAGTAAGCGCGAGTCATCAGCTCGCGTTGACTACGTCCCTGCCCTTTGTACACACCGCCCGTCGCTCCTACCGATTG | 184 (0.003526%) | | | GTTGTTATACTCAATCATACACATGACATCAAGTCATATT | 221 (0.001694%) | | | GTTTAAGTTCTTATACTCAATCATACACATGACATCAAGT | 2186 (0.016757%) | | | GTTTAAGTTGTTATACTCAATCATACACATGACAACAAGTCATATTCGACTCCAAAACACTAACCAACCTTCTTCTTGCTTCTCAAAGCTTTCATGGTGT | 147 (0.002817%) | | | GTTTAAGTTGTTATACTCAATCATACACATGACATCAAGT | 1025 (0.007857%) | | | GTTTACTCACCCGTTGACTCGCACACATGTCAGACTCCTTGGTCCGTGTTTCAAGACGGGTCGAATGGGGAGCCCACAGGCCGACGCCCGGAGCACGCTG | 27 (0.000517%) | | | GTTTCCTTGTTAGAAGACACAAAGCCAAAGACTCATATGGACTTTGGCTACACCATGAAAGCTTTGAGAAGCAAGAAGAAGGTTGGTTAGTGTTTTGGAG | 48 (0.000920%) | | | GTTTCTTTTCCTCCGCTTATTGATATGCTTAAACTCAGCGGGTAATCCCGCCTGACCTGGGGTCGCTATATGGACTTTGGGTCATCTACAGCTTCCGGAC | 26 (0.000498%) | | | GTTTGATGGTAACTACTACTCGGATAACCGTAGTAATTCTAGAGCTAATACGTGCAACAAACCCCGACTTATGGAAGGGACGCATTTATTAGATAAAAGG | 24 (0.000460%) | | | GTTTGGCTGGGGCGGCACATCTGTTAAAAGATAACGCAGGTGTCCTAAGATGAGCTCAACGAGAACAGAAATCTCGTGTGGAACAAAAGGGTAAAAGCTC | 35 (0.000671%) | | | GTTTGTTTGATGGTAACTACTACTCGGATAACCGTAGTAATTCTAGAGCTAATACGTGCAACAAACCCCGACTTATGGAAGGGACGCATTTATTAGATAA | 35 (0.000671%) | | | TAAAAGATAACGCAGGTGTCCTAAGATGAGCTCAACGAGAACAGAAATCTCGTGTGGAACAAAAGGGTAAAAGCTCGTTTGATTCTGATTTTCAGTACGA | 17 (0.000326%) | | | TAAAAGCCTAAGTAGTGTTTCCTTGTTAGAAGACACAAAGCCAAAGACTCATATGGACTTTGGCTACACCATGAAAGCTTTGAGAAGCAAGAAGAAGGTT | 37 (0.000709%) | | | TAAAGGCGTAAGAATTGTATCCTTGTTAAAAGACACAAAGCCAAAGACTCATATGGACTTTGGCTACACCATGAAAGCTTTGAGAAGCAAGAAGAAGGTT | 62 (0.001188%) | | | TAAAGGCGTAAGAATTGTATCCTTGTTAGAAGACACAAAGCCAAAGACTCATATGGACTTTGGCTACACCATGAAAGCTTTGAGAAGCAAGAAGAAGGTT | 138 (0.002645%) | | | TAAATCAGTTATAGTTTGTTTGATGGTAACTACTACTCGGATAACCGTAGTAATTCTAGAGCTAATACGTGCAACAAACCCCGACTTATGGAAGGGACGC | 9 (0.000172%) | | | TAAATTTTCATAATTTTTTGACACCTCTAGCTAGGTCATTTGACCTGATACAACATCGGATTTTCATGGTCTAGTTGGGGCTCCGTGGGCATATTTGATG | 9 (0.000172%) | | | TAACAGGGACAGTCGGGGGCATTCGTATTTCATAGTCAGAGGTGAAATTCTTGGATTTATGAAAGACGAACAACTGCGAAAGCATTTGCCAAGGATGTTT | 45 (0.000862%) | | | TAACAGGTCTGTGATGCCCTTAGATGTTCTGGGCCGCACGCGCGCTACACTGATGTATTCAACGAGTTCACACCTTGGCCGACAGGCCCGGGTAATCTTT | 37 (0.000709%) | | | TAACGCCTCGAAGAACTAATGGCAGCCCACGCAAGGCAAGCCCATTCTCCTCGACGATTCAGCAGTTTTTGTCCGAGAACTGCTGAGAAAACTCGGAAAA | 11 (0.000211%) | | | TAACGGCGTGCCTCGGCATCAGCGTGCTCCGGGCGTCGGCCTGTGGGCTCCCCATTCGACCCGTCTTGAAACACGGACCAAGGAGTCTGACATGTGTGCG | 34 (0.000652%) | | | TAACGGGTGACGGAGAATTAGGGTTCGATTCCGGAGAGGGAGCCTGAGAAACGGCTACCACATCCAAGGAAGGCAGCAGGCGCGCAAATTACCCAATCCT | 13 (0.000249%) | | | TAAGAATTGTATCCTTGTTAGAAGACACAAAGCCAAAGACTCATATGGACTTTGGCTACACCATGAAAGCTTTGAGAAGCAAGAAGAAGGTTGGTTAGTG | 101 (0.001936%) | | | TAAGCCATGCATGTGTAAGTATGAACGAATTCAGACTGTGAAACTGCGAATGGCTCATTAAATCAGTTATAGTTTGTTTGATGGTAACTACTACTCGGAT | 5 (0.000096%) | | | TAAGCGCGCGACCTATACCCGGCCGTCGGGGCAAGAGCCAGGCCTCGATGAGTAGGAGGGCGCGGCGGTCGCTGCAAAACCTAGGGCGCGAGCCCGGGCG | 10 (0.000192%) | | | TAAGGATACTAAATCCTATTTTCTGGTAAATTTTCATAATTTTTTGACACCTCTAGCTAGGTCATTTGACCTGATACAACATCGGATTTTCATGGTCTAG | 12 (0.000230%) | | | TAAGGGATTTAGATTGTACTCATTCCAATTACCAGACTCGAAAGAGCCCGGTATTGTTATTTATTGTCACTACCTCCCCGTGTCAGGATTGGGTAATTTG | 36 (0.000690%) | | | TAAGTAGTGTTTCCTTGTTAGAAGACACAAAGCCAAAGACTCATATGGACTTTGGCTACACCATGAAAGCTTTGAGAAGCAAGAAGAAGGTTGGTTAGTG | 52 (0.000997%) | | | TAAGTATGAACGAATTCAGACTGTGAAACTGCGAATGGCTCATTAAATCAGTTATAGTTTGTTTGATGGTAACTACTACTCGGATAACCGTAGTAATTCT | 5 (0.000096%) | | | TAAGTTCTTATACTCAATCATACACATGACATCAAGTCATATTCGACTCCAAAACACTAACCAACCTTCTTCTTGCTTCTCAAAGCTTTCATGGTGTAGC | 17 (0.000326%) | | | TAAGTTGTTATACTCAATCATACACATGACATCAAGTCAT | 91 (0.000698%) | | | TAATGATTAACAGGGACAGTCGGGGGCATTCGTATTTCATAGTCAGAGGTGAAATTCTTGGATTTATGAAAGACGAACAACTGCGAAAGCATTTGCCAAG | 1509 (0.028919%) | | | TAATTCCAGCTCCAATAGCGTATATTTAAGTTGTTGCAGTTAAAAAGCTCGTAGTTGAACCTTGGGATGGGTCGGCCGGTCCGCCTTTGGTGTGCATTGG | 15 (0.000287%) | | | TAATTTTTTGACACCTCTAGCTAGGTCATTTGACCTGATACAACATCGGATTTTCATGGTCTAGTTGGGGCTCCGTGGGCATATTTGATGCAAACTTGAC | 6 (0.000115%) | | | TACACATGACATCAAGTCATATTCGACTCCAAAACACTAACCAACCTTCTTCTTGCTTCTCAAAGCTTTCATGGTGTAGCCAAAGTCCATATGAGTCTTT | 11 (0.000211%) | | | TACACCATGAAAGCTTTGAGAAGCAAGAAGAAGGTTGGTTAGTGTTTTGGAGTCGAATATGACTTGATGTCATGTGTATGATTGAGTATAACAACTTAAA | 132 (0.002530%) | | | TACATTGTTCCATCGACCAGAGGCTGTTCACCTTGGAGACCTGATGCGGTTATGAGTACGACCGGGCGTGAGCGGCACTCGGTCCTCCGGATTTTCAAGG | 3 (0.000057%) | | | TACCAGACTCGAAAGAGCCCGGTATTGTTATTTATTGTCACTACCTCCCCGTGTCAGGATTGGGTAATTTGCGCGCCTGCTGCCTTCCTTGGATGTGGTA | 6 (0.000115%) | | | TACCCGGCCGTCGGGGCAAGAGCCAGGCCTCGATGAGTAGGAGGGCGCGGCGGTCGCTGCAAAACCTAGGGCGCGAGCCCGGGCGGAGCGGCCGTCGGTG | 3 (0.000057%) | | | TACGGGTTTACTCACCCGTTGACTCGCACACATGTCAGACTCCTTGGTCCGTGTTTCAAGACGGGTCGAATGGGGAGCCCACAGGCCGACGCCCGGAGCA | 3 (0.000057%) | | | TACTCAATCATACACATGACAACAAGTCATATTCGACTCCAAAACACTAACCAACCTTCTTCTTGCTTCTCAAAGCTTTCATGGTGTAGCCAAAGTCCAT | 12 (0.000230%) | | | TACTCAATCATACACATGACATCAAGTCATATTCGACTCCAAAACACTAACCAACCTTCTTCTTGCTTCTCAAAGCTTTCATGGTGTAGCCAAAGTCCAT | 17 (0.000326%) | | | TAGAAGACACAAAGCCAAAGACTCATATGGACTTTGGCTACACCATGAAAGCTTTGAGAAGCAAGAAGAAGGTTGGTTAGTGTTTTGGAGTCGAATATGA | 25 (0.000479%) | | | TAGATGTTCTGGGCCGCACGCGCGCTACACTGATGTATTCAACGAGTTCACACCTTGGCCGACAGGCCCGGGTAATCTTTGAAATTTCATCGTGATGGGG | 18 (0.000345%) | | | TAGGAGCGACGGGCGGTGTGTACAAAGGGCAGGGACGTAGTCAACGCGAGCTGATGACTCGCGCTTACTAGGAATTCCTCGTTGAAGACCAACAATTGCA | 14 (0.000268%) | | | TAGGCCACGCTTTCACGGTTCGTATTCGTACTGAAAATCAGAATCAAACGAGCTTTTACCCTTTTGTTCCACACGAGATTTCTGTTCTCGTTGAGCTCAT | 62 (0.001188%) | | | TAGGCGAGACAAGGGTTCACATTTCGTTCATCACCCTTGGCCGGCTATCGAACAGCCGGACTCCCATCAAAAGATGGTTGCCAAGAACATCTTCGTTACG | 8 (0.000153%) | | | TAGGCGAGACAAGGGTTCACATTTCGTTCATCACCCTTGGCCGGCTTTCGAACAGCCGGACTCCCATCAAAAGATGGTTGCCAAGAACATCTTCGTTACG | 6 (0.000115%) | | | TAGGCTGTCCCGAGTGTGAGCGAGGTGTGAGTGTCGCCCATGGGCATCGACACCTTGCGGCTAGGAACTGGAACGAGACGGGTAGCAAAGATTTCGAGTA | 16 (0.000307%) | | | TAGGCTGTCCCGAGTGTGAGCGAGGTGTGAGTGTCGCCCATGGGCATCGACACCTTGCGGCTAGGAACTGGAACGAGACGGGTGGCAAAGATTTCGAGTA | 21 (0.000402%) | | | TAGTCAGAGGTGAAATTCTTGGATTTATGAAAGACGAACAACTGCGAAAGCATTTGCCAAGGATGTTTTCATTAATCAAGAACGAAAGTTGGGGGCTCGA | 3 (0.000057%) | | | TAGTCATATGCTTGTCTCAAAGATTAAGCCATGCATGTGTAAGTATGAACGAATTCAGACTGTGAAACTGCGAATGGCTCATTAAATCAGTTATAGTTTG | 54 (0.001035%) | | | TAGTGGCCTACCATGGTGGTAACGGGTGACGGAGAATTAGGGTTCGATTCCGGAGAGGGAGCCTGAGAAACGGCTACCACATCCAAGGAAGGCAGCAGGC | 6 (0.000115%) | | | TAGTGTTTCCTTGTTAGAAGACACAAAGCCAAAGACTCATATGGACTTTGGCTACACCATGAAAGCTTTGAGAAGCAAGAAGAAGGTTGGTTAGTGTTTT | 10 (0.000192%) | | | TAGTTTCTTTTCCTCCGCTTATTGATATGCTTAAACTCAGCGGGTAATCCCGCCTGACCTGGGGTCGCTATATGGACTTTGGGTCATCTACAGCTTCCGG | 5 (0.000096%) | | | TATACCCGGCCGTCGGGGCAAGAGCCAGGCCTCGATGAGTAGGAGGGCGCGGCGGTCGCTGCAAAACCTAGGGCGCGAGCCCGGGCGGAGCGGCCGTCGG | 12 (0.000230%) | | | TATACTCAATCATACACATGACAACAAGTCATATTCGACTCCAAAACACTAACCAACCTTCTTCTTGCTTCTCAAAGCTTTCATGGTGTAGCCAAAGTCC | 10 (0.000192%) | | | TATACTCAATCATACACATGACATCAAGTCATATTCGACTCCAAAACACTAACCAACCTTCTTCTTGCTTCTCAAAGCTTTCATGGTGTAGCCAAAGTCC | 44 (0.000843%) | | | TATAGTTTGTTTGATGGTAACTACTACTCGGATAACCGTAGTAATTCTAGAGCTAATACGTGCAACAAACCCCGACTTATGGAAGGGACGCATTTATTAG | 4 (0.000077%) | | | TATCATGAATCATCAGAGCAACGGGCAGAGCCCGCGTCGACCTTTTATCTAATAAATGCGTCCCTTCCATAAGTCGGGGTTTGTTGCACGTATTAGCTCT | 70 (0.001342%) | | | TATCCAGAGCGTAGGCTTGCTTTGAGCACTCTAATTTCTTCAAAGTAACAGCGCCGGAGGCACGACCCGGCCAATTAAGACCAGGAGCGTATCGCCGACC | 3 (0.000057%) | | | TATCCCATGCTAATGTATCCAGAGCGTAGGCTTGCTTTGAGCACTCTAATTTCTTCAAAGTAACAGCGCCGGAGGCACGACCCGGCCAATTAAGACCAGG | 6 (0.000115%) | | | TATCCTTGTTAGAAGACACAAAGCCAAAGACTCATATGGACTTTGGCTACACCATGAAAGCTTTGAGAAGCAAGAAGAAGGTTGGTTAGTGTTTTGGAGT | 33 (0.000632%) | | | TATGACTTGATGTCATGTGTATGATTGAGTATAAGAACTT | 292 (0.002238%) | | | TATGATGTTATCCCATGCTAATGTATCCAGAGCGTAGGCTTGCTTTGAGCACTCTAATTTCTTCAAAGTAACAGCGCCGGAGGCACGACCCGGCCAATTA | 28 (0.000537%) | | | TATGCTTGTCTCAAAGATTAAGCCATGCATGTGTAAGTATGAACGAATTCAGACTGTGAAACTGCGAATGGCTCATTAAATCAGTTATAGTTTGTTTGAT | 10 (0.000192%) | | | TATGGACTTTGGCTACACCATGAAAGCTTTGAGAAGCAAGAAGAAGGTTGGTTAGTGTTTTGGAGTCGAATATGACTTGATGTCATGTGTATGATTGAGT | 23 (0.000441%) | | | TATTCGACTCCAAAACACTAACCAACCTTCTTCTTGCTTCTCAAAGCTTTCATGGTGTAGCCAAAGTCCATATGAGTCTTTGGCTTTGTGTCTTCTAACA | 41 (0.000786%) | | | TATTCGTACTGAAAATCAGAATCAAACGAGCTTTTACCCTTTTGTTCCACACGAGATTTCTGTTCTCGTTGAGCTCATCTTAGGACACCTGCGTTATCTT | 5 (0.000096%) | | | TCAAAGATTAAGCCATGCATGTGTAAGTATGAACGAATTCAGACTGTGAAACTGCGAATGGCTCATTAAATCAGTTATAGTTTGTTTGATGGTAACTACT | 4 (0.000077%) | | | TCAAAGCTTTCATGGTGTAGCCAAAGTCCATATGAGTCTTTGGCTTTGTGTCTTCTAACAAGGAAACACTACTTAGGCTTATAAGATGCGGTTGCGGTTT | 7 (0.000134%) | | | TCAAGTCATATTCGACTCCAAAACACTAACCAACCTTCTTCTTGCTTCTCAAAGCTTTCATGGTGTAGCCAAAGTCCATATGAGTCTTTGGCTTTGTGTC | 48 (0.000920%) | | | TCAAGTTCACCACCGCATGTCGGTACGCTCCAGGCGTCCTTGGCTCGGATTTAGGCCAACCGCGTGCGGTAACACACGGGAGACCAGCTTCCGTCCCGCA | 3 (0.000057%) | | | TCAATCATACACATGACAACAAGTCATATTCGACTCCAAAACACTAACCAACCTTCTTCTTGCTTCTCAAAGCTTTCATGGTGTAGCCAAAGTCCATATG | 18 (0.000345%) | | | TCAATCATACACATGACATCAAGTCATATTCGACTCCAAAACACTAACCAACCTTCTTCTTGCTTCTCAAAGCTTTCATGGTGTAGCCAAAGTCCATATG | 30 (0.000575%) | | | TCAATCGGTAGGAGCGACGGGCGGTGTGTACAAAGGGCAGGGACGTAGTCAACGCGAGCTGATGACTCGCGCTTACTAGGAATTCCTCGTTGAAGACCAA | 44 (0.000843%) | | | TCACATTTCGTTCATCACCCTTGGCCGGCTATCGAACAGCCGGACTCCCATCAAAAGATGGTTGCCAAGAACATCTTCGTTACGGTTTGCTAATTCTCGG | 5 (0.000096%) | | | TCACATTTCGTTCATCACCCTTGGCCGGCTTTCGAACAGCCGGACTCCCATCAAAAGATGGTTGCCAAGAACATCTTCGTTACGGTTTGCTAATTCTCGG | 8 (0.000153%) | | | TCACCACCGCATGTCGGTACGCTCCAGGCGTCCTTGGCTCGGATTTAGGCCAACCGCGTGCGGTAACACACGGGAGACCAGCTTCCGTCCCGCATCAGCA | 6 (0.000115%) | | | TCACCATCCTTTGCTGATGCGGGACGGAAGCTGGTCTCCCGTGTGTTACCGCACGCGGTTGGCCTAAATCCGAGCCAAGGACGCCTGGAGCGTACCGACA | 6 (0.000115%) | | | TCAGAATCAAACGAGCTTTTACCCTTTTGTTCCACACGAGATTTCTGTTCTCGTTGAGCTCATCTTAGGACACCTGCGTTATCTTTTAACAGATGTGCCG | 17 (0.000326%) | | | TCAGAGGTGAAATTCTTGGATTTATGAAAGACGAACAACTGCGAAAGCATTTGCCAAGGATGTTTTCATTAATCAAGAACGAAAGTTGGGGGCTCGAAGA | 4 (0.000077%) | | | TCAGCGTGCTCCGGGCGTCGGCCTGTGGGCTCCCCATTCGACCCGTCTTGAAACACGGACCAAGGAGTCTGACATGTGTGCGAGTCAACGGGTGAGTAAA | 4 (0.000077%) | | | TCATAGTCAGAGGTGAAATTCTTGGATTTATGAAAGACGAACAACTGCGAAAGCATTTGCCAAGGATGTTTTCATTAATCAAGAACGAAAGTTGGGGGCT | 8 (0.000153%) | | | TCATATGCTTGTCTCAAAGATTAAGCCATGCATGTGTAAGTATGAACGAATTCAGACTGTGAAACTGCGAATGGCTCATTAAATCAGTTATAGTTTGTTT | 4 (0.000077%) | | | TCATATGGACTTTGGCTACACCATGAAAGCTTTGAGAAGCAAGAAGAAGGTTGGTTAGTGTTTTGGAGTCGAATATGACTTGATGTCATGTGTATGATTG | 36 (0.000690%) | | | TCATATTCGACTCCAAAACACTAACCAACCTTCTTCTTGCTTCTCAAAGCTTTCATGGTGTAGCCAAAGTCCATATGAGTCTTTGGCTTTGTGTCTTCTA | 68 (0.001303%) | | | TCATGAATCATCAGAGCAACGGGCAGAGCCCGCGTCGACCTTTTATCTAATAAATGCGTCCCTTCCATAAGTCGGGGTTTGTTGCACGTATTAGCTCTAG | 30 (0.000575%) | | | TCATGTGTATGATTGAGTATAAGAACTTAAACCGCAACCG | 4325 (0.033155%) | | | TCATTCCAATTACCAGACTCGAAAGAGCCCGGTATTGTTATTTATTGTCACTACCTCCCCGTGTCAGGATTGGGTAATTTGCGCGCCTGCTGCCTTCCTT | 3 (0.000057%) | | | TCCAACCTAGGCGAGACAAGGGTTCACATTTCGTTCATCACCCTTGGCCGGCTATCGAACAGCCGGACTCCCATCAAAAGATGGTTGCCAAGAACATCTT | 4 (0.000077%) | | | TCCAACCTAGGCGAGACAAGGGTTCACATTTCGTTCATCACCCTTGGCCGGCTTTCGAACAGCCGGACTCCCATCAAAAGATGGTTGCCAAGAACATCTT | 3 (0.000057%) | | | TCCAACGAAGCACGCCCATCCAACCTAGGCGAGACAAGGGTTCACATTTCGTTCATCACCCTTGGCCGGCTATCGAACAGCCGGACTCCCATCAAAAGAT | 25 (0.000479%) | | | TCCAACGAAGCACGCCCATCCAACCTAGGCGAGACAAGGGTTCACATTTCGTTCATCACCCTTGGCCGGCTTTCGAACAGCCGGACTCCCATCAAAAGAT | 18 (0.000345%) | | | TCCAATTACCAGACTCGAAAGAGCCCGGTATTGTTATTTATTGTCACTACCTCCCCGTGTCAGGATTGGGTAATTTGCGCGCCTGCTGCCTTCCTTGGAT | 7 (0.000134%) | | | TCCAGAGCGTAGGCTTGCTTTGAGCACTCTAATTTCTTCAAAGTAACAGCGCCGGAGGCACGACCCGGCCAATTAAGACCAGGAGCGTATCGCCGACCGA | 4 (0.000077%) | | | TCCATATGAGTCTTTGGCTTTGTGTCTTCTAACAAGGATA | 256 (0.001962%) | | | TCCATGCTTTCCAACGAAGCACGCCCATCCAACCTAGGCGAGACAAGGGTTCACATTTCGTTCATCACCCTTGGCCGGCTATCGAACAGCCGGACTCCCA | 12 (0.000230%) | | | TCCATTGGAGGGCAAGTCTGGTGCCAGCAGCCGCGGTAATTCCAGCTCCAATAGCGTATATTTAAGTTGTTGCAGTTAAAAAGCTCGTAGTTGAACCTTG | 11 (0.000211%) | | | TCCCATGCTAATGTATCCAGAGCGTAGGCTTGCTTTGAGCACTCTAATTTCTTCAAAGTAACAGCGCCGGAGGCACGACCCGGCCAATTAAGACCAGGAG | 5 (0.000096%) | | | TCCCTTGCCTACATTGTTCCATCGACCAGAGGCTGTTCACCTTGGAGACCTGATGCGGTTATGAGTACGACCGGGCGTGAGCGGCACTCGGTCCTCCGGA | 6 (0.000115%) | | | TCCGACTTTGTGAAATGACTTGAGAGGTGTAGGATAAGTGGGAGCTTCGGCGCAAGTGAAATACCACTACTTTTAACGTTATTTTACTTACTCCGTGAAT | 6 (0.000115%) | | | TCCGCCTAGGCTGTCCCGAGTGTGAGCGAGGTGTGAGTGTCGCCCATGGGCATCGACACCTTGCGGCTAGGAACTGGAACGAGACGGGTAGCAAAGATTT | 170 (0.003258%) | | | TCCGCCTAGGCTGTCCCGAGTGTGAGCGAGGTGTGAGTGTCGCCCATGGGCATCGACACCTTGCGGCTAGGAACTGGAACGAGACGGGTGGCAAAGATTT | 283 (0.005424%) | | | TCCGCTTATTGATATGCTTAAACTCAGCGGGTAATCCCGCCTGACCTGGGGTCGCTATATGGACTTTGGGTCATCTACAGCTTCCGGACAAGAGCGACCG | 5 (0.000096%) | | | TCCGGGCGTCGGCCTGTGGGCTCCCCATTCGACCCGTCTTGAAACACGGACCAAGGAGTCTGACATGTGTGCGAGTCAACGGGTGAGTAAACCCGTAAGG | 4 (0.000077%) | | | TCCGGGGAGGCGAATGCCAGCCGTTCGTTTGCATGTTCCTTGACACTTTTCGTGCCGGGGTTTTGTGATATCCGGAAGCAACGCGCACGACAAGACCGAG | 7 (0.000134%) | | | TCCGGTTAAAATTCCGGAACCGGGACGTGGCGGTTGACGGCAACGTTAGGGAGTCCGGAGACGTCGGCGGGGGCCTCGGGAAGAGTTATCTTTTCTGTTT | 30 (0.000575%) | | | TCCGTCGAGTTATCATGAATCATCAGAGCAACGGGCAGAGCCCGCGTCGACCTTTTATCTAATAAATGCGTCCCTTCCATAAGTCGGGGTTTGTTGCACG | 17 (0.000326%) | | | TCCGTCTCCGGGGAGGCGAATGCCAGCCGTTCGTTTGCATGTTCCTTGACACTTTTCGTGCCGGGGTTTTGTGATATCCGGAAGCAACGCGCACGACAAG | 791 (0.015159%) | | | TCCTATGATGTTATCCCATGCTAATGTATCCAGAGCGTAGGCTTGCTTTGAGCACTCTAATTTCTTCAAAGTAACAGCGCCGGAGGCACGACCCGGCCAA | 7 (0.000134%) | | | TCCTCGTTAAGGGATTTAGATTGTACTCATTCCAATTACCAGACTCGAAAGAGCCCGGTATTGTTATTTATTGTCACTACCTCCCCGTGTCAGGATTGGG | 25 (0.000479%) | | | TCCTTGTTAGAAGACACAAAGCCAAAGACTCATATGGACTTTGGCTACACCATGAAAGCTTTGAGAAGCAAGAAGAAGGTTGGTTAGTGTTTTGGAGTCG | 9 (0.000172%) | | | TCGAAAGAGCCCGGTATTGTTATTTATTGTCACTACCTCCCCGTGTCAGGATTGGGTAATTTGCGCGCCTGCTGCCTTCCTTGGATGTGGTAGCCGTTTC | 8 (0.000153%) | | | TCGAAATCCGCTAAGGAGTGTGTAACAACTCACCTGCCGAATCAACTAGCCCCGAAAATGGATGGCGCTTAAGCGCGCGACCTATACCCGGCCGTCGGGG | 50 (0.000958%) | | | TCGAAATCCTATGATGTTATCCCATGCTAATGTATCCAGAGCGTAGGCTTGCTTTGAGCACTCTAATTTCTTCAAAGTAACAGCGCCGGAGGCACGACCC | 257 (0.004925%) | | | TCGAATATGACTTGATGTCATGTGTATGATTGAGTATAAG | 8791 (0.067390%) | | | TCGACCTTTTATCTAATAAATGCGTCCCTTCCATAAGTCGGGGTTTGTTGCACGTATTAGCTCTAGAATTACTACGGTTATCCGAGTAGTAGTTACCATC | 12 (0.000230%) | | | TCGACTCCAAAACACTAACCAACCTTCTTCTTGCTTCTCAAAGCTTTCATGGTGTAGCCAAAGTCCATATGAGTCTTTGGCTTTGTGTCTTCTAACAAGG | 22 (0.000422%) | | | TCGAGTTATCATGAATCATCAGAGCAACGGGCAGAGCCCGCGTCGACCTTTTATCTAATAAATGCGTCCCTTCCATAAGTCGGGGTTTGTTGCACGTATT | 62 (0.001188%) | | | TCGATCAAGTTCACCACCGCATGTCGGTACGCTCCAGGCGTCCTTGGCTCGGATTTAGGCCAACCGCGTGCGGTAACACACGGGAGACCAGCTTCCGTCC | 3 (0.000057%) | | | TCGATCCATGCTTTCCAACGAAGCACGCCCATCCAACCTAGGCGAGACAAGGGTTCACATTTCGTTCATCACCCTTGGCCGGCTATCGAACAGCCGGACT | 9 (0.000172%) | | | TCGATCCGTAACTTCGGGAAAAGGATTGGCTCTGAGGGCTGGGCTCGGGGGTCCCAGTTCCGAACCCGTCGGCTGTCAGCGGACTGCTCGAGCTGCTTCC | 12 (0.000230%) | | | TCGATCCTCGTTAAGGGATTTAGATTGTACTCATTCCAATTACCAGACTCGAAAGAGCCCGGTATTGTTATTTATTGTCACTACCTCCCCGTGTCAGGAT | 3 (0.000057%) | | | TCGATCGAAATCCTATGATGTTATCCCATGCTAATGTATCCAGAGCGTAGGCTTGCTTTGAGCACTCTAATTTCTTCAAAGTAACAGCGCCGGAGGCACG | 6 (0.000115%) | | | TCGATCGGGTTGCGGTTTAAGTTGTTATACTCAATCATAC | 10 (0.000077%) | | | TCGCCCATGGGCATCGACACCTTGCGGCTAGGAACTGGAACGAGACGGGTGGCAAAGATTTCGAGTAGCACTTCATACTACCGTGGGTTTTTTAAACCTT | 4 (0.000077%) | | | TCGCCGCCCGCGACGTCGCGAGAAGTCCACTAAACCTTATCATTTAGAGGAAGGAGAAGTCGTAACAAGGTTTCCGTAGGTGAACCTGCGGAAGGATCGA | 5 (0.000096%) | | | TCGCCGTTACTAAGGGAATCCTTGTTAGTTTCTTTTCCTCCGCTTATTGATATGCTTAAACTCAGCGGGTAATCCCGCCTGACCTGGGGTCGCTATATGG | 3 (0.000057%) | | | TCGGCATCAGCGTGCTCCGGGCGTCGGCCTGTGGGCTCCCCATTCGACCCGTCTTGAAACACGGACCAAGGAGTCTGACATGTGTGCGAGTCAACGGGTG | 28 (0.000537%) | | | TCGGCCTGTGGGCTCCCCATTCGACCCGTCTTGAAACACGGACCAAGGAGTCTGACATGTGTGCGAGTCAACGGGTGAGTAAACCCGTAAGGCGCAAGGA | 7 (0.000134%) | | | TCGGGTTGCGGTTTAAGTTGTTATACTCAATCATACACAT | 117 (0.000897%) | | | TCGGTACGCTCCAGGCGTCCTTGGCTCGGATTTAGGCCAACCGCGTGCGGTAACACACGGGAGACCAGCTTCCGTCCCGCATCAGCAAAGGATGGTGAGG | 4 (0.000077%) | | | TCGGTAGGAGCGACGGGCGGTGTGTACAAAGGGCAGGGACGTAGTCAACGCGAGCTGATGACTCGCGCTTACTAGGAATTCCTCGTTGAAGACCAACAAT | 33 (0.000632%) | | | TCGTACTGAAAATCAGAATCAAACGAGCTTTTACCCTTTTGTTCCACACGAGATTTCTGTTCTCGTTGAGCTCATCTTAGGACACCTGCGTTATCTTTTA | 5 (0.000096%) | | | TCGTATTCGTACTGAAAATCAGAATCAAACGAGCTTTTACCCTTTTGTTCCACACGAGATTTCTGTTCTCGTTGAGCTCATCTTAGGACACCTGCGTTAT | 6 (0.000115%) | | | TCGTATTTCATAGTCAGAGGTGAAATTCTTGGATTTATGAAAGACGAACAACTGCGAAAGCATTTGCCAAGGATGTTTTCATTAATCAAGAACGAAAGTT | 7 (0.000134%) | | | TCGTCCCTCACCATCCTTTGCTGATGCGGGACGGAAGCTGGTCTCCCGTGTGTTACCGCACGCGGTTGGCCTAAATCCGAGCCAAGGACGCCTGGAGCGT | 5 (0.000096%) | | | TCGTTAAGGGATTTAGATTGTACTCATTCCAATTACCAGACTCGAAAGAGCCCGGTATTGTTATTTATTGTCACTACCTCCCCGTGTCAGGATTGGGTAA | 8 (0.000153%) | | | TCTAGTCCGACTTTGTGAAATGACTTGAGAGGTGTAGGATAAGTGGGAGCTTCGGCGCAAGTGAAATACCACTACTTTTAACGTTATTTTACTTACTCCG | 19 (0.000364%) | | | TCTCAAAGCTTTCATGGTGTAGCCAAAGTCCATATGAGTCTTTGGCTTTGTGTCTTCTAACAAGGAAACACTACTTAGGCTTATAAGATGCGGTTGCGGT | 6 (0.000115%) | | | TCTCTCTCTCTCTCTCTCTCTCTCTCTCTCTCTCTCTCTC | 66 (0.000506%) | | | TCTGGGCCGCACGCGCGCTACACTGATGTATTCAACGAGTTCACACCTTGGCCGACAGGCCCGGGTAATCTTTGAAATTTCATCGTGATGGGGATAGATC | 3 (0.000057%) | | | TCTGGTGCCAGCAGCCGCGGTAATTCCAGCTCCAATAGCGTATATTTAAGTTGTTGCAGTTAAAAAGCTCGTAGTTGAACCTTGGGATGGGTCGGCCGGT | 14 (0.000268%) | | | TCTGTGATGCCCTTAGATGTTCTGGGCCGCACGCGCGCTACACTGATGTATTCAACGAGTTCACACCTTGGCCGACAGGCCCGGGTAATCTTTGAAATTT | 10 (0.000192%) | | | TCTTAAAAGCCTAAGTAGTGTTTCCTTGTTAGAAGACACAAAGCCAAAGACTCATATGGACTTTGGCTACACCATGAAAGCTTTGAGAAGCAAGAAGAAG | 24 (0.000460%) | | | TCTTAAAGGCGTAAGAATTGTATCCTTGTTAAAAGACACAAAGCCAAAGACTCATATGGACTTTGGCTACACCATGAAAGCTTTGAGAAGCAAGAAGAAG | 86 (0.001648%) | | | TCTTAAAGGCGTAAGAATTGTATCCTTGTTAGAAGACACAAAGCCAAAGACTCATATGGACTTTGGCTACACCATGAAAGCTTTGAGAAGCAAGAAGAAG | 88 (0.001686%) | | | TCTTATACTCAATCATACACATGACATCAAGTCATATTCGACTCCAAAACACTAACCAACCTTCTTCTTGCTTCTCAAAGCTTTCATGGTGTAGCCAAAG | 221 (0.004235%) | | | TCTTCAACGAGGAATTCCTAGTAAGCGCGAGTCATCAGCTCGCGTTGACTACGTCCCTGCCCTTTGTACACACCGCCCGTCGCTCCTACCGATTGAATGA | 12 (0.000230%) | | | TCTTTTCCTCCGCTTATTGATATGCTTAAACTCAGCGGGTAATCCCGCCTGACCTGGGGTCGCTATATGGACTTTGGGTCATCTACAGCTTCCGGACAAG | 4 (0.000077%) | | | TGAAAATCAGAATCAAACGAGCTTTTACCCTTTTGTTCCACACGAGATTTCTGTTCTCGTTGAGCTCATCTTAGGACACCTGCGTTATCTTTTAACAGAT | 46 (0.000882%) | | | TGAATCATCAGAGCAACGGGCAGAGCCCGCGTCGACCTTTTATCTAATAAATGCGTCCCTTCCATAAGTCGGGGTTTGTTGCACGTATTAGCTCTAGAAT | 10 (0.000192%) | | | TGACATCAAGTCATATTCGACTCCAAAACACTAACCAACCTTCTTCTTGCTTCTCAAAGCTTTCATGGTGTAGCCAAAGTCCATATGAGTCTTTGGCTTT | 31 (0.000594%) | | | TGACTCTAGTCCGACTTTGTGAAATGACTTGAGAGGTGTAGGATAAGTGGGAGCTTCGGCGCAAGTGAAATACCACTACTTTTAACGTTATTTTACTTAC | 11 (0.000211%) | | | TGACTTGATGTCATGTGTATGATTGAGTATAAGAACTTAA | 87 (0.000667%) | | | TGAGCGAGGTGTGAGTGTCGCCCATGGGCATCGACACCTTGCGGCTAGGAACTGGAACGAGACGGGTAGCAAAGATTTCGAGTAGCACTTCATACTACCG | 3 (0.000057%) | | | TGAGCGAGGTGTGAGTGTCGCCCATGGGCATCGACACCTTGCGGCTAGGAACTGGAACGAGACGGGTGGCAAAGATTTCGAGTAGCACTTCATACTACCG | 4 (0.000077%) | | | TGAGCTTGACTCTAGTCCGACTTTGTGAAATGACTTGAGAGGTGTAGGATAAGTGGGAGCTTCGGCGCAAGTGAAATACCACTACTTTTAACGTTATTTT | 9 (0.000172%) | | | TGAGTGTCGCCCATGGGCATCGACACCTTGCGGCTAGGAACTGGAACGAGACGGGTGGCAAAGATTTCGAGTAGCACTTCATACTACCGTGGGTTTTTTA | 7 (0.000134%) | | | TGATGCCCTTAGATGTTCTGGGCCGCACGCGCGCTACACTGATGTATTCAACGAGTTCACACCTTGGCCGACAGGCCCGGGTAATCTTTGAAATTTCATC | 11 (0.000211%) | | | TGATGCGGGACGGAAGCTGGTCTCCCGTGTGTTACCGCACGCGGTTGGCCTAAATCCGAGCCAAGGACGCCTGGAGCGTACCGACATGCGGTGGTGAACT | 6 (0.000115%) | | | TGATGTCATGTGTATGATTGAGTATAAGAACTTAAACCGC | 104 (0.000797%) | | | TGATGTTATCCCATGCTAATGTATCCAGAGCGTAGGCTTGCTTTGAGCACTCTAATTTCTTCAAAGTAACAGCGCCGGAGGCACGACCCGGCCAATTAAG | 31 (0.000594%) | | | TGATTAACAGGGACAGTCGGGGGCATTCGTATTTCATAGTCAGAGGTGAAATTCTTGGATTTATGAAAGACGAACAACTGCGAAAGCATTTGCCAAGGAT | 112 (0.002146%) | | | TGCAGCACGCGCCTAACGGCGTGCCTCGGCATCAGCGTGCTCCGGGCGTCGGCCTGTGGGCTCCCCATTCGACCCGTCTTGAAACACGGACCAAGGAGTC | 88 (0.001686%) | | | TGCATGTGTAAGTATGAACGAATTCAGACTGTGAAACTGCGAATGGCTCATTAAATCAGTTATAGTTTGTTTGATGGTAACTACTACTCGGATAACCGTA | 11 (0.000211%) | | | TGCCAGCAGCCGCGGTAATTCCAGCTCCAATAGCGTATATTTAAGTTGTTGCAGTTAAAAAGCTCGTAGTTGAACCTTGGGATGGGTCGGCCGGTCCGCC | 4 (0.000077%) | | | TGCCAGTAGTCATATGCTTGTCTCAAAGATTAAGCCATGCATGTGTAAGTATGAACGAATTCAGACTGTGAAACTGCGAATGGCTCATTAAATCAGTTAT | 125 (0.002396%) | | | TGCCCTTAGATGTTCTGGGCCGCACGCGCGCTACACTGATGTATTCAACGAGTTCACACCTTGGCCGACAGGCCCGGGTAATCTTTGAAATTTCATCGTG | 5 (0.000096%) | | | TGCCGACTTCCCTTGCCTACATTGTTCCATCGACCAGAGGCTGTTCACCTTGGAGACCTGATGCGGTTATGAGTACGACCGGGCGTGAGCGGCACTCGGT | 135 (0.002587%) | | | TGCCTACATTGTTCCATCGACCAGAGGCTGTTCACCTTGGAGACCTGATGCGGTTATGAGTACGACCGGGCGTGAGCGGCACTCGGTCCTCCGGATTTTC | 9 (0.000172%) | | | TGCGAATGGCTCATTAAATCAGTTATAGTTTGTTTGATGGTAACTACTACTCGGATAACCGTAGTAATTCTAGAGCTAATACGTGCAACAAACCCCGACT | 8 (0.000153%) | | | TGCGGTTTAAGTTCTTATACTCAATCATACACATGACATC | 3708 (0.028425%) | | | TGCTAATGTATCCAGAGCGTAGGCTTGCTTTGAGCACTCTAATTTCTTCAAAGTAACAGCGCCGGAGGCACGACCCGGCCAATTAAGACCAGGAGCGTAT | 21 (0.000402%) | | | TGCTGATGCGGGACGGAAGCTGGTCTCCCGTGTGTTACCGCACGCGGTTGGCCTAAATCCGAGCCAAGGACGCCTGGAGCGTACCGACATGCGGTGGTGA | 9 (0.000172%) | | | TGCTTCTCAAAGCTTTCATGGTGTAGCCAAAGTCCATATGAGTCTTTGGCTTTGTGTCTTCTAACAAGGAAACACTACTTAGGCTTATAAGATGCGGTTG | 6 (0.000115%) | | | TGCTTGTCTCAAAGATTAAGCCATGCATGTGTAAGTATGAACGAATTCAGACTGTGAAACTGCGAATGGCTCATTAAATCAGTTATAGTTTGTTTGATGG | 7 (0.000134%) | | | TGCTTTCCAACGAAGCACGCCCATCCAACCTAGGCGAGACAAGGGTTCACATTTCGTTCATCACCCTTGGCCGGCTATCGAACAGCCGGACTCCCATCAA | 26 (0.000498%) | | | TGGAAAGAGAGAAAGGACGAGGTCTTGACCGTCATCTTTTGCCCGAAGGACGGATGAGCTTTGGCGGGACTGAATCACTTCGAGTCACCGTCGACAACTT | 10 (0.000192%) | | | TGGAAGTCGAAATCCGCTAAGGAGTGTGTAACAACTCACCTGCCGAATCAACTAGCCCCGAAAATGGATGGCGCTTAAGCGCGCGACCTATACCCGGCCG | 957 (0.018340%) | | | TGGACTTTGGCTACACCATGAAAGCTTTGAGAAGCAAGAAGAAGGTTGGTTAGTGTTTTGGAGTCGAATATGACTTGATGTCATGTGTATGATTGAGTAT | 41 (0.000786%) | | | TGGAGGGCAAGTCTGGTGCCAGCAGCCGCGGTAATTCCAGCTCCAATAGCGTATATTTAAGTTGTTGCAGTTAAAAAGCTCGTAGTTGAACCTTGGGATG | 322 (0.006171%) | | | TGGCTGGGGCGGCACATCTGTTAAAAGATAACGCAGGTGTCCTAAGATGAGCTCAACGAGAACAGAAATCTCGTGTGGAACAAAAGGGTAAAAGCTCGTT | 3 (0.000057%) | | | TGGGCATCGACACCTTGCGGCTAGGAACTGGAACGAGACGGGTGGCAAAGATTTCGAGTAGCACTTCATACTACCGTGGGTTTTTTAAACCTTCCGAGTT | 28 (0.000537%) | | | TGGGCCGCACGCGCGCTACACTGATGTATTCAACGAGTTCACACCTTGGCCGACAGGCCCGGGTAATCTTTGAAATTTCATCGTGATGGGGATAGATCGA | 14 (0.000268%) | | | TGGGGCGGCACATCTGTTAAAAGATAACGCAGGTGTCCTAAGATGAGCTCAACGAGAACAGAAATCTCGTGTGGAACAAAAGGGTAAAAGCTCGTTTGAT | 12 (0.000230%) | | | TGGGTGGTTCGCCGCCCGCGACGTCGCGAGAAGTCCACTAAACCTTATCATTTAGAGGAAGGAGAAGTCGTAACAAGGTTTCCGTAGGTGAACCTGCGGA | 10 (0.000192%) | | | TGGTCTTCAACGAGGAATTCCTAGTAAGCGCGAGTCATCAGCTCGCGTTGACTACGTCCCTGCCCTTTGTACACACCGCCCGTCGCTCCTACCGATTGAA | 65 (0.001246%) | | | TGGTGCCAGCAGCCGCGGTAATTCCAGCTCCAATAGCGTATATTTAAGTTGTTGCAGTTAAAAAGCTCGTAGTTGAACCTTGGGATGGGTCGGCCGGTCC | 5 (0.000096%) | | | TGGTTCGCCGCCCGCGACGTCGCGAGAAGTCCACTAAACCTTATCATTTAGAGGAAGGAGAAGTCGTAACAAGGTTTCCGTAGGTGAACCTGCGGAAGGA | 6 (0.000115%) | | | TGTACTCATTCCAATTACCAGACTCGAAAGAGCCCGGTATTGTTATTTATTGTCACTACCTCCCCGTGTCAGGATTGGGTAATTTGCGCGCCTGCTGCCT | 5 (0.000096%) | | | TGTAGCCAAAGTCCATATGAGTCTTTGGCTTTGTGTCTTT | 6689 (0.051276%) | | | TGTATCCAGAGCGTAGGCTTGCTTTGAGCACTCTAATTTCTTCAAAGTAACAGCGCCGGAGGCACGACCCGGCCAATTAAGACCAGGAGCGTATCGCCGA | 17 (0.000326%) | | | TGTATCCTTGTTAGAAGACACAAAGCCAAAGACTCATATGGACTTTGGCTACACCATGAAAGCTTTGAGAAGCAAGAAGAAGGTTGGTTAGTGTTTTGGA | 28 (0.000537%) | | | TGTCAGGTGGGGAGTTTGGCTGGGGCGGCACATCTGTTAAAAGATAACGCAGGTGTCCTAAGATGAGCTCAACGAGAACAGAAATCTCGTGTGGAACAAA | 42 (0.000805%) | | | TGTCATGTGTATGATTGAGTATAAGAACTTAAACCGCAAC | 635 (0.004868%) | | | TGTCCCGAGTGTGAGCGAGGTGTGAGTGTCGCCCATGGGCATCGACACCTTGCGGCTAGGAACTGGAACGAGACGGGTAGCAAAGATTTCGAGTAGCACT | 5 (0.000096%) | | | TGTCCCGAGTGTGAGCGAGGTGTGAGTGTCGCCCATGGGCATCGACACCTTGCGGCTAGGAACTGGAACGAGACGGGTGGCAAAGATTTCGAGTAGCACT | 4 (0.000077%) | | | TGTCGCCCATGGGCATCGACACCTTGCGGCTAGGAACTGGAACGAGACGGGTGGCAAAGATTTCGAGTAGCACTTCATACTACCGTGGGTTTTTTAAACC | 4 (0.000077%) | | | TGTCGGTACGCTCCAGGCGTCCTTGGCTCGGATTTAGGCCAACCGCGTGCGGTAACACACGGGAGACCAGCTTCCGTCCCGCATCAGCAAAGGATGGTGA | 3 (0.000057%) | | | TGTGAGCGAGGTGTGAGTGTCGCCCATGGGCATCGACACCTTGCGGCTAGGAACTGGAACGAGACGGGTGGCAAAGATTTCGAGTAGCACTTCATACTAC | 3 (0.000057%) | | | TGTGAGTGTCGCCCATGGGCATCGACACCTTGCGGCTAGGAACTGGAACGAGACGGGTAGCAAAGATTTCGAGTAGCACTTCATACTACCGTGGGTTTTT | 9 (0.000172%) | | | TGTGAGTGTCGCCCATGGGCATCGACACCTTGCGGCTAGGAACTGGAACGAGACGGGTGGCAAAGATTTCGAGTAGCACTTCATACTACCGTGGGTTTTT | 12 (0.000230%) | | | TGTGATGCCCTTAGATGTTCTGGGCCGCACGCGCGCTACACTGATGTATTCAACGAGTTCACACCTTGGCCGACAGGCCCGGGTAATCTTTGAAATTTCA | 6 (0.000115%) | | | TGTTAGAAGACACAAAGCCAAAGACTCATATGGACTTTGGCTACACCATGAAAGCTTTGAGAAGCAAGAAGAAGGTTGGTTAGTGTTTTGGAGTCGAATA | 27 (0.000517%) | | | TGTTAGTTTCTTTTCCTCCGCTTATTGATATGCTTAAACTCAGCGGGTAATCCCGCCTGACCTGGGGTCGCTATATGGACTTTGGGTCATCTACAGCTTC | 7 (0.000134%) | | | TGTTATCCCATGCTAATGTATCCAGAGCGTAGGCTTGCTTTGAGCACTCTAATTTCTTCAAAGTAACAGCGCCGGAGGCACGACCCGGCCAATTAAGACC | 9 (0.000172%) | | | TGTTGAGCTTGACTCTAGTCCGACTTTGTGAAATGACTTGAGAGGTGTAGGATAAGTGGGAGCTTCGGCGCAAGTGAAATACCACTACTTTTAACGTTAT | 3 (0.000057%) | | | TGTTGGTCTTCAACGAGGAATTCCTAGTAAGCGCGAGTCATCAGCTCGCGTTGACTACGTCCCTGCCCTTTGTACACACCGCCCGTCGCTCCTACCGATT | 184 (0.003526%) | | | TGTTTCCTTGTTAGAAGACACAAAGCCAAAGACTCATATGGACTTTGGCTACACCATGAAAGCTTTGAGAAGCAAGAAGAAGGTTGGTTAGTGTTTTGGA | 24 (0.000460%) | | | TTAAAAGATAACGCAGGTGTCCTAAGATGAGCTCAACGAGAACAGAAATCTCGTGTGGAACAAAAGGGTAAAAGCTCGTTTGATTCTGATTTTCAGTACG | 25 (0.000479%) | | | TTAAAAGCCTAAGTAGTGTTTCCTTGTTAGAAGACACAAAGCCAAAGACTCATATGGACTTTGGCTACACCATGAAAGCTTTGAGAAGCAAGAAGAAGGT | 147 (0.002817%) | | | TTAAAGGCGTAAGAATTGTATCCTTGTTAAAAGACACAAAGCCAAAGACTCATATGGACTTTGGCTACACCATGAAAGCTTTGAGAAGCAAGAAGAAGGT | 126 (0.002415%) | | | TTAAAGGCGTAAGAATTGTATCCTTGTTAGAAGACACAAAGCCAAAGACTCATATGGACTTTGGCTACACCATGAAAGCTTTGAGAAGCAAGAAGAAGGT | 102 (0.001955%) | | | TTAAATCAGTTATAGTTTGTTTGATGGTAACTACTACTCGGATAACCGTAGTAATTCTAGAGCTAATACGTGCAACAAACCCCGACTTATGGAAGGGACG | 4 (0.000077%) | | | TTAACAGGGACAGTCGGGGGCATTCGTATTTCATAGTCAGAGGTGAAATTCTTGGATTTATGAAAGACGAACAACTGCGAAAGCATTTGCCAAGGATGTT | 202 (0.003871%) | | | TTAAGCCATGCATGTGTAAGTATGAACGAATTCAGACTGTGAAACTGCGAATGGCTCATTAAATCAGTTATAGTTTGTTTGATGGTAACTACTACTCGGA | 15 (0.000287%) | | | TTAAGCGCGCGACCTATACCCGGCCGTCGGGGCAAGAGCCAGGCCTCGATGAGTAGGAGGGCGCGGCGGTCGCTGCAAAACCTAGGGCGCGAGCCCGGGC | 13 (0.000249%) | | | TTAAGGGATTTAGATTGTACTCATTCCAATTACCAGACTCGAAAGAGCCCGGTATTGTTATTTATTGTCACTACCTCCCCGTGTCAGGATTGGGTAATTT | 19 (0.000364%) | | | TTAAGTTCTTATACTCAATCATACACATGACATCAAGTCATATTCGACTCCAAAACACTAACCAACCTTCTTCTTGCTTCTCAAAGCTTTCATGGTGTAG | 2935 (0.056248%) | | | TTAAGTTGTTATACTCAATCATACACATGACAACAAGTCATATTCGACTCCAAAACACTAACCAACCTTCTTCTTGCTTCTCAAAGCTTTCATGGTGTAG | 70 (0.001342%) | | | TTAAGTTGTTATACTCAATCATACACATGACATCAAGTCA | 474 (0.003634%) | | | TTACCAGACTCGAAAGAGCCCGGTATTGTTATTTATTGTCACTACCTCCCCGTGTCAGGATTGGGTAATTTGCGCGCCTGCTGCCTTCCTTGGATGTGGT | 30 (0.000575%) | | | TTACGGGTTTACTCACCCGTTGACTCGCACACATGTCAGACTCCTTGGTCCGTGTTTCAAGACGGGTCGAATGGGGAGCCCACAGGCCGACGCCCGGAGC | 3313 (0.063492%) | | | TTACTCACCCGTTGACTCGCACACATGTCAGACTCCTTGGTCCGTGTTTCAAGACGGGTCGAATGGGGAGCCCACAGGCCGACGCCCGGAGCACGCTGAT | 11 (0.000211%) | | | TTAGAAGACACAAAGCCAAAGACTCATATGGACTTTGGCTACACCATGAAAGCTTTGAGAAGCAAGAAGAAGGTTGGTTAGTGTTTTGGAGTCGAATATG | 83 (0.001591%) | | | TTAGATGTTCTGGGCCGCACGCGCGCTACACTGATGTATTCAACGAGTTCACACCTTGGCCGACAGGCCCGGGTAATCTTTGAAATTTCATCGTGATGGG | 15 (0.000287%) | | | TTAGATTGTACTCATTCCAATTACCAGACTCGAAAGAGCCCGGTATTGTTATTTATTGTCACTACCTCCCCGTGTCAGGATTGGGTAATTTGCGCGCCTG | 26 (0.000498%) | | | TTAGTTTCTTTTCCTCCGCTTATTGATATGCTTAAACTCAGCGGGTAATCCCGCCTGACCTGGGGTCGCTATATGGACTTTGGGTCATCTACAGCTTCCG | 4 (0.000077%) | | | TTATACTCAATCATACACATGACAACAAGTCATATTCGACTCCAAAACACTAACCAACCTTCTTCTTGCTTCTCAAAGCTTTCATGGTGTAGCCAAAGTC | 239 (0.004580%) | | | TTATACTCAATCATACACATGACATCAAGTCATATTCGACTCCAAAACACTAACCAACCTTCTTCTTGCTTCTCAAAGCTTTCATGGTGTAGCCAAAGTC | 774 (0.014833%) | | | TTATAGTTTGTTTGATGGTAACTACTACTCGGATAACCGTAGTAATTCTAGAGCTAATACGTGCAACAAACCCCGACTTATGGAAGGGACGCATTTATTA | 6 (0.000115%) | | | TTATCATGAATCATCAGAGCAACGGGCAGAGCCCGCGTCGACCTTTTATCTAATAAATGCGTCCCTTCCATAAGTCGGGGTTTGTTGCACGTATTAGCTC | 66 (0.001265%) | | | TTATCCCATGCTAATGTATCCAGAGCGTAGGCTTGCTTTGAGCACTCTAATTTCTTCAAAGTAACAGCGCCGGAGGCACGACCCGGCCAATTAAGACCAG | 5 (0.000096%) | | | TTATCTAATAAATGCGTCCCTTCCATAAGTCGGGGTTTGTTGCACGTATTAGCTCTAGAATTACTACGGTTATCCGAGTAGTAGTTACCATCAAACAAAC | 4 (0.000077%) | | | TTATTGATATGCTTAAACTCAGCGGGTAATCCCGCCTGACCTGGGGTCGCTATATGGACTTTGGGTCATCTACAGCTTCCGGACAAGAGCGACCGATAAA | 16 (0.000307%) | | | TTCAACGAGGAATTCCTAGTAAGCGCGAGTCATCAGCTCGCGTTGACTACGTCCCTGCCCTTTGTACACACCGCCCGTCGCTCCTACCGATTGAATGATC | 14 (0.000268%) | | | TTCAATCGGTAGGAGCGACGGGCGGTGTGTACAAAGGGCAGGGACGTAGTCAACGCGAGCTGATGACTCGCGCTTACTAGGAATTCCTCGTTGAAGACCA | 72 (0.001380%) | | | TTCACATTTCGTTCATCACCCTTGGCCGGCTATCGAACAGCCGGACTCCCATCAAAAGATGGTTGCCAAGAACATCTTCGTTACGGTTTGCTAATTCTCG | 24 (0.000460%) | | | TTCACATTTCGTTCATCACCCTTGGCCGGCTTTCGAACAGCCGGACTCCCATCAAAAGATGGTTGCCAAGAACATCTTCGTTACGGTTTGCTAATTCTCG | 25 (0.000479%) | | | TTCACCACCGCATGTCGGTACGCTCCAGGCGTCCTTGGCTCGGATTTAGGCCAACCGCGTGCGGTAACACACGGGAGACCAGCTTCCGTCCCGCATCAGC | 8 (0.000153%) | | | TTCACGGTTCGTATTCGTACTGAAAATCAGAATCAAACGAGCTTTTACCCTTTTGTTCCACACGAGATTTCTGTTCTCGTTGAGCTCATCTTAGGACACC | 9 (0.000172%) | | | TTCATAATTTTTTGACACCTCTAGCTAGGTCATTTGACCTGATACAACATCGGATTTTCATGGTCTAGTTGGGGCTCCGTGGGCATATTTGATGCAAACT | 3 (0.000057%) | | | TTCATAGTCAGAGGTGAAATTCTTGGATTTATGAAAGACGAACAACTGCGAAAGCATTTGCCAAGGATGTTTTCATTAATCAAGAACGAAAGTTGGGGGC | 33 (0.000632%) | | | TTCCAACGAAGCACGCCCATCCAACCTAGGCGAGACAAGGGTTCACATTTCGTTCATCACCCTTGGCCGGCTATCGAACAGCCGGACTCCCATCAAAAGA | 110 (0.002108%) | | | TTCCAACGAAGCACGCCCATCCAACCTAGGCGAGACAAGGGTTCACATTTCGTTCATCACCCTTGGCCGGCTTTCGAACAGCCGGACTCCCATCAAAAGA | 2303 (0.044136%) | | | TTCCAATTACCAGACTCGAAAGAGCCCGGTATTGTTATTTATTGTCACTACCTCCCCGTGTCAGGATTGGGTAATTTGCGCGCCTGCTGCCTTCCTTGGA | 10 (0.000192%) | | | TTCCCCGCCACATCCTCTCAAACGCAATGGAAAGAGAGAAAGGACGAGGTCTTGACCGTCATCTTTTGCCCGAAGGACGGATGAGCTTTGGCGGGACTGA | 8 (0.000153%) | | | TTCCCTTGCCTACATTGTTCCATCGACCAGAGGCTGTTCACCTTGGAGACCTGATGCGGTTATGAGTACGACCGGGCGTGAGCGGCACTCGGTCCTCCGG | 10 (0.000192%) | | | TTCCTTGTTAGAAGACACAAAGCCAAAGACTCATATGGACTTTGGCTACACCATGAAAGCTTTGAGAAGCAAGAAGAAGGTTGGTTAGTGTTTTGGAGTC | 55 (0.001054%) | | | TTCGACTCCAAAACACTAACCAACCTTCTTCTTGCTTCTCAAAGCTTTCATGGTGTAGCCAAAGTCCATATGAGTCTTTGGCTTTGTGTCTTCTAACAAG | 116 (0.002223%) | | | TTCGTACTGAAAATCAGAATCAAACGAGCTTTTACCCTTTTGTTCCACACGAGATTTCTGTTCTCGTTGAGCTCATCTTAGGACACCTGCGTTATCTTTT | 24 (0.000460%) | | | TTCGTATTCGTACTGAAAATCAGAATCAAACGAGCTTTTACCCTTTTGTTCCACACGAGATTTCTGTTCTCGTTGAGCTCATCTTAGGACACCTGCGTTA | 16 (0.000307%) | | | TTCGTATTTCATAGTCAGAGGTGAAATTCTTGGATTTATGAAAGACGAACAACTGCGAAAGCATTTGCCAAGGATGTTTTCATTAATCAAGAACGAAAGT | 18 (0.000345%) | | | TTCTCAAAGCTTTCATGGTGTAGCCAAAGTCCATATGAGTCTTTGGCTTTGTGTCTTCTAACAAGGAAACACTACTTAGGCTTATAAGATGCGGTTGCGG | 11 (0.000211%) | | | TTCTGGGCCGCACGCGCGCTACACTGATGTATTCAACGAGTTCACACCTTGGCCGACAGGCCCGGGTAATCTTTGAAATTTCATCGTGATGGGGATAGAT | 3 (0.000057%) | | | TTCTTATACTCAATCATACACATGACATCAAGTCATATTCGACTCCAAAACACTAACCAACCTTCTTCTTGCTTCTCAAAGCTTTCATGGTGTAGCCAAA | 105 (0.002012%) | | | TTCTTGCTTCTCAAAGCTTTCATGGTGTAGCCAAAGTCCATATGAGTCTTTGGCTTTGTGTCTTCTAACAAGGAAACACTACTTAGGCTTATAAGATGCG | 271 (0.005194%) | | | TTCTTTTCCTCCGCTTATTGATATGCTTAAACTCAGCGGGTAATCCCGCCTGACCTGGGGTCGCTATATGGACTTTGGGTCATCTACAGCTTCCGGACAA | 38 (0.000728%) | | | TTGACTCTAGTCCGACTTTGTGAAATGACTTGAGAGGTGTAGGATAAGTGGGAGCTTCGGCGCAAGTGAAATACCACTACTTTTAACGTTATTTTACTTA | 16 (0.000307%) | | | TTGAGCTTGACTCTAGTCCGACTTTGTGAAATGACTTGAGAGGTGTAGGATAAGTGGGAGCTTCGGCGCAAGTGAAATACCACTACTTTTAACGTTATTT | 30 (0.000575%) | | | TTGATCGATC | 180377 (0.345683%) | | | TTGATGGTAACTACTACTCGGATAACCGTAGTAATTCTAGAGCTAATACGTGCAACAAACCCCGACTTATGGAAGGGACGCATTTATTAGATAAAAGGTC | 21 (0.000402%) | | | TTGATGTCATGTGTATGATTGAGTATAAGAACTTAAACCG | 368 (0.002821%) | | | TTGCCGACTTCCCTTGCCTACATTGTTCCATCGACCAGAGGCTGTTCACCTTGGAGACCTGATGCGGTTATGAGTACGACCGGGCGTGAGCGGCACTCGG | 125 (0.002396%) | | | TTGCCTACATTGTTCCATCGACCAGAGGCTGTTCACCTTGGAGACCTGATGCGGTTATGAGTACGACCGGGCGTGAGCGGCACTCGGTCCTCCGGATTTT | 7 (0.000134%) | | | TTGCGGTTTAAGTTCTTATACTCAATCATACACATGACAT | 2025 (0.015523%) | | | TTGCGGTTTAAGTTGTTATACTCAATCATACACATGACAT | 710 (0.005443%) | | | TTGCTCCGTTCCGCATCCGACCAGGACGCATCGCCGGCCCCCATCCGCTTCCCTCCCGACAATTTCAAGCACTCTTTGACTCTCTTTTCAAAGTCCTTTT | 6 (0.000115%) | | | TTGCTGATGCGGGACGGAAGCTGGTCTCCCGTGTGTTACCGCACGCGGTTGGCCTAAATCCGAGCCAAGGACGCCTGGAGCGTACCGACATGCGGTGGTG | 4 (0.000077%) | | | TTGCTTCTCAAAGCTTTCATGGTGTAGCCAAAGTCCATATGAGTCTTTGGCTTTGTGTCTTCTAACAAGGAAACACTACTTAGGCTTATAAGATGCGGTT | 4 (0.000077%) | | | TTGCTTTGAGCACTCTAATTTCTTCAAAGTAACAGCGCCGGAGGCACGACCCGGCCAATTAAGACCAGGAGCGTATCGCCGACCGAAGGGACAAGCCGAC | 7 (0.000134%) | | | TTGGAGGGCAAGTCTGGTGCCAGCAGCCGCGGTAATTCCAGCTCCAATAGCGTATATTTAAGTTGTTGCAGTTAAAAAGCTCGTAGTTGAACCTTGGGAT | 339 (0.006497%) | | | TTGGCATGCATCATAAGGATACTAAATCCTATTTTCTGGTAAATTTTCATAATTTTTTGACACCTCTAGCTAGGTCATTTGACCTGATACAACATCGGAT | 442 (0.008471%) | | | TTGGCTGGGGCGGCACATCTGTTAAAAGATAACGCAGGTGTCCTAAGATGAGCTCAACGAGAACAGAAATCTCGTGTGGAACAAAAGGGTAAAAGCTCGT | 19 (0.000364%) | | | TTGGTCTTCAACGAGGAATTCCTAGTAAGCGCGAGTCATCAGCTCGCGTTGACTACGTCCCTGCCCTTTGTACACACCGCCCGTCGCTCCTACCGATTGA | 89 (0.001706%) | | | TTGTACTCATTCCAATTACCAGACTCGAAAGAGCCCGGTATTGTTATTTATTGTCACTACCTCCCCGTGTCAGGATTGGGTAATTTGCGCGCCTGCTGCC | 12 (0.000230%) | | | TTGTATCCTTGTTAGAAGACACAAAGCCAAAGACTCATATGGACTTTGGCTACACCATGAAAGCTTTGAGAAGCAAGAAGAAGGTTGGTTAGTGTTTTGG | 20 (0.000383%) | | | TTGTCAGGTGGGGAGTTTGGCTGGGGCGGCACATCTGTTAAAAGATAACGCAGGTGTCCTAAGATGAGCTCAACGAGAACAGAAATCTCGTGTGGAACAA | 3 (0.000057%) | | | TTGTCTCAAAGATTAAGCCATGCATGTGTAAGTATGAACGAATTCAGACTGTGAAACTGCGAATGGCTCATTAAATCAGTTATAGTTTGTTTGATGGTAA | 13 (0.000249%) | | | TTGTTAGAAGACACAAAGCCAAAGACTCATATGGACTTTGGCTACACCATGAAAGCTTTGAGAAGCAAGAAGAAGGTTGGTTAGTGTTTTGGAGTCGAAT | 59 (0.001131%) | | | TTGTTAGTTTCTTTTCCTCCGCTTATTGATATGCTTAAACTCAGCGGGTAATCCCGCCTGACCTGGGGTCGCTATATGGACTTTGGGTCATCTACAGCTT | 12 (0.000230%) | | | TTGTTGGTCTTCAACGAGGAATTCCTAGTAAGCGCGAGTCATCAGCTCGCGTTGACTACGTCCCTGCCCTTTGTACACACCGCCCGTCGCTCCTACCGAT | 195 (0.003737%) | | | TTGTTTGATGGTAACTACTACTCGGATAACCGTAGTAATTCTAGAGCTAATACGTGCAACAAACCCCGACTTATGGAAGGGACGCATTTATTAGATAAAA | 11 (0.000211%) | | | TTTAAGTTCTTATACTCAATCATACACATGACATCAAGTC | 755 (0.005788%) | | | TTTAAGTTGTTATACTCAATCATACACATGACAACAAGTCATATTCGACTCCAAAACACTAACCAACCTTCTTCTTGCTTCTCAAAGCTTTCATGGTGTA | 37 (0.000709%) | | | TTTAAGTTGTTATACTCAATCATACACATGACATCAAGTC | 325 (0.002491%) | | | TTTACTCACCCGTTGACTCGCACACATGTCAGACTCCTTGGTCCGTGTTTCAAGACGGGTCGAATGGGGAGCCCACAGGCCGACGCCCGGAGCACGCTGA | 8 (0.000153%) | | | TTTAGATTGTACTCATTCCAATTACCAGACTCGAAAGAGCCCGGTATTGTTATTTATTGTCACTACCTCCCCGTGTCAGGATTGGGTAATTTGCGCGCCT | 129 (0.002472%) | | | TTTATCTAATAAATGCGTCCCTTCCATAAGTCGGGGTTTGTTGCACGTATTAGCTCTAGAATTACTACGGTTATCCGAGTAGTAGTTACCATCAAACAAA | 4 (0.000077%) | | | TTTCACGGTTCGTATTCGTACTGAAAATCAGAATCAAACGAGCTTTTACCCTTTTGTTCCACACGAGATTTCTGTTCTCGTTGAGCTCATCTTAGGACAC | 66 (0.001265%) | | | TTTCATAATTTTTTGACACCTCTAGCTAGGTCATTTGACCTGATACAACATCGGATTTTCATGGTCTAGTTGGGGCTCCGTGGGCATATTTGATGCAAAC | 10 (0.000192%) | | | TTTCATAGTCAGAGGTGAAATTCTTGGATTTATGAAAGACGAACAACTGCGAAAGCATTTGCCAAGGATGTTTTCATTAATCAAGAACGAAAGTTGGGGG | 9 (0.000172%) | | | TTTCCAACGAAGCACGCCCATCCAACCTAGGCGAGACAAGGGTTCACATTTCGTTCATCACCCTTGGCCGGCTATCGAACAGCCGGACTCCCATCAAAAG | 304 (0.005826%) | | | TTTCCTCCGCTTATTGATATGCTTAAACTCAGCGGGTAATCCCGCCTGACCTGGGGTCGCTATATGGACTTTGGGTCATCTACAGCTTCCGGACAAGAGC | 5 (0.000096%) | | | TTTCCTTGTTAGAAGACACAAAGCCAAAGACTCATATGGACTTTGGCTACACCATGAAAGCTTTGAGAAGCAAGAAGAAGGTTGGTTAGTGTTTTGGAGT | 609 (0.011671%) | | | TTTCGTTCATCACCCTTGGCCGGCTATCGAACAGCCGGACTCCCATCAAAAGATGGTTGCCAAGAACATCTTCGTTACGGTTTGCTAATTCTCGGAATAA | 12 (0.000230%) | | | TTTCTGGTAAATTTTCATAATTTTTTGACACCTCTAGCTAGGTCATTTGACCTGATACAACATCGGATTTTCATGGTCTAGTTGGGGCTCCGTGGGCATA | 14 (0.000268%) | | | TTTCTTTTCCTCCGCTTATTGATATGCTTAAACTCAGCGGGTAATCCCGCCTGACCTGGGGTCGCTATATGGACTTTGGGTCATCTACAGCTTCCGGACA | 12 (0.000230%) | | | TTTGATGGTAACTACTACTCGGATAACCGTAGTAATTCTAGAGCTAATACGTGCAACAAACCCCGACTTATGGAAGGGACGCATTTATTAGATAAAAGGT | 8 (0.000153%) | | | TTTGCCGACTTCCCTTGCCTACATTGTTCCATCGACCAGAGGCTGTTCACCTTGGAGACCTGATGCGGTTATGAGTACGACCGGGCGTGAGCGGCACTCG | 2344 (0.044922%) | | | TTTGCTGATGCGGGACGGAAGCTGGTCTCCCGTGTGTTACCGCACGCGGTTGGCCTAAATCCGAGCCAAGGACGCCTGGAGCGTACCGACATGCGGTGGT | 30 (0.000575%) | | | TTTGGCTGGGGCGGCACATCTGTTAAAAGATAACGCAGGTGTCCTAAGATGAGCTCAACGAGAACAGAAATCTCGTGTGGAACAAAAGGGTAAAAGCTCG | 20 (0.000383%) | | | TTTGTTTGATGGTAACTACTACTCGGATAACCGTAGTAATTCTAGAGCTAATACGTGCAACAAACCCCGACTTATGGAAGGGACGCATTTATTAGATAAA | 16 (0.000307%) | | | TTTTATCTAATAAATGCGTCCCTTCCATAAGTCGGGGTTTGTTGCACGTATTAGCTCTAGAATTACTACGGTTATCCGAGTAGTAGTTACCATCAAACAA | 13 (0.000249%) | | | TTTTCATAATTTTTTGACACCTCTAGCTAGGTCATTTGACCTGATACAACATCGGATTTTCATGGTCTAGTTGGGGCTCCGTGGGCATATTTGATGCAAA | 15 (0.000287%) | | | TTTTCCTCCGCTTATTGATATGCTTAAACTCAGCGGGTAATCCCGCCTGACCTGGGGTCGCTATATGGACTTTGGGTCATCTACAGCTTCCGGACAAGAG | 47 (0.000901%) | | | TTTTCTGGTAAATTTTCATAATTTTTTGACACCTCTAGCTAGGTCATTTGACCTGATACAACATCGGATTTTCATGGTCTAGTTGGGGCTCCGTGGGCAT | 5 (0.000096%) | | | TTTTTTTTTT | 53205 (0.101965%) | | After filtering: read2: quality Value of each position will be shown on mouse over. After filtering: read2: base contents Value of each position will be shown on mouse over. After filtering: read2: KMER counting Darker background means larger counts. The count will be shown on mouse over. | | | | | | | | | | | | | | | | | | | --- | --- | --- | --- | --- | --- | --- | --- | --- | --- | --- | --- | --- | --- | --- | --- | --- | | | AA | AT | AC | AG | TA | TT | TC | TG | CA | CT | CC | CG | GA | GT | GC | GG | | AAA | AAAAA | AAAAT | AAAAC | AAAAG | AAATA | AAATT | AAATC | AAATG | AAACA | AAACT | AAACC | AAACG | AAAGA | AAAGT | AAAGC | AAAGG | | AAT | AATAA | AATAT | AATAC | AATAG | AATTA | AATTT | AATTC | AATTG | AATCA | AATCT | AATCC | AATCG | AATGA | AATGT | AATGC | AATGG | | AAC | AACAA | AACAT | AACAC | AACAG | AACTA | AACTT | AACTC | AACTG | AACCA | AACCT | AACCC | AACCG | AACGA | AACGT | AACGC | AACGG | | AAG | AAGAA | AAGAT | AAGAC | AAGAG | AAGTA | AAGTT | AAGTC | AAGTG | AAGCA | AAGCT | AAGCC | AAGCG | AAGGA | AAGGT | AAGGC | AAGGG | | ATA | ATAAA | ATAAT | ATAAC | ATAAG | ATATA | ATATT | ATATC | ATATG | ATACA | ATACT | ATACC | ATACG | ATAGA | ATAGT | ATAGC | ATAGG | | ATT | ATTAA | ATTAT | ATTAC | ATTAG | ATTTA | ATTTT | ATTTC | ATTTG | ATTCA | ATTCT | ATTCC | ATTCG | ATTGA | ATTGT | ATTGC | ATTGG | | ATC | ATCAA | ATCAT | ATCAC | ATCAG | ATCTA | ATCTT | ATCTC | ATCTG | ATCCA | ATCCT | ATCCC | ATCCG | ATCGA | ATCGT | ATCGC | ATCGG | | ATG | ATGAA | ATGAT | ATGAC | ATGAG | ATGTA | ATGTT | ATGTC | ATGTG | ATGCA | ATGCT | ATGCC | ATGCG | ATGGA | ATGGT | ATGGC | ATGGG | | ACA | ACAAA | ACAAT | ACAAC | ACAAG | ACATA | ACATT | ACATC | ACATG | ACACA | ACACT | ACACC | ACACG | ACAGA | ACAGT | ACAGC | ACAGG | | ACT | ACTAA | ACTAT | ACTAC | ACTAG | ACTTA | ACTTT | ACTTC | ACTTG | ACTCA | ACTCT | ACTCC | ACTCG | ACTGA | ACTGT | ACTGC | ACTGG | | ACC | ACCAA | ACCAT | ACCAC | ACCAG | ACCTA | ACCTT | ACCTC | ACCTG | ACCCA | ACCCT | ACCCC | ACCCG | ACCGA | ACCGT | ACCGC | ACCGG | | ACG | ACGAA | ACGAT | ACGAC | ACGAG | ACGTA | ACGTT | ACGTC | ACGTG | ACGCA | ACGCT | ACGCC | ACGCG | ACGGA | ACGGT | ACGGC | ACGGG | | AGA | AGAAA | AGAAT | AGAAC | AGAAG | AGATA | AGATT | AGATC | AGATG | AGACA | AGACT | AGACC | AGACG | AGAGA | AGAGT | AGAGC | AGAGG | | AGT | AGTAA | AGTAT | AGTAC | AGTAG | AGTTA | AGTTT | AGTTC | AGTTG | AGTCA | AGTCT | AGTCC | AGTCG | AGTGA | AGTGT | AGTGC | AGTGG | | AGC | AGCAA | AGCAT | AGCAC | AGCAG | AGCTA | AGCTT | AGCTC | AGCTG | AGCCA | AGCCT | AGCCC | AGCCG | AGCGA | AGCGT | AGCGC | AGCGG | | AGG | AGGAA | AGGAT | AGGAC | AGGAG | AGGTA | AGGTT | AGGTC | AGGTG | AGGCA | AGGCT | AGGCC | AGGCG | AGGGA | AGGGT | AGGGC | AGGGG | | TAA | TAAAA | TAAAT | TAAAC | TAAAG | TAATA | TAATT | TAATC | TAATG | TAACA | TAACT | TAACC | TAACG | TAAGA | TAAGT | TAAGC | TAAGG | | TAT | TATAA | TATAT | TATAC | TATAG | TATTA | TATTT | TATTC | TATTG | TATCA | TATCT | TATCC | TATCG | TATGA | TATGT | TATGC | TATGG | | TAC | TACAA | TACAT | TACAC | TACAG | TACTA | TACTT | TACTC | TACTG | TACCA | TACCT | TACCC | TACCG | TACGA | TACGT | TACGC | TACGG | | TAG | TAGAA | TAGAT | TAGAC | TAGAG | TAGTA | TAGTT | TAGTC | TAGTG | TAGCA | TAGCT | TAGCC | TAGCG | TAGGA | TAGGT | TAGGC | TAGGG | | TTA | TTAAA | TTAAT | TTAAC | TTAAG | TTATA | TTATT | TTATC | TTATG | TTACA | TTACT | TTACC | TTACG | TTAGA | TTAGT | TTAGC | TTAGG | | TTT | TTTAA | TTTAT | TTTAC | TTTAG | TTTTA | TTTTT | TTTTC | TTTTG | TTTCA | TTTCT | TTTCC | TTTCG | TTTGA | TTTGT | TTTGC | TTTGG | | TTC | TTCAA | TTCAT | TTCAC | TTCAG | TTCTA | TTCTT | TTCTC | TTCTG | TTCCA | TTCCT | TTCCC | TTCCG | TTCGA | TTCGT | TTCGC | TTCGG | | TTG | TTGAA | TTGAT | TTGAC | TTGAG | TTGTA | TTGTT | TTGTC | TTGTG | TTGCA | TTGCT | TTGCC | TTGCG | TTGGA | TTGGT | TTGGC | TTGGG | | TCA | TCAAA | TCAAT | TCAAC | TCAAG | TCATA | TCATT | TCATC | TCATG | TCACA | TCACT | TCACC | TCACG | TCAGA | TCAGT | TCAGC | TCAGG | | TCT | TCTAA | TCTAT | TCTAC | TCTAG | TCTTA | TCTTT | TCTTC | TCTTG | TCTCA | TCTCT | TCTCC | TCTCG | TCTGA | TCTGT | TCTGC | TCTGG | | TCC | TCCAA | TCCAT | TCCAC | TCCAG | TCCTA | TCCTT | TCCTC | TCCTG | TCCCA | TCCCT | TCCCC | TCCCG | TCCGA | TCCGT | TCCGC | TCCGG | | TCG | TCGAA | TCGAT | TCGAC | TCGAG | TCGTA | TCGTT | TCGTC | TCGTG | TCGCA | TCGCT | TCGCC | TCGCG | TCGGA | TCGGT | TCGGC | TCGGG | | TGA | TGAAA | TGAAT | TGAAC | TGAAG | TGATA | TGATT | TGATC | TGATG | TGACA | TGACT | TGACC | TGACG | TGAGA | TGAGT | TGAGC | TGAGG | | TGT | TGTAA | TGTAT | TGTAC | TGTAG | TGTTA | TGTTT | TGTTC | TGTTG | TGTCA | TGTCT | TGTCC | TGTCG | TGTGA | TGTGT | TGTGC | TGTGG | | TGC | TGCAA | TGCAT | TGCAC | TGCAG | TGCTA | TGCTT | TGCTC | TGCTG | TGCCA | TGCCT | TGCCC | TGCCG | TGCGA | TGCGT | TGCGC | TGCGG | | TGG | TGGAA | TGGAT | TGGAC | TGGAG | TGGTA | TGGTT | TGGTC | TGGTG | TGGCA | TGGCT | TGGCC | TGGCG | TGGGA | TGGGT | TGGGC | TGGGG | | CAA | CAAAA | CAAAT | CAAAC | CAAAG | CAATA | CAATT | CAATC | CAATG | CAACA | CAACT | CAACC | CAACG | CAAGA | CAAGT | CAAGC | CAAGG | | CAT | CATAA | CATAT | CATAC | CATAG | CATTA | CATTT | CATTC | CATTG | CATCA | CATCT | CATCC | CATCG | CATGA | CATGT | CATGC | CATGG | | CAC | CACAA | CACAT | CACAC | CACAG | CACTA | CACTT | CACTC | CACTG | CACCA | CACCT | CACCC | CACCG | CACGA | CACGT | CACGC | CACGG | | CAG | CAGAA | CAGAT | CAGAC | CAGAG | CAGTA | CAGTT | CAGTC | CAGTG | CAGCA | CAGCT | CAGCC | CAGCG | CAGGA | CAGGT | CAGGC | CAGGG | | CTA | CTAAA | CTAAT | CTAAC | CTAAG | CTATA | CTATT | CTATC | CTATG | CTACA | CTACT | CTACC | CTACG | CTAGA | CTAGT | CTAGC | CTAGG | | CTT | CTTAA | CTTAT | CTTAC | CTTAG | CTTTA | CTTTT | CTTTC | CTTTG | CTTCA | CTTCT | CTTCC | CTTCG | CTTGA | CTTGT | CTTGC | CTTGG | | CTC | CTCAA | CTCAT | CTCAC | CTCAG | CTCTA | CTCTT | CTCTC | CTCTG | CTCCA | CTCCT | CTCCC | CTCCG | CTCGA | CTCGT | CTCGC | CTCGG | | CTG | CTGAA | CTGAT | CTGAC | CTGAG | CTGTA | CTGTT | CTGTC | CTGTG | CTGCA | CTGCT | CTGCC | CTGCG | CTGGA | CTGGT | CTGGC | CTGGG | | CCA | CCAAA | CCAAT | CCAAC | CCAAG | CCATA | CCATT | CCATC | CCATG | CCACA | CCACT | CCACC | CCACG | CCAGA | CCAGT | CCAGC | CCAGG | | CCT | CCTAA | CCTAT | CCTAC | CCTAG | CCTTA | CCTTT | CCTTC | CCTTG | CCTCA | CCTCT | CCTCC | CCTCG | CCTGA | CCTGT | CCTGC | CCTGG | | CCC | CCCAA | CCCAT | CCCAC | CCCAG | CCCTA | CCCTT | CCCTC | CCCTG | CCCCA | CCCCT | CCCCC | CCCCG | CCCGA | CCCGT | CCCGC | CCCGG | | CCG | CCGAA | CCGAT | CCGAC | CCGAG | CCGTA | CCGTT | CCGTC | CCGTG | CCGCA | CCGCT | CCGCC | CCGCG | CCGGA | CCGGT | CCGGC | CCGGG | | CGA | CGAAA | CGAAT | CGAAC | CGAAG | CGATA | CGATT | CGATC | CGATG | CGACA | CGACT | CGACC | CGACG | CGAGA | CGAGT | CGAGC | CGAGG | | CGT | CGTAA | CGTAT | CGTAC | CGTAG | CGTTA | CGTTT | CGTTC | CGTTG | CGTCA | CGTCT | CGTCC | CGTCG | CGTGA | CGTGT | CGTGC | CGTGG | | CGC | CGCAA | CGCAT | CGCAC | CGCAG | CGCTA | CGCTT | CGCTC | CGCTG | CGCCA | CGCCT | CGCCC | CGCCG | CGCGA | CGCGT | CGCGC | CGCGG | | CGG | CGGAA | CGGAT | CGGAC | CGGAG | CGGTA | CGGTT | CGGTC | CGGTG | CGGCA | CGGCT | CGGCC | CGGCG | CGGGA | CGGGT | CGGGC | CGGGG | | GAA | GAAAA | GAAAT | GAAAC | GAAAG | GAATA | GAATT | GAATC | GAATG | GAACA | GAACT | GAACC | GAACG | GAAGA | GAAGT | GAAGC | GAAGG | | GAT | GATAA | GATAT | GATAC | GATAG | GATTA | GATTT | GATTC | GATTG | GATCA | GATCT | GATCC | GATCG | GATGA | GATGT | GATGC | GATGG | | GAC | GACAA | GACAT | GACAC | GACAG | GACTA | GACTT | GACTC | GACTG | GACCA | GACCT | GACCC | GACCG | GACGA | GACGT | GACGC | GACGG | | GAG | GAGAA | GAGAT | GAGAC | GAGAG | GAGTA | GAGTT | GAGTC | GAGTG | GAGCA | GAGCT | GAGCC | GAGCG | GAGGA | GAGGT | GAGGC | GAGGG | | GTA | GTAAA | GTAAT | GTAAC | GTAAG | GTATA | GTATT | GTATC | GTATG | GTACA | GTACT | GTACC | GTACG | GTAGA | GTAGT | GTAGC | GTAGG | | GTT | GTTAA | GTTAT | GTTAC | GTTAG | GTTTA | GTTTT | GTTTC | GTTTG | GTTCA | GTTCT | GTTCC | GTTCG | GTTGA | GTTGT | GTTGC | GTTGG | | GTC | GTCAA | GTCAT | GTCAC | GTCAG | GTCTA | GTCTT | GTCTC | GTCTG | GTCCA | GTCCT | GTCCC | GTCCG | GTCGA | GTCGT | GTCGC | GTCGG | | GTG | GTGAA | GTGAT | GTGAC | GTGAG | GTGTA | GTGTT | GTGTC | GTGTG | GTGCA | GTGCT | GTGCC | GTGCG | GTGGA | GTGGT | GTGGC | GTGGG | | GCA | GCAAA | GCAAT | GCAAC | GCAAG | GCATA | GCATT | GCATC | GCATG | GCACA | GCACT | GCACC | GCACG | GCAGA | GCAGT | GCAGC | GCAGG | | GCT | GCTAA | GCTAT | GCTAC | GCTAG | GCTTA | GCTTT | GCTTC | GCTTG | GCTCA | GCTCT | GCTCC | GCTCG | GCTGA | GCTGT | GCTGC | GCTGG | | GCC | GCCAA | GCCAT | GCCAC | GCCAG | GCCTA | GCCTT | GCCTC | GCCTG | GCCCA | GCCCT | GCCCC | GCCCG | GCCGA | GCCGT | GCCGC | GCCGG | | GCG | GCGAA | GCGAT | GCGAC | GCGAG | GCGTA | GCGTT | GCGTC | GCGTG | GCGCA | GCGCT | GCGCC | GCGCG | GCGGA | GCGGT | GCGGC | GCGGG | | GGA | GGAAA | GGAAT | GGAAC | GGAAG | GGATA | GGATT | GGATC | GGATG | GGACA | GGACT | GGACC | GGACG | GGAGA | GGAGT | GGAGC | GGAGG | | GGT | GGTAA | GGTAT | GGTAC | GGTAG | GGTTA | GGTTT | GGTTC | GGTTG | GGTCA | GGTCT | GGTCC | GGTCG | GGTGA | GGTGT | GGTGC | GGTGG | | GGC | GGCAA | GGCAT | GGCAC | GGCAG | GGCTA | GGCTT | GGCTC | GGCTG | GGCCA | GGCCT | GGCCC | GGCCG | GGCGA | GGCGT | GGCGC | GGCGG | | GGG | GGGAA | GGGAT | GGGAC | GGGAG | GGGTA | GGGTT | GGGTC | GGGTG | GGGCA | GGGCT | GGGCC | GGGCG | GGGGA | GGGGT | GGGGC | GGGGG | After filtering: read2: overrepresented sequences Sampling rate: 1 / 20 | | | | | --- | --- | --- | | overrepresented sequence | count (% of bases) | distribution: cycle 1 ~ cycle 150 | | AAAAAAAAAAAAAAAAAAAA | 5181 (0.019858%) | | | AAAAGATAACGCAGGTGTCCTAAGATGAGCTCAACGAGAACAGAAATCTCGTGTGGAACAAAAGGGTAAAAGCTCGTTTGATTCTGATTTTCAGTACGAA | 13 (0.000249%) | | | AAAAGTGTCGGAGTTTTTTCAGCAGTTCTCGGACAAAAATTGCTGAGTGGCCGAGAAGAATGGGCGTGTCATGCGTGGGCTGACATGGATTCTTCGAGGC | 3 (0.000057%) | | | AAAATCATCAATCGTTCCAACTAATCTACCGAAGTACTCGGCTAAGAAGAAAGAAGACGGACGAATCCGAGCCAAAGCCGTACAAGCGCGAGATACCTTC | 11 (0.000211%) | | | AAACACAGGCCCCGGAACTCATCATCGAGCGTAACATCGCCCGTGAATTAACTGAGAAGGATAGGTGGTAGGTAGTTCGATGCGCGAGCATGGAGCCTAC | 20 (0.000383%) | | | AAACACTTGGTGATATGAACACAAACGTTCAATATGACAAACCCATGCCAAGTAAAGAGAAAATGAAAACTGGTGATTGTTGCGGAAATCGTCCAGGATT | 312 (0.005979%) | | | AAACGTTCAATATGACAAACCCATGCCAAGTAAAGAGAAAATGAAAACTGGTGATTGTTGCGGAAATCGTCCAGGATTCCTCGACCAGGACTTGAAATCG | 7 (0.000134%) | | | AAAGCCAAAGACTCATATGGACTTTGGCTACACCATGAAAGCTTTGAGAAGCAAGAAGAAGGTTGGTTAGTGTTTTGGAGTCGAATATGACTTGATGTCA | 283 (0.005424%) | | | AAAGGCGTAAGAATTGTATCCTTGTTAGAAGACACAAAGC | 197 (0.001510%) | | | AAAGGGTGTTGGTCGATTAAGACAGCAGGACGGTGGTCATGGAAGTCGAAATCCGCTAAGGAGTGTGTAACAACTCACCTGCCGAATCAACTAGCCCCGA | 10 (0.000192%) | | | AAAGTGTCGGAGTTTTTTCAGCAGTTCTCGGACAAAAATTGCTGAGTGGCCGAGAAGAATGGGCGTGTCATGCGTGGGCTGACATGGATTCTTCGAGGCC | 3 (0.000057%) | | | AAAGTTGGGAATTCGTTAAGGAGCTGTTGCTTTGTTAGTGTAGAAACACTTGTGTAGAATTGGGGATTGTTTTTTTTGGAGTGATTTAGGGGAGGGTCGA | 10 (0.000192%) | | | AAATACGGGCGAGAGACCGATAGCGAACAAGTACCGCGAGGTAAAGATGAAAAGGACTTTGAAAAGAGAGTCAAAGAGTGCTTGAAATTGTCGGGAGGGA | 5 (0.000096%) | | | AAATCATCAATCGTTCCAACTAATCTACCGAAGTACTCGGCTAAGAAGAAAGAAGACGGACGAATCCGAGCCAAAGCCGTACAAGCGCGAGATACCTTCG | 6 (0.000115%) | | | AAATCCTATGATGTTATCCCATGCTAATGTATCCAGAGCGTAGGCTTGCTTTGAGCACTCTAATTTCTTCAAAGTAACAGCGCCGGAGGCACGACCCGGC | 21 (0.000402%) | | | AAATCGTCGTCCCTCACCATCCTTTGCTGATGCGGGACGGAAGCTGGTCTCCCGTGTGTTACCGCACGCGGTTGGCCTAAATCCGAGCCAAGGACGCCTG | 40 (0.000767%) | | | AAATTAGGTACGAAACACAGGCCCCGGAACTCATCATCGAGCGTAACATCGCCCGTGAATTAACTGAGAAGGATAGGTGGTAGGTAGTTCGATGCGCGAG | 9 (0.000172%) | | | AAATTTCTGCCCTATCAACTTTCGATGGTAGGATAGTGGCCTACCATGGTGGTAACGGGTGACGGAGAATTAGGGTTCGATTCCGGAGAGGGAGCCTGAG | 53 (0.001016%) | | | AACACAGGCCCCGGAACTCATCATCGAGCGTAACATCGCCCGTGAATTAACTGAGAAGGATAGGTGGTAGGTAGTTCGATGCGCGAGCATGGAGCCTACG | 14 (0.000268%) | | | AACACTTGGTGATATGAACACAAACGTTCAATATGACAAACCCATGCCAAGTAAAGAGAAAATGAAAACTGGTGATTGTTGCGGAAATCGTCCAGGATTC | 3 (0.000057%) | | | AACAGGGACAGTCGGGGGCATTCGTATTTCATAGTCAGAGGTGAAATTCTTGGATTTATGAAAGACGAACAACTGCGAAAGCATTTGCCAAGGATGTTTT | 24 (0.000460%) | | | AACGAAGCACGCCCATCCAACCTAGGCGAGACAAGGGTTCACATTTCGTTCATCACCCTTGGCCGGCTATCGAACAGCCGGACTCCCATCAAAAGATGGT | 18 (0.000345%) | | | AACGGCGTGCCTCGGCATCAGCGTGCTCCGGGCGTCGGCCTGTGGGCTCCCCATTCGACCCGTCTTGAAACACGGACCAAGGAGTCTGACATGTGTGCGA | 5 (0.000096%) | | | AACGGGCAGAGCCCGCGTCGACCTTTTATCTAATAAATGCGTCCCTTCCATAAGTCGGGGTTTGTTGCACGTATTAGCTCTAGAATTACTACGGTTATCC | 11 (0.000211%) | | | AACTAGCTACGTGGAGGCATCCCTTCACGGCCGGCTTCTTAGAGGGACTATGGCCGTTTAGGCCAAGGAAGTTTGAGGCAATAACAGGTCTGTGATGCCC | 6 (0.000115%) | | | AACTTTCGATGGTAGGATAGTGGCCTACCATGGTGGTAACGGGTGACGGAGAATTAGGGTTCGATTCCGGAGAGGGAGCCTGAGAAACGGCTACCACATC | 5 (0.000096%) | | | AAGAATTGTATCCTTGTTAGAAGACACAAAGCCAAAGACTCATATGGACTTTGGCTACACCATGAAAGCTTTGAGAAGCAAGAAGAAGGTTGGTTAGTGT | 19 (0.000364%) | | | AAGACACAAAGCCAAAGACTCATATGGACTTTGGCTACACCATGAAAGCTTTGAGAAGCAAGAAGAAGGTTGGTTAGTGTTTTGGAGTCGAATATGACTT | 487 (0.009333%) | | | AAGACAGCAGGACGGTGGTCATGGAAGTCGAAATCCGCTAAGGAGTGTGTAACAACTCACCTGCCGAATCAACTAGCCCCGAAAATGGATGGCGCTTAAG | 23 (0.000441%) | | | AAGATAACGCAGGTGTCCTAAGATGAGCTCAACGAGAACAGAAATCTCGTGTGGAACAAAAGGGTAAAAGCTCGTTTGATTCTGATTTTCAGTACGAATA | 5 (0.000096%) | | | AAGATCGATC | 201342 (0.385864%) | | | AAGCACGCCCATCCAACCTAGGCGAGACAAGGGTTCACATTTCGTTCATCACCCTTGGCCGGCTATCGAACAGCCGGACTCCCATCAAAAGATGGTTGCC | 6 (0.000115%) | | | AAGCCAAAGACTCATATGGACTTTGGCTACACCATGAAAGCTTTGAGAAGCAAGAAGAAGGTTGGTTAGTGTTTTGGAGTCGAATATGACTTGATGTCAT | 10 (0.000192%) | | | AAGCCTAAGTAGTGTTTCCTTGTTAGAACATACAAAGCCAAAGACTCATACGGACTTTGGCTACACCATGAAAGCTTTGAGAAGCTAGAAGAAGGTTGGT | 1075 (0.020602%) | | | AAGCCTAAGTAGTGTTTCCTTGTTAGAAGACACAAAGCCAAAGACTCATATGGACTTTGGCTACACCATGAAAGCTTTGAGAAGCAAGAAGAAGGTTGGT | 1653 (0.031679%) | | | AAGGCGTAAGAATTGTATCCTTGTTAGAAGACACAAAGCC | 67 (0.000514%) | | | AAGGGTTCACATTTCGTTCATCACCCTTGGCCGGCTTTCGAACAGCCGGACTCCCATCAAAAGATGGTTGCCAAGAACATCTTCGTTACGGTTTGCTAAT | 4 (0.000077%) | | | AAGGTATCTCGCGCTTGTACGGCTTTGGCTCGGATTCGTCCGTCTTCTTTCTTCTTAGCCGAGTACTTCGGTAGATTAGTTGGAACGATTGATGATTTTG | 9 (0.000172%) | | | AAGTAGTGTTTCCTTGTTAGAAGACACAAAGCCAAAGACTCATATGGACTTTGGCTACACCATGAAAGCTTTGAGAAGCAAGAAGAAGGTTGGTTAGTGT | 23 (0.000441%) | | | AAGTCATATTCGACTCCAAAACACTAACCAACCTTCTTCTTGCTTCTCAAAGCTTTCATGGTGTAGCCAAAGTCCATATGAGTCTTTGGCTTTGTGTCTT | 122 (0.002338%) | | | AAGTCTGGTGCCAGCAGCCGCGGTAATTCCAGCTCCAATAGCGTATATTTAAGTTGTTGCAGTTAAAAAGCTCGTAGTTGAACCTTGGGATGGGTCGGCC | 4 (0.000077%) | | | AAGTTCTTATACTCAATCATACACATGACATCAAGTCATATTCGACTCCAAAACACTAACCAACCTTCTTCTTGCTTCTCAAAGCTTTCATGGTGTAGCC | 32 (0.000613%) | | | AAGTTGTTATACTCAATCATACACATGACAACAAGTCATATTCGACTCCAAAACACTAACCAACCTTCTTCTTGCTTCTCAAAGCTTTCATGGTGTAGCC | 25 (0.000479%) | | | AAGTTGTTATACTCAATCATACACATGACATCAAGTCATA | 126 (0.000966%) | | | AATACGGGCGAGAGACCGATAGCGAACAAGTACCGCGAGGTAAAGATGAAAAGGACTTTGAAAAGAGAGTCAAAGAGTGCTTGAAATTGTCGGGAGGGAA | 8 (0.000153%) | | | AATAGCGTATATTTAAGTTGTTGCAGTTAAAAAGCTCGTAGTTGAACCTTGGGATGGGTCGGCCGGTCCGCCTTTGGTGTGCATTGGTCGGCTTGTCCCT | 6 (0.000115%) | | | AATCAGCTTCCTTGCGCCTTACGGGTTTACTCACCCGTTGACTCGCACACATGTCAGACTCCTTGGTCCGTGTTTCAAGACGGGTCGAATGGGGAGCCCA | 174 (0.003335%) | | | AATCATACACATGACATCAAGTCATATTCGACTCCAAAACACTAACCAACCTTCTTCTTGCTTCTCAAAGCTTTCATGGTGTAGCCAAAGTCCATATGAG | 56 (0.001073%) | | | AATCATCAATCGTTCCAACTAATCTACCGAAGTACTCGGCTAAGAAGAAAGAAGACGGACGAATCCGAGCCAAAGCCGTACAAGCGCGAGATACCTTCGG | 5 (0.000096%) | | | AATCATCAGAGCAACGGGCAGAGCCCGCGTCGACCTTTTATCTAATAAATGCGTCCCTTCCATAAGTCGGGGTTTGTTGCACGTATTAGCTCTAGAATTA | 48 (0.000920%) | | | AATCCTATGATGTTATCCCATGCTAATGTATCCAGAGCGTAGGCTTGCTTTGAGCACTCTAATTTCTTCAAAGTAACAGCGCCGGAGGCACGACCCGGCC | 41 (0.000786%) | | | AATCGGTAGGAGCGACGGGCGGTGTGTACAAAGGGCAGGGACGTAGTCAACGCGAGCTGATGACTCGCGCTTACTAGGAATTCCTCGTTGAAGACCAACA | 156 (0.002990%) | | | AATCGTCGTCCCTCACCATCCTTTGCTGATGCGGGACGGAAGCTGGTCTCCCGTGTGTTACCGCACGCGGTTGGCCTAAATCCGAGCCAAGGACGCCTGG | 14 (0.000268%) | | | AATCGTTCCAACTAATCTACCGAAGTACTCGGCTAAGAAGAAAGAAGACGGACGAATCCGAGCCAAAGCCGTACAAGCGCGAGATACCTTCGGGACAGCC | 17 (0.000326%) | | | AATGATTAACAGGGACAGTCGGGGGCATTCGTATTTCATAGTCAGAGGTGAAATTCTTGGATTTATGAAAGACGAACAACTGCGAAAGCATTTGCCAAGG | 515 (0.009870%) | | | AATGCTTTGTTTTAATTAAACAGTCGGATTCCCCTTGTCCGTACCAGTTCTGAGCTGACTGTTCGACGCCCGGGGAAAGCTCCCGAGAGAGCCGTTCCCA | 701 (0.013434%) | | | AATTCCAGCTCCAATAGCGTATATTTAAGTTGTTGCAGTTAAAAAGCTCGTAGTTGAACCTTGGGATGGGTCGGCCGGTCCGCCTTTGGTGTGCATTGGT | 12 (0.000230%) | | | AATTGTATCCTTGTTAGAAGACACAAAGCCAAAGACTCATATGGACTTTGGCTACACCATGAAAGCTTTGAGAAGCAAGAAGAAGGTTGGTTAGTGTTTT | 7 (0.000134%) | | | AATTGTTGGTCTTCAACGAGGAATTCCTAGTAAGCGCGAGTCATCAGCTCGCGTTGACTACGTCCCTGCCCTTTGTACACACCGCCCGTCGCTCCTACCG | 1428 (0.027367%) | | | AATTTCTGCCCTATCAACTTTCGATGGTAGGATAGTGGCCTACCATGGTGGTAACGGGTGACGGAGAATTAGGGTTCGATTCCGGAGAGGGAGCCTGAGA | 8 (0.000153%) | | | ACAAAGCCAAAGACTCATATGGACTTTGGCTACACCATGAAAGCTTTGAGAAGCAAGAAGAAGGTTGGTTAGTGTTTTGGAGTCGAATATGACTTGATGT | 161 (0.003085%) | | | ACAAAGGGTGTTGGTCGATTAAGACAGCAGGACGGTGGTCATGGAAGTCGAAATCCGCTAAGGAGTGTGTAACAACTCACCTGCCGAATCAACTAGCCCC | 13 (0.000249%) | | | ACAAATCGTCGTCCCTCACCATCCTTTGCTGATGCGGGACGGAAGCTGGTCTCCCGTGTGTTACCGCACGCGGTTGGCCTAAATCCGAGCCAAGGACGCC | 10 (0.000192%) | | | ACAAGGGTTCACATTTCGTTCATCACCCTTGGCCGGCTATCGAACAGCCGGACTCCCATCAAAAGATGGTTGCCAAGAACATCTTCGTTACGGTTTGCTA | 4 (0.000077%) | | | ACACAAAGCCAAAGACTCATATGGACTTTGGCTACACCATGAAAGCTTTGAGAAGCAAGAAGAAGGTTGGTTAGTGTTTTGGAGTCGAATATGACTTGAT | 153 (0.002932%) | | | ACACAGGCCCCGGAACTCATCATCGAGCGTAACATCGCCCGTGAATTAACTGAGAAGGATAGGTGGTAGGTAGTTCGATGCGCGAGCATGGAGCCTACGA | 8 (0.000153%) | | | ACACATGACATCAAGTCATATTCGACTCCAAAACACTAACCAACCTTCTTCTTGCTTCTCAAAGCTTTCATGGTGTAGCCAAAGTCCATATGAGTCTTTG | 22 (0.000422%) | | | ACAGGGACAGTCGGGGGCATTCGTATTTCATAGTCAGAGGTGAAATTCTTGGATTTATGAAAGACGAACAACTGCGAAAGCATTTGCCAAGGATGTTTTC | 3 (0.000057%) | | | ACATCAAGTCATATTCGACTCCAAAACACTAACCAACCTTCTTCTTGCTTCTCAAAGCTTTCATGGTGTAGCCAAAGTCCATATGAGTCTTTGGCTTTGT | 40 (0.000767%) | | | ACATGACATCAAGTCATATTCGACTCCAAAACACTAACCAACCTTCTTCTTGCTTCTCAAAGCTTTCATGGTGTAGCCAAAGTCCATATGAGTCTTTGGC | 57 (0.001092%) | | | ACATTGTCAGGTGGGGAGTTTGGCTGGGGCGGCACATCTGTTAAAAGATAACGCAGGTGTCCTAAGATGAGCTCAACGAGAACAGAAATCTCGTGTGGAA | 13 (0.000249%) | | | ACATTGTTCCATCGACCAGAGGCTGTTCACCTTGGAGACCTGATGCGGTTATGAGTACGACCGGGCGTGAGCGGCACTCGGTCCTCCGGATTTTCAAGGG | 3 (0.000057%) | | | ACATTTCGTTCATCACCCTTGGCCGGCTTTCGAACAGCCGGACTCCCATCAAAAGATGGTTGCCAAGAACATCTTCGTTACGGTTTGCTAATTCTCGGAA | 3 (0.000057%) | | | ACATTTTATCGGTCGCTCTTGTCCGGAAGCTGTAGATGACCCAAAGTCCATATAGCGACCCCAGGTCAGGCGGGATTACCCGCTGAGTTTAAGCATATCA | 3 (0.000057%) | | | ACCAGGGGTTGAAATCGTCGACCAGGTCCGAGACTTCATCGACCGGGTCCGAGGATTCGTCGACCAGGACGGCCGGATGTCCGAGAAAAAAAAATGTTGC | 6 (0.000115%) | | | ACCATCCTTTGCTGATGCGGGACGGAAGCTGGTCTCCCGTGTGTTACCGCACGCGGTTGGCCTAAATCCGAGCCAAGGACGCCTGGAGCGTACCGACATG | 9 (0.000172%) | | | ACCGGACACCACGCGACGTGCGGTGCTCTTCCAGCCGCTGGACCCTACCTCCGGCTGAGCCGTTTCCAGGGTGGGCAGGCTGTTAAACAGAAAAGATAAC | 5 (0.000096%) | | | ACCGGGACGTGGCGGTTGACGGCAACGTTAGGGAGTCCGGAGACGTCGGCGGGGGCCTCGGGAAGAGTTATCTTTTCTGTTTAACAGCCTGCCCACCCTG | 5 (0.000096%) | | | ACCTAGGCGAGACAAGGGTTCACATTTCGTTCATCACCCTTGGCCGGCTATCGAACAGCCGGACTCCCATCAAAAGATGGTTGCCAAGAACATCTTCGTT | 4 (0.000077%) | | | ACCTCAGCCTGCTAACTAGCTACGTGGAGGCATCCCTTCACGGCCGGCTTCTTAGAGGGACTATGGCCGTTTAGGCCAAGGAAGTTTGAGGCAATAACAG | 10 (0.000192%) | | | ACGAAACACAGGCCCCGGAACTCATCATCGAGCGTAACATCGCCCGTGAATTAACTGAGAAGGATAGGTGGTAGGTAGTTCGATGCGCGAGCATGGAGCC | 33 (0.000632%) | | | ACGAAGCACGCCCATCCAACCTAGGCGAGACAAGGGTTCACATTTCGTTCATCACCCTTGGCCGGCTATCGAACAGCCGGACTCCCATCAAAAGATGGTT | 4 (0.000077%) | | | ACGCATCATTCAAATTTCTGCCCTATCAACTTTCGATGGTAGGATAGTGGCCTACCATGGTGGTAACGGGTGACGGAGAATTAGGGTTCGATTCCGGAGA | 3 (0.000057%) | | | ACGCCCATCCAACCTAGGCGAGACAAGGGTTCACATTTCGTTCATCACCCTTGGCCGGCTTTCGAACAGCCGGACTCCCATCAAAAGATGGTTGCCAAGA | 2059 (0.039460%) | | | ACGCGCCTAACGGCGTGCCTCGGCATCAGCGTGCTCCGGGCGTCGGCCTGTGGGCTCCCCATTCGACCCGTCTTGAAACACGGACCAAGGAGTCTGACAT | 14 (0.000268%) | | | ACGCTTTCACGGTTCGTATTCGTACTGAAAATCAGAATCAAACGAGCTTTTACCCTTTTGTTCCACACGAGATTTCTGTTCTCGTTGAGCTCATCTTAGG | 11 (0.000211%) | | | ACGGCTTTGGCTCGGATTCGTCCGTCTTCTTTCTTCTTAGCCGAGTACTTCGGTAGATTAGTTGGAACGATTGATGATTTTGAGTTAATTGAACGTTCGG | 7 (0.000134%) | | | ACGGGCAGAGCCCGCGTCGACCTTTTATCTAATAAATGCGTCCCTTCCATAAGTCGGGGTTTGTTGCACGTATTAGCTCTAGAATTACTACGGTTATCCG | 3 (0.000057%) | | | ACGGGTTTACTCACCCGTTGACTCGCACACATGTCAGACTCCTTGGTCCGTGTTTCAAGACGGGTCGAATGGGGAGCCCACAGGCCGACGCCCGGAGCAC | 5 (0.000096%) | | | ACGGTGGTCATGGAAGTCGAAATCCGCTAAGGAGTGTGTAACAACTCACCTGCCGAATCAACTAGCCCCGAAAATGGATGGCGCTTAAGCGCGCGACCTA | 10 (0.000192%) | | | ACGTGGGTGGTTCGCCGCCCGCGACGTCGCGAGAAGTCCACTAAACCTTATCATTTAGAGGAAGGAGAAGTCGTAACAAGGTTTCCGTAGGTGAACCTGC | 5 (0.000096%) | | | ACTACCTACCACCTATCCTTCTCAGTTAATTCACGGGCGATGTTACGCTCGATGATGAGTTCCGGGGCCTGTGTTTCGTACCTAATTTGAAGGAATTGTT | 4 (0.000077%) | | | ACTAGCTACGTGGAGGCATCCCTTCACGGCCGGCTTCTTAGAGGGACTATGGCCGTTTAGGCCAAGGAAGTTTGAGGCAATAACAGGTCTGTGATGCCCT | 4 (0.000077%) | | | ACTCAATCATACACATGACATCAAGTCATATTCGACTCCAAAACACTAACCAACCTTCTTCTTGCTTCTCAAAGCTTTCATGGTGTAGCCAAAGTCCATA | 31 (0.000594%) | | | ACTCCAAAACACTAACCAACCTTCTTCTTGCTTCTCAAAGCTTTCATGGTGTAGCCAAAGTCCATATGAGTCTTTGGCTTTGTGTCTTCTAACAAGGAAA | 388 (0.007436%) | | | ACTCCAAAACACTAACCAACCTTCTTCTTGCTTCTCAAAGCTTTCATGGTGTAGCCAAAGTCCATATGAGTCTTTGGCTTTGTGTCTTCTAACAAGGATA | 3 (0.000057%) | | | ACTCGGTCCTCCGGATTTTCAAGGGCCGCCGGGGGCGCACCGGACACCACGCGACGTGCGGTGCTCTTCCAGCCGCTGGACCCTACCTCCGGCTGAGCCG | 11 (0.000211%) | | | ACTTCCCTTGCCTACATTGTTCCATCGACCAGAGGCTGTTCACCTTGGAGACCTGATGCGGTTATGAGTACGACCGGGCGTGAGCGGCACTCGGTCCTCC | 18 (0.000345%) | | | AGAAGACACAAAGCCAAAGACTCATATGGACTTTGGCTACACCATGAAAGCTTTGAGAAGCAAGAAGAAGGTTGGTTAGTGTTTTGGAGTCGAATATGAC | 64 (0.001227%) | | | AGAATTGTATCCTTGTTAGAAGACACAAAGCCAAAGACTCATATGGACTTTGGCTACACCATGAAAGCTTTGAGAAGCAAGAAGAAGGTTGGTTAGTGTT | 3 (0.000057%) | | | AGACAAGGGTTCACATTTCGTTCATCACCCTTGGCCGGCTATCGAACAGCCGGACTCCCATCAAAAGATGGTTGCCAAGAACATCTTCGTTACGGTTTGC | 6 (0.000115%) | | | AGACACAAAGCCAAAGACTCATATGGACTTTGGCTACACCATGAAAGCTTTGAGAAGCAAGAAGAAGGTTGGTTAGTGTTTTGGAGTCGAATATGACTTG | 33 (0.000632%) | | | AGACAGCAGGACGGTGGTCATGGAAGTCGAAATCCGCTAAGGAGTGTGTAACAACTCACCTGCCGAATCAACTAGCCCCGAAAATGGATGGCGCTTAAGC | 7 (0.000134%) | | | AGAGCAACGGGCAGAGCCCGCGTCGACCTTTTATCTAATAAATGCGTCCCTTCCATAAGTCGGGGTTTGTTGCACGTATTAGCTCTAGAATTACTACGGT | 31 (0.000594%) | | | AGAGCCCGCGTCGACCTTTTATCTAATAAATGCGTCCCTTCCATAAGTCGGGGTTTGTTGCACGTATTAGCTCTAGAATTACTACGGTTATCCGAGTAGT | 7 (0.000134%) | | | AGAGCGTAGGCTTGCTTTGAGCACTCTAATTTCTTCAAAGTAACAGCGCCGGAGGCACGACCCGGCCAATTAAGACCAGGAGCGTATCGCCGACCGAAGG | 15 (0.000287%) | | | AGAGGCTGTTCACCTTGGAGACCTGATGCGGTTATGAGTACGACCGGGCGTGAGCGGCACTCGGTCCTCCGGATTTTCAAGGGCCGCCGGGGGCGCACCG | 150 (0.002875%) | | | AGATCGGAAGAGCGTCGTGTAGGGAAAGAGTGTTCAGAGC | 47 (0.000360%) | | | AGCAACGGGCAGAGCCCGCGTCGACCTTTTATCTAATAAATGCGTCCCTTCCATAAGTCGGGGTTTGTTGCACGTATTAGCTCTAGAATTACTACGGTTA | 7 (0.000134%) | | | AGCACGCCCATCCAACCTAGGCGAGACAAGGGTTCACATTTCGTTCATCACCCTTGGCCGGCTATCGAACAGCCGGACTCCCATCAAAAGATGGTTGCCA | 12 (0.000230%) | | | AGCACGCGCCTAACGGCGTGCCTCGGCATCAGCGTGCTCCGGGCGTCGGCCTGTGGGCTCCCCATTCGACCCGTCTTGAAACACGGACCAAGGAGTCTGA | 31 (0.000594%) | | | AGCAGCCGCGGTAATTCCAGCTCCAATAGCGTATATTTAAGTTGTTGCAGTTAAAAAGCTCGTAGTTGAACCTTGGGATGGGTCGGCCGGTCCGCCTTTG | 6 (0.000115%) | | | AGCAGGACGGTGGTCATGGAAGTCGAAATCCGCTAAGGAGTGTGTAACAACTCACCTGCCGAATCAACTAGCCCCGAAAATGGATGGCGCTTAAGCGCGC | 6 (0.000115%) | | | AGCAGTTCTCGGACAAAAATTGCTGAGTGGCCGAGAAGAATGGGCGTGTCATGCGTGGGCTGACATGGATTCTTCGAGGCCTAGGGGTGGCGGTATATAA | 6 (0.000115%) | | | AGCCAAAGACTCATATGGACTTTGGCTACACCATGAAAGCTTTGAGAAGCAAGAAGAAGGTTGGTTAGTGTTTTGGAGTCGAATATGACTTGATGTCATG | 8 (0.000153%) | | | AGCCGCGGTAATTCCAGCTCCAATAGCGTATATTTAAGTTGTTGCAGTTAAAAAGCTCGTAGTTGAACCTTGGGATGGGTCGGCCGGTCCGCCTTTGGTG | 8 (0.000153%) | | | AGCCTAAGTAGTGTTTCCTTGTTAGAAGACACAAAGCCAAAGACTCATATGGACTTTGGCTACACCATGAAAGCTTTGAGAAGCAAGAAGAAGGTTGGTT | 33 (0.000632%) | | | AGCCTGCTAACTAGCTACGTGGAGGCATCCCTTCACGGCCGGCTTCTTAGAGGGACTATGGCCGTTTAGGCCAAGGAAGTTTGAGGCAATAACAGGTCTG | 6 (0.000115%) | | | AGCGAGGTGTGAGTGTCGCCCATGGGCATCGACACCTTGCGGCTAGGAACTGGAACGAGACGGGTGGCAAAGATTTCGAGTAGCACTTCATACTACCGTG | 7 (0.000134%) | | | AGCGGCACTCGGTCCTCCGGATTTTCAAGGGCCGCCGGGGGCGCACCGGACACCACGCGACGTGCGGTGCTCTTCCAGCCGCTGGACCCTACCTCCGGCT | 3 (0.000057%) | | | AGCGTAGGCTTGCTTTGAGCACTCTAATTTCTTCAAAGTAACAGCGCCGGAGGCACGACCCGGCCAATTAAGACCAGGAGCGTATCGCCGACCGAAGGGA | 3 (0.000057%) | | | AGCGTGCTCCGGGCGTCGGCCTGTGGGCTCCCCATTCGACCCGTCTTGAAACACGGACCAAGGAGTCTGACATGTGTGCGAGTCAACGGGTGAGTAAACC | 4 (0.000077%) | | | AGCTAGTGTTCGTAGGCTCCATGCTCGCGCATCGAACTACCTACCACCTATCCTTCTCAGTTAATTCACGGGCGATGTTACGCTCGATGATGAGTTCCGG | 12 (0.000230%) | | | AGCTCCAATAGCGTATATTTAAGTTGTTGCAGTTAAAAAGCTCGTAGTTGAACCTTGGGATGGGTCGGCCGGTCCGCCTTTGGTGTGCATTGGTCGGCTT | 3 (0.000057%) | | | AGCTTCCTTGCGCCTTACGGGTTTACTCACCCGTTGACTCGCACACATGTCAGACTCCTTGGTCCGTGTTTCAAGACGGGTCGAATGGGGAGCCCACAGG | 24 (0.000460%) | | | AGGACATTGTCAGGTGGGGAGTTTGGCTGGGGCGGCACATCTGTTAAAAGATAACGCAGGTGTCCTAAGATGAGCTCAACGAGAACAGAAATCTCGTGTG | 150 (0.002875%) | | | AGGACGGTGGTCATGGAAGTCGAAATCCGCTAAGGAGTGTGTAACAACTCACCTGCCGAATCAACTAGCCCCGAAAATGGATGGCGCTTAAGCGCGCGAC | 35 (0.000671%) | | | AGGAGCGACGGGCGGTGTGTACAAAGGGCAGGGACGTAGTCAACGCGAGCTGATGACTCGCGCTTACTAGGAATTCCTCGTTGAAGACCAACAATTGCAA | 19 (0.000364%) | | | AGGAGCTGTTGCTTTGTTAGTGTAGAAACACTTGTGTAGAATTGGGGATTGTTTTTTTTGGAGTGATTTAGGGGAGGGTCGAATCTTAGCGACAAAGGGC | 12 (0.000230%) | | | AGGCGAGACAAGGGTTCACATTTCGTTCATCACCCTTGGCCGGCTATCGAACAGCCGGACTCCCATCAAAAGATGGTTGCCAAGAACATCTTCGTTACGG | 3 (0.000057%) | | | AGGCGAGACAAGGGTTCACATTTCGTTCATCACCCTTGGCCGGCTTTCGAACAGCCGGACTCCCATCAAAAGATGGTTGCCAAGAACATCTTCGTTACGG | 4 (0.000077%) | | | AGGCGTAAGAATTGTATCCTTGTTAGAAGACACAAAGCCA | 111 (0.000851%) | | | AGGCTGTCCCGAGTGTGAGCGAGGTGTGAGTGTCGCCCATGGGCATCGACACCTTGCGGCTAGGAACTGGAACGAGACGGGTAGCAAAGATTTCGAGTAG | 1874 (0.035914%) | | | AGGCTGTCCCGAGTGTGAGCGAGGTGTGAGTGTCGCCCATGGGCATCGACACCTTGCGGCTAGGAACTGGAACGAGACGGGTGGCAAAGATTTCGAGTAG | 13 (0.000249%) | | | AGGCTGTTCACCTTGGAGACCTGATGCGGTTATGAGTACGACCGGGCGTGAGCGGCACTCGGTCCTCCGGATTTTCAAGGGCCGCCGGGGGCGCACCGGA | 33 (0.000632%) | | | AGGGACAGTCGGGGGCATTCGTATTTCATAGTCAGAGGTGAAATTCTTGGATTTATGAAAGACGAACAACTGCGAAAGCATTTGCCAAGGATGTTTTCAT | 6 (0.000115%) | | | AGGGAGTCCGGAGACGTCGGCGGGGGCCTCGGGAAGAGTTATCTTTTCTGTTTAACAGCCTGCCCACCCTGGAAACGGCTCAGCCGGAGGTAGGGTCCAG | 7 (0.000134%) | | | AGGGCAAGTCTGGTGCCAGCAGCCGCGGTAATTCCAGCTCCAATAGCGTATATTTAAGTTGTTGCAGTTAAAAAGCTCGTAGTTGAACCTTGGGATGGGT | 27 (0.000517%) | | | AGGGGTTGAAATCGTCGACCAGGTCCGAGACTTCATCGACCGGGTCCGAGGATTCGTCGACCAGGACGGCCGGATGTCCGAGAAAAAAAAATGTTGCCGA | 4 (0.000077%) | | | AGGGTGTTGGTCGATTAAGACAGCAGGACGGTGGTCATGGAAGTCGAAATCCGCTAAGGAGTGTGTAACAACTCACCTGCCGAATCAACTAGCCCCGAAA | 13 (0.000249%) | | | AGGTATCTCGCGCTTGTACGGCTTTGGCTCGGATTCGTCCGTCTTCTTTCTTCTTAGCCGAGTACTTCGGTAGATTAGTTGGAACGATTGATGATTTTGA | 6 (0.000115%) | | | AGTAGTCATATGCTTGTCTCAAAGATTAAGCCATGCATGTGTAAGTATGAACGAATTCAGACTGTGAAACTGCGAATGGCTCATTAAATCAGTTATAGTT | 2299 (0.044059%) | | | AGTAGTGTTTCCTTGTTAGAAGACACAAAGCCAAAGACTCATATGGACTTTGGCTACACCATGAAAGCTTTGAGAAGCAAGAAGAAGGTTGGTTAGTGTT | 14 (0.000268%) | | | AGTATAAGAACTTAAACCGCAACCGCATCTTATAAGCCTAAGTAGTGTTTCCTTGTTAGAAGACACAAAGCCAAAGACTCATATGGACTTTGGCTACACC | 3 (0.000057%) | | | AGTCATATGCTTGTCTCAAAGATTAAGCCATGCATGTGTAAGTATGAACGAATTCAGACTGTGAAACTGCGAATGGCTCATTAAATCAGTTATAGTTTGT | 13 (0.000249%) | | | AGTCATATTCGACTCCAAAACACTAACCAACCTTCTTCTTGCTTCTCAAAGCTTTCATGGTGTAGCCAAAGTCCATATGAGTCTTTGGCTTTGTGTCTTC | 6 (0.000115%) | | | AGTCGAATATGACTTGATGTCATGTGTATGATTGAGTATA | 1268 (0.009720%) | | | AGTCGGGGGCATTCGTATTTCATAGTCAGAGGTGAAATTCTTGGATTTATGAAAGACGAACAACTGCGAAAGCATTTGCCAAGGATGTTTTCATTAATCA | 4 (0.000077%) | | | AGTGTATCCTTGTTAGAAGACACAAAGCCAAAGACTCATA | 8146 (0.062446%) | | | AGTGTTCGTAGGCTCCATGCTCGCGCATCGAACTACCTACCACCTATCCTTCTCAGTTAATTCACGGGCGATGTTACGCTCGATGATGAGTTCCGGGGCC | 210 (0.004025%) | | | AGTGTTTCCTTGTTAGAAGACACAAAGCCAAAGACTCATATGGACTTTGGCTACACCATGAAAGCTTTGAGAAGCAAGAAGAAGGTTGGTTAGTGTTTTG | 10 (0.000192%) | | | AGTGTTTTGGAGTCGAATATGACTTGATGTCATGTGTATG | 181 (0.001388%) | | | AGTTATCATGAATCATCAGAGCAACGGGCAGAGCCCGCGTCGACCTTTTATCTAATAAATGCGTCCCTTCCATAAGTCGGGGTTTGTTGCACGTATTAGC | 12 (0.000230%) | | | AGTTCTTATACTCAATCATACACATGACATCAAGTCATATTCGACTCCAAAACACTAACCAACCTTCTTCTTGCTTCTCAAAGCTTTCATGGTGTAGCCA | 25 (0.000479%) | | | AGTTGTTATACTCAATCATACACATGACAACAAGTCATATTCGACTCCAAAACACTAACCAACCTTCTTCTTGCTTCTCAAAGCTTTCATGGTGTAGCCA | 19 (0.000364%) | | | AGTTGTTATACTCAATCATACACATGACATCAAGTCATAT | 97 (0.000744%) | | | AGTTTGGCTGGGGCGGCACATCTGTTAAAAGATAACGCAGGTGTCCTAAGATGAGCTCAACGAGAACAGAAATCTCGTGTGGAACAAAAGGGTAAAAGCT | 5 (0.000096%) | | | AGTTTTTTCAGCAGTTCTCGGACAAAAATTGCTGAGTGGCCGAGAAGAATGGGCGTGTCATGCGTGGGCTGACATGGATTCTTCGAGGCCTAGGGGTGGC | 4 (0.000077%) | | | ATAAAAGTGTCGGAGTTTTTTCAGCAGTTCTCGGACAAAAATTGCTGAGTGGCCGAGAAGAATGGGCGTGTCATGCGTGGGCTGACATGGATTCTTCGAG | 155 (0.002971%) | | | ATAAGAACTTAAACCGCAACCGCATCTTATAAGCCTAAGTAGTGTTTCCTTGTTAGAAGACACAAAGCCAAAGACTCATATGGACTTTGGCTACACCATG | 24 (0.000460%) | | | ATACACATGACATCAAGTCATATTCGACTCCAAAACACTAACCAACCTTCTTCTTGCTTCTCAAAGCTTTCATGGTGTAGCCAAAGTCCATATGAGTCTT | 286 (0.005481%) | | | ATACGGGCGAGAGACCGATAGCGAACAAGTACCGCGAGGTAAAGATGAAAAGGACTTTGAAAAGAGAGTCAAAGAGTGCTTGAAATTGTCGGGAGGGAAG | 14 (0.000268%) | | | ATACTCAATCATACACATGACATCAAGTCATATTCGACTCCAAAACACTAACCAACCTTCTTCTTGCTTCTCAAAGCTTTCATGGTGTAGCCAAAGTCCA | 29 (0.000556%) | | | ATAGCTAGTGTTCGTAGGCTCCATGCTCGCGCATCGAACTACCTACCACCTATCCTTCTCAGTTAATTCACGGGCGATGTTACGCTCGATGATGAGTTCC | 4 (0.000077%) | | | ATATGCTTGTCTCAAAGATTAAGCCATGCATGTGTAAGTATGAACGAATTCAGACTGTGAAACTGCGAATGGCTCATTAAATCAGTTATAGTTTGTTTGA | 47 (0.000901%) | | | ATATTCGACTCCAAAACACTAACCAACCTTCTTCTTGCTTCTCAAAGCTTTCATGGTGTAGCCAAAGTCCATATGAGTCTTTGGCTTTGTGTCTTCTAAC | 150 (0.002875%) | | | ATATTCGACTCCAAAACACTAACCAACCTTCTTCTTGCTTCTCAAAGCTTTCATGGTGTAGCCAAAGTCCATATGAGTCTTTGGCTTTGTGTCTTTTAAC | 61 (0.001169%) | | | ATATTTAAGTTGTTGCAGTTAAAAAGCTCGTAGTTGAACCTTGGGATGGGTCGGCCGGTCCGCCTTTGGTGTGCATTGGTCGGCTTGTCCCTTCGGTCGG | 7 (0.000134%) | | | ATCAACTTTCGATGGTAGGATAGTGGCCTACCATGGTGGTAACGGGTGACGGAGAATTAGGGTTCGATTCCGGAGAGGGAGCCTGAGAAACGGCTACCAC | 9 (0.000172%) | | | ATCAAGTCATATTCGACTCCAAAACACTAACCAACCTTCTTCTTGCTTCTCAAAGCTTTCATGGTGTAGCCAAAGTCCATATGAGTCTTTGGCTTTGTGT | 1348 (0.025834%) | | | ATCAATCGTTCCAACTAATCTACCGAAGTACTCGGCTAAGAAGAAAGAAGACGGACGAATCCGAGCCAAAGCCGTACAAGCGCGAGATACCTTCGGGACA | 5 (0.000096%) | | | ATCAGAGCAACGGGCAGAGCCCGCGTCGACCTTTTATCTAATAAATGCGTCCCTTCCATAAGTCGGGGTTTGTTGCACGTATTAGCTCTAGAATTACTAC | 36 (0.000690%) | | | ATCAGCGTGCTCCGGGCGTCGGCCTGTGGGCTCCCCATTCGACCCGTCTTGAAACACGGACCAAGGAGTCTGACATGTGTGCGAGTCAACGGGTGAGTAA | 4 (0.000077%) | | | ATCAGCTTCCTTGCGCCTTACGGGTTTACTCACCCGTTGACTCGCACACATGTCAGACTCCTTGGTCCGTGTTTCAAGACGGGTCGAATGGGGAGCCCAC | 65 (0.001246%) | | | ATCATACACATGACATCAAGTCATATTCGACTCCAAAACACTAACCAACCTTCTTCTTGCTTCTCAAAGCTTTCATGGTGTAGCCAAAGTCCATATGAGT | 24 (0.000460%) | | | ATCATCAGAGCAACGGGCAGAGCCCGCGTCGACCTTTTATCTAATAAATGCGTCCCTTCCATAAGTCGGGGTTTGTTGCACGTATTAGCTCTAGAATTAC | 12 (0.000230%) | | | ATCATGAATCATCAGAGCAACGGGCAGAGCCCGCGTCGACCTTTTATCTAATAAATGCGTCCCTTCCATAAGTCGGGGTTTGTTGCACGTATTAGCTCTA | 53 (0.001016%) | | | ATCCAACCTAGGCGAGACAAGGGTTCACATTTCGTTCATCACCCTTGGCCGGCTATCGAACAGCCGGACTCCCATCAAAAGATGGTTGCCAAGAACATCT | 25 (0.000479%) | | | ATCCAACCTAGGCGAGACAAGGGTTCACATTTCGTTCATCACCCTTGGCCGGCTTTCGAACAGCCGGACTCCCATCAAAAGATGGTTGCCAAGAACATCT | 27 (0.000517%) | | | ATCCAGAGCGTAGGCTTGCTTTGAGCACTCTAATTTCTTCAAAGTAACAGCGCCGGAGGCACGACCCGGCCAATTAAGACCAGGAGCGTATCGCCGACCG | 17 (0.000326%) | | | ATCCCATGCTAATGTATCCAGAGCGTAGGCTTGCTTTGAGCACTCTAATTTCTTCAAAGTAACAGCGCCGGAGGCACGACCCGGCCAATTAAGACCAGGA | 23 (0.000441%) | | | ATCCCGCCAATCAGCTTCCTTGCGCCTTACGGGTTTACTCACCCGTTGACTCGCACACATGTCAGACTCCTTGGTCCGTGTTTCAAGACGGGTCGAATGG | 7 (0.000134%) | | | ATCCGTCGAGTTATCATGAATCATCAGAGCAACGGGCAGAGCCCGCGTCGACCTTTTATCTAATAAATGCGTCCCTTCCATAAGTCGGGGTTTGTTGCAC | 10 (0.000192%) | | | ATCCTATGATGTTATCCCATGCTAATGTATCCAGAGCGTAGGCTTGCTTTGAGCACTCTAATTTCTTCAAAGTAACAGCGCCGGAGGCACGACCCGGCCA | 44 (0.000843%) | | | ATCCTTGTTAGAAGACACAAAGCCAAAGACTCATATGGACTTTGGCTACACCATGAAAGCTTTGAGAAGCAAGAAGAAGGTTGGTTAGTGTTTTGGAGTC | 13 (0.000249%) | | | ATCGAAATCCTATGATGTTATCCCATGCTAATGTATCCAGAGCGTAGGCTTGCTTTGAGCACTCTAATTTCTTCAAAGTAACAGCGCCGGAGGCACGACC | 15 (0.000287%) | | | ATCGACACCTTGCGGCTAGGAACTGGAACGAGACGGGTGGCAAAGATTTCGAGTAGCACTTCATACTACCGTGGGTTTTTTAAACCTTCCGAGTTTTGTT | 10 (0.000192%) | | | ATCGACCCGCCGAAGCGAGCCTTGGGACCAAAAACAGGGGTTGTACCCCGCCTCCGATTCACGGAGTAAGTAAAATAACGTTAAAAGTAGTGGTATTTCA | 6 (0.000115%) | | | ATCGATCCATTACATTTTATCGGTCGCTCTTGTCCGGAAGCTGTAGATGACCCAAAGTCCATATAGCGACCCCAGGTCAGGCGGGATTACCCGCTGAGTT | 19 (0.000364%) | | | ATCGATCCCGCCAATCAGCTTCCTTGCGCCTTACGGGTTTACTCACCCGTTGACTCGCACACATGTCAGACTCCTTGGTCCGTGTTTCAAGACGGGTCGA | 1139 (0.021828%) | | | ATCGATCCGTAACTTCGGGAAAAGGATTGGCTCTGAGGGCTGGGCTCGGGGGTCCCAGTTCCGAACCCGTCGGCTGTCAGCGGACTGCTCGAGCTGCTTC | 30 (0.000575%) | | | ATCGATCCGTCGAGTTATCATGAATCATCAGAGCAACGGGCAGAGCCCGCGTCGACCTTTTATCTAATAAATGCGTCCCTTCCATAAGTCGGGGTTTGTT | 19 (0.000364%) | | | ATCGATCGACCCGCCGAAGCGAGCCTTGGGACCAAAAACAGGGGTTGTACCCCGCCTCCGATTCACGGAGTAAGTAAAATAACGTTAAAAGTAGTGGTAT | 14 (0.000268%) | | | ATCGATCTTAAAGGCGTAAGAATTGTATCCTTGTTAAAAGACACAAAGCCAAAGACTCATATGGACTTTGGCTACACCATGAAAGCTTTGAGAAGCAAGA | 11 (0.000211%) | | | ATCGATCTTCTGAGAAGGGTTCGAGTGTGAGCATGCCTGTCGGGACCCGAAAGATGGTGAACTATGCCTGAGCGGGGTAAAGCCAGAGGAAACTCTGGTG | 20 (0.000383%) | | | ATCGGTAGGAGCGACGGGCGGTGTGTACAAAGGGCAGGGACGTAGTCAACGCGAGCTGATGACTCGCGCTTACTAGGAATTCCTCGTTGAAGACCAACAA | 65 (0.001246%) | | | ATCGTCGTCCCTCACCATCCTTTGCTGATGCGGGACGGAAGCTGGTCTCCCGTGTGTTACCGCACGCGGTTGGCCTAAATCCGAGCCAAGGACGCCTGGA | 9 (0.000172%) | | | ATCTCGCGCTTGTACGGCTTTGGCTCGGATTCGTCCGTCTTCTTTCTTCTTAGCCGAGTACTTCGGTAGATTAGTTGGAACGATTGATGATTTTGAGTTA | 3 (0.000057%) | | | ATCTTAAAGGCGTAAGAATTGTATCCTTGTTAAAAGACACAAAGCCAAAGACTCATATGGACTTTGGCTACACCATGAAAGCTTTGAGAAGCAAGAAGAA | 36 (0.000690%) | | | ATCTTAAAGGCGTAAGAATTGTATCCTTGTTAGAAGACAC | 1341 (0.010280%) | | | ATCTTCTGAGAAGGGTTCGAGTGTGAGCATGCCTGTCGGGACCCGAAAGATGGTGAACTATGCCTGAGCGGGGTAAAGCCAGAGGAAACTCTGGTGGAAG | 3 (0.000057%) | | | ATGAACACAAACGTTCAATATGACAAACCCATGCCAAGTAAAGAGAAAATGAAAACTGGTGATTGTTGCGGAAATCGTCCAGGATTCCTCGACCAGGACT | 12 (0.000230%) | | | ATGAATCATCAGAGCAACGGGCAGAGCCCGCGTCGACCTTTTATCTAATAAATGCGTCCCTTCCATAAGTCGGGGTTTGTTGCACGTATTAGCTCTAGAA | 68 (0.001303%) | | | ATGACATCAAGTCATATTCGACTCCAAAACACTAACCAACCTTCTTCTTGCTTCTCAAAGCTTTCATGGTGTAGCCAAAGTCCATATGAGTCTTTGGCTT | 116 (0.002223%) | | | ATGATGTTATCCCATGCTAATGTATCCAGAGCGTAGGCTTGCTTTGAGCACTCTAATTTCTTCAAAGTAACAGCGCCGGAGGCACGACCCGGCCAATTAA | 68 (0.001303%) | | | ATGATTAACAGGGACAGTCGGGGGCATTCGTATTTCATAGTCAGAGGTGAAATTCTTGGATTTATGAAAGACGAACAACTGCGAAAGCATTTGCCAAGGA | 98 (0.001878%) | | | ATGATTGAGTATAAGAACTTAAACCGCAACCGCATCTTATAAGCCTAAGTAGTGTTTCCTTGTTAGAAGACACAAAGCCAAAGACTCATATGGACTTTGG | 27 (0.000517%) | | | ATGCTAATGTATCCAGAGCGTAGGCTTGCTTTGAGCACTCTAATTTCTTCAAAGTAACAGCGCCGGAGGCACGACCCGGCCAATTAAGACCAGGAGCGTA | 10 (0.000192%) | | | ATGCTCGCGCATCGAACTACCTACCACCTATCCTTCTCAGTTAATTCACGGGCGATGTTACGCTCGATGATGAGTTCCGGGGCCTGTGTTTCGTACCTAA | 5 (0.000096%) | | | ATGCTTGTCTCAAAGATTAAGCCATGCATGTGTAAGTATGAACGAATTCAGACTGTGAAACTGCGAATGGCTCATTAAATCAGTTATAGTTTGTTTGATG | 20 (0.000383%) | | | ATGGAAGTCGAAATCCGCTAAGGAGTGTGTAACAACTCACCTGCCGAATCAACTAGCCCCGAAAATGGATGGCGCTTAAGCGCGCGACCTATACCCGGCC | 36 (0.000690%) | | | ATGGCCTCTGTGCTGGCGACGCATCATTCAAATTTCTGCCCTATCAACTTTCGATGGTAGGATAGTGGCCTACCATGGTGGTAACGGGTGACGGAGAATT | 392 (0.007513%) | | | ATGGGCATCGACACCTTGCGGCTAGGAACTGGAACGAGACGGGTGGCAAAGATTTCGAGTAGCACTTCATACTACCGTGGGTTTTTTAAACCTTCCGAGT | 5 (0.000096%) | | | ATGTATCCAGAGCGTAGGCTTGCTTTGAGCACTCTAATTTCTTCAAAGTAACAGCGCCGGAGGCACGACCCGGCCAATTAAGACCAGGAGCGTATCGCCG | 6 (0.000115%) | | | ATGTGTAAGTATGAACGAATTCAGACTGTGAAACTGCGAATGGCTCATTAAATCAGTTATAGTTTGTTTGATGGTAACTACTACTCGGATAACCGTAGTA | 184 (0.003526%) | | | ATGTGTATGATTGAGTATAAGAACTTAAACCGCAACCGCATCTTATAAGCCTAAGTAGTGTTTCCTTGTTAGAAGACACAAAGCCAAAGACTCATATGGA | 16 (0.000307%) | | | ATGTTATCCCATGCTAATGTATCCAGAGCGTAGGCTTGCTTTGAGCACTCTAATTTCTTCAAAGTAACAGCGCCGGAGGCACGACCCGGCCAATTAAGAC | 9 (0.000172%) | | | ATTAACAGGGACAGTCGGGGGCATTCGTATTTCATAGTCAGAGGTGAAATTCTTGGATTTATGAAAGACGAACAACTGCGAAAGCATTTGCCAAGGATGT | 85 (0.001629%) | | | ATTAAGACAGCAGGACGGTGGTCATGGAAGTCGAAATCCGCTAAGGAGTGTGTAACAACTCACCTGCCGAATCAACTAGCCCCGAAAATGGATGGCGCTT | 18 (0.000345%) | | | ATTACATTTTATCGGTCGCTCTTGTCCGGAAGCTGTAGATGACCCAAAGTCCATATAGCGACCCCAGGTCAGGCGGGATTACCCGCTGAGTTTAAGCATA | 7 (0.000134%) | | | ATTAGGTACGAAACACAGGCCCCGGAACTCATCATCGAGCGTAACATCGCCCGTGAATTAACTGAGAAGGATAGGTGGTAGGTAGTTCGATGCGCGAGCA | 16 (0.000307%) | | | ATTAGTCTTTCGCCCCTATACCCAAGTCAGACGAACGATTTGCACGTCAGTATCGCTGCGGGCTTCCACCAGAGTTTCCTCTGGCTTTACCCCGCTCAGG | 24 (0.000460%) | | | ATTCAAATTTCTGCCCTATCAACTTTCGATGGTAGGATAGTGGCCTACCATGGTGGTAACGGGTGACGGAGAATTAGGGTTCGATTCCGGAGAGGGAGCC | 17 (0.000326%) | | | ATTCAATCGGTAGGAGCGACGGGCGGTGTGTACAAAGGGCAGGGACGTAGTCAACGCGAGCTGATGACTCGCGCTTACTAGGAATTCCTCGTTGAAGACC | 1022 (0.019586%) | | | ATTCCAGCTCCAATAGCGTATATTTAAGTTGTTGCAGTTAAAAAGCTCGTAGTTGAACCTTGGGATGGGTCGGCCGGTCCGCCTTTGGTGTGCATTGGTC | 10 (0.000192%) | | | ATTCGACTCCAAAACACTAACCAACCTTCTTCTTGCTTCTCAAAGCTTTCATGGTGTAGCCAAAGTCCATATGAGTCTTTGGCTTTGTGTCTTCTAACAA | 108 (0.002070%) | | | ATTCGACTCCAAAACACTAACCAACCTTCTTCTTGCTTCTCAAAGCTTTCATGGTGTAGCCAAAGTCCATATGAGTCTTTGGCTTTGTGTCTTTTAACAA | 221 (0.004235%) | | | ATTCGTACTGAAAATCAGAATCAAACGAGCTTTTACCCTTTTGTTCCACACGAGATTTCTGTTCTCGTTGAGCTCATCTTAGGACACCTGCGTTATCTTT | 5 (0.000096%) | | | ATTCGTATTTCATAGTCAGAGGTGAAATTCTTGGATTTATGAAAGACGAACAACTGCGAAAGCATTTGCCAAGGATGTTTTCATTAATCAAGAACGAAAG | 6 (0.000115%) | | | ATTGAGTATAAGAACTTAAACCGCAACCGCATCTTATAAGCCTAAGTAGTGTTTCCTTGTTAGAAGACACAAAGCCAAAGACTCATATGGACTTTGGCTA | 26 (0.000498%) | | | ATTGGAGGGCAAGTCTGGTGCCAGCAGCCGCGGTAATTCCAGCTCCAATAGCGTATATTTAAGTTGTTGCAGTTAAAAAGCTCGTAGTTGAACCTTGGGA | 1512 (0.028977%) | | | ATTGTATCCTTGTTAGAAGACACAAAGCCAAAGACTCATATGGACTTTGGCTACACCATGAAAGCTTTGAGAAGCAAGAAGAAGGTTGGTTAGTGTTTTG | 22 (0.000422%) | | | ATTGTCAGGTGGGGAGTTTGGCTGGGGCGGCACATCTGTTAAAAGATAACGCAGGTGTCCTAAGATGAGCTCAACGAGAACAGAAATCTCGTGTGGAACA | 34 (0.000652%) | | | ATTGTTCCATCGACCAGAGGCTGTTCACCTTGGAGACCTGATGCGGTTATGAGTACGACCGGGCGTGAGCGGCACTCGGTCCTCCGGATTTTCAAGGGCC | 13 (0.000249%) | | | ATTGTTGGTCTTCAACGAGGAATTCCTAGTAAGCGCGAGTCATCAGCTCGCGTTGACTACGTCCCTGCCCTTTGTACACACCGCCCGTCGCTCCTACCGA | 31 (0.000594%) | | | ATTTCATAGTCAGAGGTGAAATTCTTGGATTTATGAAAGACGAACAACTGCGAAAGCATTTGCCAAGGATGTTTTCATTAATCAAGAACGAAAGTTGGGG | 5 (0.000096%) | | | ATTTCGTTCATCACCCTTGGCCGGCTATCGAACAGCCGGACTCCCATCAAAAGATGGTTGCCAAGAACATCTTCGTTACGGTTTGCTAATTCTCGGAATA | 18 (0.000345%) | | | ATTTCGTTCATCACCCTTGGCCGGCTTTCGAACAGCCGGACTCCCATCAAAAGATGGTTGCCAAGAACATCTTCGTTACGGTTTGCTAATTCTCGGAATA | 11 (0.000211%) | | | ATTTCTGCCCTATCAACTTTCGATGGTAGGATAGTGGCCTACCATGGTGGTAACGGGTGACGGAGAATTAGGGTTCGATTCCGGAGAGGGAGCCTGAGAA | 5 (0.000096%) | | | ATTTGCACGTCAGTATCGCTGCGGGCTTCCACCAGAGTTTCCTCTGGCTTTACCCCGCTCAGGCATAGTTCACCATCTTTCGGGTCCCGACAGGCATGCT | 35 (0.000671%) | | | ATTTTATCGGTCGCTCTTGTCCGGAAGCTGTAGATGACCCAAAGTCCATATAGCGACCCCAGGTCAGGCGGGATTACCCGCTGAGTTTAAGCATATCAAT | 18 (0.000345%) | | | ATTTTCAAGGGCCGCCGGGGGCGCACCGGACACCACGCGACGTGCGGTGCTCTTCCAGCCGCTGGACCCTACCTCCGGCTGAGCCGTTTCCAGGGTGGGC | 15 (0.000287%) | | | CAAAATCATCAATCGTTCCAACTAATCTACCGAAGTACTCGGCTAAGAAGAAAGAAGACGGACGAATCCGAGCCAAAGCCGTACAAGCGCGAGATACCTT | 21 (0.000402%) | | | CAAACGTTCAATATGACAAACCCATGCCAAGTAAAGAGAAAATGAAAACTGGTGATTGTTGCGGAAATCGTCCAGGATTCCTCGACCAGGACTTGAAATC | 21 (0.000402%) | | | CAAAGCCAAAGACTCATATGGACTTTGGCTACACCATGAAAGCTTTGAGAAGCAAGAAGAAGGTTGGTTAGTGTTTTGGAGTCGAATATGACTTGATGTC | 92 (0.001763%) | | | CAAAGGGTGTTGGTCGATTAAGACAGCAGGACGGTGGTCATGGAAGTCGAAATCCGCTAAGGAGTGTGTAACAACTCACCTGCCGAATCAACTAGCCCCG | 25 (0.000479%) | | | CAAATCGTCGTCCCTCACCATCCTTTGCTGATGCGGGACGGAAGCTGGTCTCCCGTGTGTTACCGCACGCGGTTGGCCTAAATCCGAGCCAAGGACGCCT | 7 (0.000134%) | | | CAAATTAGGTACGAAACACAGGCCCCGGAACTCATCATCGAGCGTAACATCGCCCGTGAATTAACTGAGAAGGATAGGTGGTAGGTAGTTCGATGCGCGA | 15 (0.000287%) | | | CAAATTTCTGCCCTATCAACTTTCGATGGTAGGATAGTGGCCTACCATGGTGGTAACGGGTGACGGAGAATTAGGGTTCGATTCCGGAGAGGGAGCCTGA | 154 (0.002951%) | | | CAACCTAGGCGAGACAAGGGTTCACATTTCGTTCATCACCCTTGGCCGGCTATCGAACAGCCGGACTCCCATCAAAAGATGGTTGCCAAGAACATCTTCG | 3 (0.000057%) | | | CAACGAAGCACGCCCATCCAACCTAGGCGAGACAAGGGTTCACATTTCGTTCATCACCCTTGGCCGGCTATCGAACAGCCGGACTCCCATCAAAAGATGG | 76 (0.001457%) | | | CAACGGGCAGAGCCCGCGTCGACCTTTTATCTAATAAATGCGTCCCTTCCATAAGTCGGGGTTTGTTGCACGTATTAGCTCTAGAATTACTACGGTTATC | 31 (0.000594%) | | | CAACGTTAGGGAGTCCGGAGACGTCGGCGGGGGCCTCGGGAAGAGTTATCTTTTCTGTTTAACAGCCTGCCCACCCTGGAAACGGCTCAGCCGGAGGTAG | 9 (0.000172%) | | | CAACTTTCGATGGTAGGATAGTGGCCTACCATGGTGGTAACGGGTGACGGAGAATTAGGGTTCGATTCCGGAGAGGGAGCCTGAGAAACGGCTACCACAT | 14 (0.000268%) | | | CAAGGGCCGCCGGGGGCGCACCGGACACCACGCGACGTGCGGTGCTCTTCCAGCCGCTGGACCCTACCTCCGGCTGAGCCGTTTCCAGGGTGGGCAGGCT | 12 (0.000230%) | | | CAAGGGTTCACATTTCGTTCATCACCCTTGGCCGGCTATCGAACAGCCGGACTCCCATCAAAAGATGGTTGCCAAGAACATCTTCGTTACGGTTTGCTAA | 30 (0.000575%) | | | CAAGGGTTCACATTTCGTTCATCACCCTTGGCCGGCTTTCGAACAGCCGGACTCCCATCAAAAGATGGTTGCCAAGAACATCTTCGTTACGGTTTGCTAA | 31 (0.000594%) | | | CAAGTCATATTCGACTCCAAAACACTAACCAACCTTCTTCTTGCTTCTCAAAGCTTTCATGGTGTAGCCAAAGTCCATATGAGTCTTTGGCTTTGTGTCT | 141 (0.002702%) | | | CAAGTCTGGTGCCAGCAGCCGCGGTAATTCCAGCTCCAATAGCGTATATTTAAGTTGTTGCAGTTAAAAAGCTCGTAGTTGAACCTTGGGATGGGTCGGC | 27 (0.000517%) | | | CAATAGCGTATATTTAAGTTGTTGCAGTTAAAAAGCTCGTAGTTGAACCTTGGGATGGGTCGGCCGGTCCGCCTTTGGTGTGCATTGGTCGGCTTGTCCC | 20 (0.000383%) | | | CAATCAGCTTCCTTGCGCCTTACGGGTTTACTCACCCGTTGACTCGCACACATGTCAGACTCCTTGGTCCGTGTTTCAAGACGGGTCGAATGGGGAGCCC | 157 (0.003009%) | | | CAATCATACACATGACATCAAGTCATATTCGACTCCAAAACACTAACCAACCTTCTTCTTGCTTCTCAAAGCTTTCATGGTGTAGCCAAAGTCCATATGA | 134 (0.002568%) | | | CAATCGTTCCAACTAATCTACCGAAGTACTCGGCTAAGAAGAAAGAAGACGGACGAATCCGAGCCAAAGCCGTACAAGCGCGAGATACCTTCGGGACAGC | 4 (0.000077%) | | | CACAAACGTTCAATATGACAAACCCATGCCAAGTAAAGAGAAAATGAAAACTGGTGATTGTTGCGGAAATCGTCCAGGATTCCTCGACCAGGACTTGAAA | 21 (0.000402%) | | | CACAAAGCCAAAGACTCATATGGACTTTGGCTACACCATGAAAGCTTTGAGAAGCAAGAAGAAGGTTGGTTAGTGTTTTGGAGTCGAATATGACTTGATG | 37 (0.000709%) | | | CACAAAGGGTGTTGGTCGATTAAGACAGCAGGACGGTGGTCATGGAAGTCGAAATCCGCTAAGGAGTGTGTAACAACTCACCTGCCGAATCAACTAGCCC | 47 (0.000901%) | | | CACAAATCGTCGTCCCTCACCATCCTTTGCTGATGCGGGACGGAAGCTGGTCTCCCGTGTGTTACCGCACGCGGTTGGCCTAAATCCGAGCCAAGGACGC | 38 (0.000728%) | | | CACACATGTCAGACTCCTTGGTCCGTGTTTCAAGACGGGTCGAATGGGGAGCCCACAGGCCGACGCCCGGAGCACGCTGATGCCGAGGCACGCCGTTAGG | 10 (0.000192%) | | | CACAGGCCCCGGAACTCATCATCGAGCGTAACATCGCCCGTGAATTAACTGAGAAGGATAGGTGGTAGGTAGTTCGATGCGCGAGCATGGAGCCTACGAA | 29 (0.000556%) | | | CACATCTGTTAAAAGATAACGCAGGTGTCCTAAGATGAGCTCAACGAGAACAGAAATCTCGTGTGGAACAAAAGGGTAAAAGCTCGTTTGATTCTGATTT | 6 (0.000115%) | | | CACATGACATCAAGTCATATTCGACTCCAAAACACTAACCAACCTTCTTCTTGCTTCTCAAAGCTTTCATGGTGTAGCCAAAGTCCATATGAGTCTTTGG | 33 (0.000632%) | | | CACATTTCGTTCATCACCCTTGGCCGGCTATCGAACAGCCGGACTCCCATCAAAAGATGGTTGCCAAGAACATCTTCGTTACGGTTTGCTAATTCTCGGA | 7 (0.000134%) | | | CACATTTCGTTCATCACCCTTGGCCGGCTTTCGAACAGCCGGACTCCCATCAAAAGATGGTTGCCAAGAACATCTTCGTTACGGTTTGCTAATTCTCGGA | 7 (0.000134%) | | | CACCATCCTTTGCTGATGCGGGACGGAAGCTGGTCTCCCGTGTGTTACCGCACGCGGTTGGCCTAAATCCGAGCCAAGGACGCCTGGAGCGTACCGACAT | 5 (0.000096%) | | | CACCCGTTGACTCGCACACATGTCAGACTCCTTGGTCCGTGTTTCAAGACGGGTCGAATGGGGAGCCCACAGGCCGACGCCCGGAGCACGCTGATGCCGA | 11 (0.000211%) | | | CACCGGACACCACGCGACGTGCGGTGCTCTTCCAGCCGCTGGACCCTACCTCCGGCTGAGCCGTTTCCAGGGTGGGCAGGCTGTTAAACAGAAAAGATAA | 7 (0.000134%) | | | CACGCCCATCCAACCTAGGCGAGACAAGGGTTCACATTTCGTTCATCACCCTTGGCCGGCTATCGAACAGCCGGACTCCCATCAAAAGATGGTTGCCAAG | 20 (0.000383%) | | | CACGCGCCTAACGGCGTGCCTCGGCATCAGCGTGCTCCGGGCGTCGGCCTGTGGGCTCCCCATTCGACCCGTCTTGAAACACGGACCAAGGAGTCTGACA | 82 (0.001571%) | | | CACGCTTTCACGGTTCGTATTCGTACTGAAAATCAGAATCAAACGAGCTTTTACCCTTTTGTTCCACACGAGATTTCTGTTCTCGTTGAGCTCATCTTAG | 2071 (0.039690%) | | | CACGGTTCGTATTCGTACTGAAAATCAGAATCAAACGAGCTTTTACCCTTTTGTTCCACACGAGATTTCTGTTCTCGTTGAGCTCATCTTAGGACACCTG | 5 (0.000096%) | | | CACTCGGTCCTCCGGATTTTCAAGGGCCGCCGGGGGCGCACCGGACACCACGCGACGTGCGGTGCTCTTCCAGCCGCTGGACCCTACCTCCGGCTGAGCC | 5 (0.000096%) | | | CAGAGCAACGGGCAGAGCCCGCGTCGACCTTTTATCTAATAAATGCGTCCCTTCCATAAGTCGGGGTTTGTTGCACGTATTAGCTCTAGAATTACTACGG | 17 (0.000326%) | | | CAGAGCCCGCGTCGACCTTTTATCTAATAAATGCGTCCCTTCCATAAGTCGGGGTTTGTTGCACGTATTAGCTCTAGAATTACTACGGTTATCCGAGTAG | 9 (0.000172%) | | | CAGAGCGTAGGCTTGCTTTGAGCACTCTAATTTCTTCAAAGTAACAGCGCCGGAGGCACGACCCGGCCAATTAAGACCAGGAGCGTATCGCCGACCGAAG | 38 (0.000728%) | | | CAGCACGCGCCTAACGGCGTGCCTCGGCATCAGCGTGCTCCGGGCGTCGGCCTGTGGGCTCCCCATTCGACCCGTCTTGAAACACGGACCAAGGAGTCTG | 62 (0.001188%) | | | CAGCAGCCGCGGTAATTCCAGCTCCAATAGCGTATATTTAAGTTGTTGCAGTTAAAAAGCTCGTAGTTGAACCTTGGGATGGGTCGGCCGGTCCGCCTTT | 19 (0.000364%) | | | CAGCAGGACGGTGGTCATGGAAGTCGAAATCCGCTAAGGAGTGTGTAACAACTCACCTGCCGAATCAACTAGCCCCGAAAATGGATGGCGCTTAAGCGCG | 8 (0.000153%) | | | CAGCAGTTCTCGGACAAAAATTGCTGAGTGGCCGAGAAGAATGGGCGTGTCATGCGTGGGCTGACATGGATTCTTCGAGGCCTAGGGGTGGCGGTATATA | 5 (0.000096%) | | | CAGCCGCGGTAATTCCAGCTCCAATAGCGTATATTTAAGTTGTTGCAGTTAAAAAGCTCGTAGTTGAACCTTGGGATGGGTCGGCCGGTCCGCCTTTGGT | 10 (0.000192%) | | | CAGCCTGCTAACTAGCTACGTGGAGGCATCCCTTCACGGCCGGCTTCTTAGAGGGACTATGGCCGTTTAGGCCAAGGAAGTTTGAGGCAATAACAGGTCT | 27 (0.000517%) | | | CAGCGTGCTCCGGGCGTCGGCCTGTGGGCTCCCCATTCGACCCGTCTTGAAACACGGACCAAGGAGTCTGACATGTGTGCGAGTCAACGGGTGAGTAAAC | 5 (0.000096%) | | | CAGCTCCAATAGCGTATATTTAAGTTGTTGCAGTTAAAAAGCTCGTAGTTGAACCTTGGGATGGGTCGGCCGGTCCGCCTTTGGTGTGCATTGGTCGGCT | 10 (0.000192%) | | | CAGCTTCCTTGCGCCTTACGGGTTTACTCACCCGTTGACTCGCACACATGTCAGACTCCTTGGTCCGTGTTTCAAGACGGGTCGAATGGGGAGCCCACAG | 114 (0.002185%) | | | CAGGACGGTGGTCATGGAAGTCGAAATCCGCTAAGGAGTGTGTAACAACTCACCTGCCGAATCAACTAGCCCCGAAAATGGATGGCGCTTAAGCGCGCGA | 28 (0.000537%) | | | CAGGGACAGTCGGGGGCATTCGTATTTCATAGTCAGAGGTGAAATTCTTGGATTTATGAAAGACGAACAACTGCGAAAGCATTTGCCAAGGATGTTTTCA | 36 (0.000690%) | | | CAGGGGTTGAAATCGTCGACCAGGTCCGAGACTTCATCGACCGGGTCCGAGGATTCGTCGACCAGGACGGCCGGATGTCCGAGAAAAAAAAATGTTGCCG | 21 (0.000402%) | | | CAGGTGGGGAGTTTGGCTGGGGCGGCACATCTGTTAAAAGATAACGCAGGTGTCCTAAGATGAGCTCAACGAGAACAGAAATCTCGTGTGGAACAAAAGG | 27 (0.000517%) | | | CAGTCGGGGGCATTCGTATTTCATAGTCAGAGGTGAAATTCTTGGATTTATGAAAGACGAACAACTGCGAAAGCATTTGCCAAGGATGTTTTCATTAATC | 7 (0.000134%) | | | CATACACATGACATCAAGTCATATTCGACTCCAAAACACTAACCAACCTTCTTCTTGCTTCTCAAAGCTTTCATGGTGTAGCCAAAGTCCATATGAGTCT | 47 (0.000901%) | | | CATATGAGTCTTTGGCTTTGTGTCTTCTAACAAGGATACA | 6428 (0.049276%) | | | CATATGCTTGTCTCAAAGATTAAGCCATGCATGTGTAAGTATGAACGAATTCAGACTGTGAAACTGCGAATGGCTCATTAAATCAGTTATAGTTTGTTTG | 51 (0.000977%) | | | CATATTCGACTCCAAAACACTAACCAACCTTCTTCTTGCTTCTCAAAGCTTTCATGGTGTAGCCAAAGTCCATATGAGTCTTTGGCTTTGTGTCTTCTAA | 78 (0.001495%) | | | CATATTCGACTCCAAAACACTAACCAACCTTCTTCTTGCTTCTCAAAGCTTTCATGGTGTAGCCAAAGTCCATATGAGTCTTTGGCTTTGTGTCTTTTAA | 308 (0.005903%) | | | CATCAAGTCATATTCGACTCCAAAACACTAACCAACCTTCTTCTTGCTTCTCAAAGCTTTCATGGTGTAGCCAAAGTCCATATGAGTCTTTGGCTTTGTG | 62 (0.001188%) | | | CATCAATCGTTCCAACTAATCTACCGAAGTACTCGGCTAAGAAGAAAGAAGACGGACGAATCCGAGCCAAAGCCGTACAAGCGCGAGATACCTTCGGGAC | 4 (0.000077%) | | | CATCAGAGCAACGGGCAGAGCCCGCGTCGACCTTTTATCTAATAAATGCGTCCCTTCCATAAGTCGGGGTTTGTTGCACGTATTAGCTCTAGAATTACTA | 48 (0.000920%) | | | CATCAGCGTGCTCCGGGCGTCGGCCTGTGGGCTCCCCATTCGACCCGTCTTGAAACACGGACCAAGGAGTCTGACATGTGTGCGAGTCAACGGGTGAGTA | 19 (0.000364%) | | | CATCATCGAGCGTAACATCGCCCGTGAATTAACTGAGAAGGATAGGTGGTAGGTAGTTCGATGCGCGAGCATGGAGCCTACGAACACTAGCTATCCGATC | 13 (0.000249%) | | | CATCATTCAAATTTCTGCCCTATCAACTTTCGATGGTAGGATAGTGGCCTACCATGGTGGTAACGGGTGACGGAGAATTAGGGTTCGATTCCGGAGAGGG | 11 (0.000211%) | | | CATCCAACCTAGGCGAGACAAGGGTTCACATTTCGTTCATCACCCTTGGCCGGCTTTCGAACAGCCGGACTCCCATCAAAAGATGGTTGCCAAGAACATC | 4 (0.000077%) | | | CATCCTTTGCTGATGCGGGACGGAAGCTGGTCTCCCGTGTGTTACCGCACGCGGTTGGCCTAAATCCGAGCCAAGGACGCCTGGAGCGTACCGACATGCG | 131 (0.002511%) | | | CATCGAACTACCTACCACCTATCCTTCTCAGTTAATTCACGGGCGATGTTACGCTCGATGATGAGTTCCGGGGCCTGTGTTTCGTACCTAATTTGAAGGA | 7 (0.000134%) | | | CATCTGTTAAAAGATAACGCAGGTGTCCTAAGATGAGCTCAACGAGAACAGAAATCTCGTGTGGAACAAAAGGGTAAAAGCTCGTTTGATTCTGATTTTC | 4 (0.000077%) | | | CATGAATCATCAGAGCAACGGGCAGAGCCCGCGTCGACCTTTTATCTAATAAATGCGTCCCTTCCATAAGTCGGGGTTTGTTGCACGTATTAGCTCTAGA | 173 (0.003315%) | | | CATGACATCAAGTCATATTCGACTCCAAAACACTAACCAACCTTCTTCTTGCTTCTCAAAGCTTTCATGGTGTAGCCAAAGTCCATATGAGTCTTTGGCT | 21 (0.000402%) | | | CATGGAAGTCGAAATCCGCTAAGGAGTGTGTAACAACTCACCTGCCGAATCAACTAGCCCCGAAAATGGATGGCGCTTAAGCGCGCGACCTATACCCGGC | 58 (0.001112%) | | | CATGGCCTCTGTGCTGGCGACGCATCATTCAAATTTCTGCCCTATCAACTTTCGATGGTAGGATAGTGGCCTACCATGGTGGTAACGGGTGACGGAGAAT | 258 (0.004944%) | | | CATGGGCATCGACACCTTGCGGCTAGGAACTGGAACGAGACGGGTGGCAAAGATTTCGAGTAGCACTTCATACTACCGTGGGTTTTTTAAACCTTCCGAG | 9 (0.000172%) | | | CATGTGTATGATTGAGTATAAGAACTTAAACCGCAACCGCATCTTATAAGCCTAAGTAGTGTTTCCTTGTTAGAAGACACAAAGCCAAAGACTCATATGG | 12 (0.000230%) | | | CATTGTCAGGTGGGGAGTTTGGCTGGGGCGGCACATCTGTTAAAAGATAACGCAGGTGTCCTAAGATGAGCTCAACGAGAACAGAAATCTCGTGTGGAAC | 22 (0.000422%) | | | CATTGTTCCATCGACCAGAGGCTGTTCACCTTGGAGACCTGATGCGGTTATGAGTACGACCGGGCGTGAGCGGCACTCGGTCCTCCGGATTTTCAAGGGC | 20 (0.000383%) | | | CATTTTATCGGTCGCTCTTGTCCGGAAGCTGTAGATGACCCAAAGTCCATATAGCGACCCCAGGTCAGGCGGGATTACCCGCTGAGTTTAAGCATATCAA | 4 (0.000077%) | | | CCAAAGACTCATATGGACTTTGGCTACACCATGAAAGCTTTGAGAAGCAAGAAGAAGGTTGGTTAGTGTTTTGGAGTCGAATATGACTTGATGTCATGTG | 143 (0.002741%) | | | CCAACCTAGGCGAGACAAGGGTTCACATTTCGTTCATCACCCTTGGCCGGCTATCGAACAGCCGGACTCCCATCAAAAGATGGTTGCCAAGAACATCTTC | 18 (0.000345%) | | | CCAACGAAGCACGCCCATCCAACCTAGGCGAGACAAGGGTTCACATTTCGTTCATCACCCTTGGCCGGCTATCGAACAGCCGGACTCCCATCAAAAGATG | 11 (0.000211%) | | | CCAATAGCGTATATTTAAGTTGTTGCAGTTAAAAAGCTCGTAGTTGAACCTTGGGATGGGTCGGCCGGTCCGCCTTTGGTGTGCATTGGTCGGCTTGTCC | 3 (0.000057%) | | | CCAATCAGCTTCCTTGCGCCTTACGGGTTTACTCACCCGTTGACTCGCACACATGTCAGACTCCTTGGTCCGTGTTTCAAGACGGGTCGAATGGGGAGCC | 18 (0.000345%) | | | CCACAAAGGGTGTTGGTCGATTAAGACAGCAGGACGGTGGTCATGGAAGTCGAAATCCGCTAAGGAGTGTGTAACAACTCACCTGCCGAATCAACTAGCC | 7 (0.000134%) | | | CCAGAGCGTAGGCTTGCTTTGAGCACTCTAATTTCTTCAAAGTAACAGCGCCGGAGGCACGACCCGGCCAATTAAGACCAGGAGCGTATCGCCGACCGAA | 28 (0.000537%) | | | CCAGCAGCCGCGGTAATTCCAGCTCCAATAGCGTATATTTAAGTTGTTGCAGTTAAAAAGCTCGTAGTTGAACCTTGGGATGGGTCGGCCGGTCCGCCTT | 10 (0.000192%) | | | CCAGCTCCAATAGCGTATATTTAAGTTGTTGCAGTTAAAAAGCTCGTAGTTGAACCTTGGGATGGGTCGGCCGGTCCGCCTTTGGTGTGCATTGGTCGGC | 8 (0.000153%) | | | CCATCCAACCTAGGCGAGACAAGGGTTCACATTTCGTTCATCACCCTTGGCCGGCTTTCGAACAGCCGGACTCCCATCAAAAGATGGTTGCCAAGAACAT | 6 (0.000115%) | | | CCATCCTTTGCTGATGCGGGACGGAAGCTGGTCTCCCGTGTGTTACCGCACGCGGTTGGCCTAAATCCGAGCCAAGGACGCCTGGAGCGTACCGACATGC | 8 (0.000153%) | | | CCATGCTAATGTATCCAGAGCGTAGGCTTGCTTTGAGCACTCTAATTTCTTCAAAGTAACAGCGCCGGAGGCACGACCCGGCCAATTAAGACCAGGAGCG | 8 (0.000153%) | | | CCATGCTCGCGCATCGAACTACCTACCACCTATCCTTCTCAGTTAATTCACGGGCGATGTTACGCTCGATGATGAGTTCCGGGGCCTGTGTTTCGTACCT | 4 (0.000077%) | | | CCCACAAAGGGTGTTGGTCGATTAAGACAGCAGGACGGTGGTCATGGAAGTCGAAATCCGCTAAGGAGTGTGTAACAACTCACCTGCCGAATCAACTAGC | 9 (0.000172%) | | | CCCATCCAACCTAGGCGAGACAAGGGTTCACATTTCGTTCATCACCCTTGGCCGGCTATCGAACAGCCGGACTCCCATCAAAAGATGGTTGCCAAGAACA | 8 (0.000153%) | | | CCCATCCAACCTAGGCGAGACAAGGGTTCACATTTCGTTCATCACCCTTGGCCGGCTTTCGAACAGCCGGACTCCCATCAAAAGATGGTTGCCAAGAACA | 11 (0.000211%) | | | CCCATGCTAATGTATCCAGAGCGTAGGCTTGCTTTGAGCACTCTAATTTCTTCAAAGTAACAGCGCCGGAGGCACGACCCGGCCAATTAAGACCAGGAGC | 10 (0.000192%) | | | CCCATGGGCATCGACACCTTGCGGCTAGGAACTGGAACGAGACGGGTGGCAAAGATTTCGAGTAGCACTTCATACTACCGTGGGTTTTTTAAACCTTCCG | 13 (0.000249%) | | | CCCGAAGGTATCTCGCGCTTGTACGGCTTTGGCTCGGATTCGTCCGTCTTCTTTCTTCTTAGCCGAGTACTTCGGTAGATTAGTTGGAACGATTGATGAT | 53 (0.001016%) | | | CCCGAGTGTGAGCGAGGTGTGAGTGTCGCCCATGGGCATCGACACCTTGCGGCTAGGAACTGGAACGAGACGGGTAGCAAAGATTTCGAGTAGCACTTCA | 9 (0.000172%) | | | CCCGAGTGTGAGCGAGGTGTGAGTGTCGCCCATGGGCATCGACACCTTGCGGCTAGGAACTGGAACGAGACGGGTGGCAAAGATTTCGAGTAGCACTTCA | 21 (0.000402%) | | | CCCGCCAATCAGCTTCCTTGCGCCTTACGGGTTTACTCACCCGTTGACTCGCACACATGTCAGACTCCTTGGTCCGTGTTTCAAGACGGGTCGAATGGGG | 8 (0.000153%) | | | CCCGCCGAAGCGAGCCTTGGGACCAAAAACAGGGGTTGTACCCCGCCTCCGATTCACGGAGTAAGTAAAATAACGTTAAAAGTAGTGGTATTTCACTTGC | 8 (0.000153%) | | | CCCGCGTCGACCTTTTATCTAATAAATGCGTCCCTTCCATAAGTCGGGGTTTGTTGCACGTATTAGCTCTAGAATTACTACGGTTATCCGAGTAGTAGTT | 38 (0.000728%) | | | CCCGTTGACTCGCACACATGTCAGACTCCTTGGTCCGTGTTTCAAGACGGGTCGAATGGGGAGCCCACAGGCCGACGCCCGGAGCACGCTGATGCCGAGG | 3 (0.000057%) | | | CCCTCACCATCCTTTGCTGATGCGGGACGGAAGCTGGTCTCCCGTGTGTTACCGCACGCGGTTGGCCTAAATCCGAGCCAAGGACGCCTGGAGCGTACCG | 14 (0.000268%) | | | CCCTTGCCTACATTGTTCCATCGACCAGAGGCTGTTCACCTTGGAGACCTGATGCGGTTATGAGTACGACCGGGCGTGAGCGGCACTCGGTCCTCCGGAT | 21 (0.000402%) | | | CCGAAGCGAGCCTTGGGACCAAAAACAGGGGTTGTACCCCGCCTCCGATTCACGGAGTAAGTAAAATAACGTTAAAAGTAGTGGTATTTCACTTGCGCCG | 8 (0.000153%) | | | CCGAAGGTATCTCGCGCTTGTACGGCTTTGGCTCGGATTCGTCCGTCTTCTTTCTTCTTAGCCGAGTACTTCGGTAGATTAGTTGGAACGATTGATGATT | 7 (0.000134%) | | | CCGACTTCCCTTGCCTACATTGTTCCATCGACCAGAGGCTGTTCACCTTGGAGACCTGATGCGGTTATGAGTACGACCGGGCGTGAGCGGCACTCGGTCC | 50 (0.000958%) | | | CCGAGTGTGAGCGAGGTGTGAGTGTCGCCCATGGGCATCGACACCTTGCGGCTAGGAACTGGAACGAGACGGGTAGCAAAGATTTCGAGTAGCACTTCAT | 11 (0.000211%) | | | CCGAGTGTGAGCGAGGTGTGAGTGTCGCCCATGGGCATCGACACCTTGCGGCTAGGAACTGGAACGAGACGGGTGGCAAAGATTTCGAGTAGCACTTCAT | 19 (0.000364%) | | | CCGCCAATCAGCTTCCTTGCGCCTTACGGGTTTACTCACCCGTTGACTCGCACACATGTCAGACTCCTTGGTCCGTGTTTCAAGACGGGTCGAATGGGGA | 49 (0.000939%) | | | CCGCCGGGGGCGCACCGGACACCACGCGACGTGCGGTGCTCTTCCAGCCGCTGGACCCTACCTCCGGCTGAGCCGTTTCCAGGGTGGGCAGGCTGTTAAA | 6 (0.000115%) | | | CCGCCTAGGCTGTCCCGAGTGTGAGCGAGGTGTGAGTGTCGCCCATGGGCATCGACACCTTGCGGCTAGGAACTGGAACGAGACGGGTGGCAAAGATTTC | 92 (0.001763%) | | | CCGCGGTAATTCCAGCTCCAATAGCGTATATTTAAGTTGTTGCAGTTAAAAAGCTCGTAGTTGAACCTTGGGATGGGTCGGCCGGTCCGCCTTTGGTGTG | 6 (0.000115%) | | | CCGGAACCGGGACGTGGCGGTTGACGGCAACGTTAGGGAGTCCGGAGACGTCGGCGGGGGCCTCGGGAAGAGTTATCTTTTCTGTTTAACAGCCTGCCCA | 107 (0.002051%) | | | CCGGACACCACGCGACGTGCGGTGCTCTTCCAGCCGCTGGACCCTACCTCCGGCTGAGCCGTTTCCAGGGTGGGCAGGCTGTTAAACAGAAAAGATAACT | 22 (0.000422%) | | | CCGGAGACGTCGGCGGGGGCCTCGGGAAGAGTTATCTTTTCTGTTTAACAGCCTGCCCACCCTGGAAACGGCTCAGCCGGAGGTAGGGTCCAGCGGCTGG | 8 (0.000153%) | | | CCGGATTTTCAAGGGCCGCCGGGGGCGCACCGGACACCACGCGACGTGCGGTGCTCTTCCAGCCGCTGGACCCTACCTCCGGCTGAGCCGTTTCCAGGGT | 8 (0.000153%) | | | CCGGGACGTGGCGGTTGACGGCAACGTTAGGGAGTCCGGAGACGTCGGCGGGGGCCTCGGGAAGAGTTATCTTTTCTGTTTAACAGCCTGCCCACCCTGG | 21 (0.000402%) | | | CCGGGCGTCGGCCTGTGGGCTCCCCATTCGACCCGTCTTGAAACACGGACCAAGGAGTCTGACATGTGTGCGAGTCAACGGGTGAGTAAACCCGTAAGGC | 7 (0.000134%) | | | CCGGGGGCGCACCGGACACCACGCGACGTGCGGTGCTCTTCCAGCCGCTGGACCCTACCTCCGGCTGAGCCGTTTCCAGGGTGGGCAGGCTGTTAAACAG | 7 (0.000134%) | | | CCGTCGAGTTATCATGAATCATCAGAGCAACGGGCAGAGCCCGCGTCGACCTTTTATCTAATAAATGCGTCCCTTCCATAAGTCGGGGTTTGTTGCACGT | 18 (0.000345%) | | | CCGTTGACTCGCACACATGTCAGACTCCTTGGTCCGTGTTTCAAGACGGGTCGAATGGGGAGCCCACAGGCCGACGCCCGGAGCACGCTGATGCCGAGGC | 4 (0.000077%) | | | CCTAACGGCGTGCCTCGGCATCAGCGTGCTCCGGGCGTCGGCCTGTGGGCTCCCCATTCGACCCGTCTTGAAACACGGACCAAGGAGTCTGACATGTGTG | 7 (0.000134%) | | | CCTAAGTAGTGTTTCCTTGTTAGAAGACACAAAGCCAAAGACTCATATGGACTTTGGCTACACCATGAAAGCTTTGAGAAGCAAGAAGAAGGTTGGTTAG | 67 (0.001284%) | | | CCTACATTGTTCCATCGACCAGAGGCTGTTCACCTTGGAGACCTGATGCGGTTATGAGTACGACCGGGCGTGAGCGGCACTCGGTCCTCCGGATTTTCAA | 8 (0.000153%) | | | CCTAGGCGAGACAAGGGTTCACATTTCGTTCATCACCCTTGGCCGGCTATCGAACAGCCGGACTCCCATCAAAAGATGGTTGCCAAGAACATCTTCGTTA | 10 (0.000192%) | | | CCTAGGCTGTCCCGAGTGTGAGCGAGGTGTGAGTGTCGCCCATGGGCATCGACACCTTGCGGCTAGGAACTGGAACGAGACGGGTGGCAAAGATTTCGAG | 58 (0.001112%) | | | CCTATCAACTTTCGATGGTAGGATAGTGGCCTACCATGGTGGTAACGGGTGACGGAGAATTAGGGTTCGATTCCGGAGAGGGAGCCTGAGAAACGGCTAC | 4 (0.000077%) | | | CCTATGATGTTATCCCATGCTAATGTATCCAGAGCGTAGGCTTGCTTTGAGCACTCTAATTTCTTCAAAGTAACAGCGCCGGAGGCACGACCCGGCCAAT | 24 (0.000460%) | | | CCTCACCATCCTTTGCTGATGCGGGACGGAAGCTGGTCTCCCGTGTGTTACCGCACGCGGTTGGCCTAAATCCGAGCCAAGGACGCCTGGAGCGTACCGA | 66 (0.001265%) | | | CCTCAGCCTGCTAACTAGCTACGTGGAGGCATCCCTTCACGGCCGGCTTCTTAGAGGGACTATGGCCGTTTAGGCCAAGGAAGTTTGAGGCAATAACAGG | 107 (0.002051%) | | | CCTCGGCATCAGCGTGCTCCGGGCGTCGGCCTGTGGGCTCCCCATTCGACCCGTCTTGAAACACGGACCAAGGAGTCTGACATGTGTGCGAGTCAACGGG | 5 (0.000096%) | | | CCTCTGTGCTGGCGACGCATCATTCAAATTTCTGCCCTATCAACTTTCGATGGTAGGATAGTGGCCTACCATGGTGGTAACGGGTGACGGAGAATTAGGG | 93 (0.001782%) | | | CCTGCTAACTAGCTACGTGGAGGCATCCCTTCACGGCCGGCTTCTTAGAGGGACTATGGCCGTTTAGGCCAAGGAAGTTTGAGGCAATAACAGGTCTGTG | 6 (0.000115%) | | | CCTTGCCTACATTGTTCCATCGACCAGAGGCTGTTCACCTTGGAGACCTGATGCGGTTATGAGTACGACCGGGCGTGAGCGGCACTCGGTCCTCCGGATT | 14 (0.000268%) | | | CCTTGCGCCTTACGGGTTTACTCACCCGTTGACTCGCACACATGTCAGACTCCTTGGTCCGTGTTTCAAGACGGGTCGAATGGGGAGCCCACAGGCCGAC | 3 (0.000057%) | | | CCTTGTTAGAAGACACAAAGCCAAAGACTCATATGGACTTTGGCTACACCATGAAAGCTTTGAGAAGCAAGAAGAAGGTTGGTTAGTGTTTTGGAGTCGA | 50 (0.000958%) | | | CGAAACACAGGCCCCGGAACTCATCATCGAGCGTAACATCGCCCGTGAATTAACTGAGAAGGATAGGTGGTAGGTAGTTCGATGCGCGAGCATGGAGCCT | 29 (0.000556%) | | | CGAAATCCTATGATGTTATCCCATGCTAATGTATCCAGAGCGTAGGCTTGCTTTGAGCACTCTAATTTCTTCAAAGTAACAGCGCCGGAGGCACGACCCG | 457 (0.008758%) | | | CGAACTACCTACCACCTATCCTTCTCAGTTAATTCACGGGCGATGTTACGCTCGATGATGAGTTCCGGGGCCTGTGTTTCGTACCTAATTTGAAGGAATT | 6 (0.000115%) | | | CGAAGCACGCCCATCCAACCTAGGCGAGACAAGGGTTCACATTTCGTTCATCACCCTTGGCCGGCTATCGAACAGCCGGACTCCCATCAAAAGATGGTTG | 18 (0.000345%) | | | CGAAGGTATCTCGCGCTTGTACGGCTTTGGCTCGGATTCGTCCGTCTTCTTTCTTCTTAGCCGAGTACTTCGGTAGATTAGTTGGAACGATTGATGATTT | 19 (0.000364%) | | | CGACCAGGGGTTGAAATCGTCGACCAGGTCCGAGACTTCATCGACCGGGTCCGAGGATTCGTCGACCAGGACGGCCGGATGTCCGAGAAAAAAAAATGTT | 28 (0.000537%) | | | CGACCCGCCGAAGCGAGCCTTGGGACCAAAAACAGGGGTTGTACCCCGCCTCCGATTCACGGAGTAAGTAAAATAACGTTAAAAGTAGTGGTATTTCACT | 5 (0.000096%) | | | CGACGCATCATTCAAATTTCTGCCCTATCAACTTTCGATGGTAGGATAGTGGCCTACCATGGTGGTAACGGGTGACGGAGAATTAGGGTTCGATTCCGGA | 8 (0.000153%) | | | CGACGGGCGGTGTGTACAAAGGGCAGGGACGTAGTCAACGCGAGCTGATGACTCGCGCTTACTAGGAATTCCTCGTTGAAGACCAACAATTGCAATGATC | 27 (0.000517%) | | | CGACGTGGGTGGTTCGCCGCCCGCGACGTCGCGAGAAGTCCACTAAACCTTATCATTTAGAGGAAGGAGAAGTCGTAACAAGGTTTCCGTAGGTGAACCT | 115 (0.002204%) | | | CGACTCCAAAACACTAACCAACCTTCTTCTTGCTTCTCAAAGCTTTCATGGTGTAGCCAAAGTCCATATGAGTCTTTGGCTTTGTGTCTTCTAACAAGGA | 57 (0.001092%) | | | CGACTCCAAAACACTAACCAACCTTCTTCTTGCTTCTCAAAGCTTTCATGGTGTAGCCAAAGTCCATATGAGTCTTTGGCTTTGTGTCTTTTAACAAGGA | 38 (0.000728%) | | | CGACTTCCCTTGCCTACATTGTTCCATCGACCAGAGGCTGTTCACCTTGGAGACCTGATGCGGTTATGAGTACGACCGGGCGTGAGCGGCACTCGGTCCT | 25 (0.000479%) | | | CGAGACAAGGGTTCACATTTCGTTCATCACCCTTGGCCGGCTATCGAACAGCCGGACTCCCATCAAAAGATGGTTGCCAAGAACATCTTCGTTACGGTTT | 3 (0.000057%) | | | CGAGACAAGGGTTCACATTTCGTTCATCACCCTTGGCCGGCTTTCGAACAGCCGGACTCCCATCAAAAGATGGTTGCCAAGAACATCTTCGTTACGGTTT | 9 (0.000172%) | | | CGAGCTCGTGTAAAGTTGGGAATTCGTTAAGGAGCTGTTGCTTTGTTAGTGTAGAAACACTTGTGTAGAATTGGGGATTGTTTTTTTTGGAGTGATTTAG | 753 (0.014431%) | | | CGAGGTGTGAGTGTCGCCCATGGGCATCGACACCTTGCGGCTAGGAACTGGAACGAGACGGGTGGCAAAGATTTCGAGTAGCACTTCATACTACCGTGGG | 24 (0.000460%) | | | CGAGTGTGAGCGAGGTGTGAGTGTCGCCCATGGGCATCGACACCTTGCGGCTAGGAACTGGAACGAGACGGGTAGCAAAGATTTCGAGTAGCACTTCATA | 20 (0.000383%) | | | CGAGTGTGAGCGAGGTGTGAGTGTCGCCCATGGGCATCGACACCTTGCGGCTAGGAACTGGAACGAGACGGGTGGCAAAGATTTCGAGTAGCACTTCATA | 32 (0.000613%) | | | CGAGTTATCATGAATCATCAGAGCAACGGGCAGAGCCCGCGTCGACCTTTTATCTAATAAATGCGTCCCTTCCATAAGTCGGGGTTTGTTGCACGTATTA | 143 (0.002741%) | | | CGATCCATTACATTTTATCGGTCGCTCTTGTCCGGAAGCTGTAGATGACCCAAAGTCCATATAGCGACCCCAGGTCAGGCGGGATTACCCGCTGAGTTTA | 5 (0.000096%) | | | CGATCCCGCCAATCAGCTTCCTTGCGCCTTACGGGTTTACTCACCCGTTGACTCGCACACATGTCAGACTCCTTGGTCCGTGTTTCAAGACGGGTCGAAT | 6 (0.000115%) | | | CGATCCGTCGAGTTATCATGAATCATCAGAGCAACGGGCAGAGCCCGCGTCGACCTTTTATCTAATAAATGCGTCCCTTCCATAAGTCGGGGTTTGTTGC | 177 (0.003392%) | | | CGATCGACCCGCCGAAGCGAGCCTTGGGACCAAAAACAGGGGTTGTACCCCGCCTCCGATTCACGGAGTAAGTAAAATAACGTTAAAAGTAGTGGTATTT | 7 (0.000134%) | | | CGATTAAGACAGCAGGACGGTGGTCATGGAAGTCGAAATCCGCTAAGGAGTGTGTAACAACTCACCTGCCGAATCAACTAGCCCCGAAAATGGATGGCGC | 12 (0.000230%) | | | CGATTAGTCTTTCGCCCCTATACCCAAGTCAGACGAACGATTTGCACGTCAGTATCGCTGCGGGCTTCCACCAGAGTTTCCTCTGGCTTTACCCCGCTCA | 12 (0.000230%) | | | CGATTTGCACGTCAGTATCGCTGCGGGCTTCCACCAGAGTTTCCTCTGGCTTTACCCCGCTCAGGCATAGTTCACCATCTTTCGGGTCCCGACAGGCATG | 485 (0.009295%) | | | CGCACACATGTCAGACTCCTTGGTCCGTGTTTCAAGACGGGTCGAATGGGGAGCCCACAGGCCGACGCCCGGAGCACGCTGATGCCGAGGCACGCCGTTA | 19 (0.000364%) | | | CGCACCGGACACCACGCGACGTGCGGTGCTCTTCCAGCCGCTGGACCCTACCTCCGGCTGAGCCGTTTCCAGGGTGGGCAGGCTGTTAAACAGAAAAGAT | 8 (0.000153%) | | | CGCATCATTCAAATTTCTGCCCTATCAACTTTCGATGGTAGGATAGTGGCCTACCATGGTGGTAACGGGTGACGGAGAATTAGGGTTCGATTCCGGAGAG | 5 (0.000096%) | | | CGCCAATCAGCTTCCTTGCGCCTTACGGGTTTACTCACCCGTTGACTCGCACACATGTCAGACTCCTTGGTCCGTGTTTCAAGACGGGTCGAATGGGGAG | 1227 (0.023515%) | | | CGCCCATCCAACCTAGGCGAGACAAGGGTTCACATTTCGTTCATCACCCTTGGCCGGCTATCGAACAGCCGGACTCCCATCAAAAGATGGTTGCCAAGAA | 85 (0.001629%) | | | CGCCCATCCAACCTAGGCGAGACAAGGGTTCACATTTCGTTCATCACCCTTGGCCGGCTTTCGAACAGCCGGACTCCCATCAAAAGATGGTTGCCAAGAA | 83 (0.001591%) | | | CGCCCATGGGCATCGACACCTTGCGGCTAGGAACTGGAACGAGACGGGTGGCAAAGATTTCGAGTAGCACTTCATACTACCGTGGGTTTTTTAAACCTTC | 31 (0.000594%) | | | CGCCGAAGCGAGCCTTGGGACCAAAAACAGGGGTTGTACCCCGCCTCCGATTCACGGAGTAAGTAAAATAACGTTAAAAGTAGTGGTATTTCACTTGCGC | 6 (0.000115%) | | | CGCCGCCCGCGACGTCGCGAGAAGTCCACTAAACCTTATCATTTAGAGGAAGGAGAAGTCGTAACAAGGTTTCCGTAGGTGAACCTGCGGAAGGATCGAT | 12 (0.000230%) | | | CGCCGGGGGCGCACCGGACACCACGCGACGTGCGGTGCTCTTCCAGCCGCTGGACCCTACCTCCGGCTGAGCCGTTTCCAGGGTGGGCAGGCTGTTAAAC | 9 (0.000172%) | | | CGCCTAACGGCGTGCCTCGGCATCAGCGTGCTCCGGGCGTCGGCCTGTGGGCTCCCCATTCGACCCGTCTTGAAACACGGACCAAGGAGTCTGACATGTG | 39 (0.000747%) | | | CGCCTAGGCTGTCCCGAGTGTGAGCGAGGTGTGAGTGTCGCCCATGGGCATCGACACCTTGCGGCTAGGAACTGGAACGAGACGGGTGGCAAAGATTTCG | 148 (0.002836%) | | | CGCCTTACGGGTTTACTCACCCGTTGACTCGCACACATGTCAGACTCCTTGGTCCGTGTTTCAAGACGGGTCGAATGGGGAGCCCACAGGCCGACGCCCG | 9 (0.000172%) | | | CGCGCATCGAACTACCTACCACCTATCCTTCTCAGTTAATTCACGGGCGATGTTACGCTCGATGATGAGTTCCGGGGCCTGTGTTTCGTACCTAATTTGA | 14 (0.000268%) | | | CGCGCCTAACGGCGTGCCTCGGCATCAGCGTGCTCCGGGCGTCGGCCTGTGGGCTCCCCATTCGACCCGTCTTGAAACACGGACCAAGGAGTCTGACATG | 27 (0.000517%) | | | CGCGCTTGTACGGCTTTGGCTCGGATTCGTCCGTCTTCTTTCTTCTTAGCCGAGTACTTCGGTAGATTAGTTGGAACGATTGATGATTTTGAGTTAATTG | 25 (0.000479%) | | | CGCGGTAATTCCAGCTCCAATAGCGTATATTTAAGTTGTTGCAGTTAAAAAGCTCGTAGTTGAACCTTGGGATGGGTCGGCCGGTCCGCCTTTGGTGTGC | 5 (0.000096%) | | | CGCTTGTACGGCTTTGGCTCGGATTCGTCCGTCTTCTTTCTTCTTAGCCGAGTACTTCGGTAGATTAGTTGGAACGATTGATGATTTTGAGTTAATTGAA | 6 (0.000115%) | | | CGCTTTCACGGTTCGTATTCGTACTGAAAATCAGAATCAAACGAGCTTTTACCCTTTTGTTCCACACGAGATTTCTGTTCTCGTTGAGCTCATCTTAGGA | 7 (0.000134%) | | | CGGAACCGGGACGTGGCGGTTGACGGCAACGTTAGGGAGTCCGGAGACGTCGGCGGGGGCCTCGGGAAGAGTTATCTTTTCTGTTTAACAGCCTGCCCAC | 83 (0.001591%) | | | CGGACACCACGCGACGTGCGGTGCTCTTCCAGCCGCTGGACCCTACCTCCGGCTGAGCCGTTTCCAGGGTGGGCAGGCTGTTAAACAGAAAAGATAACTC | 58 (0.001112%) | | | CGGAGACGTCGGCGGGGGCCTCGGGAAGAGTTATCTTTTCTGTTTAACAGCCTGCCCACCCTGGAAACGGCTCAGCCGGAGGTAGGGTCCAGCGGCTGGA | 41 (0.000786%) | | | CGGAGGACATTGTCAGGTGGGGAGTTTGGCTGGGGCGGCACATCTGTTAAAAGATAACGCAGGTGTCCTAAGATGAGCTCAACGAGAACAGAAATCTCGT | 45 (0.000862%) | | | CGGAGTTTTTTCAGCAGTTCTCGGACAAAAATTGCTGAGTGGCCGAGAAGAATGGGCGTGTCATGCGTGGGCTGACATGGATTCTTCGAGGCCTAGGGGT | 3 (0.000057%) | | | CGGATAGCTAGTGTTCGTAGGCTCCATGCTCGCGCATCGAACTACCTACCACCTATCCTTCTCAGTTAATTCACGGGCGATGTTACGCTCGATGATGAGT | 695 (0.013319%) | | | CGGATTTTCAAGGGCCGCCGGGGGCGCACCGGACACCACGCGACGTGCGGTGCTCTTCCAGCCGCTGGACCCTACCTCCGGCTGAGCCGTTTCCAGGGTG | 40 (0.000767%) | | | CGGCAACGTTAGGGAGTCCGGAGACGTCGGCGGGGGCCTCGGGAAGAGTTATCTTTTCTGTTTAACAGCCTGCCCACCCTGGAAACGGCTCAGCCGGAGG | 24 (0.000460%) | | | CGGCAATTCCCCGCCACATCCTCTCAAACGCAATGGAAAGAGAGAAAGGACGAGGTCTTGACCGTCATCTTTTGCCCGAAGGACGGATGAGCTTTGGCGG | 1185 (0.022710%) | | | CGGCACATCTGTTAAAAGATAACGCAGGTGTCCTAAGATGAGCTCAACGAGAACAGAAATCTCGTGTGGAACAAAAGGGTAAAAGCTCGTTTGATTCTGA | 10 (0.000192%) | | | CGGCACTCGGTCCTCCGGATTTTCAAGGGCCGCCGGGGGCGCACCGGACACCACGCGACGTGCGGTGCTCTTCCAGCCGCTGGACCCTACCTCCGGCTGA | 50 (0.000958%) | | | CGGCATCAGCGTGCTCCGGGCGTCGGCCTGTGGGCTCCCCATTCGACCCGTCTTGAAACACGGACCAAGGAGTCTGACATGTGTGCGAGTCAACGGGTGA | 10 (0.000192%) | | | CGGCCTGTGGGCTCCCCATTCGACCCGTCTTGAAACACGGACCAAGGAGTCTGACATGTGTGCGAGTCAACGGGTGAGTAAACCCGTAAGGCGCAAGGAA | 8 (0.000153%) | | | CGGCGACGTGGGTGGTTCGCCGCCCGCGACGTCGCGAGAAGTCCACTAAACCTTATCATTTAGAGGAAGGAGAAGTCGTAACAAGGTTTCCGTAGGTGAA | 502 (0.009621%) | | | CGGCGTGCCTCGGCATCAGCGTGCTCCGGGCGTCGGCCTGTGGGCTCCCCATTCGACCCGTCTTGAAACACGGACCAAGGAGTCTGACATGTGTGCGAGT | 5 (0.000096%) | | | CGGCTTTGGCTCGGATTCGTCCGTCTTCTTTCTTCTTAGCCGAGTACTTCGGTAGATTAGTTGGAACGATTGATGATTTTGAGTTAATTGAACGTTCGGC | 3 (0.000057%) | | | CGGGACGTGGCGGTTGACGGCAACGTTAGGGAGTCCGGAGACGTCGGCGGGGGCCTCGGGAAGAGTTATCTTTTCTGTTTAACAGCCTGCCCACCCTGGA | 43 (0.000824%) | | | CGGGCAGAGCCCGCGTCGACCTTTTATCTAATAAATGCGTCCCTTCCATAAGTCGGGGTTTGTTGCACGTATTAGCTCTAGAATTACTACGGTTATCCGA | 50 (0.000958%) | | | CGGGCGAGAGACCGATAGCGAACAAGTACCGCGAGGTAAAGATGAAAAGGACTTTGAAAAGAGAGTCAAAGAGTGCTTGAAATTGTCGGGAGGGAAGCGG | 3 (0.000057%) | | | CGGGCGGTGTGTACAAAGGGCAGGGACGTAGTCAACGCGAGCTGATGACTCGCGCTTACTAGGAATTCCTCGTTGAAGACCAACAATTGCAATGATCGAT | 37 (0.000709%) | | | CGGGCGTCGGCCTGTGGGCTCCCCATTCGACCCGTCTTGAAACACGGACCAAGGAGTCTGACATGTGTGCGAGTCAACGGGTGAGTAAACCCGTAAGGCG | 46 (0.000882%) | | | CGGGGGCATTCGTATTTCATAGTCAGAGGTGAAATTCTTGGATTTATGAAAGACGAACAACTGCGAAAGCATTTGCCAAGGATGTTTTCATTAATCAAGA | 29 (0.000556%) | | | CGGGGGCGCACCGGACACCACGCGACGTGCGGTGCTCTTCCAGCCGCTGGACCCTACCTCCGGCTGAGCCGTTTCCAGGGTGGGCAGGCTGTTAAACAGA | 31 (0.000594%) | | | CGGGTTTACTCACCCGTTGACTCGCACACATGTCAGACTCCTTGGTCCGTGTTTCAAGACGGGTCGAATGGGGAGCCCACAGGCCGACGCCCGGAGCACG | 17 (0.000326%) | | | CGGTAATTCCAGCTCCAATAGCGTATATTTAAGTTGTTGCAGTTAAAAAGCTCGTAGTTGAACCTTGGGATGGGTCGGCCGGTCCGCCTTTGGTGTGCAT | 15 (0.000287%) | | | CGGTAGGAGCGACGGGCGGTGTGTACAAAGGGCAGGGACGTAGTCAACGCGAGCTGATGACTCGCGCTTACTAGGAATTCCTCGTTGAAGACCAACAATT | 52 (0.000997%) | | | CGGTCCTCCGGATTTTCAAGGGCCGCCGGGGGCGCACCGGACACCACGCGACGTGCGGTGCTCTTCCAGCCGCTGGACCCTACCTCCGGCTGAGCCGTTT | 8 (0.000153%) | | | CGGTGGTCATGGAAGTCGAAATCCGCTAAGGAGTGTGTAACAACTCACCTGCCGAATCAACTAGCCCCGAAAATGGATGGCGCTTAAGCGCGCGACCTAT | 33 (0.000632%) | | | CGGTTGACGGCAACGTTAGGGAGTCCGGAGACGTCGGCGGGGGCCTCGGGAAGAGTTATCTTTTCTGTTTAACAGCCTGCCCACCCTGGAAACGGCTCAG | 5 (0.000096%) | | | CGGTTTAAGTTCTTATACTCAATCATACACATGACATCAAGTCATATTCGACTCCAAAACACTAACCAACCTTCTTCTTGCTTCTCAAAGCTTTCATGGT | 87 (0.001667%) | | | CGGTTTAAGTTGTTATACTCAATCATACACATGACAACAAGTCATATTCGACTCCAAAACACTAACCAACCTTCTTCTTGCTTCTCAAAGCTTTCATGGT | 97 (0.001859%) | | | CGTAAGAATTGTATCCTTGTTAAAAGACACAAAGCCAAAGACTCATATGGACTTTGGCTACACCATGAAAGCTTTGAGAAGCAAGAAGAAGGTTGGTTAG | 519 (0.009946%) | | | CGTAAGAATTGTATCCTTGTTAGAAGACACAAAGCCAAAG | 235 (0.001801%) | | | CGTAGGCTCCATGCTCGCGCATCGAACTACCTACCACCTATCCTTCTCAGTTAATTCACGGGCGATGTTACGCTCGATGATGAGTTCCGGGGCCTGTGTT | 3 (0.000057%) | | | CGTATATTTAAGTTGTTGCAGTTAAAAAGCTCGTAGTTGAACCTTGGGATGGGTCGGCCGGTCCGCCTTTGGTGTGCATTGGTCGGCTTGTCCCTTCGGT | 3 (0.000057%) | | | CGTATTCGTACTGAAAATCAGAATCAAACGAGCTTTTACCCTTTTGTTCCACACGAGATTTCTGTTCTCGTTGAGCTCATCTTAGGACACCTGCGTTATC | 15 (0.000287%) | | | CGTATTTCATAGTCAGAGGTGAAATTCTTGGATTTATGAAAGACGAACAACTGCGAAAGCATTTGCCAAGGATGTTTTCATTAATCAAGAACGAAAGTTG | 7 (0.000134%) | | | CGTCCCTCACCATCCTTTGCTGATGCGGGACGGAAGCTGGTCTCCCGTGTGTTACCGCACGCGGTTGGCCTAAATCCGAGCCAAGGACGCCTGGAGCGTA | 11 (0.000211%) | | | CGTCGAGTTATCATGAATCATCAGAGCAACGGGCAGAGCCCGCGTCGACCTTTTATCTAATAAATGCGTCCCTTCCATAAGTCGGGGTTTGTTGCACGTA | 104 (0.001993%) | | | CGTCGTCCCTCACCATCCTTTGCTGATGCGGGACGGAAGCTGGTCTCCCGTGTGTTACCGCACGCGGTTGGCCTAAATCCGAGCCAAGGACGCCTGGAGC | 15 (0.000287%) | | | CGTGCCTCGGCATCAGCGTGCTCCGGGCGTCGGCCTGTGGGCTCCCCATTCGACCCGTCTTGAAACACGGACCAAGGAGTCTGACATGTGTGCGAGTCAA | 20 (0.000383%) | | | CGTGCTCCGGGCGTCGGCCTGTGGGCTCCCCATTCGACCCGTCTTGAAACACGGACCAAGGAGTCTGACATGTGTGCGAGTCAACGGGTGAGTAAACCCG | 9 (0.000172%) | | | CGTGGGTGGTTCGCCGCCCGCGACGTCGCGAGAAGTCCACTAAACCTTATCATTTAGAGGAAGGAGAAGTCGTAACAAGGTTTCCGTAGGTGAACCTGCG | 35 (0.000671%) | | | CGTTAAGGAGCTGTTGCTTTGTTAGTGTAGAAACACTTGTGTAGAATTGGGGATTGTTTTTTTTGGAGTGATTTAGGGGAGGGTCGAATCTTAGCGACAA | 233 (0.004465%) | | | CGTTAGGGAGTCCGGAGACGTCGGCGGGGGCCTCGGGAAGAGTTATCTTTTCTGTTTAACAGCCTGCCCACCCTGGAAACGGCTCAGCCGGAGGTAGGGT | 3 (0.000057%) | | | CGTTCATCACCCTTGGCCGGCTTTCGAACAGCCGGACTCCCATCAAAAGATGGTTGCCAAGAACATCTTCGTTACGGTTTGCTAATTCTCGGAATAACAT | 18 (0.000345%) | | | CGTTGACTCGCACACATGTCAGACTCCTTGGTCCGTGTTTCAAGACGGGTCGAATGGGGAGCCCACAGGCCGACGCCCGGAGCACGCTGATGCCGAGGCA | 9 (0.000172%) | | | CTAAATACGGGCGAGAGACCGATAGCGAACAAGTACCGCGAGGTAAAGATGAAAAGGACTTTGAAAAGAGAGTCAAAGAGTGCTTGAAATTGTCGGGAGG | 766 (0.014680%) | | | CTAACGGCGTGCCTCGGCATCAGCGTGCTCCGGGCGTCGGCCTGTGGGCTCCCCATTCGACCCGTCTTGAAACACGGACCAAGGAGTCTGACATGTGTGC | 107 (0.002051%) | | | CTAACTAGCTACGTGGAGGCATCCCTTCACGGCCGGCTTCTTAGAGGGACTATGGCCGTTTAGGCCAAGGAAGTTTGAGGCAATAACAGGTCTGTGATGC | 38 (0.000728%) | | | CTAAGTAGTGTTTCCTTGTTAGAAGACACAAAGCCAAAGACTCATATGGACTTTGGCTACACCATGAAAGCTTTGAGAAGCAAGAAGAAGGTTGGTTAGT | 72 (0.001380%) | | | CTAATGTATCCAGAGCGTAGGCTTGCTTTGAGCACTCTAATTTCTTCAAAGTAACAGCGCCGGAGGCACGACCCGGCCAATTAAGACCAGGAGCGTATCG | 28 (0.000537%) | | | CTACATTGTTCCATCGACCAGAGGCTGTTCACCTTGGAGACCTGATGCGGTTATGAGTACGACCGGGCGTGAGCGGCACTCGGTCCTCCGGATTTTCAAG | 49 (0.000939%) | | | CTACCTACCACCTATCCTTCTCAGTTAATTCACGGGCGATGTTACGCTCGATGATGAGTTCCGGGGCCTGTGTTTCGTACCTAATTTGAAGGAATTGTTG | 20 (0.000383%) | | | CTAGCTACGTGGAGGCATCCCTTCACGGCCGGCTTCTTAGAGGGACTATGGCCGTTTAGGCCAAGGAAGTTTGAGGCAATAACAGGTCTGTGATGCCCTT | 23 (0.000441%) | | | CTAGGCGAGACAAGGGTTCACATTTCGTTCATCACCCTTGGCCGGCTATCGAACAGCCGGACTCCCATCAAAAGATGGTTGCCAAGAACATCTTCGTTAC | 28 (0.000537%) | | | CTAGGCGAGACAAGGGTTCACATTTCGTTCATCACCCTTGGCCGGCTTTCGAACAGCCGGACTCCCATCAAAAGATGGTTGCCAAGAACATCTTCGTTAC | 66 (0.001265%) | | | CTAGGCTGTCCCGAGTGTGAGCGAGGTGTGAGTGTCGCCCATGGGCATCGACACCTTGCGGCTAGGAACTGGAACGAGACGGGTGGCAAAGATTTCGAGT | 402 (0.007704%) | | | CTAGTGTTCGTAGGCTCCATGCTCGCGCATCGAACTACCTACCACCTATCCTTCTCAGTTAATTCACGGGCGATGTTACGCTCGATGATGAGTTCCGGGG | 19 (0.000364%) | | | CTATCAACTTTCGATGGTAGGATAGTGGCCTACCATGGTGGTAACGGGTGACGGAGAATTAGGGTTCGATTCCGGAGAGGGAGCCTGAGAAACGGCTACC | 3 (0.000057%) | | | CTATGATGTTATCCCATGCTAATGTATCCAGAGCGTAGGCTTGCTTTGAGCACTCTAATTTCTTCAAAGTAACAGCGCCGGAGGCACGACCCGGCCAATT | 218 (0.004178%) | | | CTCAAAATCATCAATCGTTCCAACTAATCTACCGAAGTACTCGGCTAAGAAGAAAGAAGACGGACGAATCCGAGCCAAAGCCGTACAAGCGCGAGATACC | 1983 (0.038003%) | | | CTCAAAGATTAAGCCATGCATGTGTAAGTATGAACGAATTCAGACTGTGAAACTGCGAATGGCTCATTAAATCAGTTATAGTTTGTTTGATGGTAACTAC | 19 (0.000364%) | | | CTCAATCATACACATGACATCAAGTCATATTCGACTCCAAAACACTAACCAACCTTCTTCTTGCTTCTCAAAGCTTTCATGGTGTAGCCAAAGTCCATAT | 61 (0.001169%) | | | CTCACCATCCTTTGCTGATGCGGGACGGAAGCTGGTCTCCCGTGTGTTACCGCACGCGGTTGGCCTAAATCCGAGCCAAGGACGCCTGGAGCGTACCGAC | 16 (0.000307%) | | | CTCACCCGTTGACTCGCACACATGTCAGACTCCTTGGTCCGTGTTTCAAGACGGGTCGAATGGGGAGCCCACAGGCCGACGCCCGGAGCACGCTGATGCC | 8 (0.000153%) | | | CTCAGCCTGCTAACTAGCTACGTGGAGGCATCCCTTCACGGCCGGCTTCTTAGAGGGACTATGGCCGTTTAGGCCAAGGAAGTTTGAGGCAATAACAGGT | 26 (0.000498%) | | | CTCATGTGTATGATTGAGTATAAGAACTTAAACCGCAACCGCATCTTATAAGCCTAAGTAGTGTTTCCTTGTTAGAAGACACAAAGCCAAAGACTCATAT | 110 (0.002108%) | | | CTCCAAAACACTAACCAACCTTCTTCTTGCTTCTCAAAGCTTTCATGGTGTAGCCAAAGTCCATATGAGTCTTTGGCTTTGTGTCTTCTAACAAGGATAC | 29 (0.000556%) | | | CTCCAATAGCGTATATTTAAGTTGTTGCAGTTAAAAAGCTCGTAGTTGAACCTTGGGATGGGTCGGCCGGTCCGCCTTTGGTGTGCATTGGTCGGCTTGT | 32 (0.000613%) | | | CTCCATGCTCGCGCATCGAACTACCTACCACCTATCCTTCTCAGTTAATTCACGGGCGATGTTACGCTCGATGATGAGTTCCGGGGCCTGTGTTTCGTAC | 9 (0.000172%) | | | CTCCGGGCGTCGGCCTGTGGGCTCCCCATTCGACCCGTCTTGAAACACGGACCAAGGAGTCTGACATGTGTGCGAGTCAACGGGTGAGTAAACCCGTAAG | 17 (0.000326%) | | | CTCGCACACATGTCAGACTCCTTGGTCCGTGTTTCAAGACGGGTCGAATGGGGAGCCCACAGGCCGACGCCCGGAGCACGCTGATGCCGAGGCACGCCGT | 19 (0.000364%) | | | CTCGCGCATCGAACTACCTACCACCTATCCTTCTCAGTTAATTCACGGGCGATGTTACGCTCGATGATGAGTTCCGGGGCCTGTGTTTCGTACCTAATTT | 7 (0.000134%) | | | CTCGCGCTTGTACGGCTTTGGCTCGGATTCGTCCGTCTTCTTTCTTCTTAGCCGAGTACTTCGGTAGATTAGTTGGAACGATTGATGATTTTGAGTTAAT | 10 (0.000192%) | | | CTCGGCATCAGCGTGCTCCGGGCGTCGGCCTGTGGGCTCCCCATTCGACCCGTCTTGAAACACGGACCAAGGAGTCTGACATGTGTGCGAGTCAACGGGT | 24 (0.000460%) | | | CTCGGTCCTCCGGATTTTCAAGGGCCGCCGGGGGCGCACCGGACACCACGCGACGTGCGGTGCTCTTCCAGCCGCTGGACCCTACCTCCGGCTGAGCCGT | 32 (0.000613%) | | | CTCGTGTAAAGTTGGGAATTCGTTAAGGAGCTGTTGCTTTGTTAGTGTAGAAACACTTGTGTAGAATTGGGGATTGTTTTTTTTGGAGTGATTTAGGGGA | 33 (0.000632%) | | | CTCTGTGCTGGCGACGCATCATTCAAATTTCTGCCCTATCAACTTTCGATGGTAGGATAGTGGCCTACCATGGTGGTAACGGGTGACGGAGAATTAGGGT | 57 (0.001092%) | | | CTGAGAAGGGTTCGAGTGTGAGCATGCCTGTCGGGACCCGAAAGATGGTGAACTATGCCTGAGCGGGGTAAAGCCAGAGGAAACTCTGGTGGAAGCCCGC | 12 (0.000230%) | | | CTGATGCGGGACGGAAGCTGGTCTCCCGTGTGTTACCGCACGCGGTTGGCCTAAATCCGAGCCAAGGACGCCTGGAGCGTACCGACATGCGGTGGTGAAC | 29 (0.000556%) | | | CTGCAGCACGCGCCTAACGGCGTGCCTCGGCATCAGCGTGCTCCGGGCGTCGGCCTGTGGGCTCCCCATTCGACCCGTCTTGAAACACGGACCAAGGAGT | 2045 (0.039192%) | | | CTGCCCTATCAACTTTCGATGGTAGGATAGTGGCCTACCATGGTGGTAACGGGTGACGGAGAATTAGGGTTCGATTCCGGAGAGGGAGCCTGAGAAACGG | 11 (0.000211%) | | | CTGCTAACTAGCTACGTGGAGGCATCCCTTCACGGCCGGCTTCTTAGAGGGACTATGGCCGTTTAGGCCAAGGAAGTTTGAGGCAATAACAGGTCTGTGA | 52 (0.000997%) | | | CTGGCGACGCATCATTCAAATTTCTGCCCTATCAACTTTCGATGGTAGGATAGTGGCCTACCATGGTGGTAACGGGTGACGGAGAATTAGGGTTCGATTC | 24 (0.000460%) | | | CTGGGGCGGCACATCTGTTAAAAGATAACGCAGGTGTCCTAAGATGAGCTCAACGAGAACAGAAATCTCGTGTGGAACAAAAGGGTAAAAGCTCGTTTGA | 13 (0.000249%) | | | CTGGGTGTCACAAATCGTCGTCCCTCACCATCCTTTGCTGATGCGGGACGGAAGCTGGTCTCCCGTGTGTTACCGCACGCGGTTGGCCTAAATCCGAGCC | 870 (0.016673%) | | | CTGGTGCCAGCAGCCGCGGTAATTCCAGCTCCAATAGCGTATATTTAAGTTGTTGCAGTTAAAAAGCTCGTAGTTGAACCTTGGGATGGGTCGGCCGGTC | 66 (0.001265%) | | | CTGTCCCGAAGGTATCTCGCGCTTGTACGGCTTTGGCTCGGATTCGTCCGTCTTCTTTCTTCTTAGCCGAGTACTTCGGTAGATTAGTTGGAACGATTGA | 60 (0.001150%) | | | CTGTCCCGAGTGTGAGCGAGGTGTGAGTGTCGCCCATGGGCATCGACACCTTGCGGCTAGGAACTGGAACGAGACGGGTAGCAAAGATTTCGAGTAGCAC | 21 (0.000402%) | | | CTGTCCCGAGTGTGAGCGAGGTGTGAGTGTCGCCCATGGGCATCGACACCTTGCGGCTAGGAACTGGAACGAGACGGGTGGCAAAGATTTCGAGTAGCAC | 29 (0.000556%) | | | CTGTGCTGGCGACGCATCATTCAAATTTCTGCCCTATCAACTTTCGATGGTAGGATAGTGGCCTACCATGGTGGTAACGGGTGACGGAGAATTAGGGTTC | 23 (0.000441%) | | | CTGTTGCTTTGTTAGTGTAGAAACACTTGTGTAGAATTGGGGATTGTTTTTTTTGGAGTGATTTAGGGGAGGGTCGAATCTTAGCGACAAAGGGCTGAAT | 11 (0.000211%) | | | CTTAAAGGCGTAAGAATTGTATCCTTGTTAGAAGACACAA | 943 (0.007229%) | | | CTTACGGGTTTACTCACCCGTTGACTCGCACACATGTCAGACTCCTTGGTCCGTGTTTCAAGACGGGTCGAATGGGGAGCCCACAGGCCGACGCCCGGAG | 14 (0.000268%) | | | CTTATACTCAATCATACACATGACATCAAGTCATATTCGACTCCAAAACACTAACCAACCTTCTTCTTGCTTCTCAAAGCTTTCATGGTGTAGCCAAAGT | 117 (0.002242%) | | | CTTCAAATTAGGTACGAAACACAGGCCCCGGAACTCATCATCGAGCGTAACATCGCCCGTGAATTAACTGAGAAGGATAGGTGGTAGGTAGTTCGATGCG | 778 (0.014910%) | | | CTTCAACGAGGAATTCCTAGTAAGCGCGAGTCATCAGCTCGCGTTGACTACGTCCCTGCCCTTTGTACACACCGCCCGTCGCTCCTACCGATTGAATGAT | 69 (0.001322%) | | | CTTCCCTTGCCTACATTGTTCCATCGACCAGAGGCTGTTCACCTTGGAGACCTGATGCGGTTATGAGTACGACCGGGCGTGAGCGGCACTCGGTCCTCCG | 57 (0.001092%) | | | CTTCCTTGCGCCTTACGGGTTTACTCACCCGTTGACTCGCACACATGTCAGACTCCTTGGTCCGTGTTTCAAGACGGGTCGAATGGGGAGCCCACAGGCC | 103 (0.001974%) | | | CTTCTGAGAAGGGTTCGAGTGTGAGCATGCCTGTCGGGACCCGAAAGATGGTGAACTATGCCTGAGCGGGGTAAAGCCAGAGGAAACTCTGGTGGAAGCC | 17 (0.000326%) | | | CTTGCCTACATTGTTCCATCGACCAGAGGCTGTTCACCTTGGAGACCTGATGCGGTTATGAGTACGACCGGGCGTGAGCGGCACTCGGTCCTCCGGATTT | 51 (0.000977%) | | | CTTGCGCCTTACGGGTTTACTCACCCGTTGACTCGCACACATGTCAGACTCCTTGGTCCGTGTTTCAAGACGGGTCGAATGGGGAGCCCACAGGCCGACG | 26 (0.000498%) | | | CTTGGAGACCTGATGCGGTTATGAGTACGACCGGGCGTGAGCGGCACTCGGTCCTCCGGATTTTCAAGGGCCGCCGGGGGCGCACCGGACACCACGCGAC | 118 (0.002261%) | | | CTTGGTGATATGAACACAAACGTTCAATATGACAAACCCATGCCAAGTAAAGAGAAAATGAAAACTGGTGATTGTTGCGGAAATCGTCCAGGATTCCTCG | 13 (0.000249%) | | | CTTGTACGGCTTTGGCTCGGATTCGTCCGTCTTCTTTCTTCTTAGCCGAGTACTTCGGTAGATTAGTTGGAACGATTGATGATTTTGAGTTAATTGAACG | 12 (0.000230%) | | | CTTGTCTCAAAGATTAAGCCATGCATGTGTAAGTATGAACGAATTCAGACTGTGAAACTGCGAATGGCTCATTAAATCAGTTATAGTTTGTTTGATGGTA | 49 (0.000939%) | | | CTTGTTAGAAGACACAAAGCCAAAGACTCATATGGACTTTGGCTACACCATGAAAGCTTTGAGAAGCAAGAAGAAGGTTGGTTAGTGTTTTGGAGTCGAA | 111 (0.002127%) | | | CTTTCACGGTTCGTATTCGTACTGAAAATCAGAATCAAACGAGCTTTTACCCTTTTGTTCCACACGAGATTTCTGTTCTCGTTGAGCTCATCTTAGGACA | 35 (0.000671%) | | | CTTTCGATGGTAGGATAGTGGCCTACCATGGTGGTAACGGGTGACGGAGAATTAGGGTTCGATTCCGGAGAGGGAGCCTGAGAAACGGCTACCACATCCA | 8 (0.000153%) | | | CTTTGCTGATGCGGGACGGAAGCTGGTCTCCCGTGTGTTACCGCACGCGGTTGGCCTAAATCCGAGCCAAGGACGCCTGGAGCGTACCGACATGCGGTGG | 73 (0.001399%) | | | CTTTGGCTCGGATTCGTCCGTCTTCTTTCTTCTTAGCCGAGTACTTCGGTAGATTAGTTGGAACGATTGATGATTTTGAGTTAATTGAACGTTCGGCGTA | 9 (0.000172%) | | | CTTTGTTTTAATTAAACAGTCGGATTCCCCTTGTCCGTACCAGTTCTGAGCTGACTGTTCGACGCCCGGGGAAAGCTCCCGAGAGAGCCGTTCCCAGTCC | 81 (0.001552%) | | | GAAACACAGGCCCCGGAACTCATCATCGAGCGTAACATCGCCCGTGAATTAACTGAGAAGGATAGGTGGTAGGTAGTTCGATGCGCGAGCATGGAGCCTA | 76 (0.001457%) | | | GAAATCCTATGATGTTATCCCATGCTAATGTATCCAGAGCGTAGGCTTGCTTTGAGCACTCTAATTTCTTCAAAGTAACAGCGCCGGAGGCACGACCCGG | 191 (0.003660%) | | | GAACACAAACGTTCAATATGACAAACCCATGCCAAGTAAAGAGAAAATGAAAACTGGTGATTGTTGCGGAAATCGTCCAGGATTCCTCGACCAGGACTTG | 6 (0.000115%) | | | GAACCCACAAAGGGTGTTGGTCGATTAAGACAGCAGGACGGTGGTCATGGAAGTCGAAATCCGCTAAGGAGTGTGTAACAACTCACCTGCCGAATCAACT | 898 (0.017210%) | | | GAACCGGGACGTGGCGGTTGACGGCAACGTTAGGGAGTCCGGAGACGTCGGCGGGGGCCTCGGGAAGAGTTATCTTTTCTGTTTAACAGCCTGCCCACCC | 12 (0.000230%) | | | GAACTACCTACCACCTATCCTTCTCAGTTAATTCACGGGCGATGTTACGCTCGATGATGAGTTCCGGGGCCTGTGTTTCGTACCTAATTTGAAGGAATTG | 5 (0.000096%) | | | GAAGACACAAAGCCAAAGACTCATATGGACTTTGGCTACACCATGAAAGCTTTGAGAAGCAAGAAGAAGGTTGGTTAGTGTTTTGGAGTCGAATATGACT | 35 (0.000671%) | | | GAAGCACGCCCATCCAACCTAGGCGAGACAAGGGTTCACATTTCGTTCATCACCCTTGGCCGGCTATCGAACAGCCGGACTCCCATCAAAAGATGGTTGC | 15 (0.000287%) | | | GAAGGGTTCGAGTGTGAGCATGCCTGTCGGGACCCGAAAGATGGTGAACTATGCCTGAGCGGGGTAAAGCCAGAGGAAACTCTGGTGGAAGCCCGCAGCG | 5 (0.000096%) | | | GAAGGTATCTCGCGCTTGTACGGCTTTGGCTCGGATTCGTCCGTCTTCTTTCTTCTTAGCCGAGTACTTCGGTAGATTAGTTGGAACGATTGATGATTTT | 11 (0.000211%) | | | GAAGTCGAAATCCGCTAAGGAGTGTGTAACAACTCACCTGCCGAATCAACTAGCCCCGAAAATGGATGGCGCTTAAGCGCGCGACCTATACCCGGCCGTC | 3 (0.000057%) | | | GAATCATCAGAGCAACGGGCAGAGCCCGCGTCGACCTTTTATCTAATAAATGCGTCCCTTCCATAAGTCGGGGTTTGTTGCACGTATTAGCTCTAGAATT | 97 (0.001859%) | | | GAATTGTATCCTTGTTAGAAGACACAAAGCCAAAGACTCATATGGACTTTGGCTACACCATGAAAGCTTTGAGAAGCAAGAAGAAGGTTGGTTAGTGTTT | 16 (0.000307%) | | | GACAAGGGTTCACATTTCGTTCATCACCCTTGGCCGGCTATCGAACAGCCGGACTCCCATCAAAAGATGGTTGCCAAGAACATCTTCGTTACGGTTTGCT | 6 (0.000115%) | | | GACAAGGGTTCACATTTCGTTCATCACCCTTGGCCGGCTTTCGAACAGCCGGACTCCCATCAAAAGATGGTTGCCAAGAACATCTTCGTTACGGTTTGCT | 14 (0.000268%) | | | GACACAAAGCCAAAGACTCATATGGACTTTGGCTACACCATGAAAGCTTTGAGAAGCAAGAAGAAGGTTGGTTAGTGTTTTGGAGTCGAATATGACTTGA | 51 (0.000977%) | | | GACAGCAGGACGGTGGTCATGGAAGTCGAAATCCGCTAAGGAGTGTGTAACAACTCACCTGCCGAATCAACTAGCCCCGAAAATGGATGGCGCTTAAGCG | 6 (0.000115%) | | | GACATCAAGTCATATTCGACTCCAAAACACTAACCAACCTTCTTCTTGCTTCTCAAAGCTTTCATGGTGTAGCCAAAGTCCATATGAGTCTTTGGCTTTG | 45 (0.000862%) | | | GACATTGTCAGGTGGGGAGTTTGGCTGGGGCGGCACATCTGTTAAAAGATAACGCAGGTGTCCTAAGATGAGCTCAACGAGAACAGAAATCTCGTGTGGA | 32 (0.000613%) | | | GACCAGGGGTTGAAATCGTCGACCAGGTCCGAGACTTCATCGACCGGGTCCGAGGATTCGTCGACCAGGACGGCCGGATGTCCGAGAAAAAAAAATGTTG | 10 (0.000192%) | | | GACCCGCCGAAGCGAGCCTTGGGACCAAAAACAGGGGTTGTACCCCGCCTCCGATTCACGGAGTAAGTAAAATAACGTTAAAAGTAGTGGTATTTCACTT | 6 (0.000115%) | | | GACCTCAGCCTGCTAACTAGCTACGTGGAGGCATCCCTTCACGGCCGGCTTCTTAGAGGGACTATGGCCGTTTAGGCCAAGGAAGTTTGAGGCAATAACA | 8 (0.000153%) | | | GACGCATCATTCAAATTTCTGCCCTATCAACTTTCGATGGTAGGATAGTGGCCTACCATGGTGGTAACGGGTGACGGAGAATTAGGGTTCGATTCCGGAG | 15 (0.000287%) | | | GACGGGCGGTGTGTACAAAGGGCAGGGACGTAGTCAACGCGAGCTGATGACTCGCGCTTACTAGGAATTCCTCGTTGAAGACCAACAATTGCAATGATCG | 12 (0.000230%) | | | GACGGTGGTCATGGAAGTCGAAATCCGCTAAGGAGTGTGTAACAACTCACCTGCCGAATCAACTAGCCCCGAAAATGGATGGCGCTTAAGCGCGCGACCT | 3 (0.000057%) | | | GACGTCGGCGGGGGCCTCGGGAAGAGTTATCTTTTCTGTTTAACAGCCTGCCCACCCTGGAAACGGCTCAGCCGGAGGTAGGGTCCAGCGGCTGGAAGAG | 4 (0.000077%) | | | GACGTGGGTGGTTCGCCGCCCGCGACGTCGCGAGAAGTCCACTAAACCTTATCATTTAGAGGAAGGAGAAGTCGTAACAAGGTTTCCGTAGGTGAACCTG | 42 (0.000805%) | | | GACTCCAAAACACTAACCAACCTTCTTCTTGCTTCTCAAAGCTTTCATGGTGTAGCCAAAGTCCATATGAGTCTTTGGCTTTGTGTCTTCTAACAAGGAT | 25 (0.000479%) | | | GACTTCCCTTGCCTACATTGTTCCATCGACCAGAGGCTGTTCACCTTGGAGACCTGATGCGGTTATGAGTACGACCGGGCGTGAGCGGCACTCGGTCCTC | 24 (0.000460%) | | | GAGAAGGGTTCGAGTGTGAGCATGCCTGTCGGGACCCGAAAGATGGTGAACTATGCCTGAGCGGGGTAAAGCCAGAGGAAACTCTGGTGGAAGCCCGCAG | 7 (0.000134%) | | | GAGACAAGGGTTCACATTTCGTTCATCACCCTTGGCCGGCTTTCGAACAGCCGGACTCCCATCAAAAGATGGTTGCCAAGAACATCTTCGTTACGGTTTG | 6 (0.000115%) | | | GAGACCTCAGCCTGCTAACTAGCTACGTGGAGGCATCCCTTCACGGCCGGCTTCTTAGAGGGACTATGGCCGTTTAGGCCAAGGAAGTTTGAGGCAATAA | 934 (0.017900%) | | | GAGACGTCGGCGGGGGCCTCGGGAAGAGTTATCTTTTCTGTTTAACAGCCTGCCCACCCTGGAAACGGCTCAGCCGGAGGTAGGGTCCAGCGGCTGGAAG | 15 (0.000287%) | | | GAGCAACGGGCAGAGCCCGCGTCGACCTTTTATCTAATAAATGCGTCCCTTCCATAAGTCGGGGTTTGTTGCACGTATTAGCTCTAGAATTACTACGGTT | 14 (0.000268%) | | | GAGCCCGCGTCGACCTTTTATCTAATAAATGCGTCCCTTCCATAAGTCGGGGTTTGTTGCACGTATTAGCTCTAGAATTACTACGGTTATCCGAGTAGTA | 8 (0.000153%) | | | GAGCGACGGGCGGTGTGTACAAAGGGCAGGGACGTAGTCAACGCGAGCTGATGACTCGCGCTTACTAGGAATTCCTCGTTGAAGACCAACAATTGCAATG | 12 (0.000230%) | | | GAGCGAGGTGTGAGTGTCGCCCATGGGCATCGACACCTTGCGGCTAGGAACTGGAACGAGACGGGTGGCAAAGATTTCGAGTAGCACTTCATACTACCGT | 9 (0.000172%) | | | GAGCGGCACTCGGTCCTCCGGATTTTCAAGGGCCGCCGGGGGCGCACCGGACACCACGCGACGTGCGGTGCTCTTCCAGCCGCTGGACCCTACCTCCGGC | 14 (0.000268%) | | | GAGCTCGTGTAAAGTTGGGAATTCGTTAAGGAGCTGTTGCTTTGTTAGTGTAGAAACACTTGTGTAGAATTGGGGATTGTTTTTTTTGGAGTGATTTAGG | 10 (0.000192%) | | | GAGCTGTTGCTTTGTTAGTGTAGAAACACTTGTGTAGAATTGGGGATTGTTTTTTTTGGAGTGATTTAGGGGAGGGTCGAATCTTAGCGACAAAGGGCTG | 5 (0.000096%) | | | GAGGACATTGTCAGGTGGGGAGTTTGGCTGGGGCGGCACATCTGTTAAAAGATAACGCAGGTGTCCTAAGATGAGCTCAACGAGAACAGAAATCTCGTGT | 58 (0.001112%) | | | GAGGCTGTTCACCTTGGAGACCTGATGCGGTTATGAGTACGACCGGGCGTGAGCGGCACTCGGTCCTCCGGATTTTCAAGGGCCGCCGGGGGCGCACCGG | 10 (0.000192%) | | | GAGGGCAAGTCTGGTGCCAGCAGCCGCGGTAATTCCAGCTCCAATAGCGTATATTTAAGTTGTTGCAGTTAAAAAGCTCGTAGTTGAACCTTGGGATGGG | 22 (0.000422%) | | | GAGGTGTGAGTGTCGCCCATGGGCATCGACACCTTGCGGCTAGGAACTGGAACGAGACGGGTAGCAAAGATTTCGAGTAGCACTTCATACTACCGTGGGT | 66 (0.001265%) | | | GAGGTGTGAGTGTCGCCCATGGGCATCGACACCTTGCGGCTAGGAACTGGAACGAGACGGGTGGCAAAGATTTCGAGTAGCACTTCATACTACCGTGGGT | 11 (0.000211%) | | | GAGTATAAGAACTTAAACCGCAACCGCATCTTATAAGCCTAAGTAGTGTTTCCTTGTTAGAAGACACAAAGCCAAAGACTCATATGGACTTTGGCTACAC | 6 (0.000115%) | | | GAGTCGAATATGACTTGATGTCATGTGTATGATTGAGTAT | 1382 (0.010594%) | | | GAGTGTCGCCCATGGGCATCGACACCTTGCGGCTAGGAACTGGAACGAGACGGGTGGCAAAGATTTCGAGTAGCACTTCATACTACCGTGGGTTTTTTAA | 4 (0.000077%) | | | GAGTGTGAGCGAGGTGTGAGTGTCGCCCATGGGCATCGACACCTTGCGGCTAGGAACTGGAACGAGACGGGTAGCAAAGATTTCGAGTAGCACTTCATAC | 3 (0.000057%) | | | GAGTTATCATGAATCATCAGAGCAACGGGCAGAGCCCGCGTCGACCTTTTATCTAATAAATGCGTCCCTTCCATAAGTCGGGGTTTGTTGCACGTATTAG | 84 (0.001610%) | | | GAGTTTGGCTGGGGCGGCACATCTGTTAAAAGATAACGCAGGTGTCCTAAGATGAGCTCAACGAGAACAGAAATCTCGTGTGGAACAAAAGGGTAAAAGC | 4 (0.000077%) | | | GATAGCTAGTGTTCGTAGGCTCCATGCTCGCGCATCGAACTACCTACCACCTATCCTTCTCAGTTAATTCACGGGCGATGTTACGCTCGATGATGAGTTC | 13 (0.000249%) | | | GATCCATTACATTTTATCGGTCGCTCTTGTCCGGAAGCTGTAGATGACCCAAAGTCCATATAGCGACCCCAGGTCAGGCGGGATTACCCGCTGAGTTTAA | 12 (0.000230%) | | | GATCCCGCCAATCAGCTTCCTTGCGCCTTACGGGTTTACTCACCCGTTGACTCGCACACATGTCAGACTCCTTGGTCCGTGTTTCAAGACGGGTCGAATG | 523 (0.010023%) | | | GATCCGTCGAGTTATCATGAATCATCAGAGCAACGGGCAGAGCCCGCGTCGACCTTTTATCTAATAAATGCGTCCCTTCCATAAGTCGGGGTTTGTTGCA | 131 (0.002511%) | | | GATCGAAATCCTATGATGTTATCCCATGCTAATGTATCCAGAGCGTAGGCTTGCTTTGAGCACTCTAATTTCTTCAAAGTAACAGCGCCGGAGGCACGAC | 1612 (0.030893%) | | | GATCGACCCGCCGAAGCGAGCCTTGGGACCAAAAACAGGGGTTGTACCCCGCCTCCGATTCACGGAGTAAGTAAAATAACGTTAAAAGTAGTGGTATTTC | 41 (0.000786%) | | | GATCGATCAA | 168754 (0.323410%) | | | GATCGATCCATTACATTTTATCGGTCGCTCTTGTCCGGAAGCTGTAGATGACCCAAAGTCCATATAGCGACCCCAGGTCAGGCGGGATTACCCGCTGAGT | 1202 (0.023036%) | | | GATCGATCCGTAACTTCGGGAAAAGGATTGGCTCTGAGGGCTGGGCTCGGGGGTCCCAGTTCCGAACCCGTCGGCTGTCAGCGGACTGCTCGAGCTGCTT | 1224 (0.023457%) | | | GATCGATCCGTCGAGTTATCATGAATCATCAGAGCAACGGGCAGAGCCCGCGTCGACCTTTTATCTAATAAATGCGTCCCTTCCATAAGTCGGGGTTTGT | 1215 (0.023285%) | | | GATCGATCGACCCGCCGAAGCGAGCCTTGGGACCAAAAACAGGGGTTGTACCCCGCCTCCGATTCACGGAGTAAGTAAAATAACGTTAAAAGTAGTGGTA | 989 (0.018954%) | | | GATCGATCTCATGTGTATGATTGAGTATAAGAACTTAAACCGCAACCGCATCTTATAAGCCTAAGTAGTGTTTCCTTGTTAGAAGACACAAAGCCAAAGA | 233 (0.004465%) | | | GATCGATCTT | 187116 (0.358600%) | | | GATCGATCTTAAAGGCGTAAGAATTGTATCCTTGTTAAAAGACACAAAGCCAAAGACTCATATGGACTTTGGCTACACCATGAAAGCTTTGAGAAGCAAG | 467 (0.008950%) | | | GATCGATCTTAAAGGCGTAAGAATTGTATCCTTGTTAGAA | 4228 (0.032411%) | | | GATCGATCTTCTGAGAAGGGTTCGAGTGTGAGCATGCCTGTCGGGACCCGAAAGATGGTGAACTATGCCTGAGCGGGGTAAAGCCAGAGGAAACTCTGGT | 1125 (0.021560%) | | | GATCTCATGTGTATGATTGAGTATAAGAACTTAAACCGCAACCGCATCTTATAAGCCTAAGTAGTGTTTCCTTGTTAGAAGACACAAAGCCAAAGACTCA | 49 (0.000939%) | | | GATCTTAAAGGCGTAAGAATTGTATCCTTGTTAAAAGACACAAAGCCAAAGACTCATATGGACTTTGGCTACACCATGAAAGCTTTGAGAAGCAAGAAGA | 253 (0.004849%) | | | GATCTTCTGAGAAGGGTTCGAGTGTGAGCATGCCTGTCGGGACCCGAAAGATGGTGAACTATGCCTGAGCGGGGTAAAGCCAGAGGAAACTCTGGTGGAA | 134 (0.002568%) | | | GATGTTATCCCATGCTAATGTATCCAGAGCGTAGGCTTGCTTTGAGCACTCTAATTTCTTCAAAGTAACAGCGCCGGAGGCACGACCCGGCCAATTAAGA | 17 (0.000326%) | | | GATTAACAGGGACAGTCGGGGGCATTCGTATTTCATAGTCAGAGGTGAAATTCTTGGATTTATGAAAGACGAACAACTGCGAAAGCATTTGCCAAGGATG | 150 (0.002875%) | | | GATTAAGACAGCAGGACGGTGGTCATGGAAGTCGAAATCCGCTAAGGAGTGTGTAACAACTCACCTGCCGAATCAACTAGCCCCGAAAATGGATGGCGCT | 37 (0.000709%) | | | GATTAGTCTTTCGCCCCTATACCCAAGTCAGACGAACGATTTGCACGTCAGTATCGCTGCGGGCTTCCACCAGAGTTTCCTCTGGCTTTACCCCGCTCAG | 21 (0.000402%) | | | GATTGAGTATAAGAACTTAAACCGCAACCGCATCTTATAAGCCTAAGTAGTGTTTCCTTGTTAGAAGACACAAAGCCAAAGACTCATATGGACTTTGGCT | 35 (0.000671%) | | | GATTTGCACGTCAGTATCGCTGCGGGCTTCCACCAGAGTTTCCTCTGGCTTTACCCCGCTCAGGCATAGTTCACCATCTTTCGGGTCCCGACAGGCATGC | 29 (0.000556%) | | | GATTTTCAAGGGCCGCCGGGGGCGCACCGGACACCACGCGACGTGCGGTGCTCTTCCAGCCGCTGGACCCTACCTCCGGCTGAGCCGTTTCCAGGGTGGG | 9 (0.000172%) | | | GCAACGGGCAGAGCCCGCGTCGACCTTTTATCTAATAAATGCGTCCCTTCCATAAGTCGGGGTTTGTTGCACGTATTAGCTCTAGAATTACTACGGTTAT | 34 (0.000652%) | | | GCAACGTTAGGGAGTCCGGAGACGTCGGCGGGGGCCTCGGGAAGAGTTATCTTTTCTGTTTAACAGCCTGCCCACCCTGGAAACGGCTCAGCCGGAGGTA | 18 (0.000345%) | | | GCAAGTCTGGTGCCAGCAGCCGCGGTAATTCCAGCTCCAATAGCGTATATTTAAGTTGTTGCAGTTAAAAAGCTCGTAGTTGAACCTTGGGATGGGTCGG | 26 (0.000498%) | | | GCACCGGACACCACGCGACGTGCGGTGCTCTTCCAGCCGCTGGACCCTACCTCCGGCTGAGCCGTTTCCAGGGTGGGCAGGCTGTTAAACAGAAAAGATA | 22 (0.000422%) | | | GCACGCCCATCCAACCTAGGCGAGACAAGGGTTCACATTTCGTTCATCACCCTTGGCCGGCTATCGAACAGCCGGACTCCCATCAAAAGATGGTTGCCAA | 38 (0.000728%) | | | GCACGCGCCTAACGGCGTGCCTCGGCATCAGCGTGCTCCGGGCGTCGGCCTGTGGGCTCCCCATTCGACCCGTCTTGAAACACGGACCAAGGAGTCTGAC | 22 (0.000422%) | | | GCACTCGGTCCTCCGGATTTTCAAGGGCCGCCGGGGGCGCACCGGACACCACGCGACGTGCGGTGCTCTTCCAGCCGCTGGACCCTACCTCCGGCTGAGC | 6 (0.000115%) | | | GCAGAGCCCGCGTCGACCTTTTATCTAATAAATGCGTCCCTTCCATAAGTCGGGGTTTGTTGCACGTATTAGCTCTAGAATTACTACGGTTATCCGAGTA | 20 (0.000383%) | | | GCAGCACGCGCCTAACGGCGTGCCTCGGCATCAGCGTGCTCCGGGCGTCGGCCTGTGGGCTCCCCATTCGACCCGTCTTGAAACACGGACCAAGGAGTCT | 419 (0.008030%) | | | GCAGCCGCGGTAATTCCAGCTCCAATAGCGTATATTTAAGTTGTTGCAGTTAAAAAGCTCGTAGTTGAACCTTGGGATGGGTCGGCCGGTCCGCCTTTGG | 5 (0.000096%) | | | GCAGGACGGTGGTCATGGAAGTCGAAATCCGCTAAGGAGTGTGTAACAACTCACCTGCCGAATCAACTAGCCCCGAAAATGGATGGCGCTTAAGCGCGCG | 46 (0.000882%) | | | GCATCAGCGTGCTCCGGGCGTCGGCCTGTGGGCTCCCCATTCGACCCGTCTTGAAACACGGACCAAGGAGTCTGACATGTGTGCGAGTCAACGGGTGAGT | 7 (0.000134%) | | | GCATCATTCAAATTTCTGCCCTATCAACTTTCGATGGTAGGATAGTGGCCTACCATGGTGGTAACGGGTGACGGAGAATTAGGGTTCGATTCCGGAGAGG | 45 (0.000862%) | | | GCATCGACACCTTGCGGCTAGGAACTGGAACGAGACGGGTGGCAAAGATTTCGAGTAGCACTTCATACTACCGTGGGTTTTTTAAACCTTCCGAGTTTTG | 7 (0.000134%) | | | GCATGGCCTCTGTGCTGGCGACGCATCATTCAAATTTCTGCCCTATCAACTTTCGATGGTAGGATAGTGGCCTACCATGGTGGTAACGGGTGACGGAGAA | 1617 (0.030989%) | | | GCATTCGTATTTCATAGTCAGAGGTGAAATTCTTGGATTTATGAAAGACGAACAACTGCGAAAGCATTTGCCAAGGATGTTTTCATTAATCAAGAACGAA | 3 (0.000057%) | | | GCCAAAGACTCATATGGACTTTGGCTACACCATGAAAGCTTTGAGAAGCAAGAAGAAGGTTGGTTAGTGTTTTGGAGTCGAATATGACTTGATGTCATGT | 51 (0.000977%) | | | GCCAATCAGCTTCCTTGCGCCTTACGGGTTTACTCACCCGTTGACTCGCACACATGTCAGACTCCTTGGTCCGTGTTTCAAGACGGGTCGAATGGGGAGC | 128 (0.002453%) | | | GCCAGCAGCCGCGGTAATTCCAGCTCCAATAGCGTATATTTAAGTTGTTGCAGTTAAAAAGCTCGTAGTTGAACCTTGGGATGGGTCGGCCGGTCCGCCT | 25 (0.000479%) | | | GCCCATCCAACCTAGGCGAGACAAGGGTTCACATTTCGTTCATCACCCTTGGCCGGCTATCGAACAGCCGGACTCCCATCAAAAGATGGTTGCCAAGAAC | 60 (0.001150%) | | | GCCCATCCAACCTAGGCGAGACAAGGGTTCACATTTCGTTCATCACCCTTGGCCGGCTTTCGAACAGCCGGACTCCCATCAAAAGATGGTTGCCAAGAAC | 34 (0.000652%) | | | GCCCATGGGCATCGACACCTTGCGGCTAGGAACTGGAACGAGACGGGTGGCAAAGATTTCGAGTAGCACTTCATACTACCGTGGGTTTTTTAAACCTTCC | 8 (0.000153%) | | | GCCCTATCAACTTTCGATGGTAGGATAGTGGCCTACCATGGTGGTAACGGGTGACGGAGAATTAGGGTTCGATTCCGGAGAGGGAGCCTGAGAAACGGCT | 23 (0.000441%) | | | GCCGAAGCGAGCCTTGGGACCAAAAACAGGGGTTGTACCCCGCCTCCGATTCACGGAGTAAGTAAAATAACGTTAAAAGTAGTGGTATTTCACTTGCGCC | 29 (0.000556%) | | | GCCGACTTCCCTTGCCTACATTGTTCCATCGACCAGAGGCTGTTCACCTTGGAGACCTGATGCGGTTATGAGTACGACCGGGCGTGAGCGGCACTCGGTC | 204 (0.003910%) | | | GCCGCCCGCGACGTCGCGAGAAGTCCACTAAACCTTATCATTTAGAGGAAGGAGAAGTCGTAACAAGGTTTCCGTAGGTGAACCTGCGGAAGGATCGATC | 23 (0.000441%) | | | GCCGCCGGGGGCGCACCGGACACCACGCGACGTGCGGTGCTCTTCCAGCCGCTGGACCCTACCTCCGGCTGAGCCGTTTCCAGGGTGGGCAGGCTGTTAA | 9 (0.000172%) | | | GCCGCGGTAATTCCAGCTCCAATAGCGTATATTTAAGTTGTTGCAGTTAAAAAGCTCGTAGTTGAACCTTGGGATGGGTCGGCCGGTCCGCCTTTGGTGT | 11 (0.000211%) | | | GCCGGGGGCGCACCGGACACCACGCGACGTGCGGTGCTCTTCCAGCCGCTGGACCCTACCTCCGGCTGAGCCGTTTCCAGGGTGGGCAGGCTGTTAAACA | 17 (0.000326%) | | | GCCTAACGGCGTGCCTCGGCATCAGCGTGCTCCGGGCGTCGGCCTGTGGGCTCCCCATTCGACCCGTCTTGAAACACGGACCAAGGAGTCTGACATGTGT | 34 (0.000652%) | | | GCCTAAGTAGTGTTTCCTTGTTAGAACATACAAAGCCAAAGACTCATACGGACTTTGGCTACACCATGAAAGCTTTGAGAAGCTAGAAGAAGGTTGGTTA | 54 (0.001035%) | | | GCCTAAGTAGTGTTTCCTTGTTAGAAGACACAAAGCCAAAGACTCATATGGACTTTGGCTACACCATGAAAGCTTTGAGAAGCAAGAAGAAGGTTGGTTA | 34 (0.000652%) | | | GCCTACATTGTTCCATCGACCAGAGGCTGTTCACCTTGGAGACCTGATGCGGTTATGAGTACGACCGGGCGTGAGCGGCACTCGGTCCTCCGGATTTTCA | 3 (0.000057%) | | | GCCTAGGCTGTCCCGAGTGTGAGCGAGGTGTGAGTGTCGCCCATGGGCATCGACACCTTGCGGCTAGGAACTGGAACGAGACGGGTGGCAAAGATTTCGA | 358 (0.006861%) | | | GCCTCGGCATCAGCGTGCTCCGGGCGTCGGCCTGTGGGCTCCCCATTCGACCCGTCTTGAAACACGGACCAAGGAGTCTGACATGTGTGCGAGTCAACGG | 17 (0.000326%) | | | GCCTCTGTGCTGGCGACGCATCATTCAAATTTCTGCCCTATCAACTTTCGATGGTAGGATAGTGGCCTACCATGGTGGTAACGGGTGACGGAGAATTAGG | 13 (0.000249%) | | | GCCTGCTAACTAGCTACGTGGAGGCATCCCTTCACGGCCGGCTTCTTAGAGGGACTATGGCCGTTTAGGCCAAGGAAGTTTGAGGCAATAACAGGTCTGT | 5 (0.000096%) | | | GCCTTACGGGTTTACTCACCCGTTGACTCGCACACATGTCAGACTCCTTGGTCCGTGTTTCAAGACGGGTCGAATGGGGAGCCCACAGGCCGACGCCCGG | 31 (0.000594%) | | | GCGACGCATCATTCAAATTTCTGCCCTATCAACTTTCGATGGTAGGATAGTGGCCTACCATGGTGGTAACGGGTGACGGAGAATTAGGGTTCGATTCCGG | 35 (0.000671%) | | | GCGACGGGCGGTGTGTACAAAGGGCAGGGACGTAGTCAACGCGAGCTGATGACTCGCGCTTACTAGGAATTCCTCGTTGAAGACCAACAATTGCAATGAT | 46 (0.000882%) | | | GCGACGTGGGTGGTTCGCCGCCCGCGACGTCGCGAGAAGTCCACTAAACCTTATCATTTAGAGGAAGGAGAAGTCGTAACAAGGTTTCCGTAGGTGAACC | 347 (0.006650%) | | | GCGAGACAAGGGTTCACATTTCGTTCATCACCCTTGGCCGGCTATCGAACAGCCGGACTCCCATCAAAAGATGGTTGCCAAGAACATCTTCGTTACGGTT | 29 (0.000556%) | | | GCGAGACAAGGGTTCACATTTCGTTCATCACCCTTGGCCGGCTTTCGAACAGCCGGACTCCCATCAAAAGATGGTTGCCAAGAACATCTTCGTTACGGTT | 39 (0.000747%) | | | GCGAGGTGTGAGTGTCGCCCATGGGCATCGACACCTTGCGGCTAGGAACTGGAACGAGACGGGTGGCAAAGATTTCGAGTAGCACTTCATACTACCGTGG | 48 (0.000920%) | | | GCGCACCGGACACCACGCGACGTGCGGTGCTCTTCCAGCCGCTGGACCCTACCTCCGGCTGAGCCGTTTCCAGGGTGGGCAGGCTGTTAAACAGAAAAGA | 27 (0.000517%) | | | GCGCATCGAACTACCTACCACCTATCCTTCTCAGTTAATTCACGGGCGATGTTACGCTCGATGATGAGTTCCGGGGCCTGTGTTTCGTACCTAATTTGAA | 5 (0.000096%) | | | GCGCCTAACGGCGTGCCTCGGCATCAGCGTGCTCCGGGCGTCGGCCTGTGGGCTCCCCATTCGACCCGTCTTGAAACACGGACCAAGGAGTCTGACATGT | 15 (0.000287%) | | | GCGCCTTACGGGTTTACTCACCCGTTGACTCGCACACATGTCAGACTCCTTGGTCCGTGTTTCAAGACGGGTCGAATGGGGAGCCCACAGGCCGACGCCC | 20 (0.000383%) | | | GCGCTTGTACGGCTTTGGCTCGGATTCGTCCGTCTTCTTTCTTCTTAGCCGAGTACTTCGGTAGATTAGTTGGAACGATTGATGATTTTGAGTTAATTGA | 10 (0.000192%) | | | GCGGAGGACATTGTCAGGTGGGGAGTTTGGCTGGGGCGGCACATCTGTTAAAAGATAACGCAGGTGTCCTAAGATGAGCTCAACGAGAACAGAAATCTCG | 1475 (0.028268%) | | | GCGGCACATCTGTTAAAAGATAACGCAGGTGTCCTAAGATGAGCTCAACGAGAACAGAAATCTCGTGTGGAACAAAAGGGTAAAAGCTCGTTTGATTCTG | 8 (0.000153%) | | | GCGGCACTCGGTCCTCCGGATTTTCAAGGGCCGCCGGGGGCGCACCGGACACCACGCGACGTGCGGTGCTCTTCCAGCCGCTGGACCCTACCTCCGGCTG | 60 (0.001150%) | | | GCGGCGACGTGGGTGGTTCGCCGCCCGCGACGTCGCGAGAAGTCCACTAAACCTTATCATTTAGAGGAAGGAGAAGTCGTAACAAGGTTTCCGTAGGTGA | 1261 (0.024167%) | | | GCGGTAATTCCAGCTCCAATAGCGTATATTTAAGTTGTTGCAGTTAAAAAGCTCGTAGTTGAACCTTGGGATGGGTCGGCCGGTCCGCCTTTGGTGTGCA | 18 (0.000345%) | | | GCGGTTGACGGCAACGTTAGGGAGTCCGGAGACGTCGGCGGGGGCCTCGGGAAGAGTTATCTTTTCTGTTTAACAGCCTGCCCACCCTGGAAACGGCTCA | 10 (0.000192%) | | | GCGGTTTAAGTTCTTATACTCAATCATACACATGACATCAAGTCATATTCGACTCCAAAACACTAACCAACCTTCTTCTTGCTTCTCAAAGCTTTCATGG | 139 (0.002664%) | | | GCGGTTTAAGTTGTTATACTCAATCATACACATGACAACAAGTCATATTCGACTCCAAAACACTAACCAACCTTCTTCTTGCTTCTCAAAGCTTTCATGG | 45 (0.000862%) | | | GCGTAAGAATTGTATCCTTGTTAGAAGACACAAAGCCAAA | 170 (0.001303%) | | | GCGTGCCTCGGCATCAGCGTGCTCCGGGCGTCGGCCTGTGGGCTCCCCATTCGACCCGTCTTGAAACACGGACCAAGGAGTCTGACATGTGTGCGAGTCA | 10 (0.000192%) | | | GCGTGCTCCGGGCGTCGGCCTGTGGGCTCCCCATTCGACCCGTCTTGAAACACGGACCAAGGAGTCTGACATGTGTGCGAGTCAACGGGTGAGTAAACCC | 11 (0.000211%) | | | GCTAACTAGCTACGTGGAGGCATCCCTTCACGGCCGGCTTCTTAGAGGGACTATGGCCGTTTAGGCCAAGGAAGTTTGAGGCAATAACAGGTCTGTGATG | 31 (0.000594%) | | | GCTAATGTATCCAGAGCGTAGGCTTGCTTTGAGCACTCTAATTTCTTCAAAGTAACAGCGCCGGAGGCACGACCCGGCCAATTAAGACCAGGAGCGTATC | 19 (0.000364%) | | | GCTAGTGTTCGTAGGCTCCATGCTCGCGCATCGAACTACCTACCACCTATCCTTCTCAGTTAATTCACGGGCGATGTTACGCTCGATGATGAGTTCCGGG | 20 (0.000383%) | | | GCTCCAATAGCGTATATTTAAGTTGTTGCAGTTAAAAAGCTCGTAGTTGAACCTTGGGATGGGTCGGCCGGTCCGCCTTTGGTGTGCATTGGTCGGCTTG | 13 (0.000249%) | | | GCTCCATGCTCGCGCATCGAACTACCTACCACCTATCCTTCTCAGTTAATTCACGGGCGATGTTACGCTCGATGATGAGTTCCGGGGCCTGTGTTTCGTA | 4 (0.000077%) | | | GCTCCGGGCGTCGGCCTGTGGGCTCCCCATTCGACCCGTCTTGAAACACGGACCAAGGAGTCTGACATGTGTGCGAGTCAACGGGTGAGTAAACCCGTAA | 10 (0.000192%) | | | GCTCGCGCATCGAACTACCTACCACCTATCCTTCTCAGTTAATTCACGGGCGATGTTACGCTCGATGATGAGTTCCGGGGCCTGTGTTTCGTACCTAATT | 6 (0.000115%) | | | GCTCGTGTAAAGTTGGGAATTCGTTAAGGAGCTGTTGCTTTGTTAGTGTAGAAACACTTGTGTAGAATTGGGGATTGTTTTTTTTGGAGTGATTTAGGGG | 5 (0.000096%) | | | GCTGATGCGGGACGGAAGCTGGTCTCCCGTGTGTTACCGCACGCGGTTGGCCTAAATCCGAGCCAAGGACGCCTGGAGCGTACCGACATGCGGTGGTGAA | 6 (0.000115%) | | | GCTGGCGACGCATCATTCAAATTTCTGCCCTATCAACTTTCGATGGTAGGATAGTGGCCTACCATGGTGGTAACGGGTGACGGAGAATTAGGGTTCGATT | 29 (0.000556%) | | | GCTGGGGCGGCACATCTGTTAAAAGATAACGCAGGTGTCCTAAGATGAGCTCAACGAGAACAGAAATCTCGTGTGGAACAAAAGGGTAAAAGCTCGTTTG | 9 (0.000172%) | | | GCTGTCCCGAAGGTATCTCGCGCTTGTACGGCTTTGGCTCGGATTCGTCCGTCTTCTTTCTTCTTAGCCGAGTACTTCGGTAGATTAGTTGGAACGATTG | 55 (0.001054%) | | | GCTGTCCCGAGTGTGAGCGAGGTGTGAGTGTCGCCCATGGGCATCGACACCTTGCGGCTAGGAACTGGAACGAGACGGGTAGCAAAGATTTCGAGTAGCA | 66 (0.001265%) | | | GCTGTCCCGAGTGTGAGCGAGGTGTGAGTGTCGCCCATGGGCATCGACACCTTGCGGCTAGGAACTGGAACGAGACGGGTGGCAAAGATTTCGAGTAGCA | 99 (0.001897%) | | | GCTGTTGCTTTGTTAGTGTAGAAACACTTGTGTAGAATTGGGGATTGTTTTTTTTGGAGTGATTTAGGGGAGGGTCGAATCTTAGCGACAAAGGGCTGAA | 5 (0.000096%) | | | GCTTCCTTGCGCCTTACGGGTTTACTCACCCGTTGACTCGCACACATGTCAGACTCCTTGGTCCGTGTTTCAAGACGGGTCGAATGGGGAGCCCACAGGC | 44 (0.000843%) | | | GCTTGTACGGCTTTGGCTCGGATTCGTCCGTCTTCTTTCTTCTTAGCCGAGTACTTCGGTAGATTAGTTGGAACGATTGATGATTTTGAGTTAATTGAAC | 18 (0.000345%) | | | GCTTGTCTCAAAGATTAAGCCATGCATGTGTAAGTATGAACGAATTCAGACTGTGAAACTGCGAATGGCTCATTAAATCAGTTATAGTTTGTTTGATGGT | 20 (0.000383%) | | | GCTTTCACGGTTCGTATTCGTACTGAAAATCAGAATCAAACGAGCTTTTACCCTTTTGTTCCACACGAGATTTCTGTTCTCGTTGAGCTCATCTTAGGAC | 10 (0.000192%) | | | GCTTTGGCTCGGATTCGTCCGTCTTCTTTCTTCTTAGCCGAGTACTTCGGTAGATTAGTTGGAACGATTGATGATTTTGAGTTAATTGAACGTTCGGCGT | 5 (0.000096%) | | | GCTTTGTTTTAATTAAACAGTCGGATTCCCCTTGTCCGTACCAGTTCTGAGCTGACTGTTCGACGCCCGGGGAAAGCTCCCGAGAGAGCCGTTCCCAGTC | 16 (0.000307%) | | | GGAACCGGGACGTGGCGGTTGACGGCAACGTTAGGGAGTCCGGAGACGTCGGCGGGGGCCTCGGGAAGAGTTATCTTTTCTGTTTAACAGCCTGCCCACC | 72 (0.001380%) | | | GGAAGTCGAAATCCGCTAAGGAGTGTGTAACAACTCACCTGCCGAATCAACTAGCCCCGAAAATGGATGGCGCTTAAGCGCGCGACCTATACCCGGCCGT | 24 (0.000460%) | | | GGACAGTCGGGGGCATTCGTATTTCATAGTCAGAGGTGAAATTCTTGGATTTATGAAAGACGAACAACTGCGAAAGCATTTGCCAAGGATGTTTTCATTA | 63 (0.001207%) | | | GGACATTGTCAGGTGGGGAGTTTGGCTGGGGCGGCACATCTGTTAAAAGATAACGCAGGTGTCCTAAGATGAGCTCAACGAGAACAGAAATCTCGTGTGG | 183 (0.003507%) | | | GGACGGTGGTCATGGAAGTCGAAATCCGCTAAGGAGTGTGTAACAACTCACCTGCCGAATCAACTAGCCCCGAAAATGGATGGCGCTTAAGCGCGCGACC | 50 (0.000958%) | | | GGACGTGGCGGTTGACGGCAACGTTAGGGAGTCCGGAGACGTCGGCGGGGGCCTCGGGAAGAGTTATCTTTTCTGTTTAACAGCCTGCCCACCCTGGAAA | 3 (0.000057%) | | | GGAGACGTCGGCGGGGGCCTCGGGAAGAGTTATCTTTTCTGTTTAACAGCCTGCCCACCCTGGAAACGGCTCAGCCGGAGGTAGGGTCCAGCGGCTGGAA | 19 (0.000364%) | | | GGAGCGACGGGCGGTGTGTACAAAGGGCAGGGACGTAGTCAACGCGAGCTGATGACTCGCGCTTACTAGGAATTCCTCGTTGAAGACCAACAATTGCAAT | 54 (0.001035%) | | | GGAGCTGTTGCTTTGTTAGTGTAGAAACACTTGTGTAGAATTGGGGATTGTTTTTTTTGGAGTGATTTAGGGGAGGGTCGAATCTTAGCGACAAAGGGCT | 7 (0.000134%) | | | GGAGGACATTGTCAGGTGGGGAGTTTGGCTGGGGCGGCACATCTGTTAAAAGATAACGCAGGTGTCCTAAGATGAGCTCAACGAGAACAGAAATCTCGTG | 368 (0.007053%) | | | GGAGGGCAAGTCTGGTGCCAGCAGCCGCGGTAATTCCAGCTCCAATAGCGTATATTTAAGTTGTTGCAGTTAAAAAGCTCGTAGTTGAACCTTGGGATGG | 85 (0.001629%) | | | GGAGTAATGATTAACAGGGACAGTCGGGGGCATTCGTATTTCATAGTCAGAGGTGAAATTCTTGGATTTATGAAAGACGAACAACTGCGAAAGCATTTGC | 960 (0.018398%) | | | GGAGTCCGGAGACGTCGGCGGGGGCCTCGGGAAGAGTTATCTTTTCTGTTTAACAGCCTGCCCACCCTGGAAACGGCTCAGCCGGAGGTAGGGTCCAGCG | 5 (0.000096%) | | | GGAGTTTGGCTGGGGCGGCACATCTGTTAAAAGATAACGCAGGTGTCCTAAGATGAGCTCAACGAGAACAGAAATCTCGTGTGGAACAAAAGGGTAAAAG | 5 (0.000096%) | | | GGAGTTTTTTCAGCAGTTCTCGGACAAAAATTGCTGAGTGGCCGAGAAGAATGGGCGTGTCATGCGTGGGCTGACATGGATTCTTCGAGGCCTAGGGGTG | 5 (0.000096%) | | | GGATAGCTAGTGTTCGTAGGCTCCATGCTCGCGCATCGAACTACCTACCACCTATCCTTCTCAGTTAATTCACGGGCGATGTTACGCTCGATGATGAGTT | 33 (0.000632%) | | | GGATTTTCAAGGGCCGCCGGGGGCGCACCGGACACCACGCGACGTGCGGTGCTCTTCCAGCCGCTGGACCCTACCTCCGGCTGAGCCGTTTCCAGGGTGG | 43 (0.000824%) | | | GGCAACGTTAGGGAGTCCGGAGACGTCGGCGGGGGCCTCGGGAAGAGTTATCTTTTCTGTTTAACAGCCTGCCCACCCTGGAAACGGCTCAGCCGGAGGT | 8 (0.000153%) | | | GGCAAGTCTGGTGCCAGCAGCCGCGGTAATTCCAGCTCCAATAGCGTATATTTAAGTTGTTGCAGTTAAAAAGCTCGTAGTTGAACCTTGGGATGGGTCG | 44 (0.000843%) | | | GGCAATTCCCCGCCACATCCTCTCAAACGCAATGGAAAGAGAGAAAGGACGAGGTCTTGACCGTCATCTTTTGCCCGAAGGACGGATGAGCTTTGGCGGG | 64 (0.001227%) | | | GGCACATCTGTTAAAAGATAACGCAGGTGTCCTAAGATGAGCTCAACGAGAACAGAAATCTCGTGTGGAACAAAAGGGTAAAAGCTCGTTTGATTCTGAT | 16 (0.000307%) | | | GGCACTCGGTCCTCCGGATTTTCAAGGGCCGCCGGGGGCGCACCGGACACCACGCGACGTGCGGTGCTCTTCCAGCCGCTGGACCCTACCTCCGGCTGAG | 21 (0.000402%) | | | GGCAGAGCCCGCGTCGACCTTTTATCTAATAAATGCGTCCCTTCCATAAGTCGGGGTTTGTTGCACGTATTAGCTCTAGAATTACTACGGTTATCCGAGT | 29 (0.000556%) | | | GGCATCAGCGTGCTCCGGGCGTCGGCCTGTGGGCTCCCCATTCGACCCGTCTTGAAACACGGACCAAGGAGTCTGACATGTGTGCGAGTCAACGGGTGAG | 12 (0.000230%) | | | GGCATCGACACCTTGCGGCTAGGAACTGGAACGAGACGGGTGGCAAAGATTTCGAGTAGCACTTCATACTACCGTGGGTTTTTTAAACCTTCCGAGTTTT | 6 (0.000115%) | | | GGCATTCGTATTTCATAGTCAGAGGTGAAATTCTTGGATTTATGAAAGACGAACAACTGCGAAAGCATTTGCCAAGGATGTTTTCATTAATCAAGAACGA | 4 (0.000077%) | | | GGCCGCCGGGGGCGCACCGGACACCACGCGACGTGCGGTGCTCTTCCAGCCGCTGGACCCTACCTCCGGCTGAGCCGTTTCCAGGGTGGGCAGGCTGTTA | 7 (0.000134%) | | | GGCCTCTGTGCTGGCGACGCATCATTCAAATTTCTGCCCTATCAACTTTCGATGGTAGGATAGTGGCCTACCATGGTGGTAACGGGTGACGGAGAATTAG | 147 (0.002817%) | | | GGCCTGTGGGCTCCCCATTCGACCCGTCTTGAAACACGGACCAAGGAGTCTGACATGTGTGCGAGTCAACGGGTGAGTAAACCCGTAAGGCGCAAGGAAG | 8 (0.000153%) | | | GGCGACGCATCATTCAAATTTCTGCCCTATCAACTTTCGATGGTAGGATAGTGGCCTACCATGGTGGTAACGGGTGACGGAGAATTAGGGTTCGATTCCG | 29 (0.000556%) | | | GGCGACGTGGGTGGTTCGCCGCCCGCGACGTCGCGAGAAGTCCACTAAACCTTATCATTTAGAGGAAGGAGAAGTCGTAACAAGGTTTCCGTAGGTGAAC | 348 (0.006669%) | | | GGCGAGACAAGGGTTCACATTTCGTTCATCACCCTTGGCCGGCTATCGAACAGCCGGACTCCCATCAAAAGATGGTTGCCAAGAACATCTTCGTTACGGT | 12 (0.000230%) | | | GGCGAGACAAGGGTTCACATTTCGTTCATCACCCTTGGCCGGCTTTCGAACAGCCGGACTCCCATCAAAAGATGGTTGCCAAGAACATCTTCGTTACGGT | 25 (0.000479%) | | | GGCGCACCGGACACCACGCGACGTGCGGTGCTCTTCCAGCCGCTGGACCCTACCTCCGGCTGAGCCGTTTCCAGGGTGGGCAGGCTGTTAAACAGAAAAG | 11 (0.000211%) | | | GGCGGCACATCTGTTAAAAGATAACGCAGGTGTCCTAAGATGAGCTCAACGAGAACAGAAATCTCGTGTGGAACAAAAGGGTAAAAGCTCGTTTGATTCT | 6 (0.000115%) | | | GGCGGTTGACGGCAACGTTAGGGAGTCCGGAGACGTCGGCGGGGGCCTCGGGAAGAGTTATCTTTTCTGTTTAACAGCCTGCCCACCCTGGAAACGGCTC | 14 (0.000268%) | | | GGCGTAAGAATTGTATCCTTGTTAGAAGACACAAAGCCAA | 55 (0.000422%) | | | GGCGTGCCTCGGCATCAGCGTGCTCCGGGCGTCGGCCTGTGGGCTCCCCATTCGACCCGTCTTGAAACACGGACCAAGGAGTCTGACATGTGTGCGAGTC | 18 (0.000345%) | | | GGCTGGGGCGGCACATCTGTTAAAAGATAACGCAGGTGTCCTAAGATGAGCTCAACGAGAACAGAAATCTCGTGTGGAACAAAAGGGTAAAAGCTCGTTT | 10 (0.000192%) | | | GGCTGTCCCGAGTGTGAGCGAGGTGTGAGTGTCGCCCATGGGCATCGACACCTTGCGGCTAGGAACTGGAACGAGACGGGTAGCAAAGATTTCGAGTAGC | 19 (0.000364%) | | | GGCTGTCCCGAGTGTGAGCGAGGTGTGAGTGTCGCCCATGGGCATCGACACCTTGCGGCTAGGAACTGGAACGAGACGGGTGGCAAAGATTTCGAGTAGC | 16 (0.000307%) | | | GGCTTTGGCTCGGATTCGTCCGTCTTCTTTCTTCTTAGCCGAGTACTTCGGTAGATTAGTTGGAACGATTGATGATTTTGAGTTAATTGAACGTTCGGCG | 8 (0.000153%) | | | GGGACAGTCGGGGGCATTCGTATTTCATAGTCAGAGGTGAAATTCTTGGATTTATGAAAGACGAACAACTGCGAAAGCATTTGCCAAGGATGTTTTCATT | 17 (0.000326%) | | | GGGACGTGGCGGTTGACGGCAACGTTAGGGAGTCCGGAGACGTCGGCGGGGGCCTCGGGAAGAGTTATCTTTTCTGTTTAACAGCCTGCCCACCCTGGAA | 20 (0.000383%) | | | GGGAGTCCGGAGACGTCGGCGGGGGCCTCGGGAAGAGTTATCTTTTCTGTTTAACAGCCTGCCCACCCTGGAAACGGCTCAGCCGGAGGTAGGGTCCAGC | 4 (0.000077%) | | | GGGAGTTTGGCTGGGGCGGCACATCTGTTAAAAGATAACGCAGGTGTCCTAAGATGAGCTCAACGAGAACAGAAATCTCGTGTGGAACAAAAGGGTAAAA | 32 (0.000613%) | | | GGGCAAGTCTGGTGCCAGCAGCCGCGGTAATTCCAGCTCCAATAGCGTATATTTAAGTTGTTGCAGTTAAAAAGCTCGTAGTTGAACCTTGGGATGGGTC | 148 (0.002836%) | | | GGGCAGAGCCCGCGTCGACCTTTTATCTAATAAATGCGTCCCTTCCATAAGTCGGGGTTTGTTGCACGTATTAGCTCTAGAATTACTACGGTTATCCGAG | 53 (0.001016%) | | | GGGCATCGACACCTTGCGGCTAGGAACTGGAACGAGACGGGTGGCAAAGATTTCGAGTAGCACTTCATACTACCGTGGGTTTTTTAAACCTTCCGAGTTT | 13 (0.000249%) | | | GGGCATTCGTATTTCATAGTCAGAGGTGAAATTCTTGGATTTATGAAAGACGAACAACTGCGAAAGCATTTGCCAAGGATGTTTTCATTAATCAAGAACG | 29 (0.000556%) | | | GGGCCGCCGGGGGCGCACCGGACACCACGCGACGTGCGGTGCTCTTCCAGCCGCTGGACCCTACCTCCGGCTGAGCCGTTTCCAGGGTGGGCAGGCTGTT | 9 (0.000172%) | | | GGGCGCACCGGACACCACGCGACGTGCGGTGCTCTTCCAGCCGCTGGACCCTACCTCCGGCTGAGCCGTTTCCAGGGTGGGCAGGCTGTTAAACAGAAAA | 4 (0.000077%) | | | GGGCGGCACATCTGTTAAAAGATAACGCAGGTGTCCTAAGATGAGCTCAACGAGAACAGAAATCTCGTGTGGAACAAAAGGGTAAAAGCTCGTTTGATTC | 7 (0.000134%) | | | GGGCGGTGTGTACAAAGGGCAGGGACGTAGTCAACGCGAGCTGATGACTCGCGCTTACTAGGAATTCCTCGTTGAAGACCAACAATTGCAATGATCGATC | 22 (0.000422%) | | | GGGGAGTTTGGCTGGGGCGGCACATCTGTTAAAAGATAACGCAGGTGTCCTAAGATGAGCTCAACGAGAACAGAAATCTCGTGTGGAACAAAAGGGTAAA | 31 (0.000594%) | | | GGGGCATTCGTATTTCATAGTCAGAGGTGAAATTCTTGGATTTATGAAAGACGAACAACTGCGAAAGCATTTGCCAAGGATGTTTTCATTAATCAAGAAC | 24 (0.000460%) | | | GGGGCGCACCGGACACCACGCGACGTGCGGTGCTCTTCCAGCCGCTGGACCCTACCTCCGGCTGAGCCGTTTCCAGGGTGGGCAGGCTGTTAAACAGAAA | 9 (0.000172%) | | | GGGGCGGCACATCTGTTAAAAGATAACGCAGGTGTCCTAAGATGAGCTCAACGAGAACAGAAATCTCGTGTGGAACAAAAGGGTAAAAGCTCGTTTGATT | 5 (0.000096%) | | | GGGGGCATTCGTATTTCATAGTCAGAGGTGAAATTCTTGGATTTATGAAAGACGAACAACTGCGAAAGCATTTGCCAAGGATGTTTTCATTAATCAAGAA | 17 (0.000326%) | | | GGGGGCGCACCGGACACCACGCGACGTGCGGTGCTCTTCCAGCCGCTGGACCCTACCTCCGGCTGAGCCGTTTCCAGGGTGGGCAGGCTGTTAAACAGAA | 21 (0.000402%) | | | GGGGTTGAAATCGTCGACCAGGTCCGAGACTTCATCGACCGGGTCCGAGGATTCGTCGACCAGGACGGCCGGATGTCCGAGAAAAAAAAATGTTGCCGAA | 5 (0.000096%) | | | GGGTGGTTCGCCGCCCGCGACGTCGCGAGAAGTCCACTAAACCTTATCATTTAGAGGAAGGAGAAGTCGTAACAAGGTTTCCGTAGGTGAACCTGCGGAA | 146 (0.002798%) | | | GGGTGTCACAAATCGTCGTCCCTCACCATCCTTTGCTGATGCGGGACGGAAGCTGGTCTCCCGTGTGTTACCGCACGCGGTTGGCCTAAATCCGAGCCAA | 3 (0.000057%) | | | GGGTGTTGGTCGATTAAGACAGCAGGACGGTGGTCATGGAAGTCGAAATCCGCTAAGGAGTGTGTAACAACTCACCTGCCGAATCAACTAGCCCCGAAAA | 17 (0.000326%) | | | GGGTTGCGGTTTAAGTTGTTATACTCAATCATACACATGACAACAAGTCATATTCGACTCCAAAACACTAACCAACCTTCTTCTTGCTTCTCAAAGCTTT | 453 (0.008682%) | | | GGGTTTACTCACCCGTTGACTCGCACACATGTCAGACTCCTTGGTCCGTGTTTCAAGACGGGTCGAATGGGGAGCCCACAGGCCGACGCCCGGAGCACGC | 8 (0.000153%) | | | GGTAATTCCAGCTCCAATAGCGTATATTTAAGTTGTTGCAGTTAAAAAGCTCGTAGTTGAACCTTGGGATGGGTCGGCCGGTCCGCCTTTGGTGTGCATT | 96 (0.001840%) | | | GGTACGAAACACAGGCCCCGGAACTCATCATCGAGCGTAACATCGCCCGTGAATTAACTGAGAAGGATAGGTGGTAGGTAGTTCGATGCGCGAGCATGGA | 6 (0.000115%) | | | GGTAGGAGCGACGGGCGGTGTGTACAAAGGGCAGGGACGTAGTCAACGCGAGCTGATGACTCGCGCTTACTAGGAATTCCTCGTTGAAGACCAACAATTG | 39 (0.000747%) | | | GGTATCTCGCGCTTGTACGGCTTTGGCTCGGATTCGTCCGTCTTCTTTCTTCTTAGCCGAGTACTTCGGTAGATTAGTTGGAACGATTGATGATTTTGAG | 18 (0.000345%) | | | GGTCCTCCGGATTTTCAAGGGCCGCCGGGGGCGCACCGGACACCACGCGACGTGCGGTGCTCTTCCAGCCGCTGGACCCTACCTCCGGCTGAGCCGTTTC | 6 (0.000115%) | | | GGTCGACCAGGGGTTGAAATCGTCGACCAGGTCCGAGACTTCATCGACCGGGTCCGAGGATTCGTCGACCAGGACGGCCGGATGTCCGAGAAAAAAAAAT | 800 (0.015332%) | | | GGTCGATTAAGACAGCAGGACGGTGGTCATGGAAGTCGAAATCCGCTAAGGAGTGTGTAACAACTCACCTGCCGAATCAACTAGCCCCGAAAATGGATGG | 43 (0.000824%) | | | GGTCTTCAACGAGGAATTCCTAGTAAGCGCGAGTCATCAGCTCGCGTTGACTACGTCCCTGCCCTTTGTACACACCGCCCGTCGCTCCTACCGATTGAAT | 22 (0.000422%) | | | GGTGCCAGCAGCCGCGGTAATTCCAGCTCCAATAGCGTATATTTAAGTTGTTGCAGTTAAAAAGCTCGTAGTTGAACCTTGGGATGGGTCGGCCGGTCCG | 9 (0.000172%) | | | GGTGGTCATGGAAGTCGAAATCCGCTAAGGAGTGTGTAACAACTCACCTGCCGAATCAACTAGCCCCGAAAATGGATGGCGCTTAAGCGCGCGACCTATA | 13 (0.000249%) | | | GGTGGTTCGCCGCCCGCGACGTCGCGAGAAGTCCACTAAACCTTATCATTTAGAGGAAGGAGAAGTCGTAACAAGGTTTCCGTAGGTGAACCTGCGGAAG | 22 (0.000422%) | | | GGTGTCACAAATCGTCGTCCCTCACCATCCTTTGCTGATGCGGGACGGAAGCTGGTCTCCCGTGTGTTACCGCACGCGGTTGGCCTAAATCCGAGCCAAG | 6 (0.000115%) | | | GGTGTGAGTGTCGCCCATGGGCATCGACACCTTGCGGCTAGGAACTGGAACGAGACGGGTAGCAAAGATTTCGAGTAGCACTTCATACTACCGTGGGTTT | 9 (0.000172%) | | | GGTGTGAGTGTCGCCCATGGGCATCGACACCTTGCGGCTAGGAACTGGAACGAGACGGGTGGCAAAGATTTCGAGTAGCACTTCATACTACCGTGGGTTT | 5 (0.000096%) | | | GGTGTTGGTCGATTAAGACAGCAGGACGGTGGTCATGGAAGTCGAAATCCGCTAAGGAGTGTGTAACAACTCACCTGCCGAATCAACTAGCCCCGAAAAT | 3 (0.000057%) | | | GGTTCACATTTCGTTCATCACCCTTGGCCGGCTTTCGAACAGCCGGACTCCCATCAAAAGATGGTTGCCAAGAACATCTTCGTTACGGTTTGCTAATTCT | 3 (0.000057%) | | | GGTTCGAGTGTGAGCATGCCTGTCGGGACCCGAAAGATGGTGAACTATGCCTGAGCGGGGTAAAGCCAGAGGAAACTCTGGTGGAAGCCCGCAGCGATAC | 3 (0.000057%) | | | GGTTCGCCGCCCGCGACGTCGCGAGAAGTCCACTAAACCTTATCATTTAGAGGAAGGAGAAGTCGTAACAAGGTTTCCGTAGGTGAACCTGCGGAAGGAT | 18 (0.000345%) | | | GGTTCGTATTCGTACTGAAAATCAGAATCAAACGAGCTTTTACCCTTTTGTTCCACACGAGATTTCTGTTCTCGTTGAGCTCATCTTAGGACACCTGCGT | 11 (0.000211%) | | | GGTTGAAATCGTCGACCAGGTCCGAGACTTCATCGACCGGGTCCGAGGATTCGTCGACCAGGACGGCCGGATGTCCGAGAAAAAAAAATGTTGCCGAATA | 17 (0.000326%) | | | GGTTGACGGCAACGTTAGGGAGTCCGGAGACGTCGGCGGGGGCCTCGGGAAGAGTTATCTTTTCTGTTTAACAGCCTGCCCACCCTGGAAACGGCTCAGC | 18 (0.000345%) | | | GGTTGCGGTTTAAGTTCTTATACTCAATCATACACATGACATCAAGTCATATTCGACTCCAAAACACTAACCAACCTTCTTCTTGCTTCTCAAAGCTTTC | 1203 (0.023055%) | | | GGTTGCGGTTTAAGTTGTTATACTCAATCATACACATGACAACAAGTCATATTCGACTCCAAAACACTAACCAACCTTCTTCTTGCTTCTCAAAGCTTTC | 21 (0.000402%) | | | GGTTTAAGTTCTTATACTCAATCATACACATGACATCAAGTCATATTCGACTCCAAAACACTAACCAACCTTCTTCTTGCTTCTCAAAGCTTTCATGGTG | 253 (0.004849%) | | | GGTTTAAGTTGTTATACTCAATCATACACATGACAACAAGTCATATTCGACTCCAAAACACTAACCAACCTTCTTCTTGCTTCTCAAAGCTTTCATGGTG | 116 (0.002223%) | | | GGTTTACTCACCCGTTGACTCGCACACATGTCAGACTCCTTGGTCCGTGTTTCAAGACGGGTCGAATGGGGAGCCCACAGGCCGACGCCCGGAGCACGCT | 24 (0.000460%) | | | GTAAAGTTGGGAATTCGTTAAGGAGCTGTTGCTTTGTTAGTGTAGAAACACTTGTGTAGAATTGGGGATTGTTTTTTTTGGAGTGATTTAGGGGAGGGTC | 46 (0.000882%) | | | GTAAGAATTGTATCCTTGTTAGAAGACACAAAGCCAAAGA | 320 (0.002453%) | | | GTAATTCCAGCTCCAATAGCGTATATTTAAGTTGTTGCAGTTAAAAAGCTCGTAGTTGAACCTTGGGATGGGTCGGCCGGTCCGCCTTTGGTGTGCATTG | 14 (0.000268%) | | | GTACGAAACACAGGCCCCGGAACTCATCATCGAGCGTAACATCGCCCGTGAATTAACTGAGAAGGATAGGTGGTAGGTAGTTCGATGCGCGAGCATGGAG | 16 (0.000307%) | | | GTACGGCTTTGGCTCGGATTCGTCCGTCTTCTTTCTTCTTAGCCGAGTACTTCGGTAGATTAGTTGGAACGATTGATGATTTTGAGTTAATTGAACGTTC | 6 (0.000115%) | | | GTAGGAGCGACGGGCGGTGTGTACAAAGGGCAGGGACGTAGTCAACGCGAGCTGATGACTCGCGCTTACTAGGAATTCCTCGTTGAAGACCAACAATTGC | 100 (0.001916%) | | | GTAGGCTCCATGCTCGCGCATCGAACTACCTACCACCTATCCTTCTCAGTTAATTCACGGGCGATGTTACGCTCGATGATGAGTTCCGGGGCCTGTGTTT | 13 (0.000249%) | | | GTAGTCATATGCTTGTCTCAAAGATTAAGCCATGCATGTGTAAGTATGAACGAATTCAGACTGTGAAACTGCGAATGGCTCATTAAATCAGTTATAGTTT | 79 (0.001514%) | | | GTATAAGAACTTAAACCGCAACCGCATCTTATAAGCCTAAGTAGTGTTTCCTTGTTAGAAGACACAAAGCCAAAGACTCATATGGACTTTGGCTACACCA | 6 (0.000115%) | | | GTATATTTAAGTTGTTGCAGTTAAAAAGCTCGTAGTTGAACCTTGGGATGGGTCGGCCGGTCCGCCTTTGGTGTGCATTGGTCGGCTTGTCCCTTCGGTC | 23 (0.000441%) | | | GTATCCAGAGCGTAGGCTTGCTTTGAGCACTCTAATTTCTTCAAAGTAACAGCGCCGGAGGCACGACCCGGCCAATTAAGACCAGGAGCGTATCGCCGAC | 24 (0.000460%) | | | GTATCCTTGTTAGAAGACACAAAGCCAAAGACTCATATGGACTTTGGCTACACCATGAAAGCTTTGAGAAGCAAGAAGAAGGTTGGTTAGTGTTTTGGAG | 17 (0.000326%) | | | GTATGATTGAGTATAAGAACTTAAACCGCAACCGCATCTTATAAGCCTAAGTAGTGTTTCCTTGTTAGAAGACACAAAGCCAAAGACTCATATGGACTTT | 46 (0.000882%) | | | GTATTCGTACTGAAAATCAGAATCAAACGAGCTTTTACCCTTTTGTTCCACACGAGATTTCTGTTCTCGTTGAGCTCATCTTAGGACACCTGCGTTATCT | 5 (0.000096%) | | | GTCACAAATCGTCGTCCCTCACCATCCTTTGCTGATGCGGGACGGAAGCTGGTCTCCCGTGTGTTACCGCACGCGGTTGGCCTAAATCCGAGCCAAGGAC | 13 (0.000249%) | | | GTCAGGTGGGGAGTTTGGCTGGGGCGGCACATCTGTTAAAAGATAACGCAGGTGTCCTAAGATGAGCTCAACGAGAACAGAAATCTCGTGTGGAACAAAA | 8 (0.000153%) | | | GTCATATGCTTGTCTCAAAGATTAAGCCATGCATGTGTAAGTATGAACGAATTCAGACTGTGAAACTGCGAATGGCTCATTAAATCAGTTATAGTTTGTT | 38 (0.000728%) | | | GTCATATTCGACTCCAAAACACTAACCAACCTTCTTCTTGCTTCTCAAAGCTTTCATGGTGTAGCCAAAGTCCATATGAGTCTTTGGCTTTGTGTCTTCT | 188 (0.003603%) | | | GTCCCGAAGGTATCTCGCGCTTGTACGGCTTTGGCTCGGATTCGTCCGTCTTCTTTCTTCTTAGCCGAGTACTTCGGTAGATTAGTTGGAACGATTGATG | 10 (0.000192%) | | | GTCCCGAGTGTGAGCGAGGTGTGAGTGTCGCCCATGGGCATCGACACCTTGCGGCTAGGAACTGGAACGAGACGGGTAGCAAAGATTTCGAGTAGCACTT | 8 (0.000153%) | | | GTCCCGAGTGTGAGCGAGGTGTGAGTGTCGCCCATGGGCATCGACACCTTGCGGCTAGGAACTGGAACGAGACGGGTGGCAAAGATTTCGAGTAGCACTT | 12 (0.000230%) | | | GTCCCTCACCATCCTTTGCTGATGCGGGACGGAAGCTGGTCTCCCGTGTGTTACCGCACGCGGTTGGCCTAAATCCGAGCCAAGGACGCCTGGAGCGTAC | 11 (0.000211%) | | | GTCCGGAGACGTCGGCGGGGGCCTCGGGAAGAGTTATCTTTTCTGTTTAACAGCCTGCCCACCCTGGAAACGGCTCAGCCGGAGGTAGGGTCCAGCGGCT | 4 (0.000077%) | | | GTCCTCCGGATTTTCAAGGGCCGCCGGGGGCGCACCGGACACCACGCGACGTGCGGTGCTCTTCCAGCCGCTGGACCCTACCTCCGGCTGAGCCGTTTCC | 4 (0.000077%) | | | GTCGAATATGACTTGATGTCATGTGTATGATTGAGTATAA | 466 (0.003572%) | | | GTCGACCAGGGGTTGAAATCGTCGACCAGGTCCGAGACTTCATCGACCGGGTCCGAGGATTCGTCGACCAGGACGGCCGGATGTCCGAGAAAAAAAAATG | 58 (0.001112%) | | | GTCGAGTTATCATGAATCATCAGAGCAACGGGCAGAGCCCGCGTCGACCTTTTATCTAATAAATGCGTCCCTTCCATAAGTCGGGGTTTGTTGCACGTAT | 443 (0.008490%) | | | GTCGATTAAGACAGCAGGACGGTGGTCATGGAAGTCGAAATCCGCTAAGGAGTGTGTAACAACTCACCTGCCGAATCAACTAGCCCCGAAAATGGATGGC | 27 (0.000517%) | | | GTCGCCCATGGGCATCGACACCTTGCGGCTAGGAACTGGAACGAGACGGGTGGCAAAGATTTCGAGTAGCACTTCATACTACCGTGGGTTTTTTAAACCT | 18 (0.000345%) | | | GTCGGGGGCATTCGTATTTCATAGTCAGAGGTGAAATTCTTGGATTTATGAAAGACGAACAACTGCGAAAGCATTTGCCAAGGATGTTTTCATTAATCAA | 7 (0.000134%) | | | GTCGTCCCTCACCATCCTTTGCTGATGCGGGACGGAAGCTGGTCTCCCGTGTGTTACCGCACGCGGTTGGCCTAAATCCGAGCCAAGGACGCCTGGAGCG | 33 (0.000632%) | | | GTCTCAAAGATTAAGCCATGCATGTGTAAGTATGAACGAATTCAGACTGTGAAACTGCGAATGGCTCATTAAATCAGTTATAGTTTGTTTGATGGTAACT | 5 (0.000096%) | | | GTCTGGTGCCAGCAGCCGCGGTAATTCCAGCTCCAATAGCGTATATTTAAGTTGTTGCAGTTAAAAAGCTCGTAGTTGAACCTTGGGATGGGTCGGCCGG | 20 (0.000383%) | | | GTCTTCAACGAGGAATTCCTAGTAAGCGCGAGTCATCAGCTCGCGTTGACTACGTCCCTGCCCTTTGTACACACCGCCCGTCGCTCCTACCGATTGAATG | 9 (0.000172%) | | | GTGAGCGAGGTGTGAGTGTCGCCCATGGGCATCGACACCTTGCGGCTAGGAACTGGAACGAGACGGGTGGCAAAGATTTCGAGTAGCACTTCATACTACC | 22 (0.000422%) | | | GTGAGCGGCACTCGGTCCTCCGGATTTTCAAGGGCCGCCGGGGGCGCACCGGACACCACGCGACGTGCGGTGCTCTTCCAGCCGCTGGACCCTACCTCCG | 377 (0.007225%) | | | GTGAGTGTCGCCCATGGGCATCGACACCTTGCGGCTAGGAACTGGAACGAGACGGGTGGCAAAGATTTCGAGTAGCACTTCATACTACCGTGGGTTTTTT | 33 (0.000632%) | | | GTGCCAGCAGCCGCGGTAATTCCAGCTCCAATAGCGTATATTTAAGTTGTTGCAGTTAAAAAGCTCGTAGTTGAACCTTGGGATGGGTCGGCCGGTCCGC | 31 (0.000594%) | | | GTGCCTCGGCATCAGCGTGCTCCGGGCGTCGGCCTGTGGGCTCCCCATTCGACCCGTCTTGAAACACGGACCAAGGAGTCTGACATGTGTGCGAGTCAAC | 15 (0.000287%) | | | GTGCTCCGGGCGTCGGCCTGTGGGCTCCCCATTCGACCCGTCTTGAAACACGGACCAAGGAGTCTGACATGTGTGCGAGTCAACGGGTGAGTAAACCCGT | 6 (0.000115%) | | | GTGCTGGCGACGCATCATTCAAATTTCTGCCCTATCAACTTTCGATGGTAGGATAGTGGCCTACCATGGTGGTAACGGGTGACGGAGAATTAGGGTTCGA | 28 (0.000537%) | | | GTGGCGGTTGACGGCAACGTTAGGGAGTCCGGAGACGTCGGCGGGGGCCTCGGGAAGAGTTATCTTTTCTGTTTAACAGCCTGCCCACCCTGGAAACGGC | 18 (0.000345%) | | | GTGGGGAGTTTGGCTGGGGCGGCACATCTGTTAAAAGATAACGCAGGTGTCCTAAGATGAGCTCAACGAGAACAGAAATCTCGTGTGGAACAAAAGGGTA | 13 (0.000249%) | | | GTGGGTGGTTCGCCGCCCGCGACGTCGCGAGAAGTCCACTAAACCTTATCATTTAGAGGAAGGAGAAGTCGTAACAAGGTTTCCGTAGGTGAACCTGCGG | 77 (0.001476%) | | | GTGGTCATGGAAGTCGAAATCCGCTAAGGAGTGTGTAACAACTCACCTGCCGAATCAACTAGCCCCGAAAATGGATGGCGCTTAAGCGCGCGACCTATAC | 13 (0.000249%) | | | GTGGTTCGCCGCCCGCGACGTCGCGAGAAGTCCACTAAACCTTATCATTTAGAGGAAGGAGAAGTCGTAACAAGGTTTCCGTAGGTGAACCTGCGGAAGG | 12 (0.000230%) | | | GTGTAAAGTTGGGAATTCGTTAAGGAGCTGTTGCTTTGTTAGTGTAGAAACACTTGTGTAGAATTGGGGATTGTTTTTTTTGGAGTGATTTAGGGGAGGG | 38 (0.000728%) | | | GTGTATCCTTGTTAGAAGACACAAAGCCAAAGACTCATAT | 66 (0.000506%) | | | GTGTATGATTGAGTATAAGAACTTAAACCGCAACCGCATCTTATAAGCCTAAGTAGTGTTTCCTTGTTAGAAGACACAAAGCCAAAGACTCATATGGACT | 6 (0.000115%) | | | GTGTCACAAATCGTCGTCCCTCACCATCCTTTGCTGATGCGGGACGGAAGCTGGTCTCCCGTGTGTTACCGCACGCGGTTGGCCTAAATCCGAGCCAAGG | 6 (0.000115%) | | | GTGTCGCCCATGGGCATCGACACCTTGCGGCTAGGAACTGGAACGAGACGGGTGGCAAAGATTTCGAGTAGCACTTCATACTACCGTGGGTTTTTTAAAC | 10 (0.000192%) | | | GTGTGAGCATGCCTGTCGGGACCCGAAAGATGGTGAACTATGCCTGAGCGGGGTAAAGCCAGAGGAAACTCTGGTGGAAGCCCGCAGCGATACTGACGTG | 4 (0.000077%) | | | GTGTGAGCGAGGTGTGAGTGTCGCCCATGGGCATCGACACCTTGCGGCTAGGAACTGGAACGAGACGGGTAGCAAAGATTTCGAGTAGCACTTCATACTA | 19 (0.000364%) | | | GTGTGAGCGAGGTGTGAGTGTCGCCCATGGGCATCGACACCTTGCGGCTAGGAACTGGAACGAGACGGGTGGCAAAGATTTCGAGTAGCACTTCATACTA | 29 (0.000556%) | | | GTGTGAGTGTCGCCCATGGGCATCGACACCTTGCGGCTAGGAACTGGAACGAGACGGGTAGCAAAGATTTCGAGTAGCACTTCATACTACCGTGGGTTTT | 11 (0.000211%) | | | GTGTGAGTGTCGCCCATGGGCATCGACACCTTGCGGCTAGGAACTGGAACGAGACGGGTGGCAAAGATTTCGAGTAGCACTTCATACTACCGTGGGTTTT | 15 (0.000287%) | | | GTGTTCGTAGGCTCCATGCTCGCGCATCGAACTACCTACCACCTATCCTTCTCAGTTAATTCACGGGCGATGTTACGCTCGATGATGAGTTCCGGGGCCT | 8 (0.000153%) | | | GTGTTGGTCGATTAAGACAGCAGGACGGTGGTCATGGAAGTCGAAATCCGCTAAGGAGTGTGTAACAACTCACCTGCCGAATCAACTAGCCCCGAAAATG | 6 (0.000115%) | | | GTGTTTCCTTGTTAGAAGACACAAAGCCAAAGACTCATATGGACTTTGGCTACACCATGAAAGCTTTGAGAAGCAAGAAGAAGGTTGGTTAGTGTTTTGG | 25 (0.000479%) | | | GTGTTTTGGAGTCGAATATGACTTGATGTCATGTGTATGA | 209 (0.001602%) | | | GTTAAAAGATAACGCAGGTGTCCTAAGATGAGCTCAACGAGAACAGAAATCTCGTGTGGAACAAAAGGGTAAAAGCTCGTTTGATTCTGATTTTCAGTAC | 30 (0.000575%) | | | GTTAAGGAGCTGTTGCTTTGTTAGTGTAGAAACACTTGTGTAGAATTGGGGATTGTTTTTTTTGGAGTGATTTAGGGGAGGGTCGAATCTTAGCGACAAA | 47 (0.000901%) | | | GTTAGAAGACACAAAGCCAAAGACTCATATGGACTTTGGCTACACCATGAAAGCTTTGAGAAGCAAGAAGAAGGTTGGTTAGTGTTTTGGAGTCGAATAT | 73 (0.001399%) | | | GTTAGGGAGTCCGGAGACGTCGGCGGGGGCCTCGGGAAGAGTTATCTTTTCTGTTTAACAGCCTGCCCACCCTGGAAACGGCTCAGCCGGAGGTAGGGTC | 13 (0.000249%) | | | GTTAGTGTTTTGGAGTCGAATATGACTTGATGTCATGTGT | 15393 (0.118000%) | | | GTTATCATGAATCATCAGAGCAACGGGCAGAGCCCGCGTCGACCTTTTATCTAATAAATGCGTCCCTTCCATAAGTCGGGGTTTGTTGCACGTATTAGCT | 264 (0.005059%) | | | GTTATCCCATGCTAATGTATCCAGAGCGTAGGCTTGCTTTGAGCACTCTAATTTCTTCAAAGTAACAGCGCCGGAGGCACGACCCGGCCAATTAAGACCA | 49 (0.000939%) | | | GTTCAATATGACAAACCCATGCCAAGTAAAGAGAAAATGAAAACTGGTGATTGTTGCGGAAATCGTCCAGGATTCCTCGACCAGGACTTGAAATCGTCGA | 23 (0.000441%) | | | GTTCACATTTCGTTCATCACCCTTGGCCGGCTATCGAACAGCCGGACTCCCATCAAAAGATGGTTGCCAAGAACATCTTCGTTACGGTTTGCTAATTCTC | 12 (0.000230%) | | | GTTCACATTTCGTTCATCACCCTTGGCCGGCTTTCGAACAGCCGGACTCCCATCAAAAGATGGTTGCCAAGAACATCTTCGTTACGGTTTGCTAATTCTC | 19 (0.000364%) | | | GTTCATCACCCTTGGCCGGCTTTCGAACAGCCGGACTCCCATCAAAAGATGGTTGCCAAGAACATCTTCGTTACGGTTTGCTAATTCTCGGAATAACATC | 6 (0.000115%) | | | GTTCGATTAGTCTTTCGCCCCTATACCCAAGTCAGACGAACGATTTGCACGTCAGTATCGCTGCGGGCTTCCACCAGAGTTTCCTCTGGCTTTACCCCGC | 775 (0.014853%) | | | GTTCGCCGCCCGCGACGTCGCGAGAAGTCCACTAAACCTTATCATTTAGAGGAAGGAGAAGTCGTAACAAGGTTTCCGTAGGTGAACCTGCGGAAGGATC | 33 (0.000632%) | | | GTTCGTAGGCTCCATGCTCGCGCATCGAACTACCTACCACCTATCCTTCTCAGTTAATTCACGGGCGATGTTACGCTCGATGATGAGTTCCGGGGCCTGT | 8 (0.000153%) | | | GTTCGTATTCGTACTGAAAATCAGAATCAAACGAGCTTTTACCCTTTTGTTCCACACGAGATTTCTGTTCTCGTTGAGCTCATCTTAGGACACCTGCGTT | 12 (0.000230%) | | | GTTCTCGGACAAAAATTGCTGAGTGGCCGAGAAGAATGGGCGTGTCATGCGTGGGCTGACATGGATTCTTCGAGGCCTAGGGGTGGCGGTATATAACTTG | 3 (0.000057%) | | | GTTCTTATACTCAATCATACACATGACATCAAGTCATATTCGACTCCAAAACACTAACCAACCTTCTTCTTGCTTCTCAAAGCTTTCATGGTGTAGCCAA | 92 (0.001763%) | | | GTTGAAATCGTCGACCAGGTCCGAGACTTCATCGACCGGGTCCGAGGATTCGTCGACCAGGACGGCCGGATGTCCGAGAAAAAAAAATGTTGCCGAATAA | 41 (0.000786%) | | | GTTGACGGCAACGTTAGGGAGTCCGGAGACGTCGGCGGGGGCCTCGGGAAGAGTTATCTTTTCTGTTTAACAGCCTGCCCACCCTGGAAACGGCTCAGCC | 41 (0.000786%) | | | GTTGACTCGCACACATGTCAGACTCCTTGGTCCGTGTTTCAAGACGGGTCGAATGGGGAGCCCACAGGCCGACGCCCGGAGCACGCTGATGCCGAGGCAC | 54 (0.001035%) | | | GTTGCGGTTTAAGTTCTTATACTCAATCATACACATGACATCAAGTCATATTCGACTCCAAAACACTAACCAACCTTCTTCTTGCTTCTCAAAGCTTTCA | 292 (0.005596%) | | | GTTGCGGTTTAAGTTGTTATACTCAATCATACACATGACAACAAGTCATATTCGACTCCAAAACACTAACCAACCTTCTTCTTGCTTCTCAAAGCTTTCA | 146 (0.002798%) | | | GTTGGTCGATTAAGACAGCAGGACGGTGGTCATGGAAGTCGAAATCCGCTAAGGAGTGTGTAACAACTCACCTGCCGAATCAACTAGCCCCGAAAATGGA | 15 (0.000287%) | | | GTTGGTCTTCAACGAGGAATTCCTAGTAAGCGCGAGTCATCAGCTCGCGTTGACTACGTCCCTGCCCTTTGTACACACCGCCCGTCGCTCCTACCGATTG | 175 (0.003354%) | | | GTTGTTATACTCAATCATACACATGACAACAAGTCATATTCGACTCCAAAACACTAACCAACCTTCTTCTTGCTTCTCAAAGCTTTCATGGTGTAGCCAA | 14 (0.000268%) | | | GTTGTTATACTCAATCATACACATGACATCAAGTCATATT | 240 (0.001840%) | | | GTTTAAGTTCTTATACTCAATCATACACATGACATCAAGTCATATTCGACTCCAAAACACTAACCAACCTTCTTCTTGCTTCTCAAAGCTTTCATGGTGT | 348 (0.006669%) | | | GTTTAAGTTGTTATACTCAATCATACACATGACAACAAGTCATATTCGACTCCAAAACACTAACCAACCTTCTTCTTGCTTCTCAAAGCTTTCATGGTGT | 127 (0.002434%) | | | GTTTACTCACCCGTTGACTCGCACACATGTCAGACTCCTTGGTCCGTGTTTCAAGACGGGTCGAATGGGGAGCCCACAGGCCGACGCCCGGAGCACGCTG | 19 (0.000364%) | | | GTTTGGCTGGGGCGGCACATCTGTTAAAAGATAACGCAGGTGTCCTAAGATGAGCTCAACGAGAACAGAAATCTCGTGTGGAACAAAAGGGTAAAAGCTC | 33 (0.000632%) | | | GTTTTAATTAAACAGTCGGATTCCCCTTGTCCGTACCAGTTCTGAGCTGACTGTTCGACGCCCGGGGAAAGCTCCCGAGAGAGCCGTTCCCAGTCCGTCC | 14 (0.000268%) | | | GTTTTGGAGTCGAATATGACTTGATGTCATGTGTATGATT | 459 (0.003519%) | | | GTTTTTTCAGCAGTTCTCGGACAAAAATTGCTGAGTGGCCGAGAAGAATGGGCGTGTCATGCGTGGGCTGACATGGATTCTTCGAGGCCTAGGGGTGGCG | 16 (0.000307%) | | | TAAAAGATAACGCAGGTGTCCTAAGATGAGCTCAACGAGAACAGAAATCTCGTGTGGAACAAAAGGGTAAAAGCTCGTTTGATTCTGATTTTCAGTACGA | 19 (0.000364%) | | | TAAAAGTGTCGGAGTTTTTTCAGCAGTTCTCGGACAAAAATTGCTGAGTGGCCGAGAAGAATGGGCGTGTCATGCGTGGGCTGACATGGATTCTTCGAGG | 8 (0.000153%) | | | TAAAGGCGTAAGAATTGTATCCTTGTTAGAAGACACAAAG | 572 (0.004385%) | | | TAAAGTTGGGAATTCGTTAAGGAGCTGTTGCTTTGTTAGTGTAGAAACACTTGTGTAGAATTGGGGATTGTTTTTTTTGGAGTGATTTAGGGGAGGGTCG | 3 (0.000057%) | | | TAAATACGGGCGAGAGACCGATAGCGAACAAGTACCGCGAGGTAAAGATGAAAAGGACTTTGAAAAGAGAGTCAAAGAGTGCTTGAAATTGTCGGGAGGG | 24 (0.000460%) | | | TAACAGGGACAGTCGGGGGCATTCGTATTTCATAGTCAGAGGTGAAATTCTTGGATTTATGAAAGACGAACAACTGCGAAAGCATTTGCCAAGGATGTTT | 61 (0.001169%) | | | TAACGGCGTGCCTCGGCATCAGCGTGCTCCGGGCGTCGGCCTGTGGGCTCCCCATTCGACCCGTCTTGAAACACGGACCAAGGAGTCTGACATGTGTGCG | 41 (0.000786%) | | | TAACTAGCTACGTGGAGGCATCCCTTCACGGCCGGCTTCTTAGAGGGACTATGGCCGTTTAGGCCAAGGAAGTTTGAGGCAATAACAGGTCTGTGATGCC | 33 (0.000632%) | | | TAAGAACTTAAACCGCAACCGCATCTTATAAGCCTAAGTAGTGTTTCCTTGTTAGAAGACACAAAGCCAAAGACTCATATGGACTTTGGCTACACCATGA | 38 (0.000728%) | | | TAAGAATTGTATCCTTGTTAGAAGACACAAAGCCAAAGACTCATATGGACTTTGGCTACACCATGAAAGCTTTGAGAAGCAAGAAGAAGGTTGGTTAGTG | 1020 (0.019548%) | | | TAAGACAGCAGGACGGTGGTCATGGAAGTCGAAATCCGCTAAGGAGTGTGTAACAACTCACCTGCCGAATCAACTAGCCCCGAAAATGGATGGCGCTTAA | 45 (0.000862%) | | | TAAGGAGCTGTTGCTTTGTTAGTGTAGAAACACTTGTGTAGAATTGGGGATTGTTTTTTTTGGAGTGATTTAGGGGAGGGTCGAATCTTAGCGACAAAGG | 26 (0.000498%) | | | TAAGTAGTGTTTCCTTGTTAGAAGACACAAAGCCAAAGACTCATATGGACTTTGGCTACACCATGAAAGCTTTGAGAAGCAAGAAGAAGGTTGGTTAGTG | 41 (0.000786%) | | | TAAGTTCTTATACTCAATCATACACATGACATCAAGTCATATTCGACTCCAAAACACTAACCAACCTTCTTCTTGCTTCTCAAAGCTTTCATGGTGTAGC | 23 (0.000441%) | | | TAAGTTGTTATACTCAATCATACACATGACAACAAGTCATATTCGACTCCAAAACACTAACCAACCTTCTTCTTGCTTCTCAAAGCTTTCATGGTGTAGC | 15 (0.000287%) | | | TAAGTTGTTATACTCAATCATACACATGACATCAAGTCAT | 9364 (0.071783%) | | | TAATGTATCCAGAGCGTAGGCTTGCTTTGAGCACTCTAATTTCTTCAAAGTAACAGCGCCGGAGGCACGACCCGGCCAATTAAGACCAGGAGCGTATCGC | 5 (0.000096%) | | | TAATTCCAGCTCCAATAGCGTATATTTAAGTTGTTGCAGTTAAAAAGCTCGTAGTTGAACCTTGGGATGGGTCGGCCGGTCCGCCTTTGGTGTGCATTGG | 10 (0.000192%) | | | TACACATGACATCAAGTCATATTCGACTCCAAAACACTAACCAACCTTCTTCTTGCTTCTCAAAGCTTTCATGGTGTAGCCAAAGTCCATATGAGTCTTT | 17 (0.000326%) | | | TACATTTTATCGGTCGCTCTTGTCCGGAAGCTGTAGATGACCCAAAGTCCATATAGCGACCCCAGGTCAGGCGGGATTACCCGCTGAGTTTAAGCATATC | 5 (0.000096%) | | | TACCTACCACCTATCCTTCTCAGTTAATTCACGGGCGATGTTACGCTCGATGATGAGTTCCGGGGCCTGTGTTTCGTACCTAATTTGAAGGAATTGTTGA | 4 (0.000077%) | | | TACGAAACACAGGCCCCGGAACTCATCATCGAGCGTAACATCGCCCGTGAATTAACTGAGAAGGATAGGTGGTAGGTAGTTCGATGCGCGAGCATGGAGC | 10 (0.000192%) | | | TACGGGTTTACTCACCCGTTGACTCGCACACATGTCAGACTCCTTGGTCCGTGTTTCAAGACGGGTCGAATGGGGAGCCCACAGGCCGACGCCCGGAGCA | 4 (0.000077%) | | | TACTCAATCATACACATGACATCAAGTCATATTCGACTCCAAAACACTAACCAACCTTCTTCTTGCTTCTCAAAGCTTTCATGGTGTAGCCAAAGTCCAT | 27 (0.000517%) | | | TACTCACCCGTTGACTCGCACACATGTCAGACTCCTTGGTCCGTGTTTCAAGACGGGTCGAATGGGGAGCCCACAGGCCGACGCCCGGAGCACGCTGATG | 3 (0.000057%) | | | TAGAAGACACAAAGCCAAAGACTCATATGGACTTTGGCTACACCATGAAAGCTTTGAGAAGCAAGAAGAAGGTTGGTTAGTGTTTTGGAGTCGAATATGA | 38 (0.000728%) | | | TAGCTAGTGTTCGTAGGCTCCATGCTCGCGCATCGAACTACCTACCACCTATCCTTCTCAGTTAATTCACGGGCGATGTTACGCTCGATGATGAGTTCCG | 6 (0.000115%) | | | TAGGAGCGACGGGCGGTGTGTACAAAGGGCAGGGACGTAGTCAACGCGAGCTGATGACTCGCGCTTACTAGGAATTCCTCGTTGAAGACCAACAATTGCA | 11 (0.000211%) | | | TAGGCGAGACAAGGGTTCACATTTCGTTCATCACCCTTGGCCGGCTTTCGAACAGCCGGACTCCCATCAAAAGATGGTTGCCAAGAACATCTTCGTTACG | 4 (0.000077%) | | | TAGGCTCCATGCTCGCGCATCGAACTACCTACCACCTATCCTTCTCAGTTAATTCACGGGCGATGTTACGCTCGATGATGAGTTCCGGGGCCTGTGTTTC | 4 (0.000077%) | | | TAGGCTGTCCCGAAGGTATCTCGCGCTTGTACGGCTTTGGCTCGGATTCGTCCGTCTTCTTTCTTCTTAGCCGAGTACTTCGGTAGATTAGTTGGAACGA | 1761 (0.033749%) | | | TAGGCTGTCCCGAGTGTGAGCGAGGTGTGAGTGTCGCCCATGGGCATCGACACCTTGCGGCTAGGAACTGGAACGAGACGGGTGGCAAAGATTTCGAGTA | 40 (0.000767%) | | | TAGGGAGTCCGGAGACGTCGGCGGGGGCCTCGGGAAGAGTTATCTTTTCTGTTTAACAGCCTGCCCACCCTGGAAACGGCTCAGCCGGAGGTAGGGTCCA | 14 (0.000268%) | | | TAGGTACGAAACACAGGCCCCGGAACTCATCATCGAGCGTAACATCGCCCGTGAATTAACTGAGAAGGATAGGTGGTAGGTAGTTCGATGCGCGAGCATG | 21 (0.000402%) | | | TAGTCATATGCTTGTCTCAAAGATTAAGCCATGCATGTGTAAGTATGAACGAATTCAGACTGTGAAACTGCGAATGGCTCATTAAATCAGTTATAGTTTG | 46 (0.000882%) | | | TAGTGTTCGTAGGCTCCATGCTCGCGCATCGAACTACCTACCACCTATCCTTCTCAGTTAATTCACGGGCGATGTTACGCTCGATGATGAGTTCCGGGGC | 11 (0.000211%) | | | TAGTGTTTCCTTGTTAGAAGACACAAAGCCAAAGACTCATATGGACTTTGGCTACACCATGAAAGCTTTGAGAAGCAAGAAGAAGGTTGGTTAGTGTTTT | 19 (0.000364%) | | | TAGTGTTTTGGAGTCGAATATGACTTGATGTCATGTGTAT | 277 (0.002123%) | | | TATAAGAACTTAAACCGCAACCGCATCTTATAAGCCTAAGTAGTGTTTCCTTGTTAGAAGACACAAAGCCAAAGACTCATATGGACTTTGGCTACACCAT | 28 (0.000537%) | | | TATACTCAATCATACACATGACATCAAGTCATATTCGACTCCAAAACACTAACCAACCTTCTTCTTGCTTCTCAAAGCTTTCATGGTGTAGCCAAAGTCC | 36 (0.000690%) | | | TATCAACTTTCGATGGTAGGATAGTGGCCTACCATGGTGGTAACGGGTGACGGAGAATTAGGGTTCGATTCCGGAGAGGGAGCCTGAGAAACGGCTACCA | 12 (0.000230%) | | | TATCATGAATCATCAGAGCAACGGGCAGAGCCCGCGTCGACCTTTTATCTAATAAATGCGTCCCTTCCATAAGTCGGGGTTTGTTGCACGTATTAGCTCT | 66 (0.001265%) | | | TATCCAGAGCGTAGGCTTGCTTTGAGCACTCTAATTTCTTCAAAGTAACAGCGCCGGAGGCACGACCCGGCCAATTAAGACCAGGAGCGTATCGCCGACC | 4 (0.000077%) | | | TATCCCATGCTAATGTATCCAGAGCGTAGGCTTGCTTTGAGCACTCTAATTTCTTCAAAGTAACAGCGCCGGAGGCACGACCCGGCCAATTAAGACCAGG | 3 (0.000057%) | | | TATCCTTGTTAGAAGACACAAAGCCAAAGACTCATATGGACTTTGGCTACACCATGAAAGCTTTGAGAAGCAAGAAGAAGGTTGGTTAGTGTTTTGGAGT | 24 (0.000460%) | | | TATGAACACAAACGTTCAATATGACAAACCCATGCCAAGTAAAGAGAAAATGAAAACTGGTGATTGTTGCGGAAATCGTCCAGGATTCCTCGACCAGGAC | 5 (0.000096%) | | | TATGATGTTATCCCATGCTAATGTATCCAGAGCGTAGGCTTGCTTTGAGCACTCTAATTTCTTCAAAGTAACAGCGCCGGAGGCACGACCCGGCCAATTA | 23 (0.000441%) | | | TATGATTGAGTATAAGAACTTAAACCGCAACCGCATCTTATAAGCCTAAGTAGTGTTTCCTTGTTAGAAGACACAAAGCCAAAGACTCATATGGACTTTG | 16 (0.000307%) | | | TATGCTTGTCTCAAAGATTAAGCCATGCATGTGTAAGTATGAACGAATTCAGACTGTGAAACTGCGAATGGCTCATTAAATCAGTTATAGTTTGTTTGAT | 14 (0.000268%) | | | TATTCGACTCCAAAACACTAACCAACCTTCTTCTTGCTTCTCAAAGCTTTCATGGTGTAGCCAAAGTCCATATGAGTCTTTGGCTTTGTGTCTTCTAACA | 38 (0.000728%) | | | TATTCGACTCCAAAACACTAACCAACCTTCTTCTTGCTTCTCAAAGCTTTCATGGTGTAGCCAAAGTCCATATGAGTCTTTGGCTTTGTGTCTTTTAACA | 6 (0.000115%) | | | TATTTAAGTTGTTGCAGTTAAAAAGCTCGTAGTTGAACCTTGGGATGGGTCGGCCGGTCCGCCTTTGGTGTGCATTGGTCGGCTTGTCCCTTCGGTCGGC | 4 (0.000077%) | | | TCAAAATCATCAATCGTTCCAACTAATCTACCGAAGTACTCGGCTAAGAAGAAAGAAGACGGACGAATCCGAGCCAAAGCCGTACAAGCGCGAGATACCT | 14 (0.000268%) | | | TCAAAGATTAAGCCATGCATGTGTAAGTATGAACGAATTCAGACTGTGAAACTGCGAATGGCTCATTAAATCAGTTATAGTTTGTTTGATGGTAACTACT | 3 (0.000057%) | | | TCAAATTAGGTACGAAACACAGGCCCCGGAACTCATCATCGAGCGTAACATCGCCCGTGAATTAACTGAGAAGGATAGGTGGTAGGTAGTTCGATGCGCG | 7 (0.000134%) | | | TCAAATTTCTGCCCTATCAACTTTCGATGGTAGGATAGTGGCCTACCATGGTGGTAACGGGTGACGGAGAATTAGGGTTCGATTCCGGAGAGGGAGCCTG | 12 (0.000230%) | | | TCAAGGGCCGCCGGGGGCGCACCGGACACCACGCGACGTGCGGTGCTCTTCCAGCCGCTGGACCCTACCTCCGGCTGAGCCGTTTCCAGGGTGGGCAGGC | 15 (0.000287%) | | | TCAAGTCATATTCGACTCCAAAACACTAACCAACCTTCTTCTTGCTTCTCAAAGCTTTCATGGTGTAGCCAAAGTCCATATGAGTCTTTGGCTTTGTGTC | 44 (0.000843%) | | | TCAATCATACACATGACATCAAGTCATATTCGACTCCAAAACACTAACCAACCTTCTTCTTGCTTCTCAAAGCTTTCATGGTGTAGCCAAAGTCCATATG | 30 (0.000575%) | | | TCAATCGGTAGGAGCGACGGGCGGTGTGTACAAAGGGCAGGGACGTAGTCAACGCGAGCTGATGACTCGCGCTTACTAGGAATTCCTCGTTGAAGACCAA | 63 (0.001207%) | | | TCACAAATCGTCGTCCCTCACCATCCTTTGCTGATGCGGGACGGAAGCTGGTCTCCCGTGTGTTACCGCACGCGGTTGGCCTAAATCCGAGCCAAGGACG | 4 (0.000077%) | | | TCACATTTCGTTCATCACCCTTGGCCGGCTATCGAACAGCCGGACTCCCATCAAAAGATGGTTGCCAAGAACATCTTCGTTACGGTTTGCTAATTCTCGG | 4 (0.000077%) | | | TCACATTTCGTTCATCACCCTTGGCCGGCTTTCGAACAGCCGGACTCCCATCAAAAGATGGTTGCCAAGAACATCTTCGTTACGGTTTGCTAATTCTCGG | 8 (0.000153%) | | | TCACCATCCTTTGCTGATGCGGGACGGAAGCTGGTCTCCCGTGTGTTACCGCACGCGGTTGGCCTAAATCCGAGCCAAGGACGCCTGGAGCGTACCGACA | 7 (0.000134%) | | | TCACGGTTCGTATTCGTACTGAAAATCAGAATCAAACGAGCTTTTACCCTTTTGTTCCACACGAGATTTCTGTTCTCGTTGAGCTCATCTTAGGACACCT | 4 (0.000077%) | | | TCAGCCTGCTAACTAGCTACGTGGAGGCATCCCTTCACGGCCGGCTTCTTAGAGGGACTATGGCCGTTTAGGCCAAGGAAGTTTGAGGCAATAACAGGTC | 7 (0.000134%) | | | TCAGCGTGCTCCGGGCGTCGGCCTGTGGGCTCCCCATTCGACCCGTCTTGAAACACGGACCAAGGAGTCTGACATGTGTGCGAGTCAACGGGTGAGTAAA | 5 (0.000096%) | | | TCAGCTTCCTTGCGCCTTACGGGTTTACTCACCCGTTGACTCGCACACATGTCAGACTCCTTGGTCCGTGTTTCAAGACGGGTCGAATGGGGAGCCCACA | 35 (0.000671%) | | | TCATACACATGACATCAAGTCATATTCGACTCCAAAACACTAACCAACCTTCTTCTTGCTTCTCAAAGCTTTCATGGTGTAGCCAAAGTCCATATGAGTC | 10 (0.000192%) | | | TCATATTCGACTCCAAAACACTAACCAACCTTCTTCTTGCTTCTCAAAGCTTTCATGGTGTAGCCAAAGTCCATATGAGTCTTTGGCTTTGTGTCTTCTA | 60 (0.001150%) | | | TCATCAGAGCAACGGGCAGAGCCCGCGTCGACCTTTTATCTAATAAATGCGTCCCTTCCATAAGTCGGGGTTTGTTGCACGTATTAGCTCTAGAATTACT | 3 (0.000057%) | | | TCATCATCGAGCGTAACATCGCCCGTGAATTAACTGAGAAGGATAGGTGGTAGGTAGTTCGATGCGCGAGCATGGAGCCTACGAACACTAGCTATCCGAT | 246 (0.004714%) | | | TCATGAATCATCAGAGCAACGGGCAGAGCCCGCGTCGACCTTTTATCTAATAAATGCGTCCCTTCCATAAGTCGGGGTTTGTTGCACGTATTAGCTCTAG | 26 (0.000498%) | | | TCATGTGTATGATTGAGTATAAGAACTTAAACCGCAACCGCATCTTATAAGCCTAAGTAGTGTTTCCTTGTTAGAAGACACAAAGCCAAAGACTCATATG | 11 (0.000211%) | | | TCCAACCTAGGCGAGACAAGGGTTCACATTTCGTTCATCACCCTTGGCCGGCTTTCGAACAGCCGGACTCCCATCAAAAGATGGTTGCCAAGAACATCTT | 4 (0.000077%) | | | TCCAACGAAGCACGCCCATCCAACCTAGGCGAGACAAGGGTTCACATTTCGTTCATCACCCTTGGCCGGCTATCGAACAGCCGGACTCCCATCAAAAGAT | 18 (0.000345%) | | | TCCAGAGCGTAGGCTTGCTTTGAGCACTCTAATTTCTTCAAAGTAACAGCGCCGGAGGCACGACCCGGCCAATTAAGACCAGGAGCGTATCGCCGACCGA | 5 (0.000096%) | | | TCCAGCTCCAATAGCGTATATTTAAGTTGTTGCAGTTAAAAAGCTCGTAGTTGAACCTTGGGATGGGTCGGCCGGTCCGCCTTTGGTGTGCATTGGTCGG | 53 (0.001016%) | | | TCCATGCTCGCGCATCGAACTACCTACCACCTATCCTTCTCAGTTAATTCACGGGCGATGTTACGCTCGATGATGAGTTCCGGGGCCTGTGTTTCGTACC | 6 (0.000115%) | | | TCCCATGCTAATGTATCCAGAGCGTAGGCTTGCTTTGAGCACTCTAATTTCTTCAAAGTAACAGCGCCGGAGGCACGACCCGGCCAATTAAGACCAGGAG | 4 (0.000077%) | | | TCCCGAAGGTATCTCGCGCTTGTACGGCTTTGGCTCGGATTCGTCCGTCTTCTTTCTTCTTAGCCGAGTACTTCGGTAGATTAGTTGGAACGATTGATGA | 5 (0.000096%) | | | TCCCGCCAATCAGCTTCCTTGCGCCTTACGGGTTTACTCACCCGTTGACTCGCACACATGTCAGACTCCTTGGTCCGTGTTTCAAGACGGGTCGAATGGG | 3 (0.000057%) | | | TCCCTTGCCTACATTGTTCCATCGACCAGAGGCTGTTCACCTTGGAGACCTGATGCGGTTATGAGTACGACCGGGCGTGAGCGGCACTCGGTCCTCCGGA | 5 (0.000096%) | | | TCCGCCTAGGCTGTCCCGAGTGTGAGCGAGGTGTGAGTGTCGCCCATGGGCATCGACACCTTGCGGCTAGGAACTGGAACGAGACGGGTGGCAAAGATTT | 2024 (0.038789%) | | | TCCGGAACCGGGACGTGGCGGTTGACGGCAACGTTAGGGAGTCCGGAGACGTCGGCGGGGGCCTCGGGAAGAGTTATCTTTTCTGTTTAACAGCCTGCCC | 31 (0.000594%) | | | TCCGGATTTTCAAGGGCCGCCGGGGGCGCACCGGACACCACGCGACGTGCGGTGCTCTTCCAGCCGCTGGACCCTACCTCCGGCTGAGCCGTTTCCAGGG | 3 (0.000057%) | | | TCCGGGCGTCGGCCTGTGGGCTCCCCATTCGACCCGTCTTGAAACACGGACCAAGGAGTCTGACATGTGTGCGAGTCAACGGGTGAGTAAACCCGTAAGG | 5 (0.000096%) | | | TCCGTCGAGTTATCATGAATCATCAGAGCAACGGGCAGAGCCCGCGTCGACCTTTTATCTAATAAATGCGTCCCTTCCATAAGTCGGGGTTTGTTGCACG | 13 (0.000249%) | | | TCCTATGATGTTATCCCATGCTAATGTATCCAGAGCGTAGGCTTGCTTTGAGCACTCTAATTTCTTCAAAGTAACAGCGCCGGAGGCACGACCCGGCCAA | 4 (0.000077%) | | | TCCTTGCGCCTTACGGGTTTACTCACCCGTTGACTCGCACACATGTCAGACTCCTTGGTCCGTGTTTCAAGACGGGTCGAATGGGGAGCCCACAGGCCGA | 7 (0.000134%) | | | TCCTTGTTAGAAGACACAAAGCCAAAGACTCATATGGACTTTGGCTACACCATGAAAGCTTTGAGAAGCAAGAAGAAGGTTGGTTAGTGTTTTGGAGTCG | 250 (0.004791%) | | | TCGAAATCCTATGATGTTATCCCATGCTAATGTATCCAGAGCGTAGGCTTGCTTTGAGCACTCTAATTTCTTCAAAGTAACAGCGCCGGAGGCACGACCC | 203 (0.003890%) | | | TCGACCCGCCGAAGCGAGCCTTGGGACCAAAAACAGGGGTTGTACCCCGCCTCCGATTCACGGAGTAAGTAAAATAACGTTAAAAGTAGTGGTATTTCAC | 3 (0.000057%) | | | TCGACTCCAAAACACTAACCAACCTTCTTCTTGCTTCTCAAAGCTTTCATGGTGTAGCCAAAGTCCATATGAGTCTTTGGCTTTGTGTCTTCTAACAAGG | 25 (0.000479%) | | | TCGACTCCAAAACACTAACCAACCTTCTTCTTGCTTCTCAAAGCTTTCATGGTGTAGCCAAAGTCCATATGAGTCTTTGGCTTTGTGTCTTTTAACAAGG | 11 (0.000211%) | | | TCGAGTTATCATGAATCATCAGAGCAACGGGCAGAGCCCGCGTCGACCTTTTATCTAATAAATGCGTCCCTTCCATAAGTCGGGGTTTGTTGCACGTATT | 58 (0.001112%) | | | TCGATCCATTACATTTTATCGGTCGCTCTTGTCCGGAAGCTGTAGATGACCCAAAGTCCATATAGCGACCCCAGGTCAGGCGGGATTACCCGCTGAGTTT | 7 (0.000134%) | | | TCGATCCGTCGAGTTATCATGAATCATCAGAGCAACGGGCAGAGCCCGCGTCGACCTTTTATCTAATAAATGCGTCCCTTCCATAAGTCGGGGTTTGTTG | 4 (0.000077%) | | | TCGATCGACCCGCCGAAGCGAGCCTTGGGACCAAAAACAGGGGTTGTACCCCGCCTCCGATTCACGGAGTAAGTAAAATAACGTTAAAAGTAGTGGTATT | 3 (0.000057%) | | | TCGATCTTAAAGGCGTAAGAATTGTATCCTTGTTAAAAGACACAAAGCCAAAGACTCATATGGACTTTGGCTACACCATGAAAGCTTTGAGAAGCAAGAA | 4 (0.000077%) | | | TCGATTAAGACAGCAGGACGGTGGTCATGGAAGTCGAAATCCGCTAAGGAGTGTGTAACAACTCACCTGCCGAATCAACTAGCCCCGAAAATGGATGGCG | 3 (0.000057%) | | | TCGATTAGTCTTTCGCCCCTATACCCAAGTCAGACGAACGATTTGCACGTCAGTATCGCTGCGGGCTTCCACCAGAGTTTCCTCTGGCTTTACCCCGCTC | 8 (0.000153%) | | | TCGCACACATGTCAGACTCCTTGGTCCGTGTTTCAAGACGGGTCGAATGGGGAGCCCACAGGCCGACGCCCGGAGCACGCTGATGCCGAGGCACGCCGTT | 3 (0.000057%) | | | TCGCCCATGGGCATCGACACCTTGCGGCTAGGAACTGGAACGAGACGGGTGGCAAAGATTTCGAGTAGCACTTCATACTACCGTGGGTTTTTTAAACCTT | 3 (0.000057%) | | | TCGCCGCCCGCGACGTCGCGAGAAGTCCACTAAACCTTATCATTTAGAGGAAGGAGAAGTCGTAACAAGGTTTCCGTAGGTGAACCTGCGGAAGGATCGA | 8 (0.000153%) | | | TCGGCATCAGCGTGCTCCGGGCGTCGGCCTGTGGGCTCCCCATTCGACCCGTCTTGAAACACGGACCAAGGAGTCTGACATGTGTGCGAGTCAACGGGTG | 17 (0.000326%) | | | TCGGCCTGTGGGCTCCCCATTCGACCCGTCTTGAAACACGGACCAAGGAGTCTGACATGTGTGCGAGTCAACGGGTGAGTAAACCCGTAAGGCGCAAGGA | 65 (0.001246%) | | | TCGGTAGGAGCGACGGGCGGTGTGTACAAAGGGCAGGGACGTAGTCAACGCGAGCTGATGACTCGCGCTTACTAGGAATTCCTCGTTGAAGACCAACAAT | 30 (0.000575%) | | | TCGGTCCTCCGGATTTTCAAGGGCCGCCGGGGGCGCACCGGACACCACGCGACGTGCGGTGCTCTTCCAGCCGCTGGACCCTACCTCCGGCTGAGCCGTT | 11 (0.000211%) | | | TCGTACTGAAAATCAGAATCAAACGAGCTTTTACCCTTTTGTTCCACACGAGATTTCTGTTCTCGTTGAGCTCATCTTAGGACACCTGCGTTATCTTTTA | 4 (0.000077%) | | | TCGTAGGCTCCATGCTCGCGCATCGAACTACCTACCACCTATCCTTCTCAGTTAATTCACGGGCGATGTTACGCTCGATGATGAGTTCCGGGGCCTGTGT | 7 (0.000134%) | | | TCGTATTCGTACTGAAAATCAGAATCAAACGAGCTTTTACCCTTTTGTTCCACACGAGATTTCTGTTCTCGTTGAGCTCATCTTAGGACACCTGCGTTAT | 4 (0.000077%) | | | TCGTATTTCATAGTCAGAGGTGAAATTCTTGGATTTATGAAAGACGAACAACTGCGAAAGCATTTGCCAAGGATGTTTTCATTAATCAAGAACGAAAGTT | 9 (0.000172%) | | | TCGTCCCTCACCATCCTTTGCTGATGCGGGACGGAAGCTGGTCTCCCGTGTGTTACCGCACGCGGTTGGCCTAAATCCGAGCCAAGGACGCCTGGAGCGT | 5 (0.000096%) | | | TCGTCGTCCCTCACCATCCTTTGCTGATGCGGGACGGAAGCTGGTCTCCCGTGTGTTACCGCACGCGGTTGGCCTAAATCCGAGCCAAGGACGCCTGGAG | 5 (0.000096%) | | | TCTCATGTGTATGATTGAGTATAAGAACTTAAACCGCAACCGCATCTTATAAGCCTAAGTAGTGTTTCCTTGTTAGAAGACACAAAGCCAAAGACTCATA | 7 (0.000134%) | | | TCTCGCGCTTGTACGGCTTTGGCTCGGATTCGTCCGTCTTCTTTCTTCTTAGCCGAGTACTTCGGTAGATTAGTTGGAACGATTGATGATTTTGAGTTAA | 15 (0.000287%) | | | TCTGAGAAGGGTTCGAGTGTGAGCATGCCTGTCGGGACCCGAAAGATGGTGAACTATGCCTGAGCGGGGTAAAGCCAGAGGAAACTCTGGTGGAAGCCCG | 8 (0.000153%) | | | TCTGCCCTATCAACTTTCGATGGTAGGATAGTGGCCTACCATGGTGGTAACGGGTGACGGAGAATTAGGGTTCGATTCCGGAGAGGGAGCCTGAGAAACG | 10 (0.000192%) | | | TCTGGTGCCAGCAGCCGCGGTAATTCCAGCTCCAATAGCGTATATTTAAGTTGTTGCAGTTAAAAAGCTCGTAGTTGAACCTTGGGATGGGTCGGCCGGT | 11 (0.000211%) | | | TCTGTGCTGGCGACGCATCATTCAAATTTCTGCCCTATCAACTTTCGATGGTAGGATAGTGGCCTACCATGGTGGTAACGGGTGACGGAGAATTAGGGTT | 24 (0.000460%) | | | TCTGTTAAAAGATAACGCAGGTGTCCTAAGATGAGCTCAACGAGAACAGAAATCTCGTGTGGAACAAAAGGGTAAAAGCTCGTTTGATTCTGATTTTCAG | 4 (0.000077%) | | | TCTTAAAGGCGTAAGAATTGTATCCTTGTTAGAAGACACA | 426 (0.003266%) | | | TCTTATACTCAATCATACACATGACATCAAGTCATATTCGACTCCAAAACACTAACCAACCTTCTTCTTGCTTCTCAAAGCTTTCATGGTGTAGCCAAAG | 199 (0.003814%) | | | TCTTCAACGAGGAATTCCTAGTAAGCGCGAGTCATCAGCTCGCGTTGACTACGTCCCTGCCCTTTGTACACACCGCCCGTCGCTCCTACCGATTGAATGA | 12 (0.000230%) | | | TGAACACAAACGTTCAATATGACAAACCCATGCCAAGTAAAGAGAAAATGAAAACTGGTGATTGTTGCGGAAATCGTCCAGGATTCCTCGACCAGGACTT | 19 (0.000364%) | | | TGAATCATCAGAGCAACGGGCAGAGCCCGCGTCGACCTTTTATCTAATAAATGCGTCCCTTCCATAAGTCGGGGTTTGTTGCACGTATTAGCTCTAGAAT | 7 (0.000134%) | | | TGACATCAAGTCATATTCGACTCCAAAACACTAACCAACCTTCTTCTTGCTTCTCAAAGCTTTCATGGTGTAGCCAAAGTCCATATGAGTCTTTGGCTTT | 32 (0.000613%) | | | TGACGGCAACGTTAGGGAGTCCGGAGACGTCGGCGGGGGCCTCGGGAAGAGTTATCTTTTCTGTTTAACAGCCTGCCCACCCTGGAAACGGCTCAGCCGG | 22 (0.000422%) | | | TGACTCGCACACATGTCAGACTCCTTGGTCCGTGTTTCAAGACGGGTCGAATGGGGAGCCCACAGGCCGACGCCCGGAGCACGCTGATGCCGAGGCACGC | 5 (0.000096%) | | | TGAGCGAGGTGTGAGTGTCGCCCATGGGCATCGACACCTTGCGGCTAGGAACTGGAACGAGACGGGTGGCAAAGATTTCGAGTAGCACTTCATACTACCG | 6 (0.000115%) | | | TGAGTATAAGAACTTAAACCGCAACCGCATCTTATAAGCCTAAGTAGTGTTTCCTTGTTAGAAGACACAAAGCCAAAGACTCATATGGACTTTGGCTACA | 23 (0.000441%) | | | TGAGTGTCGCCCATGGGCATCGACACCTTGCGGCTAGGAACTGGAACGAGACGGGTGGCAAAGATTTCGAGTAGCACTTCATACTACCGTGGGTTTTTTA | 7 (0.000134%) | | | TGATGTTATCCCATGCTAATGTATCCAGAGCGTAGGCTTGCTTTGAGCACTCTAATTTCTTCAAAGTAACAGCGCCGGAGGCACGACCCGGCCAATTAAG | 21 (0.000402%) | | | TGATTAACAGGGACAGTCGGGGGCATTCGTATTTCATAGTCAGAGGTGAAATTCTTGGATTTATGAAAGACGAACAACTGCGAAAGCATTTGCCAAGGAT | 126 (0.002415%) | | | TGATTGAGTATAAGAACTTAAACCGCAACCGCATCTTATAAGCCTAAGTAGTGTTTCCTTGTTAGAAGACACAAAGCCAAAGACTCATATGGACTTTGGC | 5 (0.000096%) | | | TGCAGCACGCGCCTAACGGCGTGCCTCGGCATCAGCGTGCTCCGGGCGTCGGCCTGTGGGCTCCCCATTCGACCCGTCTTGAAACACGGACCAAGGAGTC | 58 (0.001112%) | | | TGCCAGCAGCCGCGGTAATTCCAGCTCCAATAGCGTATATTTAAGTTGTTGCAGTTAAAAAGCTCGTAGTTGAACCTTGGGATGGGTCGGCCGGTCCGCC | 5 (0.000096%) | | | TGCCGACTTCCCTTGCCTACATTGTTCCATCGACCAGAGGCTGTTCACCTTGGAGACCTGATGCGGTTATGAGTACGACCGGGCGTGAGCGGCACTCGGT | 2358 (0.045190%) | | | TGCCTACATTGTTCCATCGACCAGAGGCTGTTCACCTTGGAGACCTGATGCGGTTATGAGTACGACCGGGCGTGAGCGGCACTCGGTCCTCCGGATTTTC | 8 (0.000153%) | | | TGCGCCTTACGGGTTTACTCACCCGTTGACTCGCACACATGTCAGACTCCTTGGTCCGTGTTTCAAGACGGGTCGAATGGGGAGCCCACAGGCCGACGCC | 6 (0.000115%) | | | TGCGGTTTAAGTTCTTATACTCAATCATACACATGACATCAAGTCATATTCGACTCCAAAACACTAACCAACCTTCTTCTTGCTTCTCAAAGCTTTCATG | 392 (0.007513%) | | | TGCGGTTTAAGTTGTTATACTCAATCATACACATGACAACAAGTCATATTCGACTCCAAAACACTAACCAACCTTCTTCTTGCTTCTCAAAGCTTTCATG | 92 (0.001763%) | | | TGCTAACTAGCTACGTGGAGGCATCCCTTCACGGCCGGCTTCTTAGAGGGACTATGGCCGTTTAGGCCAAGGAAGTTTGAGGCAATAACAGGTCTGTGAT | 26 (0.000498%) | | | TGCTAATGTATCCAGAGCGTAGGCTTGCTTTGAGCACTCTAATTTCTTCAAAGTAACAGCGCCGGAGGCACGACCCGGCCAATTAAGACCAGGAGCGTAT | 13 (0.000249%) | | | TGCTCCGGGCGTCGGCCTGTGGGCTCCCCATTCGACCCGTCTTGAAACACGGACCAAGGAGTCTGACATGTGTGCGAGTCAACGGGTGAGTAAACCCGTA | 5 (0.000096%) | | | TGCTGATGCGGGACGGAAGCTGGTCTCCCGTGTGTTACCGCACGCGGTTGGCCTAAATCCGAGCCAAGGACGCCTGGAGCGTACCGACATGCGGTGGTGA | 6 (0.000115%) | | | TGCTTGTCTCAAAGATTAAGCCATGCATGTGTAAGTATGAACGAATTCAGACTGTGAAACTGCGAATGGCTCATTAAATCAGTTATAGTTTGTTTGATGG | 5 (0.000096%) | | | TGCTTTGTTTTAATTAAACAGTCGGATTCCCCTTGTCCGTACCAGTTCTGAGCTGACTGTTCGACGCCCGGGGAAAGCTCCCGAGAGAGCCGTTCCCAGT | 4 (0.000077%) | | | TGGAAGTCGAAATCCGCTAAGGAGTGTGTAACAACTCACCTGCCGAATCAACTAGCCCCGAAAATGGATGGCGCTTAAGCGCGCGACCTATACCCGGCCG | 62 (0.001188%) | | | TGGAGGGCAAGTCTGGTGCCAGCAGCCGCGGTAATTCCAGCTCCAATAGCGTATATTTAAGTTGTTGCAGTTAAAAAGCTCGTAGTTGAACCTTGGGATG | 311 (0.005960%) | | | TGGCCTCTGTGCTGGCGACGCATCATTCAAATTTCTGCCCTATCAACTTTCGATGGTAGGATAGTGGCCTACCATGGTGGTAACGGGTGACGGAGAATTA | 32 (0.000613%) | | | TGGCGACGCATCATTCAAATTTCTGCCCTATCAACTTTCGATGGTAGGATAGTGGCCTACCATGGTGGTAACGGGTGACGGAGAATTAGGGTTCGATTCC | 33 (0.000632%) | | | TGGCGGTTGACGGCAACGTTAGGGAGTCCGGAGACGTCGGCGGGGGCCTCGGGAAGAGTTATCTTTTCTGTTTAACAGCCTGCCCACCCTGGAAACGGCT | 7 (0.000134%) | | | TGGCTGGGGCGGCACATCTGTTAAAAGATAACGCAGGTGTCCTAAGATGAGCTCAACGAGAACAGAAATCTCGTGTGGAACAAAAGGGTAAAAGCTCGTT | 4 (0.000077%) | | | TGGGCATCGACACCTTGCGGCTAGGAACTGGAACGAGACGGGTGGCAAAGATTTCGAGTAGCACTTCATACTACCGTGGGTTTTTTAAACCTTCCGAGTT | 18 (0.000345%) | | | TGGGGAGTTTGGCTGGGGCGGCACATCTGTTAAAAGATAACGCAGGTGTCCTAAGATGAGCTCAACGAGAACAGAAATCTCGTGTGGAACAAAAGGGTAA | 8 (0.000153%) | | | TGGGGCGGCACATCTGTTAAAAGATAACGCAGGTGTCCTAAGATGAGCTCAACGAGAACAGAAATCTCGTGTGGAACAAAAGGGTAAAAGCTCGTTTGAT | 17 (0.000326%) | | | TGGGTGGTTCGCCGCCCGCGACGTCGCGAGAAGTCCACTAAACCTTATCATTTAGAGGAAGGAGAAGTCGTAACAAGGTTTCCGTAGGTGAACCTGCGGA | 8 (0.000153%) | | | TGGGTGTCACAAATCGTCGTCCCTCACCATCCTTTGCTGATGCGGGACGGAAGCTGGTCTCCCGTGTGTTACCGCACGCGGTTGGCCTAAATCCGAGCCA | 3 (0.000057%) | | | TGGTCATGGAAGTCGAAATCCGCTAAGGAGTGTGTAACAACTCACCTGCCGAATCAACTAGCCCCGAAAATGGATGGCGCTTAAGCGCGCGACCTATACC | 17 (0.000326%) | | | TGGTCGATTAAGACAGCAGGACGGTGGTCATGGAAGTCGAAATCCGCTAAGGAGTGTGTAACAACTCACCTGCCGAATCAACTAGCCCCGAAAATGGATG | 7 (0.000134%) | | | TGGTCTTCAACGAGGAATTCCTAGTAAGCGCGAGTCATCAGCTCGCGTTGACTACGTCCCTGCCCTTTGTACACACCGCCCGTCGCTCCTACCGATTGAA | 55 (0.001054%) | | | TGGTGCCAGCAGCCGCGGTAATTCCAGCTCCAATAGCGTATATTTAAGTTGTTGCAGTTAAAAAGCTCGTAGTTGAACCTTGGGATGGGTCGGCCGGTCC | 8 (0.000153%) | | | TGGTTCGCCGCCCGCGACGTCGCGAGAAGTCCACTAAACCTTATCATTTAGAGGAAGGAGAAGTCGTAACAAGGTTTCCGTAGGTGAACCTGCGGAAGGA | 3 (0.000057%) | | | TGTAAAGTTGGGAATTCGTTAAGGAGCTGTTGCTTTGTTAGTGTAGAAACACTTGTGTAGAATTGGGGATTGTTTTTTTTGGAGTGATTTAGGGGAGGGT | 10 (0.000192%) | | | TGTATCCAGAGCGTAGGCTTGCTTTGAGCACTCTAATTTCTTCAAAGTAACAGCGCCGGAGGCACGACCCGGCCAATTAAGACCAGGAGCGTATCGCCGA | 22 (0.000422%) | | | TGTATCCTTGTTAGAAGACACAAAGCCAAAGACTCATATGGACTTTGGCTACACCATGAAAGCTTTGAGAAGCAAGAAGAAGGTTGGTTAGTGTTTTGGA | 702 (0.013454%) | | | TGTATGATTGAGTATAAGAACTTAAACCGCAACCGCATCTTATAAGCCTAAGTAGTGTTTCCTTGTTAGAAGACACAAAGCCAAAGACTCATATGGACTT | 3 (0.000057%) | | | TGTCAGGTGGGGAGTTTGGCTGGGGCGGCACATCTGTTAAAAGATAACGCAGGTGTCCTAAGATGAGCTCAACGAGAACAGAAATCTCGTGTGGAACAAA | 49 (0.000939%) | | | TGTCCCGAAGGTATCTCGCGCTTGTACGGCTTTGGCTCGGATTCGTCCGTCTTCTTTCTTCTTAGCCGAGTACTTCGGTAGATTAGTTGGAACGATTGAT | 3 (0.000057%) | | | TGTCCCGAGTGTGAGCGAGGTGTGAGTGTCGCCCATGGGCATCGACACCTTGCGGCTAGGAACTGGAACGAGACGGGTAGCAAAGATTTCGAGTAGCACT | 3 (0.000057%) | | | TGTCCCGAGTGTGAGCGAGGTGTGAGTGTCGCCCATGGGCATCGACACCTTGCGGCTAGGAACTGGAACGAGACGGGTGGCAAAGATTTCGAGTAGCACT | 5 (0.000096%) | | | TGTCGCCCATGGGCATCGACACCTTGCGGCTAGGAACTGGAACGAGACGGGTGGCAAAGATTTCGAGTAGCACTTCATACTACCGTGGGTTTTTTAAACC | 4 (0.000077%) | | | TGTCTCAAAGATTAAGCCATGCATGTGTAAGTATGAACGAATTCAGACTGTGAAACTGCGAATGGCTCATTAAATCAGTTATAGTTTGTTTGATGGTAAC | 3 (0.000057%) | | | TGTGAGCGAGGTGTGAGTGTCGCCCATGGGCATCGACACCTTGCGGCTAGGAACTGGAACGAGACGGGTGGCAAAGATTTCGAGTAGCACTTCATACTAC | 3 (0.000057%) | | | TGTGAGTGTCGCCCATGGGCATCGACACCTTGCGGCTAGGAACTGGAACGAGACGGGTAGCAAAGATTTCGAGTAGCACTTCATACTACCGTGGGTTTTT | 7 (0.000134%) | | | TGTGAGTGTCGCCCATGGGCATCGACACCTTGCGGCTAGGAACTGGAACGAGACGGGTGGCAAAGATTTCGAGTAGCACTTCATACTACCGTGGGTTTTT | 7 (0.000134%) | | | TGTGCTGGCGACGCATCATTCAAATTTCTGCCCTATCAACTTTCGATGGTAGGATAGTGGCCTACCATGGTGGTAACGGGTGACGGAGAATTAGGGTTCG | 8 (0.000153%) | | | TGTTAGAAGACACAAAGCCAAAGACTCATATGGACTTTGGCTACACCATGAAAGCTTTGAGAAGCAAGAAGAAGGTTGGTTAGTGTTTTGGAGTCGAATA | 24 (0.000460%) | | | TGTTATACTCAATCATACACATGACAACAAGTCATATTCGACTCCAAAACACTAACCAACCTTCTTCTTGCTTCTCAAAGCTTTCATGGTGTAGCCAAAG | 3 (0.000057%) | | | TGTTATCCCATGCTAATGTATCCAGAGCGTAGGCTTGCTTTGAGCACTCTAATTTCTTCAAAGTAACAGCGCCGGAGGCACGACCCGGCCAATTAAGACC | 12 (0.000230%) | | | TGTTGCTTTGTTAGTGTAGAAACACTTGTGTAGAATTGGGGATTGTTTTTTTTGGAGTGATTTAGGGGAGGGTCGAATCTTAGCGACAAAGGGCTGAATC | 15 (0.000287%) | | | TGTTGGTCGATTAAGACAGCAGGACGGTGGTCATGGAAGTCGAAATCCGCTAAGGAGTGTGTAACAACTCACCTGCCGAATCAACTAGCCCCGAAAATGG | 8 (0.000153%) | | | TGTTGGTCTTCAACGAGGAATTCCTAGTAAGCGCGAGTCATCAGCTCGCGTTGACTACGTCCCTGCCCTTTGTACACACCGCCCGTCGCTCCTACCGATT | 196 (0.003756%) | | | TGTTTTAATTAAACAGTCGGATTCCCCTTGTCCGTACCAGTTCTGAGCTGACTGTTCGACGCCCGGGGAAAGCTCCCGAGAGAGCCGTTCCCAGTCCGTC | 3 (0.000057%) | | | TGTTTTGGAGTCGAATATGACTTGATGTCATGTGTATGAT | 252 (0.001932%) | | | TTAAAAGATAACGCAGGTGTCCTAAGATGAGCTCAACGAGAACAGAAATCTCGTGTGGAACAAAAGGGTAAAAGCTCGTTTGATTCTGATTTTCAGTACG | 24 (0.000460%) | | | TTAAAGGCGTAAGAATTGTATCCTTGTTAGAAGACACAAA | 519 (0.003979%) | | | TTAACAGGGACAGTCGGGGGCATTCGTATTTCATAGTCAGAGGTGAAATTCTTGGATTTATGAAAGACGAACAACTGCGAAAGCATTTGCCAAGGATGTT | 191 (0.003660%) | | | TTAAGACAGCAGGACGGTGGTCATGGAAGTCGAAATCCGCTAAGGAGTGTGTAACAACTCACCTGCCGAATCAACTAGCCCCGAAAATGGATGGCGCTTA | 86 (0.001648%) | | | TTAAGGAGCTGTTGCTTTGTTAGTGTAGAAACACTTGTGTAGAATTGGGGATTGTTTTTTTTGGAGTGATTTAGGGGAGGGTCGAATCTTAGCGACAAAG | 15 (0.000287%) | | | TTAAGTTCTTATACTCAATCATACACATGACATCAAGTCATATTCGACTCCAAAACACTAACCAACCTTCTTCTTGCTTCTCAAAGCTTTCATGGTGTAG | 166 (0.003181%) | | | TTAAGTTGTTATACTCAATCATACACATGACAACAAGTCATATTCGACTCCAAAACACTAACCAACCTTCTTCTTGCTTCTCAAAGCTTTCATGGTGTAG | 63 (0.001207%) | | | TTACATTTTATCGGTCGCTCTTGTCCGGAAGCTGTAGATGACCCAAAGTCCATATAGCGACCCCAGGTCAGGCGGGATTACCCGCTGAGTTTAAGCATAT | 12 (0.000230%) | | | TTACGGGTTTACTCACCCGTTGACTCGCACACATGTCAGACTCCTTGGTCCGTGTTTCAAGACGGGTCGAATGGGGAGCCCACAGGCCGACGCCCGGAGC | 24 (0.000460%) | | | TTACTCACCCGTTGACTCGCACACATGTCAGACTCCTTGGTCCGTGTTTCAAGACGGGTCGAATGGGGAGCCCACAGGCCGACGCCCGGAGCACGCTGAT | 9 (0.000172%) | | | TTAGAAGACACAAAGCCAAAGACTCATATGGACTTTGGCTACACCATGAAAGCTTTGAGAAGCAAGAAGAAGGTTGGTTAGTGTTTTGGAGTCGAATATG | 73 (0.001399%) | | | TTAGGGAGTCCGGAGACGTCGGCGGGGGCCTCGGGAAGAGTTATCTTTTCTGTTTAACAGCCTGCCCACCCTGGAAACGGCTCAGCCGGAGGTAGGGTCC | 3 (0.000057%) | | | TTAGGTACGAAACACAGGCCCCGGAACTCATCATCGAGCGTAACATCGCCCGTGAATTAACTGAGAAGGATAGGTGGTAGGTAGTTCGATGCGCGAGCAT | 14 (0.000268%) | | | TTAGTGTTTTGGAGTCGAATATGACTTGATGTCATGTGTA | 340 (0.002606%) | | | TTATACTCAATCATACACATGACAACAAGTCATATTCGACTCCAAAACACTAACCAACCTTCTTCTTGCTTCTCAAAGCTTTCATGGTGTAGCCAAAGTC | 276 (0.005289%) | | | TTATACTCAATCATACACATGACATCAAGTCATATTCGACTCCAAAACACTAACCAACCTTCTTCTTGCTTCTCAAAGCTTTCATGGTGTAGCCAAAGTC | 697 (0.013358%) | | | TTATCATGAATCATCAGAGCAACGGGCAGAGCCCGCGTCGACCTTTTATCTAATAAATGCGTCCCTTCCATAAGTCGGGGTTTGTTGCACGTATTAGCTC | 55 (0.001054%) | | | TTATCCCATGCTAATGTATCCAGAGCGTAGGCTTGCTTTGAGCACTCTAATTTCTTCAAAGTAACAGCGCCGGAGGCACGACCCGGCCAATTAAGACCAG | 6 (0.000115%) | | | TTATCGGTCGCTCTTGTCCGGAAGCTGTAGATGACCCAAAGTCCATATAGCGACCCCAGGTCAGGCGGGATTACCCGCTGAGTTTAAGCATATCAATAAG | 6 (0.000115%) | | | TTCAAATTAGGTACGAAACACAGGCCCCGGAACTCATCATCGAGCGTAACATCGCCCGTGAATTAACTGAGAAGGATAGGTGGTAGGTAGTTCGATGCGC | 36 (0.000690%) | | | TTCAAATTTCTGCCCTATCAACTTTCGATGGTAGGATAGTGGCCTACCATGGTGGTAACGGGTGACGGAGAATTAGGGTTCGATTCCGGAGAGGGAGCCT | 20 (0.000383%) | | | TTCAACGAGGAATTCCTAGTAAGCGCGAGTCATCAGCTCGCGTTGACTACGTCCCTGCCCTTTGTACACACCGCCCGTCGCTCCTACCGATTGAATGATC | 7 (0.000134%) | | | TTCAAGGGCCGCCGGGGGCGCACCGGACACCACGCGACGTGCGGTGCTCTTCCAGCCGCTGGACCCTACCTCCGGCTGAGCCGTTTCCAGGGTGGGCAGG | 14 (0.000268%) | | | TTCAATCGGTAGGAGCGACGGGCGGTGTGTACAAAGGGCAGGGACGTAGTCAACGCGAGCTGATGACTCGCGCTTACTAGGAATTCCTCGTTGAAGACCA | 84 (0.001610%) | | | TTCAATGTCTGTTGGTGCCAAGAGGGAAAAGGGCTATTAAGCTATATAGGGGGGTGGGTGTTGAGGGAGTCTGGGCAGTCCGTGGGGAACCCCCTTTTTC | 469 (0.008988%) | | | TTCACATTTCGTTCATCACCCTTGGCCGGCTATCGAACAGCCGGACTCCCATCAAAAGATGGTTGCCAAGAACATCTTCGTTACGGTTTGCTAATTCTCG | 17 (0.000326%) | | | TTCACATTTCGTTCATCACCCTTGGCCGGCTTTCGAACAGCCGGACTCCCATCAAAAGATGGTTGCCAAGAACATCTTCGTTACGGTTTGCTAATTCTCG | 17 (0.000326%) | | | TTCACGGTTCGTATTCGTACTGAAAATCAGAATCAAACGAGCTTTTACCCTTTTGTTCCACACGAGATTTCTGTTCTCGTTGAGCTCATCTTAGGACACC | 14 (0.000268%) | | | TTCAGCAGTTCTCGGACAAAAATTGCTGAGTGGCCGAGAAGAATGGGCGTGTCATGCGTGGGCTGACATGGATTCTTCGAGGCCTAGGGGTGGCGGTATA | 8 (0.000153%) | | | TTCATCACCCTTGGCCGGCTTTCGAACAGCCGGACTCCCATCAAAAGATGGTTGCCAAGAACATCTTCGTTACGGTTTGCTAATTCTCGGAATAACATCA | 3 (0.000057%) | | | TTCCAACGAAGCACGCCCATCCAACCTAGGCGAGACAAGGGTTCACATTTCGTTCATCACCCTTGGCCGGCTATCGAACAGCCGGACTCCCATCAAAAGA | 105 (0.002012%) | | | TTCCCTTGCCTACATTGTTCCATCGACCAGAGGCTGTTCACCTTGGAGACCTGATGCGGTTATGAGTACGACCGGGCGTGAGCGGCACTCGGTCCTCCGG | 5 (0.000096%) | | | TTCCGGAACCGGGACGTGGCGGTTGACGGCAACGTTAGGGAGTCCGGAGACGTCGGCGGGGGCCTCGGGAAGAGTTATCTTTTCTGTTTAACAGCCTGCC | 2365 (0.045324%) | | | TTCCTTGCGCCTTACGGGTTTACTCACCCGTTGACTCGCACACATGTCAGACTCCTTGGTCCGTGTTTCAAGACGGGTCGAATGGGGAGCCCACAGGCCG | 8 (0.000153%) | | | TTCGACTCCAAAACACTAACCAACCTTCTTCTTGCTTCTCAAAGCTTTCATGGTGTAGCCAAAGTCCATATGAGTCTTTGGCTTTGTGTCTTCTAACAAG | 99 (0.001897%) | | | TTCGACTCCAAAACACTAACCAACCTTCTTCTTGCTTCTCAAAGCTTTCATGGTGTAGCCAAAGTCCATATGAGTCTTTGGCTTTGTGTCTTTTAACAAG | 85 (0.001629%) | | | TTCGAGTGTGAGCATGCCTGTCGGGACCCGAAAGATGGTGAACTATGCCTGAGCGGGGTAAAGCCAGAGGAAACTCTGGTGGAAGCCCGCAGCGATACTG | 3 (0.000057%) | | | TTCGATTAGTCTTTCGCCCCTATACCCAAGTCAGACGAACGATTTGCACGTCAGTATCGCTGCGGGCTTCCACCAGAGTTTCCTCTGGCTTTACCCCGCT | 21 (0.000402%) | | | TTCGCCGCCCGCGACGTCGCGAGAAGTCCACTAAACCTTATCATTTAGAGGAAGGAGAAGTCGTAACAAGGTTTCCGTAGGTGAACCTGCGGAAGGATCG | 3 (0.000057%) | | | TTCGTACTGAAAATCAGAATCAAACGAGCTTTTACCCTTTTGTTCCACACGAGATTTCTGTTCTCGTTGAGCTCATCTTAGGACACCTGCGTTATCTTTT | 18 (0.000345%) | | | TTCGTAGGCTCCATGCTCGCGCATCGAACTACCTACCACCTATCCTTCTCAGTTAATTCACGGGCGATGTTACGCTCGATGATGAGTTCCGGGGCCTGTG | 9 (0.000172%) | | | TTCGTATTCGTACTGAAAATCAGAATCAAACGAGCTTTTACCCTTTTGTTCCACACGAGATTTCTGTTCTCGTTGAGCTCATCTTAGGACACCTGCGTTA | 16 (0.000307%) | | | TTCGTATTTCATAGTCAGAGGTGAAATTCTTGGATTTATGAAAGACGAACAACTGCGAAAGCATTTGCCAAGGATGTTTTCATTAATCAAGAACGAAAGT | 22 (0.000422%) | | | TTCGTTCATCACCCTTGGCCGGCTATCGAACAGCCGGACTCCCATCAAAAGATGGTTGCCAAGAACATCTTCGTTACGGTTTGCTAATTCTCGGAATAAC | 12 (0.000230%) | | | TTCGTTCATCACCCTTGGCCGGCTTTCGAACAGCCGGACTCCCATCAAAAGATGGTTGCCAAGAACATCTTCGTTACGGTTTGCTAATTCTCGGAATAAC | 11 (0.000211%) | | | TTCTGAGAAGGGTTCGAGTGTGAGCATGCCTGTCGGGACCCGAAAGATGGTGAACTATGCCTGAGCGGGGTAAAGCCAGAGGAAACTCTGGTGGAAGCCC | 10 (0.000192%) | | | TTCTGCCCTATCAACTTTCGATGGTAGGATAGTGGCCTACCATGGTGGTAACGGGTGACGGAGAATTAGGGTTCGATTCCGGAGAGGGAGCCTGAGAAAC | 24 (0.000460%) | | | TTCTTATACTCAATCATACACATGACATCAAGTCATATTCGACTCCAAAACACTAACCAACCTTCTTCTTGCTTCTCAAAGCTTTCATGGTGTAGCCAAA | 76 (0.001457%) | | | TTGAAATCGTCGACCAGGTCCGAGACTTCATCGACCGGGTCCGAGGATTCGTCGACCAGGACGGCCGGATGTCCGAGAAAAAAAAATGTTGCCGAATAAC | 46 (0.000882%) | | | TTGACGGCAACGTTAGGGAGTCCGGAGACGTCGGCGGGGGCCTCGGGAAGAGTTATCTTTTCTGTTTAACAGCCTGCCCACCCTGGAAACGGCTCAGCCG | 10 (0.000192%) | | | TTGACTCGCACACATGTCAGACTCCTTGGTCCGTGTTTCAAGACGGGTCGAATGGGGAGCCCACAGGCCGACGCCCGGAGCACGCTGATGCCGAGGCACG | 15 (0.000287%) | | | TTGAGTATAAGAACTTAAACCGCAACCGCATCTTATAAGCCTAAGTAGTGTTTCCTTGTTAGAAGACACAAAGCCAAAGACTCATATGGACTTTGGCTAC | 8 (0.000153%) | | | TTGCGCCTTACGGGTTTACTCACCCGTTGACTCGCACACATGTCAGACTCCTTGGTCCGTGTTTCAAGACGGGTCGAATGGGGAGCCCACAGGCCGACGC | 20 (0.000383%) | | | TTGCGGTTTAAGTTCTTATACTCAATCATACACATGACATCAAGTCATATTCGACTCCAAAACACTAACCAACCTTCTTCTTGCTTCTCAAAGCTTTCAT | 444 (0.008509%) | | | TTGCGGTTTAAGTTGTTATACTCAATCATACACATGACAACAAGTCATATTCGACTCCAAAACACTAACCAACCTTCTTCTTGCTTCTCAAAGCTTTCAT | 85 (0.001629%) | | | TTGCTGATGCGGGACGGAAGCTGGTCTCCCGTGTGTTACCGCACGCGGTTGGCCTAAATCCGAGCCAAGGACGCCTGGAGCGTACCGACATGCGGTGGTG | 5 (0.000096%) | | | TTGGAGACCTGATGCGGTTATGAGTACGACCGGGCGTGAGCGGCACTCGGTCCTCCGGATTTTCAAGGGCCGCCGGGGGCGCACCGGACACCACGCGACG | 22 (0.000422%) | | | TTGGAGGGCAAGTCTGGTGCCAGCAGCCGCGGTAATTCCAGCTCCAATAGCGTATATTTAAGTTGTTGCAGTTAAAAAGCTCGTAGTTGAACCTTGGGAT | 314 (0.006018%) | | | TTGGAGTCGAATATGACTTGATGTCATGTGTATGATTGAG | 274 (0.002100%) | | | TTGGCTCGGATTCGTCCGTCTTCTTTCTTCTTAGCCGAGTACTTCGGTAGATTAGTTGGAACGATTGATGATTTTGAGTTAATTGAACGTTCGGCGTATG | 6 (0.000115%) | | | TTGGCTGGGGCGGCACATCTGTTAAAAGATAACGCAGGTGTCCTAAGATGAGCTCAACGAGAACAGAAATCTCGTGTGGAACAAAAGGGTAAAAGCTCGT | 19 (0.000364%) | | | TTGGTCGATTAAGACAGCAGGACGGTGGTCATGGAAGTCGAAATCCGCTAAGGAGTGTGTAACAACTCACCTGCCGAATCAACTAGCCCCGAAAATGGAT | 7 (0.000134%) | | | TTGGTCTTCAACGAGGAATTCCTAGTAAGCGCGAGTCATCAGCTCGCGTTGACTACGTCCCTGCCCTTTGTACACACCGCCCGTCGCTCCTACCGATTGA | 110 (0.002108%) | | | TTGGTGATATGAACACAAACGTTCAATATGACAAACCCATGCCAAGTAAAGAGAAAATGAAAACTGGTGATTGTTGCGGAAATCGTCCAGGATTCCTCGA | 6 (0.000115%) | | | TTGTACGGCTTTGGCTCGGATTCGTCCGTCTTCTTTCTTCTTAGCCGAGTACTTCGGTAGATTAGTTGGAACGATTGATGATTTTGAGTTAATTGAACGT | 6 (0.000115%) | | | TTGTATCCTTGTTAGAAGACACAAAGCCAAAGACTCATATGGACTTTGGCTACACCATGAAAGCTTTGAGAAGCAAGAAGAAGGTTGGTTAGTGTTTTGG | 11 (0.000211%) | | | TTGTCAGGTGGGGAGTTTGGCTGGGGCGGCACATCTGTTAAAAGATAACGCAGGTGTCCTAAGATGAGCTCAACGAGAACAGAAATCTCGTGTGGAACAA | 17 (0.000326%) | | | TTGTCTCAAAGATTAAGCCATGCATGTGTAAGTATGAACGAATTCAGACTGTGAAACTGCGAATGGCTCATTAAATCAGTTATAGTTTGTTTGATGGTAA | 15 (0.000287%) | | | TTGTTAGAAGACACAAAGCCAAAGACTCATATGGACTTTGGCTACACCATGAAAGCTTTGAGAAGCAAGAAGAAGGTTGGTTAGTGTTTTGGAGTCGAAT | 53 (0.001016%) | | | TTGTTATACTCAATCATACACATGACAACAAGTCATATTCGACTCCAAAACACTAACCAACCTTCTTCTTGCTTCTCAAAGCTTTCATGGTGTAGCCAAA | 58 (0.001112%) | | | TTGTTGGTCTTCAACGAGGAATTCCTAGTAAGCGCGAGTCATCAGCTCGCGTTGACTACGTCCCTGCCCTTTGTACACACCGCCCGTCGCTCCTACCGAT | 187 (0.003584%) | | | TTGTTTTAATTAAACAGTCGGATTCCCCTTGTCCGTACCAGTTCTGAGCTGACTGTTCGACGCCCGGGGAAAGCTCCCGAGAGAGCCGTTCCCAGTCCGT | 12 (0.000230%) | | | TTTAAGTTCTTATACTCAATCATACACATGACATCAAGTCATATTCGACTCCAAAACACTAACCAACCTTCTTCTTGCTTCTCAAAGCTTTCATGGTGTA | 105 (0.002012%) | | | TTTAAGTTGTTATACTCAATCATACACATGACAACAAGTCATATTCGACTCCAAAACACTAACCAACCTTCTTCTTGCTTCTCAAAGCTTTCATGGTGTA | 26 (0.000498%) | | | TTTAATTAAACAGTCGGATTCCCCTTGTCCGTACCAGTTCTGAGCTGACTGTTCGACGCCCGGGGAAAGCTCCCGAGAGAGCCGTTCCCAGTCCGTCCCC | 14 (0.000268%) | | | TTTACTCACCCGTTGACTCGCACACATGTCAGACTCCTTGGTCCGTGTTTCAAGACGGGTCGAATGGGGAGCCCACAGGCCGACGCCCGGAGCACGCTGA | 11 (0.000211%) | | | TTTATCGGTCGCTCTTGTCCGGAAGCTGTAGATGACCCAAAGTCCATATAGCGACCCCAGGTCAGGCGGGATTACCCGCTGAGTTTAAGCATATCAATAA | 12 (0.000230%) | | | TTTCAAGGGCCGCCGGGGGCGCACCGGACACCACGCGACGTGCGGTGCTCTTCCAGCCGCTGGACCCTACCTCCGGCTGAGCCGTTTCCAGGGTGGGCAG | 22 (0.000422%) | | | TTTCACGGTTCGTATTCGTACTGAAAATCAGAATCAAACGAGCTTTTACCCTTTTGTTCCACACGAGATTTCTGTTCTCGTTGAGCTCATCTTAGGACAC | 51 (0.000977%) | | | TTTCAGCAGTTCTCGGACAAAAATTGCTGAGTGGCCGAGAAGAATGGGCGTGTCATGCGTGGGCTGACATGGATTCTTCGAGGCCTAGGGGTGGCGGTAT | 5 (0.000096%) | | | TTTCCAACGAAGCACGCCCATCCAACCTAGGCGAGACAAGGGTTCACATTTCGTTCATCACCCTTGGCCGGCTATCGAACAGCCGGACTCCCATCAAAAG | 1817 (0.034822%) | | | TTTCGTTCATCACCCTTGGCCGGCTATCGAACAGCCGGACTCCCATCAAAAGATGGTTGCCAAGAACATCTTCGTTACGGTTTGCTAATTCTCGGAATAA | 12 (0.000230%) | | | TTTCGTTCATCACCCTTGGCCGGCTTTCGAACAGCCGGACTCCCATCAAAAGATGGTTGCCAAGAACATCTTCGTTACGGTTTGCTAATTCTCGGAATAA | 9 (0.000172%) | | | TTTCTGCCCTATCAACTTTCGATGGTAGGATAGTGGCCTACCATGGTGGTAACGGGTGACGGAGAATTAGGGTTCGATTCCGGAGAGGGAGCCTGAGAAA | 20 (0.000383%) | | | TTTGCTGATGCGGGACGGAAGCTGGTCTCCCGTGTGTTACCGCACGCGGTTGGCCTAAATCCGAGCCAAGGACGCCTGGAGCGTACCGACATGCGGTGGT | 32 (0.000613%) | | | TTTGGAGTCGAATATGACTTGATGTCATGTGTATGATTGA | 327 (0.002507%) | | | TTTGGCTCGGATTCGTCCGTCTTCTTTCTTCTTAGCCGAGTACTTCGGTAGATTAGTTGGAACGATTGATGATTTTGAGTTAATTGAACGTTCGGCGTAT | 26 (0.000498%) | | | TTTGGCTGGGGCGGCACATCTGTTAAAAGATAACGCAGGTGTCCTAAGATGAGCTCAACGAGAACAGAAATCTCGTGTGGAACAAAAGGGTAAAAGCTCG | 16 (0.000307%) | | | TTTGGCTTTGTGTCTTCTAACAAGGAAACACTACTTAGGC | 6775 (0.051936%) | | | TTTGTTTTAATTAAACAGTCGGATTCCCCTTGTCCGTACCAGTTCTGAGCTGACTGTTCGACGCCCGGGGAAAGCTCCCGAGAGAGCCGTTCCCAGTCCG | 30 (0.000575%) | | | TTTTAATTAAACAGTCGGATTCCCCTTGTCCGTACCAGTTCTGAGCTGACTGTTCGACGCCCGGGGAAAGCTCCCGAGAGAGCCGTTCCCAGTCCGTCCC | 13 (0.000249%) | | | TTTTATCGGTCGCTCTTGTCCGGAAGCTGTAGATGACCCAAAGTCCATATAGCGACCCCAGGTCAGGCGGGATTACCCGCTGAGTTTAAGCATATCAATA | 8 (0.000153%) | | | TTTTCAAGGGCCGCCGGGGGCGCACCGGACACCACGCGACGTGCGGTGCTCTTCCAGCCGCTGGACCCTACCTCCGGCTGAGCCGTTTCCAGGGTGGGCA | 23 (0.000441%) | | | TTTTCAGCAGTTCTCGGACAAAAATTGCTGAGTGGCCGAGAAGAATGGGCGTGTCATGCGTGGGCTGACATGGATTCTTCGAGGCCTAGGGGTGGCGGTA | 4 (0.000077%) | | | TTTTGGAGTCGAATATGACTTGATGTCATGTGTATGATTG | 480 (0.003680%) | | | TTTTTCAGCAGTTCTCGGACAAAAATTGCTGAGTGGCCGAGAAGAATGGGCGTGTCATGCGTGGGCTGACATGGATTCTTCGAGGCCTAGGGGTGGCGGT | 21 (0.000402%) | | | TTTTTTCAGCAGTTCTCGGACAAAAATTGCTGAGTGGCCGAGAAGAATGGGCGTGTCATGCGTGGGCTGACATGGATTCTTCGAGGCCTAGGGGTGGCGG | 3 (0.000057%) | | | TTTTTTTTTT | 52066 (0.099782%) | |

fastp -p -i resources/raw\_hic/CRR302669\_1.fastq.gz -I resources/raw\_hic/CRR302669\_2.fastq.gz -o results/fastp/hic\_trim\_1.fastq.gz -O results/fastp/hic\_trim\_2.fastq.gz --detect\_adapter\_for\_pe --json results/fastp/hic\_report\_fastp.HiC.json --html results/fastp/hic\_report\_fastp.HiC.html --thread 20

fastp 0.23.4, at 2024-04-29 23:10:53
