## Additional files for "Colora: A Snakemake Workflow for Complete Chromosome-scale *De Novo* Genome Assembly": S9_fastp_M.domestica.html

fastp report at 2024-05-07 10:48:51 

### fastp report Summary General | | | | --- | --- | | fastp version: | 0.23.4 (https://github.com/OpenGene/fastp) | | sequencing: | paired end (150 cycles + 150 cycles) | | mean length before filtering: | 148bp, 148bp | | mean length after filtering: | 148bp, 148bp | | duplication rate: | 15.932978% | | Insert size peak: | 269 | Before filtering | | | | --- | --- | | total reads: | 676.416866 M | | total bases: | 100.522162 G | | Q20 bases: | 97.141880 G (96.637277%) | | Q30 bases: | 91.747105 G (91.270525%) | | GC content: | 38.625093% | After filtering | | | | --- | --- | | total reads: | 670.577270 M | | total bases: | 99.631921 G | | Q20 bases: | 96.542553 G (96.899218%) | | Q30 bases: | 91.267955 G (91.605134%) | | GC content: | 38.614515% | Filtering result | | | | --- | --- | | reads passed filters: | 670.577270 M (99.136687%) | | reads with low quality: | 5.839586 M (0.863312%) | | reads with too many N: | 10 (0.000001%) | | reads too short: | 0 (0.000000%) | Adapters Adapter or bad ligation of read1 The input has little adapter percentage (~0.021127%), probably it's trimmed before. | | | | --- | --- | | Sequence | Occurrences | | A | 13923 | | AG | 10829 | | AGA | 10090 | | AGAT | 9126 | | AGATC | 7589 | | AGATCG | 6282 | | AGATCGG | 5910 | | AGATCGGA | 5459 | | AGATCGGAA | 4996 | | AGATCGGAAG | 4442 | | other adapter sequences | 350838 | Adapter or bad ligation of read2 The input has little adapter percentage (~0.019260%), probably it's trimmed before. | | | | --- | --- | | Sequence | Occurrences | | A | 14491 | | AG | 12850 | | AGA | 12305 | | AGAT | 11612 | | AGATC | 10336 | | AGATCG | 9425 | | AGATCGG | 9149 | | AGATCGGA | 8601 | | AGATCGGAA | 8016 | | AGATCGGAAG | 7382 | | AGATCGGAAGA | 7323 | | AGATCGGAAGAG | 6966 | | AGATCGGAAGAGC | 6487 | | AGATCGGAAGAGCG | 5979 | | AGATCGGAAGAGCGT | 5416 | | AGATCGGAAGAGCGTC | 6472 | | AGATCGGAAGAGCGTCG | 4983 | | AGATCGGAAGAGCGTCGT | 5269 | | AGATCGGAAGAGCGTCGTGTA | 5089 | | AGATCGGAAGAGCGTCGTGTAGGGAAA | 5615 | | other adapter sequences | 266342 | Insert size estimation This estimation is based on paired-end overlap analysis, and there are 65.406428% reads found not overlapped. The nonoverlapped read pairs may have insert size <30 or >270, or contain too much sequencing errors to be detected as overlapped. Before filtering Before filtering: read1: quality Value of each position will be shown on mouse over. Before filtering: read1: base contents Value of each position will be shown on mouse over. Before filtering: read1: KMER counting Darker background means larger counts. The count will be shown on mouse over. | | | | | | | | | | | | | | | | | | | --- | --- | --- | --- | --- | --- | --- | --- | --- | --- | --- | --- | --- | --- | --- | --- | --- | | | AA | AT | AC | AG | TA | TT | TC | TG | CA | CT | CC | CG | GA | GT | GC | GG | | AAA | AAAAA | AAAAT | AAAAC | AAAAG | AAATA | AAATT | AAATC | AAATG | AAACA | AAACT | AAACC | AAACG | AAAGA | AAAGT | AAAGC | AAAGG | | AAT | AATAA | AATAT | AATAC | AATAG | AATTA | AATTT | AATTC | AATTG | AATCA | AATCT | AATCC | AATCG | AATGA | AATGT | AATGC | AATGG | | AAC | AACAA | AACAT | AACAC | AACAG | AACTA | AACTT | AACTC | AACTG | AACCA | AACCT | AACCC | AACCG | AACGA | AACGT | AACGC | AACGG | | AAG | AAGAA | AAGAT | AAGAC | AAGAG | AAGTA | AAGTT | AAGTC | AAGTG | AAGCA | AAGCT | AAGCC | AAGCG | AAGGA | AAGGT | AAGGC | AAGGG | | ATA | ATAAA | ATAAT | ATAAC | ATAAG | ATATA | ATATT | ATATC | ATATG | ATACA | ATACT | ATACC | ATACG | ATAGA | ATAGT | ATAGC | ATAGG | | ATT | ATTAA | ATTAT | ATTAC | ATTAG | ATTTA | ATTTT | ATTTC | ATTTG | ATTCA | ATTCT | ATTCC | ATTCG | ATTGA | ATTGT | ATTGC | ATTGG | | ATC | ATCAA | ATCAT | ATCAC | ATCAG | ATCTA | ATCTT | ATCTC | ATCTG | ATCCA | ATCCT | ATCCC | ATCCG | ATCGA | ATCGT | ATCGC | ATCGG | | ATG | ATGAA | ATGAT | ATGAC | ATGAG | ATGTA | ATGTT | ATGTC | ATGTG | ATGCA | ATGCT | ATGCC | ATGCG | ATGGA | ATGGT | ATGGC | ATGGG | | ACA | ACAAA | ACAAT | ACAAC | ACAAG | ACATA | ACATT | ACATC | ACATG | ACACA | ACACT | ACACC | ACACG | ACAGA | ACAGT | ACAGC | ACAGG | | ACT | ACTAA | ACTAT | ACTAC | ACTAG | ACTTA | ACTTT | ACTTC | ACTTG | ACTCA | ACTCT | ACTCC | ACTCG | ACTGA | ACTGT | ACTGC | ACTGG | | ACC | ACCAA | ACCAT | ACCAC | ACCAG | ACCTA | ACCTT | ACCTC | ACCTG | ACCCA | ACCCT | ACCCC | ACCCG | ACCGA | ACCGT | ACCGC | ACCGG | | ACG | ACGAA | ACGAT | ACGAC | ACGAG | ACGTA | ACGTT | ACGTC | ACGTG | ACGCA | ACGCT | ACGCC | ACGCG | ACGGA | ACGGT | ACGGC | ACGGG | | AGA | AGAAA | AGAAT | AGAAC | AGAAG | AGATA | AGATT | AGATC | AGATG | AGACA | AGACT | AGACC | AGACG | AGAGA | AGAGT | AGAGC | AGAGG | | AGT | AGTAA | AGTAT | AGTAC | AGTAG | AGTTA | AGTTT | AGTTC | AGTTG | AGTCA | AGTCT | AGTCC | AGTCG | AGTGA | AGTGT | AGTGC | AGTGG | | AGC | AGCAA | AGCAT | AGCAC | AGCAG | AGCTA | AGCTT | AGCTC | AGCTG | AGCCA | AGCCT | AGCCC | AGCCG | AGCGA | AGCGT | AGCGC | AGCGG | | AGG | AGGAA | AGGAT | AGGAC | AGGAG | AGGTA | AGGTT | AGGTC | AGGTG | AGGCA | AGGCT | AGGCC | AGGCG | AGGGA | AGGGT | AGGGC | AGGGG | | TAA | TAAAA | TAAAT | TAAAC | TAAAG | TAATA | TAATT | TAATC | TAATG | TAACA | TAACT | TAACC | TAACG | TAAGA | TAAGT | TAAGC | TAAGG | | TAT | TATAA | TATAT | TATAC | TATAG | TATTA | TATTT | TATTC | TATTG | TATCA | TATCT | TATCC | TATCG | TATGA | TATGT | TATGC | TATGG | | TAC | TACAA | TACAT | TACAC | TACAG | TACTA | TACTT | TACTC | TACTG | TACCA | TACCT | TACCC | TACCG | TACGA | TACGT | TACGC | TACGG | | TAG | TAGAA | TAGAT | TAGAC | TAGAG | TAGTA | TAGTT | TAGTC | TAGTG | TAGCA | TAGCT | TAGCC | TAGCG | TAGGA | TAGGT | TAGGC | TAGGG | | TTA | TTAAA | TTAAT | TTAAC | TTAAG | TTATA | TTATT | TTATC | TTATG | TTACA | TTACT | TTACC | TTACG | TTAGA | TTAGT | TTAGC | TTAGG | | TTT | TTTAA | TTTAT | TTTAC | TTTAG | TTTTA | TTTTT | TTTTC | TTTTG | TTTCA | TTTCT | TTTCC | TTTCG | TTTGA | TTTGT | TTTGC | TTTGG | | TTC | TTCAA | TTCAT | TTCAC | TTCAG | TTCTA | TTCTT | TTCTC | TTCTG | TTCCA | TTCCT | TTCCC | TTCCG | TTCGA | TTCGT | TTCGC | TTCGG | | TTG | TTGAA | TTGAT | TTGAC | TTGAG | TTGTA | TTGTT | TTGTC | TTGTG | TTGCA | TTGCT | TTGCC | TTGCG | TTGGA | TTGGT | TTGGC | TTGGG | | TCA | TCAAA | TCAAT | TCAAC | TCAAG | TCATA | TCATT | TCATC | TCATG | TCACA | TCACT | TCACC | TCACG | TCAGA | TCAGT | TCAGC | TCAGG | | TCT | TCTAA | TCTAT | TCTAC | TCTAG | TCTTA | TCTTT | TCTTC | TCTTG | TCTCA | TCTCT | TCTCC | TCTCG | TCTGA | TCTGT | TCTGC | TCTGG | | TCC | TCCAA | TCCAT | TCCAC | TCCAG | TCCTA | TCCTT | TCCTC | TCCTG | TCCCA | TCCCT | TCCCC | TCCCG | TCCGA | TCCGT | TCCGC | TCCGG | | TCG | TCGAA | TCGAT | TCGAC | TCGAG | TCGTA | TCGTT | TCGTC | TCGTG | TCGCA | TCGCT | TCGCC | TCGCG | TCGGA | TCGGT | TCGGC | TCGGG | | TGA | TGAAA | TGAAT | TGAAC | TGAAG | TGATA | TGATT | TGATC | TGATG | TGACA | TGACT | TGACC | TGACG | TGAGA | TGAGT | TGAGC | TGAGG | | TGT | TGTAA | TGTAT | TGTAC | TGTAG | TGTTA | TGTTT | TGTTC | TGTTG | TGTCA | TGTCT | TGTCC | TGTCG | TGTGA | TGTGT | TGTGC | TGTGG | | TGC | TGCAA | TGCAT | TGCAC | TGCAG | TGCTA | TGCTT | TGCTC | TGCTG | TGCCA | TGCCT | TGCCC | TGCCG | TGCGA | TGCGT | TGCGC | TGCGG | | TGG | TGGAA | TGGAT | TGGAC | TGGAG | TGGTA | TGGTT | TGGTC | TGGTG | TGGCA | TGGCT | TGGCC | TGGCG | TGGGA | TGGGT | TGGGC | TGGGG | | CAA | CAAAA | CAAAT | CAAAC | CAAAG | CAATA | CAATT | CAATC | CAATG | CAACA | CAACT | CAACC | CAACG | CAAGA | CAAGT | CAAGC | CAAGG | | CAT | CATAA | CATAT | CATAC | CATAG | CATTA | CATTT | CATTC | CATTG | CATCA | CATCT | CATCC | CATCG | CATGA | CATGT | CATGC | CATGG | | CAC | CACAA | CACAT | CACAC | CACAG | CACTA | CACTT | CACTC | CACTG | CACCA | CACCT | CACCC | CACCG | CACGA | CACGT | CACGC | CACGG | | CAG | CAGAA | CAGAT | CAGAC | CAGAG | CAGTA | CAGTT | CAGTC | CAGTG | CAGCA | CAGCT | CAGCC | CAGCG | CAGGA | CAGGT | CAGGC | CAGGG | | CTA | CTAAA | CTAAT | CTAAC | CTAAG | CTATA | CTATT | CTATC | CTATG | CTACA | CTACT | CTACC | CTACG | CTAGA | CTAGT | CTAGC | CTAGG | | CTT | CTTAA | CTTAT | CTTAC | CTTAG | CTTTA | CTTTT | CTTTC | CTTTG | CTTCA | CTTCT | CTTCC | CTTCG | CTTGA | CTTGT | CTTGC | CTTGG | | CTC | CTCAA | CTCAT | CTCAC | CTCAG | CTCTA | CTCTT | CTCTC | CTCTG | CTCCA | CTCCT | CTCCC | CTCCG | CTCGA | CTCGT | CTCGC | CTCGG | | CTG | CTGAA | CTGAT | CTGAC | CTGAG | CTGTA | CTGTT | CTGTC | CTGTG | CTGCA | CTGCT | CTGCC | CTGCG | CTGGA | CTGGT | CTGGC | CTGGG | | CCA | CCAAA | CCAAT | CCAAC | CCAAG | CCATA | CCATT | CCATC | CCATG | CCACA | CCACT | CCACC | CCACG | CCAGA | CCAGT | CCAGC | CCAGG | | CCT | CCTAA | CCTAT | CCTAC | CCTAG | CCTTA | CCTTT | CCTTC | CCTTG | CCTCA | CCTCT | CCTCC | CCTCG | CCTGA | CCTGT | CCTGC | CCTGG | | CCC | CCCAA | CCCAT | CCCAC | CCCAG | CCCTA | CCCTT | CCCTC | CCCTG | CCCCA | CCCCT | CCCCC | CCCCG | CCCGA | CCCGT | CCCGC | CCCGG | | CCG | CCGAA | CCGAT | CCGAC | CCGAG | CCGTA | CCGTT | CCGTC | CCGTG | CCGCA | CCGCT | CCGCC | CCGCG | CCGGA | CCGGT | CCGGC | CCGGG | | CGA | CGAAA | CGAAT | CGAAC | CGAAG | CGATA | CGATT | CGATC | CGATG | CGACA | CGACT | CGACC | CGACG | CGAGA | CGAGT | CGAGC | CGAGG | | CGT | CGTAA | CGTAT | CGTAC | CGTAG | CGTTA | CGTTT | CGTTC | CGTTG | CGTCA | CGTCT | CGTCC | CGTCG | CGTGA | CGTGT | CGTGC | CGTGG | | CGC | CGCAA | CGCAT | CGCAC | CGCAG | CGCTA | CGCTT | CGCTC | CGCTG | CGCCA | CGCCT | CGCCC | CGCCG | CGCGA | CGCGT | CGCGC | CGCGG | | CGG | CGGAA | CGGAT | CGGAC | CGGAG | CGGTA | CGGTT | CGGTC | CGGTG | CGGCA | CGGCT | CGGCC | CGGCG | CGGGA | CGGGT | CGGGC | CGGGG | | GAA | GAAAA | GAAAT | GAAAC | GAAAG | GAATA | GAATT | GAATC | GAATG | GAACA | GAACT | GAACC | GAACG | GAAGA | GAAGT | GAAGC | GAAGG | | GAT | GATAA | GATAT | GATAC | GATAG | GATTA | GATTT | GATTC | GATTG | GATCA | GATCT | GATCC | GATCG | GATGA | GATGT | GATGC | GATGG | | GAC | GACAA | GACAT | GACAC | GACAG | GACTA | GACTT | GACTC | GACTG | GACCA | GACCT | GACCC | GACCG | GACGA | GACGT | GACGC | GACGG | | GAG | GAGAA | GAGAT | GAGAC | GAGAG | GAGTA | GAGTT | GAGTC | GAGTG | GAGCA | GAGCT | GAGCC | GAGCG | GAGGA | GAGGT | GAGGC | GAGGG | | GTA | GTAAA | GTAAT | GTAAC | GTAAG | GTATA | GTATT | GTATC | GTATG | GTACA | GTACT | GTACC | GTACG | GTAGA | GTAGT | GTAGC | GTAGG | | GTT | GTTAA | GTTAT | GTTAC | GTTAG | GTTTA | GTTTT | GTTTC | GTTTG | GTTCA | GTTCT | GTTCC | GTTCG | GTTGA | GTTGT | GTTGC | GTTGG | | GTC | GTCAA | GTCAT | GTCAC | GTCAG | GTCTA | GTCTT | GTCTC | GTCTG | GTCCA | GTCCT | GTCCC | GTCCG | GTCGA | GTCGT | GTCGC | GTCGG | | GTG | GTGAA | GTGAT | GTGAC | GTGAG | GTGTA | GTGTT | GTGTC | GTGTG | GTGCA | GTGCT | GTGCC | GTGCG | GTGGA | GTGGT | GTGGC | GTGGG | | GCA | GCAAA | GCAAT | GCAAC | GCAAG | GCATA | GCATT | GCATC | GCATG | GCACA | GCACT | GCACC | GCACG | GCAGA | GCAGT | GCAGC | GCAGG | | GCT | GCTAA | GCTAT | GCTAC | GCTAG | GCTTA | GCTTT | GCTTC | GCTTG | GCTCA | GCTCT | GCTCC | GCTCG | GCTGA | GCTGT | GCTGC | GCTGG | | GCC | GCCAA | GCCAT | GCCAC | GCCAG | GCCTA | GCCTT | GCCTC | GCCTG | GCCCA | GCCCT | GCCCC | GCCCG | GCCGA | GCCGT | GCCGC | GCCGG | | GCG | GCGAA | GCGAT | GCGAC | GCGAG | GCGTA | GCGTT | GCGTC | GCGTG | GCGCA | GCGCT | GCGCC | GCGCG | GCGGA | GCGGT | GCGGC | GCGGG | | GGA | GGAAA | GGAAT | GGAAC | GGAAG | GGATA | GGATT | GGATC | GGATG | GGACA | GGACT | GGACC | GGACG | GGAGA | GGAGT | GGAGC | GGAGG | | GGT | GGTAA | GGTAT | GGTAC | GGTAG | GGTTA | GGTTT | GGTTC | GGTTG | GGTCA | GGTCT | GGTCC | GGTCG | GGTGA | GGTGT | GGTGC | GGTGG | | GGC | GGCAA | GGCAT | GGCAC | GGCAG | GGCTA | GGCTT | GGCTC | GGCTG | GGCCA | GGCCT | GGCCC | GGCCG | GGCGA | GGCGT | GGCGC | GGCGG | | GGG | GGGAA | GGGAT | GGGAC | GGGAG | GGGTA | GGGTT | GGGTC | GGGTG | GGGCA | GGGCT | GGGCC | GGGCG | GGGGA | GGGGT | GGGGC | GGGGG | Before filtering: read1: overrepresented sequences Sampling rate: 1 / 20 | | | | | --- | --- | --- | | overrepresented sequence | count (% of bases) | distribution: cycle 1 ~ cycle 150 | | AAAAAAAAAA | 274736 (0.109319%) | | | AAACTAACCTGTCTCACGACGGTCTAAACCCAGCTCACGTTCCCTATTGGTGGGTGAACAATCCAACACTTGGTGAATTCTGCTTCACAATGATAGGAAG | 323 (0.001285%) | | | AACCTGTCTCACGACGGTCTAAACCCAGCTCACGTTCCCTATTGGTGGGTGAACAATCCAACACTTGGTGAATTCTGCTTCACAATGATAGGAAGAGCCG | 122 (0.000485%) | | | AACTAACCTGTCTCACGACGGTCTAAACCCAGCTCACGTTCCCTATTGGTGGGTGAACAATCCAACACTTGGTGAATTCTGCTTCACAATGATAGGAAGA | 120 (0.000477%) | | | AAGATCGATC | 1125372 (0.447793%) | | | ACCTGTCTCACGACGGTCTAAACCCAGCTCACGTTCCCTATTGGTGGGTGAACAATCCAACACTTGGTGAATTCTGCTTCACAATGATAGGAAGAGCCGA | 97 (0.000386%) | | | ACGACGGTCTAAACCCAGCTCACGTTCCCTATTGGTGGGTGAACAATCCAACACTTGGTGAATTCTGCTTCACAATGATAGGAAGAGCCGACATCGAAGG | 103 (0.000410%) | | | ACGGTCTAAACCCAGCTCACGTTCCCTATTGGTGGGTGAACAATCCAACACTTGGTGAATTCTGCTTCACAATGATAGGAAGAGCCGACATCGAAGGATC | 91 (0.000362%) | | | ACTAACCTGTCTCACGACGGTCTAAACCCAGCTCACGTTCCCTATTGGTGGGTGAACAATCCAACACTTGGTGAATTCTGCTTCACAATGATAGGAAGAG | 106 (0.000422%) | | | AGGATCGATC | 891180 (0.354606%) | | | AGGGTAAAACTAACCTGTCTCACGACGGTCTAAACCCAGCTCACGTTCCCTATTGGTGGGTGAACAATCCAACACTTGGTGAATTCTGCTTCACAATGAT | 121 (0.000481%) | | | AGTAGGGTAAAACTAACCTGTCTCACGACGGTCTAAACCCAGCTCACGTTCCCTATTGGTGGGTGAACAATCCAACACTTGGTGAATTCTGCTTCACAAT | 188 (0.000748%) | | | ATCAGTAGGGTAAAACTAACCTGTCTCACGACGGTCTAAACCCAGCTCACGTTCCCTATTGGTGGGTGAACAATCCAACACTTGGTGAATTCTGCTTCAC | 4519 (0.017981%) | | | ATGATCGATC | 1064344 (0.423510%) | | | CACGACGGTCTAAACCCAGCTCACGTTCCCTATTGGTGGGTGAACAATCCAACACTTGGTGAATTCTGCTTCACAATGATAGGAAGAGCCGACATCGAAG | 163 (0.000649%) | | | CAGTAGGGTAAAACTAACCTGTCTCACGACGGTCTAAACCCAGCTCACGTTCCCTATTGGTGGGTGAACAATCCAACACTTGGTGAATTCTGCTTCACAA | 80 (0.000318%) | | | CCTGTCTCACGACGGTCTAAACCCAGCTCACGTTCCCTATTGGTGGGTGAACAATCCAACACTTGGTGAATTCTGCTTCACAATGATAGGAAGAGCCGAC | 201 (0.000800%) | | | CGACGGTCTAAACCCAGCTCACGTTCCCTATTGGTGGGTGAACAATCCAACACTTGGTGAATTCTGCTTCACAATGATAGGAAGAGCCGACATCGAAGGA | 87 (0.000346%) | | | CGGTCTAAACCCAGCTCACGTTCCCTATTGGTGGGTGAACAATCCAACACTTGGTGAATTCTGCTTCACAATGATAGGAAGAGCCGACATCGAAGGATCG | 39 (0.000155%) | | | CTAACCTGTCTCACGACGGTCTAAACCCAGCTCACGTTCCCTATTGGTGGGTGAACAATCCAACACTTGGTGAATTCTGCTTCACAATGATAGGAAGAGC | 207 (0.000824%) | | | CTCACGACGGTCTAAACCCAGCTCACGTTCCCTATTGGTGGGTGAACAATCCAACACTTGGTGAATTCTGCTTCACAATGATAGGAAGAGCCGACATCGA | 221 (0.000879%) | | | CTGTCTCACGACGGTCTAAACCCAGCTCACGTTCCCTATTGGTGGGTGAACAATCCAACACTTGGTGAATTCTGCTTCACAATGATAGGAAGAGCCGACA | 140 (0.000557%) | | | GACGGTCTAAACCCAGCTCACGTTCCCTATTGGTGGGTGAACAATCCAACACTTGGTGAATTCTGCTTCACAATGATAGGAAGAGCCGACATCGAAGGAT | 167 (0.000665%) | | | GATCGATCAA | 438317 (0.174409%) | | | GATCGATCAT | 467813 (0.186146%) | | | GATCGATCCT | 446159 (0.177530%) | | | GATCGATCTT | 570665 (0.227071%) | | | GGGTAAAACTAACCTGTCTCACGACGGTCTAAACCCAGCTCACGTTCCCTATTGGTGGGTGAACAATCCAACACTTGGTGAATTCTGCTTCACAATGATA | 40 (0.000159%) | | | GGTAAAACTAACCTGTCTCACGACGGTCTAAACCCAGCTCACGTTCCCTATTGGTGGGTGAACAATCCAACACTTGGTGAATTCTGCTTCACAATGATAG | 85 (0.000338%) | | | GGTCTAAACCCAGCTCACGTTCCCTATTGGTGGGTGAACAATCCAACACTTGGTGAATTCTGCTTCACAATGATAGGAAGAGCCGACATCGAAGGATCGA | 144 (0.000573%) | | | GTAGGGTAAAACTAACCTGTCTCACGACGGTCTAAACCCAGCTCACGTTCCCTATTGGTGGGTGAACAATCCAACACTTGGTGAATTCTGCTTCACAATG | 36 (0.000143%) | | | GTCTAAACCCAGCTCACGTTCCCTATTGGTGGGTGAACAATCCAACACTTGGTGAATTCTGCTTCACAATGATAGGAAGAGCCGACATCGAAGGATCGAT | 113 (0.000450%) | | | GTCTCACGACGGTCTAAACCCAGCTCACGTTCCCTATTGGTGGGTGAACAATCCAACACTTGGTGAATTCTGCTTCACAATGATAGGAAGAGCCGACATC | 75 (0.000298%) | | | TAACCTGTCTCACGACGGTCTAAACCCAGCTCACGTTCCCTATTGGTGGGTGAACAATCCAACACTTGGTGAATTCTGCTTCACAATGATAGGAAGAGCC | 45 (0.000179%) | | | TAGGGTAAAACTAACCTGTCTCACGACGGTCTAAACCCAGCTCACGTTCCCTATTGGTGGGTGAACAATCCAACACTTGGTGAATTCTGCTTCACAATGA | 7 (0.000028%) | | | TCACGACGGTCTAAACCCAGCTCACGTTCCCTATTGGTGGGTGAACAATCCAACACTTGGTGAATTCTGCTTCACAATGATAGGAAGAGCCGACATCGAA | 10 (0.000040%) | | | TCAGTAGGGTAAAACTAACCTGTCTCACGACGGTCTAAACCCAGCTCACGTTCCCTATTGGTGGGTGAACAATCCAACACTTGGTGAATTCTGCTTCACA | 12 (0.000048%) | | | TCTAAACCCAGCTCACGTTCCCTATTGGTGGGTGAACAATCCAACACTTGGTGAATTCTGCTTCACAATGATAGGAAGAGCCGACATCGAAGGATCGATC | 39 (0.000155%) | | | TCTCACGACGGTCTAAACCCAGCTCACGTTCCCTATTGGTGGGTGAACAATCCAACACTTGGTGAATTCTGCTTCACAATGATAGGAAGAGCCGACATCG | 27 (0.000107%) | | | TGATCGATCA | 170240 (0.067740%) | | | TGTCTCACGACGGTCTAAACCCAGCTCACGTTCCCTATTGGTGGGTGAACAATCCAACACTTGGTGAATTCTGCTTCACAATGATAGGAAGAGCCGACAT | 118 (0.000470%) | | | TTGATCGATC | 1024205 (0.407538%) | | | TTTTTTTTTTTTTTTTTTTT | 13665 (0.010875%) | | Before filtering: read2: quality Value of each position will be shown on mouse over. Before filtering: read2: base contents Value of each position will be shown on mouse over. Before filtering: read2: KMER counting Darker background means larger counts. The count will be shown on mouse over. | | | | | | | | | | | | | | | | | | | --- | --- | --- | --- | --- | --- | --- | --- | --- | --- | --- | --- | --- | --- | --- | --- | --- | | | AA | AT | AC | AG | TA | TT | TC | TG | CA | CT | CC | CG | GA | GT | GC | GG | | AAA | AAAAA | AAAAT | AAAAC | AAAAG | AAATA | AAATT | AAATC | AAATG | AAACA | AAACT | AAACC | AAACG | AAAGA | AAAGT | AAAGC | AAAGG | | AAT | AATAA | AATAT | AATAC | AATAG | AATTA | AATTT | AATTC | AATTG | AATCA | AATCT | AATCC | AATCG | AATGA | AATGT | AATGC | AATGG | | AAC | AACAA | AACAT | AACAC | AACAG | AACTA | AACTT | AACTC | AACTG | AACCA | AACCT | AACCC | AACCG | AACGA | AACGT | AACGC | AACGG | | AAG | AAGAA | AAGAT | AAGAC | AAGAG | AAGTA | AAGTT | AAGTC | AAGTG | AAGCA | AAGCT | AAGCC | AAGCG | AAGGA | AAGGT | AAGGC | AAGGG | | ATA | ATAAA | ATAAT | ATAAC | ATAAG | ATATA | ATATT | ATATC | ATATG | ATACA | ATACT | ATACC | ATACG | ATAGA | ATAGT | ATAGC | ATAGG | | ATT | ATTAA | ATTAT | ATTAC | ATTAG | ATTTA | ATTTT | ATTTC | ATTTG | ATTCA | ATTCT | ATTCC | ATTCG | ATTGA | ATTGT | ATTGC | ATTGG | | ATC | ATCAA | ATCAT | ATCAC | ATCAG | ATCTA | ATCTT | ATCTC | ATCTG | ATCCA | ATCCT | ATCCC | ATCCG | ATCGA | ATCGT | ATCGC | ATCGG | | ATG | ATGAA | ATGAT | ATGAC | ATGAG | ATGTA | ATGTT | ATGTC | ATGTG | ATGCA | ATGCT | ATGCC | ATGCG | ATGGA | ATGGT | ATGGC | ATGGG | | ACA | ACAAA | ACAAT | ACAAC | ACAAG | ACATA | ACATT | ACATC | ACATG | ACACA | ACACT | ACACC | ACACG | ACAGA | ACAGT | ACAGC | ACAGG | | ACT | ACTAA | ACTAT | ACTAC | ACTAG | ACTTA | ACTTT | ACTTC | ACTTG | ACTCA | ACTCT | ACTCC | ACTCG | ACTGA | ACTGT | ACTGC | ACTGG | | ACC | ACCAA | ACCAT | ACCAC | ACCAG | ACCTA | ACCTT | ACCTC | ACCTG | ACCCA | ACCCT | ACCCC | ACCCG | ACCGA | ACCGT | ACCGC | ACCGG | | ACG | ACGAA | ACGAT | ACGAC | ACGAG | ACGTA | ACGTT | ACGTC | ACGTG | ACGCA | ACGCT | ACGCC | ACGCG | ACGGA | ACGGT | ACGGC | ACGGG | | AGA | AGAAA | AGAAT | AGAAC | AGAAG | AGATA | AGATT | AGATC | AGATG | AGACA | AGACT | AGACC | AGACG | AGAGA | AGAGT | AGAGC | AGAGG | | AGT | AGTAA | AGTAT | AGTAC | AGTAG | AGTTA | AGTTT | AGTTC | AGTTG | AGTCA | AGTCT | AGTCC | AGTCG | AGTGA | AGTGT | AGTGC | AGTGG | | AGC | AGCAA | AGCAT | AGCAC | AGCAG | AGCTA | AGCTT | AGCTC | AGCTG | AGCCA | AGCCT | AGCCC | AGCCG | AGCGA | AGCGT | AGCGC | AGCGG | | AGG | AGGAA | AGGAT | AGGAC | AGGAG | AGGTA | AGGTT | AGGTC | AGGTG | AGGCA | AGGCT | AGGCC | AGGCG | AGGGA | AGGGT | AGGGC | AGGGG | | TAA | TAAAA | TAAAT | TAAAC | TAAAG | TAATA | TAATT | TAATC | TAATG | TAACA | TAACT | TAACC | TAACG | TAAGA | TAAGT | TAAGC | TAAGG | | TAT | TATAA | TATAT | TATAC | TATAG | TATTA | TATTT | TATTC | TATTG | TATCA | TATCT | TATCC | TATCG | TATGA | TATGT | TATGC | TATGG | | TAC | TACAA | TACAT | TACAC | TACAG | TACTA | TACTT | TACTC | TACTG | TACCA | TACCT | TACCC | TACCG | TACGA | TACGT | TACGC | TACGG | | TAG | TAGAA | TAGAT | TAGAC | TAGAG | TAGTA | TAGTT | TAGTC | TAGTG | TAGCA | TAGCT | TAGCC | TAGCG | TAGGA | TAGGT | TAGGC | TAGGG | | TTA | TTAAA | TTAAT | TTAAC | TTAAG | TTATA | TTATT | TTATC | TTATG | TTACA | TTACT | TTACC | TTACG | TTAGA | TTAGT | TTAGC | TTAGG | | TTT | TTTAA | TTTAT | TTTAC | TTTAG | TTTTA | TTTTT | TTTTC | TTTTG | TTTCA | TTTCT | TTTCC | TTTCG | TTTGA | TTTGT | TTTGC | TTTGG | | TTC | TTCAA | TTCAT | TTCAC | TTCAG | TTCTA | TTCTT | TTCTC | TTCTG | TTCCA | TTCCT | TTCCC | TTCCG | TTCGA | TTCGT | TTCGC | TTCGG | | TTG | TTGAA | TTGAT | TTGAC | TTGAG | TTGTA | TTGTT | TTGTC | TTGTG | TTGCA | TTGCT | TTGCC | TTGCG | TTGGA | TTGGT | TTGGC | TTGGG | | TCA | TCAAA | TCAAT | TCAAC | TCAAG | TCATA | TCATT | TCATC | TCATG | TCACA | TCACT | TCACC | TCACG | TCAGA | TCAGT | TCAGC | TCAGG | | TCT | TCTAA | TCTAT | TCTAC | TCTAG | TCTTA | TCTTT | TCTTC | TCTTG | TCTCA | TCTCT | TCTCC | TCTCG | TCTGA | TCTGT | TCTGC | TCTGG | | TCC | TCCAA | TCCAT | TCCAC | TCCAG | TCCTA | TCCTT | TCCTC | TCCTG | TCCCA | TCCCT | TCCCC | TCCCG | TCCGA | TCCGT | TCCGC | TCCGG | | TCG | TCGAA | TCGAT | TCGAC | TCGAG | TCGTA | TCGTT | TCGTC | TCGTG | TCGCA | TCGCT | TCGCC | TCGCG | TCGGA | TCGGT | TCGGC | TCGGG | | TGA | TGAAA | TGAAT | TGAAC | TGAAG | TGATA | TGATT | TGATC | TGATG | TGACA | TGACT | TGACC | TGACG | TGAGA | TGAGT | TGAGC | TGAGG | | TGT | TGTAA | TGTAT | TGTAC | TGTAG | TGTTA | TGTTT | TGTTC | TGTTG | TGTCA | TGTCT | TGTCC | TGTCG | TGTGA | TGTGT | TGTGC | TGTGG | | TGC | TGCAA | TGCAT | TGCAC | TGCAG | TGCTA | TGCTT | TGCTC | TGCTG | TGCCA | TGCCT | TGCCC | TGCCG | TGCGA | TGCGT | TGCGC | TGCGG | | TGG | TGGAA | TGGAT | TGGAC | TGGAG | TGGTA | TGGTT | TGGTC | TGGTG | TGGCA | TGGCT | TGGCC | TGGCG | TGGGA | TGGGT | TGGGC | TGGGG | | CAA | CAAAA | CAAAT | CAAAC | CAAAG | CAATA | CAATT | CAATC | CAATG | CAACA | CAACT | CAACC | CAACG | CAAGA | CAAGT | CAAGC | CAAGG | | CAT | CATAA | CATAT | CATAC | CATAG | CATTA | CATTT | CATTC | CATTG | CATCA | CATCT | CATCC | CATCG | CATGA | CATGT | CATGC | CATGG | | CAC | CACAA | CACAT | CACAC | CACAG | CACTA | CACTT | CACTC | CACTG | CACCA | CACCT | CACCC | CACCG | CACGA | CACGT | CACGC | CACGG | | CAG | CAGAA | CAGAT | CAGAC | CAGAG | CAGTA | CAGTT | CAGTC | CAGTG | CAGCA | CAGCT | CAGCC | CAGCG | CAGGA | CAGGT | CAGGC | CAGGG | | CTA | CTAAA | CTAAT | CTAAC | CTAAG | CTATA | CTATT | CTATC | CTATG | CTACA | CTACT | CTACC | CTACG | CTAGA | CTAGT | CTAGC | CTAGG | | CTT | CTTAA | CTTAT | CTTAC | CTTAG | CTTTA | CTTTT | CTTTC | CTTTG | CTTCA | CTTCT | CTTCC | CTTCG | CTTGA | CTTGT | CTTGC | CTTGG | | CTC | CTCAA | CTCAT | CTCAC | CTCAG | CTCTA | CTCTT | CTCTC | CTCTG | CTCCA | CTCCT | CTCCC | CTCCG | CTCGA | CTCGT | CTCGC | CTCGG | | CTG | CTGAA | CTGAT | CTGAC | CTGAG | CTGTA | CTGTT | CTGTC | CTGTG | CTGCA | CTGCT | CTGCC | CTGCG | CTGGA | CTGGT | CTGGC | CTGGG | | CCA | CCAAA | CCAAT | CCAAC | CCAAG | CCATA | CCATT | CCATC | CCATG | CCACA | CCACT | CCACC | CCACG | CCAGA | CCAGT | CCAGC | CCAGG | | CCT | CCTAA | CCTAT | CCTAC | CCTAG | CCTTA | CCTTT | CCTTC | CCTTG | CCTCA | CCTCT | CCTCC | CCTCG | CCTGA | CCTGT | CCTGC | CCTGG | | CCC | CCCAA | CCCAT | CCCAC | CCCAG | CCCTA | CCCTT | CCCTC | CCCTG | CCCCA | CCCCT | CCCCC | CCCCG | CCCGA | CCCGT | CCCGC | CCCGG | | CCG | CCGAA | CCGAT | CCGAC | CCGAG | CCGTA | CCGTT | CCGTC | CCGTG | CCGCA | CCGCT | CCGCC | CCGCG | CCGGA | CCGGT | CCGGC | CCGGG | | CGA | CGAAA | CGAAT | CGAAC | CGAAG | CGATA | CGATT | CGATC | CGATG | CGACA | CGACT | CGACC | CGACG | CGAGA | CGAGT | CGAGC | CGAGG | | CGT | CGTAA | CGTAT | CGTAC | CGTAG | CGTTA | CGTTT | CGTTC | CGTTG | CGTCA | CGTCT | CGTCC | CGTCG | CGTGA | CGTGT | CGTGC | CGTGG | | CGC | CGCAA | CGCAT | CGCAC | CGCAG | CGCTA | CGCTT | CGCTC | CGCTG | CGCCA | CGCCT | CGCCC | CGCCG | CGCGA | CGCGT | CGCGC | CGCGG | | CGG | CGGAA | CGGAT | CGGAC | CGGAG | CGGTA | CGGTT | CGGTC | CGGTG | CGGCA | CGGCT | CGGCC | CGGCG | CGGGA | CGGGT | CGGGC | CGGGG | | GAA | GAAAA | GAAAT | GAAAC | GAAAG | GAATA | GAATT | GAATC | GAATG | GAACA | GAACT | GAACC | GAACG | GAAGA | GAAGT | GAAGC | GAAGG | | GAT | GATAA | GATAT | GATAC | GATAG | GATTA | GATTT | GATTC | GATTG | GATCA | GATCT | GATCC | GATCG | GATGA | GATGT | GATGC | GATGG | | GAC | GACAA | GACAT | GACAC | GACAG | GACTA | GACTT | GACTC | GACTG | GACCA | GACCT | GACCC | GACCG | GACGA | GACGT | GACGC | GACGG | | GAG | GAGAA | GAGAT | GAGAC | GAGAG | GAGTA | GAGTT | GAGTC | GAGTG | GAGCA | GAGCT | GAGCC | GAGCG | GAGGA | GAGGT | GAGGC | GAGGG | | GTA | GTAAA | GTAAT | GTAAC | GTAAG | GTATA | GTATT | GTATC | GTATG | GTACA | GTACT | GTACC | GTACG | GTAGA | GTAGT | GTAGC | GTAGG | | GTT | GTTAA | GTTAT | GTTAC | GTTAG | GTTTA | GTTTT | GTTTC | GTTTG | GTTCA | GTTCT | GTTCC | GTTCG | GTTGA | GTTGT | GTTGC | GTTGG | | GTC | GTCAA | GTCAT | GTCAC | GTCAG | GTCTA | GTCTT | GTCTC | GTCTG | GTCCA | GTCCT | GTCCC | GTCCG | GTCGA | GTCGT | GTCGC | GTCGG | | GTG | GTGAA | GTGAT | GTGAC | GTGAG | GTGTA | GTGTT | GTGTC | GTGTG | GTGCA | GTGCT | GTGCC | GTGCG | GTGGA | GTGGT | GTGGC | GTGGG | | GCA | GCAAA | GCAAT | GCAAC | GCAAG | GCATA | GCATT | GCATC | GCATG | GCACA | GCACT | GCACC | GCACG | GCAGA | GCAGT | GCAGC | GCAGG | | GCT | GCTAA | GCTAT | GCTAC | GCTAG | GCTTA | GCTTT | GCTTC | GCTTG | GCTCA | GCTCT | GCTCC | GCTCG | GCTGA | GCTGT | GCTGC | GCTGG | | GCC | GCCAA | GCCAT | GCCAC | GCCAG | GCCTA | GCCTT | GCCTC | GCCTG | GCCCA | GCCCT | GCCCC | GCCCG | GCCGA | GCCGT | GCCGC | GCCGG | | GCG | GCGAA | GCGAT | GCGAC | GCGAG | GCGTA | GCGTT | GCGTC | GCGTG | GCGCA | GCGCT | GCGCC | GCGCG | GCGGA | GCGGT | GCGGC | GCGGG | | GGA | GGAAA | GGAAT | GGAAC | GGAAG | GGATA | GGATT | GGATC | GGATG | GGACA | GGACT | GGACC | GGACG | GGAGA | GGAGT | GGAGC | GGAGG | | GGT | GGTAA | GGTAT | GGTAC | GGTAG | GGTTA | GGTTT | GGTTC | GGTTG | GGTCA | GGTCT | GGTCC | GGTCG | GGTGA | GGTGT | GGTGC | GGTGG | | GGC | GGCAA | GGCAT | GGCAC | GGCAG | GGCTA | GGCTT | GGCTC | GGCTG | GGCCA | GGCCT | GGCCC | GGCCG | GGCGA | GGCGT | GGCGC | GGCGG | | GGG | GGGAA | GGGAT | GGGAC | GGGAG | GGGTA | GGGTT | GGGTC | GGGTG | GGGCA | GGGCT | GGGCC | GGGCG | GGGGA | GGGGT | GGGGC | GGGGG | Before filtering: read2: overrepresented sequences Sampling rate: 1 / 20 | | | | | --- | --- | --- | | overrepresented sequence | count (% of bases) | distribution: cycle 1 ~ cycle 150 | | AAAAAAAAAAAAAAAAAAAAAAAAAAAAAAAAAAAAAAAAAAAAAAAAAAAAAAAAAAAAAAAAAAAAAAAAAAAAAAAAAAAAAAAAAAAAAAAAAAAA | 773 (0.003076%) | | | AAGATCGATC | 1129958 (0.449653%) | | | AGGATCGATC | 887147 (0.353029%) | | | ATGATCGATC | 1062878 (0.422959%) | | | GATCGATCAA | 436416 (0.173666%) | | | GATCGATCAT | 464150 (0.184703%) | | | GATCGATCCT | 440620 (0.175339%) | | | GATCGATCTT | 558367 (0.222195%) | | | TGATCGATCA | 170756 (0.067950%) | | | TTGATCGATC | 1011707 (0.402596%) | | | TTTTTTTTTT | 245359 (0.097638%) | | After filtering After filtering: read1: quality Value of each position will be shown on mouse over. After filtering: read1: base contents Value of each position will be shown on mouse over. After filtering: read1: KMER counting Darker background means larger counts. The count will be shown on mouse over. | | | | | | | | | | | | | | | | | | | --- | --- | --- | --- | --- | --- | --- | --- | --- | --- | --- | --- | --- | --- | --- | --- | --- | | | AA | AT | AC | AG | TA | TT | TC | TG | CA | CT | CC | CG | GA | GT | GC | GG | | AAA | AAAAA | AAAAT | AAAAC | AAAAG | AAATA | AAATT | AAATC | AAATG | AAACA | AAACT | AAACC | AAACG | AAAGA | AAAGT | AAAGC | AAAGG | | AAT | AATAA | AATAT | AATAC | AATAG | AATTA | AATTT | AATTC | AATTG | AATCA | AATCT | AATCC | AATCG | AATGA | AATGT | AATGC | AATGG | | AAC | AACAA | AACAT | AACAC | AACAG | AACTA | AACTT | AACTC | AACTG | AACCA | AACCT | AACCC | AACCG | AACGA | AACGT | AACGC | AACGG | | AAG | AAGAA | AAGAT | AAGAC | AAGAG | AAGTA | AAGTT | AAGTC | AAGTG | AAGCA | AAGCT | AAGCC | AAGCG | AAGGA | AAGGT | AAGGC | AAGGG | | ATA | ATAAA | ATAAT | ATAAC | ATAAG | ATATA | ATATT | ATATC | ATATG | ATACA | ATACT | ATACC | ATACG | ATAGA | ATAGT | ATAGC | ATAGG | | ATT | ATTAA | ATTAT | ATTAC | ATTAG | ATTTA | ATTTT | ATTTC | ATTTG | ATTCA | ATTCT | ATTCC | ATTCG | ATTGA | ATTGT | ATTGC | ATTGG | | ATC | ATCAA | ATCAT | ATCAC | ATCAG | ATCTA | ATCTT | ATCTC | ATCTG | ATCCA | ATCCT | ATCCC | ATCCG | ATCGA | ATCGT | ATCGC | ATCGG | | ATG | ATGAA | ATGAT | ATGAC | ATGAG | ATGTA | ATGTT | ATGTC | ATGTG | ATGCA | ATGCT | ATGCC | ATGCG | ATGGA | ATGGT | ATGGC | ATGGG | | ACA | ACAAA | ACAAT | ACAAC | ACAAG | ACATA | ACATT | ACATC | ACATG | ACACA | ACACT | ACACC | ACACG | ACAGA | ACAGT | ACAGC | ACAGG | | ACT | ACTAA | ACTAT | ACTAC | ACTAG | ACTTA | ACTTT | ACTTC | ACTTG | ACTCA | ACTCT | ACTCC | ACTCG | ACTGA | ACTGT | ACTGC | ACTGG | | ACC | ACCAA | ACCAT | ACCAC | ACCAG | ACCTA | ACCTT | ACCTC | ACCTG | ACCCA | ACCCT | ACCCC | ACCCG | ACCGA | ACCGT | ACCGC | ACCGG | | ACG | ACGAA | ACGAT | ACGAC | ACGAG | ACGTA | ACGTT | ACGTC | ACGTG | ACGCA | ACGCT | ACGCC | ACGCG | ACGGA | ACGGT | ACGGC | ACGGG | | AGA | AGAAA | AGAAT | AGAAC | AGAAG | AGATA | AGATT | AGATC | AGATG | AGACA | AGACT | AGACC | AGACG | AGAGA | AGAGT | AGAGC | AGAGG | | AGT | AGTAA | AGTAT | AGTAC | AGTAG | AGTTA | AGTTT | AGTTC | AGTTG | AGTCA | AGTCT | AGTCC | AGTCG | AGTGA | AGTGT | AGTGC | AGTGG | | AGC | AGCAA | AGCAT | AGCAC | AGCAG | AGCTA | AGCTT | AGCTC | AGCTG | AGCCA | AGCCT | AGCCC | AGCCG | AGCGA | AGCGT | AGCGC | AGCGG | | AGG | AGGAA | AGGAT | AGGAC | AGGAG | AGGTA | AGGTT | AGGTC | AGGTG | AGGCA | AGGCT | AGGCC | AGGCG | AGGGA | AGGGT | AGGGC | AGGGG | | TAA | TAAAA | TAAAT | TAAAC | TAAAG | TAATA | TAATT | TAATC | TAATG | TAACA | TAACT | TAACC | TAACG | TAAGA | TAAGT | TAAGC | TAAGG | | TAT | TATAA | TATAT | TATAC | TATAG | TATTA | TATTT | TATTC | TATTG | TATCA | TATCT | TATCC | TATCG | TATGA | TATGT | TATGC | TATGG | | TAC | TACAA | TACAT | TACAC | TACAG | TACTA | TACTT | TACTC | TACTG | TACCA | TACCT | TACCC | TACCG | TACGA | TACGT | TACGC | TACGG | | TAG | TAGAA | TAGAT | TAGAC | TAGAG | TAGTA | TAGTT | TAGTC | TAGTG | TAGCA | TAGCT | TAGCC | TAGCG | TAGGA | TAGGT | TAGGC | TAGGG | | TTA | TTAAA | TTAAT | TTAAC | TTAAG | TTATA | TTATT | TTATC | TTATG | TTACA | TTACT | TTACC | TTACG | TTAGA | TTAGT | TTAGC | TTAGG | | TTT | TTTAA | TTTAT | TTTAC | TTTAG | TTTTA | TTTTT | TTTTC | TTTTG | TTTCA | TTTCT | TTTCC | TTTCG | TTTGA | TTTGT | TTTGC | TTTGG | | TTC | TTCAA | TTCAT | TTCAC | TTCAG | TTCTA | TTCTT | TTCTC | TTCTG | TTCCA | TTCCT | TTCCC | TTCCG | TTCGA | TTCGT | TTCGC | TTCGG | | TTG | TTGAA | TTGAT | TTGAC | TTGAG | TTGTA | TTGTT | TTGTC | TTGTG | TTGCA | TTGCT | TTGCC | TTGCG | TTGGA | TTGGT | TTGGC | TTGGG | | TCA | TCAAA | TCAAT | TCAAC | TCAAG | TCATA | TCATT | TCATC | TCATG | TCACA | TCACT | TCACC | TCACG | TCAGA | TCAGT | TCAGC | TCAGG | | TCT | TCTAA | TCTAT | TCTAC | TCTAG | TCTTA | TCTTT | TCTTC | TCTTG | TCTCA | TCTCT | TCTCC | TCTCG | TCTGA | TCTGT | TCTGC | TCTGG | | TCC | TCCAA | TCCAT | TCCAC | TCCAG | TCCTA | TCCTT | TCCTC | TCCTG | TCCCA | TCCCT | TCCCC | TCCCG | TCCGA | TCCGT | TCCGC | TCCGG | | TCG | TCGAA | TCGAT | TCGAC | TCGAG | TCGTA | TCGTT | TCGTC | TCGTG | TCGCA | TCGCT | TCGCC | TCGCG | TCGGA | TCGGT | TCGGC | TCGGG | | TGA | TGAAA | TGAAT | TGAAC | TGAAG | TGATA | TGATT | TGATC | TGATG | TGACA | TGACT | TGACC | TGACG | TGAGA | TGAGT | TGAGC | TGAGG | | TGT | TGTAA | TGTAT | TGTAC | TGTAG | TGTTA | TGTTT | TGTTC | TGTTG | TGTCA | TGTCT | TGTCC | TGTCG | TGTGA | TGTGT | TGTGC | TGTGG | | TGC | TGCAA | TGCAT | TGCAC | TGCAG | TGCTA | TGCTT | TGCTC | TGCTG | TGCCA | TGCCT | TGCCC | TGCCG | TGCGA | TGCGT | TGCGC | TGCGG | | TGG | TGGAA | TGGAT | TGGAC | TGGAG | TGGTA | TGGTT | TGGTC | TGGTG | TGGCA | TGGCT | TGGCC | TGGCG | TGGGA | TGGGT | TGGGC | TGGGG | | CAA | CAAAA | CAAAT | CAAAC | CAAAG | CAATA | CAATT | CAATC | CAATG | CAACA | CAACT | CAACC | CAACG | CAAGA | CAAGT | CAAGC | CAAGG | | CAT | CATAA | CATAT | CATAC | CATAG | CATTA | CATTT | CATTC | CATTG | CATCA | CATCT | CATCC | CATCG | CATGA | CATGT | CATGC | CATGG | | CAC | CACAA | CACAT | CACAC | CACAG | CACTA | CACTT | CACTC | CACTG | CACCA | CACCT | CACCC | CACCG | CACGA | CACGT | CACGC | CACGG | | CAG | CAGAA | CAGAT | CAGAC | CAGAG | CAGTA | CAGTT | CAGTC | CAGTG | CAGCA | CAGCT | CAGCC | CAGCG | CAGGA | CAGGT | CAGGC | CAGGG | | CTA | CTAAA | CTAAT | CTAAC | CTAAG | CTATA | CTATT | CTATC | CTATG | CTACA | CTACT | CTACC | CTACG | CTAGA | CTAGT | CTAGC | CTAGG | | CTT | CTTAA | CTTAT | CTTAC | CTTAG | CTTTA | CTTTT | CTTTC | CTTTG | CTTCA | CTTCT | CTTCC | CTTCG | CTTGA | CTTGT | CTTGC | CTTGG | | CTC | CTCAA | CTCAT | CTCAC | CTCAG | CTCTA | CTCTT | CTCTC | CTCTG | CTCCA | CTCCT | CTCCC | CTCCG | CTCGA | CTCGT | CTCGC | CTCGG | | CTG | CTGAA | CTGAT | CTGAC | CTGAG | CTGTA | CTGTT | CTGTC | CTGTG | CTGCA | CTGCT | CTGCC | CTGCG | CTGGA | CTGGT | CTGGC | CTGGG | | CCA | CCAAA | CCAAT | CCAAC | CCAAG | CCATA | CCATT | CCATC | CCATG | CCACA | CCACT | CCACC | CCACG | CCAGA | CCAGT | CCAGC | CCAGG | | CCT | CCTAA | CCTAT | CCTAC | CCTAG | CCTTA | CCTTT | CCTTC | CCTTG | CCTCA | CCTCT | CCTCC | CCTCG | CCTGA | CCTGT | CCTGC | CCTGG | | CCC | CCCAA | CCCAT | CCCAC | CCCAG | CCCTA | CCCTT | CCCTC | CCCTG | CCCCA | CCCCT | CCCCC | CCCCG | CCCGA | CCCGT | CCCGC | CCCGG | | CCG | CCGAA | CCGAT | CCGAC | CCGAG | CCGTA | CCGTT | CCGTC | CCGTG | CCGCA | CCGCT | CCGCC | CCGCG | CCGGA | CCGGT | CCGGC | CCGGG | | CGA | CGAAA | CGAAT | CGAAC | CGAAG | CGATA | CGATT | CGATC | CGATG | CGACA | CGACT | CGACC | CGACG | CGAGA | CGAGT | CGAGC | CGAGG | | CGT | CGTAA | CGTAT | CGTAC | CGTAG | CGTTA | CGTTT | CGTTC | CGTTG | CGTCA | CGTCT | CGTCC | CGTCG | CGTGA | CGTGT | CGTGC | CGTGG | | CGC | CGCAA | CGCAT | CGCAC | CGCAG | CGCTA | CGCTT | CGCTC | CGCTG | CGCCA | CGCCT | CGCCC | CGCCG | CGCGA | CGCGT | CGCGC | CGCGG | | CGG | CGGAA | CGGAT | CGGAC | CGGAG | CGGTA | CGGTT | CGGTC | CGGTG | CGGCA | CGGCT | CGGCC | CGGCG | CGGGA | CGGGT | CGGGC | CGGGG | | GAA | GAAAA | GAAAT | GAAAC | GAAAG | GAATA | GAATT | GAATC | GAATG | GAACA | GAACT | GAACC | GAACG | GAAGA | GAAGT | GAAGC | GAAGG | | GAT | GATAA | GATAT | GATAC | GATAG | GATTA | GATTT | GATTC | GATTG | GATCA | GATCT | GATCC | GATCG | GATGA | GATGT | GATGC | GATGG | | GAC | GACAA | GACAT | GACAC | GACAG | GACTA | GACTT | GACTC | GACTG | GACCA | GACCT | GACCC | GACCG | GACGA | GACGT | GACGC | GACGG | | GAG | GAGAA | GAGAT | GAGAC | GAGAG | GAGTA | GAGTT | GAGTC | GAGTG | GAGCA | GAGCT | GAGCC | GAGCG | GAGGA | GAGGT | GAGGC | GAGGG | | GTA | GTAAA | GTAAT | GTAAC | GTAAG | GTATA | GTATT | GTATC | GTATG | GTACA | GTACT | GTACC | GTACG | GTAGA | GTAGT | GTAGC | GTAGG | | GTT | GTTAA | GTTAT | GTTAC | GTTAG | GTTTA | GTTTT | GTTTC | GTTTG | GTTCA | GTTCT | GTTCC | GTTCG | GTTGA | GTTGT | GTTGC | GTTGG | | GTC | GTCAA | GTCAT | GTCAC | GTCAG | GTCTA | GTCTT | GTCTC | GTCTG | GTCCA | GTCCT | GTCCC | GTCCG | GTCGA | GTCGT | GTCGC | GTCGG | | GTG | GTGAA | GTGAT | GTGAC | GTGAG | GTGTA | GTGTT | GTGTC | GTGTG | GTGCA | GTGCT | GTGCC | GTGCG | GTGGA | GTGGT | GTGGC | GTGGG | | GCA | GCAAA | GCAAT | GCAAC | GCAAG | GCATA | GCATT | GCATC | GCATG | GCACA | GCACT | GCACC | GCACG | GCAGA | GCAGT | GCAGC | GCAGG | | GCT | GCTAA | GCTAT | GCTAC | GCTAG | GCTTA | GCTTT | GCTTC | GCTTG | GCTCA | GCTCT | GCTCC | GCTCG | GCTGA | GCTGT | GCTGC | GCTGG | | GCC | GCCAA | GCCAT | GCCAC | GCCAG | GCCTA | GCCTT | GCCTC | GCCTG | GCCCA | GCCCT | GCCCC | GCCCG | GCCGA | GCCGT | GCCGC | GCCGG | | GCG | GCGAA | GCGAT | GCGAC | GCGAG | GCGTA | GCGTT | GCGTC | GCGTG | GCGCA | GCGCT | GCGCC | GCGCG | GCGGA | GCGGT | GCGGC | GCGGG | | GGA | GGAAA | GGAAT | GGAAC | GGAAG | GGATA | GGATT | GGATC | GGATG | GGACA | GGACT | GGACC | GGACG | GGAGA | GGAGT | GGAGC | GGAGG | | GGT | GGTAA | GGTAT | GGTAC | GGTAG | GGTTA | GGTTT | GGTTC | GGTTG | GGTCA | GGTCT | GGTCC | GGTCG | GGTGA | GGTGT | GGTGC | GGTGG | | GGC | GGCAA | GGCAT | GGCAC | GGCAG | GGCTA | GGCTT | GGCTC | GGCTG | GGCCA | GGCCT | GGCCC | GGCCG | GGCGA | GGCGT | GGCGC | GGCGG | | GGG | GGGAA | GGGAT | GGGAC | GGGAG | GGGTA | GGGTT | GGGTC | GGGTG | GGGCA | GGGCT | GGGCC | GGGCG | GGGGA | GGGGT | GGGGC | GGGGG | After filtering: read1: overrepresented sequences Sampling rate: 1 / 20 | | | | | --- | --- | --- | | overrepresented sequence | count (% of bases) | distribution: cycle 1 ~ cycle 150 | | AAAAAAAAAA | 263012 (0.105592%) | | | AAACTAACCTGTCTCACGACGGTCTAAACCCAGCTCACGTTCCCTATTGGTGGGTGAACAATCCAACACTTGGTGAATTCTGCTTCACAATGATAGGAAG | 299 (0.001200%) | | | AACCTGTCTCACGACGGTCTAAACCCAGCTCACGTTCCCTATTGGTGGGTGAACAATCCAACACTTGGTGAATTCTGCTTCACAATGATAGGAAGAGCCG | 100 (0.000401%) | | | AACTAACCTGTCTCACGACGGTCTAAACCCAGCTCACGTTCCCTATTGGTGGGTGAACAATCCAACACTTGGTGAATTCTGCTTCACAATGATAGGAAGA | 127 (0.000510%) | | | AAGATCGATC | 1118795 (0.449167%) | | | ACCTGTCTCACGACGGTCTAAACCCAGCTCACGTTCCCTATTGGTGGGTGAACAATCCAACACTTGGTGAATTCTGCTTCACAATGATAGGAAGAGCCGA | 98 (0.000393%) | | | ACGACGGTCTAAACCCAGCTCACGTTCCCTATTGGTGGGTGAACAATCCAACACTTGGTGAATTCTGCTTCACAATGATAGGAAGAGCCGACATCGAAGG | 96 (0.000385%) | | | ACGGTCTAAACCCAGCTCACGTTCCCTATTGGTGGGTGAACAATCCAACACTTGGTGAATTCTGCTTCACAATGATAGGAAGAGCCGACATCGAAGGATC | 95 (0.000381%) | | | ACTAACCTGTCTCACGACGGTCTAAACCCAGCTCACGTTCCCTATTGGTGGGTGAACAATCCAACACTTGGTGAATTCTGCTTCACAATGATAGGAAGAG | 114 (0.000458%) | | | AGGATCGATC | 887151 (0.356168%) | | | AGGGTAAAACTAACCTGTCTCACGACGGTCTAAACCCAGCTCACGTTCCCTATTGGTGGGTGAACAATCCAACACTTGGTGAATTCTGCTTCACAATGAT | 121 (0.000486%) | | | AGTAGGGTAAAACTAACCTGTCTCACGACGGTCTAAACCCAGCTCACGTTCCCTATTGGTGGGTGAACAATCCAACACTTGGTGAATTCTGCTTCACAAT | 181 (0.000727%) | | | ATCAGTAGGGTAAAACTAACCTGTCTCACGACGGTCTAAACCCAGCTCACGTTCCCTATTGGTGGGTGAACAATCCAACACTTGGTGAATTCTGCTTCAC | 4534 (0.018203%) | | | ATGATCGATC | 1058833 (0.425093%) | | | CACGACGGTCTAAACCCAGCTCACGTTCCCTATTGGTGGGTGAACAATCCAACACTTGGTGAATTCTGCTTCACAATGATAGGAAGAGCCGACATCGAAG | 139 (0.000558%) | | | CAGTAGGGTAAAACTAACCTGTCTCACGACGGTCTAAACCCAGCTCACGTTCCCTATTGGTGGGTGAACAATCCAACACTTGGTGAATTCTGCTTCACAA | 89 (0.000357%) | | | CCTGTCTCACGACGGTCTAAACCCAGCTCACGTTCCCTATTGGTGGGTGAACAATCCAACACTTGGTGAATTCTGCTTCACAATGATAGGAAGAGCCGAC | 213 (0.000855%) | | | CGACGGTCTAAACCCAGCTCACGTTCCCTATTGGTGGGTGAACAATCCAACACTTGGTGAATTCTGCTTCACAATGATAGGAAGAGCCGACATCGAAGGA | 77 (0.000309%) | | | CGGTCTAAACCCAGCTCACGTTCCCTATTGGTGGGTGAACAATCCAACACTTGGTGAATTCTGCTTCACAATGATAGGAAGAGCCGACATCGAAGGATCG | 48 (0.000193%) | | | CTAACCTGTCTCACGACGGTCTAAACCCAGCTCACGTTCCCTATTGGTGGGTGAACAATCCAACACTTGGTGAATTCTGCTTCACAATGATAGGAAGAGC | 173 (0.000695%) | | | CTCACGACGGTCTAAACCCAGCTCACGTTCCCTATTGGTGGGTGAACAATCCAACACTTGGTGAATTCTGCTTCACAATGATAGGAAGAGCCGACATCGA | 240 (0.000964%) | | | CTGTCTCACGACGGTCTAAACCCAGCTCACGTTCCCTATTGGTGGGTGAACAATCCAACACTTGGTGAATTCTGCTTCACAATGATAGGAAGAGCCGACA | 144 (0.000578%) | | | GACGGTCTAAACCCAGCTCACGTTCCCTATTGGTGGGTGAACAATCCAACACTTGGTGAATTCTGCTTCACAATGATAGGAAGAGCCGACATCGAAGGAT | 171 (0.000687%) | | | GATCGATCAA | 436700 (0.175324%) | | | GATCGATCAT | 463213 (0.185968%) | | | GATCGATCCT | 443416 (0.178020%) | | | GATCGATCTT | 566315 (0.227360%) | | | GGGTAAAACTAACCTGTCTCACGACGGTCTAAACCCAGCTCACGTTCCCTATTGGTGGGTGAACAATCCAACACTTGGTGAATTCTGCTTCACAATGATA | 53 (0.000213%) | | | GGTAAAACTAACCTGTCTCACGACGGTCTAAACCCAGCTCACGTTCCCTATTGGTGGGTGAACAATCCAACACTTGGTGAATTCTGCTTCACAATGATAG | 74 (0.000297%) | | | GGTCTAAACCCAGCTCACGTTCCCTATTGGTGGGTGAACAATCCAACACTTGGTGAATTCTGCTTCACAATGATAGGAAGAGCCGACATCGAAGGATCGA | 161 (0.000646%) | | | GTAGGGTAAAACTAACCTGTCTCACGACGGTCTAAACCCAGCTCACGTTCCCTATTGGTGGGTGAACAATCCAACACTTGGTGAATTCTGCTTCACAATG | 42 (0.000169%) | | | GTCTAAACCCAGCTCACGTTCCCTATTGGTGGGTGAACAATCCAACACTTGGTGAATTCTGCTTCACAATGATAGGAAGAGCCGACATCGAAGGATCGAT | 120 (0.000482%) | | | GTCTCACGACGGTCTAAACCCAGCTCACGTTCCCTATTGGTGGGTGAACAATCCAACACTTGGTGAATTCTGCTTCACAATGATAGGAAGAGCCGACATC | 75 (0.000301%) | | | TAACCTGTCTCACGACGGTCTAAACCCAGCTCACGTTCCCTATTGGTGGGTGAACAATCCAACACTTGGTGAATTCTGCTTCACAATGATAGGAAGAGCC | 45 (0.000181%) | | | TAGGGTAAAACTAACCTGTCTCACGACGGTCTAAACCCAGCTCACGTTCCCTATTGGTGGGTGAACAATCCAACACTTGGTGAATTCTGCTTCACAATGA | 14 (0.000056%) | | | TCACGACGGTCTAAACCCAGCTCACGTTCCCTATTGGTGGGTGAACAATCCAACACTTGGTGAATTCTGCTTCACAATGATAGGAAGAGCCGACATCGAA | 7 (0.000028%) | | | TCAGTAGGGTAAAACTAACCTGTCTCACGACGGTCTAAACCCAGCTCACGTTCCCTATTGGTGGGTGAACAATCCAACACTTGGTGAATTCTGCTTCACA | 11 (0.000044%) | | | TCTAAACCCAGCTCACGTTCCCTATTGGTGGGTGAACAATCCAACACTTGGTGAATTCTGCTTCACAATGATAGGAAGAGCCGACATCGAAGGATCGATC | 48 (0.000193%) | | | TCTCACGACGGTCTAAACCCAGCTCACGTTCCCTATTGGTGGGTGAACAATCCAACACTTGGTGAATTCTGCTTCACAATGATAGGAAGAGCCGACATCG | 41 (0.000165%) | | | TGATCGATCA | 168554 (0.067670%) | | | TGTCTCACGACGGTCTAAACCCAGCTCACGTTCCCTATTGGTGGGTGAACAATCCAACACTTGGTGAATTCTGCTTCACAATGATAGGAAGAGCCGACAT | 115 (0.000462%) | | | TTGATCGATC | 1018395 (0.408859%) | | | TTTTTTTTTTTTTTTTTTTT | 12294 (0.009871%) | | After filtering: read2: quality Value of each position will be shown on mouse over. After filtering: read2: base contents Value of each position will be shown on mouse over. After filtering: read2: KMER counting Darker background means larger counts. The count will be shown on mouse over. | | | | | | | | | | | | | | | | | | | --- | --- | --- | --- | --- | --- | --- | --- | --- | --- | --- | --- | --- | --- | --- | --- | --- | | | AA | AT | AC | AG | TA | TT | TC | TG | CA | CT | CC | CG | GA | GT | GC | GG | | AAA | AAAAA | AAAAT | AAAAC | AAAAG | AAATA | AAATT | AAATC | AAATG | AAACA | AAACT | AAACC | AAACG | AAAGA | AAAGT | AAAGC | AAAGG | | AAT | AATAA | AATAT | AATAC | AATAG | AATTA | AATTT | AATTC | AATTG | AATCA | AATCT | AATCC | AATCG | AATGA | AATGT | AATGC | AATGG | | AAC | AACAA | AACAT | AACAC | AACAG | AACTA | AACTT | AACTC | AACTG | AACCA | AACCT | AACCC | AACCG | AACGA | AACGT | AACGC | AACGG | | AAG | AAGAA | AAGAT | AAGAC | AAGAG | AAGTA | AAGTT | AAGTC | AAGTG | AAGCA | AAGCT | AAGCC | AAGCG | AAGGA | AAGGT | AAGGC | AAGGG | | ATA | ATAAA | ATAAT | ATAAC | ATAAG | ATATA | ATATT | ATATC | ATATG | ATACA | ATACT | ATACC | ATACG | ATAGA | ATAGT | ATAGC | ATAGG | | ATT | ATTAA | ATTAT | ATTAC | ATTAG | ATTTA | ATTTT | ATTTC | ATTTG | ATTCA | ATTCT | ATTCC | ATTCG | ATTGA | ATTGT | ATTGC | ATTGG | | ATC | ATCAA | ATCAT | ATCAC | ATCAG | ATCTA | ATCTT | ATCTC | ATCTG | ATCCA | ATCCT | ATCCC | ATCCG | ATCGA | ATCGT | ATCGC | ATCGG | | ATG | ATGAA | ATGAT | ATGAC | ATGAG | ATGTA | ATGTT | ATGTC | ATGTG | ATGCA | ATGCT | ATGCC | ATGCG | ATGGA | ATGGT | ATGGC | ATGGG | | ACA | ACAAA | ACAAT | ACAAC | ACAAG | ACATA | ACATT | ACATC | ACATG | ACACA | ACACT | ACACC | ACACG | ACAGA | ACAGT | ACAGC | ACAGG | | ACT | ACTAA | ACTAT | ACTAC | ACTAG | ACTTA | ACTTT | ACTTC | ACTTG | ACTCA | ACTCT | ACTCC | ACTCG | ACTGA | ACTGT | ACTGC | ACTGG | | ACC | ACCAA | ACCAT | ACCAC | ACCAG | ACCTA | ACCTT | ACCTC | ACCTG | ACCCA | ACCCT | ACCCC | ACCCG | ACCGA | ACCGT | ACCGC | ACCGG | | ACG | ACGAA | ACGAT | ACGAC | ACGAG | ACGTA | ACGTT | ACGTC | ACGTG | ACGCA | ACGCT | ACGCC | ACGCG | ACGGA | ACGGT | ACGGC | ACGGG | | AGA | AGAAA | AGAAT | AGAAC | AGAAG | AGATA | AGATT | AGATC | AGATG | AGACA | AGACT | AGACC | AGACG | AGAGA | AGAGT | AGAGC | AGAGG | | AGT | AGTAA | AGTAT | AGTAC | AGTAG | AGTTA | AGTTT | AGTTC | AGTTG | AGTCA | AGTCT | AGTCC | AGTCG | AGTGA | AGTGT | AGTGC | AGTGG | | AGC | AGCAA | AGCAT | AGCAC | AGCAG | AGCTA | AGCTT | AGCTC | AGCTG | AGCCA | AGCCT | AGCCC | AGCCG | AGCGA | AGCGT | AGCGC | AGCGG | | AGG | AGGAA | AGGAT | AGGAC | AGGAG | AGGTA | AGGTT | AGGTC | AGGTG | AGGCA | AGGCT | AGGCC | AGGCG | AGGGA | AGGGT | AGGGC | AGGGG | | TAA | TAAAA | TAAAT | TAAAC | TAAAG | TAATA | TAATT | TAATC | TAATG | TAACA | TAACT | TAACC | TAACG | TAAGA | TAAGT | TAAGC | TAAGG | | TAT | TATAA | TATAT | TATAC | TATAG | TATTA | TATTT | TATTC | TATTG | TATCA | TATCT | TATCC | TATCG | TATGA | TATGT | TATGC | TATGG | | TAC | TACAA | TACAT | TACAC | TACAG | TACTA | TACTT | TACTC | TACTG | TACCA | TACCT | TACCC | TACCG | TACGA | TACGT | TACGC | TACGG | | TAG | TAGAA | TAGAT | TAGAC | TAGAG | TAGTA | TAGTT | TAGTC | TAGTG | TAGCA | TAGCT | TAGCC | TAGCG | TAGGA | TAGGT | TAGGC | TAGGG | | TTA | TTAAA | TTAAT | TTAAC | TTAAG | TTATA | TTATT | TTATC | TTATG | TTACA | TTACT | TTACC | TTACG | TTAGA | TTAGT | TTAGC | TTAGG | | TTT | TTTAA | TTTAT | TTTAC | TTTAG | TTTTA | TTTTT | TTTTC | TTTTG | TTTCA | TTTCT | TTTCC | TTTCG | TTTGA | TTTGT | TTTGC | TTTGG | | TTC | TTCAA | TTCAT | TTCAC | TTCAG | TTCTA | TTCTT | TTCTC | TTCTG | TTCCA | TTCCT | TTCCC | TTCCG | TTCGA | TTCGT | TTCGC | TTCGG | | TTG | TTGAA | TTGAT | TTGAC | TTGAG | TTGTA | TTGTT | TTGTC | TTGTG | TTGCA | TTGCT | TTGCC | TTGCG | TTGGA | TTGGT | TTGGC | TTGGG | | TCA | TCAAA | TCAAT | TCAAC | TCAAG | TCATA | TCATT | TCATC | TCATG | TCACA | TCACT | TCACC | TCACG | TCAGA | TCAGT | TCAGC | TCAGG | | TCT | TCTAA | TCTAT | TCTAC | TCTAG | TCTTA | TCTTT | TCTTC | TCTTG | TCTCA | TCTCT | TCTCC | TCTCG | TCTGA | TCTGT | TCTGC | TCTGG | | TCC | TCCAA | TCCAT | TCCAC | TCCAG | TCCTA | TCCTT | TCCTC | TCCTG | TCCCA | TCCCT | TCCCC | TCCCG | TCCGA | TCCGT | TCCGC | TCCGG | | TCG | TCGAA | TCGAT | TCGAC | TCGAG | TCGTA | TCGTT | TCGTC | TCGTG | TCGCA | TCGCT | TCGCC | TCGCG | TCGGA | TCGGT | TCGGC | TCGGG | | TGA | TGAAA | TGAAT | TGAAC | TGAAG | TGATA | TGATT | TGATC | TGATG | TGACA | TGACT | TGACC | TGACG | TGAGA | TGAGT | TGAGC | TGAGG | | TGT | TGTAA | TGTAT | TGTAC | TGTAG | TGTTA | TGTTT | TGTTC | TGTTG | TGTCA | TGTCT | TGTCC | TGTCG | TGTGA | TGTGT | TGTGC | TGTGG | | TGC | TGCAA | TGCAT | TGCAC | TGCAG | TGCTA | TGCTT | TGCTC | TGCTG | TGCCA | TGCCT | TGCCC | TGCCG | TGCGA | TGCGT | TGCGC | TGCGG | | TGG | TGGAA | TGGAT | TGGAC | TGGAG | TGGTA | TGGTT | TGGTC | TGGTG | TGGCA | TGGCT | TGGCC | TGGCG | TGGGA | TGGGT | TGGGC | TGGGG | | CAA | CAAAA | CAAAT | CAAAC | CAAAG | CAATA | CAATT | CAATC | CAATG | CAACA | CAACT | CAACC | CAACG | CAAGA | CAAGT | CAAGC | CAAGG | | CAT | CATAA | CATAT | CATAC | CATAG | CATTA | CATTT | CATTC | CATTG | CATCA | CATCT | CATCC | CATCG | CATGA | CATGT | CATGC | CATGG | | CAC | CACAA | CACAT | CACAC | CACAG | CACTA | CACTT | CACTC | CACTG | CACCA | CACCT | CACCC | CACCG | CACGA | CACGT | CACGC | CACGG | | CAG | CAGAA | CAGAT | CAGAC | CAGAG | CAGTA | CAGTT | CAGTC | CAGTG | CAGCA | CAGCT | CAGCC | CAGCG | CAGGA | CAGGT | CAGGC | CAGGG | | CTA | CTAAA | CTAAT | CTAAC | CTAAG | CTATA | CTATT | CTATC | CTATG | CTACA | CTACT | CTACC | CTACG | CTAGA | CTAGT | CTAGC | CTAGG | | CTT | CTTAA | CTTAT | CTTAC | CTTAG | CTTTA | CTTTT | CTTTC | CTTTG | CTTCA | CTTCT | CTTCC | CTTCG | CTTGA | CTTGT | CTTGC | CTTGG | | CTC | CTCAA | CTCAT | CTCAC | CTCAG | CTCTA | CTCTT | CTCTC | CTCTG | CTCCA | CTCCT | CTCCC | CTCCG | CTCGA | CTCGT | CTCGC | CTCGG | | CTG | CTGAA | CTGAT | CTGAC | CTGAG | CTGTA | CTGTT | CTGTC | CTGTG | CTGCA | CTGCT | CTGCC | CTGCG | CTGGA | CTGGT | CTGGC | CTGGG | | CCA | CCAAA | CCAAT | CCAAC | CCAAG | CCATA | CCATT | CCATC | CCATG | CCACA | CCACT | CCACC | CCACG | CCAGA | CCAGT | CCAGC | CCAGG | | CCT | CCTAA | CCTAT | CCTAC | CCTAG | CCTTA | CCTTT | CCTTC | CCTTG | CCTCA | CCTCT | CCTCC | CCTCG | CCTGA | CCTGT | CCTGC | CCTGG | | CCC | CCCAA | CCCAT | CCCAC | CCCAG | CCCTA | CCCTT | CCCTC | CCCTG | CCCCA | CCCCT | CCCCC | CCCCG | CCCGA | CCCGT | CCCGC | CCCGG | | CCG | CCGAA | CCGAT | CCGAC | CCGAG | CCGTA | CCGTT | CCGTC | CCGTG | CCGCA | CCGCT | CCGCC | CCGCG | CCGGA | CCGGT | CCGGC | CCGGG | | CGA | CGAAA | CGAAT | CGAAC | CGAAG | CGATA | CGATT | CGATC | CGATG | CGACA | CGACT | CGACC | CGACG | CGAGA | CGAGT | CGAGC | CGAGG | | CGT | CGTAA | CGTAT | CGTAC | CGTAG | CGTTA | CGTTT | CGTTC | CGTTG | CGTCA | CGTCT | CGTCC | CGTCG | CGTGA | CGTGT | CGTGC | CGTGG | | CGC | CGCAA | CGCAT | CGCAC | CGCAG | CGCTA | CGCTT | CGCTC | CGCTG | CGCCA | CGCCT | CGCCC | CGCCG | CGCGA | CGCGT | CGCGC | CGCGG | | CGG | CGGAA | CGGAT | CGGAC | CGGAG | CGGTA | CGGTT | CGGTC | CGGTG | CGGCA | CGGCT | CGGCC | CGGCG | CGGGA | CGGGT | CGGGC | CGGGG | | GAA | GAAAA | GAAAT | GAAAC | GAAAG | GAATA | GAATT | GAATC | GAATG | GAACA | GAACT | GAACC | GAACG | GAAGA | GAAGT | GAAGC | GAAGG | | GAT | GATAA | GATAT | GATAC | GATAG | GATTA | GATTT | GATTC | GATTG | GATCA | GATCT | GATCC | GATCG | GATGA | GATGT | GATGC | GATGG | | GAC | GACAA | GACAT | GACAC | GACAG | GACTA | GACTT | GACTC | GACTG | GACCA | GACCT | GACCC | GACCG | GACGA | GACGT | GACGC | GACGG | | GAG | GAGAA | GAGAT | GAGAC | GAGAG | GAGTA | GAGTT | GAGTC | GAGTG | GAGCA | GAGCT | GAGCC | GAGCG | GAGGA | GAGGT | GAGGC | GAGGG | | GTA | GTAAA | GTAAT | GTAAC | GTAAG | GTATA | GTATT | GTATC | GTATG | GTACA | GTACT | GTACC | GTACG | GTAGA | GTAGT | GTAGC | GTAGG | | GTT | GTTAA | GTTAT | GTTAC | GTTAG | GTTTA | GTTTT | GTTTC | GTTTG | GTTCA | GTTCT | GTTCC | GTTCG | GTTGA | GTTGT | GTTGC | GTTGG | | GTC | GTCAA | GTCAT | GTCAC | GTCAG | GTCTA | GTCTT | GTCTC | GTCTG | GTCCA | GTCCT | GTCCC | GTCCG | GTCGA | GTCGT | GTCGC | GTCGG | | GTG | GTGAA | GTGAT | GTGAC | GTGAG | GTGTA | GTGTT | GTGTC | GTGTG | GTGCA | GTGCT | GTGCC | GTGCG | GTGGA | GTGGT | GTGGC | GTGGG | | GCA | GCAAA | GCAAT | GCAAC | GCAAG | GCATA | GCATT | GCATC | GCATG | GCACA | GCACT | GCACC | GCACG | GCAGA | GCAGT | GCAGC | GCAGG | | GCT | GCTAA | GCTAT | GCTAC | GCTAG | GCTTA | GCTTT | GCTTC | GCTTG | GCTCA | GCTCT | GCTCC | GCTCG | GCTGA | GCTGT | GCTGC | GCTGG | | GCC | GCCAA | GCCAT | GCCAC | GCCAG | GCCTA | GCCTT | GCCTC | GCCTG | GCCCA | GCCCT | GCCCC | GCCCG | GCCGA | GCCGT | GCCGC | GCCGG | | GCG | GCGAA | GCGAT | GCGAC | GCGAG | GCGTA | GCGTT | GCGTC | GCGTG | GCGCA | GCGCT | GCGCC | GCGCG | GCGGA | GCGGT | GCGGC | GCGGG | | GGA | GGAAA | GGAAT | GGAAC | GGAAG | GGATA | GGATT | GGATC | GGATG | GGACA | GGACT | GGACC | GGACG | GGAGA | GGAGT | GGAGC | GGAGG | | GGT | GGTAA | GGTAT | GGTAC | GGTAG | GGTTA | GGTTT | GGTTC | GGTTG | GGTCA | GGTCT | GGTCC | GGTCG | GGTGA | GGTGT | GGTGC | GGTGG | | GGC | GGCAA | GGCAT | GGCAC | GGCAG | GGCTA | GGCTT | GGCTC | GGCTG | GGCCA | GGCCT | GGCCC | GGCCG | GGCGA | GGCGT | GGCGC | GGCGG | | GGG | GGGAA | GGGAT | GGGAC | GGGAG | GGGTA | GGGTT | GGGTC | GGGTG | GGGCA | GGGCT | GGGCC | GGGCG | GGGGA | GGGGT | GGGGC | GGGGG | After filtering: read2: overrepresented sequences Sampling rate: 1 / 20 | | | | | --- | --- | --- | | overrepresented sequence | count (% of bases) | distribution: cycle 1 ~ cycle 150 | | AAAAAAAAAAAAAAAAAAAAAAAAAAAAAAAAAAAAAAAAAAAAAAAAAAAAAAAAAAAAAAAAAAAAAAAAAAAAAAAAAAAAAAAAAAAAAAAAAAAA | 749 (0.003007%) | | | AAGATCGATC | 1127304 (0.452592%) | | | AGGATCGATC | 885316 (0.355438%) | | | ATGATCGATC | 1057436 (0.424541%) | | | GATCGATCAA | 435102 (0.174686%) | | | GATCGATCAT | 462890 (0.185842%) | | | GATCGATCCT | 439461 (0.176436%) | | | GATCGATCTT | 557663 (0.223892%) | | | TGATCGATCA | 170869 (0.068601%) | | | TTGATCGATC | 1010176 (0.405567%) | | | TTTTTTTTTT | 235186 (0.094423%) | |

fastp -p -i resources/raw\_hic/SRR18311512\_1.fastq.gz -I resources/raw\_hic/SRR18311512\_2.fastq.gz -o results/fastp/hic\_trim\_1.fastq.gz -O results/fastp/hic\_trim\_2.fastq.gz --detect\_adapter\_for\_pe --json results/fastp/hic\_report\_fastp.HiC.json --html results/fastp/hic\_report\_fastp.HiC.html --thread 20

fastp 0.23.4, at 2024-05-07 10:48:51
