## Additional files for "Colora: A Snakemake Workflow for Complete Chromosome-scale *De Novo* Genome Assembly": S10_NanoPlot_A.thaliana_ONT.html

NanoPlot Report


### NanoPlot statistics report

#### Menu

- Summary Statistics
- Plots
  - Weighted histogram of read lengths
  - Weighted histogram of read lengths after log transformation
  - Non weighted histogram of read lengths
  - Non weighted histogram of read lengths after log transformation
  - Yield by length
  - Read lengths vs Average read quality plot using dots
- Report issue on Github


#### NanoPlot reports

##### Summary statistics

|  |  |
| --- | --- |
| General summary |  |
| Mean read length | 18,541.3 |
| Mean read quality | 11.1 |
| Median read length | 7,818.0 |
| Median read quality | 11.2 |
| Number of reads | 3,064,191.0 |
| Read length N50 | 46,452.0 |
| STDEV read length | 26,536.0 |
| Total bases | 56,814,196,989.0 |
| Number, percentage and megabases of reads above quality cutoffs |  |
| >Q5 | 3064191 (100.0%) 56814.2Mb |
| >Q7 | 3064123 (100.0%) 56814.2Mb |
| >Q10 | 2168595 (70.8%) 40456.1Mb |
| >Q12 | 1055916 (34.5%) 19383.7Mb |
| >Q15 | 6640 (0.2%) 12.3Mb |
| Top 5 highest mean basecall quality scores and their read lengths |  |
| 1 | 21.0 (1) |
| 2 | 19.0 (1) |
| 3 | 19.0 (1) |
| 4 | 19.0 (1) |
| 5 | 18.9 (358) |
| Top 5 longest reads and their mean basecall quality score |  |
| 1 | 495032 (12.4) |
| 2 | 457760 (8.7) |
| 3 | 439434 (9.1) |
| 4 | 438143 (8.7) |
| 5 | 431286 (9.7) |

##### Plots

Weighted histogram of read lengths

###### Weighted histogram of read lengths

Weighted histogram of read lengths after log transformation

###### Weighted histogram of read lengths after log transformation

Non weighted histogram of read lengths

###### Non weighted histogram of read lengths

Non weighted histogram of read lengths after log transformation

###### Non weighted histogram of read lengths after log transformation

Yield by length

###### Yield by length

Read lengths vs Average read quality plot using dots

###### Read lengths vs Average read quality plot using dots
